# Defining glycoproteoform landscapes through an integrated glycoproteomics approach enabled by high-resolving power proton transfer charge reduction tandem mass spectrometry

**DOI:** 10.64898/2026.08.27.747529

**Authors:** Tim S. Veth, Emmajay Sutherland, Joshua D Hinkle, David Bergen, Rafael D. Melani, Graeme C McAlister, Christopher Mullen, Nicholas M. Riley

## Abstract

Glycan heterogeneity is a fundamental property of glycoproteins. A holistic understanding of glycan modification states is critical to translating glycoproteome regulation to biological function, but the high degree of glycosite-level heterogeneity leads to technical challenges in measuring glycoproteoforms. Common bottom-up glycoproteomics provide some insights but cannot recapitulate the full ensemble of glycoproteoforms from glycopeptide measurements alone. Promising efforts to profile masses of intact glycoproteins have recently explored data-independent acquisition (DIA) coupled with proton-transfer charge reduction (PTCR) or electron-capture-induced charge reduction mass spectrometry (MS). While valuable for generating broad glycoproteoform mass distributions, these approaches have remained limited in their ability to generate discrete glycoproteoform mass measurements, largely because they rely on low-resolving power measurements and deconvolution that does not account for isotopic information. Here, we develop a DIA-PTCR workflow that couples high-resolving power (Rp ∼240,000 at m/z 200) tandem mass spectra with an open-source processing suite to define glycoproteoform populations within 20 ppm mass accuracy thresholds. We demonstrate the glycoproteoform characterization capabilities of this platform using a collection of glycoproteins with well-described translational interests (EpCAM, TIGIT, CD40, PDL1, and CD24). With a focus on EpCAM, we showcase how intact glycoproteoform masses acquired using our high-resolving power DIA-PTCR (hRp-DIA-PTCR) approach can be integrated with bottom-up intact glycoproteomics and Direct-Mass Technology (i.e., Orbitrap-based charge-detection MS) acquisitions to inform structural and biological insights. Altogether, our hRp-DIA-PTCR method extends the current capabilities of intact glycoprotein analyses by enabling robust characterization of isotopically resolved proteoforms and facilitating deep biological interpretation of glycosylation heterogeneity. Our open-source informatics platform includes a GUI-based tool called PTsliCR to clean PTCR spectra directly from DIA-PTCR raw files and a deconvolution R package called IsoTrac, both of which are freely available on GitHub at https://github.com/riley-research.

## INTRODUCTION

Protein glycosylation describes a diverse family of post-translational modifications comprising multiple classes of glycoproteins (e.g., N- and O-glycoproteins) that harbor hundreds of different glycan (carbohydrate) structures.^1^ Glycoproteins exist as a collection of heterogeneous modification states called glycoproteoforms that vary both in glycosite occupancy (macroheterogeneity) and also in composition and structure of the glycans that occupy any given site (microheterogeneity).^2^ While critical for tuning biological functionality, glycoprotein heterogeneity dramatically increases the chemical complexity of the glycoproteome relative to non-modified proteins. Several decades of analytical method development have been dedicated to improving our ability to characterize glycoproteins with site-specific resolution to better understand the biological consequences of specific glycoproteoforms. A majority of these efforts have focused on bottom-up approaches that characterize proteolytically-derived glycopeptides that harbor a single glycosite or a small collection of glycosites (generally fewer than five). Indeed, improvements in enrichment strategies, mass spectrometry instrumentation, and informatic tools have enabled site-specific characterization of glycopeptides with increasing breadth and depth.^3–9^

Even as bottom-up glycoproteomic methods advance, however, they remain fundamentally limited by their reliance on proteolysis to generate glycopeptides from glycoproteoforms. The glycocode is a language of combinational modification states^10,11^, so this limitation in capturing glycoproteoforms impedes our ability to capture functional units of the glycoproteome. An understanding of how information is encrypted in a holistic manner is critical for translating glycoproteome regulation into biological function^12^, and bottom-up glycoproteomics cannot recapitulate the full ensemble of glycoproteoforms. To address this shortcoming, several efforts have sought to complement bottom-up workflows by measuring intact glycoproteins with native mass spectrometry (MS) and related approaches.^13–20^ This strategy can be useful for relatively simple or highly purified glycoproteins, but glycoproteoform heterogeneity often yields highly complex mass spectra that cannot be deconvoluted into useful glycoprotein information, even under native MS conditions. Charge detection mass spectrometry (CDMS) has been combined with native MS to attempt to resolve glycoproteoform masses with some degree of success,^21–25^ but CDMS of complex glycoproteins often generates broad mass distributions that do not provide discrete glycoproteoform masses that connect to glycosite-level measurements.

Recently, several groups have explored spectral decongestion through gas-phase charge reduction to simplify glycoprotein mass spectra.^26–29^ Charge reduction has been valued for its ability to simplify spectral interpretation for decades^30–43^, and as proton-transfer charge reduction (PTCR) has become more widely accessible on commercial instruments^44–49^, it has emerged as a potentially useful tool for charting glycoproteoform modification states.^26,27^ A related approach, electron capture charge reduction (ECCR), has also demonstrated utility toward this goal,^28,29^ although ECCR (more commonly known as non-dissociative electron capture, or ECnoD) is typically only successful on native proteins with low enough charge density to not undergo electron-driven fragmentation under normal conditions (any species susceptible to dissociation upon electron capture will undergo ECD in addition to ECCR. PTCR, on the other hand, does not drive protein backbone fragmentation and generates stable, non-radical product ions, giving it significant advantages as a more universal strategy. Recently, data-independent acquisition (DIA) strategies have used quadrupole isolation of specific m/z regions for PTCR or ECCR reactions (instead of charge reducing the entire ion population) to reduce complexity and improve signal-to-noise.^26^ In these DIA schemes, quadrupole isolation iterates through adjacent windows across a defined m/z range to generate a collection of charge-reduced spectra that are stitched back together into glycoproteoform populations.

Despite the advantages DIA-PTCR and DIA-ECCR have demonstrated thus far, significant challenges in current iterations include relatively wide isolation windows used to select precursor ions and a reliance on low-to-moderate resolving powers (∼10,000-30,000) for spectra collected following the charge-reduction reaction. Wider isolation windows yield more complex precursor-ion populations, resulting in a lower signal-to-noise ratio after charge reduction. Low resolving power translates into low mass accuracy, resulting in poor resolution of glycoforms and large mass errors that hurt specificity. These choices have in part been guided by general practices used in native MS workflows^50^, including spectral deconvolution with UniDec,^51^ which excels at processing low-resolution data lacking isotopic information. When applied to complex glycoprotein spectra or isotopically resolved data, however, current software solutions generate imprecise deconvoluted glycoproteoform masses with broad mass distributions that are difficult to identify. Furthermore, low-resolving-power acquisitions suffer from broad peaks comprised of merged signals of closely spaced m/z values for which only the average mass can be obtained. This merged peak information can be valuable for defining general trends in glycoproteoform mass distributions, but assigning individual glycoproteoform species from these relatively low mass accuracy peaks remains nearly impossible. One DIA-PTCR study noted an example of a glycoproteoform resolved within a ±5 Da window using PTCR could still be explained by more than 600,000 discrete glycoproteoforms predicted from site-specific glycosite information derived from bottom-up glycoproteomic data.^26^ They further noted that the lack of discrete glycoproteoform assignment also lowered precision in determining the number of monosaccharides present for each glycoform. Ultimately, this mismatch in intact glycoproteoform mass assignments and potential glycoproteoform composition limits the sensitivity and specificity with which we can connect glycocode regulation to biological significance.

Here, we demonstrate an improved approach to map glycoproteoform landscapes with ∼10ppm mass accuracy that leverages narrow quadrupole isolation (5 m/z windows) for DIA-PTCR acquisition and high-resolving power (hRp) Orbitrap m/z analysis (Rp ∼ 240,000 at m/z 200), a strategy we refer to as hRp-DIA-PTCR. This effort required us to develop a new informatics strategy to process hRp-PTCR spectra, mainly because previous studies have used resolving powers insufficient to resolve isotopic envelopes. Moreover, isotopically resolved data are not readily compatible with the existing deconvolution tools used in these workflows, largely due to limited control over spurious identifications. We, however, found isotopically resolved spectra to be highly beneficial, as they avoid peak merging of proteoforms with similar masses or incorrect peak summation due to insufficient ion statistics (as recently noted by Fornelli and co-workers^52^, as well). To process this data, we created a GUI-based tool called PTsliCR to clean PTCR spectra directly from DIA-PTCR raw files, and we built an R package called IsoTrac for spectral deconvolution. Both informatic tools are freely available on GitHub at https://github.com/riley-research. Altogether, we demonstrate the analytical power of hRp-DIA-PTCR by characterizing translationally relevant glycoproteins (EpCAM, TIGIT, CD40, PDL1, and CD24), and we use EpCAM as an example to show how hRp-DIA-PTCR can be used in concert with bottom-up glycoproteomics and Direct-Mass Technology (DMT, i.e., Orbitrap-based CDMS) to capture heterogeneous populations of glycoproteoforms that can guide biological interpretation.

## EXPERIMENTAL PROCEDURES

### Native glycoprotein preparation

Ten µg each of recombinant human glycoproteins [EpCAM (ACROBiosystems, #EPM-H5223, AAH14785.1), TIGIT (R&D Systems, #9525-TG, Q495A1-1), CD40 (ACROBiosystems, #CD0-H5228, P25942), PDL1 (ACROBiosystems, #PD1-H5223, Q9NZQ7), and CD24 (ACROBiosystems, #CD4-H52H3, P25063-1)] were individually buffer-exchanged into 150 mM aqueous ultrapure ammonium acetate (pH 7.2) by ultrafiltration (Millipore Amicon) with a 10 kDa cutoff filter. The protein concentration was adjusted to 2–3 μM prior to native mass spectrometry analysis.

### Glycopeptide sample preparation

Ten µg of each glycoprotein was prepared separately in parallel in a 100 mM HEPES (4-(2-hydroxyethyl)-1-piperazineethanesulfonic acid) buffer solution^53^, subjected to a 5-minute heat shock at 95 °C, reduced with 5 mM tris(2-carboxyethyl)phosphine hydrochloride (TCEP) for 30 min at 60 °C, and alkylated in the dark with 20 mM chloroacetamide for 30 min at room temperature, followed by a 15-min quench using TCEP. Sodium deoxycholate (SDC) was added to a final concentration of 1%. Overnight digestions were performed at 37 °C with a) trypsin (Promega, 1:50 μg/μg protease:protein) for EpCAM, CD40, PDL1; b) chymotrypsin (Promega, 1:70 µg/µg protease:protein) for EpCAM and PDL1; c) sequential trypsin and chymotrypsin for CD40 and TIGIT; or d) sequential trypsin and IMPa (NEB, 1:10 μg/μg protease:protein) for CD24. One CD24 sample underwent N-glycosylation removal with 500 units of PNGase F (New England Biolabs) for 4 h at 37 °C after proteolytic digestion. The SDC in all samples was precipitated twice with 2% formic acid (FA), after which the samples were desalted using Strata-X 33 µm 10 mg/1 mL Polymeric Reversed Phase SPE cartridges (Phenomenex) by conditioning the cartridge with 1 mL acetonitrile (ACN) followed by 1 mL 0.2% formic acid in water. Peptides were then loaded onto the cartridge, followed by a 1 mL wash with 0.2% FA in water. Peptides were eluted with 400 μL of 0.2% FA in 80% ACN and dried via vacuum centrifugation. After drying, all samples were stored at −70 °C until mass spectrometry analysis.

### Bottom-up glycoproteomics acquisition

Glycopeptides were analyzed on an Orbitrap Ascend Tribrid Mass Spectrometer (Thermo Fisher Scientific) coupled to a Vanquish Neo UHPLC (Thermo Fisher Scientific) using the Orbitrap Ascend Tune Application (v4.2.4321). First, samples were reconstituted in 0.1% FA and loaded on a C18 trap column (300 µm × 5 mm, 5 µm particles, PepMap Neo) using a 300 nL/min flow using buffer A (0.1% formic acid). Subsequently, samples were linearly eluted using a 55-min gradient ranging from 2.2% buffer B (99.9% acetonitrile with 0.1% FA) to 28% buffer B on a C18 analytical column (75 µm × 25 cm, 1.7 µm particles, IonOptics). The column was washed for 5 min with 99% B after separation and equilibrated with 100% buffer A. Precursors were ionized using a nanospray flex ionization source (Thermo Fisher Scientific) held at +2.0 kV compared to ground, and the inlet capillary temperature was held at 275 °C. The instrument method collected spectra in a data-dependent fashion using full scan (MS1) settings of 120,000 resolving power at 200 m/z, a normalized AGC target of 250% (1,000,000 charges), and an MS1 mass range from 400 to 1800 m/z. Dynamic exclusion was set at 15 s, and only charges 2–6 were selected for fragmentation. Precursor ions were isolated for MS/MS spectra using a 0.7 m/z quadrupole isolation width, and product ions were mass analyzed in the Orbitrap with an MS2 resolution of 30,000 at 200 m/z using a normalized AGC target set to 200% (100,000 charges) and a maximum injection time of 59 ms. Selected precursor ions were fragmented with stepped collision energy/higher-energy collisional dissociation (sceHCD) performed using 20, 30, and 40% normalized collision energies. sceHCD MS/MS spectra containing at least four glycan-specific oxonium ions (126.055, 138.0549, 144.0655, 168.0654, 186.076, 204.0865, 274.0921, 292.1027, and 366.1395 m/z) triggered collection of electron-transfer/higher-energy collision dissociation (EThcD) MS/MS spectra, where precursor ion isolation was performed with the quadrupole using a 1.6 m/z isolation window, an AGC target of 200% (100,000 charges), and a maximum injection time of 251 ms. EThcD fragmentation was performed using a 50 ms reaction time, a 2e6 reagent target, and a m/z range from m/z 120-3,600 based on previous studies.^54–56^

### Direct mass technology measurements (DMT) acquisition

Samples were analyzed using an Orbitrap Ascend Tribrid Mass Spectrometer operating in DMT mode at low pressure (3 mTorr N2 as measured in the IRM). Samples were ionized using borosilicate nanoelectrospray emitters coated with gold (Thermo Fisher Scientific). The spray voltage was set to 1.5 kV, source fragmentation was 60 V, and source temperature was 300 °C. For DMT collection, automated ion control (AIC) and 500% target density were used with an Orbitrap transient length was set to 1 second.

### High-resolving power data-independent proton transfer charge (hRp-DIA-PTCR) acquisition

Prior to DIA-PTCR acquisitions, MS1 spectra were acquired to inform DIA-PTCR acquisition windows (i.e., spectra at different m/z positions) using an identical system setup to that used for the DMT acquisitions. Using these acquisitions, the DIA-PTCR scan ranges were set to 2600-2950 m/z for CD40, 2800-3350 m/z for EpCAM, 2200-2600 m/z for TIGIT, 2600-3800 m/z for PDL1, and 2400-3200 m/z for CD24. The spray voltage was set to 1.5 kV, the source fragmentation to 60 V at 60% RF, and the source temperature to 300 °C. Isolation widths were set to 5 Th using a 0.5 Th overlap between windows, and isolation was performed using an extended-range quadrupole, which can isolate up to m/z 8,000. Orbitrap resolving power was set to 240,000 at m/z 200, the PTR reagent target to 2e6, the PTR reaction time to 35 ms, a maximum injection time of 1 s, and an AGC target of 4000% (2,000,000 charges).

### Glycopeptide data analysis

All samples, except CD24, were analyzed separately with FragPipe (v24.0)^57^ using the Glyco-N workflow, with the protein sequence to which decoys and contaminants were added as part of the FragPipe workflow. Protein sequences were used based on vendor supplied sequence information. Default settings were used unless specified otherwise. In short, the search was performed using a 20 ppm precursor mass error and fragment mass tolerance, two allowed miscleavages (KR), a maximum of 3 variable modifications (M oxidation, N-terminal acetylation), and fixed modifications set to carbamidomethylated C. The ‘Human_N-glycans-medium-253’ glycan database was used with a 1% glycan FDR. Modified peptide output files are available as supplemental material.

CD24 raw files were searched separately using Byonic (v5.5.2) using the CD24 sequence and common contaminants as supplied by Byonic. The PNGase F-treated samples were searched against the Byonic O-glycan database consisting of 78 glycans, and the non-PNGase F-treated sample was searched against the O-glycan database combined with Byonic’s N-glycan database, which consisted of 309 unique mammalian N-glycan compositions. Carbamidomethylation was set as a fixed modification, and oxidation on methionine, deamidation on N, and acetylation on the N-terminus were used as variable modifications. A maximum of 3 miscleavages was allowed, and a maximum of three glycosites was allowed for any one glycopeptide. Filtering metrics included a Byonic score greater than or equal to 200 and a logProb value greater than or equal to 2.

We manually verified all identified glycopeptide spectra for accurate identification. Only glycopeptides were retained that met the following criteria: at least a 50% sequence coverage after removal of unreliable fragment annotations (e.g., too large mass error, c- or z-ions n-terminal of proline, or improbable isotope intensities), an unambiguous glycan assignment (e.g., biologically plausible glycan compositions, or no multiple fucosylation with high Neu5Ac oxonium signal), an oxonium ion signal in line with the glycan composition (e.g., >10% of TIC is oxonium ion-derived, Neu5Ac diagnostic ions found when composition has Neu5Ac), and, where observed, glycan-loss fragments in line with the glycopeptide identification, such as HexNAc-only residues, and Y-ion series. The absence of Neu5Gc was verified by oxonium-ion mining in MS/MS spectra with GlyCounter.^58^ Visualization and analysis of the bottom-up glycoproteomic data were performed using GlycoDiveR.^59^

### DMT analysis

DMT analysis was performed using STORIboard (build 1.0.21204.1) using the ‘Voting’ algorithm.^60^ Settings used included a 0.9 r2 threshold, 0.2 duration threshold, 0 minimum time of death, 0.5 maximum time of birth, 1 S/N threshold, frequency correction is false, 3 bin size (ppm), minimum ions in bin are 1, 2 charge neighbors, and 5 isotope neighbors. The data were exported from STORIboard and replotted using R (version 4.5.3).

### DIA-PTCR analysis

The first step in analyzing the DIA-PTCR data is to extract and clean the PTCR spectra using the in-house-built tool PTsliCR, available on GitHub. PTsliCR is an open-source tool written in C# that uses the C# Mass Spectrometry Language (CSMSL, https://github.com/dbrademan/CSMSL/) to access spectral information. It reads PTCR spectra directly from raw files and determines the regions of m/z where ions could appear based on the precursor m/z, the isolation width, and the possible charge states within the isolation window. For example, a precursor window of *m/z* 2000-2005, with a possible charge set to 3, can only charge-reduce to a 3000-3007.5 m/z and a 6000-6015 m/z PTCR slice. Precursor charge states for the m/z range interrogated in the PTCR analyses were determined using the DMT data and set to 7-13 for PDL1, 5-9 for TIGIT, 5-15 for CD24, 7-11 for CD40, and 6-12 for EpCAM. All intensities outside these windows are set to 0; subsequently, only one flanking 0 is retained next to the intensity values to allow centroiding. The output for each .raw file consists of a cleaned .mgf file and quality-control plots.

The .mgf output file is directly compatible with our in-house, open-source R analysis workflow with integrated C++ called IsoTrac, available on GitHub. IsoTrac reads the mgf files, performs an initial clean by applying a user-set peak intensity threshold and removes empty PTCR slices. It then converts profile data to centroided data by fitting a cubic spline function through each isotope peak and determines the charge state of each slice using the deltas between m/z values. Subsequently, per spectrum and PTCR slice, the average glycan mass is determined using:

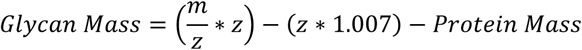

The protein mass is the average mass calculated using the protein’s amino acid composition. The total chemical composition is determined using the protein’s amino acid composition and by using the glycan’s mass times the average glycan composition normalized to one Dalton (C = 0.038321192, H = 0.062956245, N = 0.002737228, O = 0.027372280), conceptually similar to an average protein isotope distribution.^61^ The resulting atomic composition of the glycan is rounded to the nearest integer. The composite isotope distribution of protein and glycan is obtained using the EnviPath R package.^62^

IsoTrac calculates cosine score, Pearson correlation, and normalized root-mean-square error (NRMSE) between the theoretical isotope envelope and the measured isotope distribution. To enable calculation of the false discovery rate (FDR), an equal number of decoy spectra are generated as the number of isotope matches (targets). Each decoy spectrum has the same m/z values as its template target spectrum. The peak intensities are randomized using the lower and upper limits of the intensities of the template target spectrum, except for every nth peak (where n is the number of isotopes next to the highest isotope peak used for matching, e.g., it will be four for a nine-isotope matching approach), which is limited to the lowest one-third of the intensity range. This prevents the generation of valid isotope patterns in the decoy spectra (because even random peak intensity variations can create valid isotope pattern peak shapes that would then exist in the “decoy” set), which decrease the reliability of the target-decoy approach. The correlation values are sorted, and the FDR is calculated using:

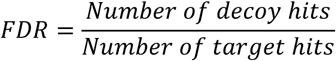

To allow for the deconvolution of overlapping isotope envelopes, the number of isotope peaks used by the matching algorithm can be set; for example, the unmodified EpCAM protein corresponds to the chemical formula C_1221_H_1936_N_348_O_388_S_16_, which has 25 theoretical isotopes with intensities above a 1% threshold of the maximum isotope intensity. Using only the nine most intense isotopes allows accurate deconvolution even when the measured isotope distributions partly overlap. To ensure reliable matches, IsoTrac does not allow any gaps in the matched isotope sequence and uses a user-modifiable minimum number of isotope peaks to be matched, ensuring that cutoff isotope envelopes are not considered. Settings used for CD40/TIGIT analysis were: a Pearson cutoff of 0.9 and an NRMSE cutoff of 45, at least isotopes matched from -3 to 3 (where -3 is the third isotope left of the most intense isotope peak), and 9 isotope peaks in total were used for matching. Settings for EpCAM analysis used the same values, but with a Pearson cutoff of 0.88. Settings for CD24 analysis used seven isotope peaks for matching. Summarization of extracted intact masses is done by taking the median in 0.3 Da bins per spectrum, and subsequently the median in 0.03 Da bins per raw file.

IsoTrac generates a PDF file that displays the empirical spectrum overlaid with the theoretical isotope match for all matches. It also outputs detailed information for each isotope pattern match, including correlation scores, spectrum number, and peak m/z value, making it easy to link extracted intact masses to the isotope matches from which they were deduced. Isotope pattern matches were manually verified to ensure accurate intact mass extraction for each sample. These files are attached as as supplemental file S1 and S2.

#### Glycoproteomic-informed site occupancy prediction of the DMT data

The simulated intact spectrum derived from the bottom-up data and the DMT data was used to determine the macroform distribution (i.e., EpCAM with one, two, or three occupied sites). To this end, a LOESS model was fitted to the simulated spectrum for each glycan occupancy group (e.g., EpCAM with one site occupied) to determine the relative contribution of each macroform per 10 m/z bin. The total signal per bin in the DMT data was then multiplied by the contribution of each macroform. This provided the macroform distribution, which was visualized using the DMT data.

#### UniDec deconvolution of TIGIT

Intact MS1 measurements of TIGIT at a 7.5 Orbitrap resolution were deconvoluted using UniDec (v8.1.2).^51^ Mass ranges of *m/z* 1750-3200 were used for deconvolution, charge states of 4-12, and sampled at 1 Da intervals. We used split G/L with a peak FWHM of 2.0, a beta of 40, a charge-smooth width of 1.0, and a point-smooth width of 40. The peak detection range was set to 5.0 Da, and the peak detection threshold to 0.01.

#### Glycoprotein modeling

The crystal structure of a mutated EpCAM construct had been previously determined (PDB ID: 4MZV).^63^ In that structure, glycosylation sites were mutated to glutamine residues whereas for the present glycoprotein modeling, these positions were reverted to asparagine residues. Glycans were selected based on their relative signal intensities observed in DIA-PTCR experiments. Glycan structures were modeled on sites N74 (GlyTouCan ID: G28959FS) and N198 (GlyTouCan ID: G27169KC) using the Re-Glyco tool implemented in the GlycoShape platform, with glycans selected from the GLYCAM database and inserted using default parameters.^64^ Structures were rendered with VMD,^65^ and the SNFG representations of the glycan structures were visualized using a plugin from Glycam-Web.^66,67^

## RESULTS AND DISCUSSION

### High-resolving power DIA-PTCR as a tool to access the glycoproteoforms

Understanding the glycocode and its biology requires studying glycoproteins at the intact level, a frontier in glycobiology that remains underdeveloped. Consider, for instance, Epithelial Cell Adhesion Molecule (EpCAM), a type I membrane protein of 314 amino acids that readily forms cis-dimers and regulates cell adhesion through interactions with numerous cell adhesion molecules.^68,69^ EpCAM has long been known to be involved in physiological processes, such as the organization of epithelial tissue architecture and cell proliferation, and is also implicated in numerous cancer types.^69,70^ These diverse functions are coordinated by its three N-glycosylation sites (N74, N111, N198). For example, tumor-implicated EpCAM exhibits distinct glycosylation patterns, knockout of glycosylation sites leads to EpCAM-mediated regulation of proliferation by enhancing autophagy in breast cancer cells via the PI3K/Akt/mTOR pathway, and hypoxia induces epithelial-to-mesenchymal transition through targeted N-glycosylation of EpCAM in breast cancer cells.^71–73^ Fully understanding EpCAM’s glycocode requires measuring glycosylation in its intact molecular context, because biological function emerges from the complete glycoprotein proteoform rather than the isolated glycosylation sites alone.^74^

To understand EpCAM’s glycoforms, we first quantified the site-specific heterogeneity of human EpCAM by digesting EpCAM in parallel using trypsin and chymotrypsin and used bottom-up glycoproteomics to quantify intact glycopeptides. Glycopeptides covering all three putative N-glycosylation sites were quantified, with the glycan intensities spanning a ∼500-fold dynamic range, with notably a high degree of fucosylation (**Figure 1a**). There were seven, fourteen, and seventeen distinct glycan compositions detected for N198, N74, and N111, respectively. To bridge this site-specific glycan quantification with combinatorial glycosylation profiles, we next measured intact EpCAM natively using high-resolution Orbitrap mass spectrometry and Direct Mass Technology (DMT), also known as charge-detection mass spectrometry (CDMS), a leading approach for intact measurements of highly complex samples.^22,75^. Unsurprisingly, the high sample complexity due to glycan heterogeneity prevented accurate mass inference for glycosylated EpCAM (**Figure 1** and **Figure S1**). No intact mass information could be gleaned from native MS alone, and only the exact mass of unglycosylated EpCAM could be readily extracted by native MS with DMT. The mass distribution in **Figure 1b** obtained from DMT measurements shows that the glycoforms appeared as a “mass smear” ranging from approximately 29.5 to 35.5 kDa, where there are effectively peaks at every mass value instead of distinct glycoproteoform masses that could be realistically coupled with bottom-up glycosite-level data. This complexity highlights the need for accessible methods that can infer intact masses from highly complex glycoproteins.

**Figure 1.**
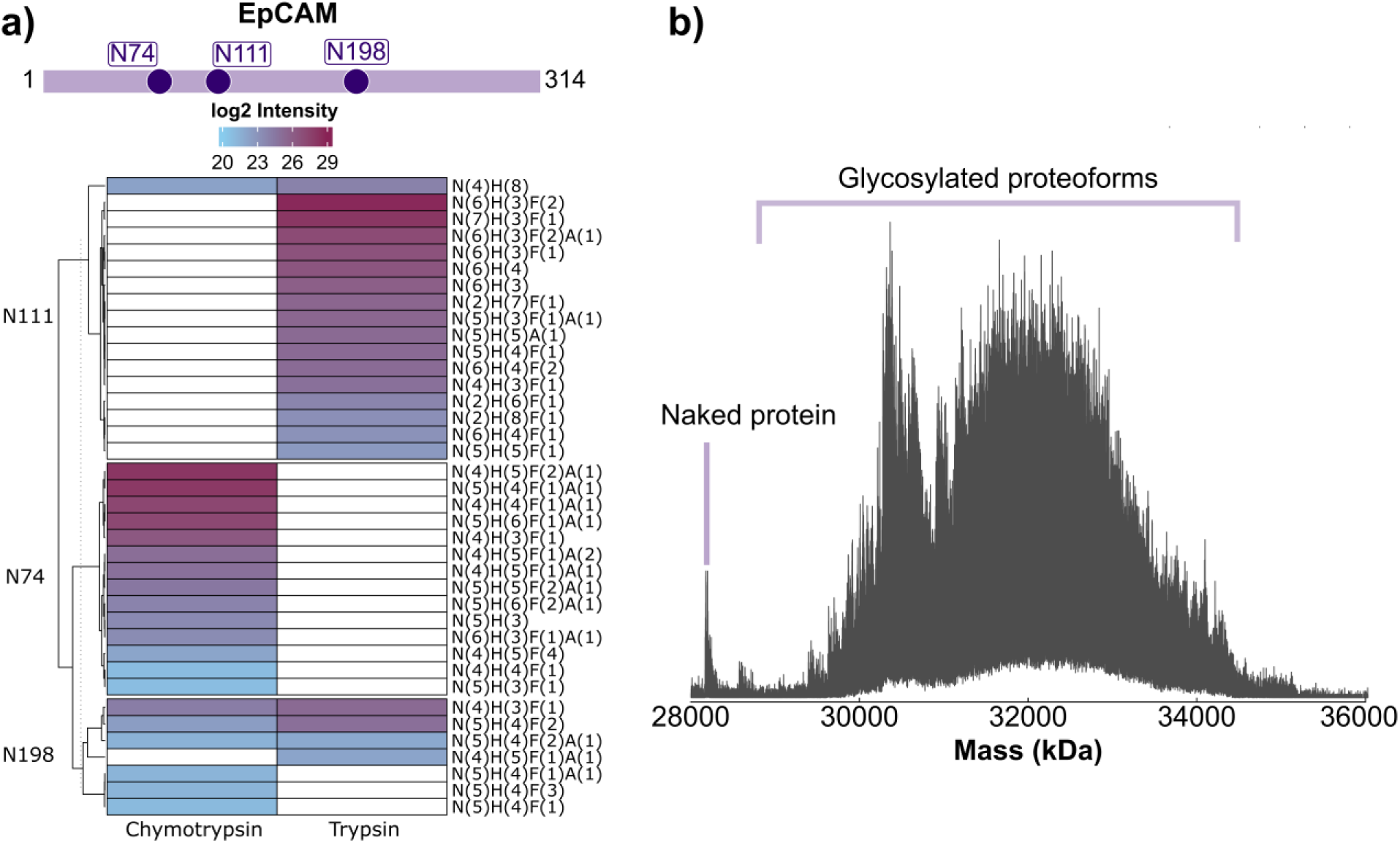
Heterogeneous glycosylation of EpCAM. (**a**) EpCAM’s site-specific glycosylation patterns were quantified using a bottom-up intact glycopeptide mass spectrometry workflow. Glycans were detected on all three potential N-glycosylation sites, and their intensities are shown in the heatmap. ( **b**) Intact EpCAM was subjected to DMT measurements, revealing the naked protein and a ∼5 k Da distribution of EpCAM glycoforms.

To approach glycoproteoform measurements from a different angle, we turned to data-independent acquisition coupled with proton transfer charge reduction (DIA-PTCR), which has recently emerged as a powerful approach for elucidating intact masses of glycoproteins.^26^ Enabled by the newly developed high-mass-range quadrupole of the Orbitrap Ascend Structural Biology Tribrid Mass Spectrometer that can isolate up to m/z 8000, we used 5 mz isolation windows for our DIA-PTCR data collection, i.e., the narrowest isolation that can be performed with that quadrupole. After isolation, the PTCR ion-ion reaction was conducted in the high-pressure linear ion trap, where precursor cations were charge-reduced using perfluoroperhydrophenanthrene (PFPP) as a PTCR reagent anion. Subsequent mass analysis was performed with the Orbitrap (**Figure 2a**). We compared DIA-PTCR acquisitions of EpCAM using 100 spectral averages at various Orbitrap resolving powers ranging from 7.5k to 480k (all defined at m/z 200). At lower resolving powers (mostly 7.5k and 15k), we observed broad, unresolved peaks consisting of multiple underlying glycoforms (**Figure 2b**), which is detrimental to discerning individual glycporoteoforms. To illustrate this challenge, Schachner et al. report a mass of 98,224 Da of IL22-Fc binned in a 5 Da window they could resolve with 7k Orbitrap resolving power. They calculated >600,000 glycoproteoforms that could be explained within that mass window, showing a clear limitation of intact glycoproteoform measurements with these lower-resolving-power approaches.^26^ Furthermore, intermediate resolving powers (30k, 60k and 120k), were insufficient to afford isotopic resolution (**Figure 2b**). Importantly, while the 30k spectrum showed good peak m/z inference in the representative EpCAM DIA-PTCR slice, this was not consistent throughout a full DIA-PTCR acquisition, as it also suffered from sporadic unresolved peaks arising from low-abundance analyte signal and peaks covering part of the isotope envelope. The only consistently reliable acquisitions were those using higher Orbitrap resolving powers (240k and 480k), in which we observed resolved isotopes that provided reliable peak information. Additionally, higher resolving powers can increase the signal-to-noise, particularly if ion trajectories are stable during Orbitrap measurement.^76^ To balance acquisition speed and reliable peak reconstruction, we selected a resolving power of 240k for subsequent high resolving power (hRp)-DIA-PTCR acquisitions.

**Figure 2.**
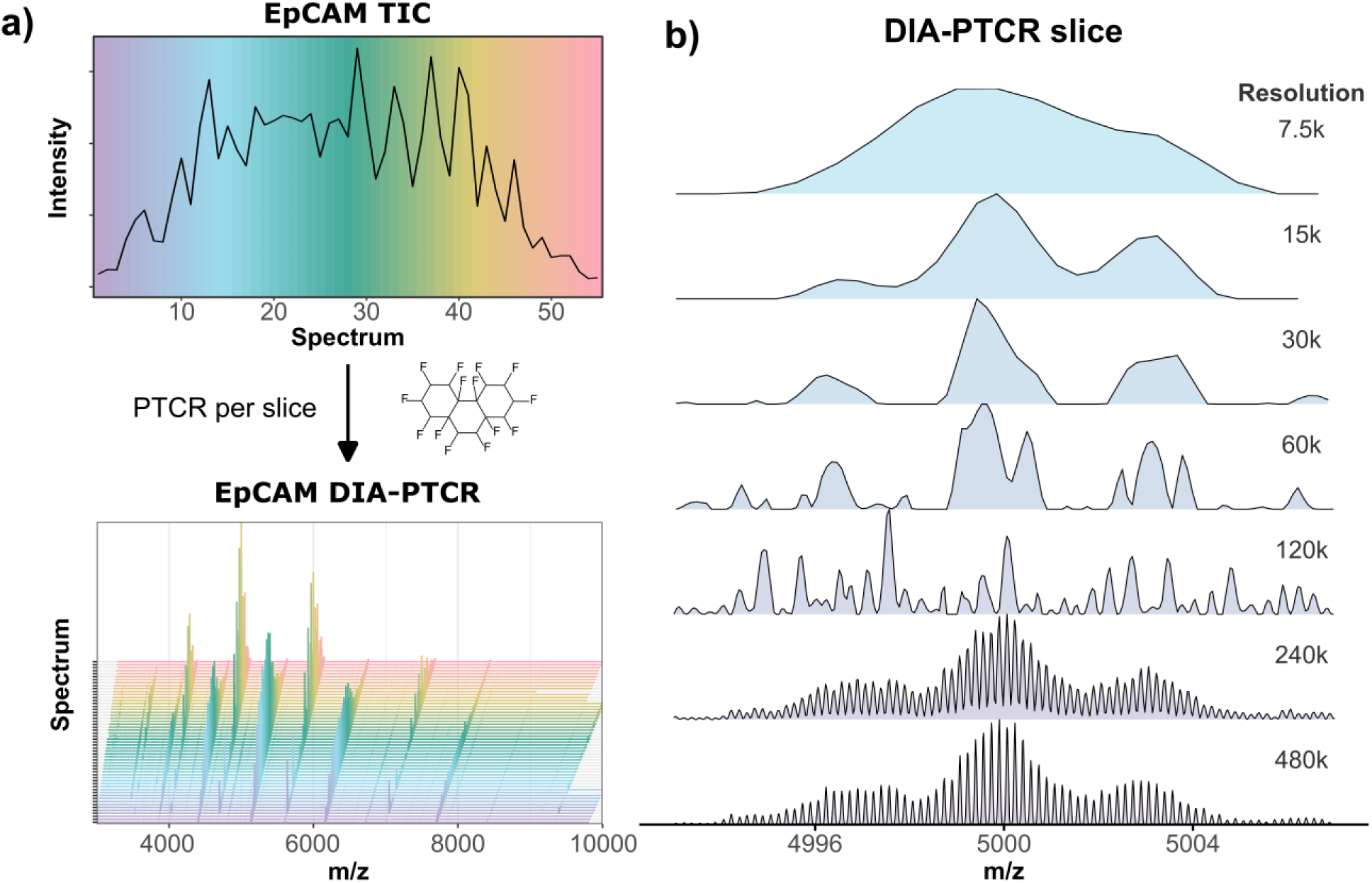
hRp- DIA-PTCR reveals isotopically resolved glycoproteoform peaks in highly complex samples. (**a**) In the hRp-DIA-PTCR workflow, consecutive 5 m/z windows are quadrupole-isolated across a defined mass range, and all precursors within the isolation window are subjected to PTCR using the perfluoroperhydrophenanthrene reagent anion. Overlapping precursors with different charge states occupy distinct m/z regions after charge reduction to disentangle overlapping species. (**b**) A single EpCAM hRp-DIA-PTCR slice ( i.e., defined narrow m/z region within an PTCR MS/MS spectrum, 100 spectra averaged) was compared after acquisition at various Orbitrap resolving powers (at m/z 200). Lower resolving powers resulted in either peak merging or peak distortion, whereas resolving powers of 240k and 480k yielded reliable, isotopically resolved peaks.

### Isotope envelope guided spectral deconvolution

Numerous algorithms have been developed for charge deconvolution of intact mass spectra,^51,77–82^ yet none were able to satisfactorily deconvolve our isotopically resolved hRp-DIA-PTCR data, with the main bottlenecks being difficulty handling overlapping isotope envelopes and erroneous peak assignments. We therefore developed a dedicated charge-deconvolution workflow tailored to hRp-DIA-PTCR data, leveraging isotopically resolved peaks and enforcing parsimonious peak assignment. This is critical for highly heterogeneous glycoproteins, where the amount of closely related masses makes spurious deconvolution events difficult to detect. The workflow is directly compatible with raw data (i.e., .raw files) and consists of an initial cleaning step using PTsliCR, followed by deconvolution using IsoTrac (**Figure S2**).

The first step is extracting and cleaning hRp-DIA-PTCR spectra using our purpose-built, open-source C# tool, PTsliCR. For each MS/MS spectrum, PTsliCR determines allowable m/z ranges to consider charge-reduced product ions (i.e., “slices” of the full m/z range) based on precursor m/z, quadrupole isolation width, and possible charge states. Only peaks within these slices are retained (**Figure 3a** and **Figure S3**), providing a data-driven strategy to remove chemical noise that can produce deconvoluted masses that do not correspond to genuine precursor ions. For each input .raw file, PTsliCR outputs a cleaned .mgf file and quality-control plots. The exported .mgf is directly compatible with our second tool, the IsoTrac R package. For each PTCR slice, which corresponds to a single m/z range calculated from charge-reduced values of the precursor ion, elemental compositions of possible glycoprotein species are derived from the protein sequence and the inferred glycan contribution. The glycan mass is determined as the residual mass, which is the total neutral mass (m/z × z – z × proton mass) minus the protein mass. Next, the protein’s elemental composition is directly calculated from its amino acid sequence. To model glycan composition, an average glycan elemental formula normalized per Dalton is used and scaled by the inferred glycan mass to determine each PTCR slice’s glycan-specific elemental contribution (**Figure 3a**). IsoTrac generates a theoretical isotope distribution using the Envipath package^62^ and calculates a Pearson correlation, cosine score, and normalized root mean square error (NRMSE) using a sliding-window approach (i.e., a correlation score is calculated for each position in the spectrum).

**Figure 3.**
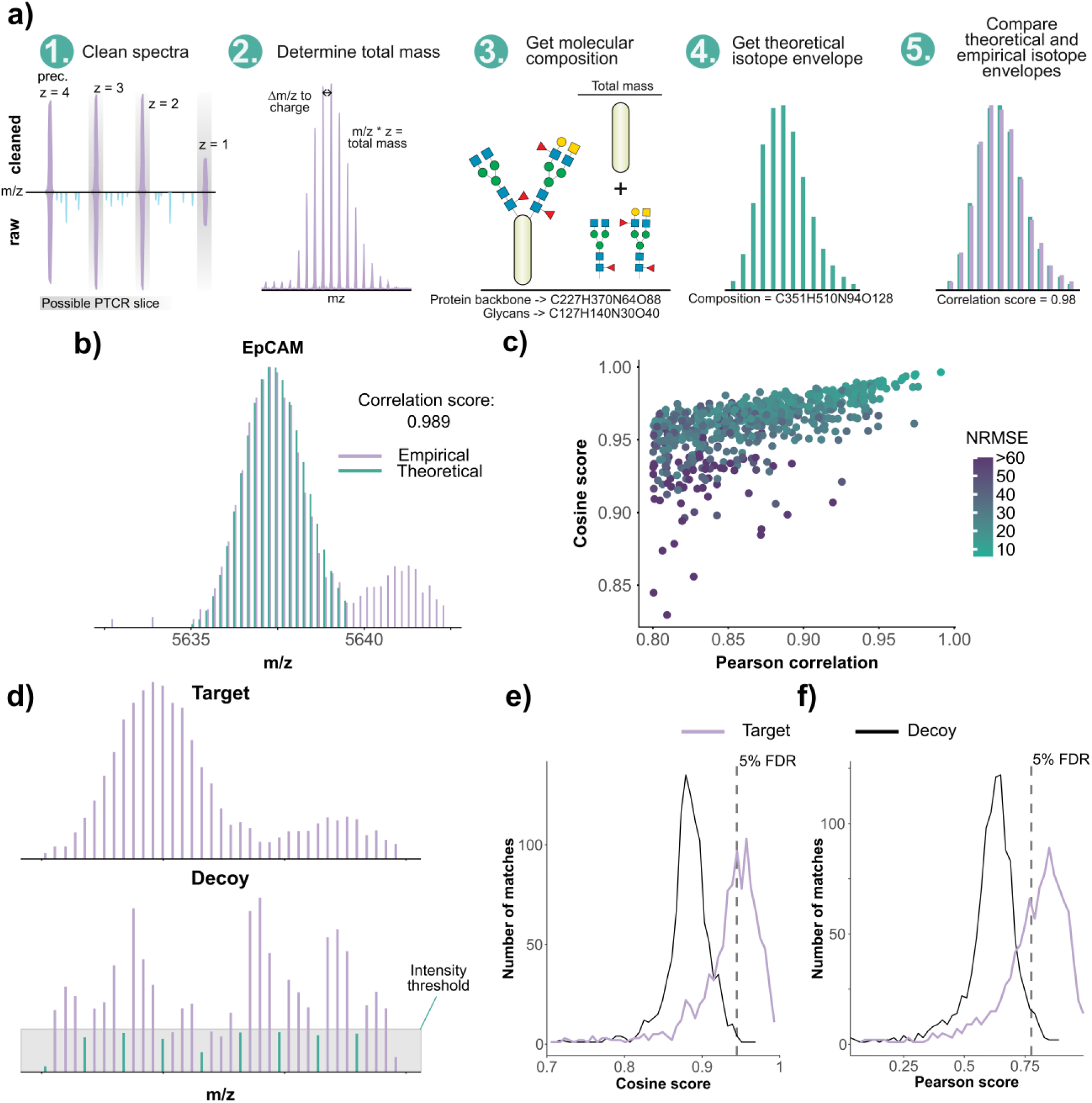
Isotope matching using theoretical isotope envelopes allows peak extraction. (**a**) Overview of the IsoTrac workflow for theoretical isotope-based peak extraction. ( **b**) An empirical EpCAM isotope envelope measured using hRp-DIA-PTCR (green) and the theoretical isotope pattern (purple). The correlation score is the Pearson correlation between the empirical and theoretical isotope patterns. ( **c**) The correlation between Pearson correlation and Cosine scoring when using isotope-based peak extraction. Points are color-coded by Normalized Root Mean Square Error (NRMSE) of the theoretical and empirical peak intensities. (**d**) A decoy spectrum is generated for each empirical spectrum by retaining the same m/z values but applying supervised randomization to the intensity values. The top spectrum shows the empirical spectrum, and the bottom shows the deduced decoy spectrum. Every nth peak (green) (n is the number of isotope peaks adjacent to the base peak in the theoretical spectrum used for matching), depending on the number of isotope peaks used for matching, has an artificial upper intensity limit (gray box) to prevent the generation of true isotope patterns. (**e**/**f**) The decoy spectra are used to calculate Pearson and cosine score cutoffs for numerous FDRs using a target-decoy approach. Empirical isotope matches are presented in purple and decoy hits in grey.

To benchmark the IsoTrac approach, we first analyzed a native MS spectrum of ubiquitin, an 8.6 kDa protein that is not glycosylated. This analysis yielded a Pearson correlation of 0.999 (**Figure S4**), indicating that the isotope matching approach could generate expected results that could be manually verified. IsoTrac also yielded high quality matches to glycoprotein spectra, as shown by a representative single EpCAM hRp-DIA-PTCR envelope that yielded a Pearson correlation of 0.989 (**Figure 3b**). Importantly, Pearson and cosine scores correlate (**Figure 3c**), but can diverge because Pearson correlation captures a linear relationship between the empirical and theoretical isotope envelope intensities, whereas cosine similarity reflects similarity in the shape of the intensities, independent of magnitude. To further improve the reliability of isotopic pattern identifications, we implemented a target-decoy approach to estimate FDR values.^83,84^ One decoy PTCR slice is generated for each empirical PTCR slice and subjected to the same sliding-window isotope pattern-matching approach as the empirical spectrum (**Figure 3d**). Correlation cutoffs are returned for different FDR values (**Figure 3e** and **3f**), but users are encouraged to verify isotope match quality with all isotope matches and adjust the correlation cutoffs accordingly. IsoTrac enables this evaluation by automatically exporting a PDF file with each isotope pattern match.

We next set out to benchmark IsoTrac deconvolution using T cell immunoreceptor with Ig and ITIM domains (TIGIT). TIGIT is a 26.3 kDa protein with two N-glycosylation sites that, in contrast to EpCAM, has isotopically resolved charge states in the MS1 spectrum amenable to deconvolution. We found strong agreement (Pearson correlation of 0.879) between intact masses determined by our hRp-DIA-PTCR workflow and those obtained by MS1-based deconvolution using UniDec (**Figure 4a** and **Figure S5**).^51^ Interestingly, only the hRp-DIA-PTCR workflow identified intact TIGIT without glycosylation. To further verify robustness, two hRp-DIA-PTCR cycles covering *m/z* 2200-2600 were acquired and the resulting IsoTrac-deconvoluted spectra compared, yielding a Pearson correlation of 0.952 (**Figure S6**). An identical two-cycle analysis of EpCAM yielded a Pearson correlation of 0.785 (**Figure S7a/b**). The lower score is the result of fluctuations in isotope-envelope intensities due to low ion statistics (**Figure S7c/d**)^85^, which reduces sensitivity by excluding true-positive assignments. Appropriately setting the maximum injection time, AGC target, and using spectral averaging are effective options to ensure sufficient ion statistics for accurate quantitative isotope envelopes.

**Figure 4.**
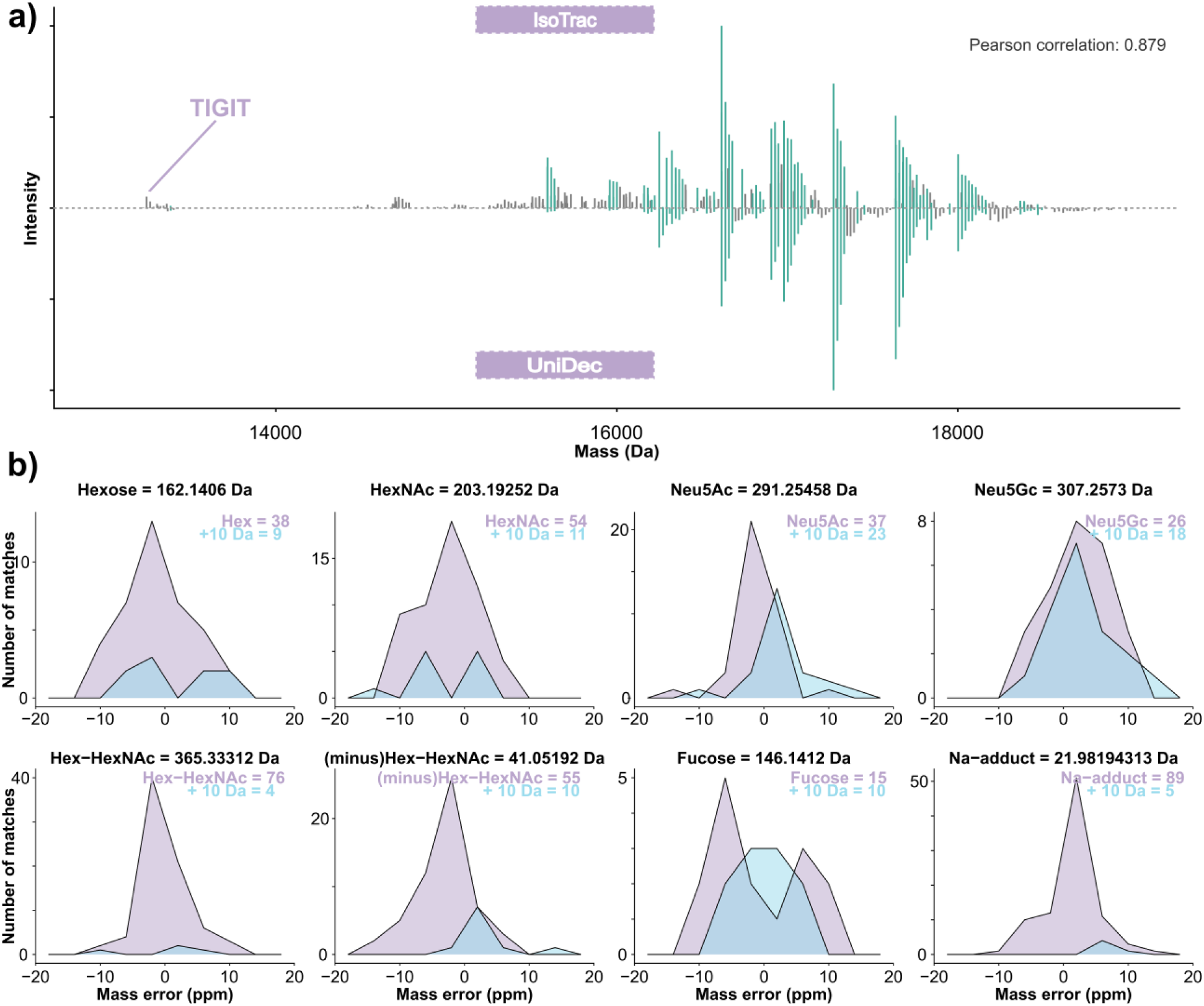
IsoTrac deconvolution of TIGIT with h Rp-DIA-PTCR. (**a**) The h Rp-DIA-PTCR workflow was used to measure intact glycoproteoform masses of TIGIT, and intact masses generated from isotope envelope matches in IsoTrac are compared to the intact masses acquired using native MS (MS1-only) with UniDec deconvolution. ^51^ The two deconvolution approaches agree well, as shown by the deconvoluted neutral masses found in both acquisitions (green lines), compared with those found in only one acquisition (grey lines). The non-glycosylated form of TIGIT, highlighted, was measured using the hRp-DIA-PTCR workflow but not by native MS with UniDec deconvolution. (**b**) Intact glycoproteins exhibit distinct mass differences arising from glycosylation. Common delta masses were extracted within a 10 ppm tolerance from deconvoluted masses following hRp -DIA-PTCR acquisition and PTsliCR and IsoTrac analyses . To rule out random matches as the driver of these hits, a decoy search encompassing a delta mass of +10 Da was performed. The target delta masses corresponding to real glycan shifts were more prevalent than the decoys, except for Neu5Gc, which is expected, as TIGIT is produced in EXPI29 3 cells that do not readily synthesize Neu5Gc.

Because the quadrupole isolation efficiency has a non-square transmission profile,^86^ we hypothesized that suboptimal isolation at the edges of the isolation windows might lead to artificial isotope patterns and thus erroneous deconvoluted neutral masses. To verify this, we collected two full DIA-PTCR cycles of TIGIT for *m/z* 2200-2600 and *m/z* 2205.5-2602.5. The two acquisitions showed good agreement (Pearson correlation: 0.896), with only a small portion of peaks identified in one acquisition but not the other (**Figure S8a**). Furthermore, spectral validation of these peaks showed that they arise from genuine signals present between adjacent DIA windows (**Figure S8b** and **Figure S8c**), highlighting the importance of window overlap and indicating that artificial isotope envelopes are unlikely to produce spurious deconvoluted neutral masses.

Finally, we calculated expected monosaccharide mass differences between deconvoluted intact mass peaks using a 20 ppm tolerance. Mass shifts we matched corresponded to hexose (Hex), N-Acetyl-Hexosamine (HexNAc), Sialic acid (Neu5Ac and Neu5Gc), Fucose, Hex-HexNAc, [minus a Hex plus a HexNAc], and sodium adducts (Na). We also used a decoy mass shift corresponding to target masses + 10 Da. The Hex, HexNAc, Neu5Ac, Hex-HexNAc, (minus)Hex-HexNAc, and Na-adduct mass shifts were much more prevalent compared to the decoys (**Figure 4b**). The fucose mass shift occurred 15 times compared to 10 times for the decoy, yet the distinct mass error distributions show this is likely due to coincidental hits to the decoy mass shift. The Neu5Gc mass shift is the only mass shift with a similar frequency and mass error distribution to its decoy, as expected, since TIGIT was produced in Expi293 cells that do not readily produce Neu5Gc-containing glycans.^87^ Altogether, IsoTrac deconvoluted spectra agree with technical and biological expectations, highlighting its utility as a deconvolution tool.

### Glycoproteoform fingerprinting with hRp-DIA-PTCR

IsoTrac should enable accurate intact-mass assignment of glycoproteins with multiple glycosylation sites and glycan variant, so we applied our hRp-DIA-PTCR workflow using PTsliCR and IsoTrac deconvolution with stringent correlation scores and NRMSE cutoffs to increasingly complex glycoproteins. A previous deep sequencing study of intact glycopeptides showed an average of 18 glycan compositions per site with an average of two N-glycosylation sites per protein across the glycoproteome.^88^ Given this level of glycan heterogeneity, intact-mass measurements are theoretically feasible for a large portion glycoproteins. We collected bottom-up intact glycopeptide-level measurements and DMT acquisitions to complement our hRP-DIA-PTCR data for CD40 (two N-glycosylation sites), EpCAM (three N-glycosylation sites), PDL1 (four N-glycosylation sites), and CD24 (sixteen O-glycosylation sites and two N-glycosylation sites) (**Figure S2**). All glycoproteins were highly complex and could not be satisfactorily deconvoluted at the MS1 level (**Figure S1** and **Figure S9**). hRp-DIA-PTCR data provided intact masses of discrete glycoforms and showed high similarity to DMT mass distributions (**Figure 5** and **Figure S10**). Each deconvoluted spectrum exhibited numerous glycan-specific mass differences, further highlighting the accuracy of the intact mass determination (**Figure 5** and **Figure S11**). We used the bottom-up glycoproteomics data to assess the complexity of each glycoprotein, where we calculated the number of potential glycoforms to be 60 for CD40; 5,280 for EpCAM; 4,802,280 for PDL1; and 10^24^ for CD24. Comparing the theoretical intact masses and the deconvoluted intact masses highlights the difficulty of using bottom-up data alone to infer glycoproteoforms, as incomplete glycan coverage or unknown modifications can lead to unexplained masses, and the same intact protein mass can often be explained by multiple glycans (**Figure 5**). Using 0.5 Da mass bins instead of 5 Da, a specificity that is enabled by our hRp-DIA-PTCR workflow, reduces the number of potential glycoforms per intact mass by about 50%. For PD-L1, the number of potential glycoforms associated with the intact mass with the greatest ambiguity decreases from 18,000 to only 9,000 (**Figure S12**). Despite this immense combinatorial complexity, IsoTrac defined intact glycoform masses with distinct glycan-specific mass deltas, supporting the use of hRp-DIA-PTCR for glycoproteoform fingerprinting for a broad range of highly glycosylated proteins.

**Figure 5.**
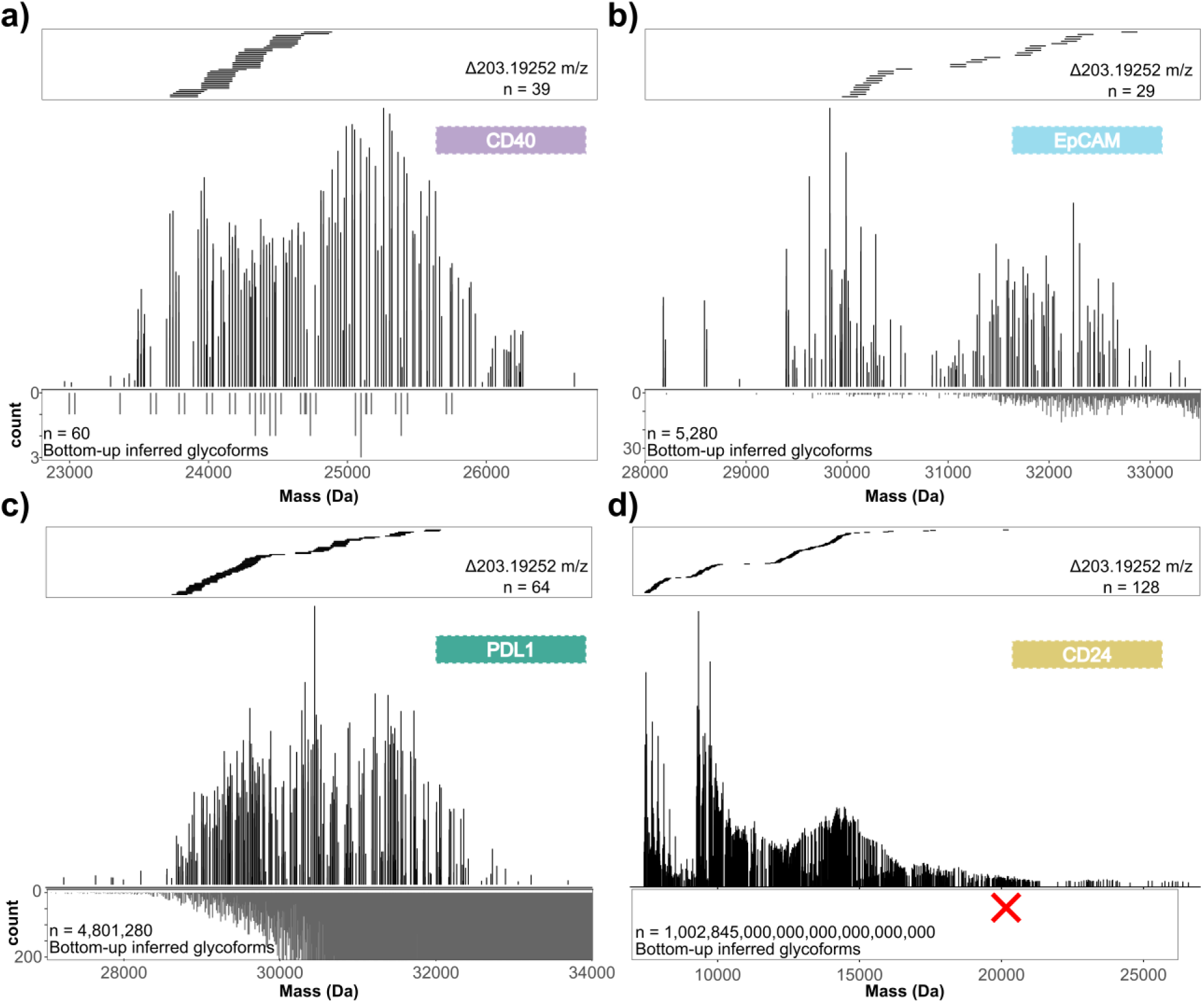
Glycoproteoform fingerprinting with hRp -DIA-PTCR. hRp- DIA-PTCR acquisitions were collected for translationally relevant glycoproteins **a)** CD40, **b)** EpCAM, **c)** PDL1, and **d)** CD24, and intact masses were extracted using the PTsliCR and IsoTrac worflow. Hex NAc delta masses between discrete glycoproteoform masses are visualized above each glycoproteoform profile. Below each profile, a histogram of the theoretical number of glyco proteoforms, as inferred from bottom-up glycoproteomics data, is shown using 0.5 Da bins. The PDL1 histogram is scaled to show up to 200 glycoforms per bin. If not scaled to the 15,000 glycoforms per bin, the lower - mass ranges would be difficult to visualize. Calculating the CD24 histogram was not feasible due to the exceptionally high memory requirements.

### Multimodal integration to decode EpCAM’s glycoproteoform profile

With this platform developed, we return to our inquiry about EpCAM glycoproteoforms. Integration of complementary analytical techniques is critical for understanding the combinatorial glycocode, as orthogonal measurements provide distinct layers of information, enable independent validation, and reduce the risk of false conclusions. We therefore compared an *in silico*-predicted EpCAM spectrum simulated from the site-specific glycopeptide-level data, the hRp-DIA-PTCR data of intact glycoproteoforms, and the DMT data from native MS (**Figure 6**). The simulated glycoproteoform fingerprint derived from bottom-up data reveals macroheterogeneity by mapping which mass ranges are occupied by specific glycoforms with defined glycan occupancies. The IsoTrac analysis of hRp-DIA-PTCR data provided distinct intact masses with relative abundances that, together with bottom-up data, can be used to infer the most likely glycan type or composition of a glycoproteoform. The DMT data functions as a valuable quality control tool and an additional quantitative layer of information. Importantly, the three techniques show good agreement of quantified glycoforms over the mass range. hRp-DIA-PTCR serves an important complementary role in generating relative abundance profiles of discrete glycoproteoform masses that match general distributions of DMT data and can be directly matched with glycosite-level information from bottom-up data to infer glycoproteoforms profiles.

**Figure 6.**
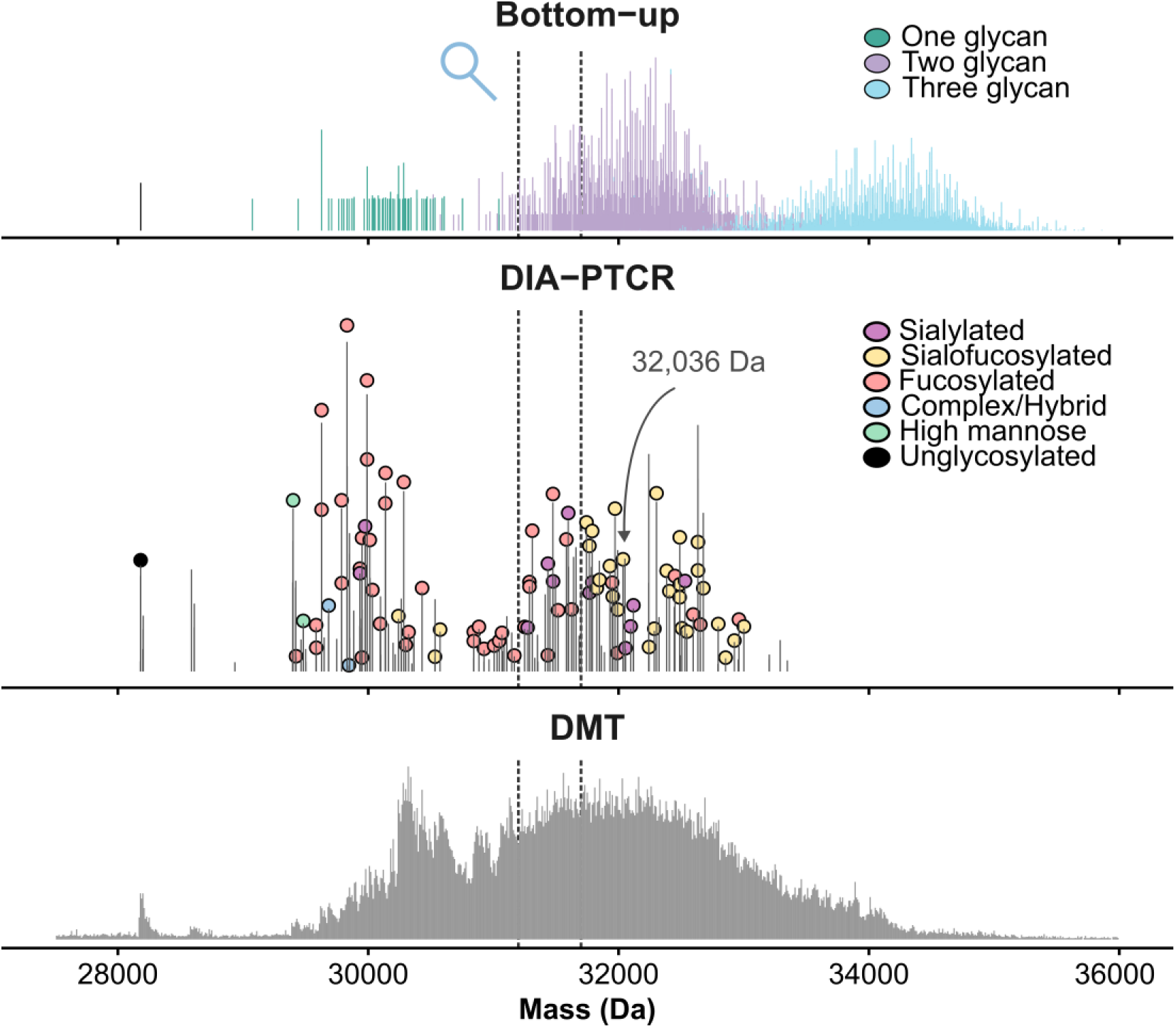
Integrating hRp-DIA-PTCR data with bottom-up glycoproteomics and DMT measurements provides complementary insights for glycoproteoform profiling. Three different approaches were used to get intact mass information of EpCAM. Glycopeptide intensities were used to generate a predicted intact spectrum (top). hRp-DIA-PTCR acquisition with PTsliCR spectral cleaning and isotope-templated deconvolution using IsoTrac generated exact intact masses (middle). EpCAM glycoproteoform masses are annotated with the most likely glycan class assignments, as informed by bottom-up glycoproteomics data. EpCAM was also measured using native MS DMT (bottom). The right side inset shows a zoom-in of the 31.2 -31.7 k Da mass range, highlighting the peak density across the different approaches.

To semi-quantitatively assess macroheterogeneity (i.e., un-, mono-, di-, and tri-glycosylated EpCAM), we estimated the relative contribution of each occupancy state per mass bin by applying a LOESS fit to the bottom-up-derived intact mass distributions for each occupancy. These simulated glycoforms were compared to the quantitative DMT data, showing that un-, mono-, di-, and tri-glycosylated EpCAM comprised 0.7%, 22.8%, 70.4%, and 6.1% of signal, respectively (**Figure 7a** and **7b**). We categorized the glycan classes of each glycoproteoform mass using glycoform-resolved monosaccharide fingerprints that integrate bottom-up data and hRp-DIA-PTCR data, which generated a probability plot for the glycan compositions per deconvoluted intact mass (**Figure 7c**).^26^ This analysis showed especially high levels of fucosylation and sialylation, which is an observation corroborated by quantification of the hRp-DIA-PTCR masses and site-specific quantification of the bottom-up data (**Figure 7 d** and **7e**). N111 showed the highest micro-heterogeneity, with about 50 distinct glycan compositions detected compared to 10 and 15 for N198 and N74, respectively. The majority of the signal for N111 and N198 was from fucosylated glycan compositions, while this was mostly for sialofucosylated compositions for N74. We note that bottom-up quantification of glycan types, while useful, is likely an imperfect estimation of absolute glycan quantity, as numerous factors influence bottom-up glycopeptide quantification, such as ionization efficiency, efficient glycopeptide generation by proteases, and glycopeptide stability.^89^

**Figure 7.**
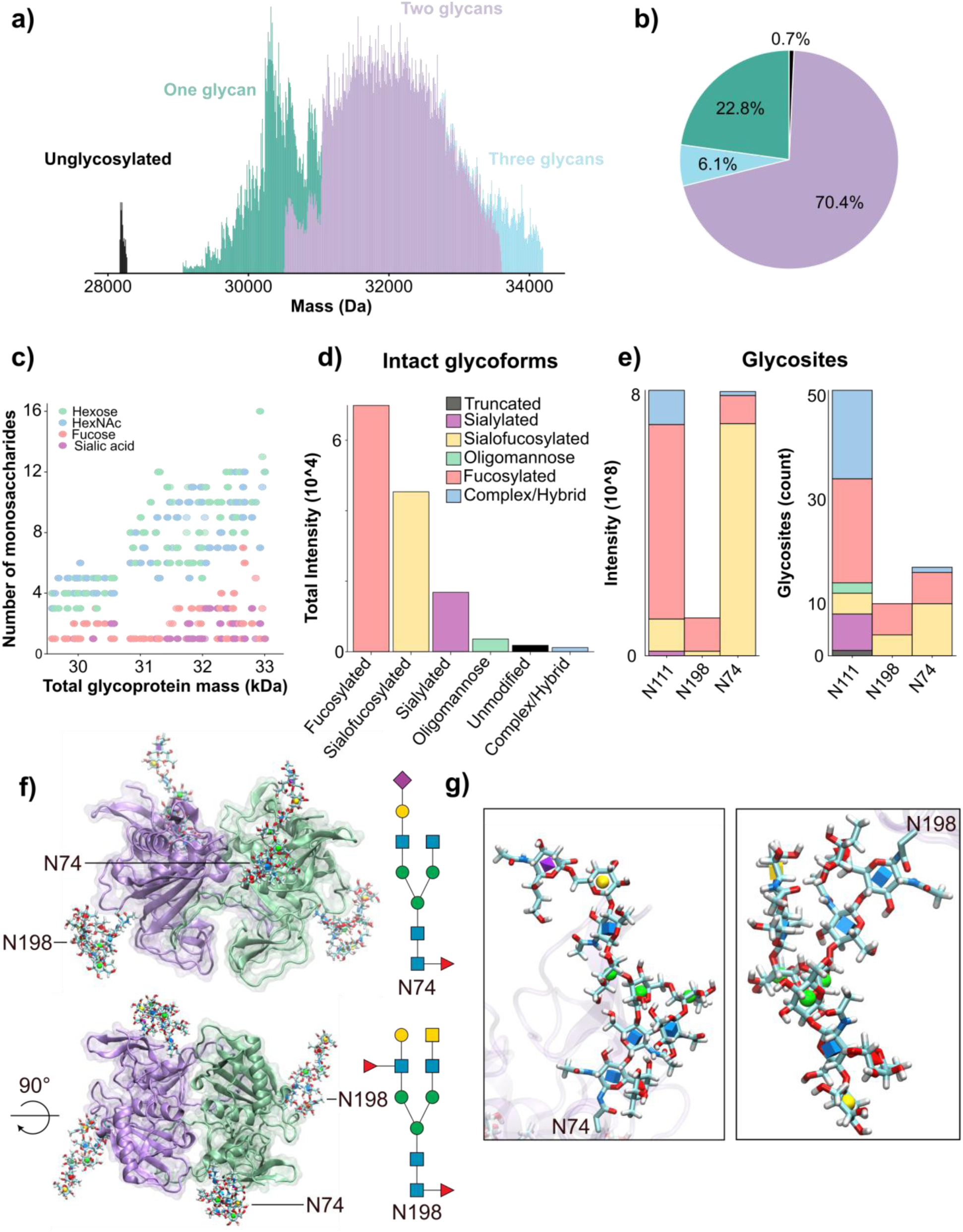
Further refining glycoproteoform information for EpCAM through an integrated glycoproteomics approach. (**a**) The bottom-up glycoproteomics data of EpCAM were used to statistically determine the percentage of each intact mass bin attributable to each macro - heterogeneity (i.e., the number of glycans). The resulting intact DMT mass distribution is visualized and color-coded per number of glycans. ( **b**) The repertoire of EpCAM glycoproteoforms from this recombinantly expressed population consists mostly of doubly glycosylated proteoforms. The pie chart is derived from the data in panel (a). (**c**) Glycan barcodes were generated using 0.5 Da bins as previously described.^26, 95^ In short, the glycoproteomics data were used to statistically infer the percentage of intact signal attributable to each possible intact glycoform. Each bin is plotted according to the total number of monosaccharides in the glycoform s, with alpha scaled by the percentage of signal attributed to the specific glycoform. (**d**) EpCAM’s intact glycoproteoforms were derived from the hRp-DIA-PTCR data, and the resulting glycan categories were visualized, showing a high degree of fucosylation. ( **e**) Site-specific glycosylation abundances are presented across EpCAM’s three glycosylation sites, derived from the bottom -up glycoproteomics data. ( **f**) A glycosylated proteoglycan of EpCAM visualized in 3D with glycans modeled using Re-Glyco from GlycoShape. N-glycans were present on both N 74 and N198, as supported by hRp-DIA-PTCR measurements. (**g**) A magnified view of both glycosites with the modeled conformations from Glyco Shape.

Taken together, our hRp-DIA-PTCR workflow and integration with bottom-up glycoproteomics and native MS DMT provide a direct avenue to further interrogating the biology of glycoproteoforms. To exemplify how insights from our workflow aid biological interpretation, we structurally modeled the identified glycoproteoform with a mass of 32.026 kDa corresponding to EpCAM with a sialofucosylated glycan at N74 (Hex(4)HexNAc(4)Fuc(1)Neu5Ac(1), GlyTouCan ID: G28959FS) and a doubly fucosylated glycan at N198 (Hex(4)HexNAc(5)Fuc(2), GlyTouCan ID: G27169KC). A previously solved crystal structure of EpCAM (PDB: 4MZV) was used as a template, however, all three native N-glycosylation sites had been mutated to remove glycosylation and reduce sample heterogeneity for crystallization.^68^ We therefore restored the glycosylation sites to their original asparagine residues and glycan structures identified by DIA-PTCR were modelled using Re-Glyco through the GlycoShape platform (**Figure 7f**).^90^ This model highlighted both N-glycans projecting away from the protein surface into the solvent. Specifically, N74 is positioned closest to the dimer interface and yet the glycan itself extends away from said interface, consistent with previous observations that N-glycosylation is not required for EpCAM dimerization. Glycosite N198 is situated within the C-terminal domain, proximal to the transmembrane region, where its solvent-exposed glycan may influence the orientation of EpCAM at the cell surface and its accessibility with neighboring membrane proteins.^91,92^ Close-up views of the modelled glycans revealed intra-glycan interactions which contort the flexible monosaccharide units for participation in hydrogen bonding and van der Waals interactions (**Figure 7g**). In both glycans, the core fucose residue is positioned to interact with either GlcNAc (blue square) or GalNAc (yellow square) residues, as well as being within hydrogen-bond distance from the protein backbone. Additionally, the N74 glycan contains a terminal sialic acid (NeuAc, purple diamond), which extends into the solvent accessible space. This solvent-exposed orientation is compatible with a potential role in glycan-mediated molecule recognition and is a known function of terminal sialylation in cell-cell communication, with reports linking EpCAM sialylated glycosylation states to tumour progression.^93,94^ Together, these findings highlight the importance of resolving intact glycoproteoforms as their complete glycan complement has the potential to influence protein structure and molecular function. By enabling the confident identification of native glycoproteoforms, our workflow provides a foundation for generating biologically relevant structural models that more accurately capture the functional consequences of protein glycosylation.

## CONCLUSIONS

Understanding the glycoproteoform landscape of any given glycoprotein has important implications for interpreting its biological function, but capturing these complex combinatorial modification states has remained too technically challenging for most glycoproteins. Here we present a strategy to directly measure intact glycoproteoform masses using gas-phase chemistry to simplify complex mass spectra. Our approach combines isolation of narrow m/z ranges, proton transfer charge reduction ion-ion reactions, and high resolving power Orbitrap mass analysis with open-source informatics solutions for processing isotopically-resolved glycoproteoform measurements. Using this hRp-DIA-PTCR approach, we characterized hundreds of intact glycoproteoform masses for five therapeutically relevant glycoproteins, TIGIT, CD40, EpCAM, PDL1, and CD24. Furthermore, we integrated this glycoproteoform profiling with bottom-up glycoproteomics and native MS DMT measurements to further refine interpretations of the glycoproteoform landscape of EpCAM. This work introduces several advances that now make glycoproteoform measurements an approachable addition to glycoproteoform characterization.

Our approach can be transferred to other related approaches on other instrument platforms, including those that rely on electron-based charge reduction (ECCR) rather than proton transfer charge reduction. While ECCR will introduce complications with hydrogen atom transfers and potentially unstable radical charge reduced species, our PTsliCR and IsoTrac informatics tools can still be applied to ECCR spectra. This informatics solution also works for native MS approaches for simple glycoproteins where PTCR may not be required or other measurements that generate isotopically resolved intact glycoprotein mass spectra. Our current iteration requires a priori knowledge of the glycoprotein being analyzed in order to properly account for protein and glycan mass contributions, but future work will look to integrate top-down fragmentation or other approaches to experimentally determine protein identity to extend this approach to discovery-based workflows. While hRp-DIA-PTCR significantly improves ambiguity in connecting measured intact glycoproteoforms to site-specific glycan microheterogeneity from bottom-up glycoproteomics data, future solutions to further reduce potential glycoproteoforms to be considered for any given intact mass include middle glycoproteomics, targeted use of proteases and glycosidases, and front-end separations either prior to or online with hRp-DIA-PTCR data acquisition. In all, this hRp-DIA-PTCR workflow represents a significant step toward decoding heterogeneous glycoprotein landscapes that will be critical for prescribing function to the glycoproteome.

## NOTES

D.B., R.D.M., C.M., J.D.H., and G.C.M. are employees of Thermo Fisher Scientific. N.M.R. receives support from Thermo Fisher Scientific under a collaborative research agreement and is a consultant for Augment Biologics.

## Supporting information

FilesS1_matchedPeaks

FileS2_cleanedSpectra

## SUPPORTING INFORMATION

Supporting Information is available free of charge online.

## ACKNOWLEDGEMENTS

Research reported in this publication was supported by the U.S. National Science Foundation under CAREER Award CHE-2540459 (N.M.R.) and by a Searle Scholar Fellowship from the Kinship Foundation (N.M.R.). The mass spectrometry raw data, fasta files, and search results have been deposited to the ProteomeXchange Consortium via the PRIDE partner repository^96^ with the dataset identifier PXD083184.

