## Supporting Material for "Defining glycoproteoform landscapes through an integrated glycoproteomics approach enabled by high-resolving power proton transfer charge reduction tandem mass spectrometry"

The mass spectrometry raw data, fasta files, and search results have been deposited to the ProteomeXchange Consortium via the PRIDE partner repository with the dataset identifier PXD083184.

**Table of Contents**

**Supplemental Files**
These will be the PDFs with the isotope pattern matches.

**Supplemental Figures**
**Figure S1**. High-resolution intact EpCAM spectrum suffers from high peak density.
**Figure S2**. Workflow used in this manuscript.
**Figure S3**. Cleaning PTCR spectra using predicted charge-reduced m/z ranges.
**Figure S4**. Isotope-guided deconvolution of Ubiquitin.
**Figure S5**: UniDec deconvolution of TIGIT.
**Figure S6**. Isotope-guided deconvolution of two DIA-PTCR cycles of TIGIT shows high similarity.
**Figure S7**. The importance of ion statistics for optimal sequencing depth.
**Figure S8**. Exploring the effects of DIA-PTCR window offsets.
**Figure S9**. MS1 spectra of TIGIT, CD40, PDL1, and CD24.
**Figure S10**. Comparison of intact mass profiles obtained from DIA-PTCR and DMT analyses.
**Figure S11**. Glycan-specific delta masses in isotope-guided deconvoluted DIA-PTCR spectra.
**Figure S12**. Number of theoretical intact glycoforms based on bottom-up glycoproteomics data.

**SUPPLEMENTAL FIGURES**

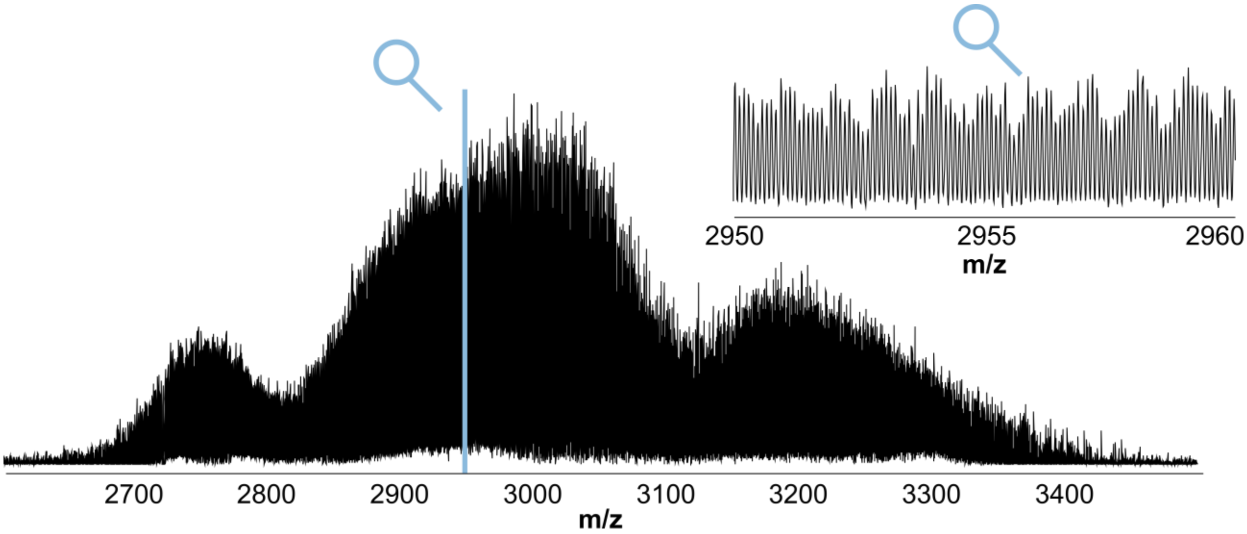


**Figure S1. High-resolution intact EpCAM spectrum suffers from high peak density.** Intact spectra were acquired at 240k resolution. Three charge clusters are clearly visible, yet a zoom-in of the spectra reveals high peak density that hinders spectral interpretation.

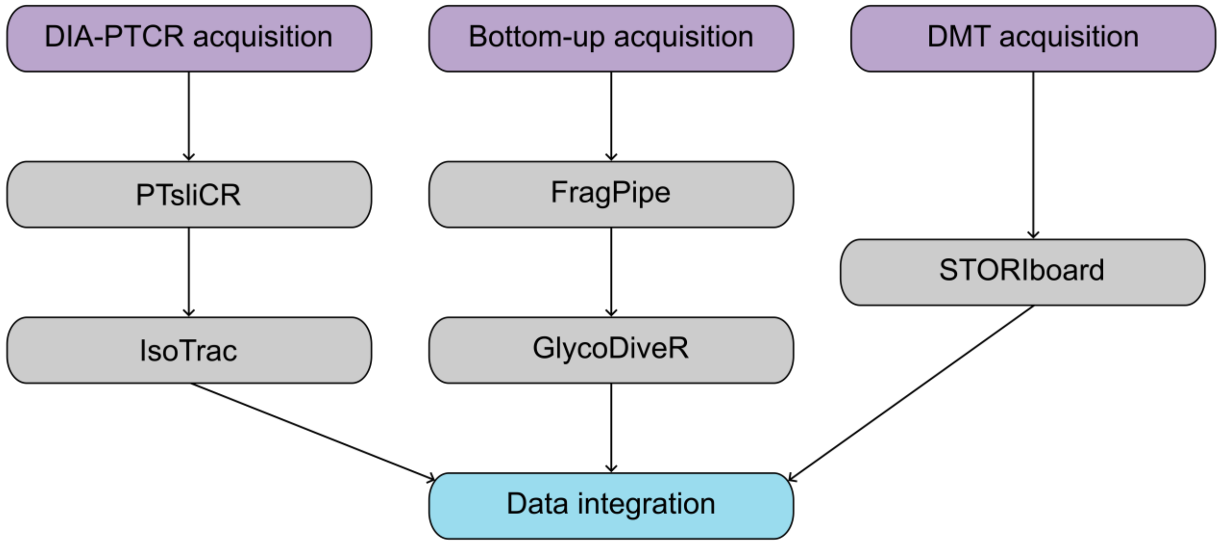


**Figure S2. Workflow used in this manuscript.** Each acquisition type was first analyzed using the appropriate software tools, and the generated outputs were used for the integrated comparison. Importantly, no manual data preparation was required for any step, as each tool was directly compatible with the untouched generated input. IsoTrac, GlycoDiveR, and STORIboard output detailed CSV files allowing customizable downstream analysis.

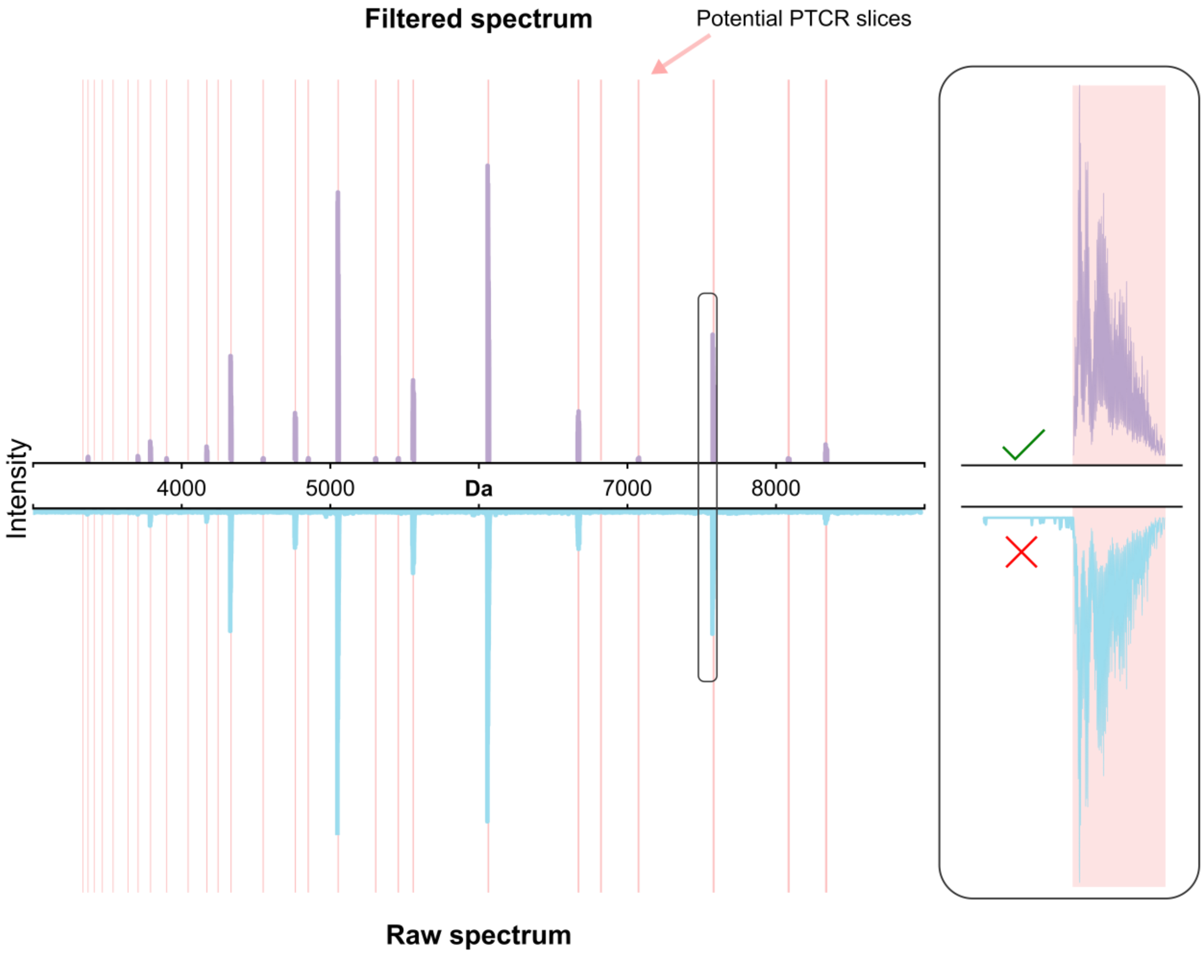


**Figure S3. Cleaning PTCR spectra using predicted charge-reduced m/z ranges**. Only a limited number of mass ranges in PTCR scans potentially hold spectral information. These ranges were determined using the isolation window and the expected precursor charge states (shown in pink). All peaks outside these slices are non-informational and can interfere with charge deconvolution of the spectra. Therefore, only spectral information within the potential PTCR is retained and used for deconvolution.

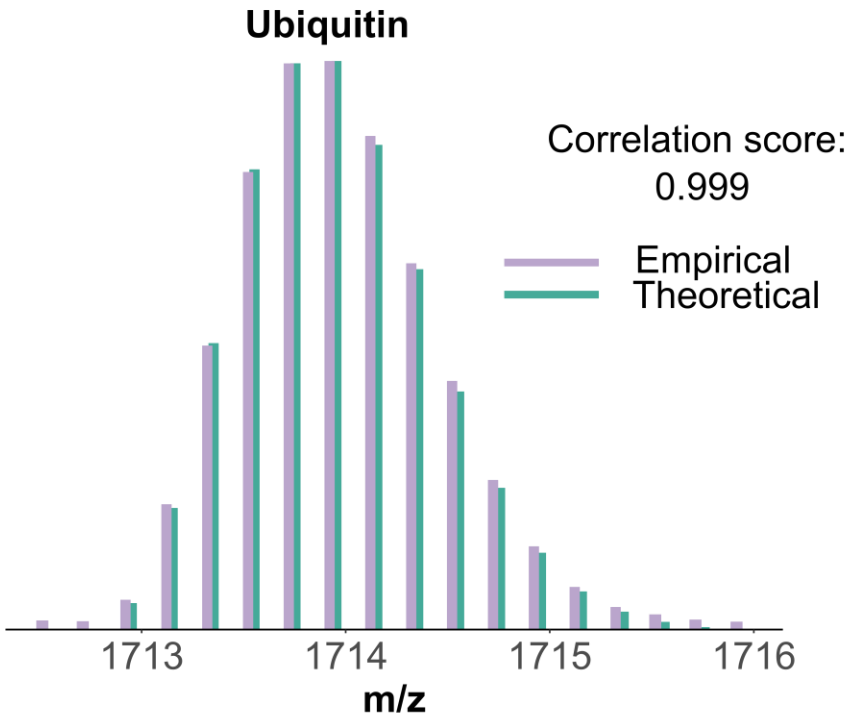


**Figure S4: Isotope-guided deconvolution of Ubiquitin.** To verify the accuracy of the isotope calculation, we overlaid the theoretically determined isotope envelope onto an empirically measured unglycosylated Ubiquitin. The overlay showed a high similarity with a Pearson correlation of 0.999. The theoretical envelope was slightly shifted to the right for better visualization.

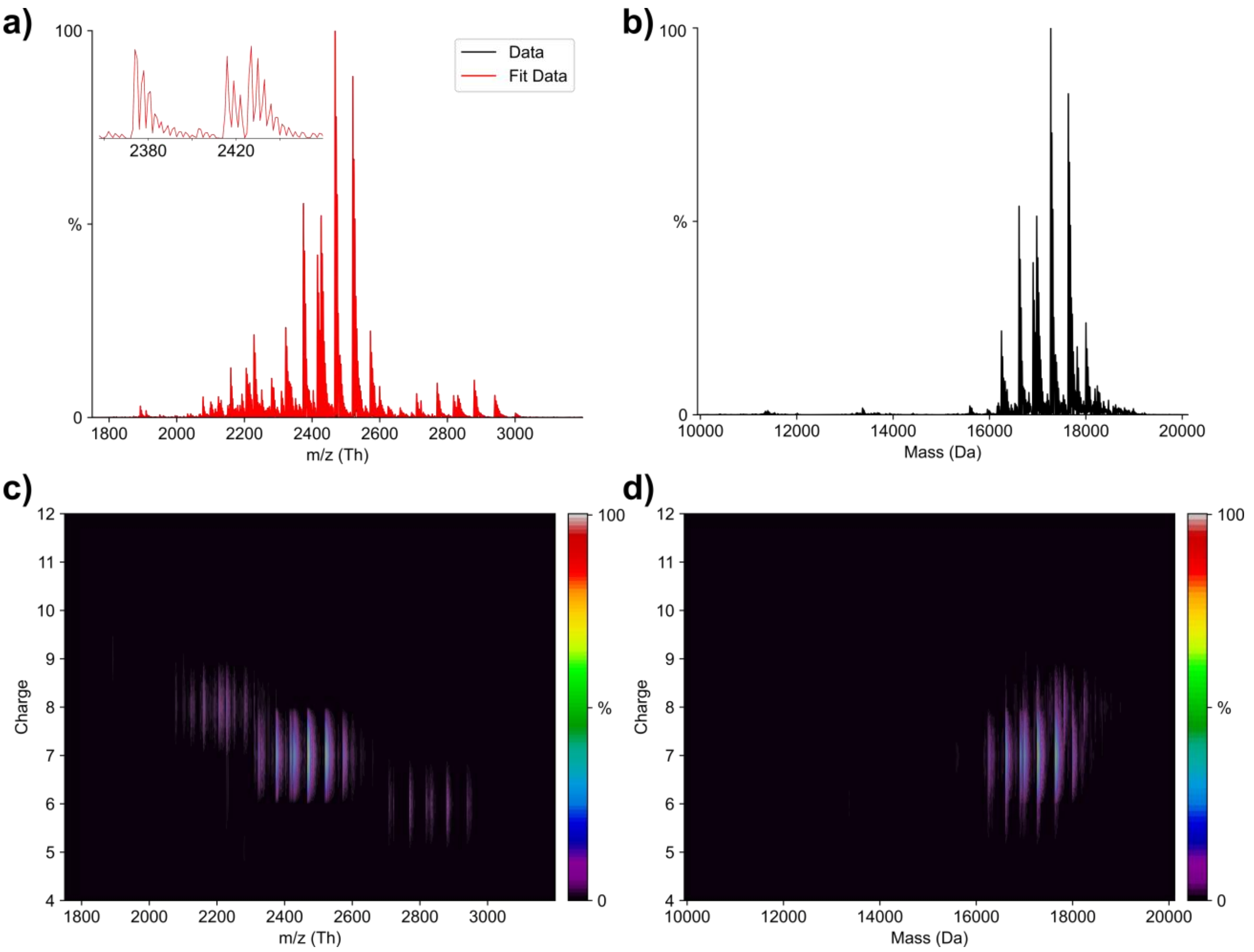


**Figure S5: UniDec deconvolution of TIGIT.** (**a**) Intact MS1 spectra of TIGIT were acquired and deconvoluted using UniDec. The empirical spectrum (black) is overlaid with the fitted data (red). (**b**) The charge deconvoluted spectrum of TIGIT. (**c**) The charge-to-m/z map used for deconvolution. (**d**) The mass-to-m/z map after deconvolution.

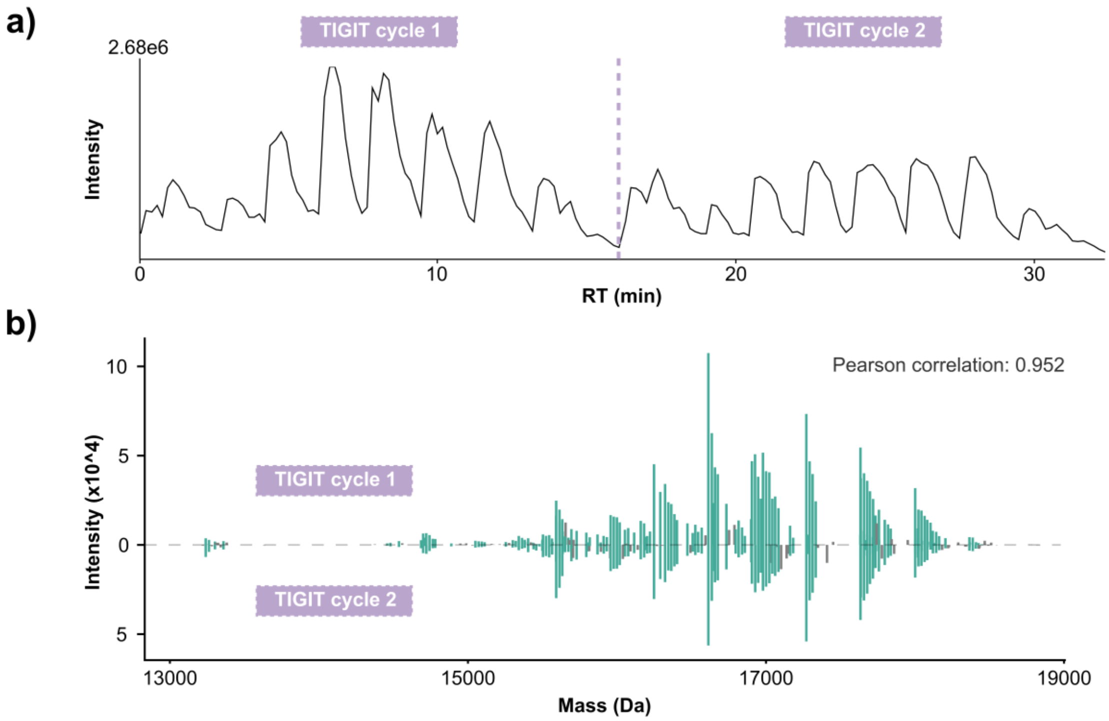


**Figure S6. Isotope-guided deconvolution of two DIA-PTCR cycles of TIGIT shows high similarity**. Two DIA-PTCR cycles were acquired over the m/z 2200-2600 mass range in a single acquisition. The TIC shows a similar pattern (**a**), and the Pearson correlation of the isotope-guided deconvolved spectra revealed a high correlation (**b**).

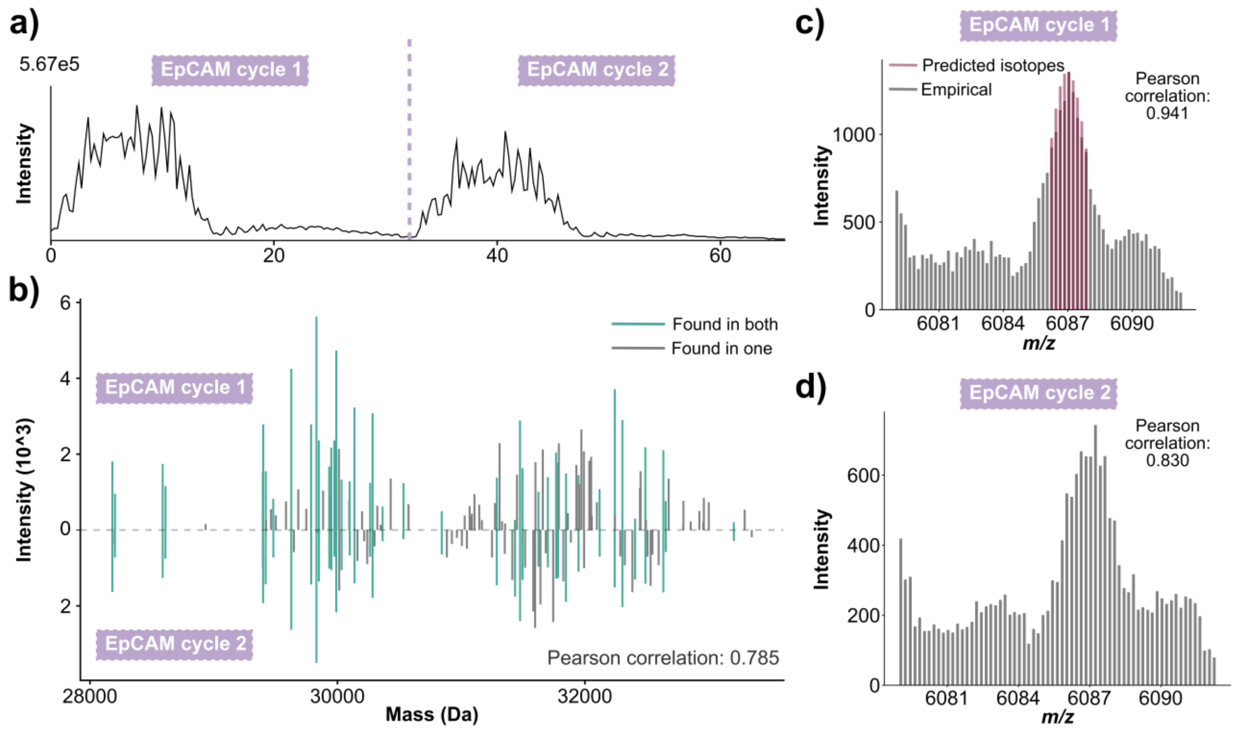


**Figure S7. The importance of ion statistics for optimal sequencing depth.** (**a**) Two loops (m/z 2800-3350) of DIA-PTCR scans were acquired of EpCAM, with the TIC revealing a slightly lower intensity in the second pass than the first pass. (**b**) The isotope-guided deconvoluted spectra showed decent agreement and Pearson correlation; however, the correlation was lower than the two DIA-PTCR loops of TIGIT. (**c/d**) Isotope pattern overlays of the empirical data (grey) and the theoretical envelope (red) revealed that this result reflects quantitative variability in the empirical isotope patterns due to insufficient ion statistics. Sufficient ion signal allows for quantitatively accurate isotope patterns that are matched with the empirical isotope envelope (panel c), while the same PTCR slice with a lower ion signal shows too much quantitative variability for the theoretical isotope patterns to be matched (panel d).

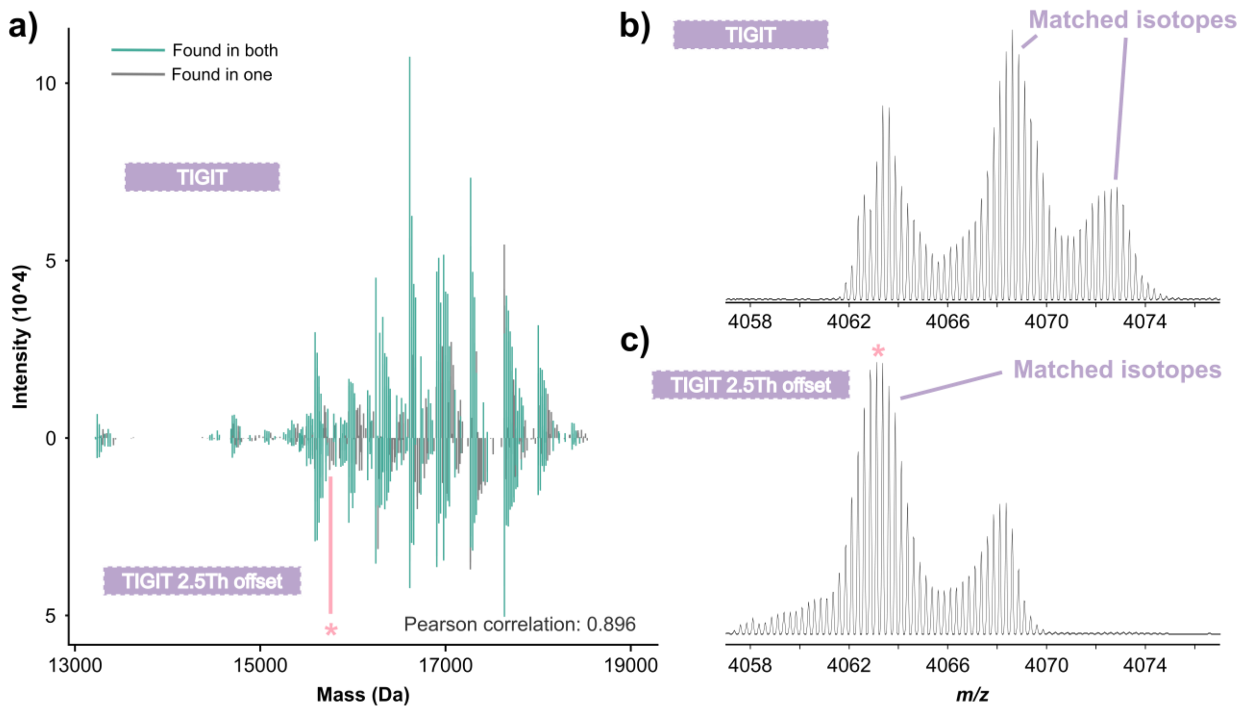


**Figure S8. Exploring the effects of DIA-PTCR window offsets.** TIGIT was measured using DIA-PTCR with 5 Th windows, with two separate acquisitions (m/z 2200-2600 and m/z 2202.5-2602.5), to assess whether the edges of the isolation window could generate isotope-like peak shapes that lead to false hits. (**a**) The two acquisitions showed a high correlation, with only a small portion of peaks identified in one acquisition but not the other. (**b/c**) This isn't the result of artificial isotope-like peak shapes generated at the edges of the extraction window, but by genuine signals present between adjacent DIA windows.

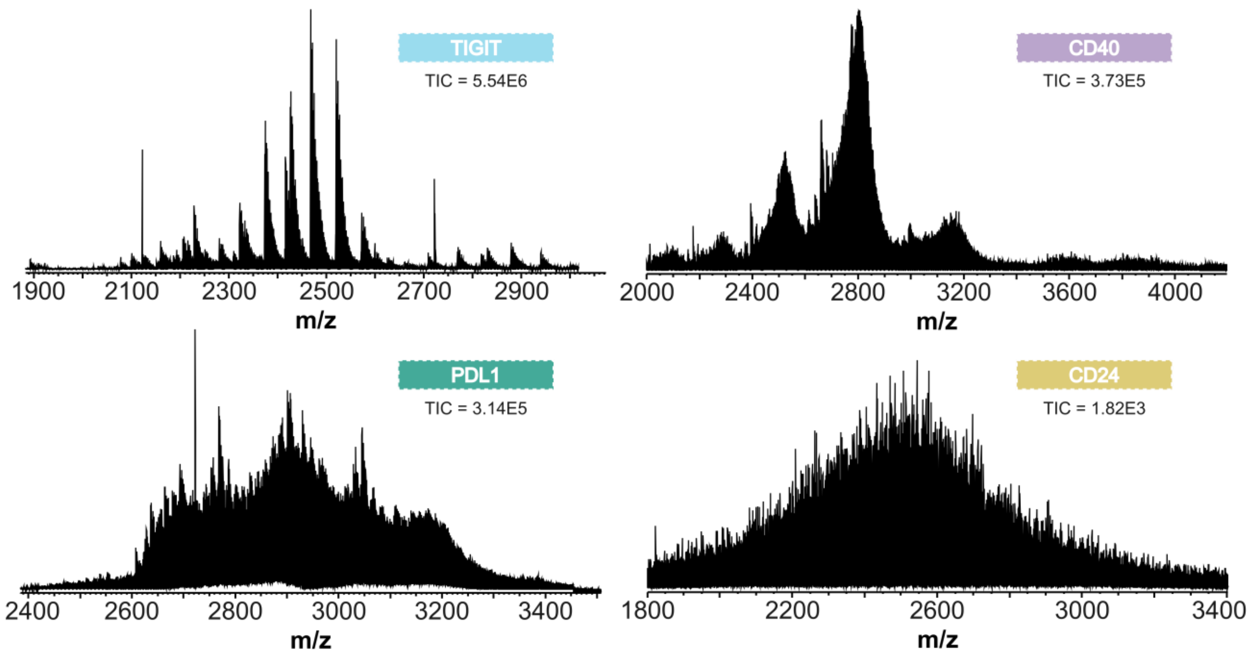


**Figure S9. MS1 spectra of TIGIT, CD40, PDL1, and CD24.** MS1 spectra were acquired at 240k resolution for TIGIT, CD40, PDL1, and CD24, sorted from low to high glycan heterogeneity. The increasing complexity is reflected by greater spectral density and broader, more Gaussian-like spectral features.

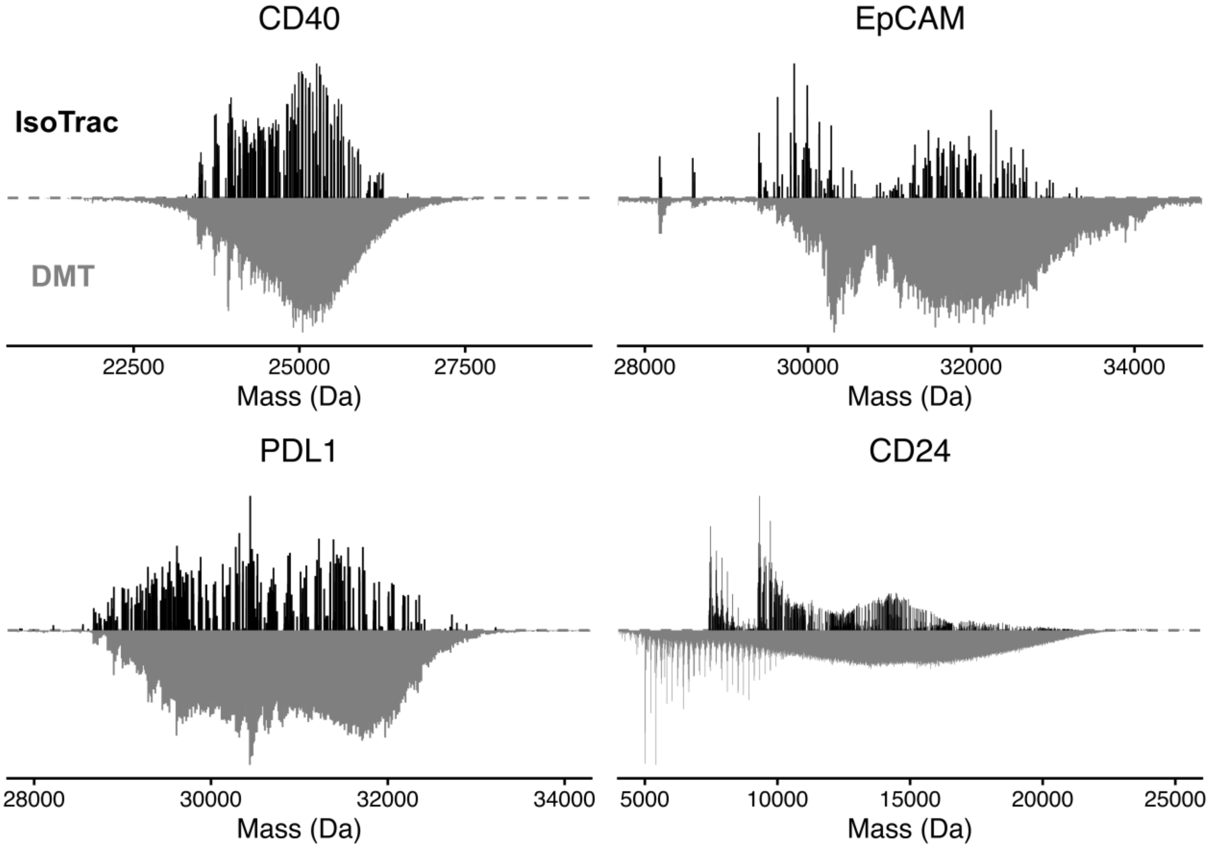


**Figure S10: Comparison of intact mass profiles obtained from DIA-PTCR and DMT analyses.** Intact masses were determined using DIA-PTCR and DMT, and visualized in a mirror plot (top: DIA-PTCR; bottom: DMT). The two intact mass profiles show strong agreement.

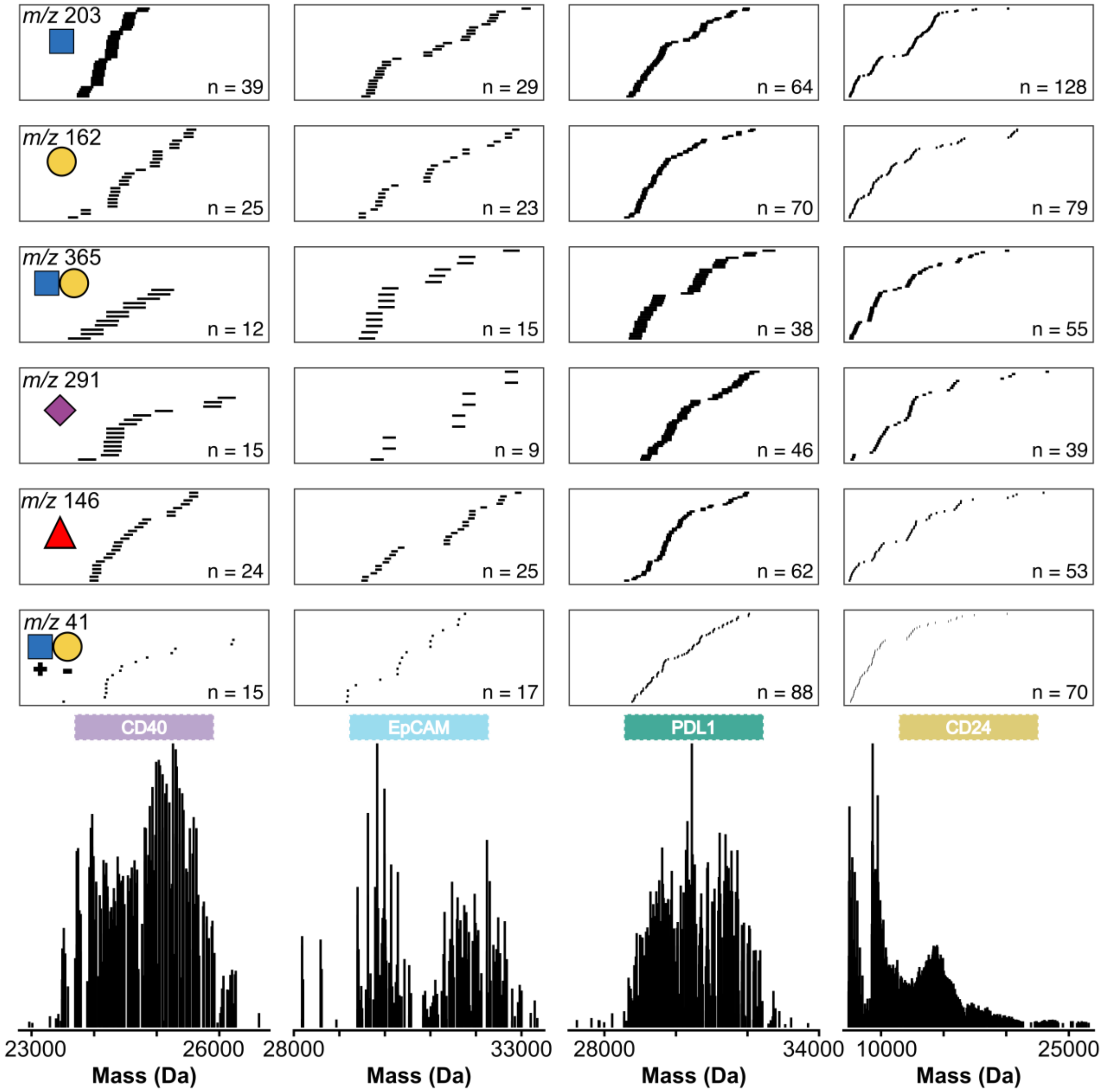


**Figure S11. Glycan-specific delta masses in isotope-guided deconvoluted DIA-PTCR spectra.** The heterogeneity of glycoproteins results in characteristic mass differences among intact glycoproteins. These characteristic mass differences were extracted using a 20 ppm tolerance and visualized above the intact chromatograms. The exact delta masses included were 203.19252, 162.1406, 365.33312, 291.25458, 146.1412, and 41.05192.

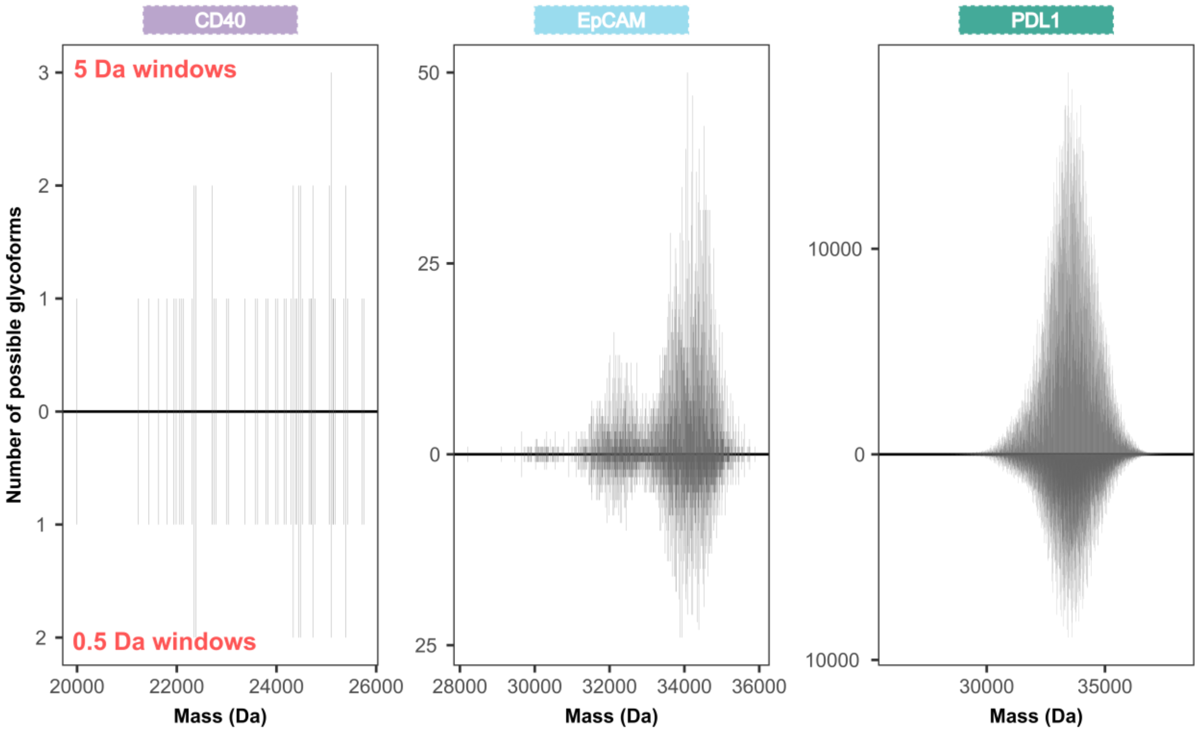


**Figure S12. Number of theoretical intact glycoforms based on bottom-up glycoproteomics data.** The theoretical number of glycoforms was calculated in 5 Da (top) and 0.5 Da (bottom) bins using the glycoproteomics data. Quantitative glycoproteomics data are a central data layer for inferring intact glycoforms, yet they often suffer from insufficient sequencing depth, as exemplified by CD40, or from many potential glycoforms matching each mass, as exemplified by PDL1.
