## Supplementary material for "Defining glycoproteoform landscapes through an integrated glycoproteomics approach enabled by high-resolving power proton transfer charge reduction tandem mass spectrometry": FilesS1_matchedPeaks

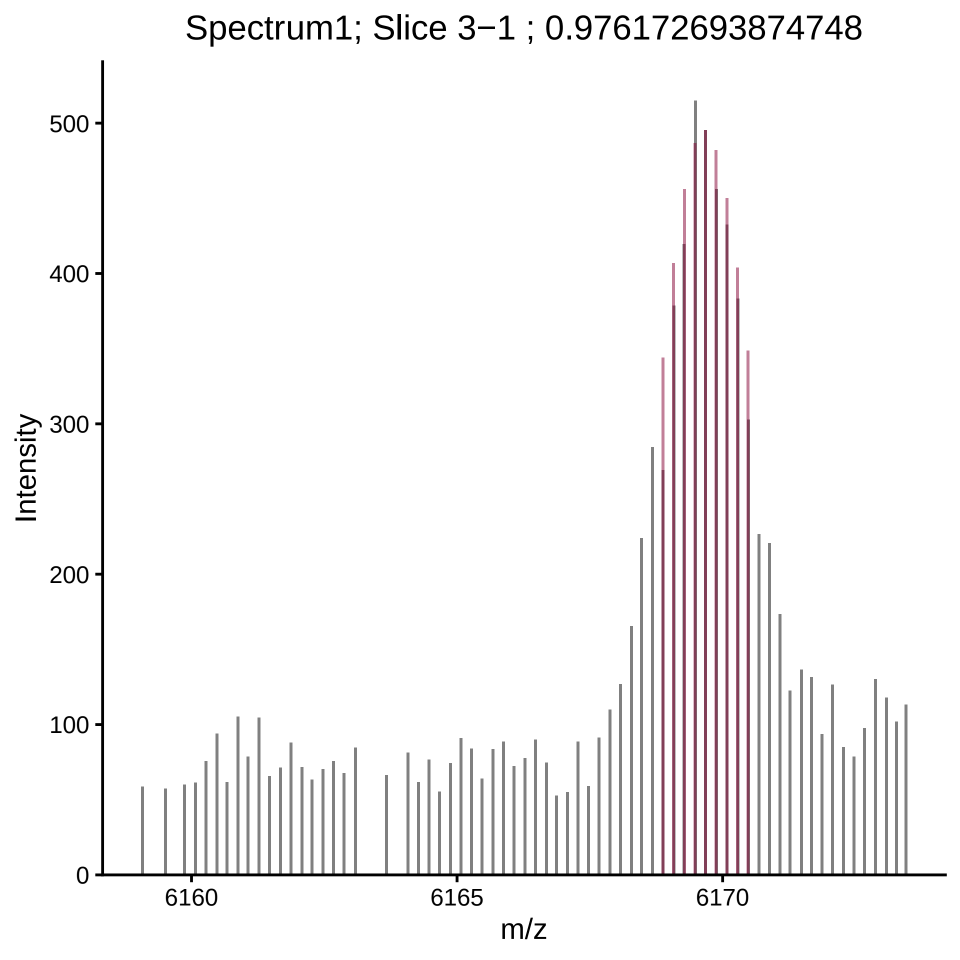

Spectrum1; Slice 3−1 ; 0.976172693874748
Intensity
500
400
300
200
100
0
6160
6165
6170
m/z

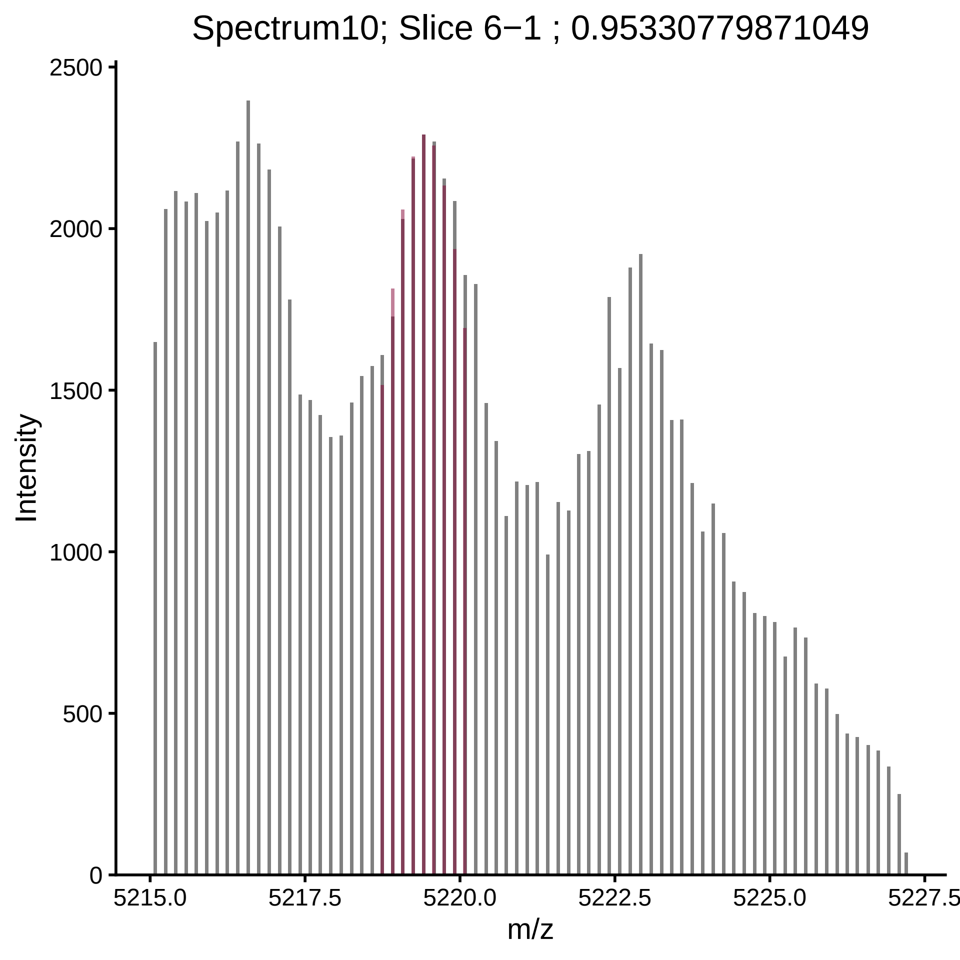

Spectrum10; Slice 6−1 ; 0.95330779871049
Intensity
2500
2000
1500
1000
500
0
5215.0
5217.5
5220.0
5222.5
5225.0
5227.5
m/z

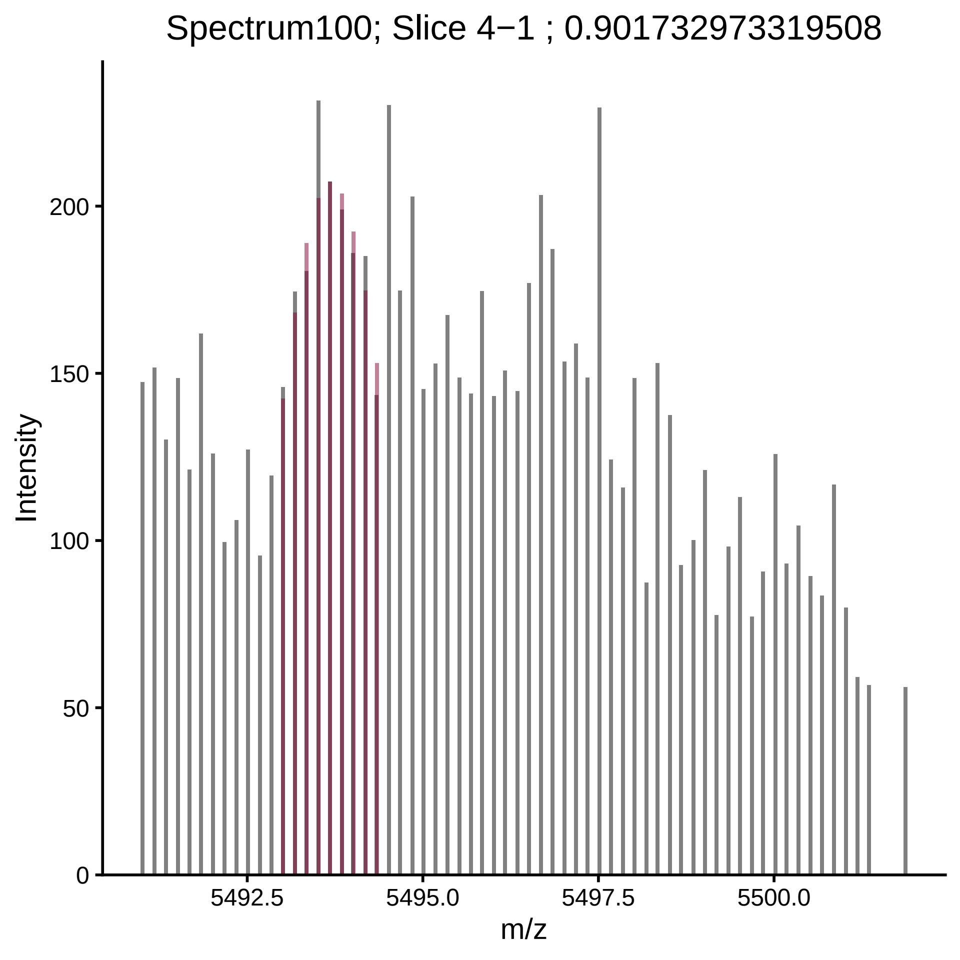

Spectrum100; Slice 4−1 ; 0.901732973319508
Intensity
200
150
100
50
0
5492.5
5495.0
5497.5
5500.0
m/z

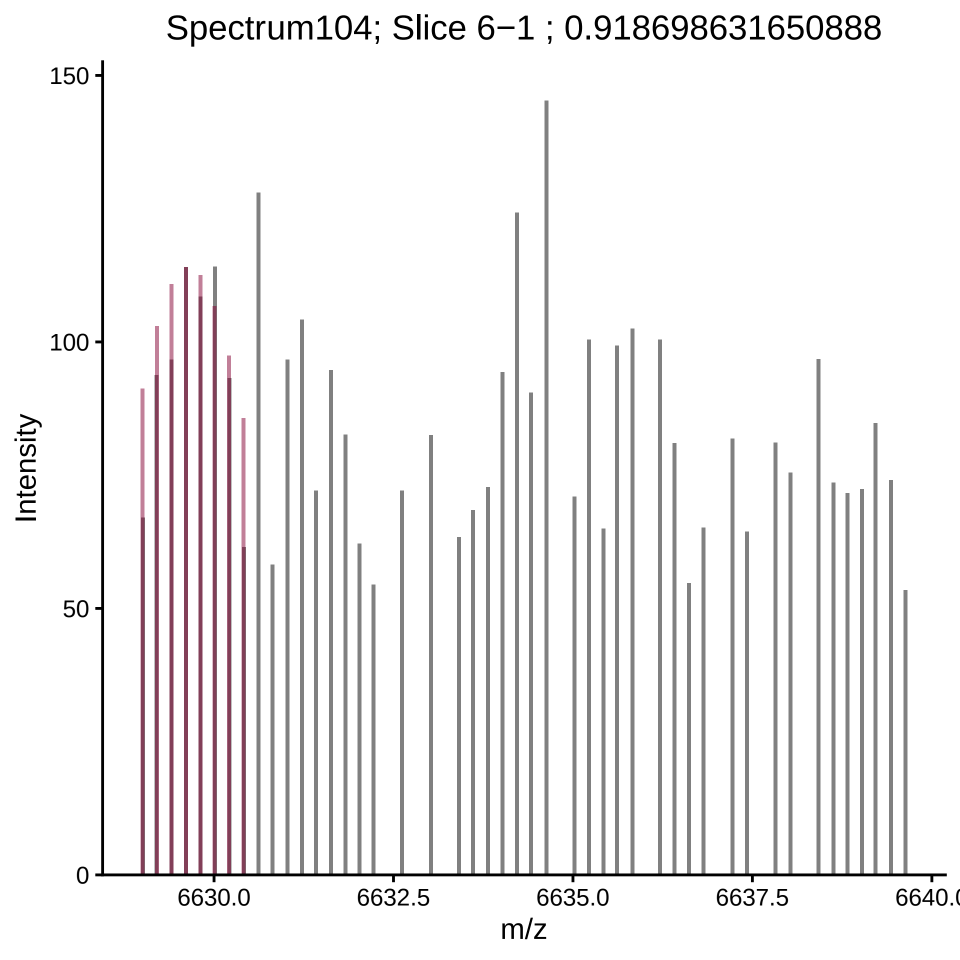
Spectrum104; Slice 6−1 ; 0.918698631650888
Intensity
150
100
50
0
6630.0
6632.5
6635.0
6637.5
6640.0
m/z

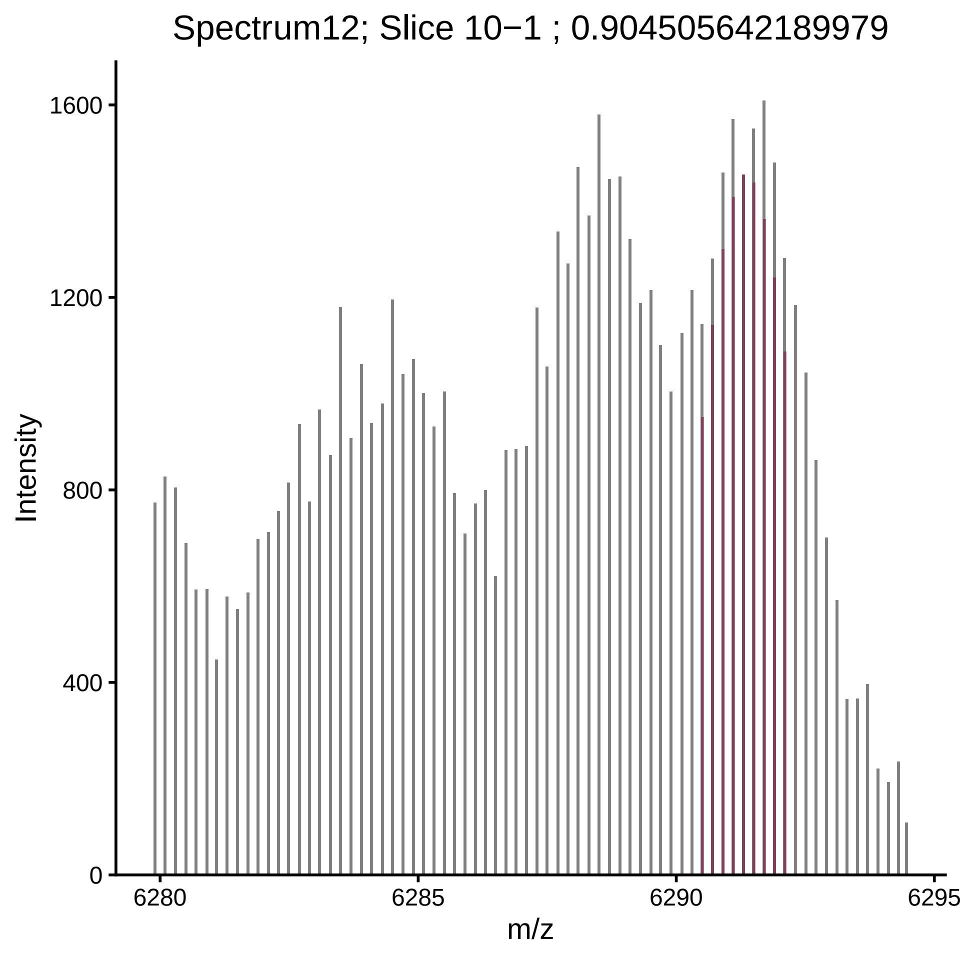

Spectrum12; Slice 10−1 ; 0.904505642189979
Intensity
1600
1200
800
400
0
6280
6285
6290
6295
m/z

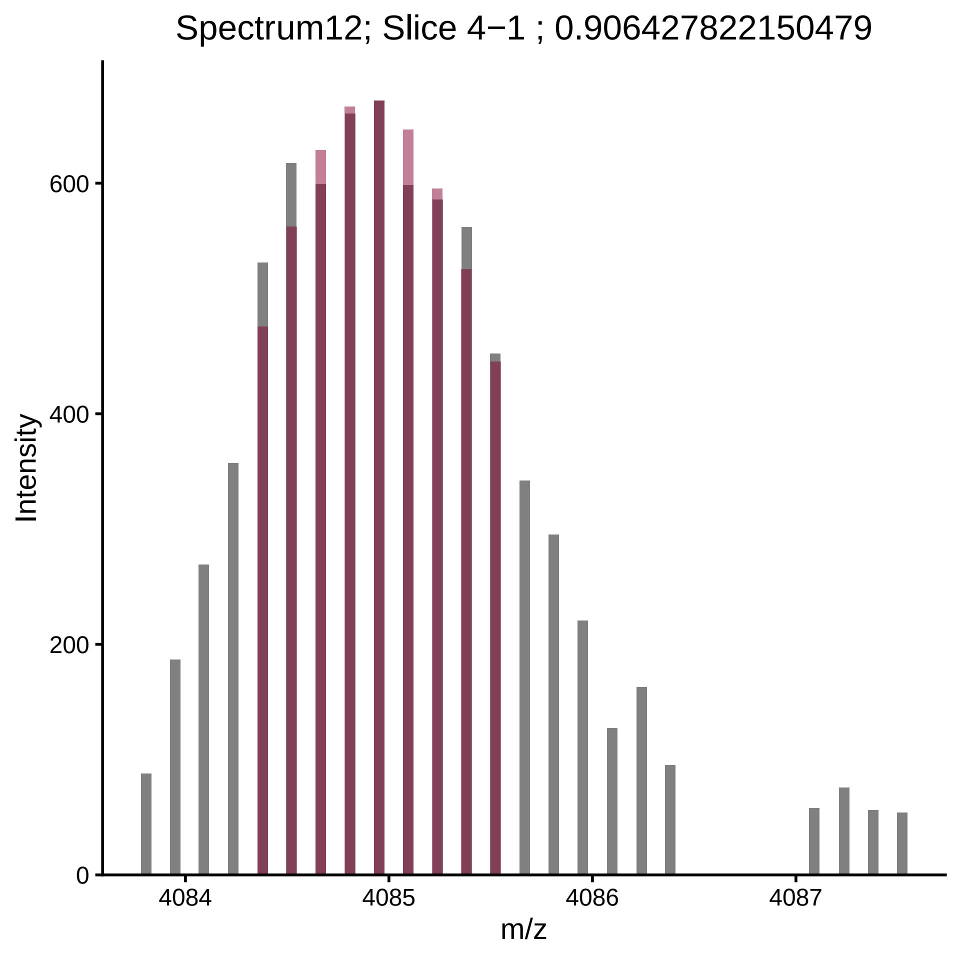
Spectrum12; Slice 4−1 ; 0.906427822150479
Intensity
m/z
0
200
400
600
4084
4085
4086
4087

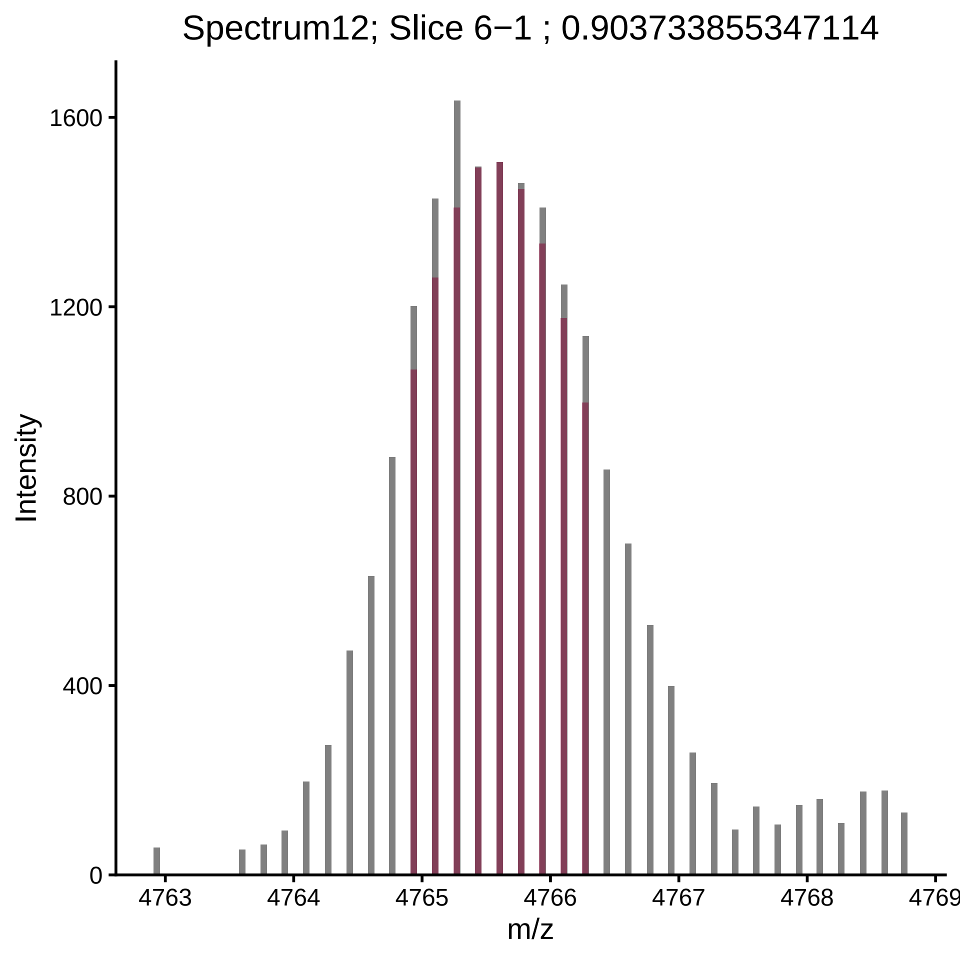

Spectrum12; Slice 6−1 ; 0.903733855347114
Intensity
1600
1200
800
400
0
4763
4764
4765
4766
4767
4768
4769
m/z

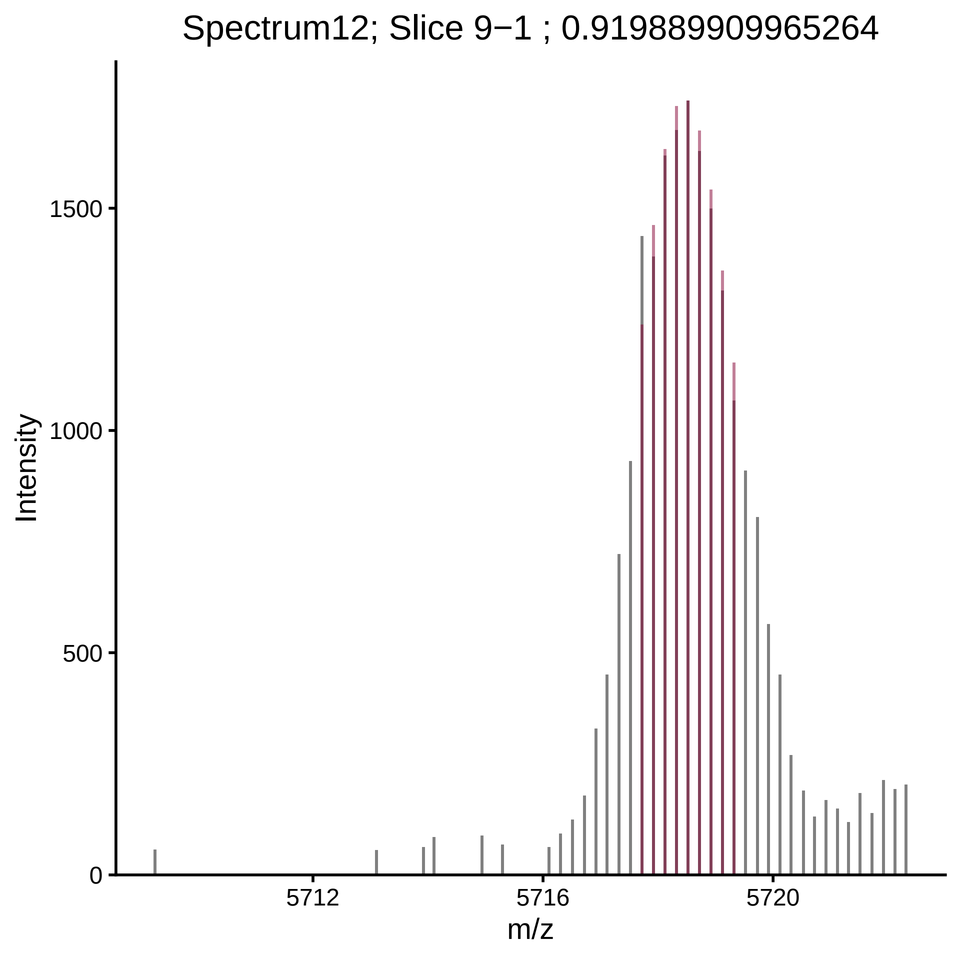

Spectrum12; Slice 9−1 ; 0.919889909965264
Intensity
0
500
1000
1500
5712
5716
5720
m/z

Spectrum121; Slice 3−1 ; 0.95054644317926

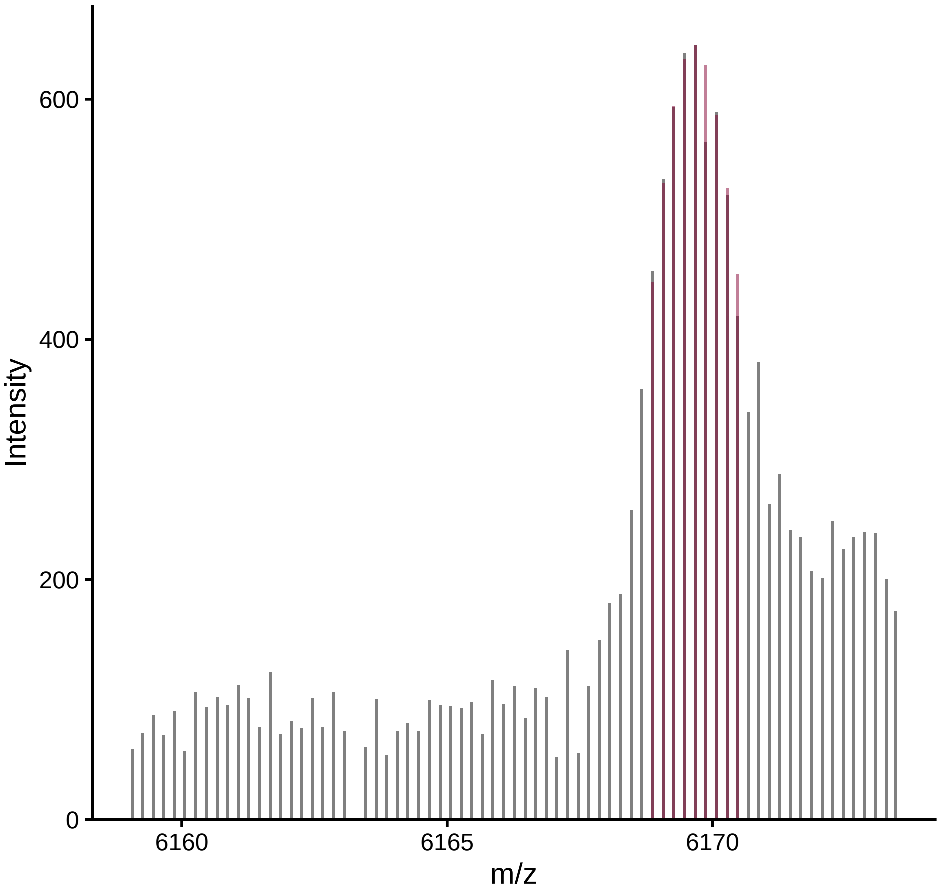

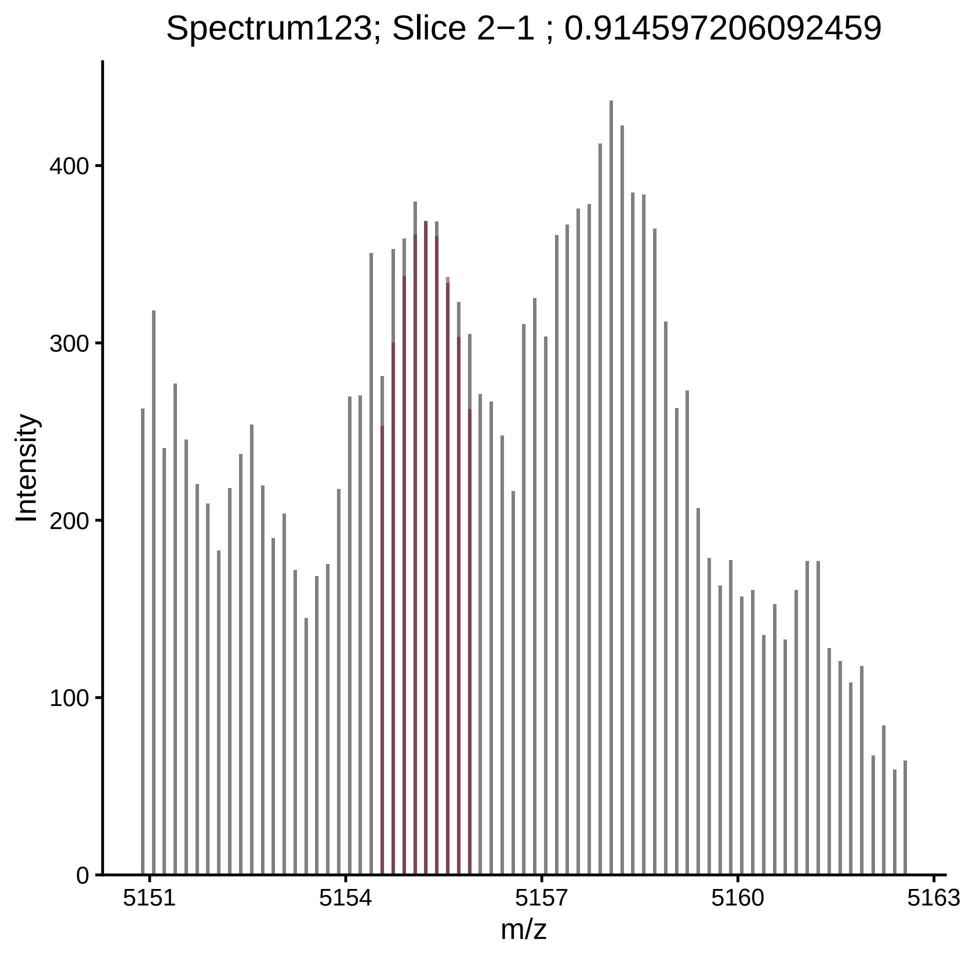
Spectrum123; Slice 2−1 ; 0.914597206092459
Intensity
400
300
200
100
0
5151
5154
5157
5160
5163
m/z

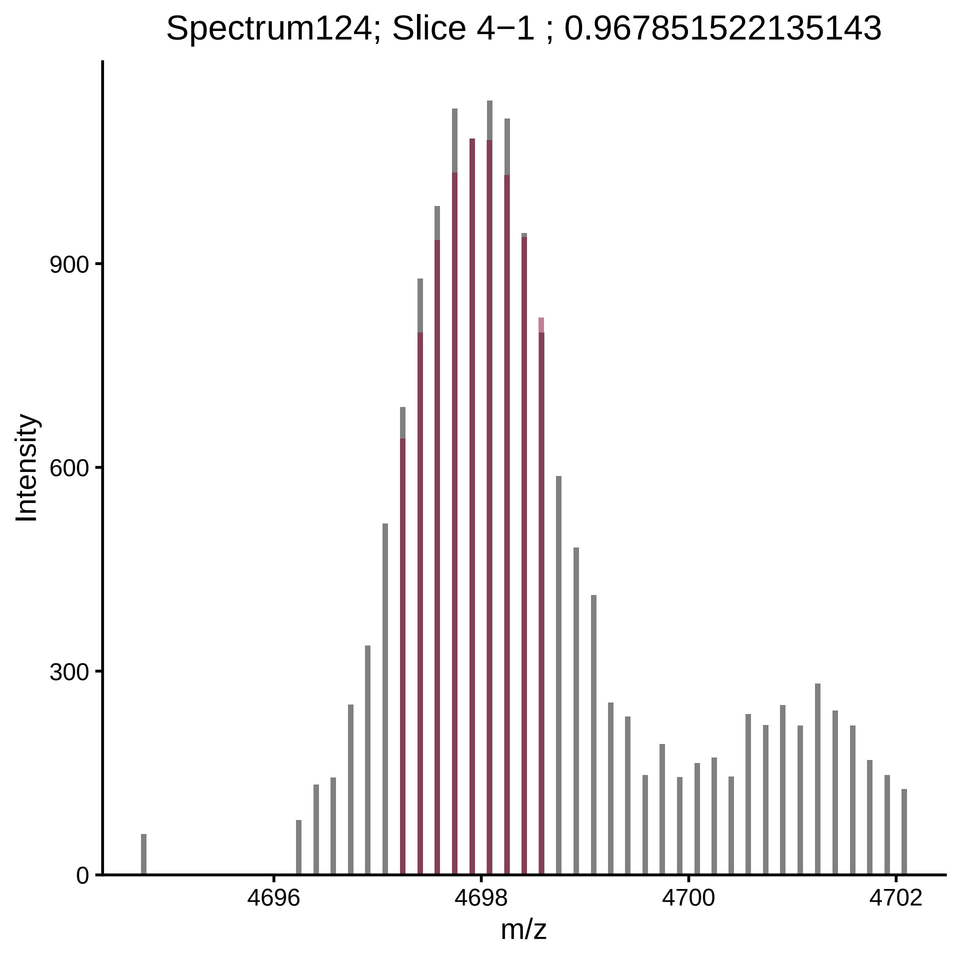

Spectrum124; Slice 4−1 ; 0.967851522135143
Intensity
900
600
300
0
4696
4698
4700
4702
m/z

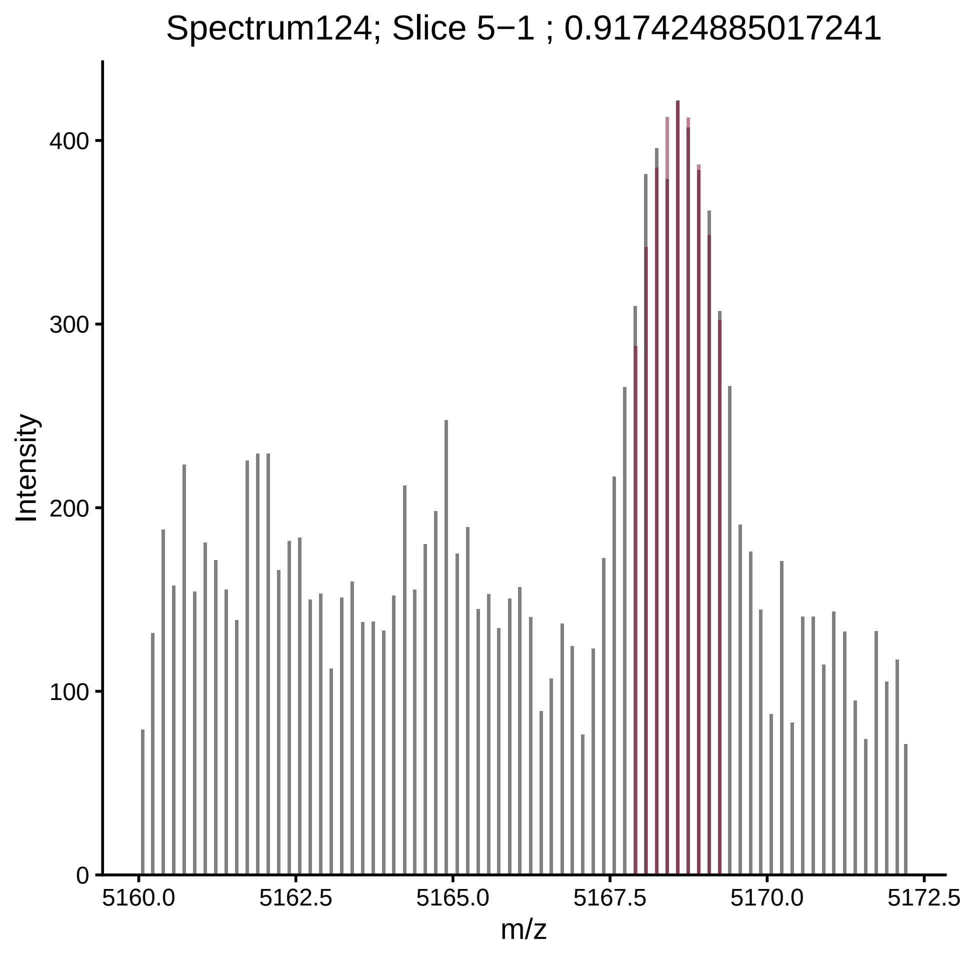
Spectrum124; Slice 5−1 ; 0.917424885017241
Intensity
m/z
0
100
200
300
400
5160.0
5162.5
5165.0
5167.5
5170.0
5172.5

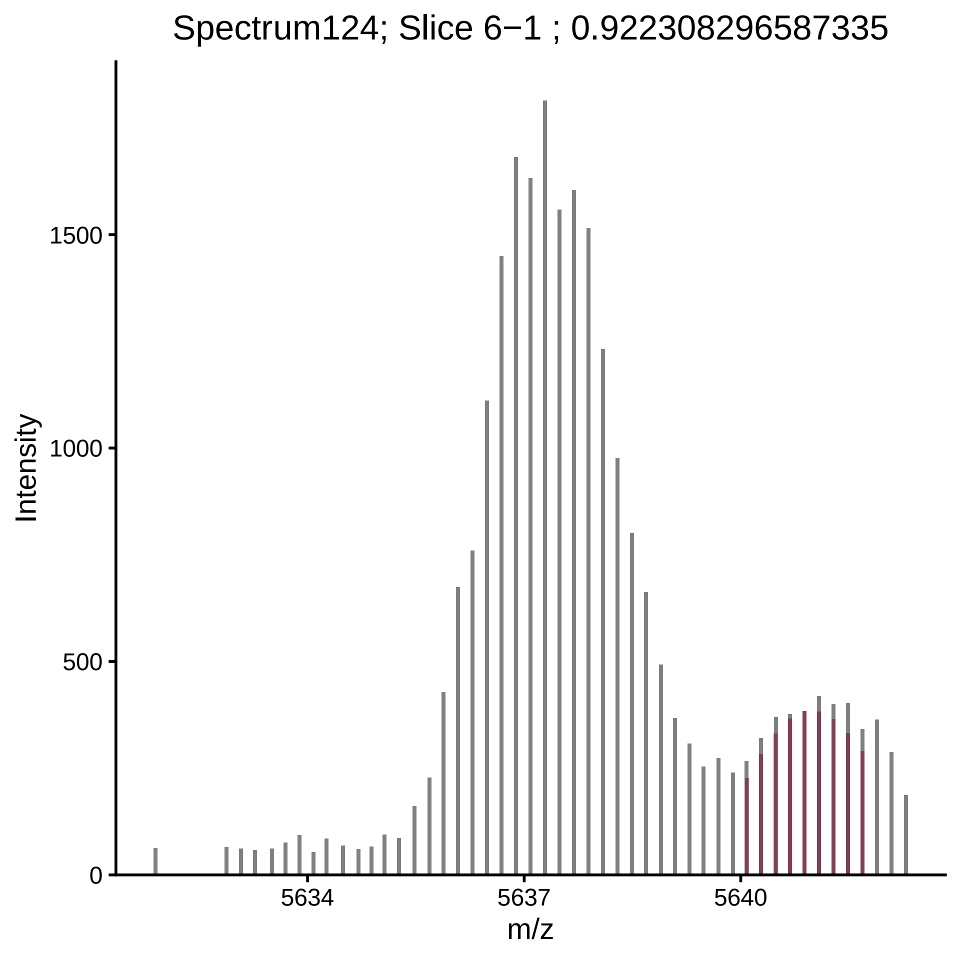

Spectrum124; Slice 6−1 ; 0.922308296587335
Intensity
0
500
1000
1500
5634
5637
5640
m/z

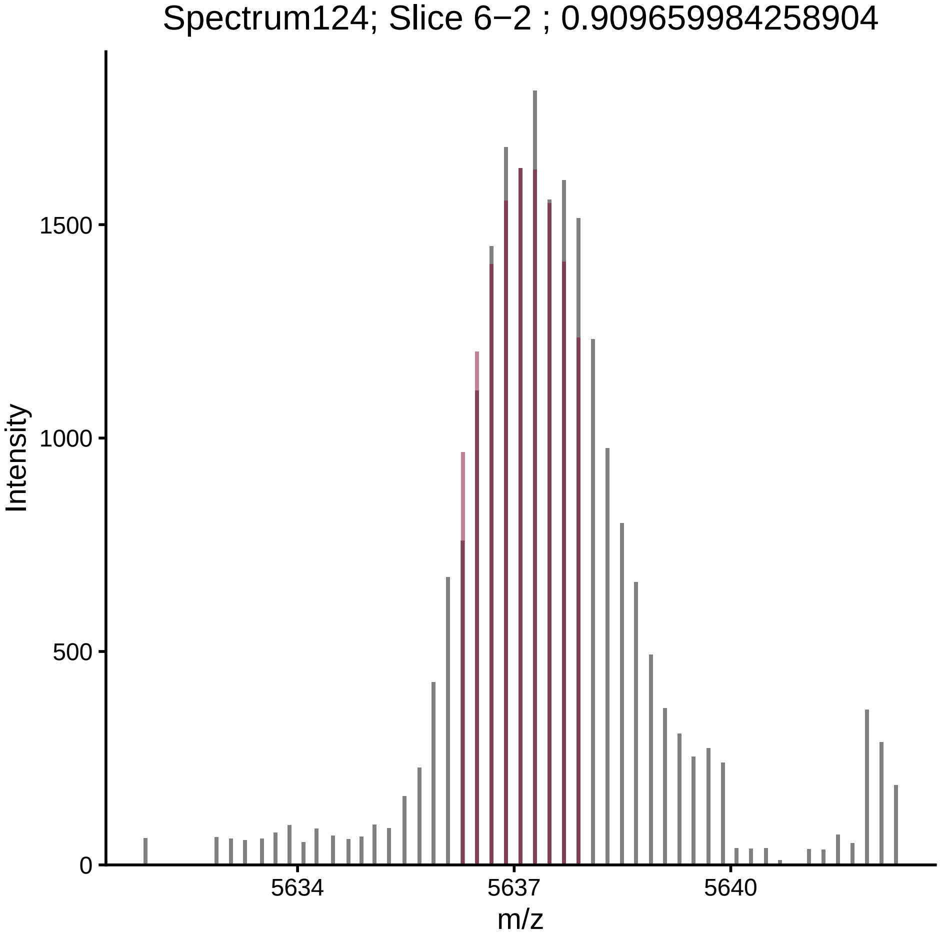
Spectrum124; Slice 6−2 ; 0.909659984258904
Intensity
1500
1000
500
0
5634
5637
5640
m/z

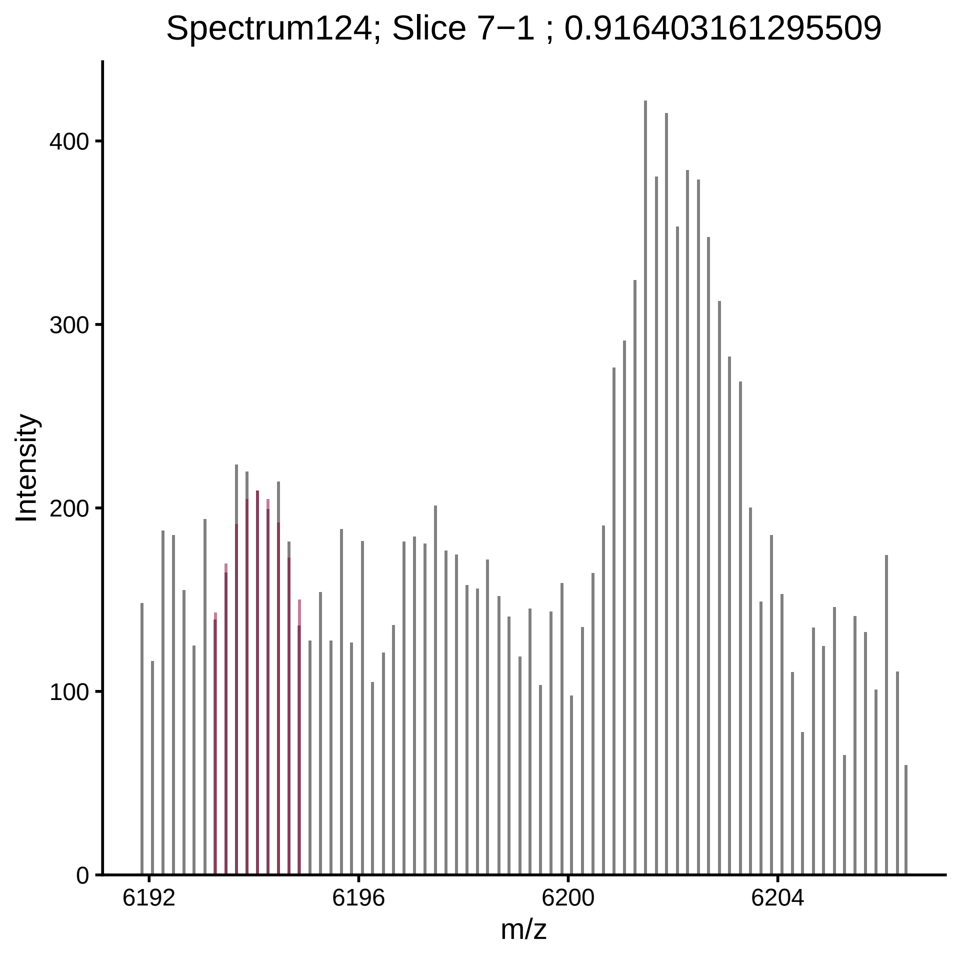

Spectrum124; Slice 7−1 ; 0.916403161295509
Intensity
0
100
200
300
400
6192
6196
6200
6204
m/z

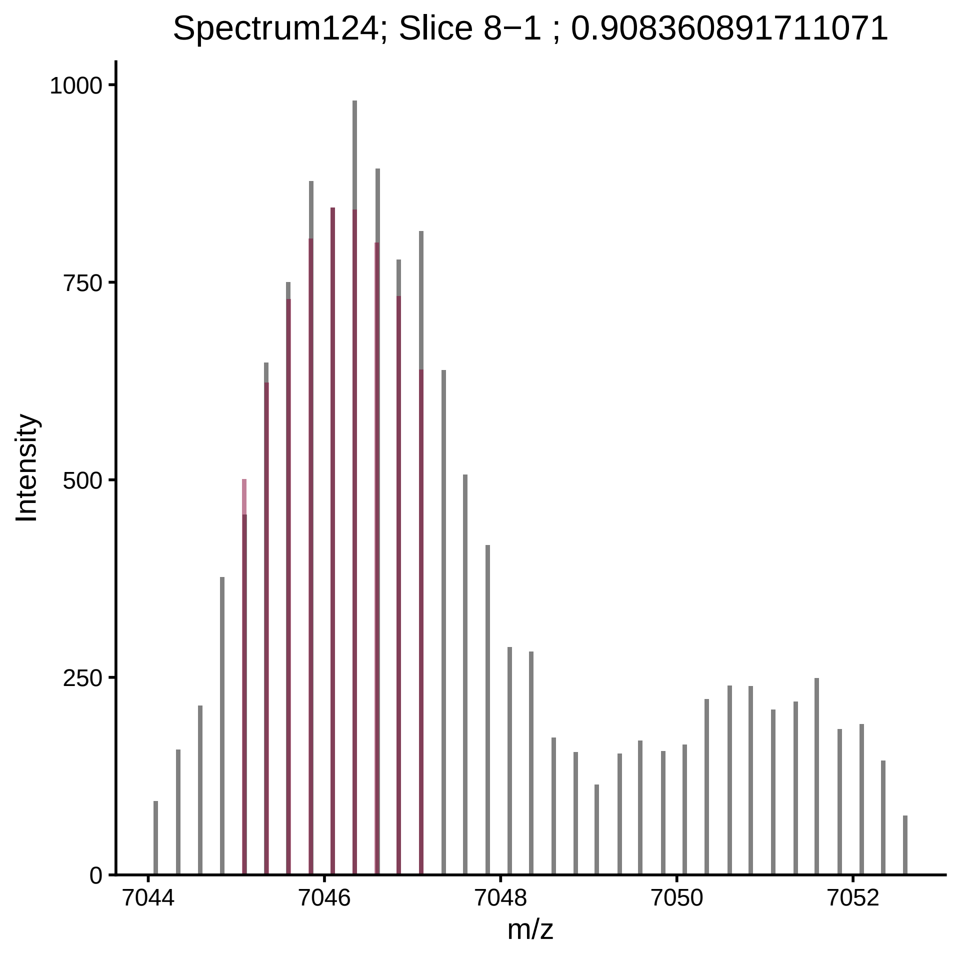

Spectrum124; Slice 8−1 ; 0.908360891711071
Intensity
1000
750
500
250
0
7044
7046
7048
7050
7052
m/z

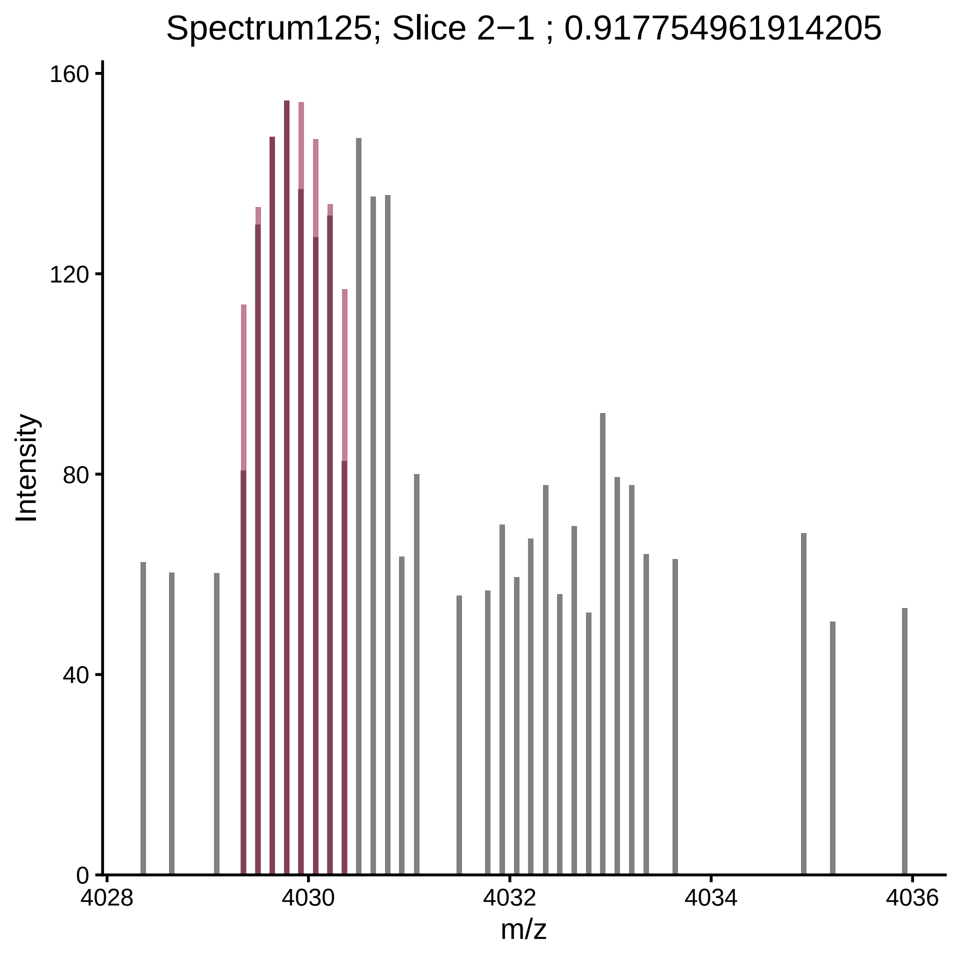

Spectrum125; Slice 2−1 ; 0.917754961914205
Intensity
0
40
80
120
160
m/z
4028
4030
4032
4034
4036

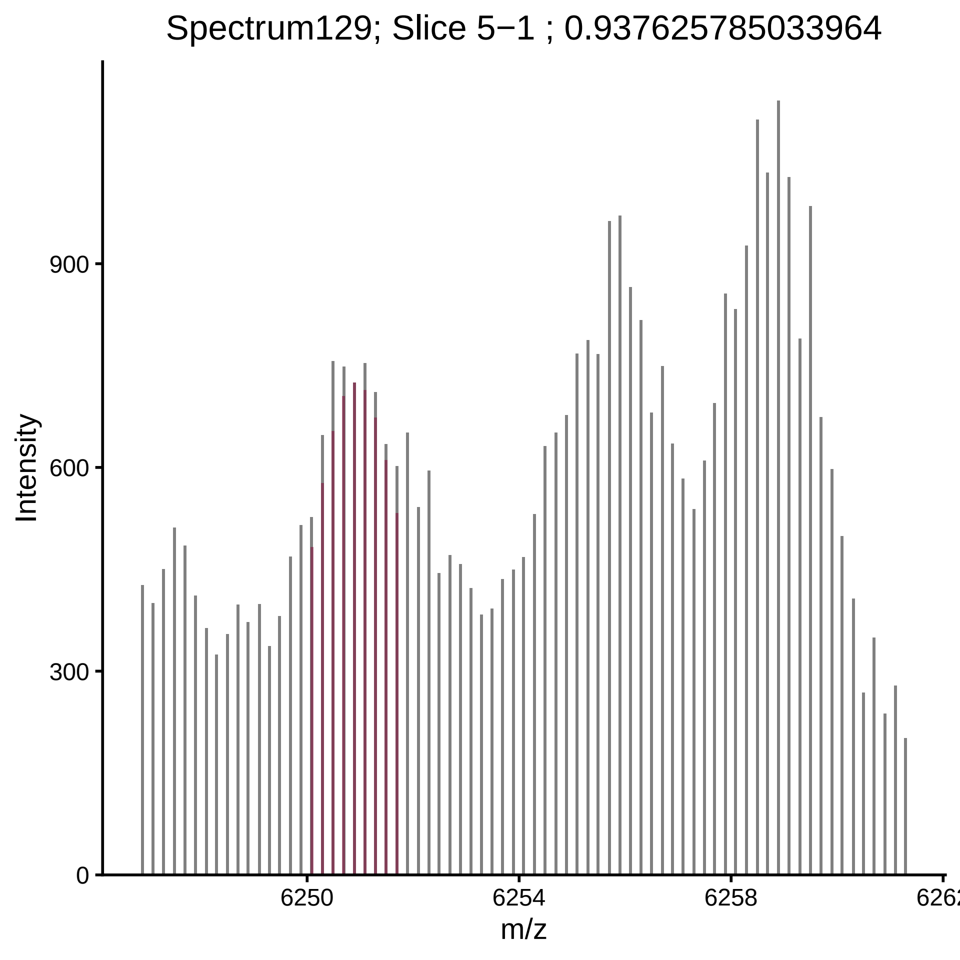

Spectrum129; Slice 5−1 ; 0.937625785033964
Intensity
0
300
600
900
6250
6254
6258
m/z

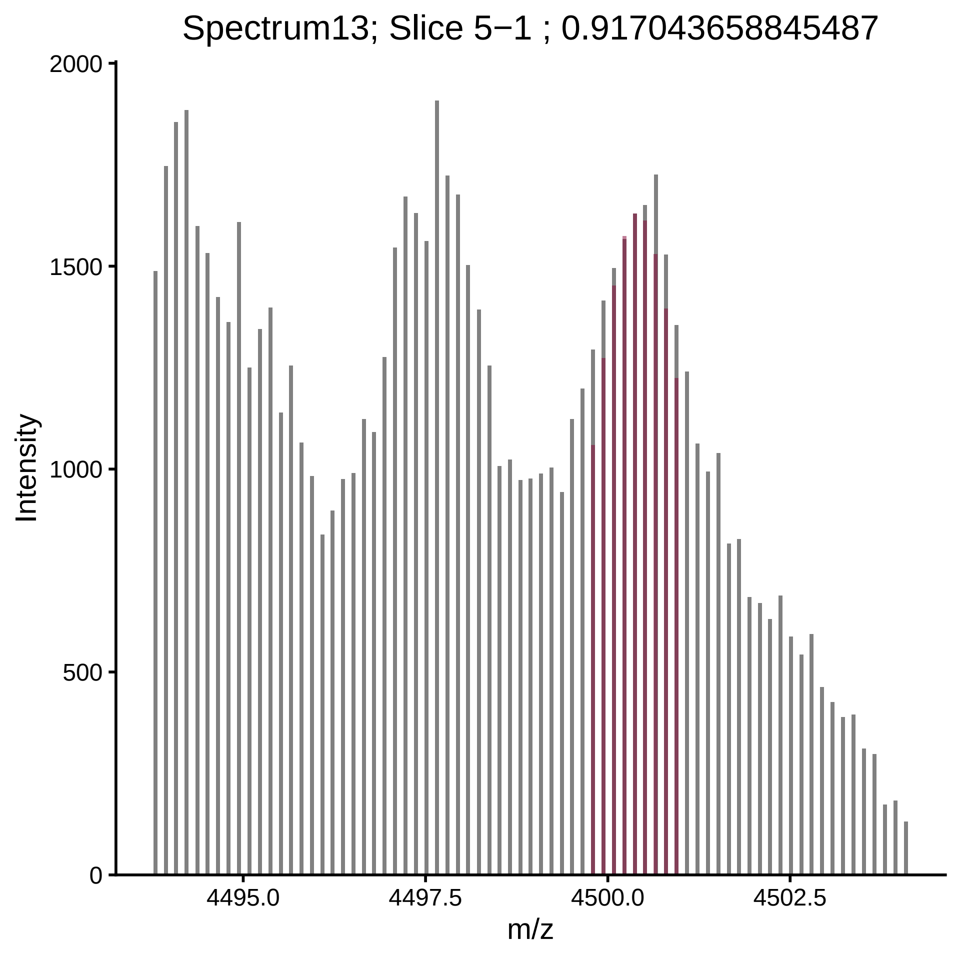
Spectrum13; Slice 5−1 ; 0.917043658845487
Intensity
2000
1500
1000
500
0
4495.0
4497.5
4500.0
4502.5
m/z

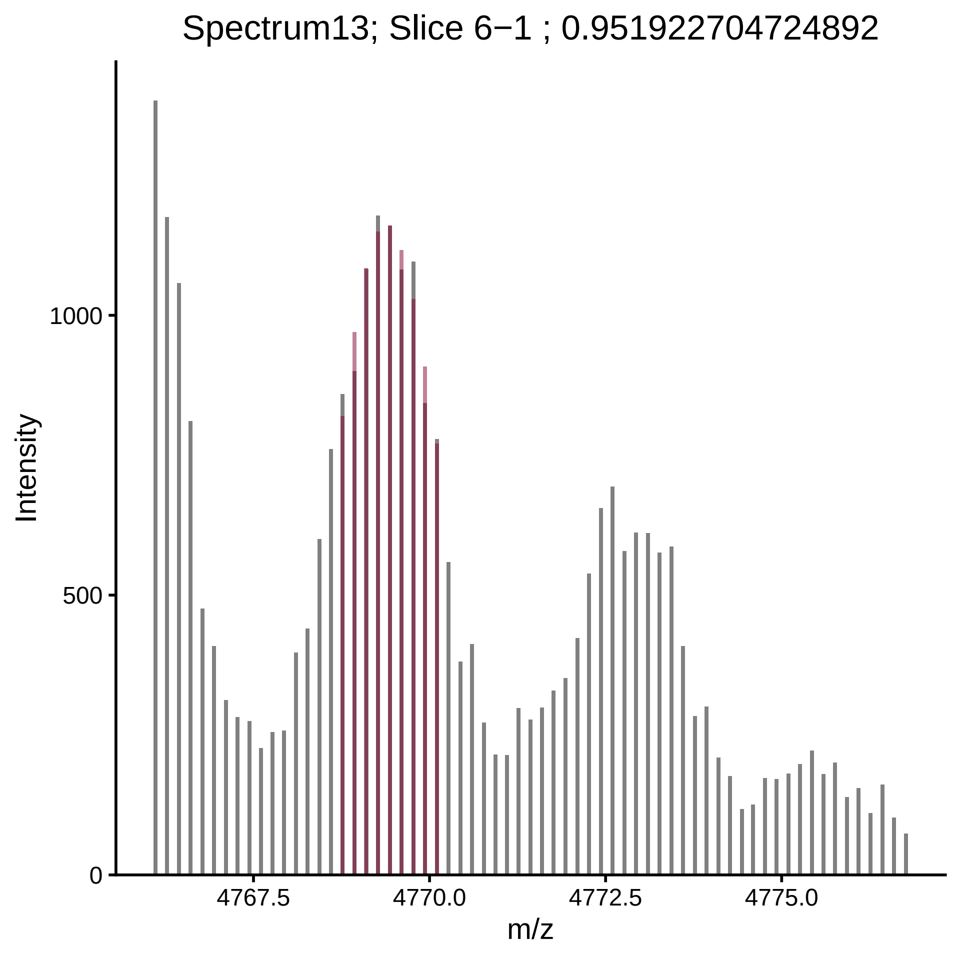

Spectrum13; Slice 6−1 ; 0.951922704724892
Intensity
1000
500
0
4767.5
4770.0
4772.5
4775.0
m/z

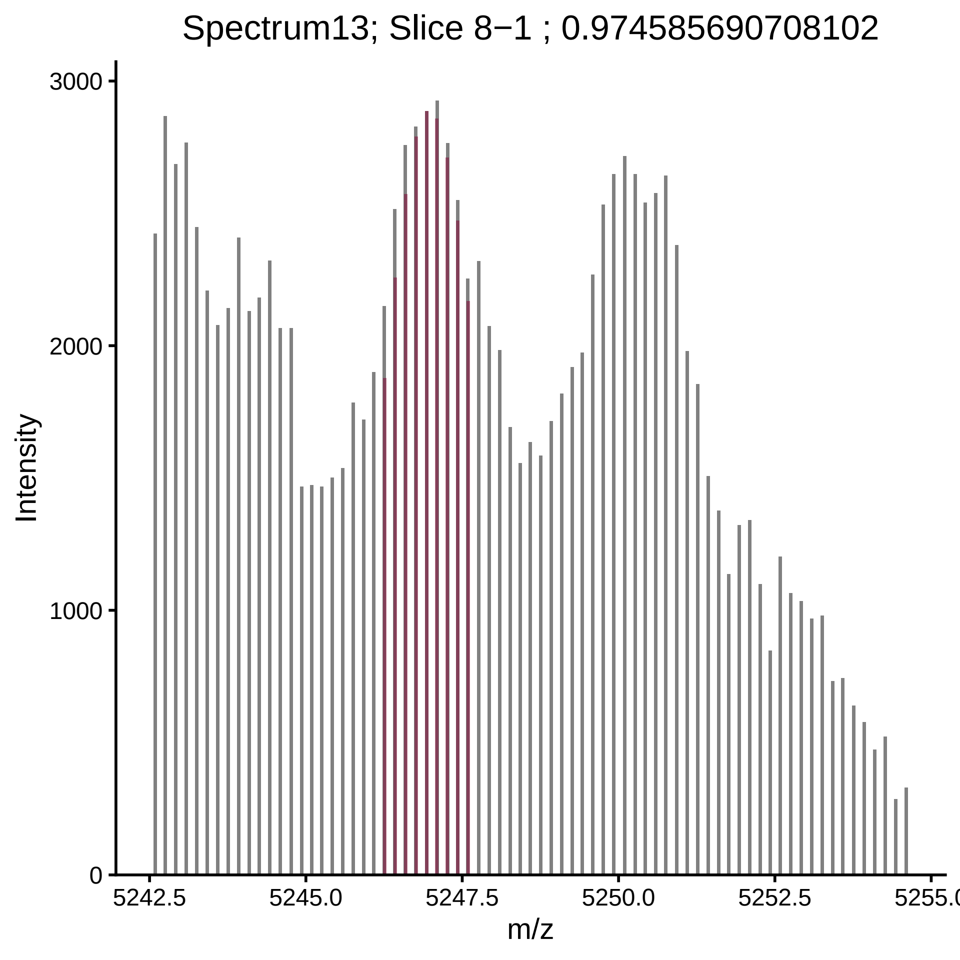
Spectrum13; Slice 8−1 ; 0.974585690708102
Intensity
3000
2000
1000
0
5242.5
5245.0
5247.5
5250.0
5252.5
m/z

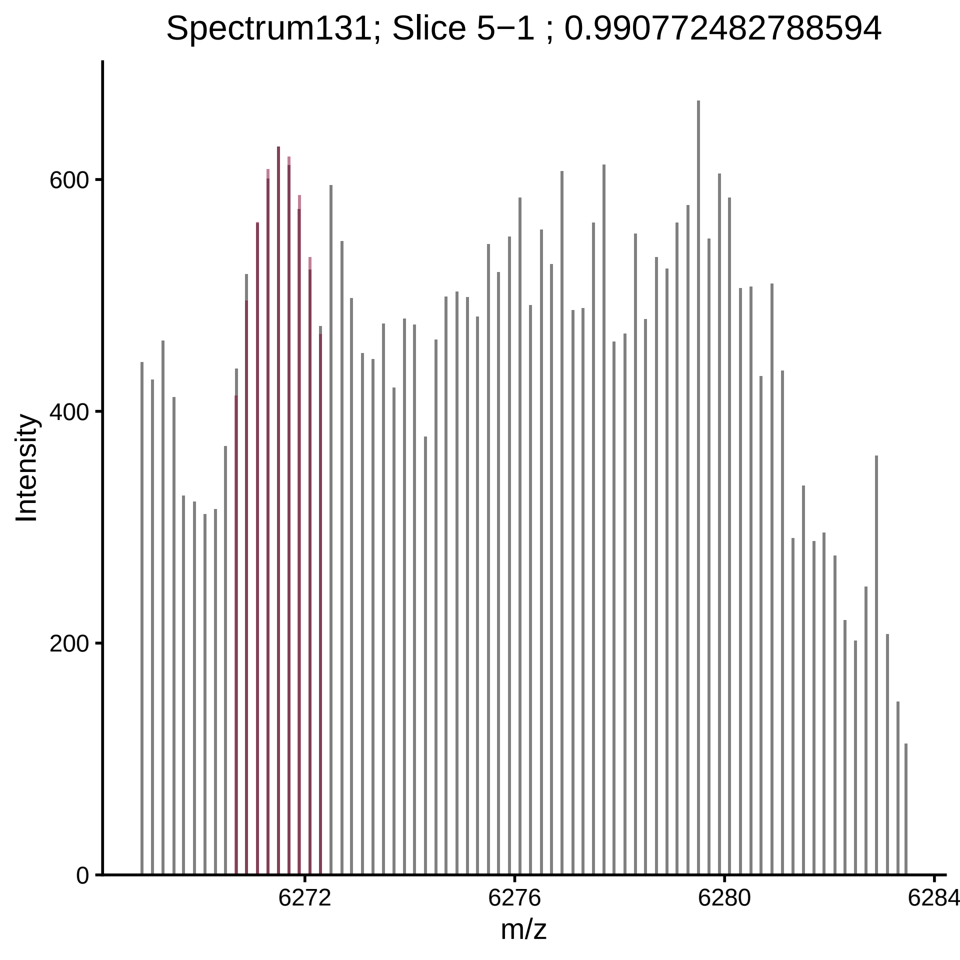

Spectrum131; Slice 5−1 ; 0.990772482788594
Intensity
600
400
200
0
6272
6276
6280
6284
m/z

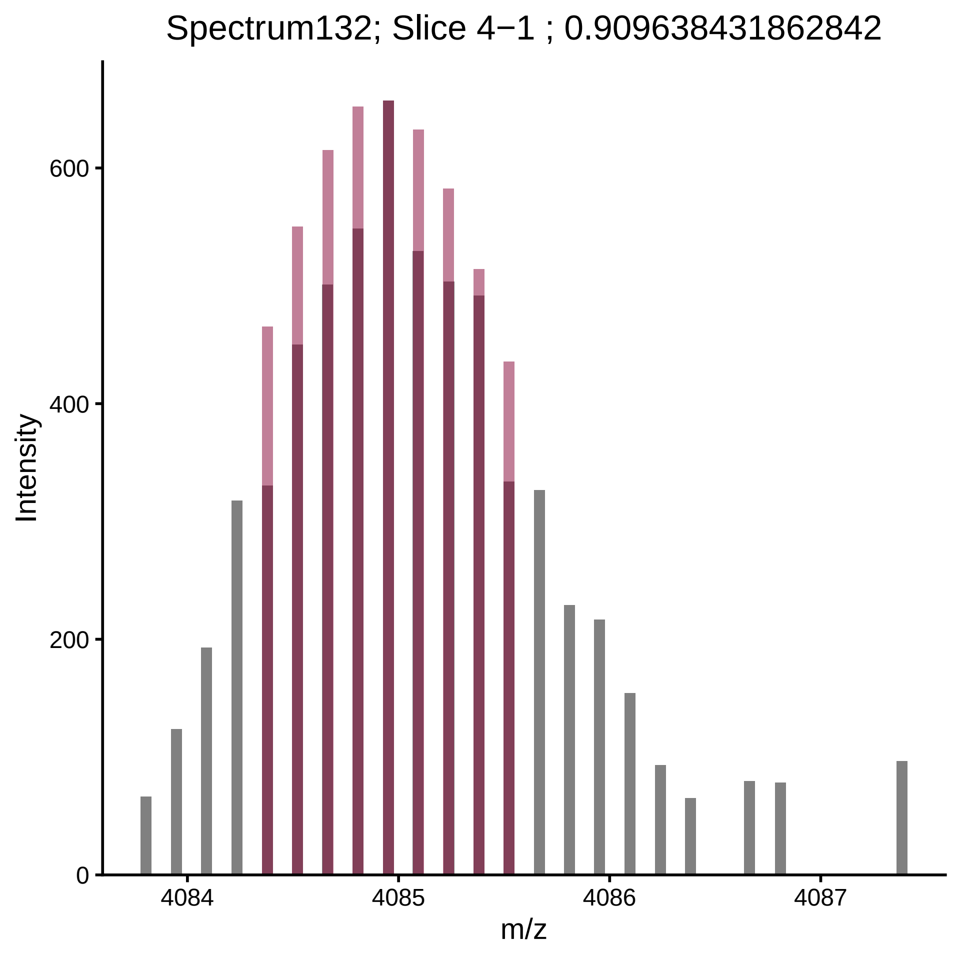

Spectrum132; Slice 4−1 ; 0.909638431862842
Intensity
600
400
200
0
4084
4085
4086
4087
m/z

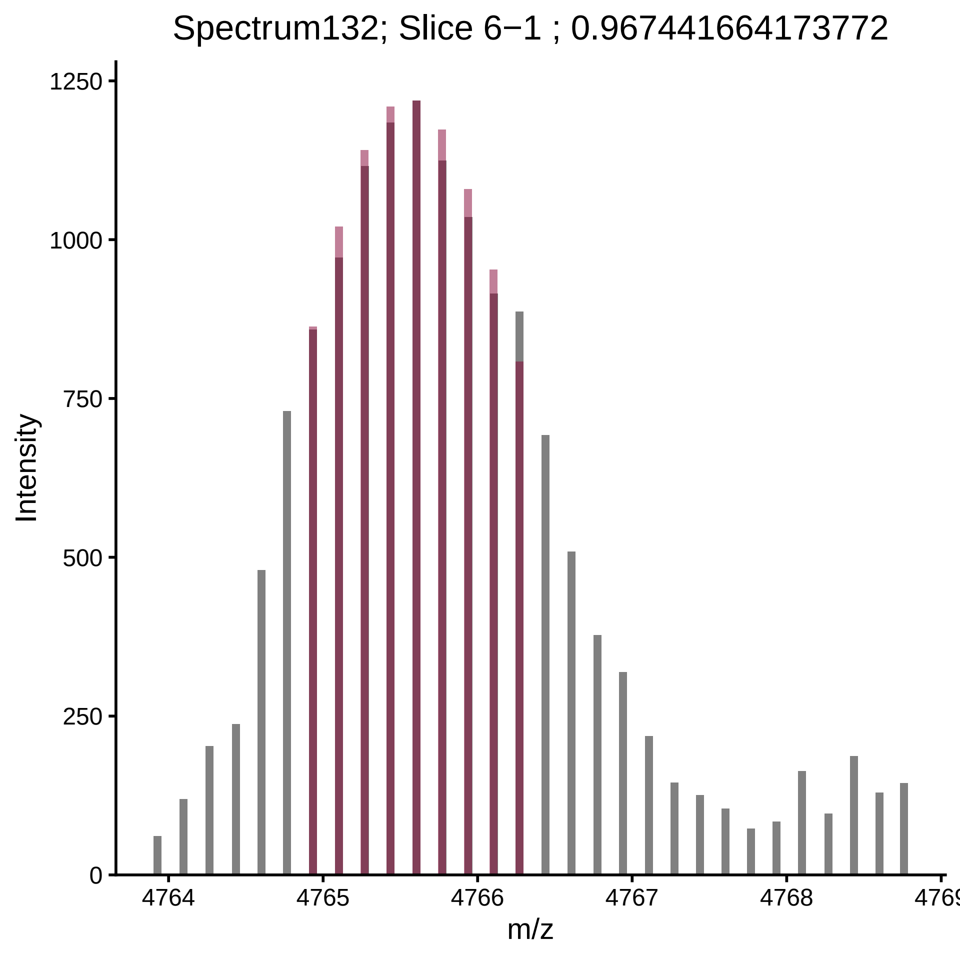

Spectrum132; Slice 6−1 ; 0.967441664173772
Intensity
1250
1000
750
500
250
0
4764
4765
4766
4767
4768
4769
m/z

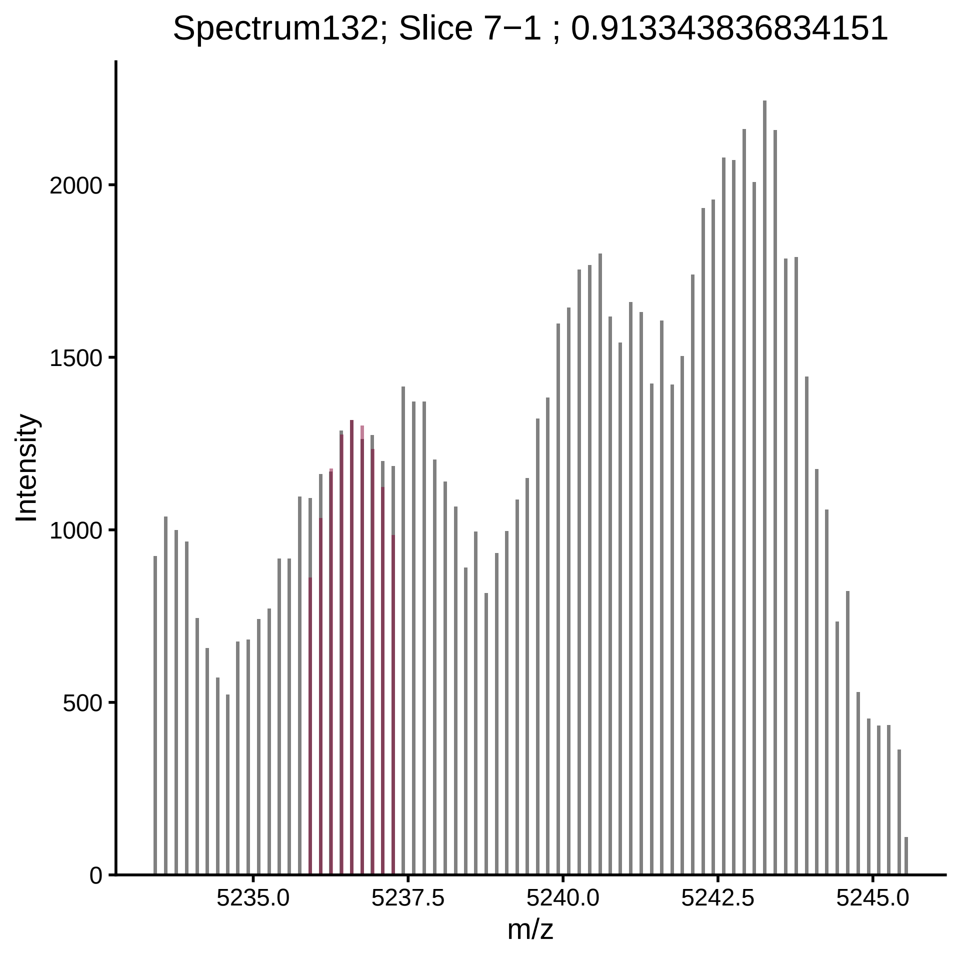

Spectrum132; Slice 7−1 ; 0.913343836834151
Intensity
2000
1500
1000
500
0
5235.0
5237.5
5240.0
5242.5
5245.0
m/z

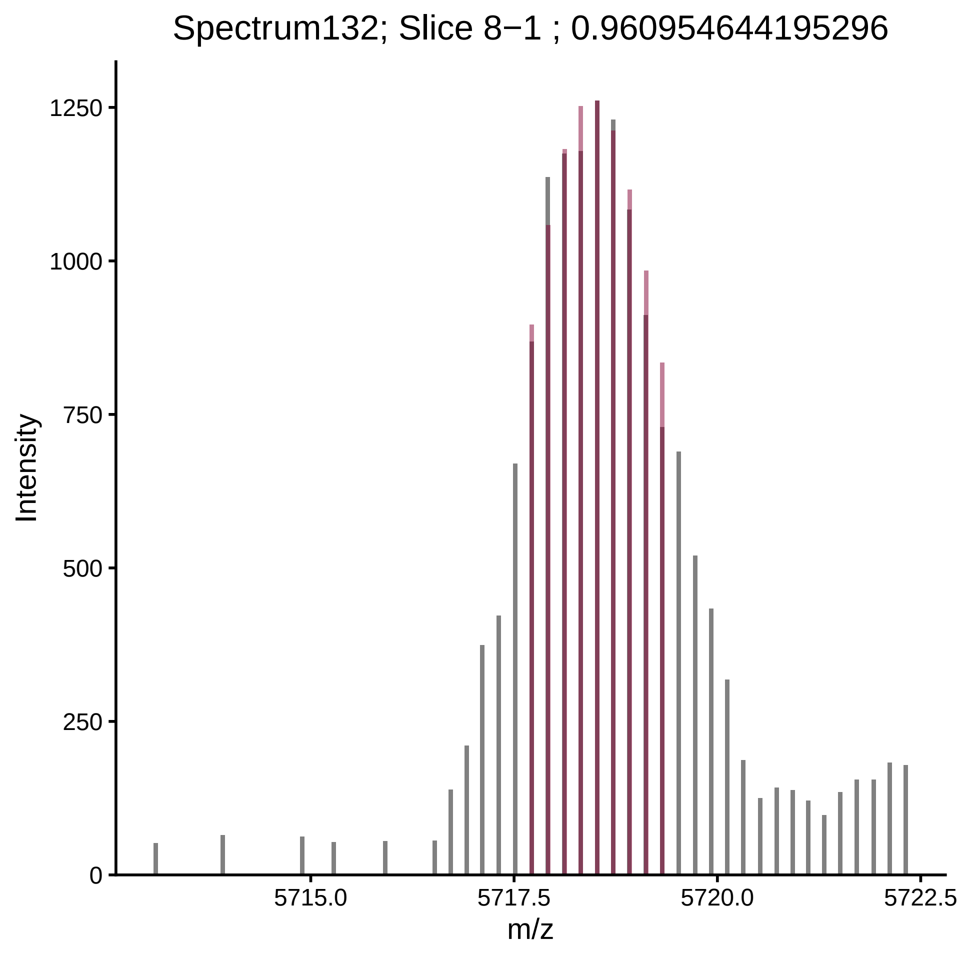
Spectrum132; Slice 8−1 ; 0.960954644195296
Intensity
1250
1000
750
500
250
0
5715.0
5717.5
5720.0
5722.5
m/z

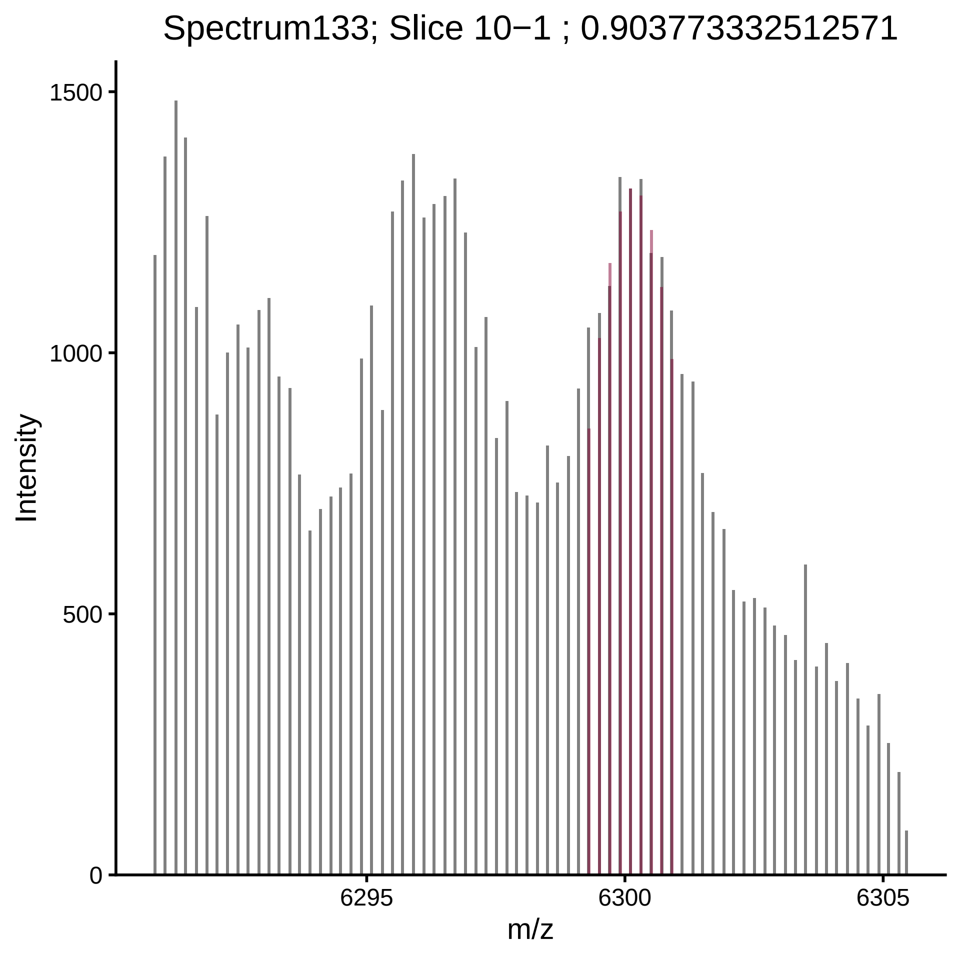

Spectrum133; Slice 10−1 ; 0.903773332512571
Intensity
1500
1000
500
0
6295
6300
6305
m/z

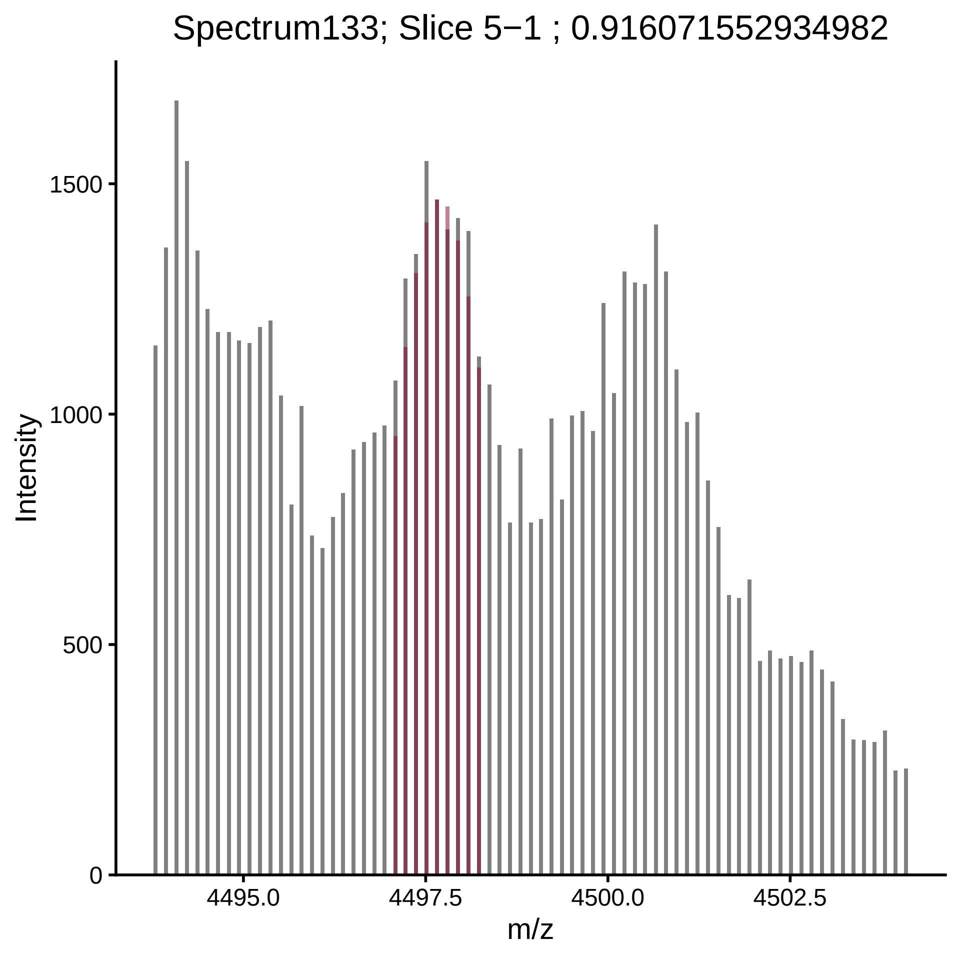
Spectrum133; Slice 5−1 ; 0.916071552934982
Intensity
1500
1000
500
0
4495.0
4497.5
4500.0
4502.5
m/z

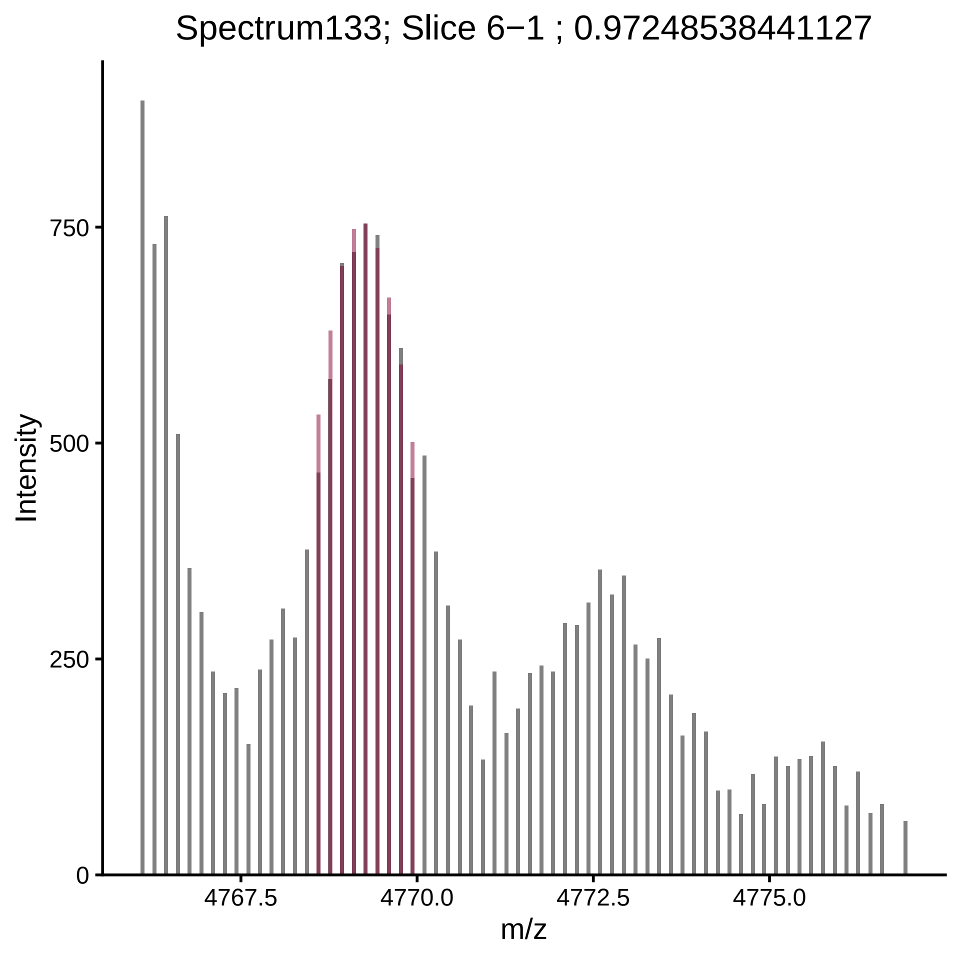
Spectrum133; Slice 6−1 ; 0.97248538441127
Intensity
750
500
250
0
4767.5
4770.0
4772.5
4775.0
m/z

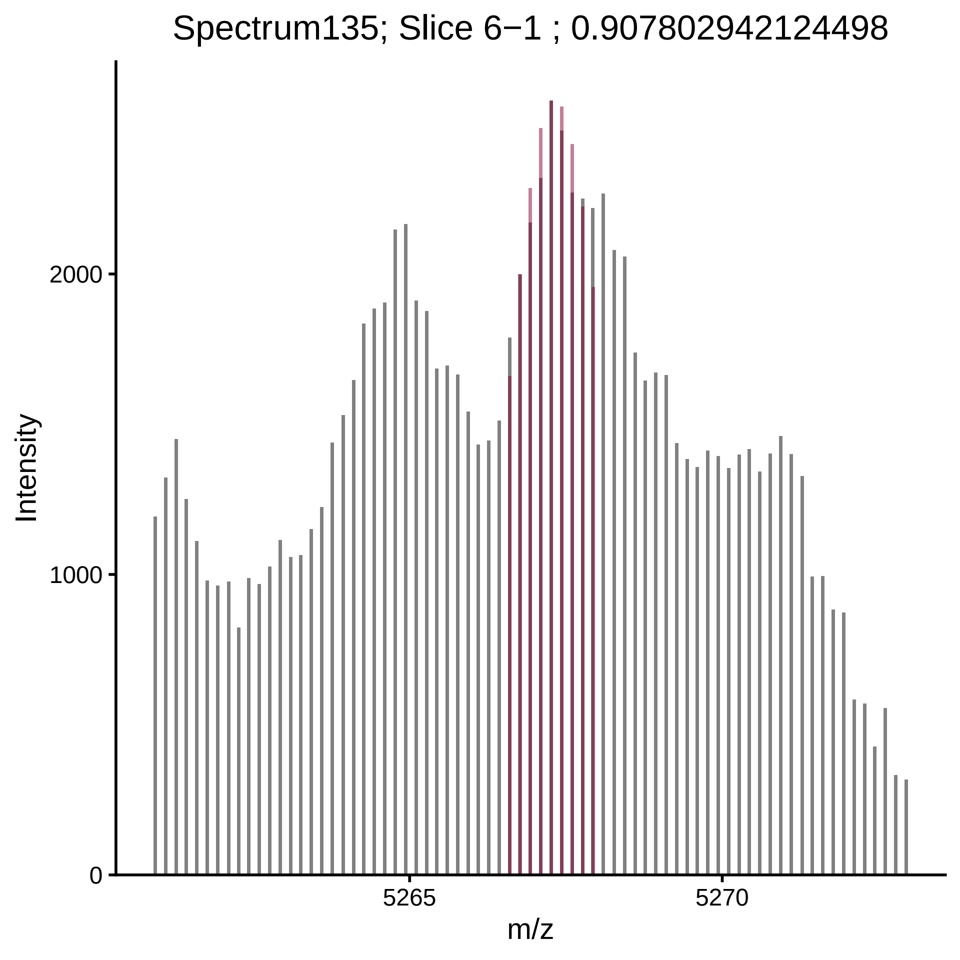

Spectrum135; Slice 6−1 ; 0.907802942124498
Intensity
2000
1000
0
5265
5270
m/z

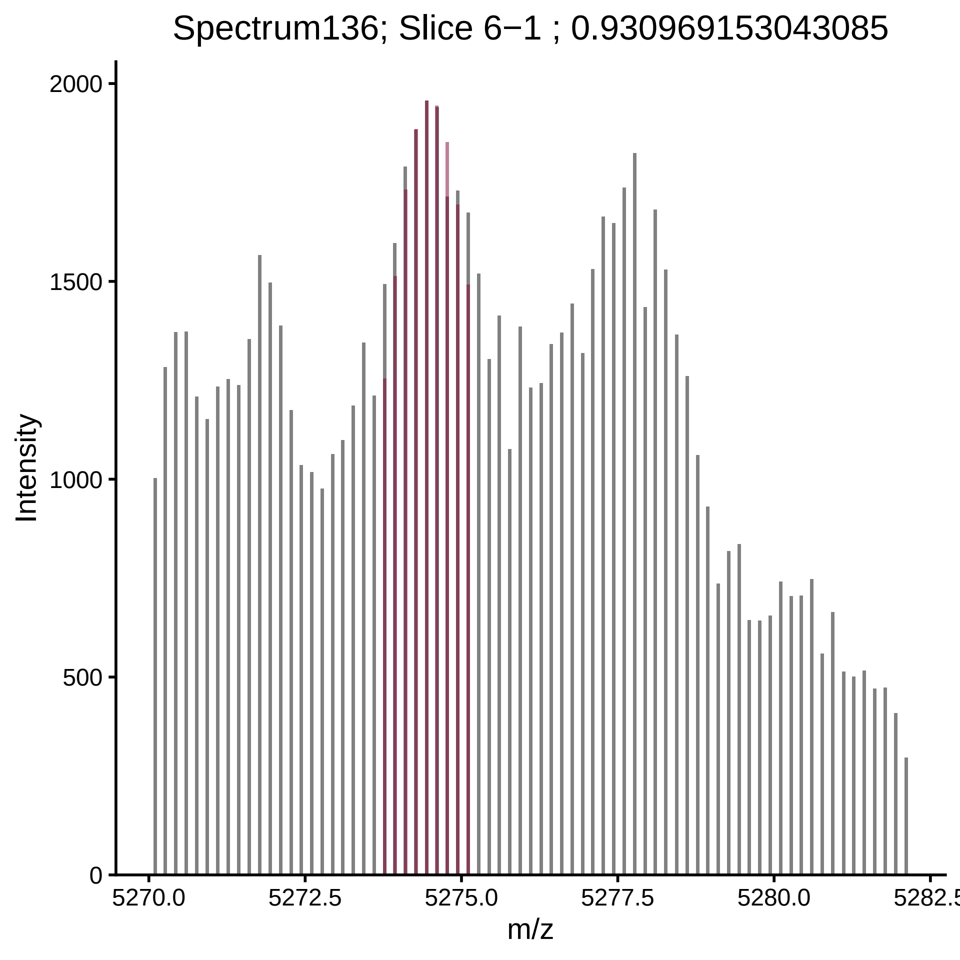

Spectrum136; Slice 6−1 ; 0.930969153043085
Intensity
2000
1500
1000
500
0
5270.0
5272.5
5275.0
5277.5
5280.0
5282.5
m/z

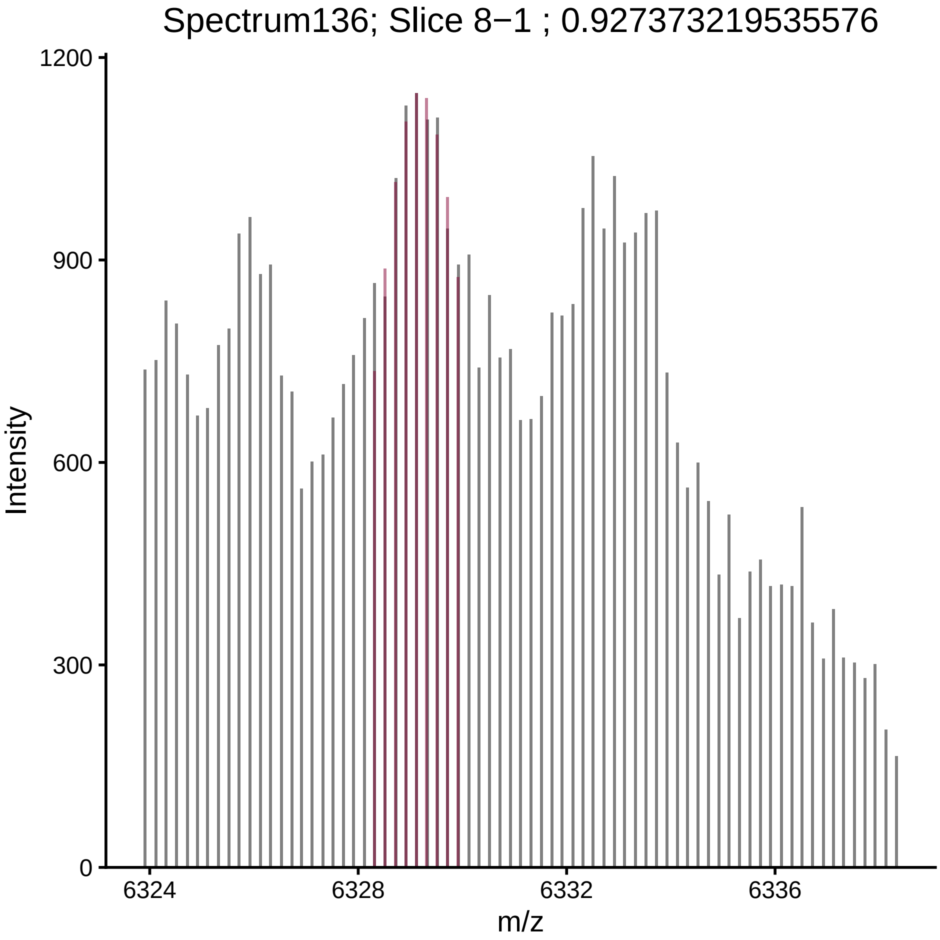

Spectrum136; Slice 8−1 ; 0.927373219535576
Intensity
0
300
600
900
1200
6324
6328
6332
6336
m/z

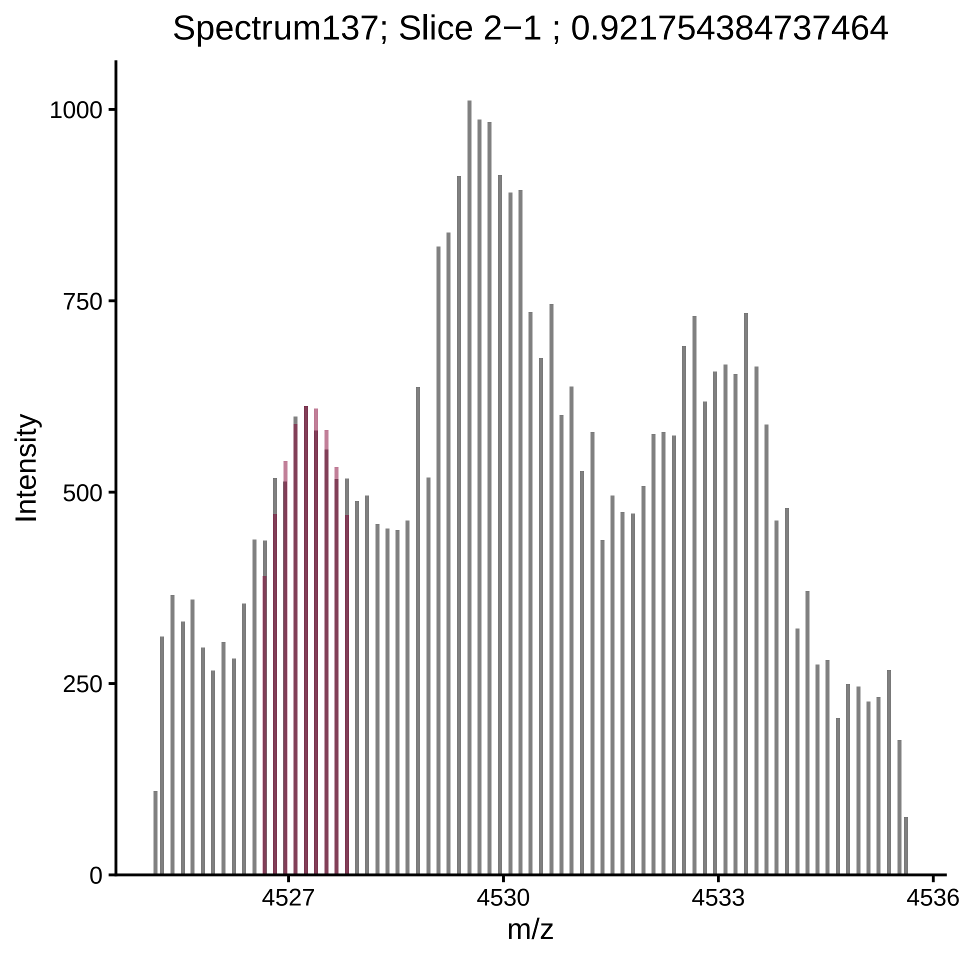

Spectrum137; Slice 2−1 ; 0.921754384737464
Intensity
1000
750
500
250
0
4527
4530
4533
4536
m/z

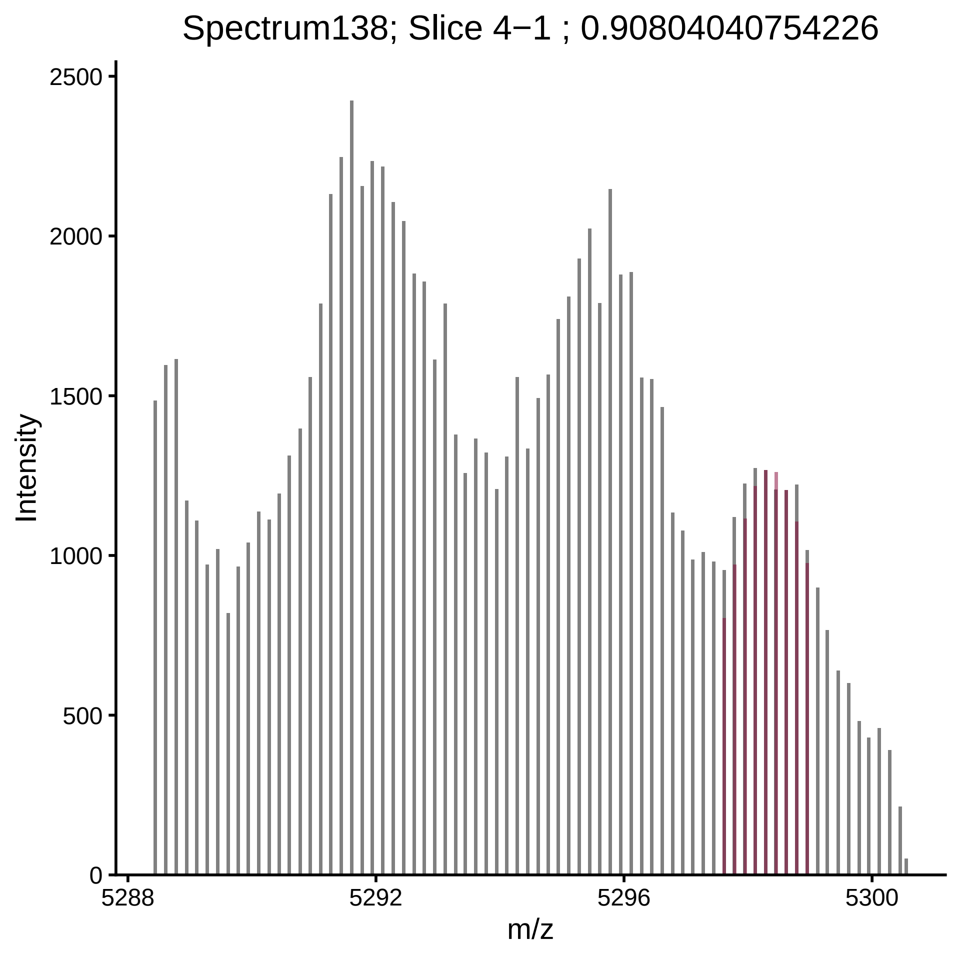

Spectrum138; Slice 4−1 ; 0.90804040754226
Intensity
0
500
1000
1500
2000
2500
5288
5292
5296
5300
m/z

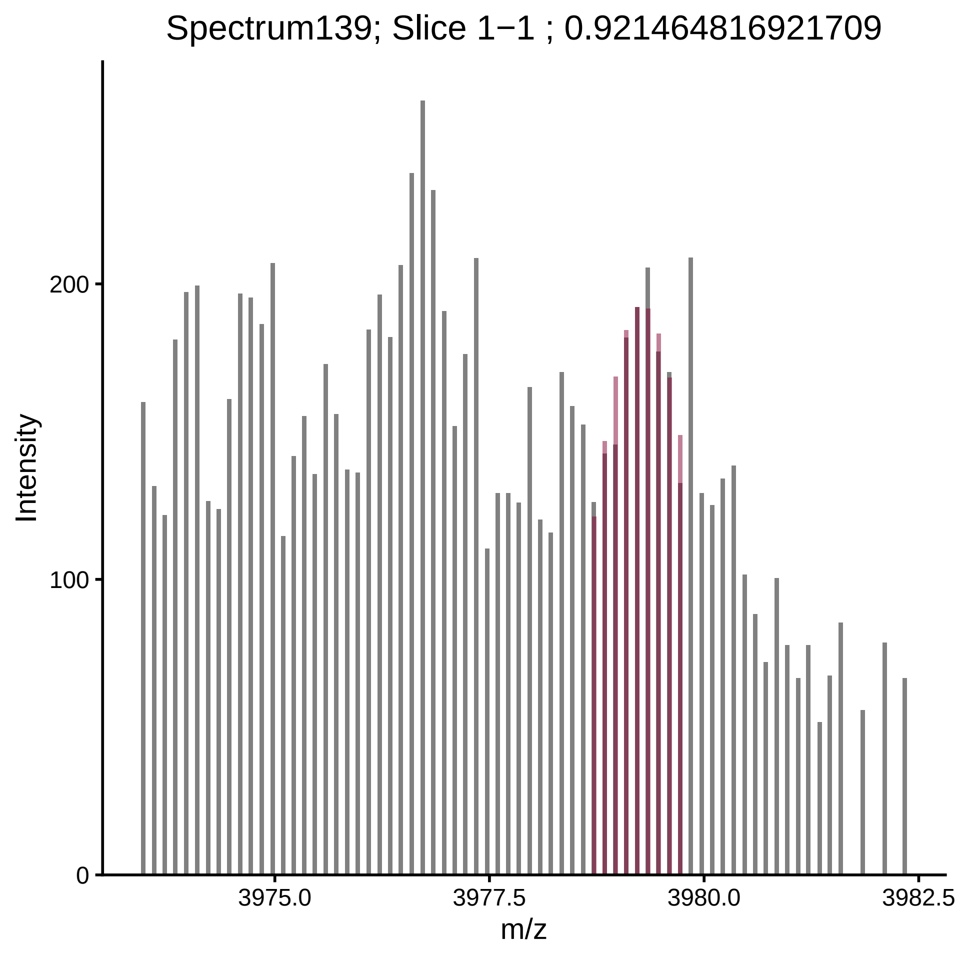

Spectrum139; Slice 1−1 ; 0.921464816921709
Intensity
0
100
200
3975.0
3977.5
3980.0
3982.5
m/z

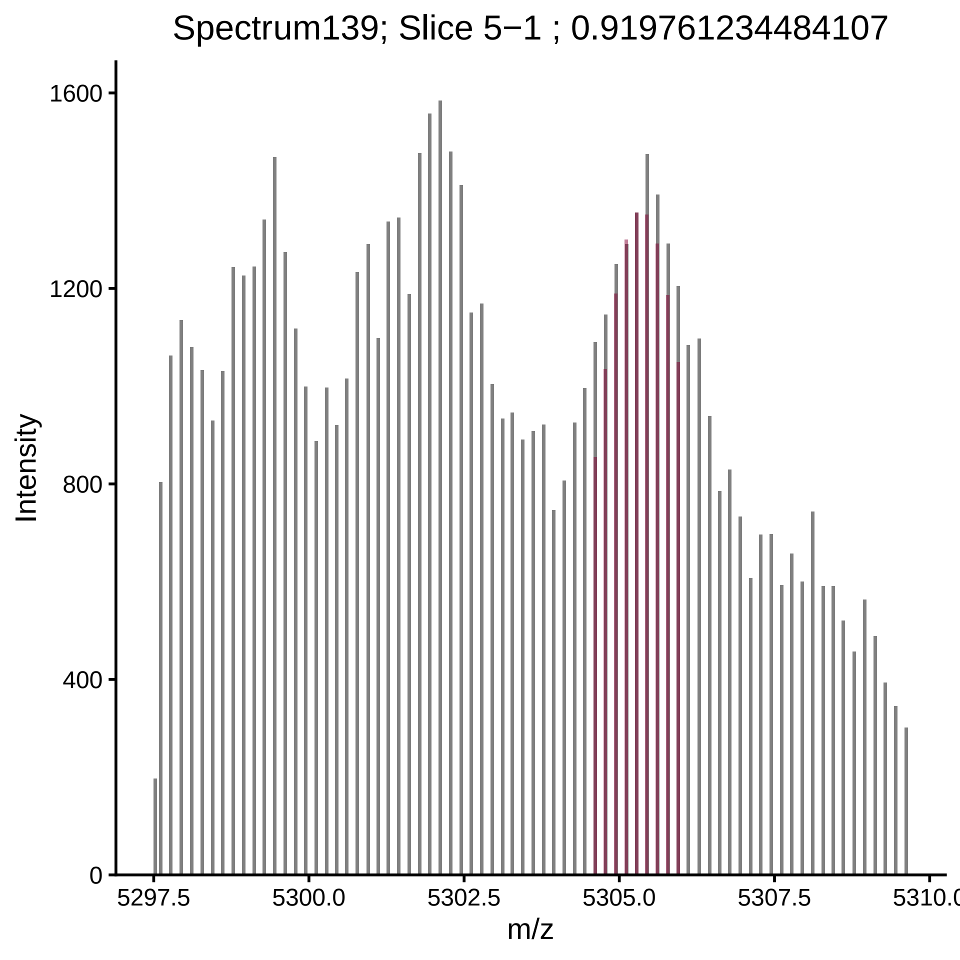

Spectrum139; Slice 5−1 ; 0.919761234484107
Intensity
0
400
800
1200
1600
5297.5
5300.0
5302.5
5305.0
5307.5
5310.0
m/z

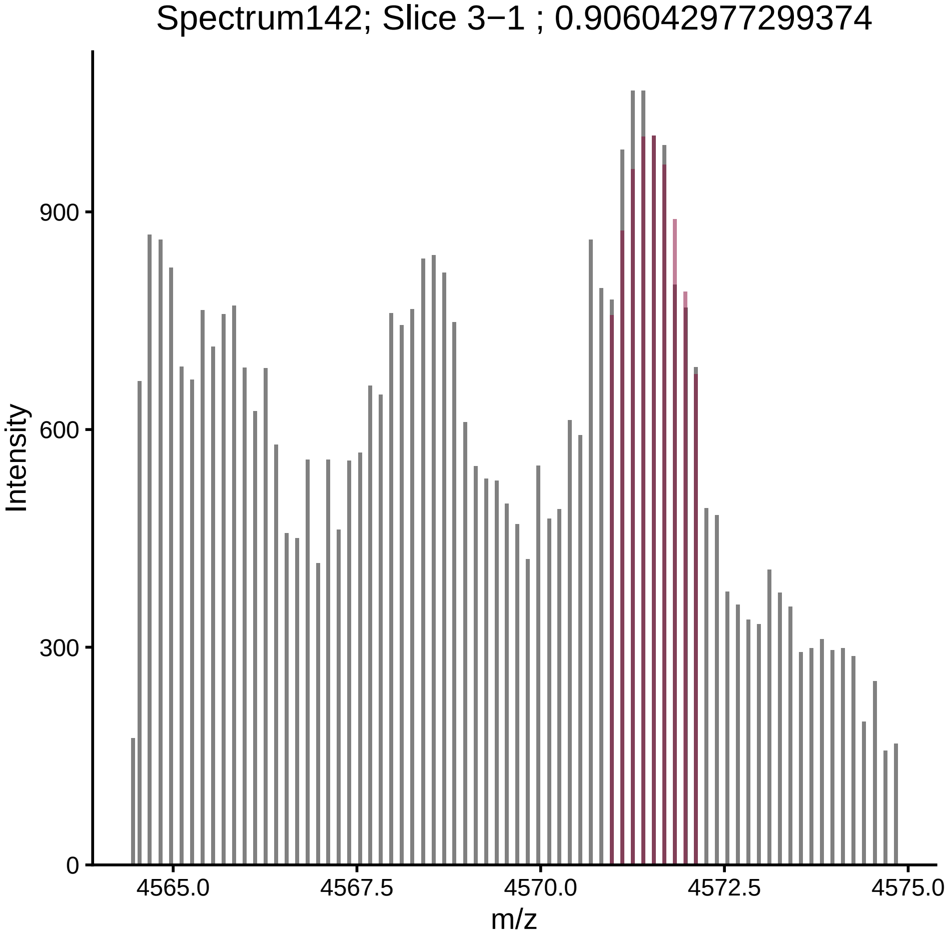

Spectrum142; Slice 3−1 ; 0.906042977299374
Intensity
900
600
300
0
4565.0
4567.5
4570.0
4572.5
4575.0
m/z

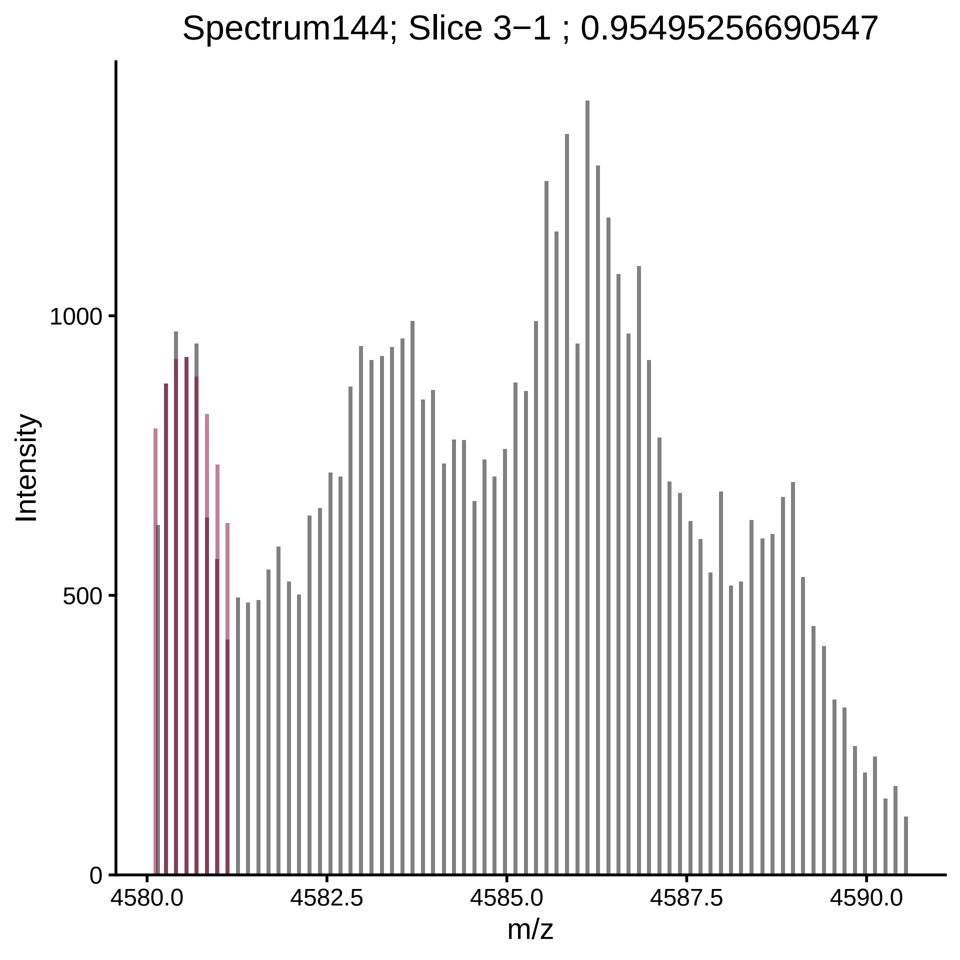

Spectrum144; Slice 3−1 ; 0.95495256690547
Intensity
1000
500
0
4580.0
4582.5
4585.0
4587.5
4590.0
m/z

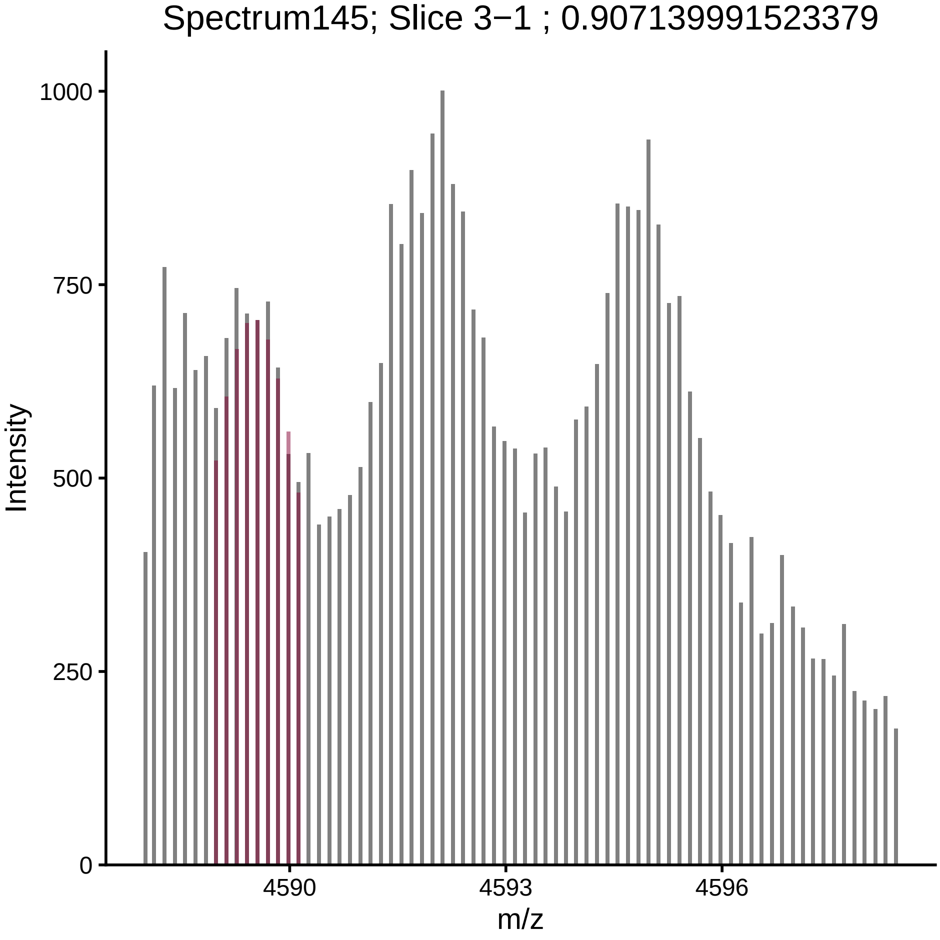

Spectrum145; Slice 3−1 ; 0.907139991523379
Intensity
1000
750
500
250
0
4590
4593
4596
m/z

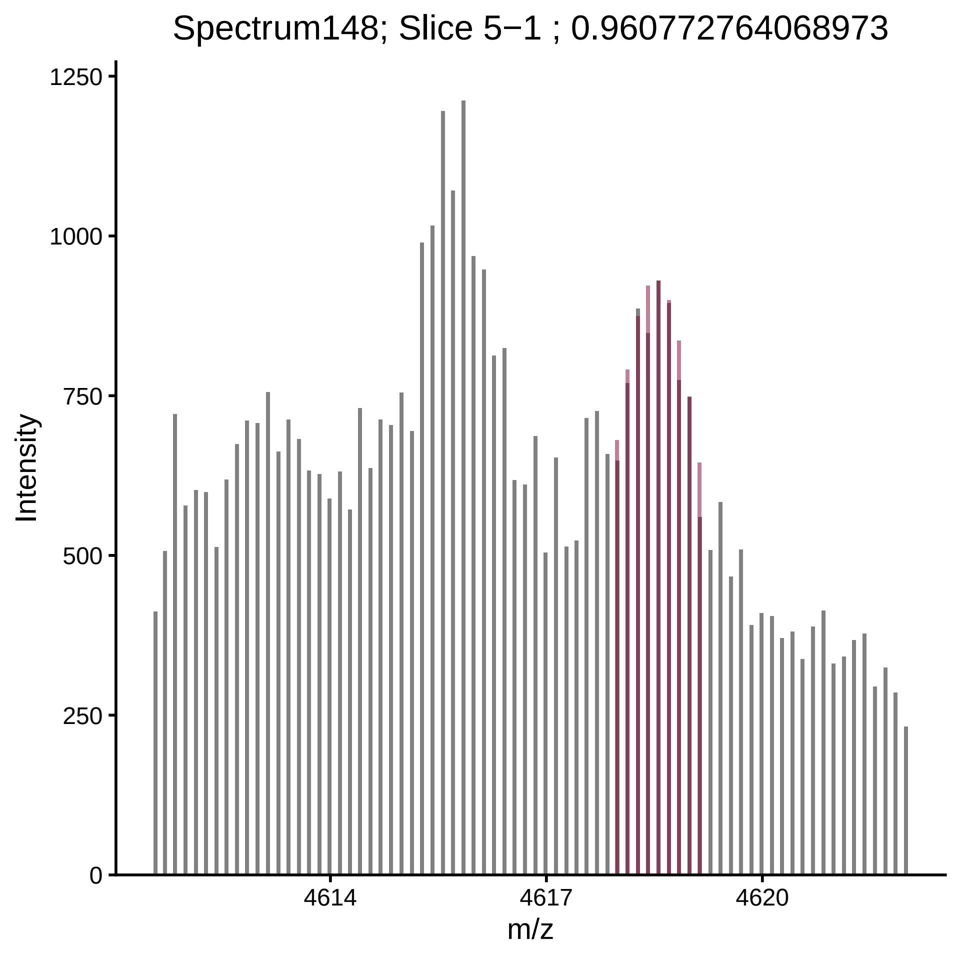

Spectrum148; Slice 5−1 ; 0.960772764068973
Intensity
1250
1000
750
500
250
0
4614
4617
4620
m/z

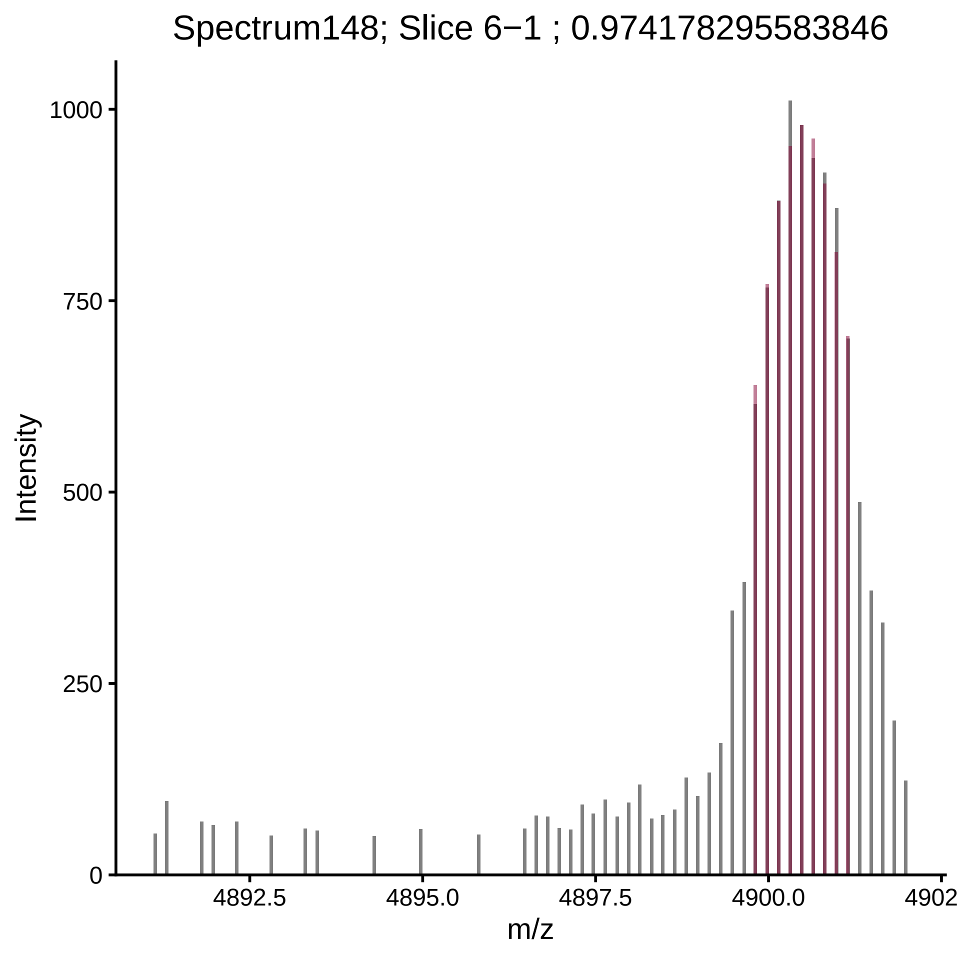

Spectrum148; Slice 6−1 ; 0.974178295583846
Intensity
1000
750
500
250
0
4892.5
4895.0
4897.5
4900.0
4902
m/z

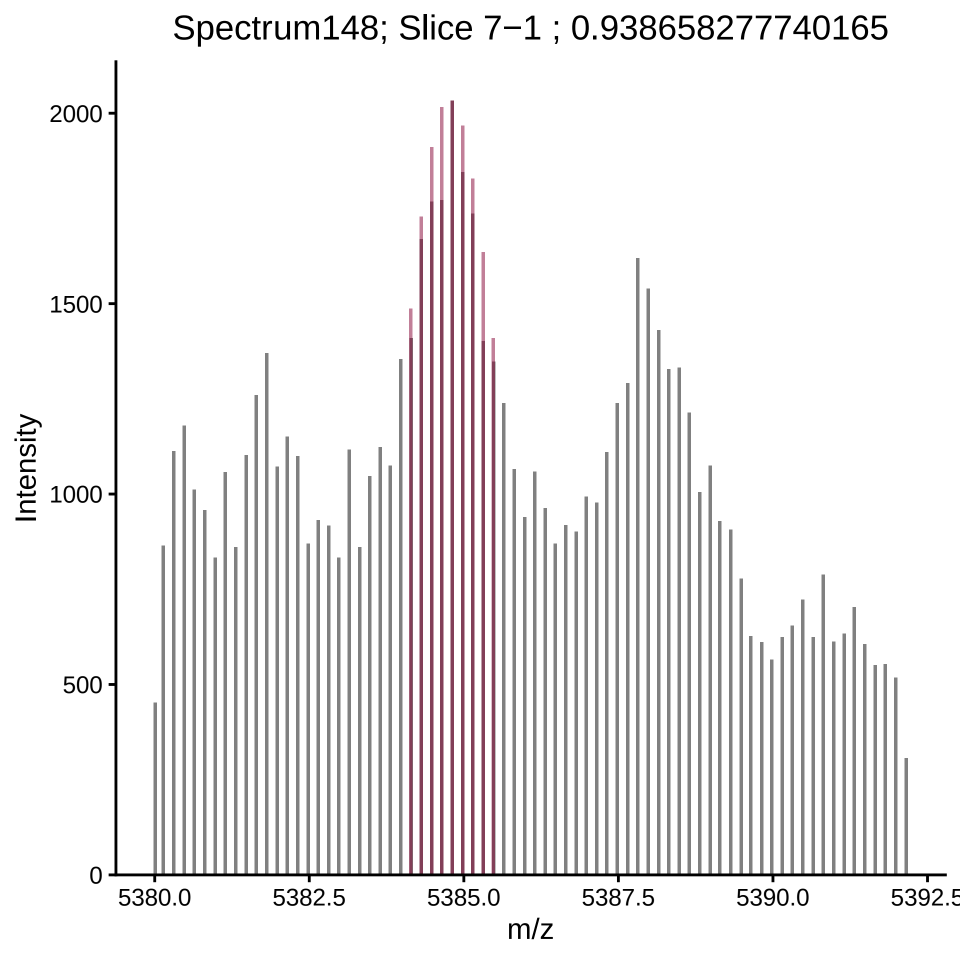

Spectrum148; Slice 7−1 ; 0.938658277740165
Intensity
2000
1500
1000
500
0
5380.0
5382.5
5385.0
5387.5
5390.0
5392.5
m/z

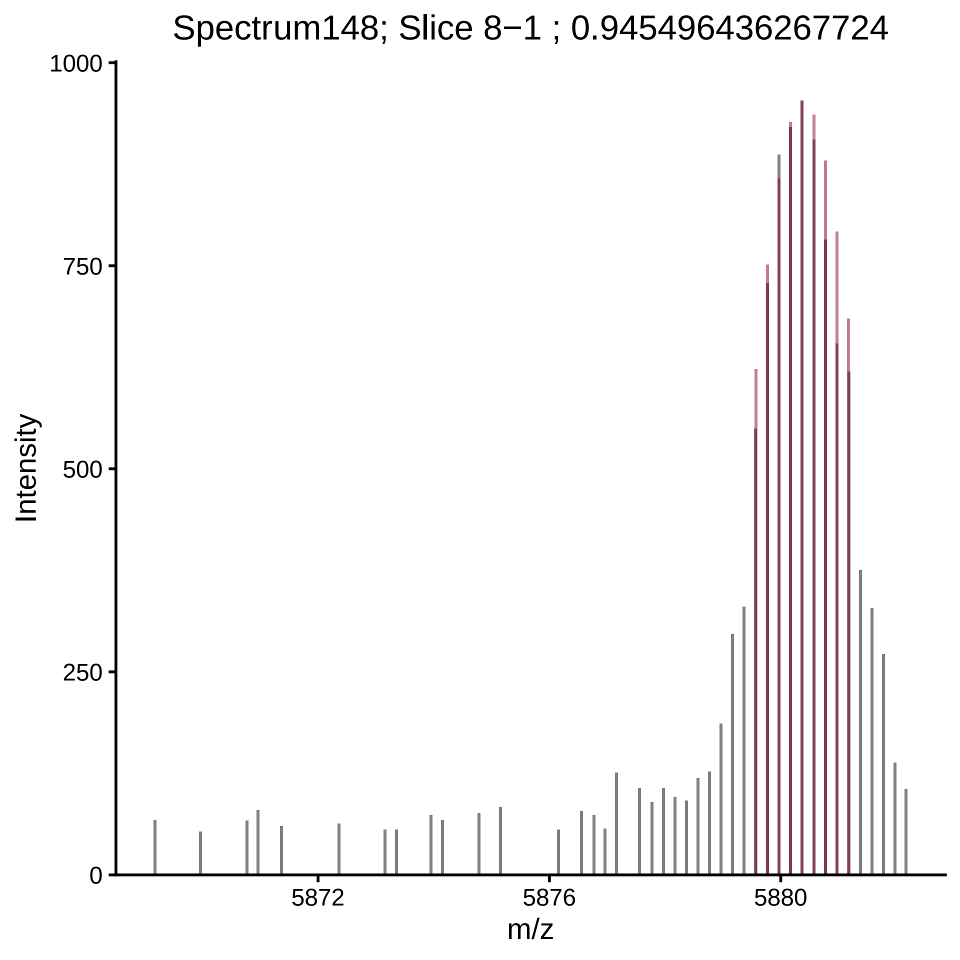

Spectrum148; Slice 8−1 ; 0.945496436267724
Intensity
1000
750
500
250
0
5872
5876
5880
m/z

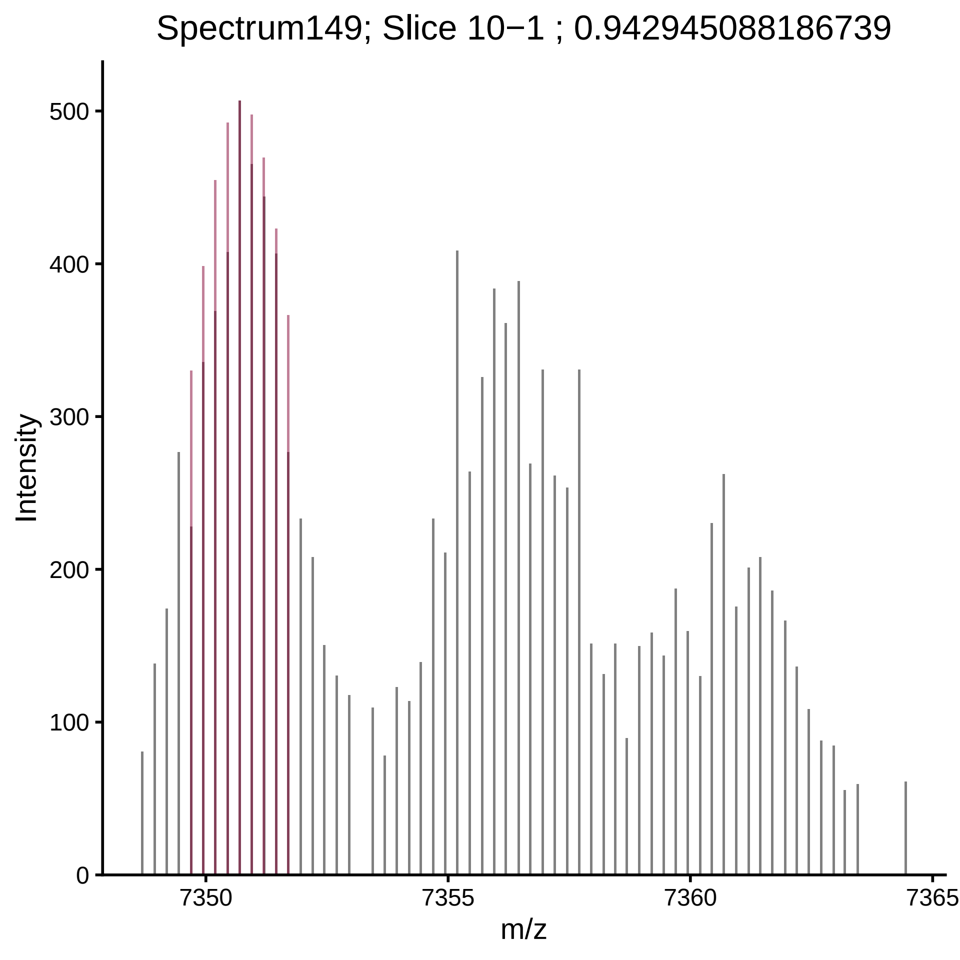

Spectrum149; Slice 10−1 ; 0.942945088186739
Intensity
0
100
200
300
400
500
7350
7355
7360
7365
m/z

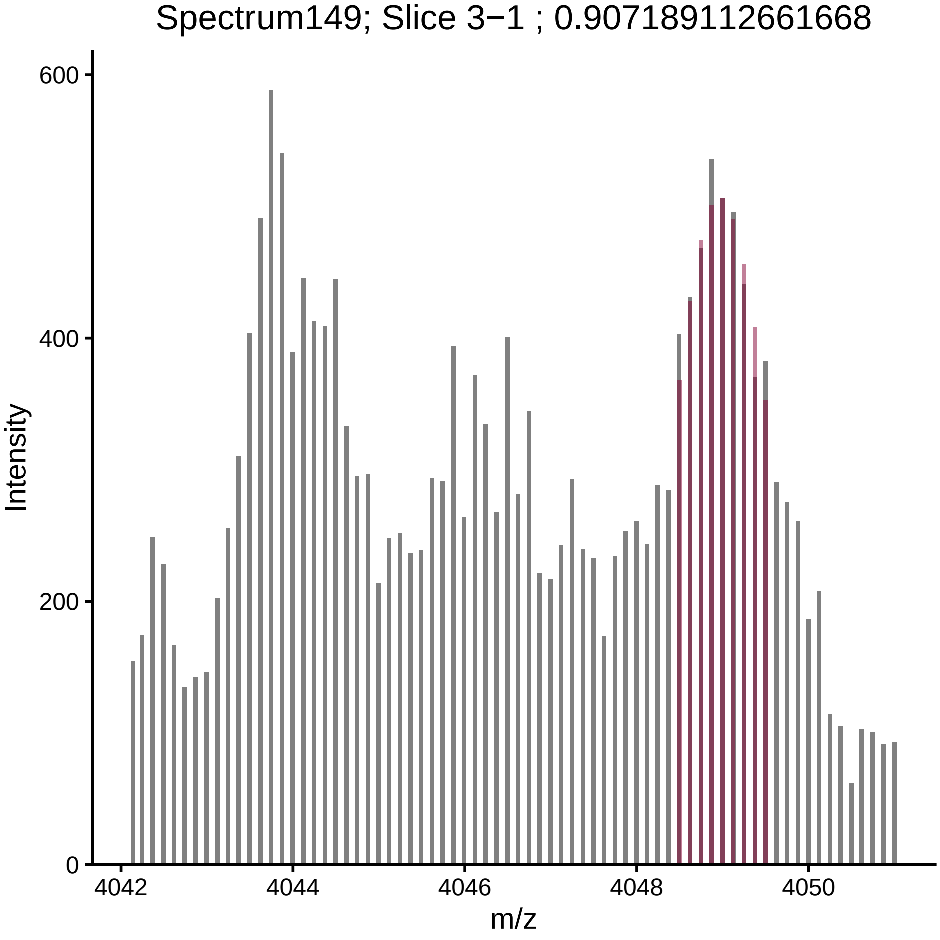

Spectrum149; Slice 3−1 ; 0.907189112661668
Intensity
600
400
200
0
4042
4044
4046
4048
4050
m/z

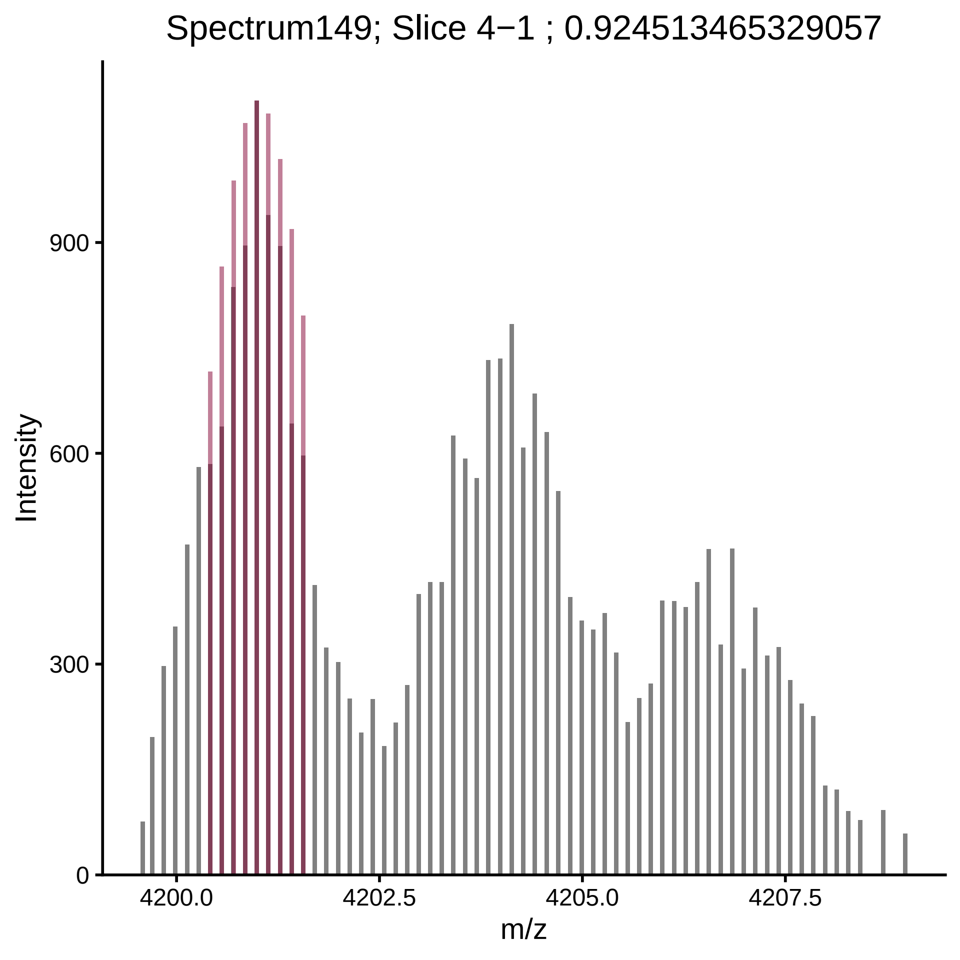
Spectrum149; Slice 4−1 ; 0.924513465329057
Intensity
0
300
600
900
4200.0
4202.5
4205.0
4207.5
m/z

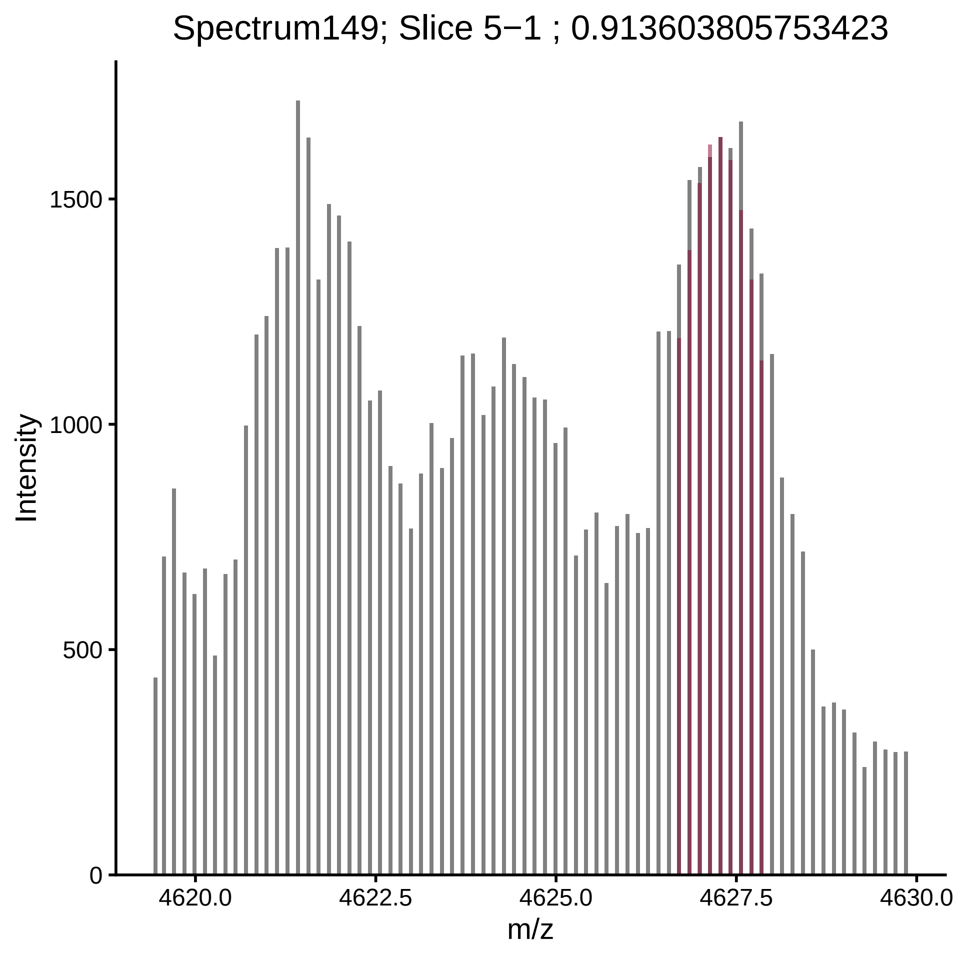

Spectrum149; Slice 5−1 ; 0.913603805753423
Intensity
m/z
0
500
1000
1500
4620.0
4622.5
4625.0
4627.5
4630.0

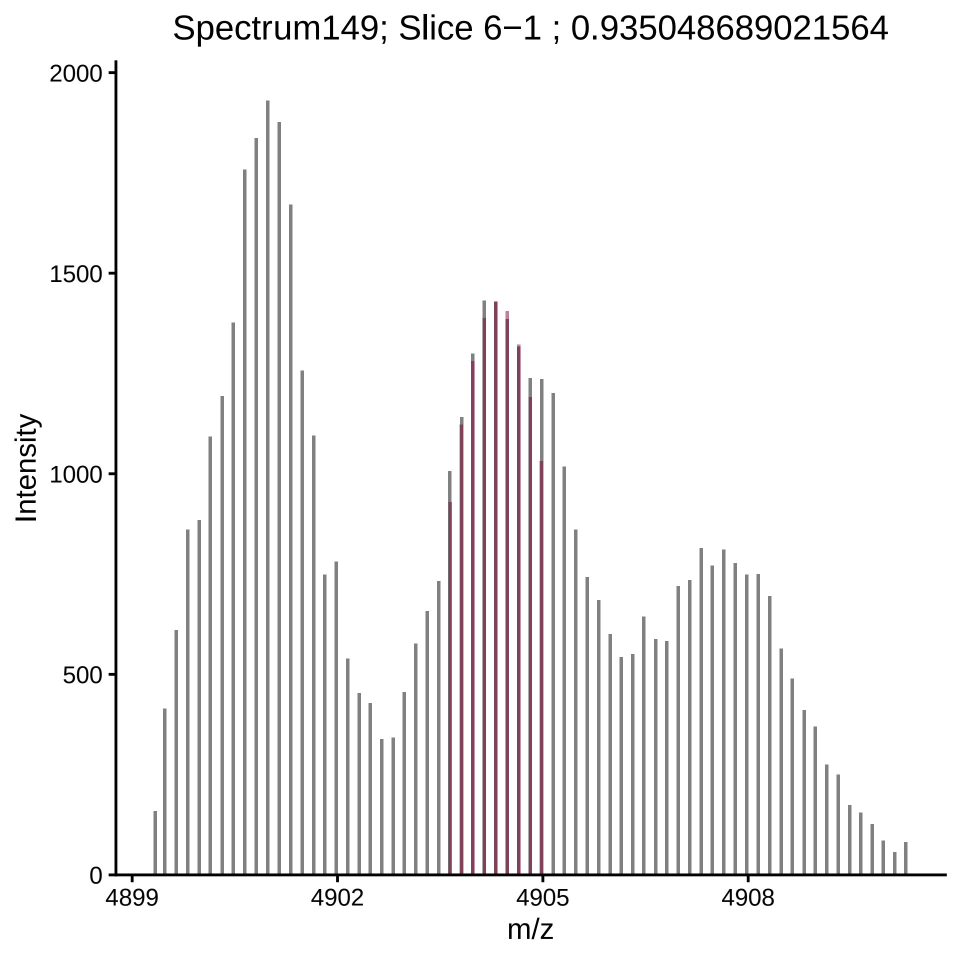
Spectrum149; Slice 6−1 ; 0.935048689021564
Intensity
2000
1500
1000
500
0
4899
4902
4905
4908
m/z

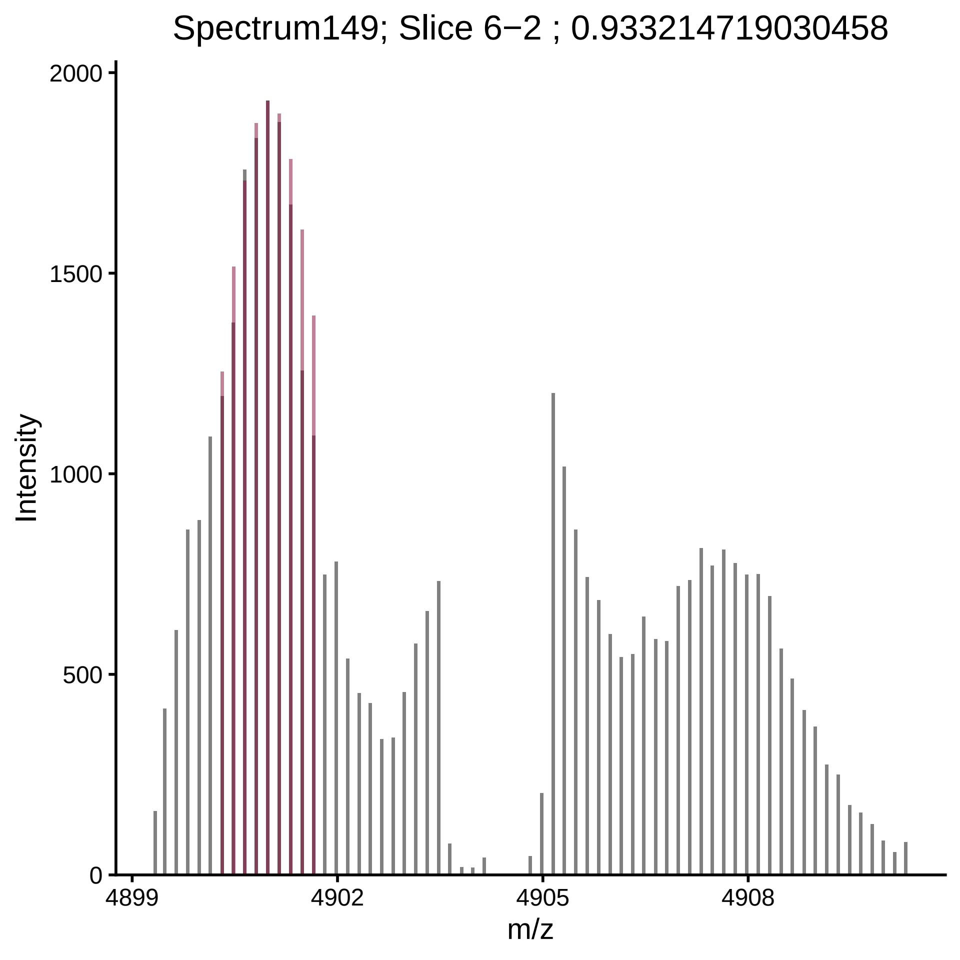

Spectrum149; Slice 6−2 ; 0.933214719030458
Intensity
2000
1500
1000
500
0
4899
4902
4905
4908
m/z

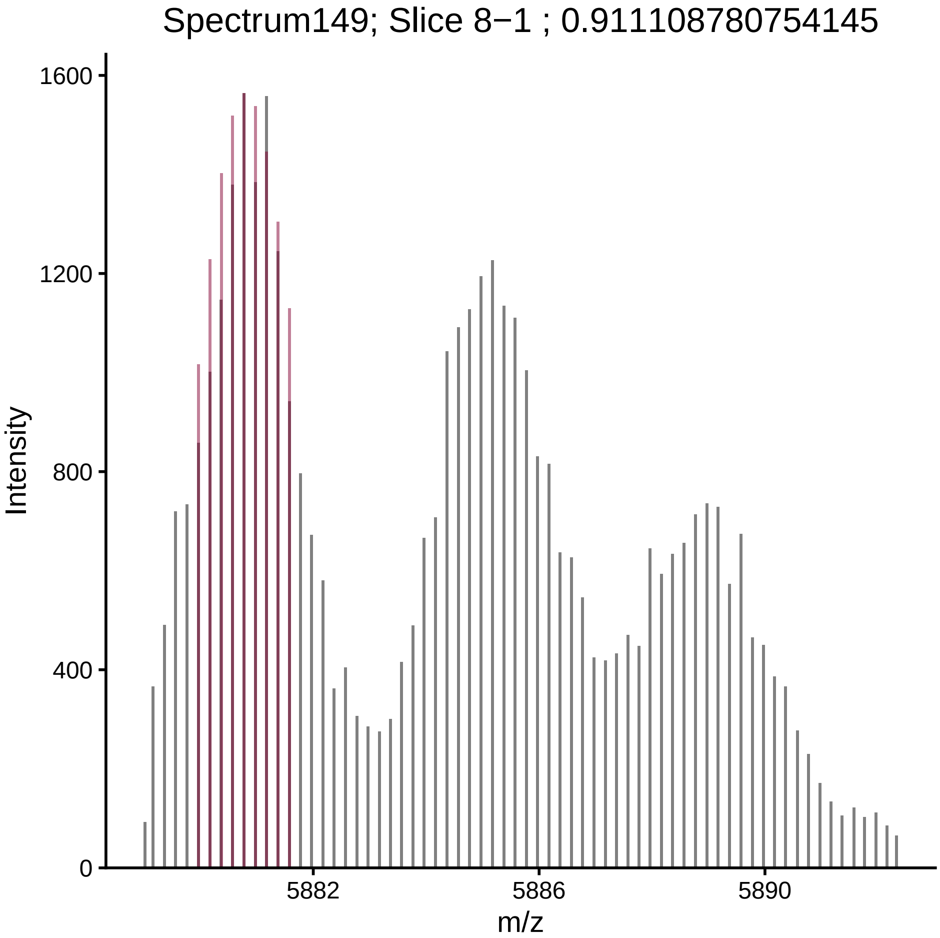

Spectrum149; Slice 8−1 ; 0.911108780754145
Intensity
1600
1200
800
400
0
5882
5886
5890
m/z

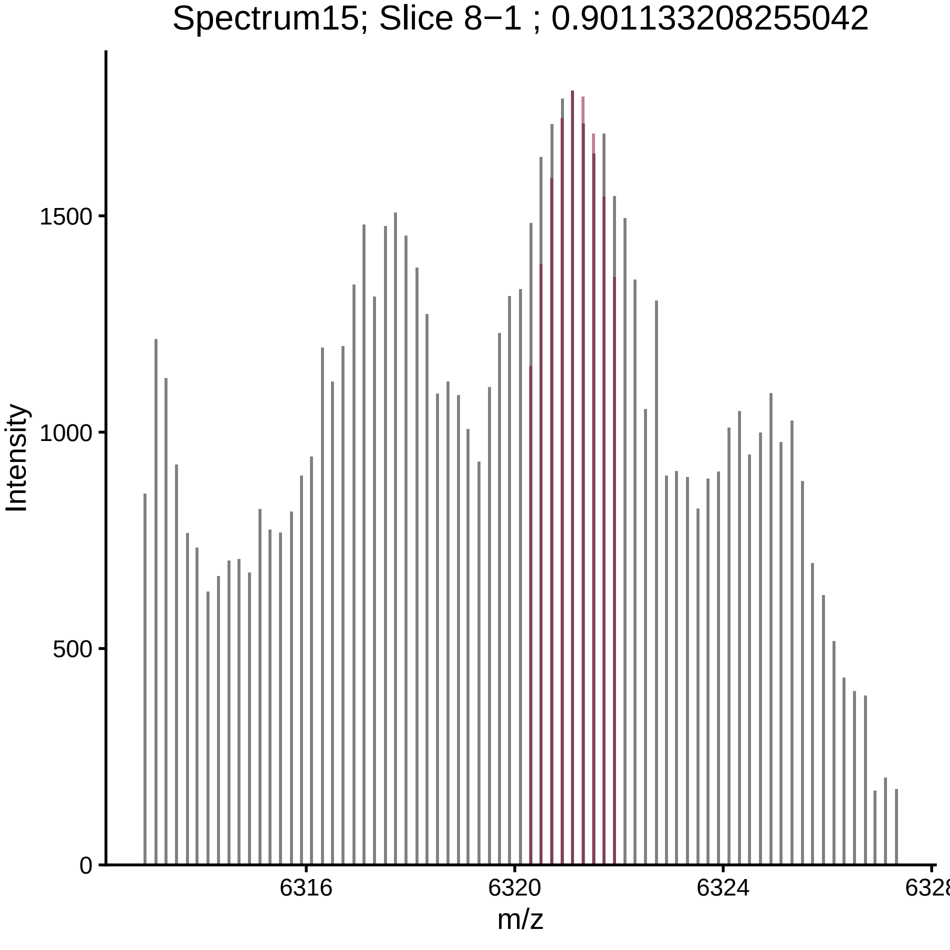

Spectrum15; Slice 8−1 ; 0.901133208255042
Intensity
m/z
0
500
1000
1500
6316
6320
6324
6328

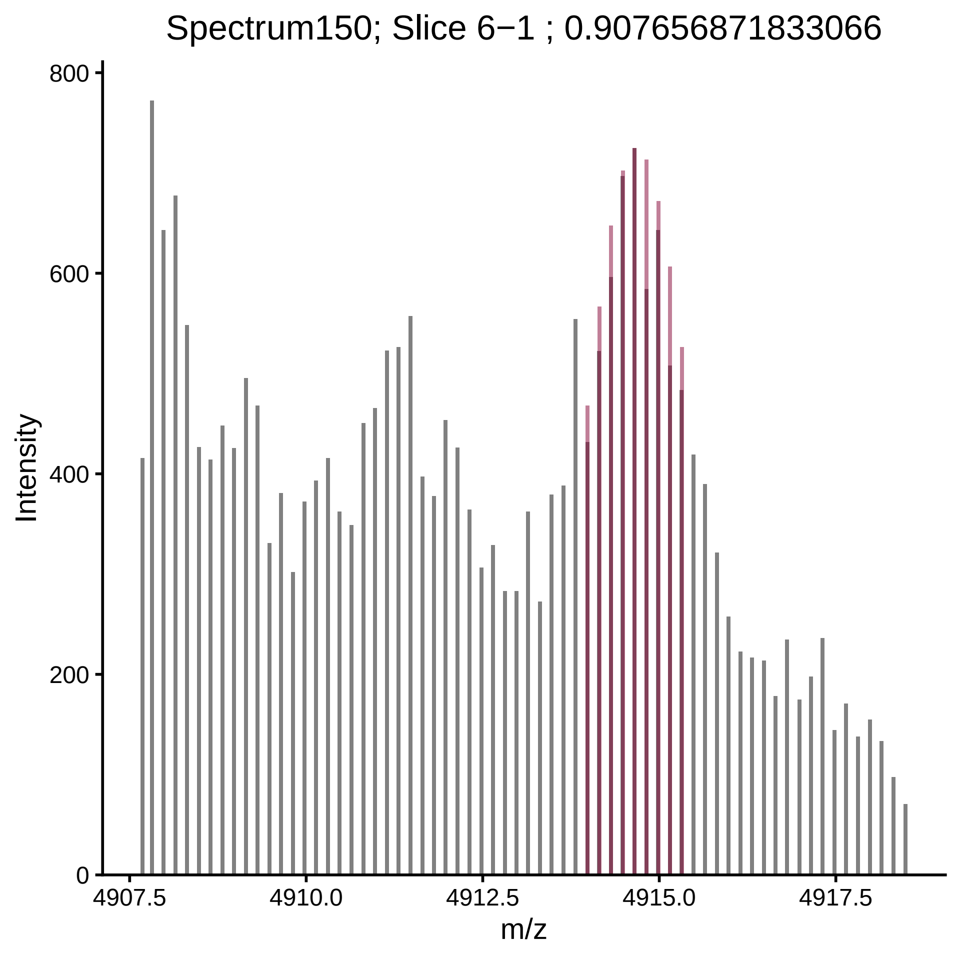

Spectrum150; Slice 6−1 ; 0.907656871833066
Intensity
800
600
400
200
0
4907.5
4910.0
4912.5
4915.0
4917.5
m/z

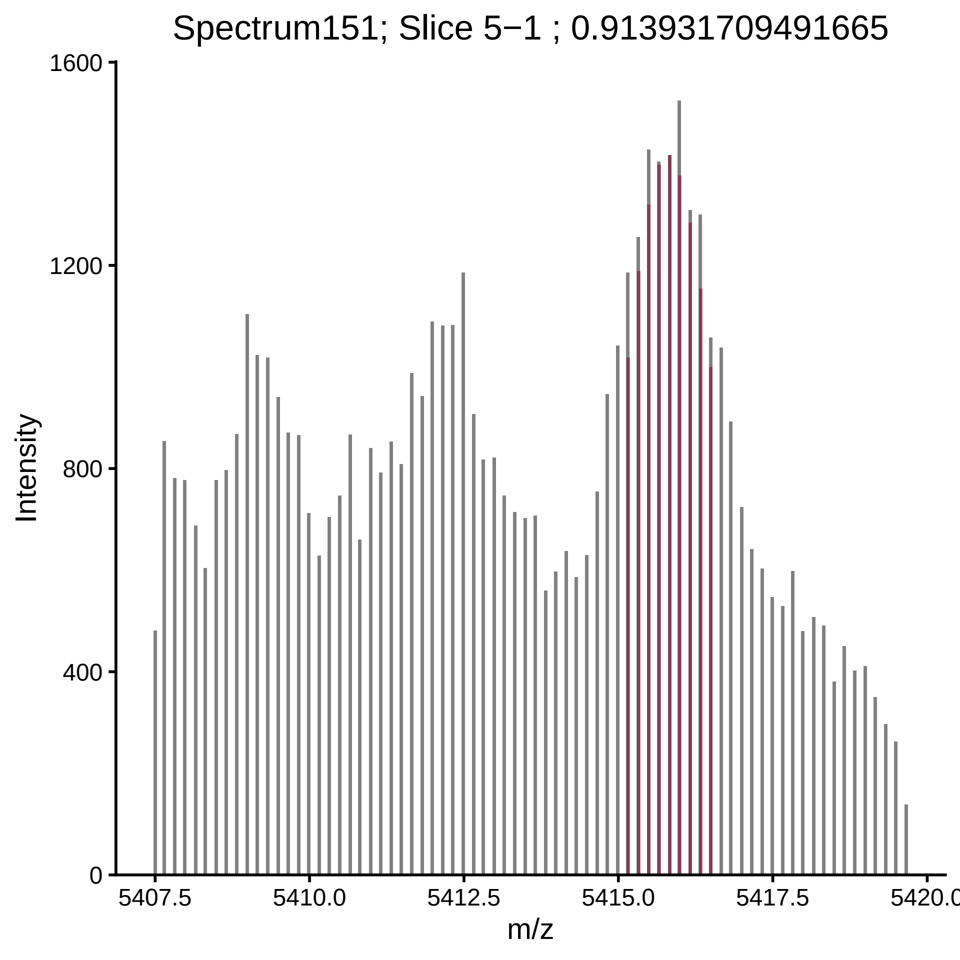

Spectrum151; Slice 5−1 ; 0.913931709491665
Intensity
1600
1200
800
400
0
m/z
5407.5
5410.0
5412.5
5415.0
5417.5
5420.0

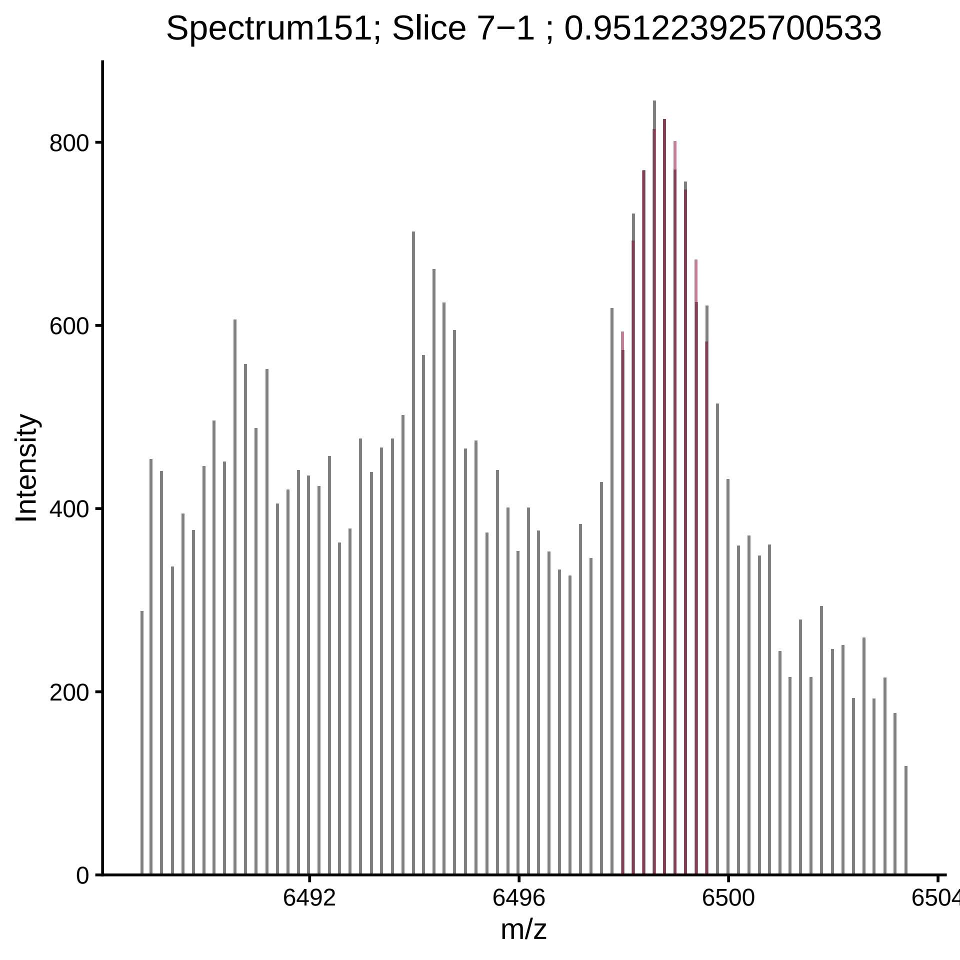

Spectrum151; Slice 7−1 ; 0.951223925700533
Intensity
m/z
0
200
400
600
800
6492
6496
6500
6504

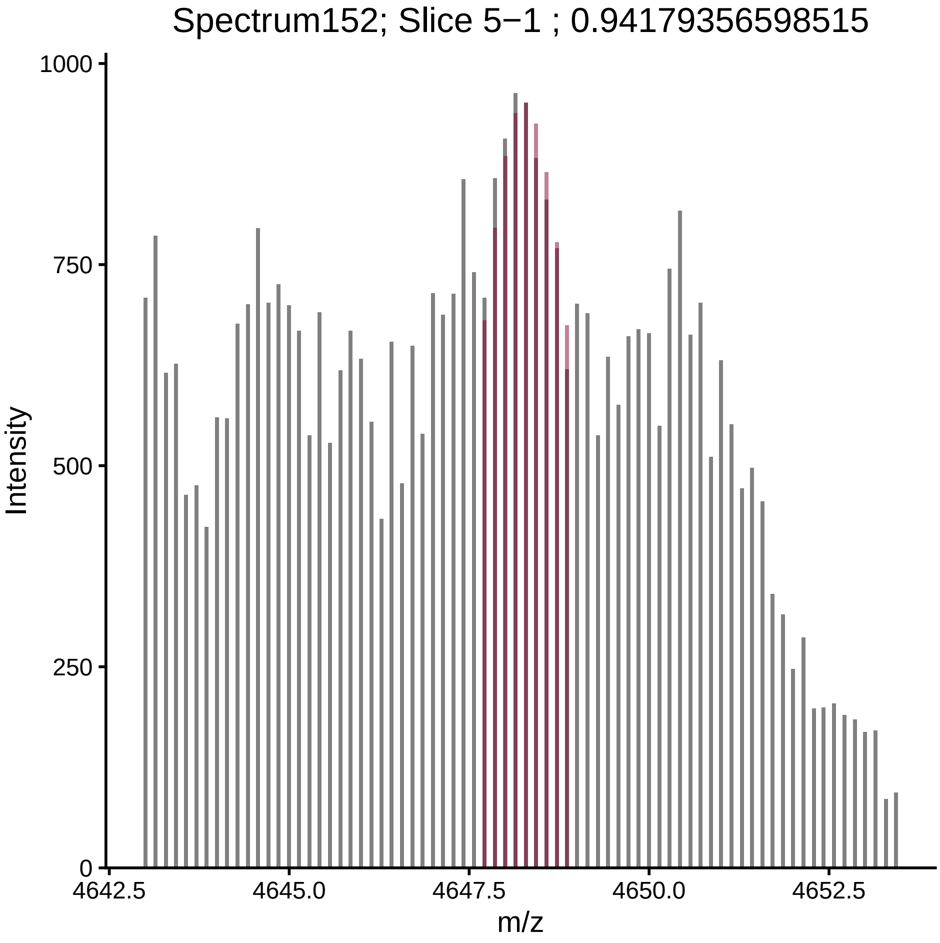
Spectrum152; Slice 5−1 ; 0.94179356598515
Intensity
m/z
1000
750
500
250
0
4642.5
4645.0
4647.5
4650.0
4652.5

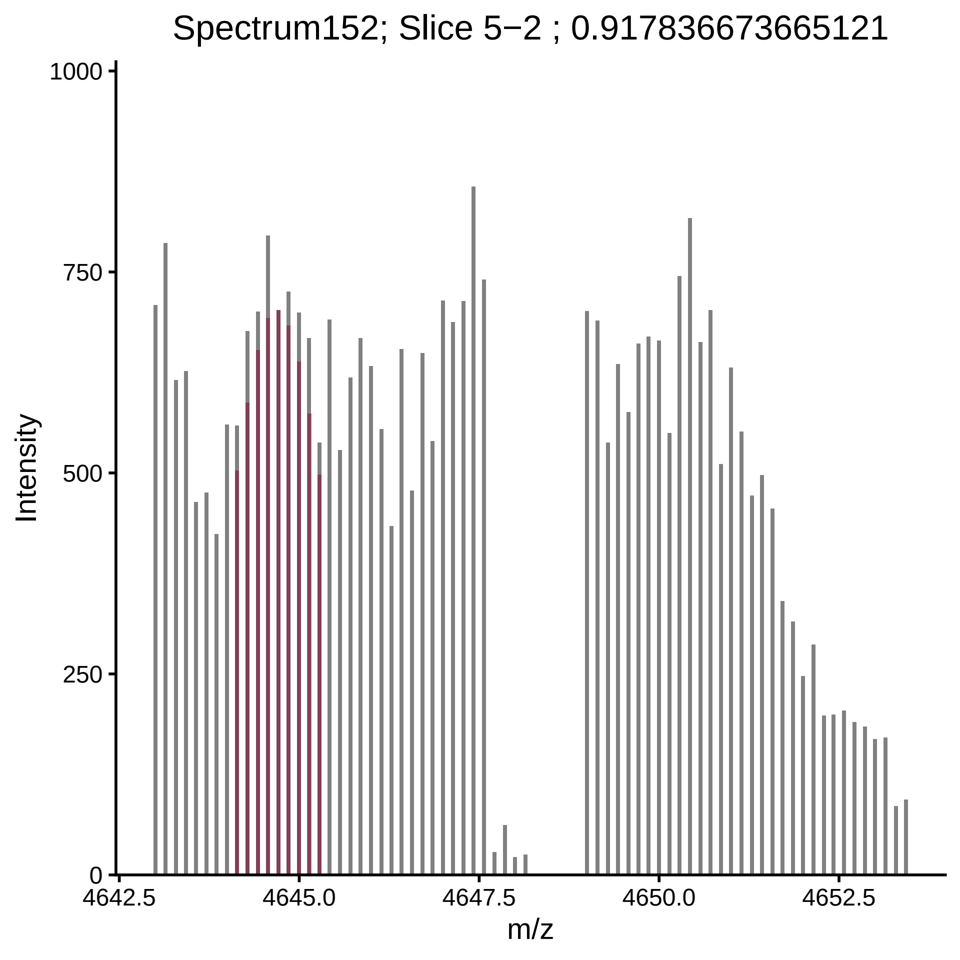
Spectrum152; Slice 5−2 ; 0.917836673665121
Intensity
1000
750
500
250
0
4642.5
4645.0
4647.5
4650.0
4652.5
m/z

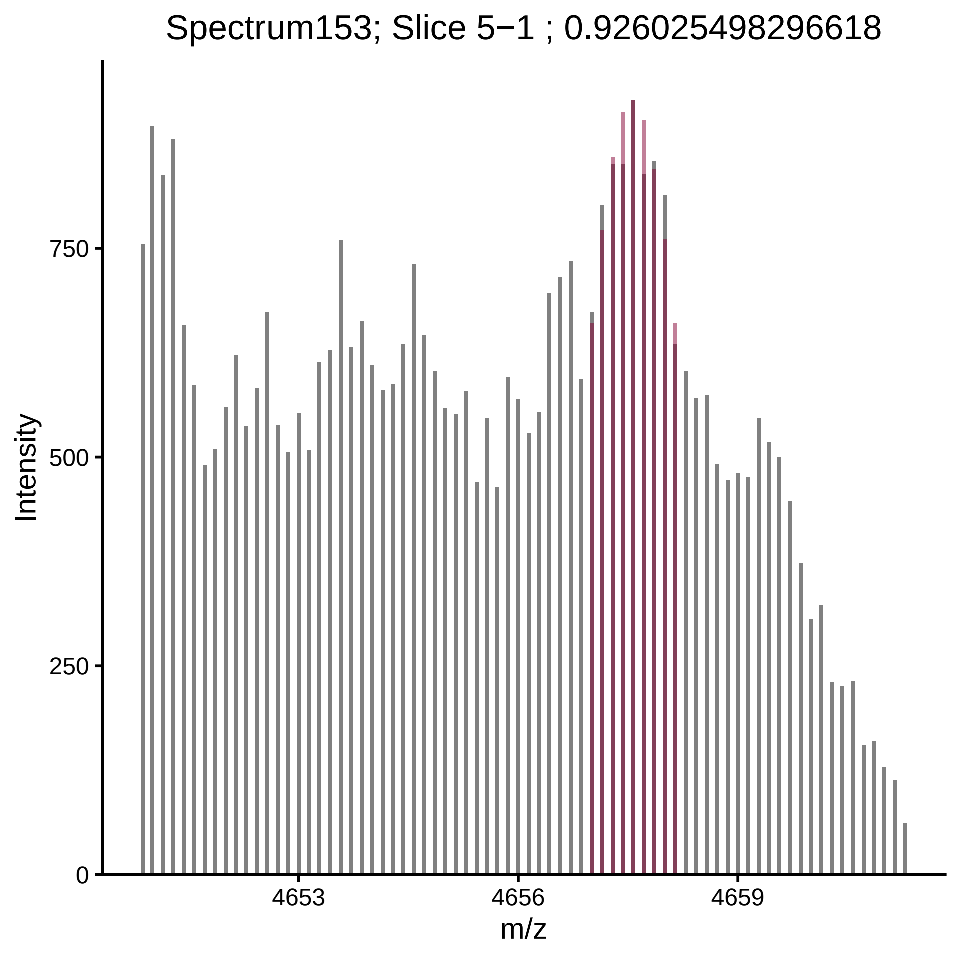

Spectrum153; Slice 5−1 ; 0.926025498296618
Intensity
750
500
250
0
4653
4656
4659
m/z

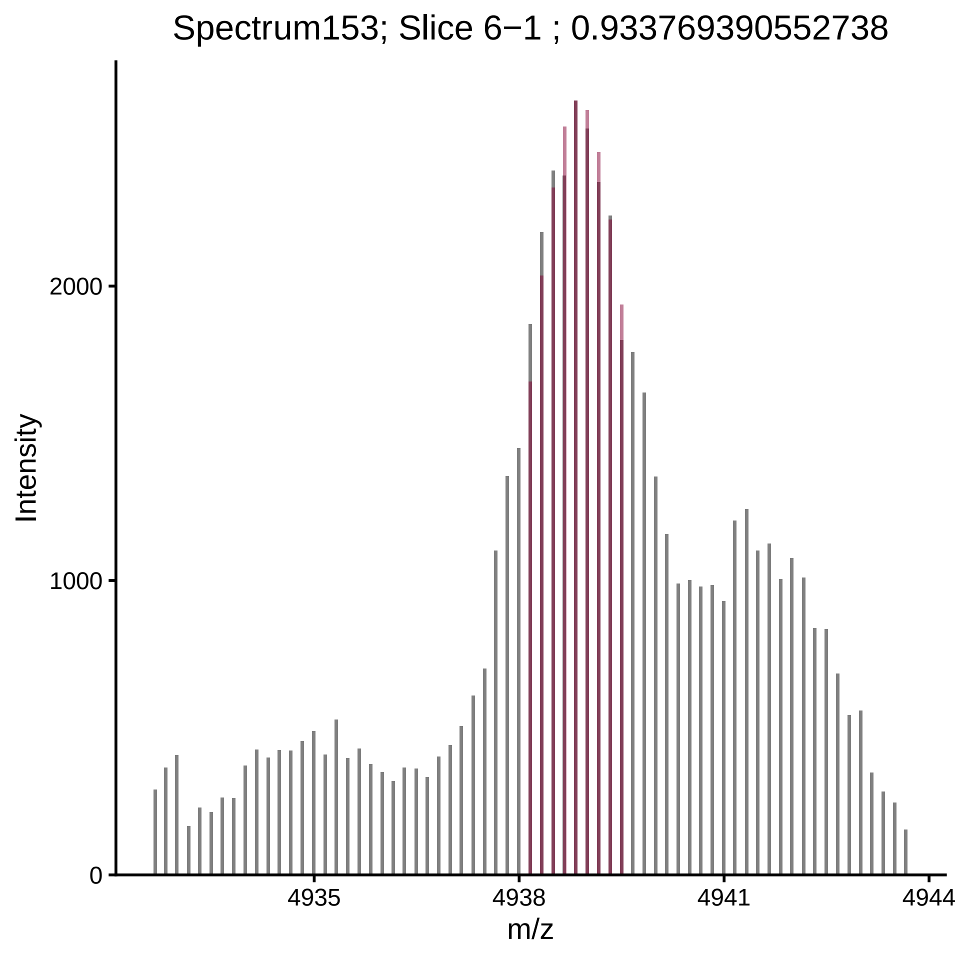

Spectrum153; Slice 6−1 ; 0.933769390552738
Intensity
2000
1000
0
4935
4938
4941
4944
m/z

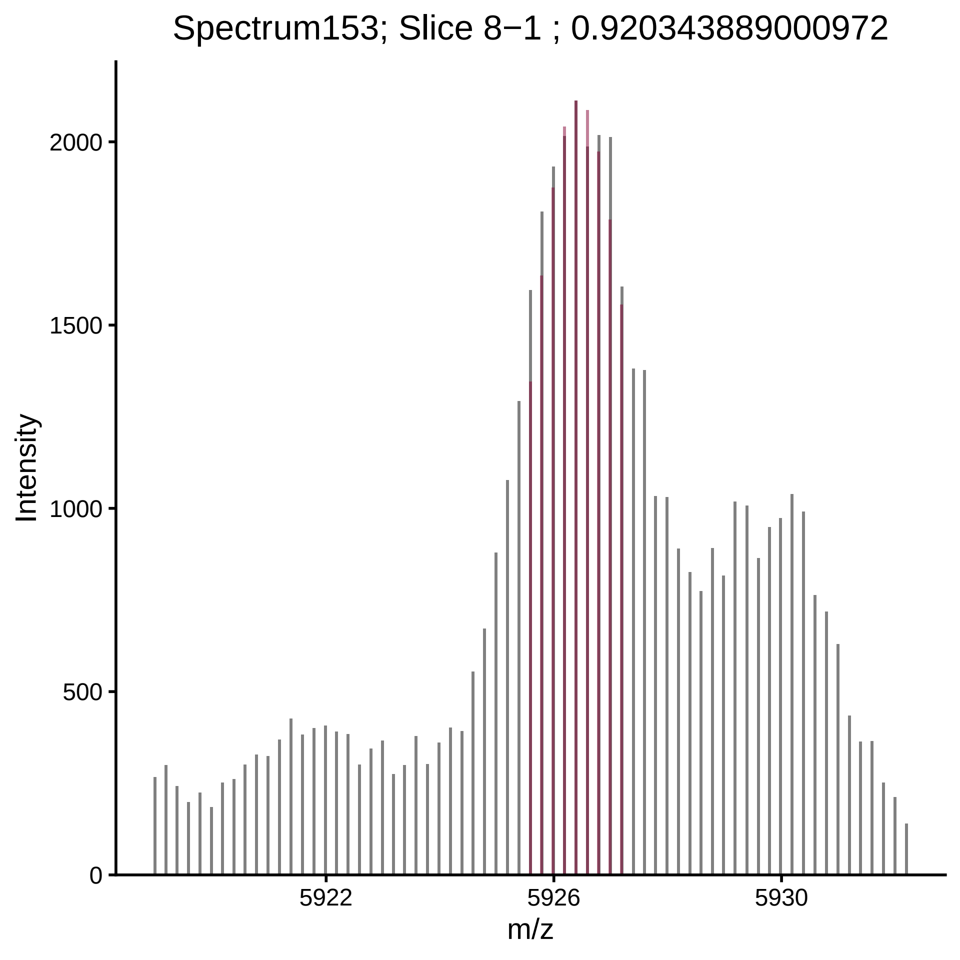
Spectrum153; Slice 8−1 ; 0.920343889000972
Intensity
0
500
1000
1500
2000
5922
5926
5930
m/z

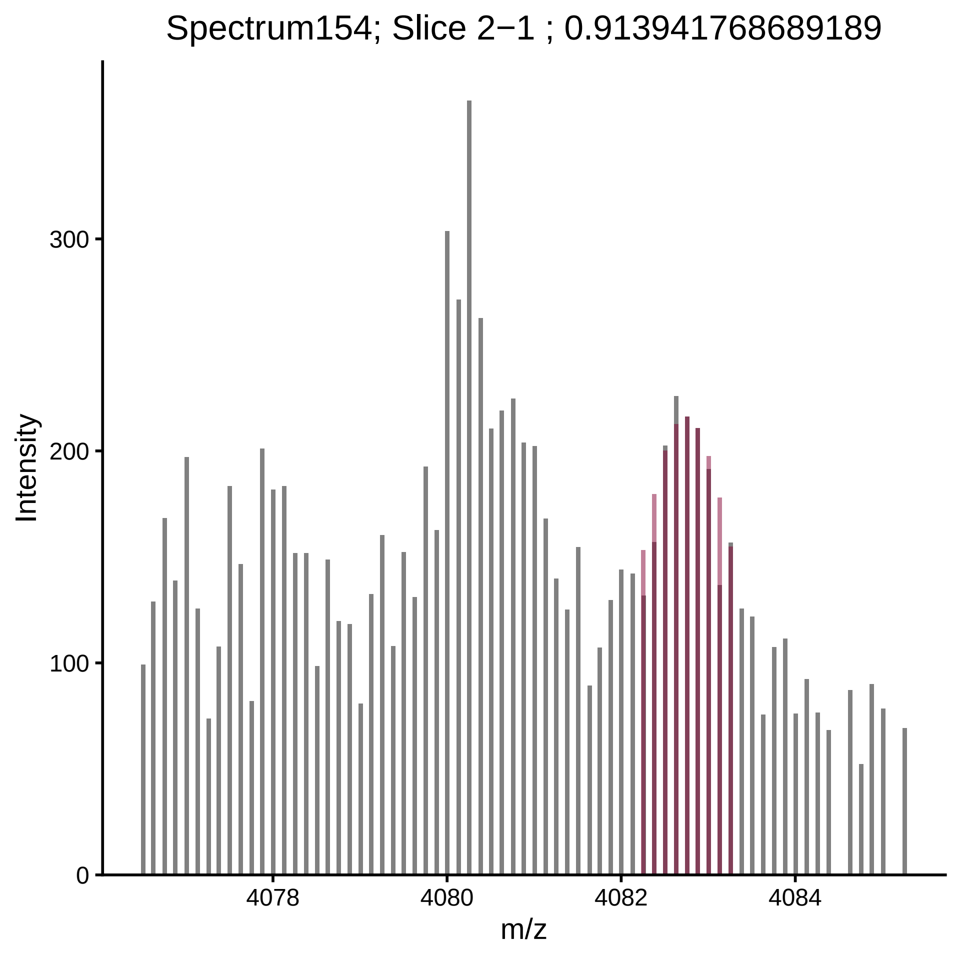

Spectrum154; Slice 2−1 ; 0.913941768689189
Intensity
300
200
100
0
4078
4080
4082
4084
m/z

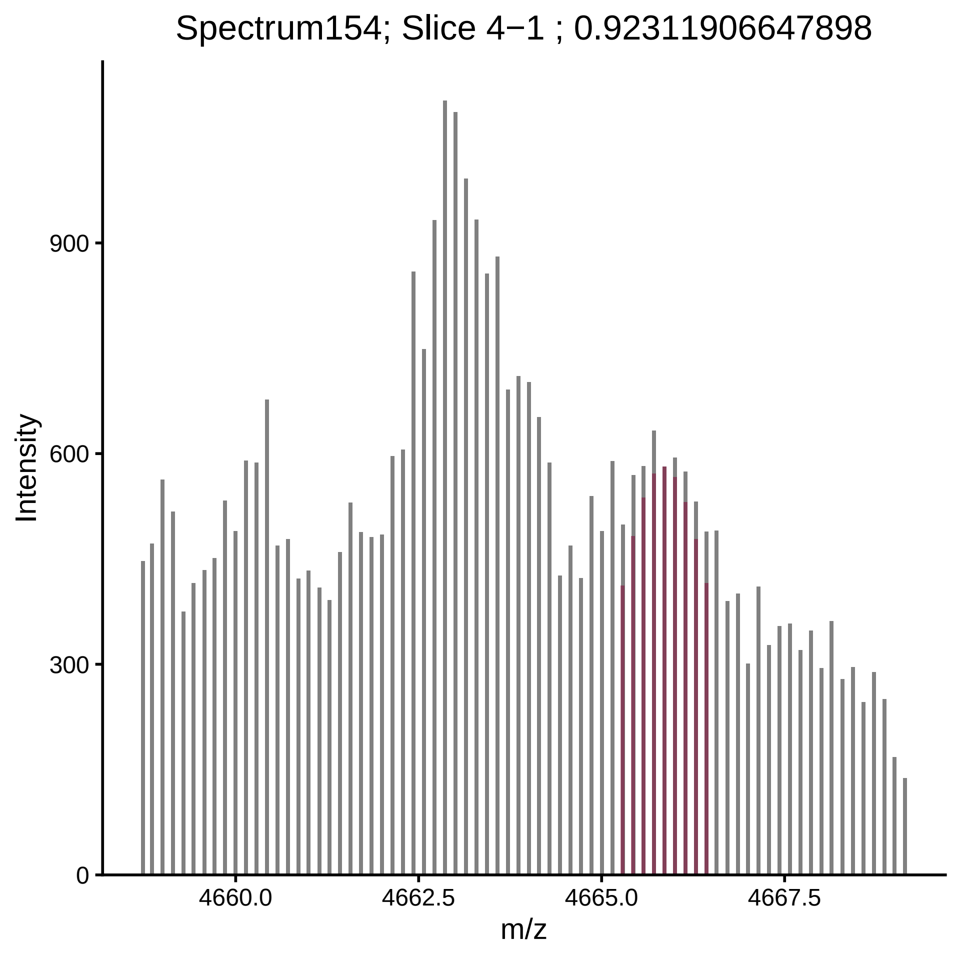
Spectrum154; Slice 4−1 ; 0.92311906647898
Intensity
0
300
600
900
4660.0
4662.5
4665.0
4667.5
m/z

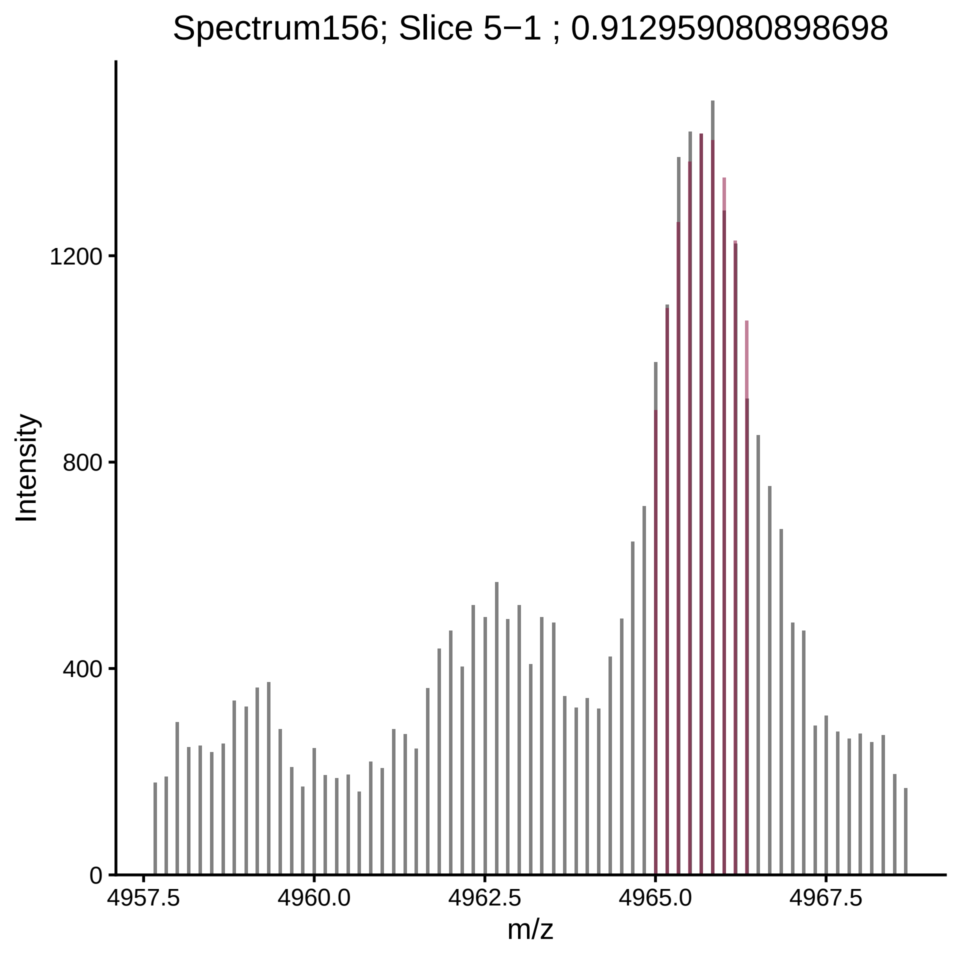
Spectrum156; Slice 5−1 ; 0.912959080898698
Intensity
0
400
800
1200
4957.5
4960.0
4962.5
4965.0
4967.5
m/z

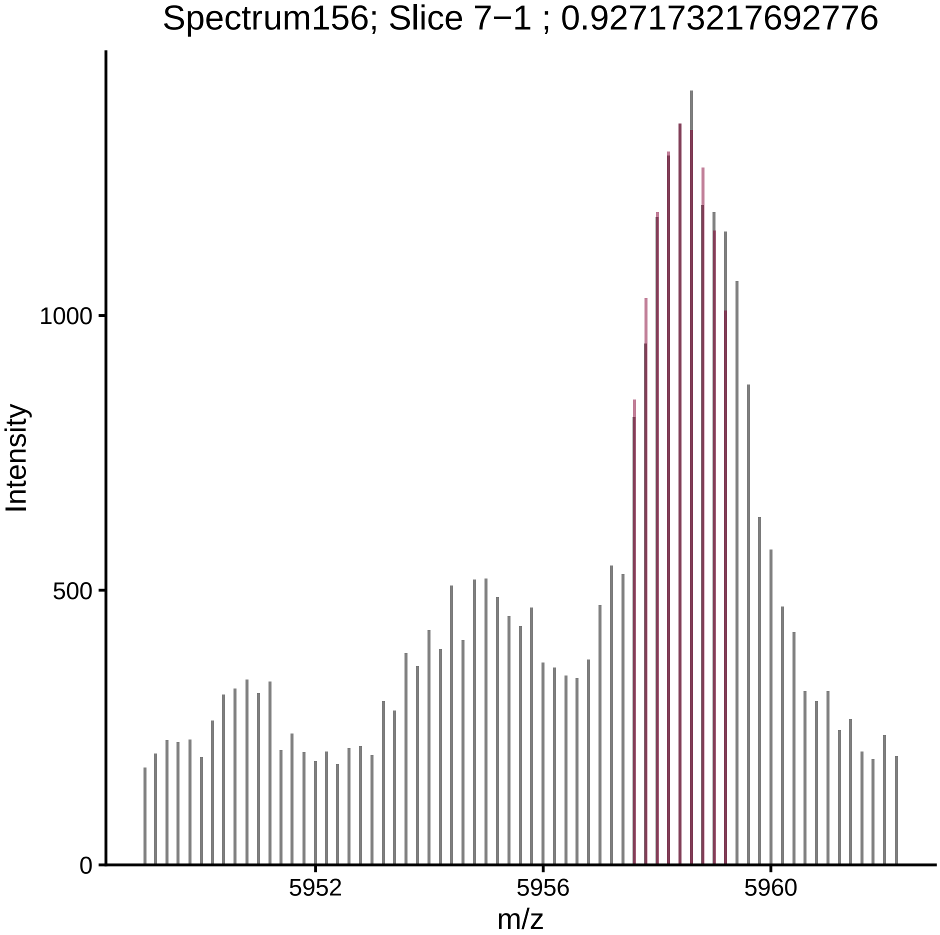
Spectrum156; Slice 7−1 ; 0.927173217692776
Intensity
1000
500
0
5952
5956
5960
m/z

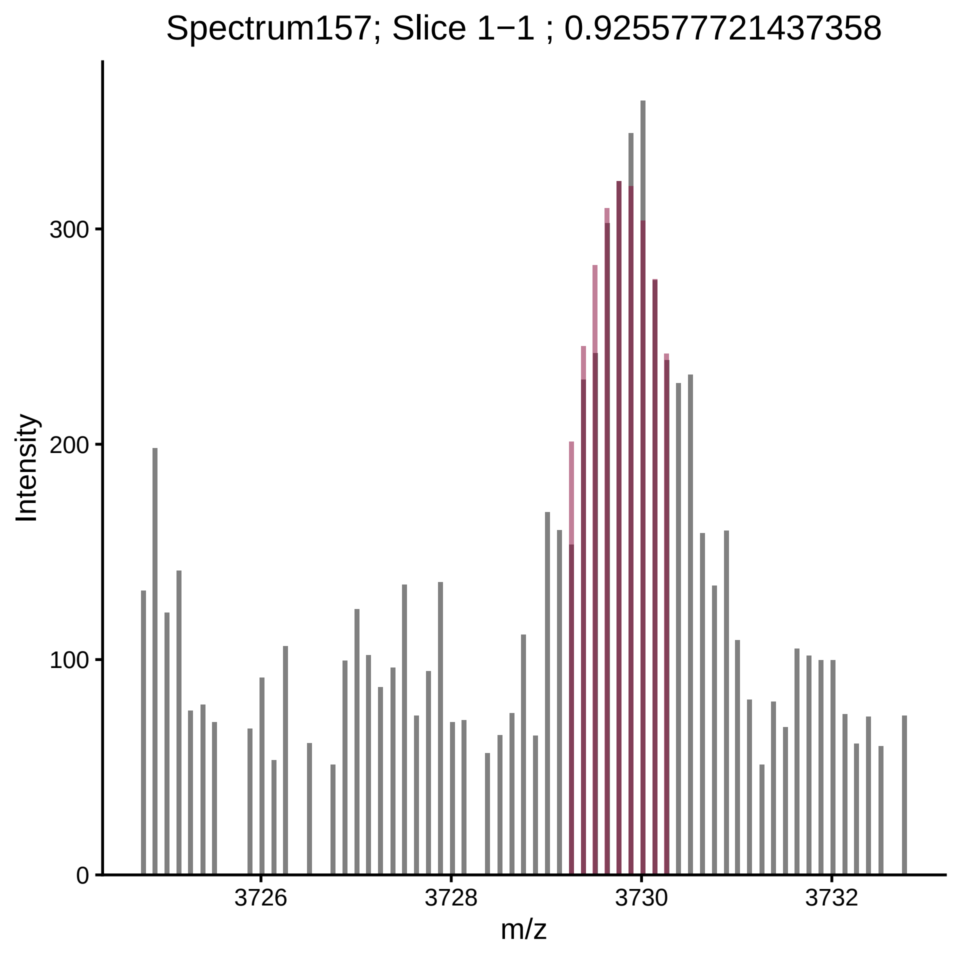
Spectrum157; Slice 1−1 ; 0.925577721437358
Intensity
0
100
200
300
3726
3728
3730
3732
m/z

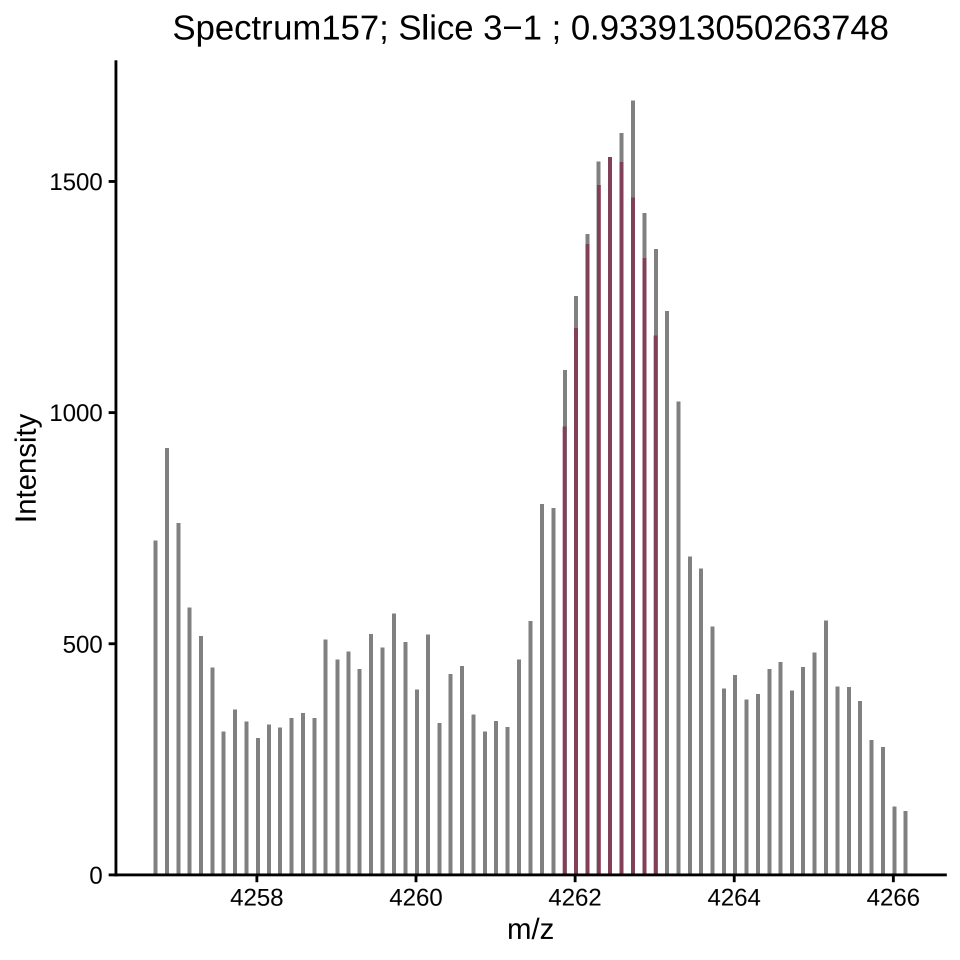

Spectrum157; Slice 3−1 ; 0.933913050263748
Intensity
0
500
1000
1500
4258
4260
4262
4264
4266
m/z

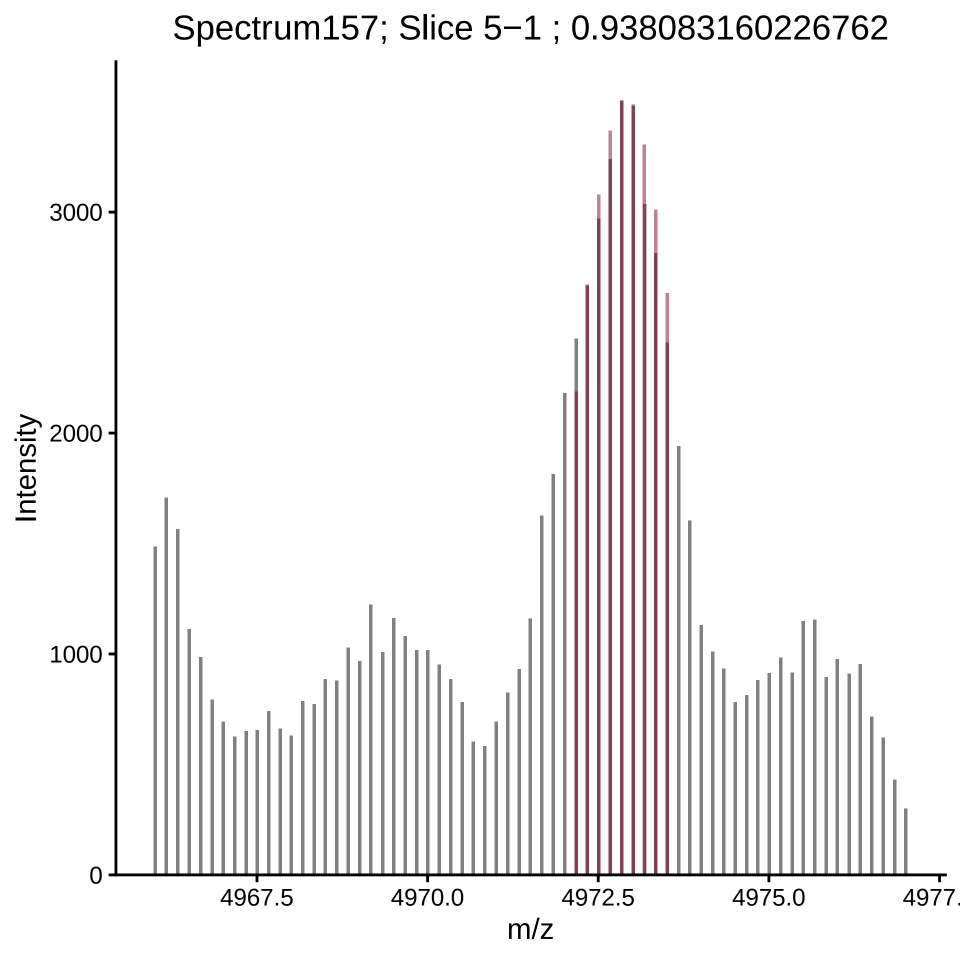

Spectrum157; Slice 5−1 ; 0.938083160226762
Intensity
3000
2000
1000
0
4967.5
4970.0
4972.5
4975.0
4977.
m/z

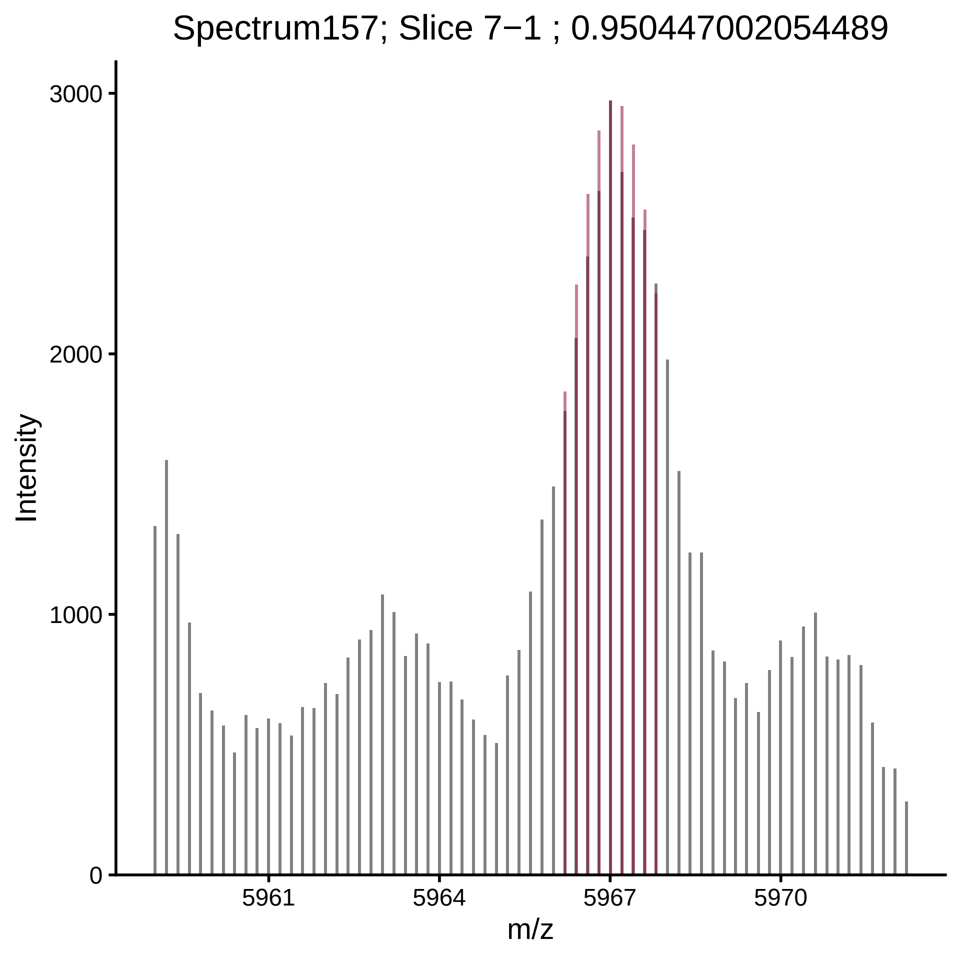
Spectrum157; Slice 7−1 ; 0.950447002054489
Intensity
3000
2000
1000
0
5961
5964
5967
5970
m/z

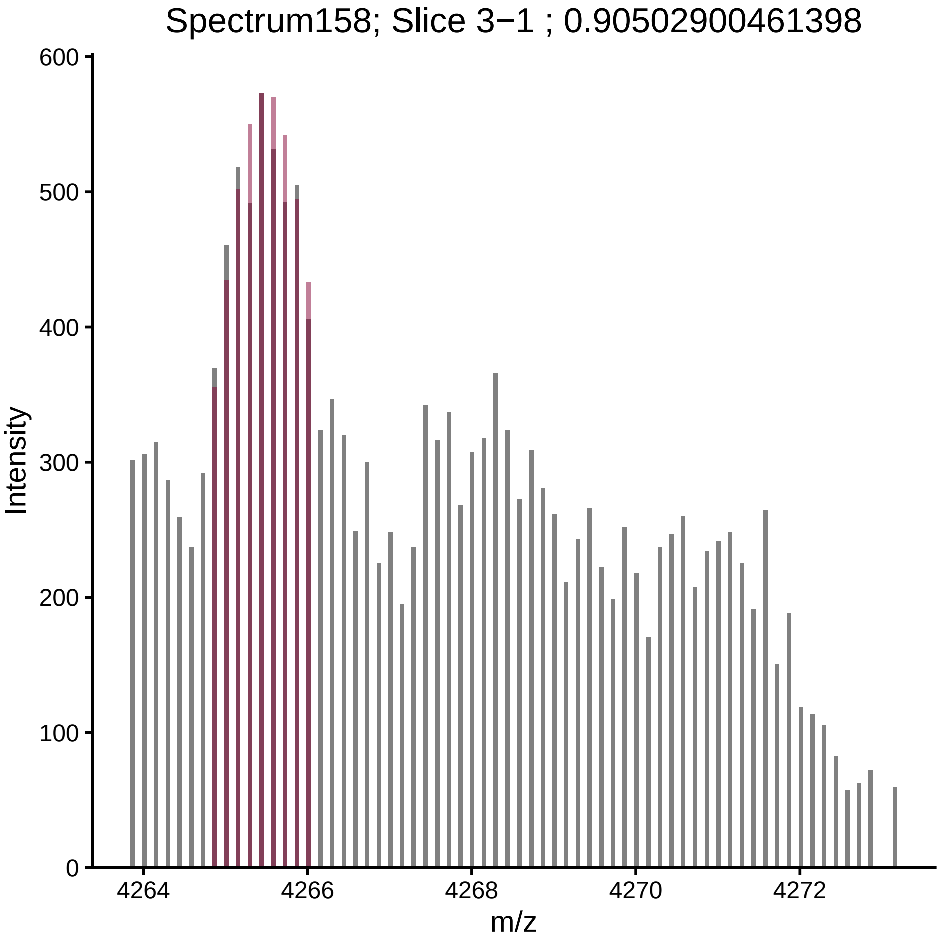

Spectrum158; Slice 3−1 ; 0.90502900461398
Intensity
m/z
0
100
200
300
400
500
600
4264
4266
4268
4270
4272

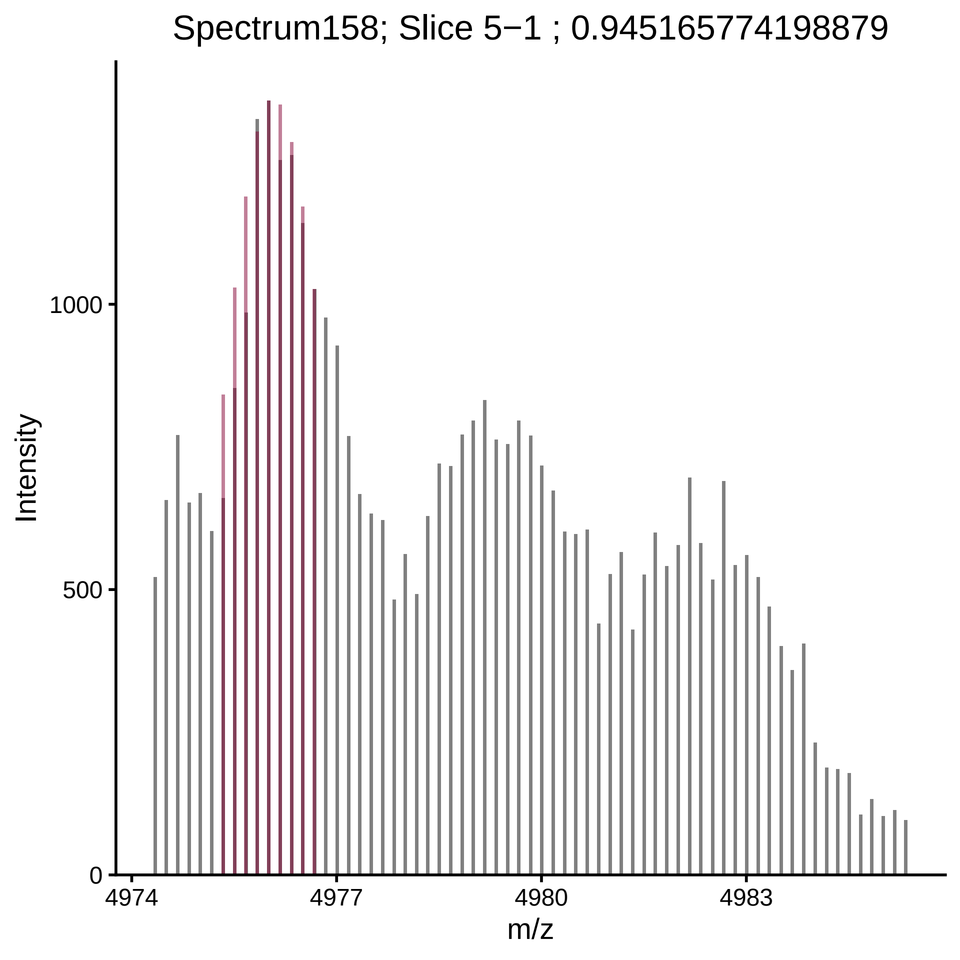
Spectrum158; Slice 5−1 ; 0.945165774198879
Intensity
1000
500
0
4974
4977
4980
4983
m/z

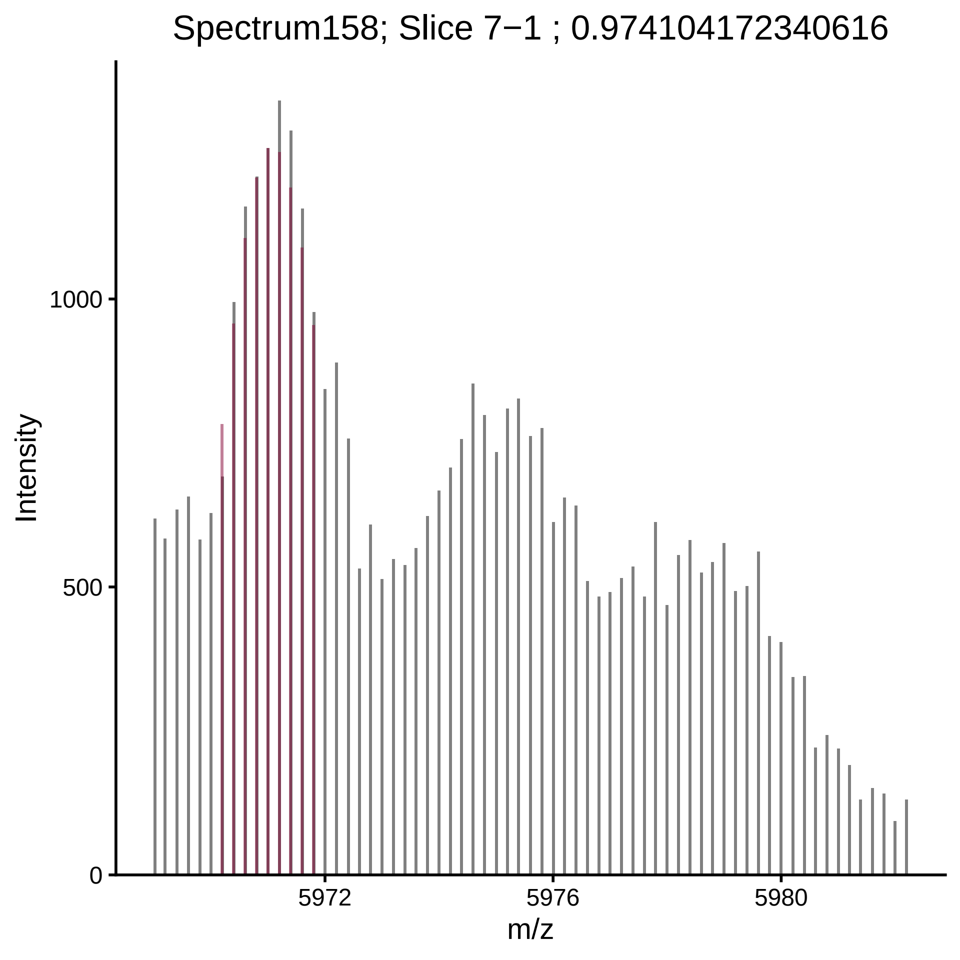
Spectrum158; Slice 7−1 ; 0.974104172340616
Intensity
1000
500
0
5972
5976
5980
m/z

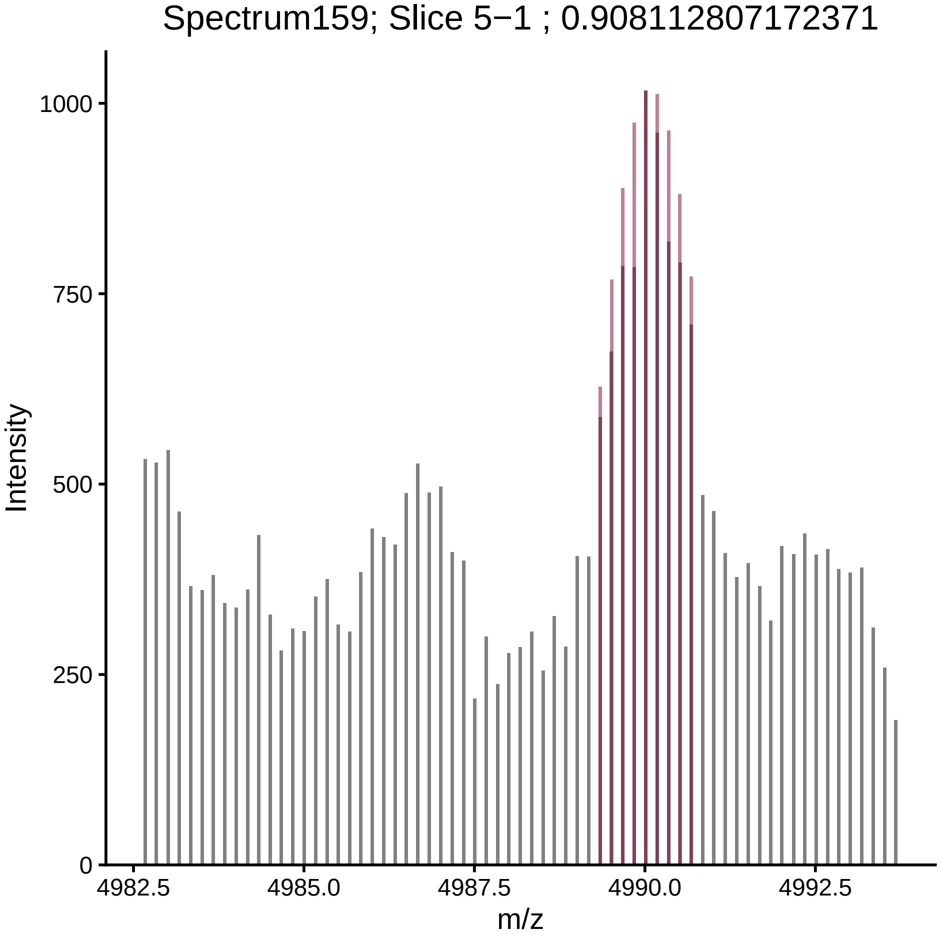

Spectrum159; Slice 5−1 ; 0.908112807172371
Intensity
0
250
500
750
1000
4982.5
4985.0
4987.5
4990.0
4992.5
m/z

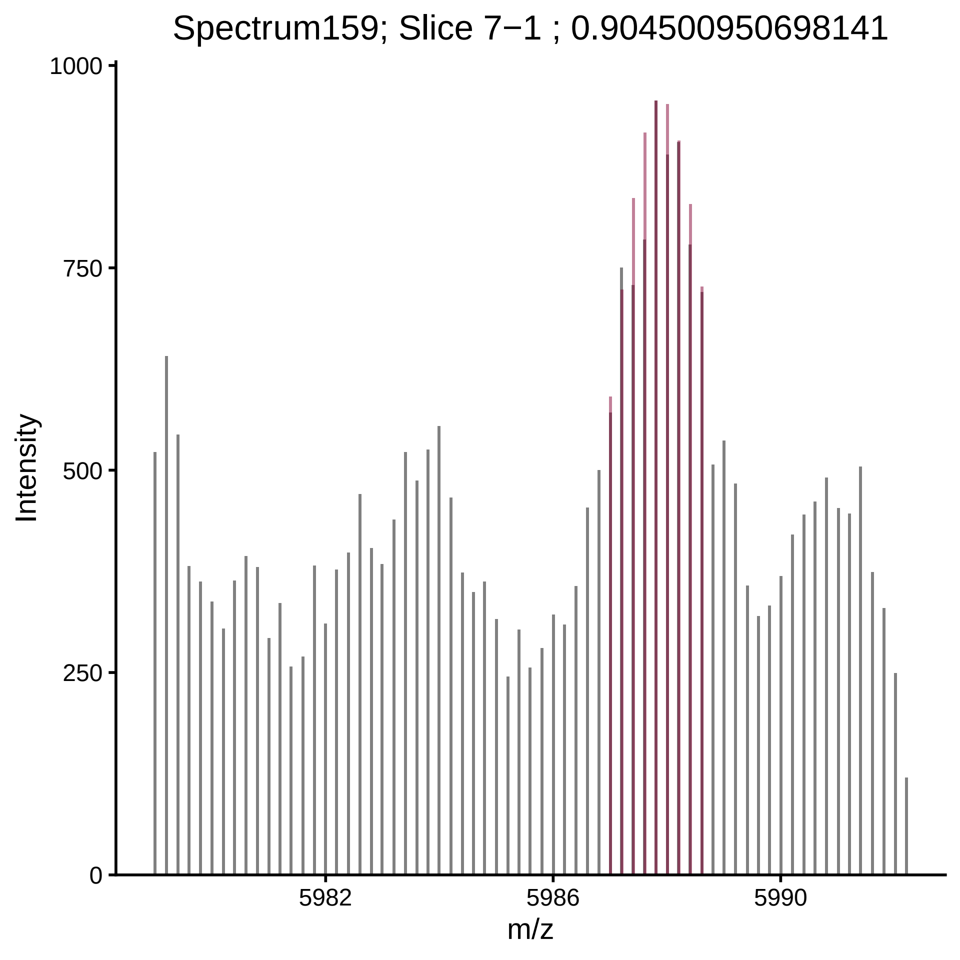

Spectrum159; Slice 7−1 ; 0.904500950698141
Intensity
1000
750
500
250
0
5982
5986
5990
m/z

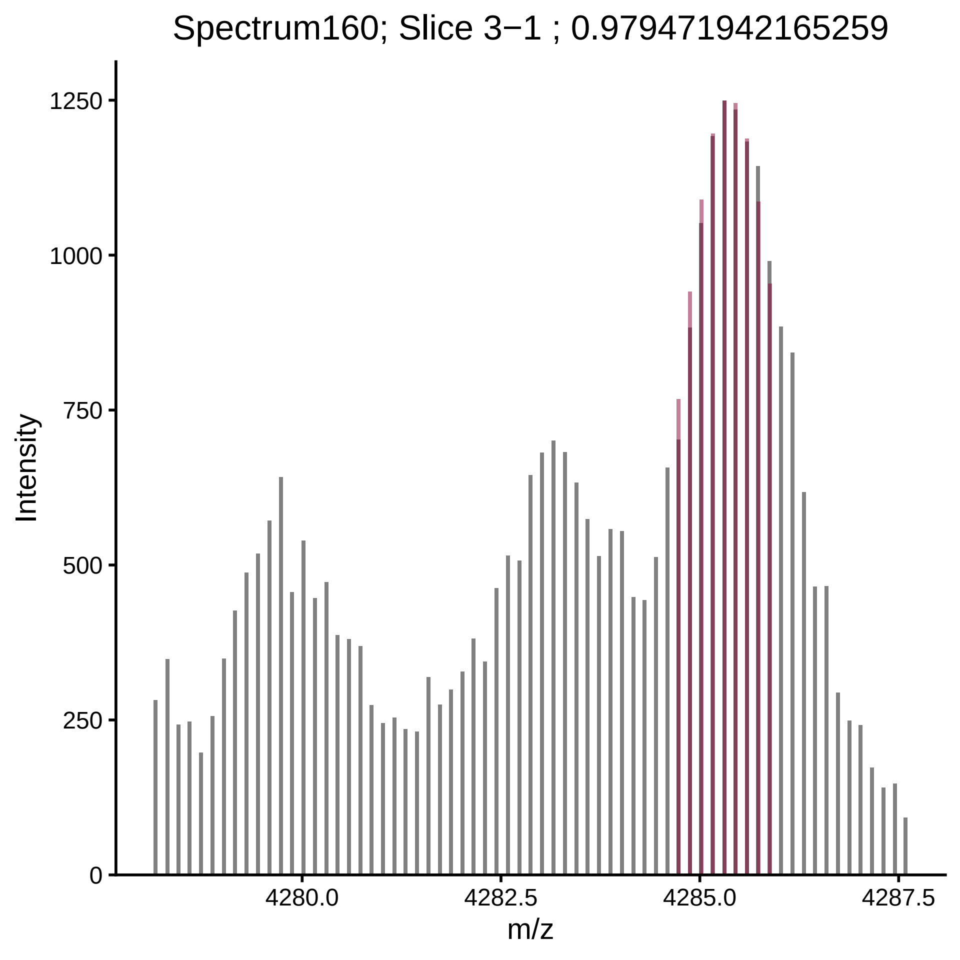

Spectrum160; Slice 3−1 ; 0.979471942165259
Intensity
1250
1000
750
500
250
0
4280.0
4282.5
4285.0
4287.5
m/z

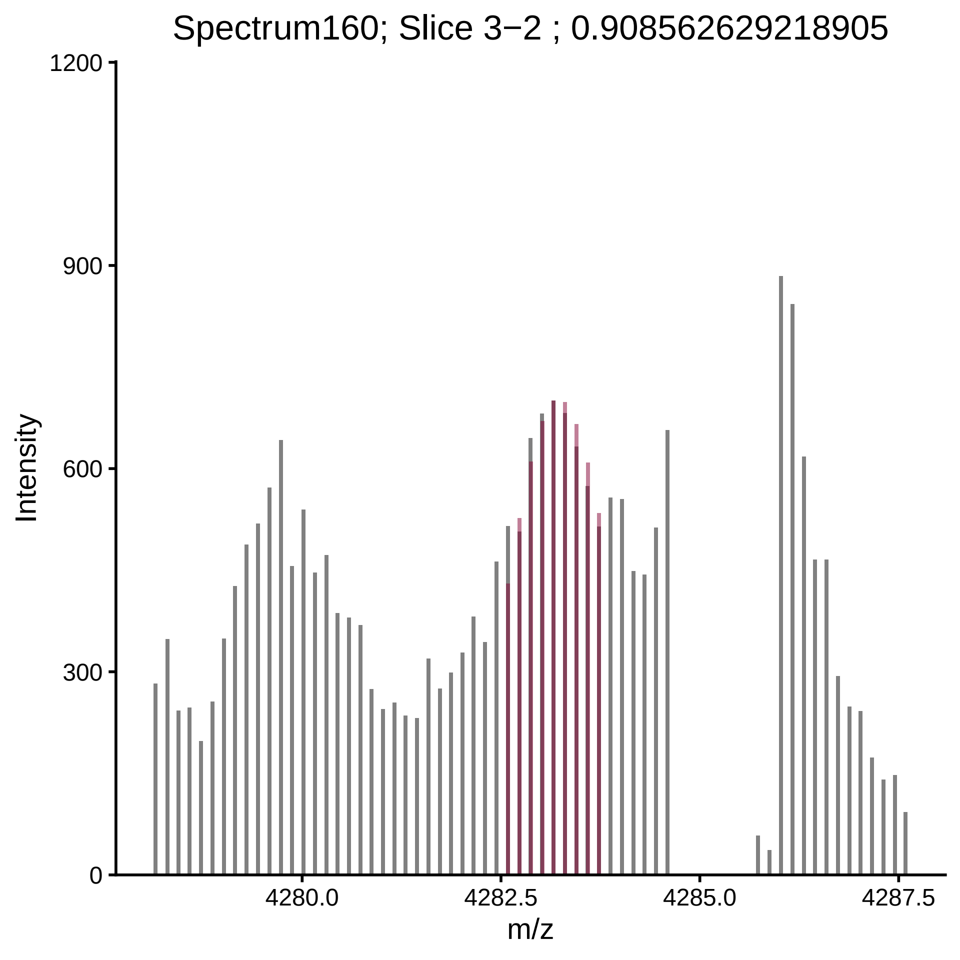

Spectrum160; Slice 3−2 ; 0.908562629218905
Intensity
1200
900
600
300
0
4280.0
4282.5
4285.0
4287.5
m/z

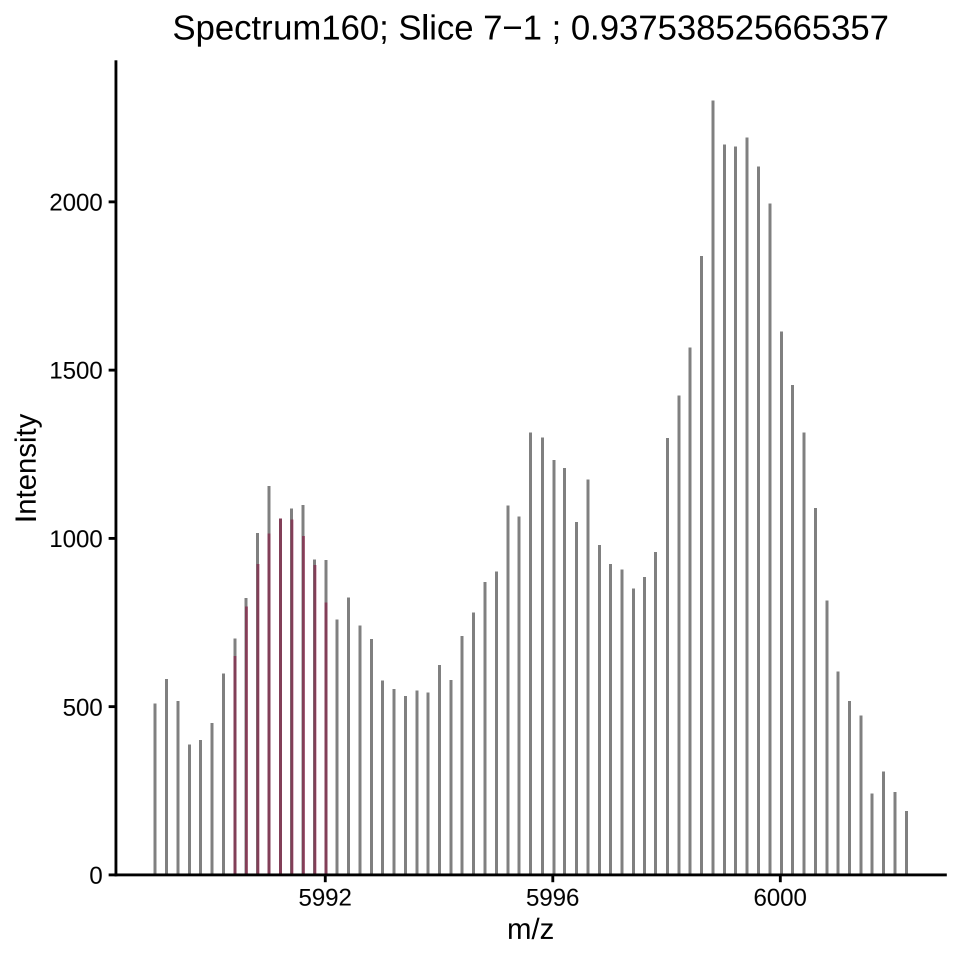
Spectrum160; Slice 7−1 ; 0.937538525665357
Intensity
2000
1500
1000
500
0
5992
5996
6000
m/z

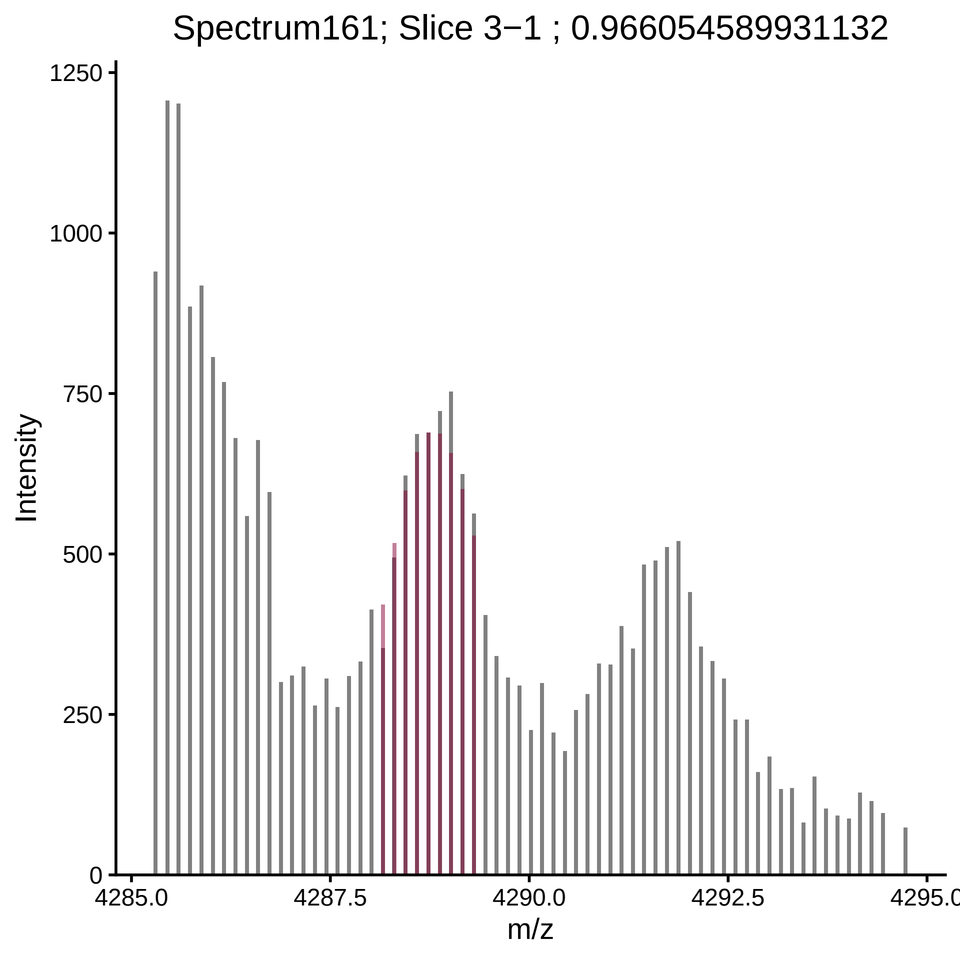

Spectrum161; Slice 3−1 ; 0.966054589931132
Intensity
1250
1000
750
500
250
0
4285.0
4287.5
4290.0
4292.5
4295.0
m/z

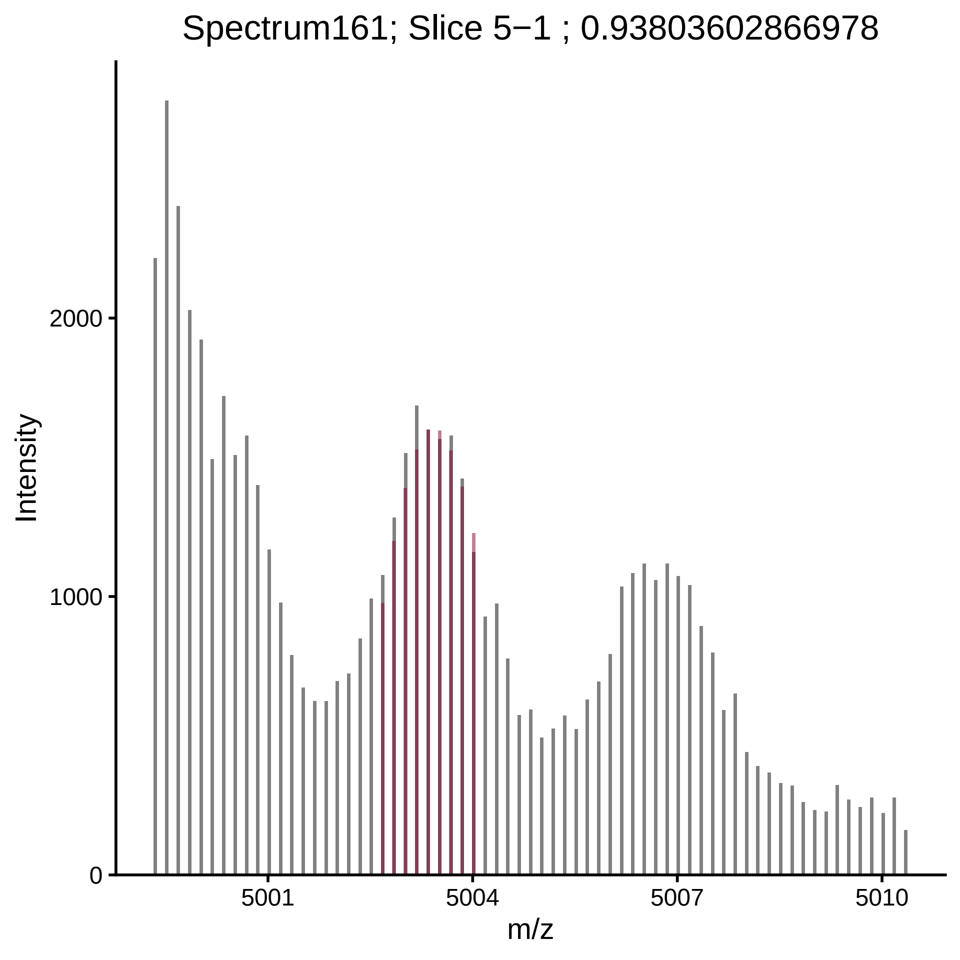

Spectrum161; Slice 5−1 ; 0.93803602866978
Intensity
2000
1000
0
5001
5004
5007
5010
m/z

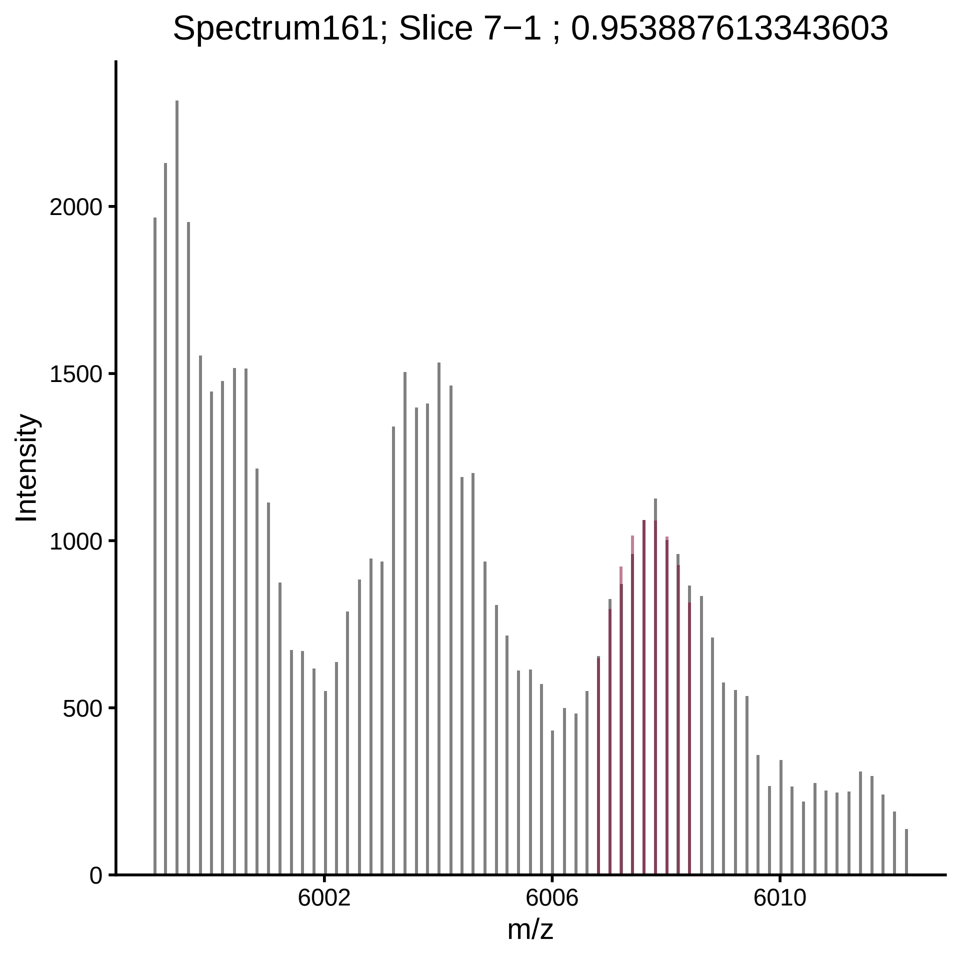
Spectrum161; Slice 7−1 ; 0.953887613343603
Intensity
0
500
1000
1500
2000
6002
6006
6010
m/z

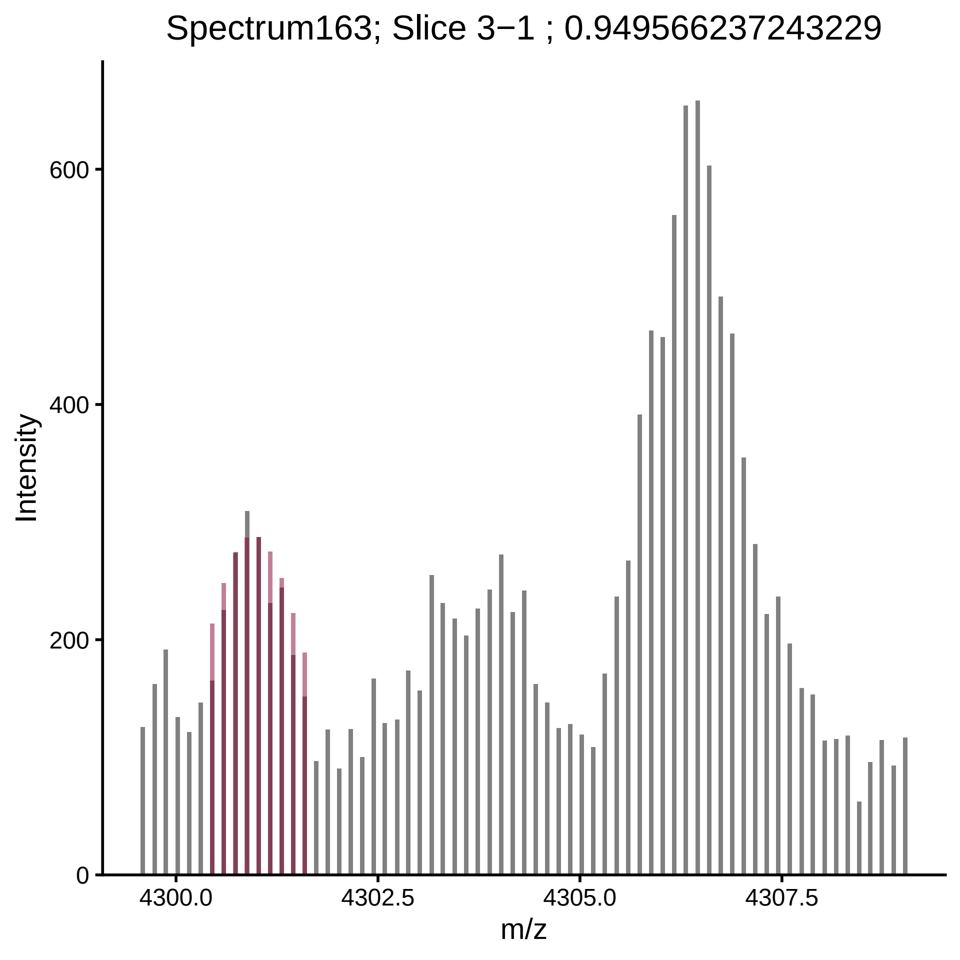

Spectrum163; Slice 3−1 ; 0.949566237243229
Intensity
600
400
200
0
4300.0
4302.5
4305.0
4307.5
m/z

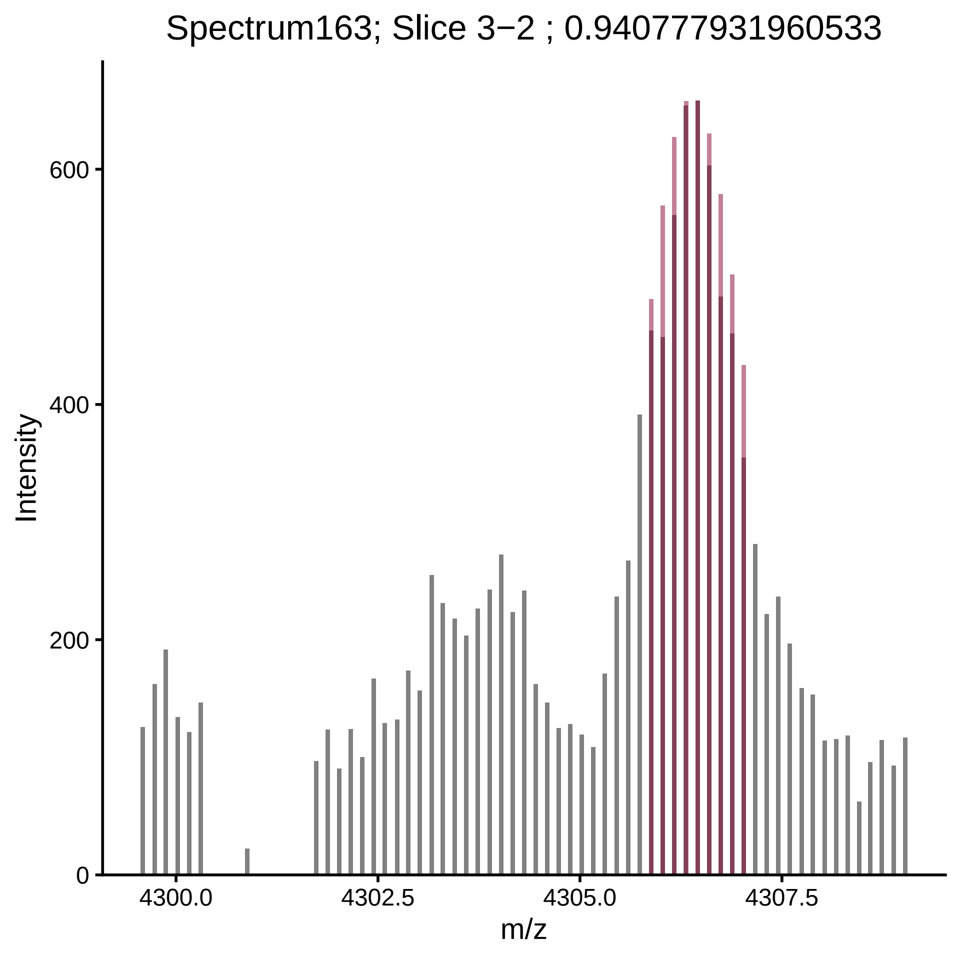
Spectrum163; Slice 3−2 ; 0.940777931960533
Intensity
0
200
400
600
4300.0
4302.5
4305.0
4307.5
m/z

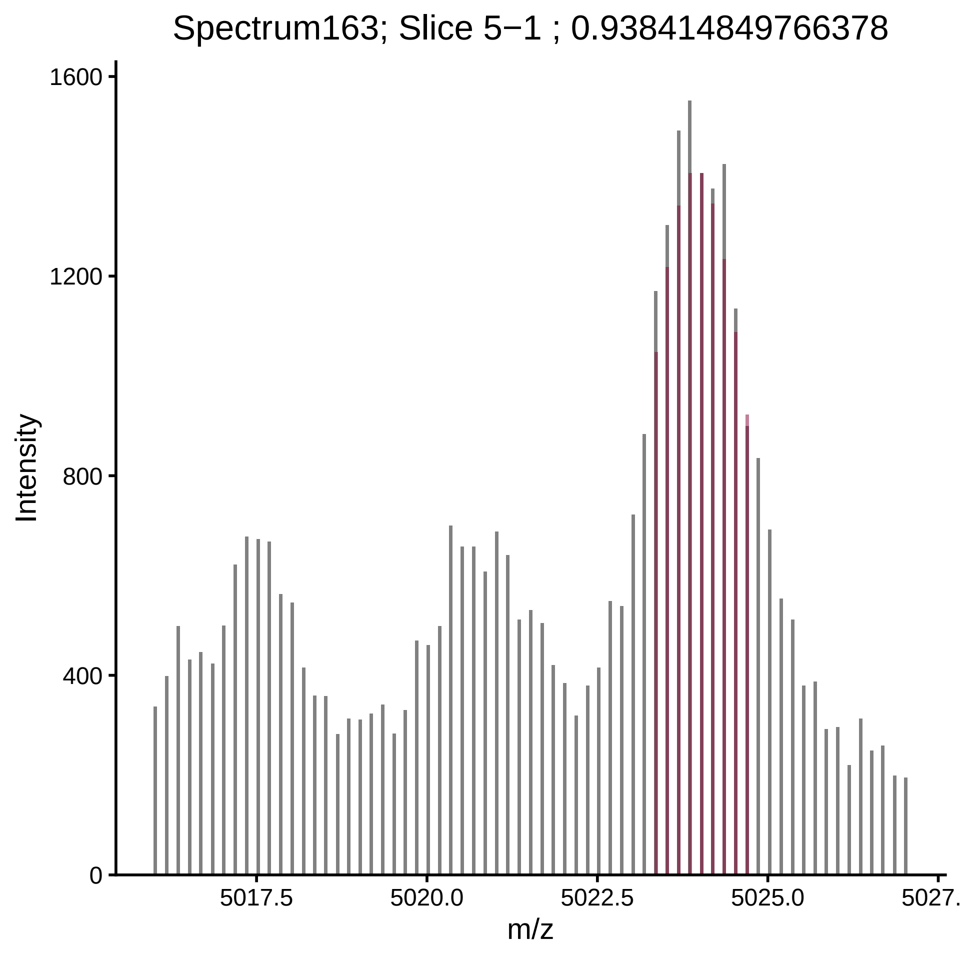

Spectrum163; Slice 5−1 ; 0.938414849766378
Intensity
1600
1200
800
400
0
5017.5
5020.0
5022.5
5025.0
5027.
m/z

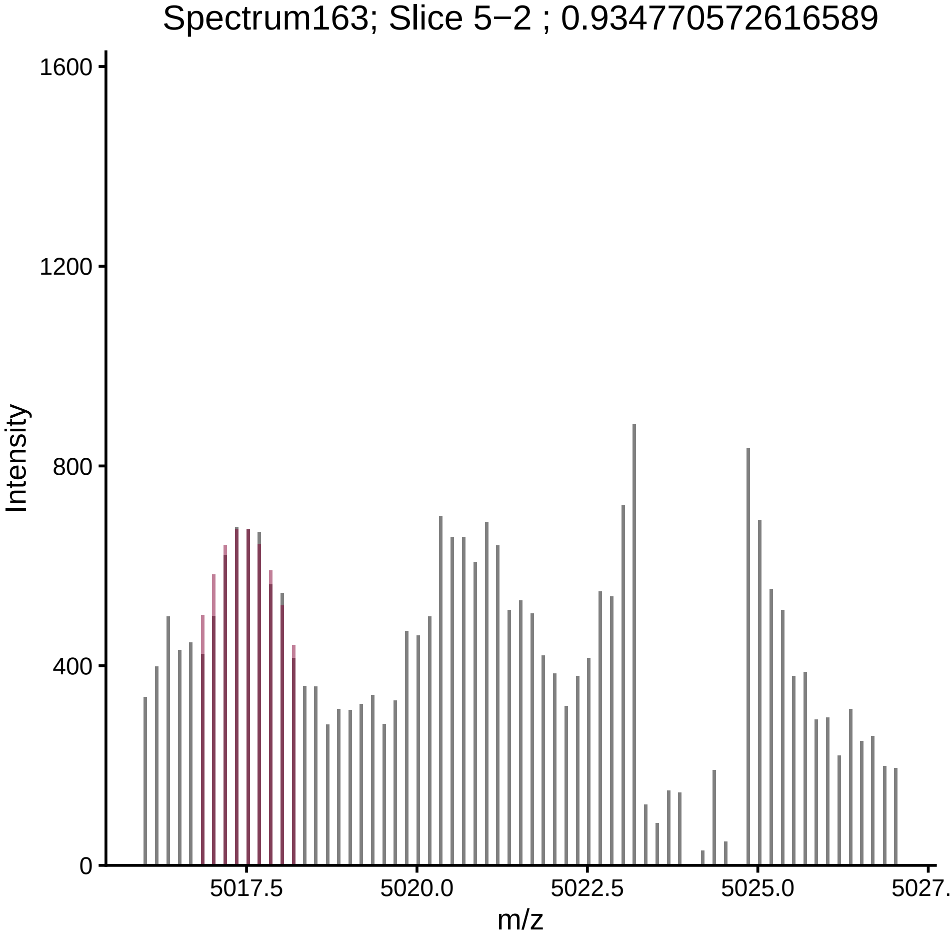

Spectrum163; Slice 5−2 ; 0.934770572616589
Intensity
1600
1200
800
400
0
5017.5
5020.0
5022.5
5025.0
m/z

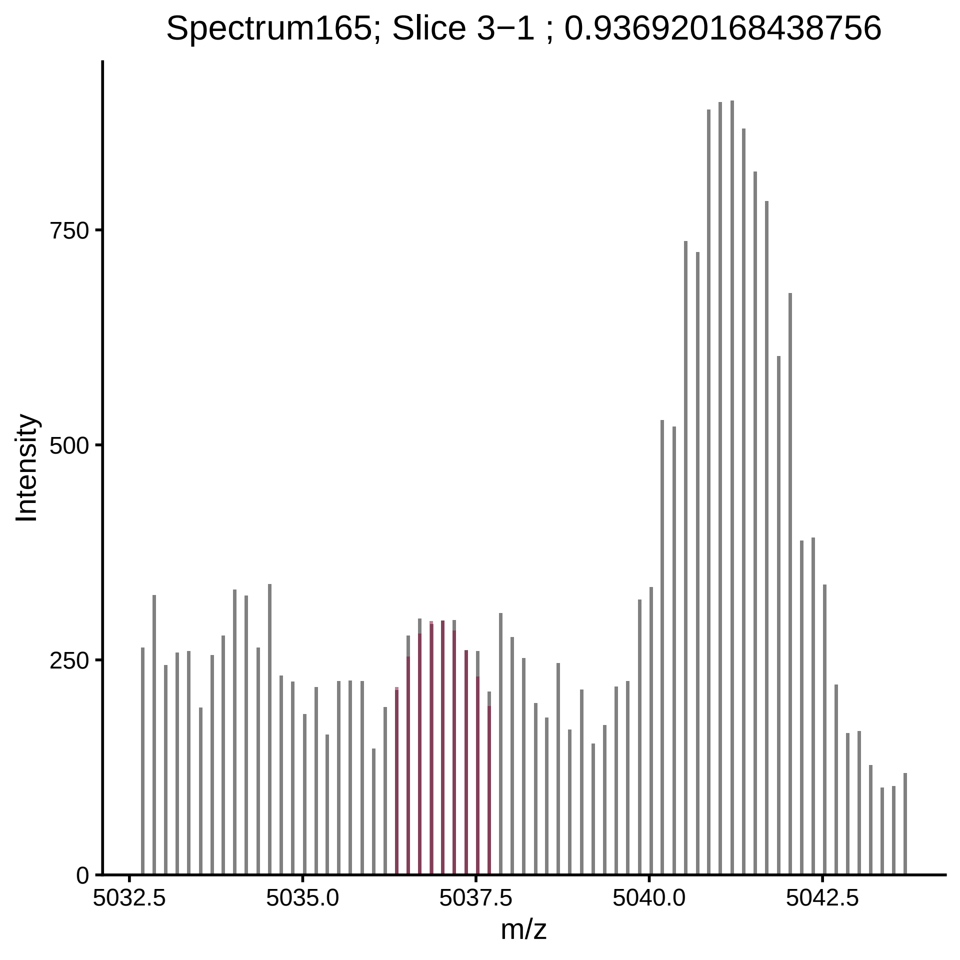

Spectrum165; Slice 3−1 ; 0.936920168438756
Intensity
m/z
0
250
500
750
5032.5
5035.0
5037.5
5040.0
5042.5

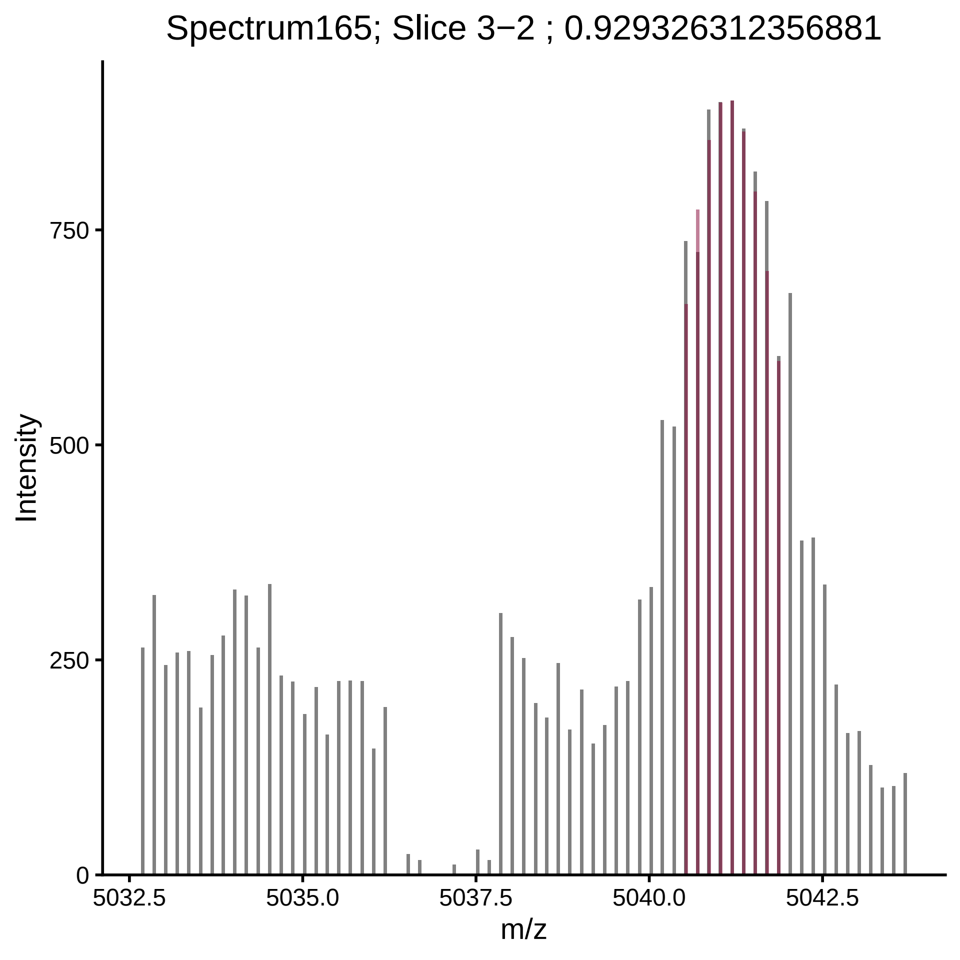
Spectrum165; Slice 3−2 ; 0.929326312356881
Intensity
750
500
250
0
5032.5
5035.0
5037.5
5040.0
5042.5
m/z

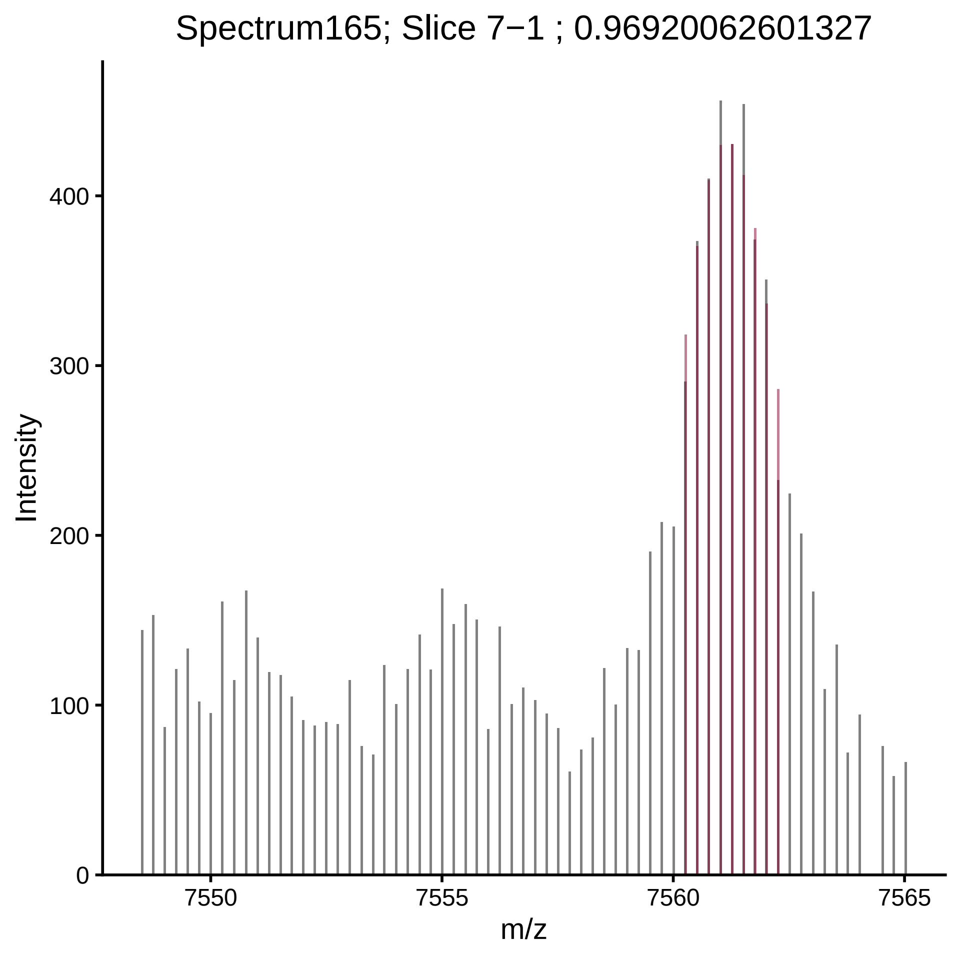

Spectrum165; Slice 7−1 ; 0.96920062601327
Intensity
400
300
200
100
0
7550
7555
7560
7565
m/z

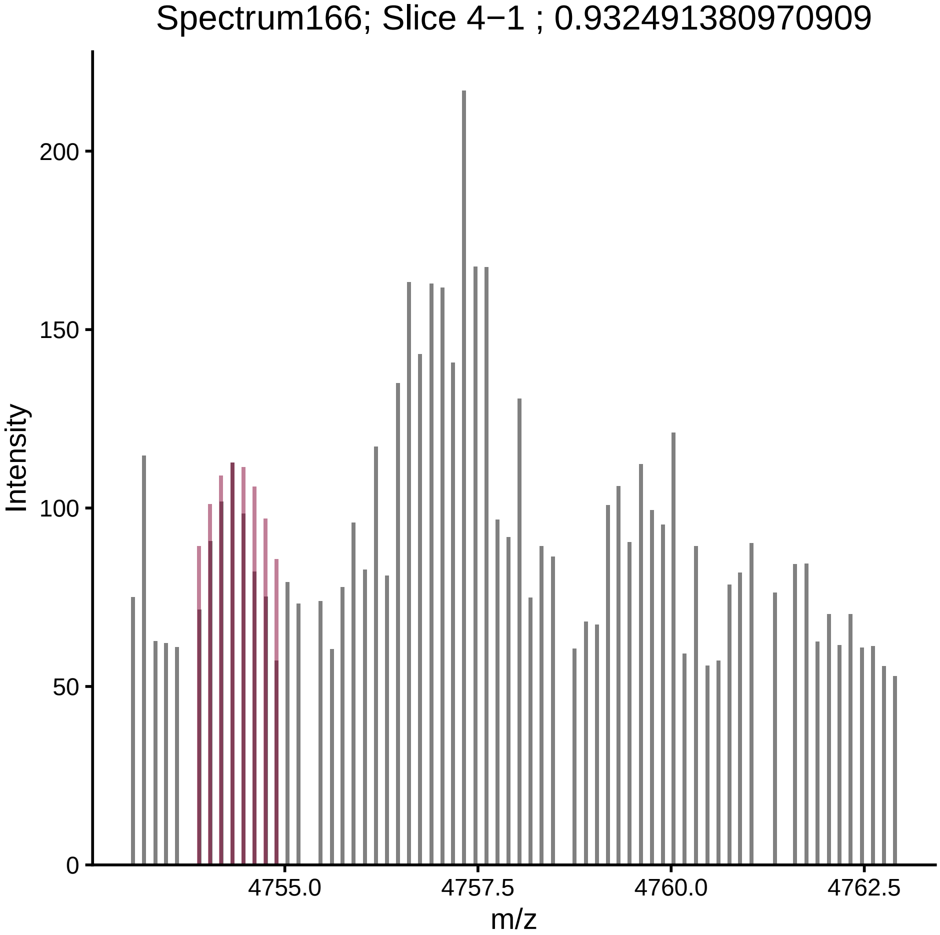

Spectrum166; Slice 4−1 ; 0.932491380970909
Intensity
200
150
100
50
0
4755.0
4757.5
4760.0
4762.5
m/z

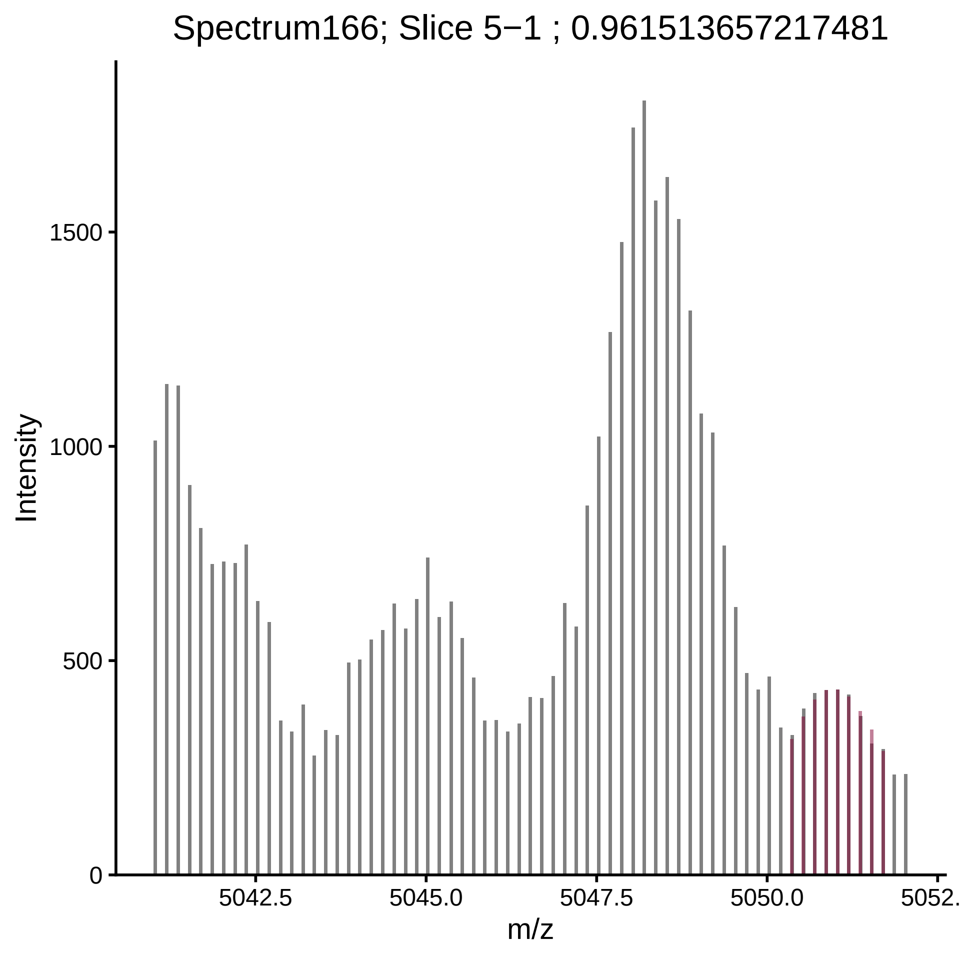
Spectrum166; Slice 5−1 ; 0.961513657217481
Intensity
0
500
1000
1500
5042.5
5045.0
5047.5
5050.0
5052.
m/z

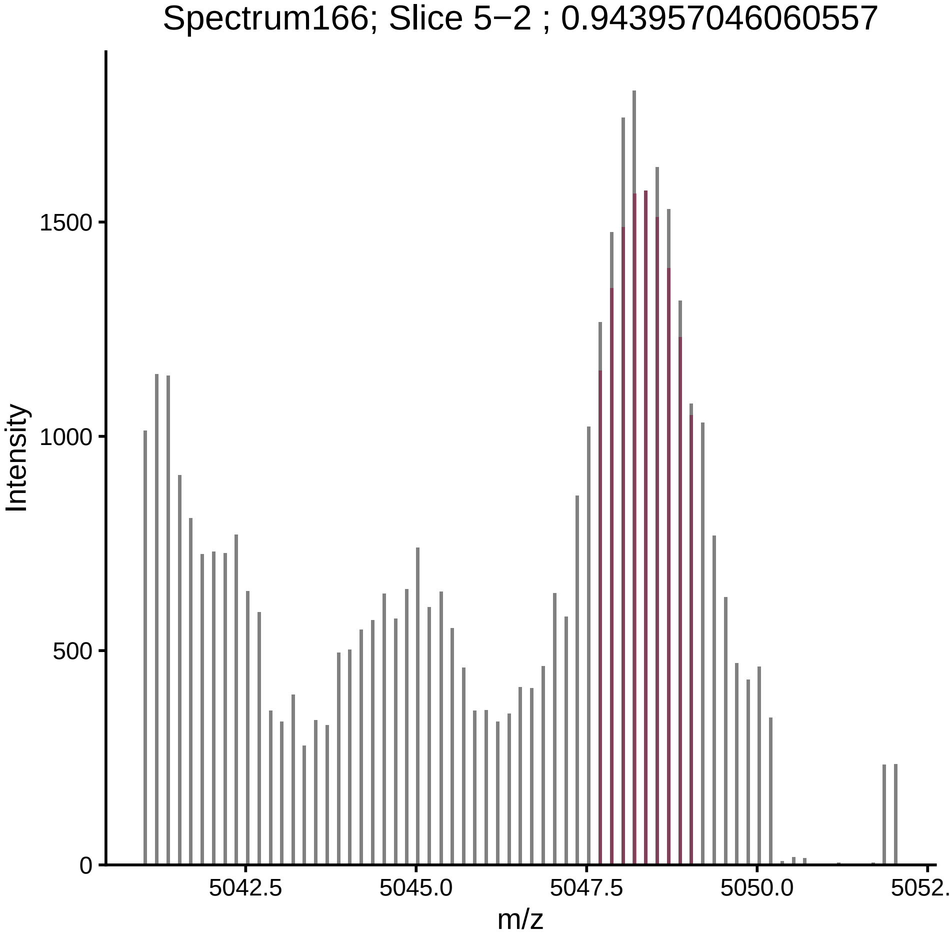

Spectrum166; Slice 5−2 ; 0.943957046060557
Intensity
1500
1000
500
0
5042.5
5045.0
5047.5
5050.0
5052.
m/z

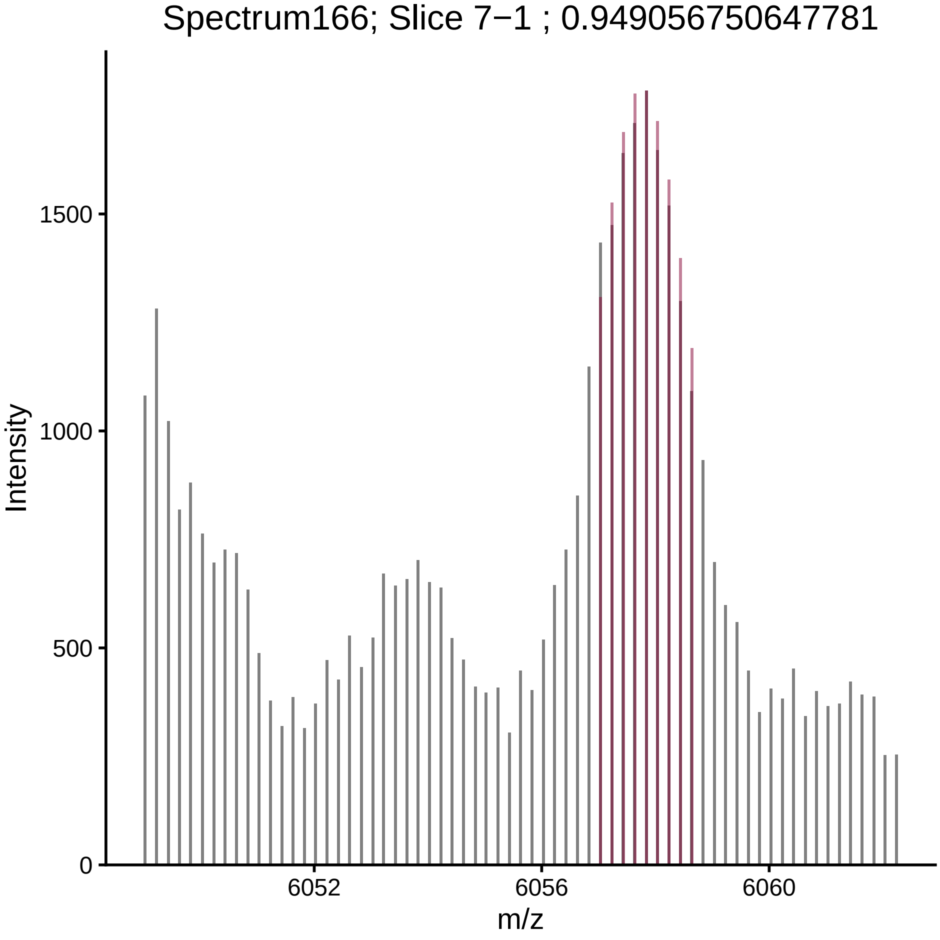
Spectrum166; Slice 7−1 ; 0.949056750647781
Intensity
1500
1000
500
0
6052
6056
6060
m/z

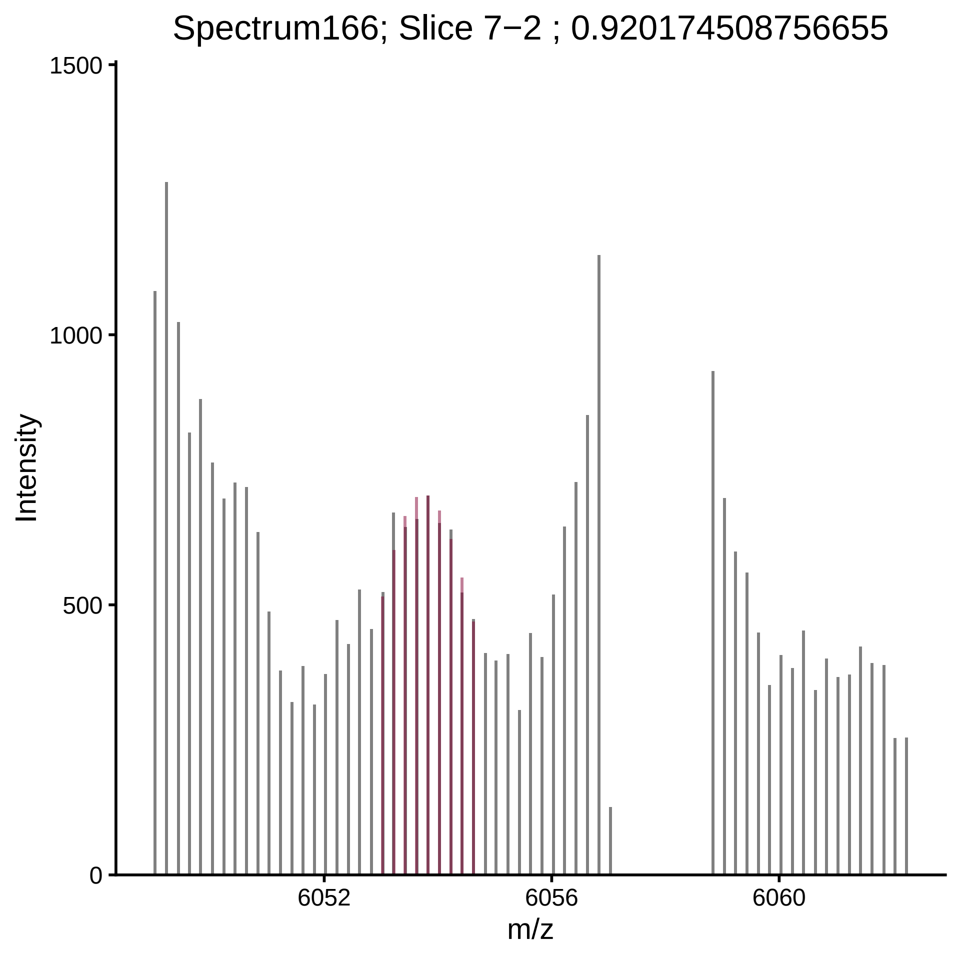

Spectrum166; Slice 7−2 ; 0.920174508756655
Intensity
1500
1000
500
0
6052
6056
6060
m/z

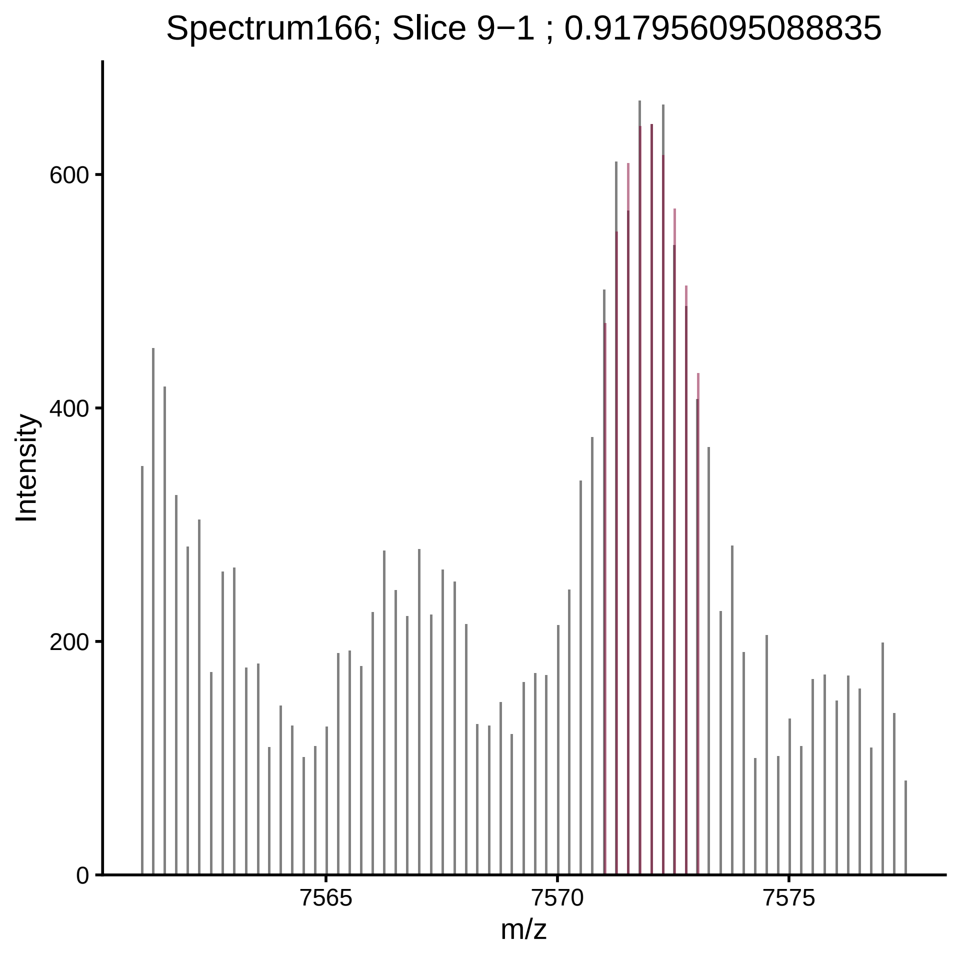
Spectrum166; Slice 9−1 ; 0.917956095088835
Intensity
0
200
400
600
7565
7570
7575
m/z

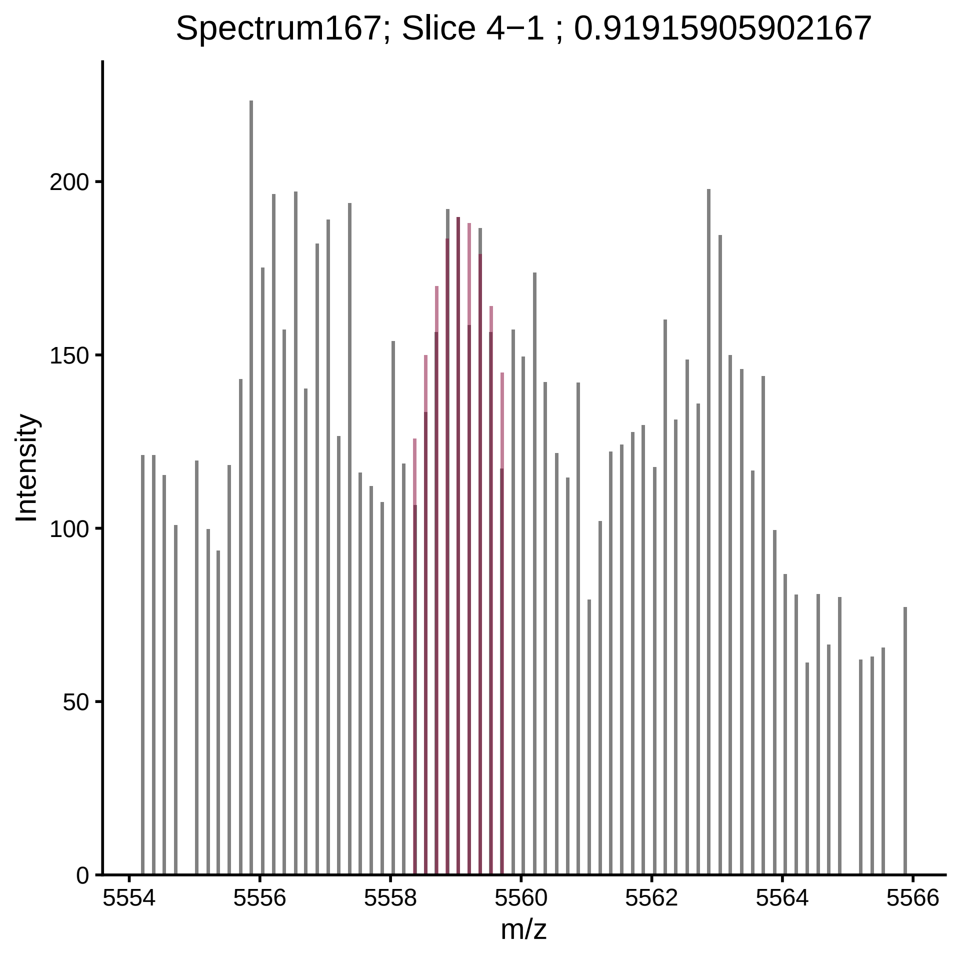

Spectrum167; Slice 4−1 ; 0.91915905902167
Intensity
0
50
100
150
200
5554
5556
5558
5560
5562
5564
5566
m/z

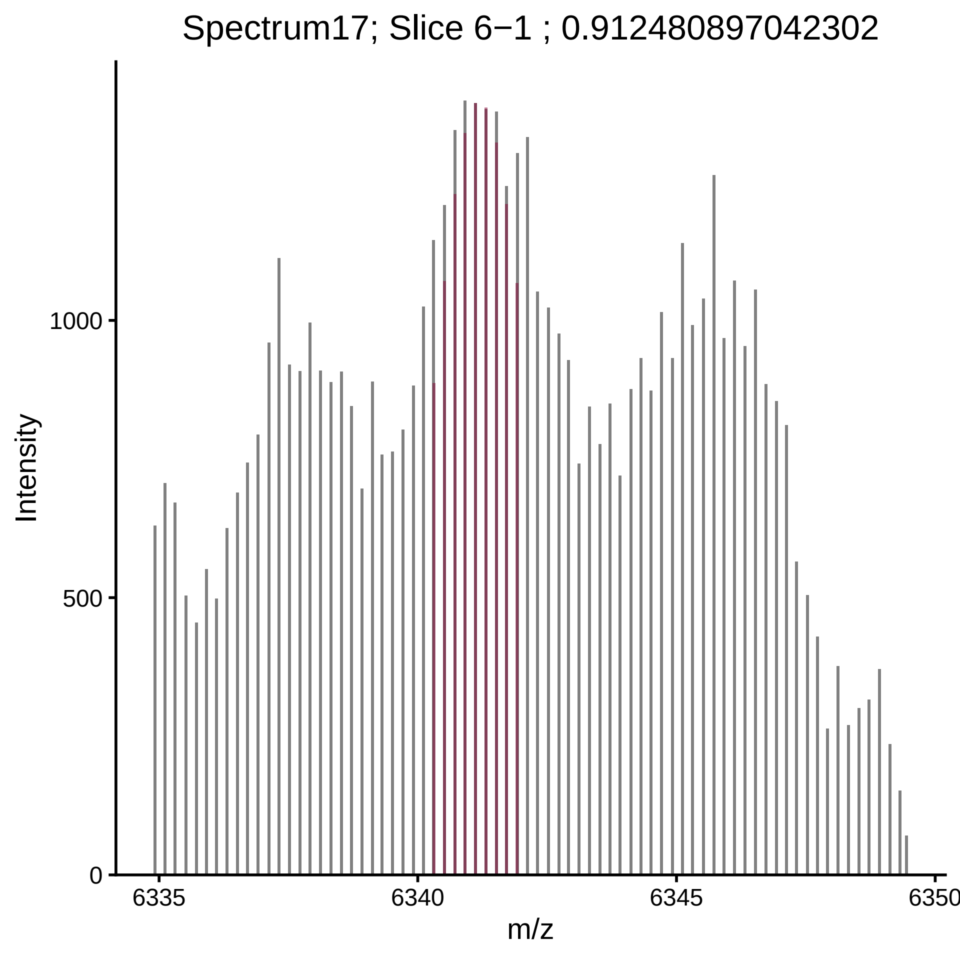

Spectrum17; Slice 6−1 ; 0.912480897042302
Intensity
1000
500
0
6335
6340
6345
6350
m/z

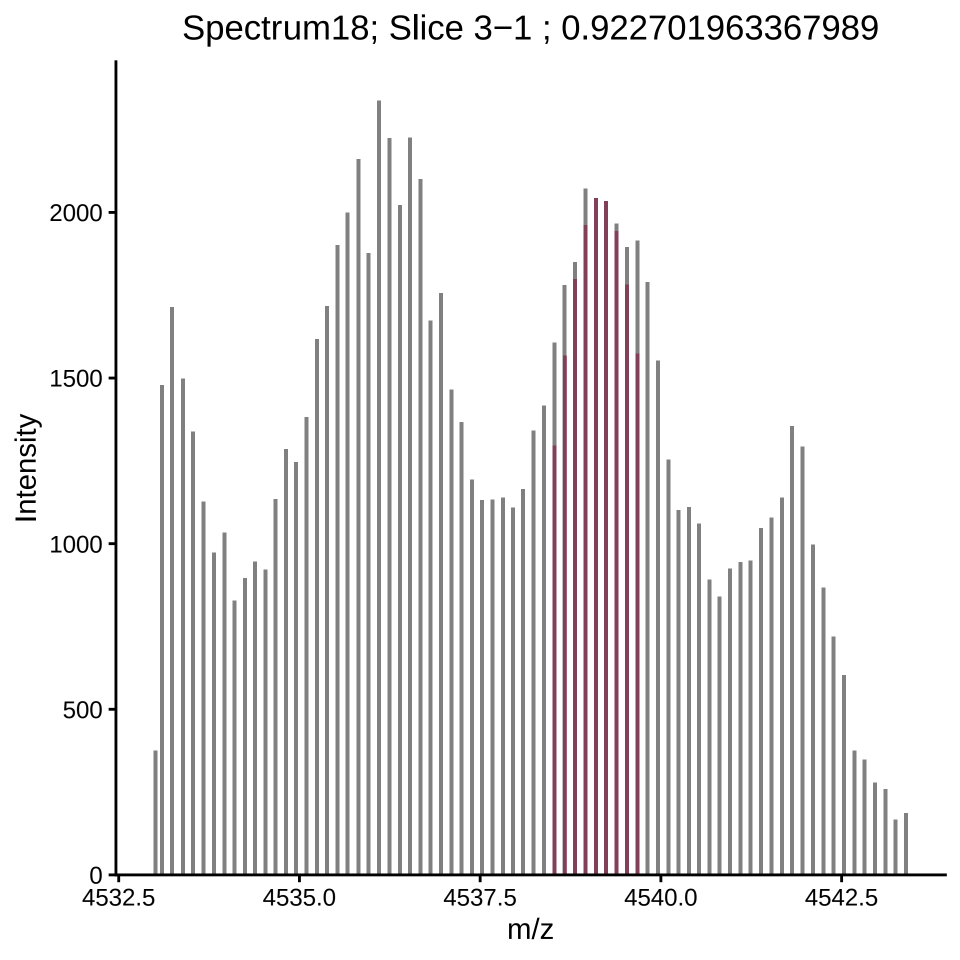
Spectrum18; Slice 3−1 ; 0.922701963367989
Intensity
2000
1500
1000
500
0
4532.5
4535.0
4537.5
4540.0
4542.5
m/z

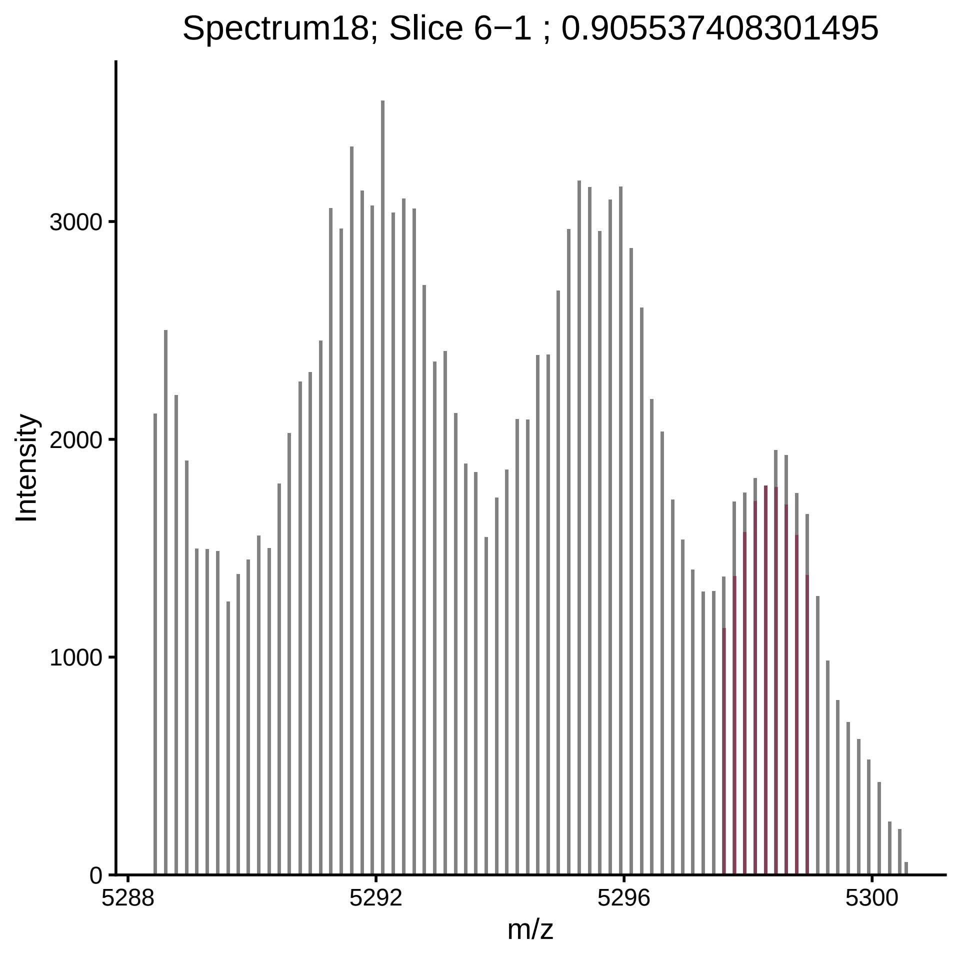

Spectrum18; Slice 6−1 ; 0.905537408301495
Intensity
3000
2000
1000
0
5288
5292
5296
5300
m/z

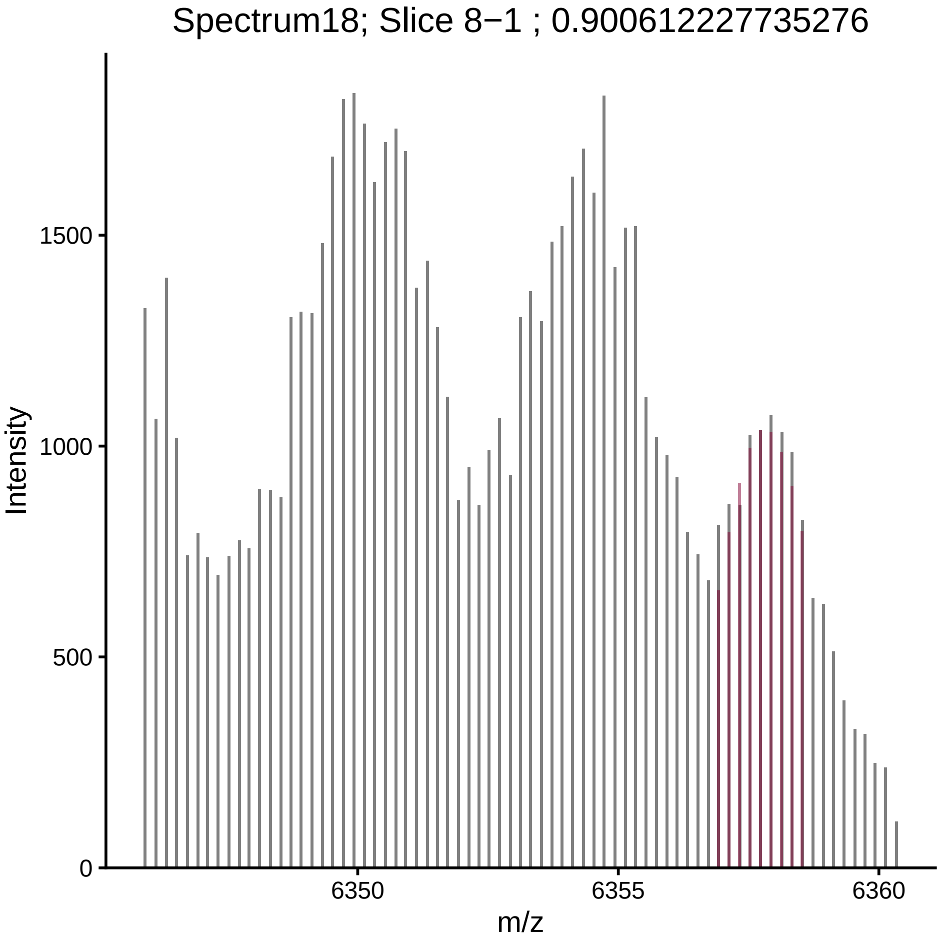

Spectrum18; Slice 8−1 ; 0.900612227735276
Intensity
m/z
0
500
1000
1500
6350
6355
6360

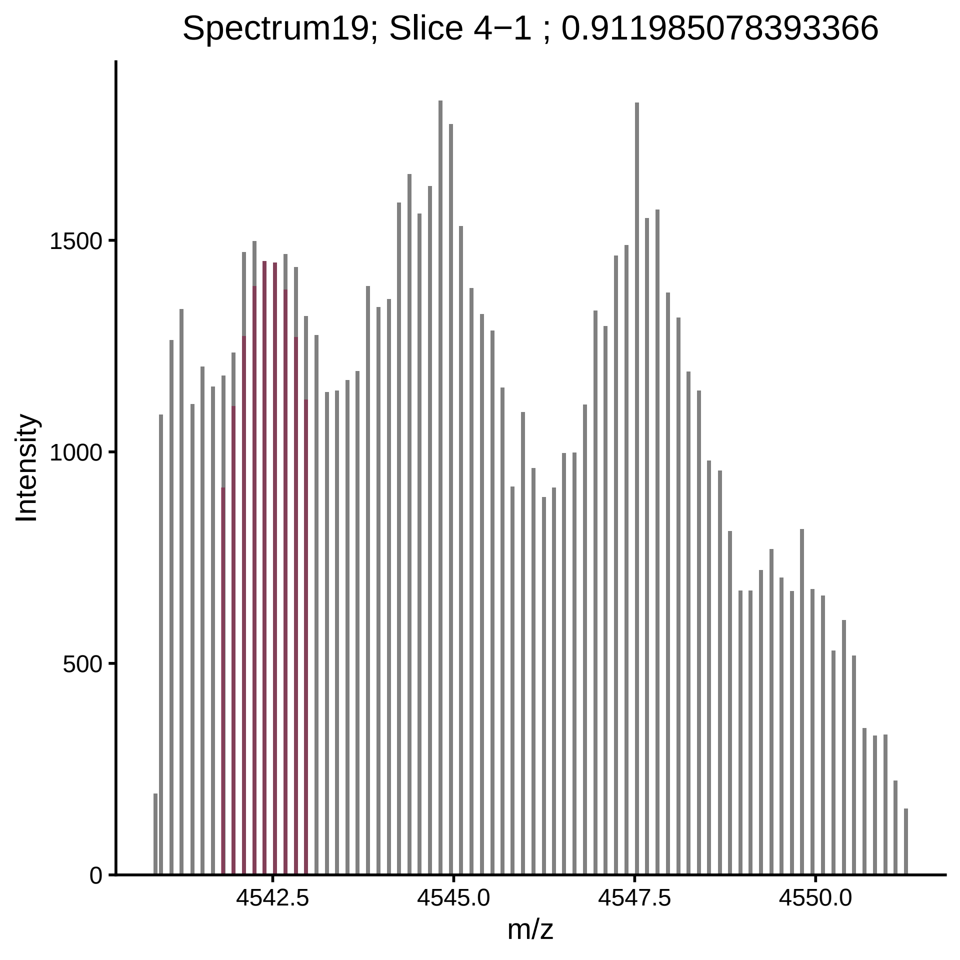

Spectrum19; Slice 4−1 ; 0.911985078393366
Intensity
1500
1000
500
0
4542.5
4545.0
4547.5
4550.0
m/z

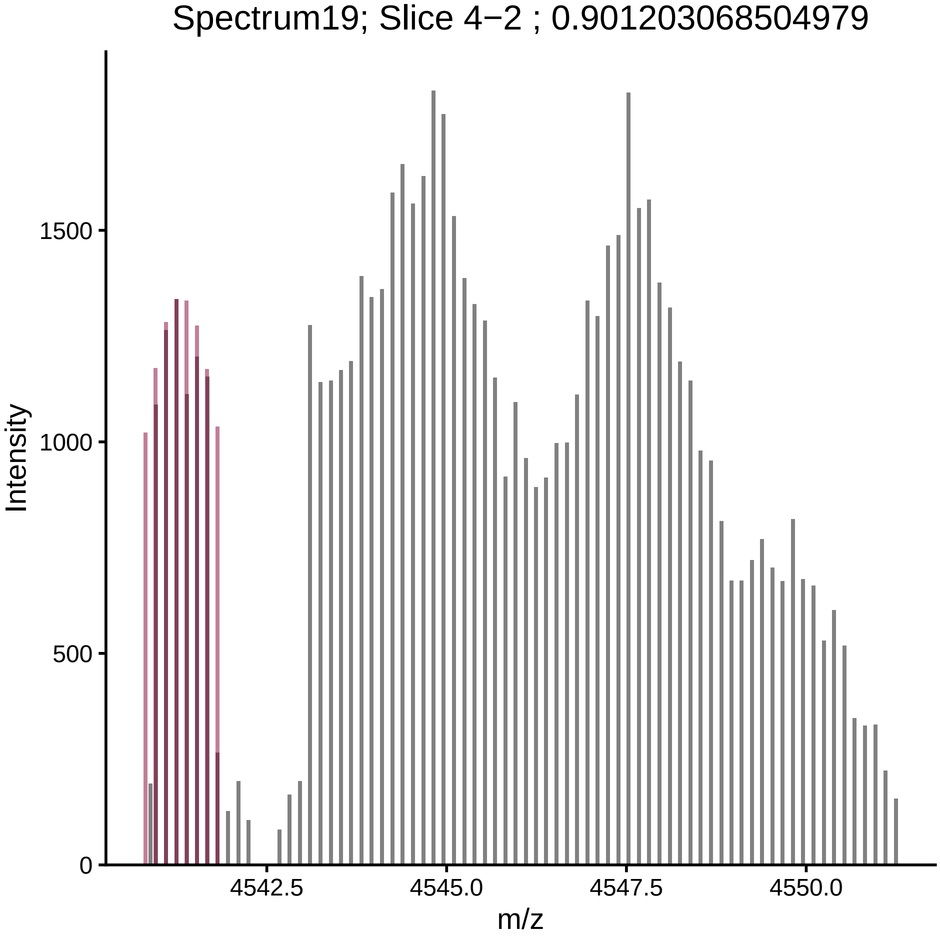

Spectrum19; Slice 4−2 ; 0.901203068504979
Intensity
m/z
1500
1000
500
0
4542.5
4545.0
4547.5
4550.0

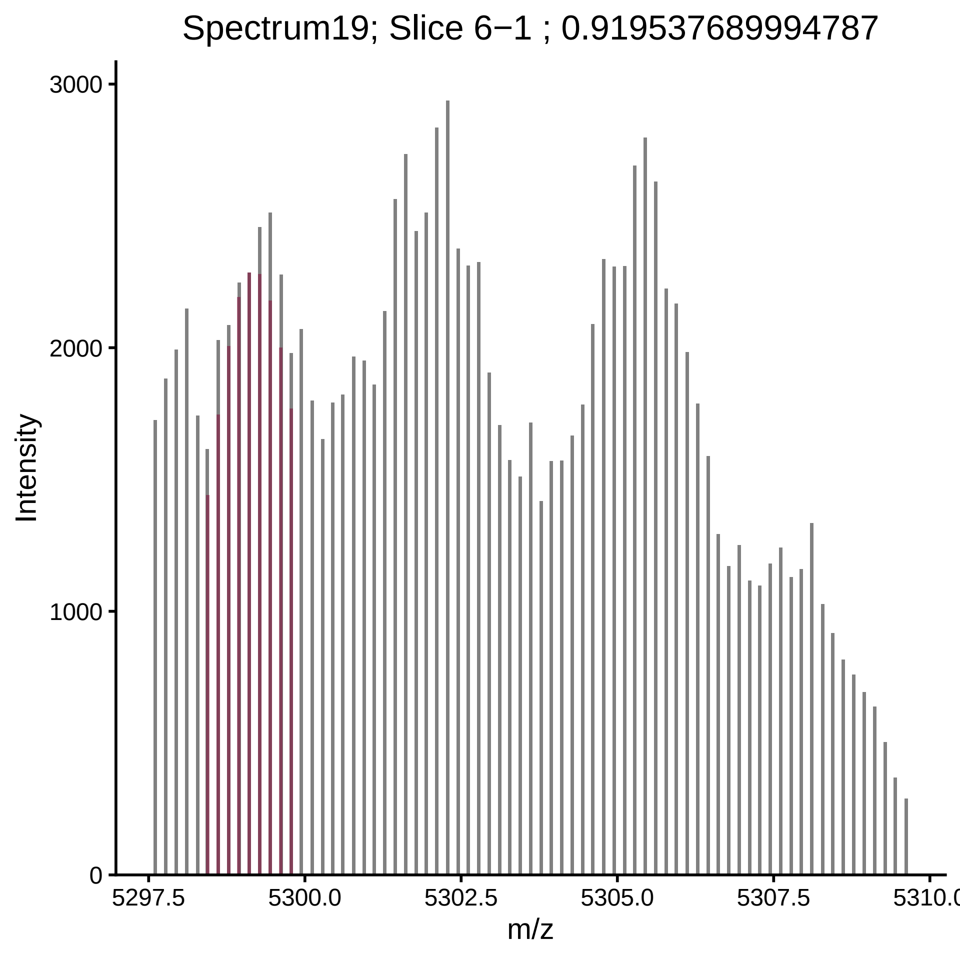

Spectrum19; Slice 6−1 ; 0.919537689994787
Intensity
3000
2000
1000
0
5297.5
5300.0
5302.5
5305.0
5307.5
5310.0
m/z

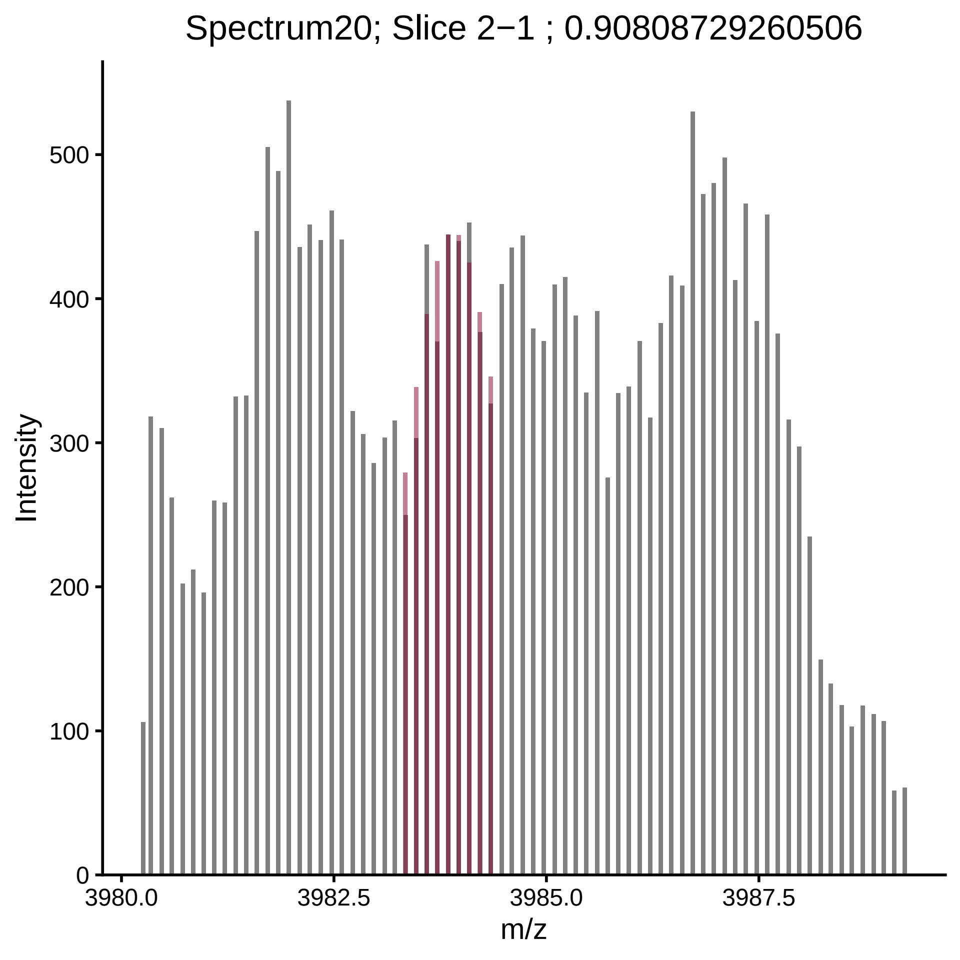

Spectrum20; Slice 2−1 ; 0.90808729260506
Intensity
0
100
200
300
400
500
3980.0
3982.5
3985.0
3987.5
m/z

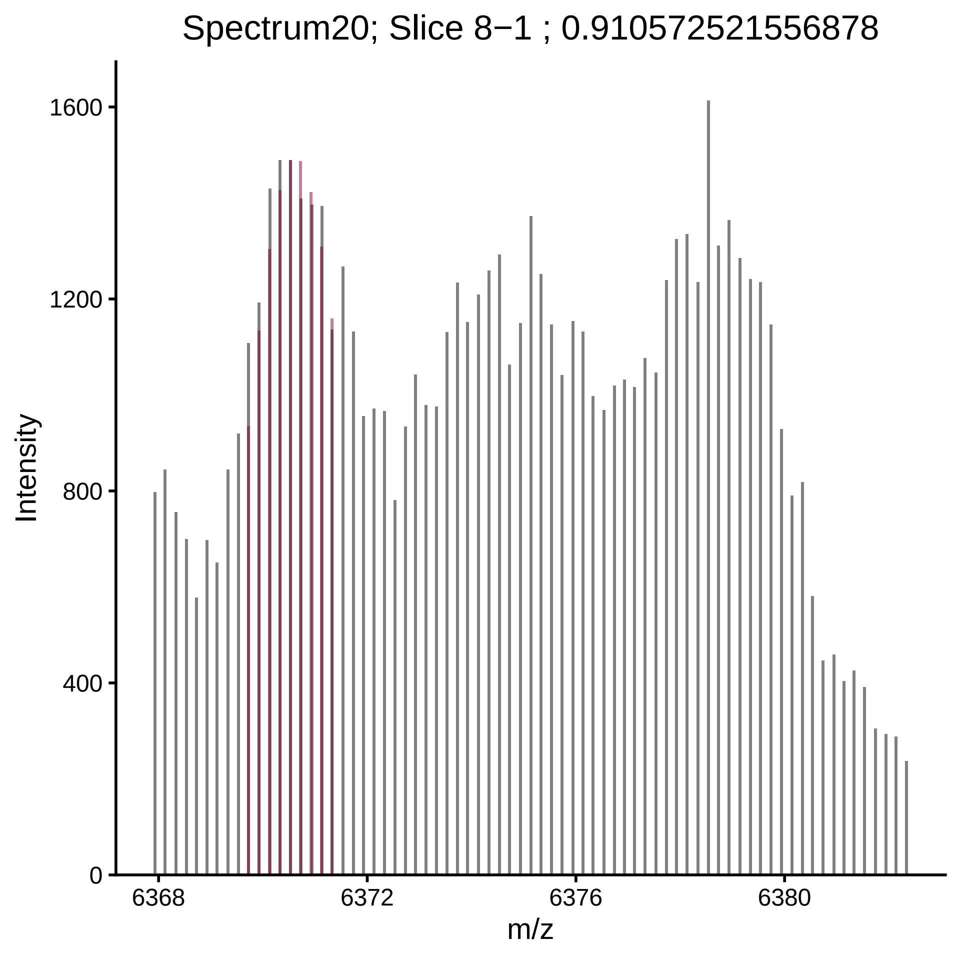

Spectrum20; Slice 8−1 ; 0.910572521556878
Intensity
1600
1200
800
400
0
6368
6372
6376
6380
m/z

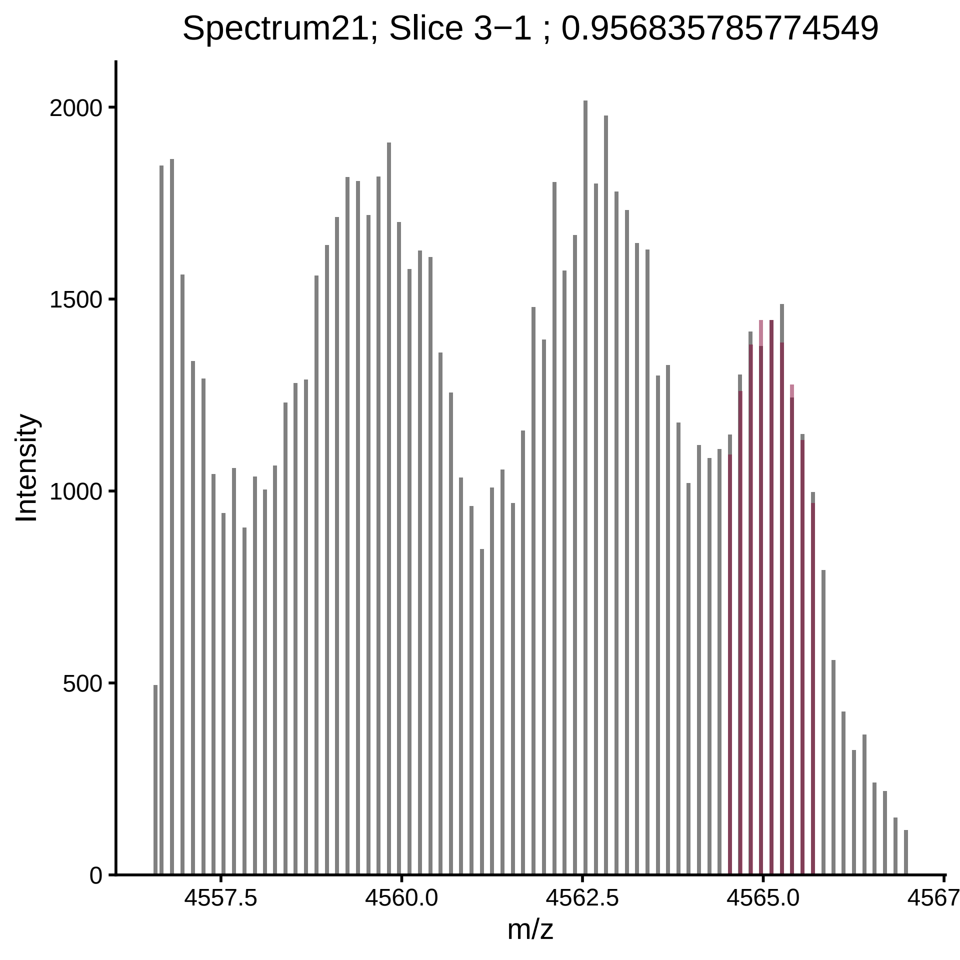
Spectrum21; Slice 3−1 ; 0.956835785774549
Intensity
2000
1500
1000
500
0
4557.5
4560.0
4562.5
4565.0
4567
m/z

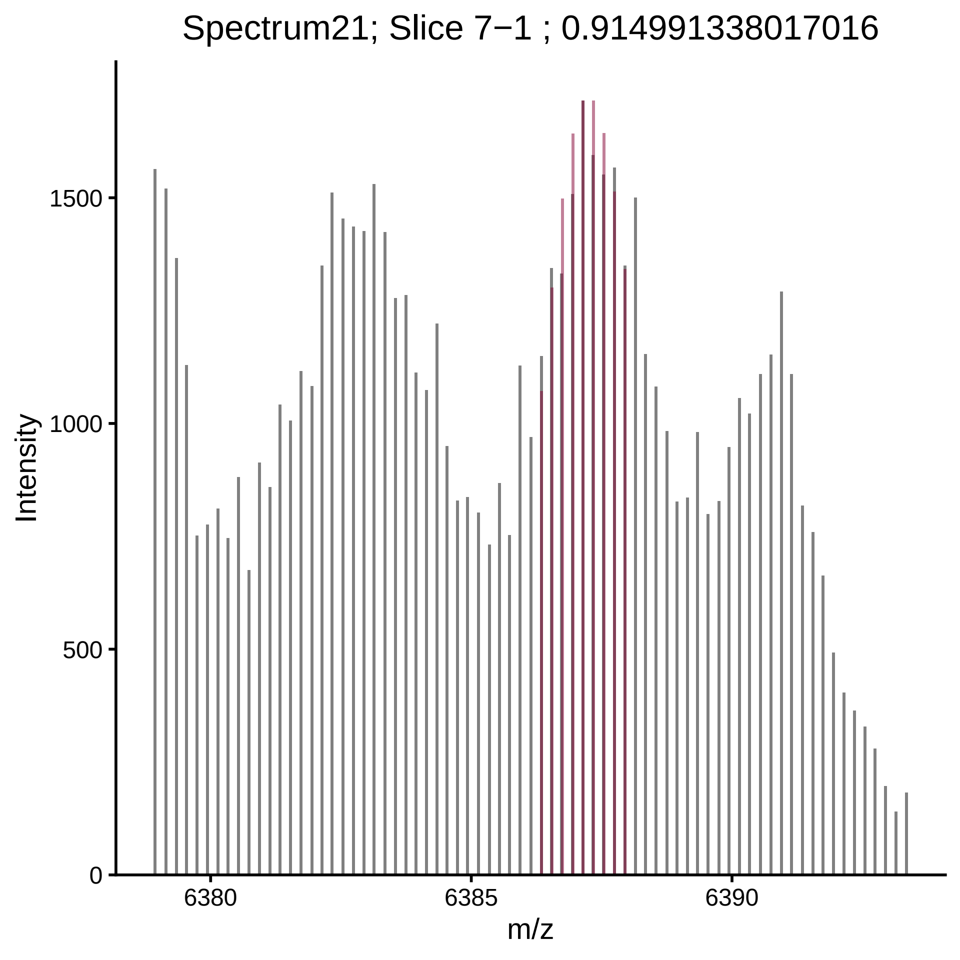

Spectrum21; Slice 7−1 ; 0.914991338017016
Intensity
1500
1000
500
0
6380
6385
6390
m/z

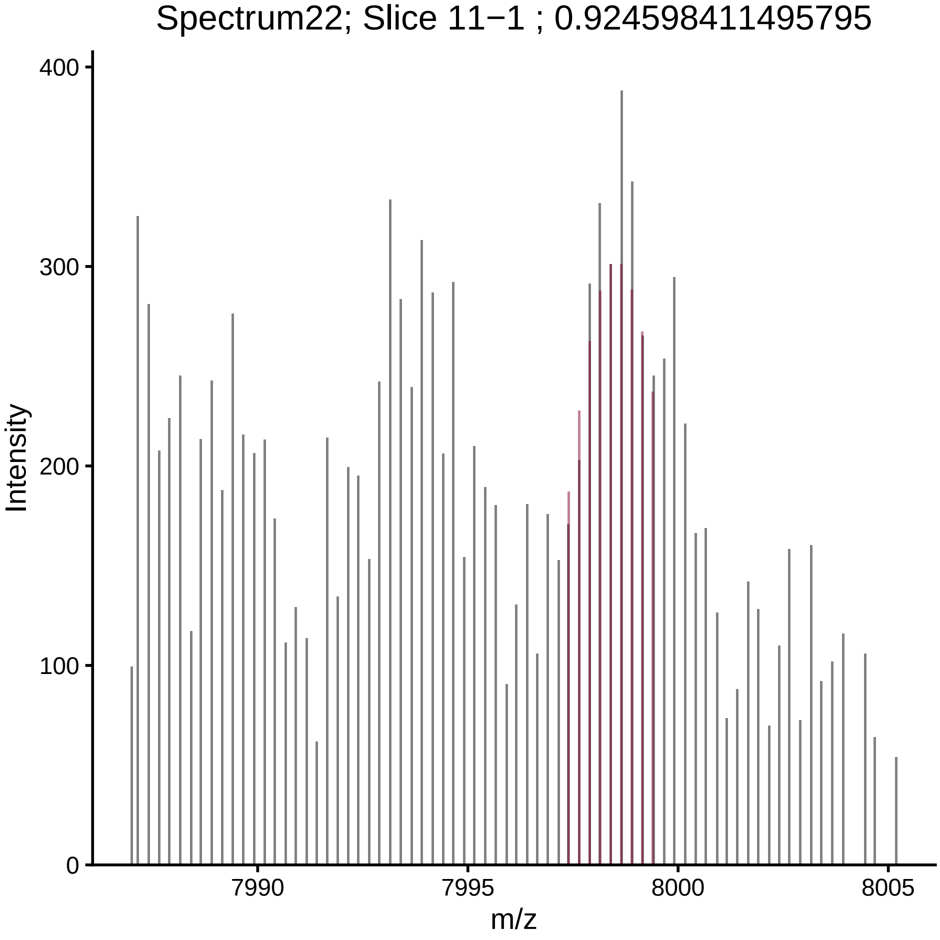

Spectrum22; Slice 11−1 ; 0.924598411495795
Intensity
m/z
0
100
200
300
400
7990
7995
8000
8005

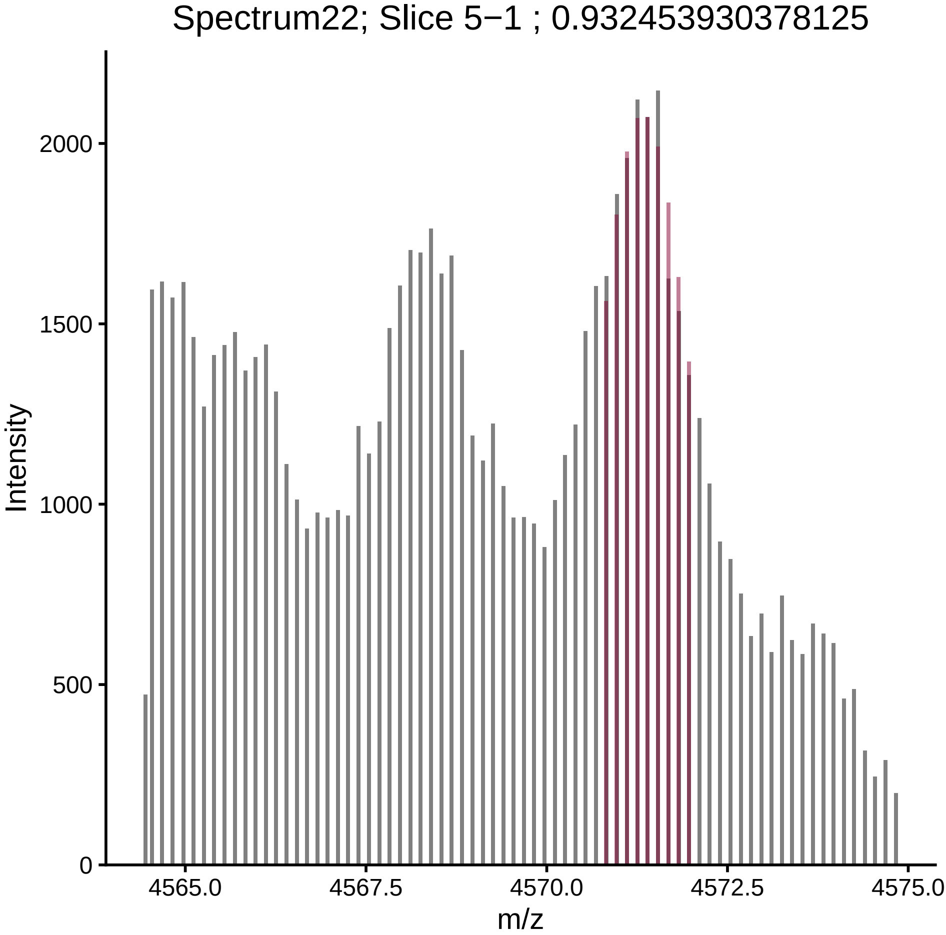

Spectrum22; Slice 5−1 ; 0.932453930378125
Intensity
m/z
0
500
1000
1500
2000
4565.0
4567.5
4570.0
4572.5
4575.0

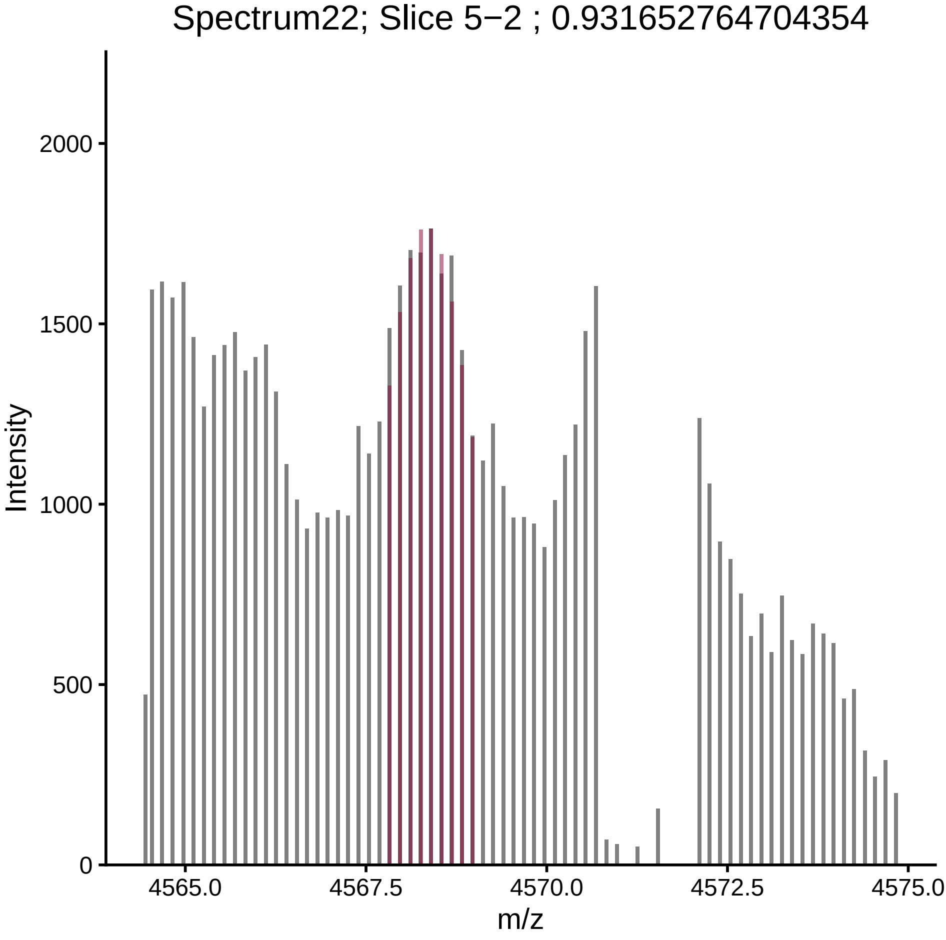

Spectrum22; Slice 5−2 ; 0.931652764704354
Intensity
2000
1500
1000
500
0
4565.0
4567.5
4570.0
4572.5
4575.0
m/z

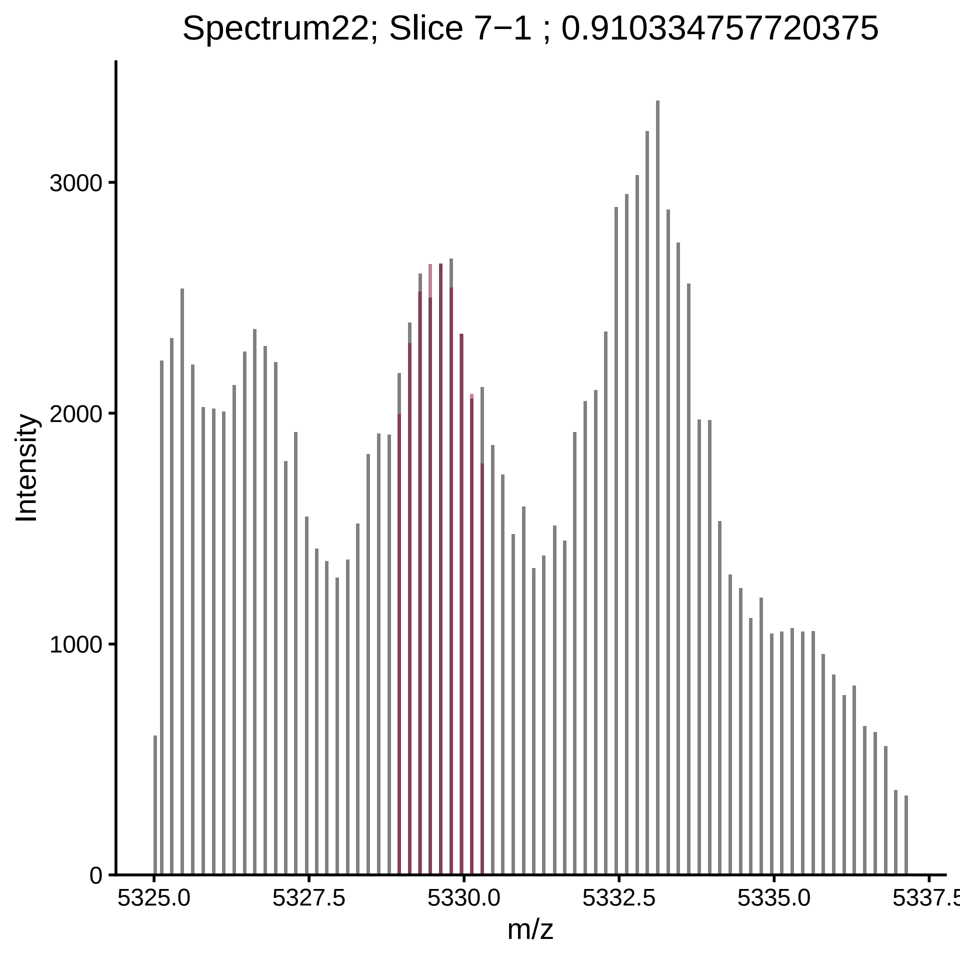

Spectrum22; Slice 7−1 ; 0.910334757720375
Intensity
3000
2000
1000
0
5325.0
5327.5
5330.0
5332.5
5335.0
5337.5
m/z

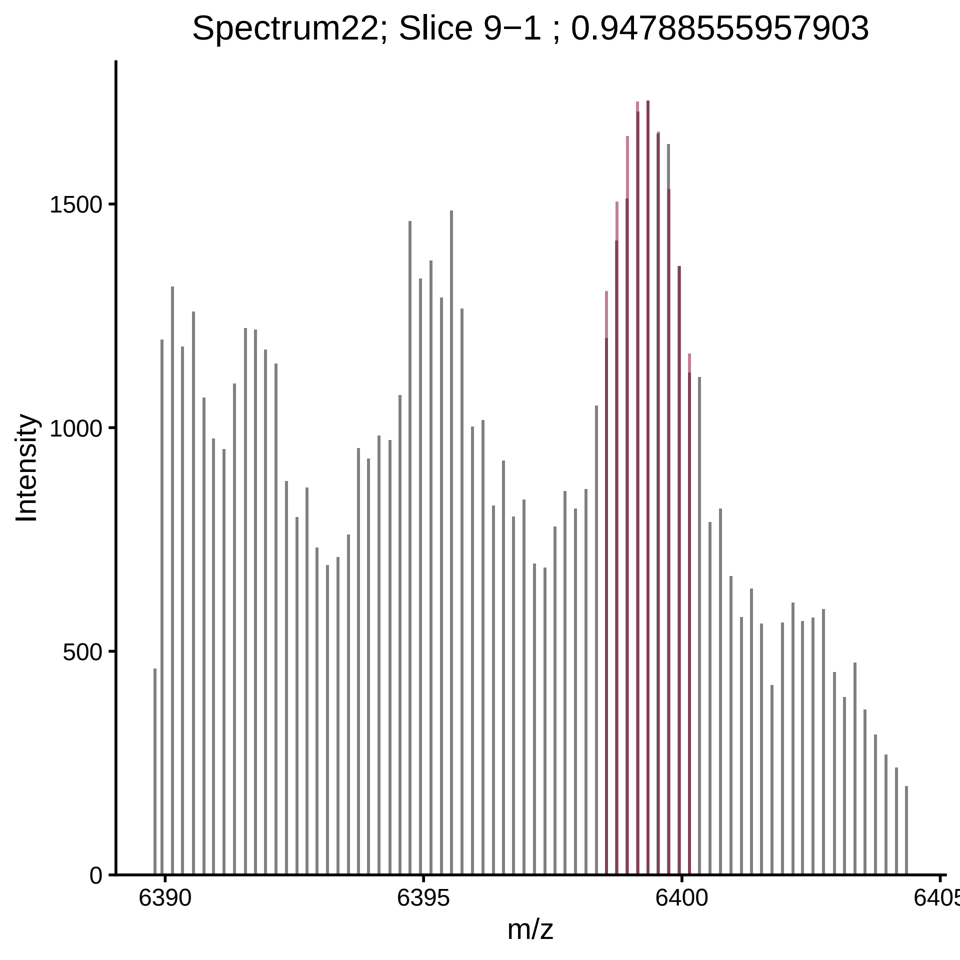
Spectrum22; Slice 9−1 ; 0.947885559579033
Intensity
1500
1000
500
0
6390
6395
6400
6405
m/z

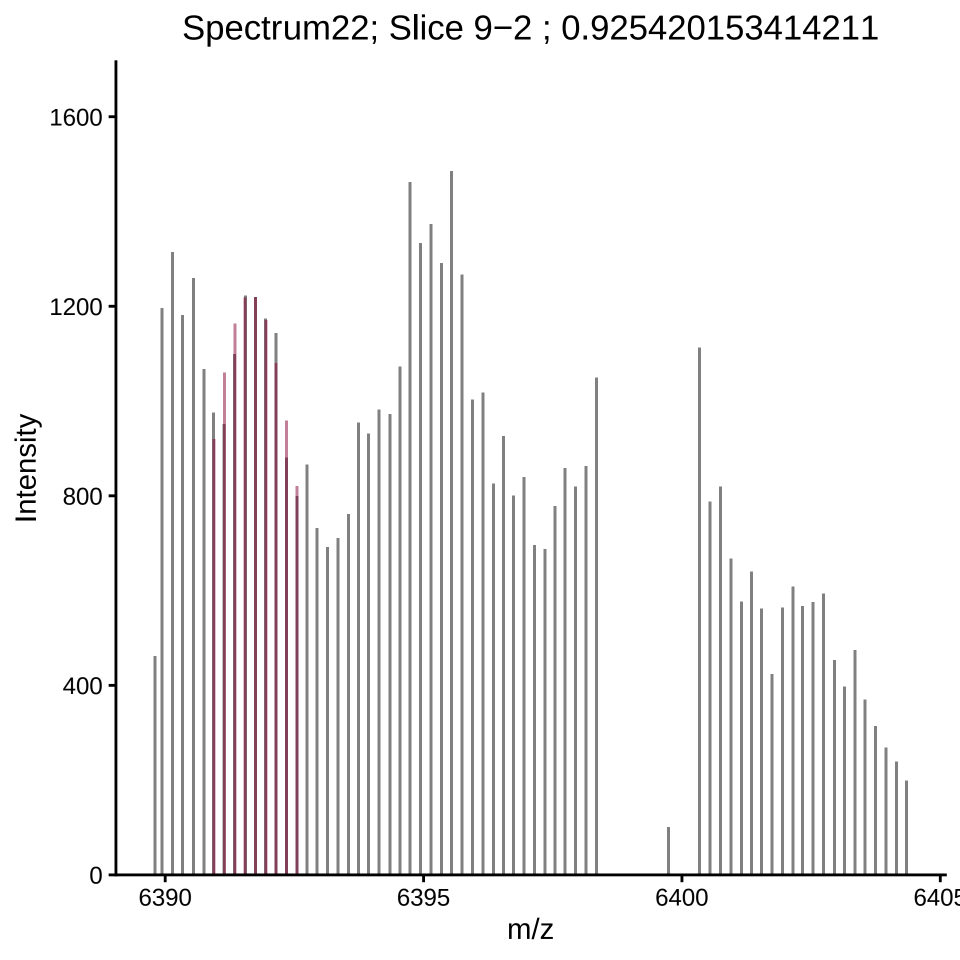

Spectrum22; Slice 9−2 ; 0.925420153414211
Intensity
1600
1200
800
400
0
6390
6395
6400
6405
m/z

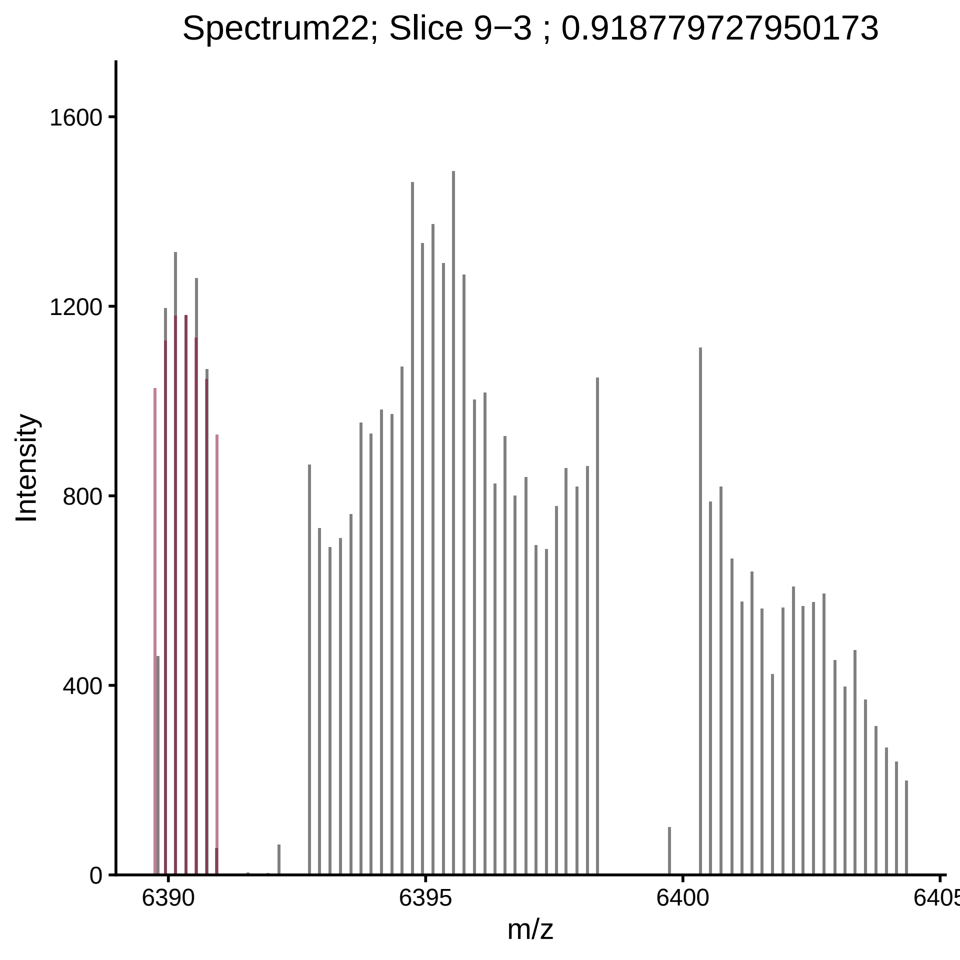

Spectrum22; Slice 9−3 ; 0.918779727950173
Intensity
1600
1200
800
400
0
6390
6395
6400
6405
m/z

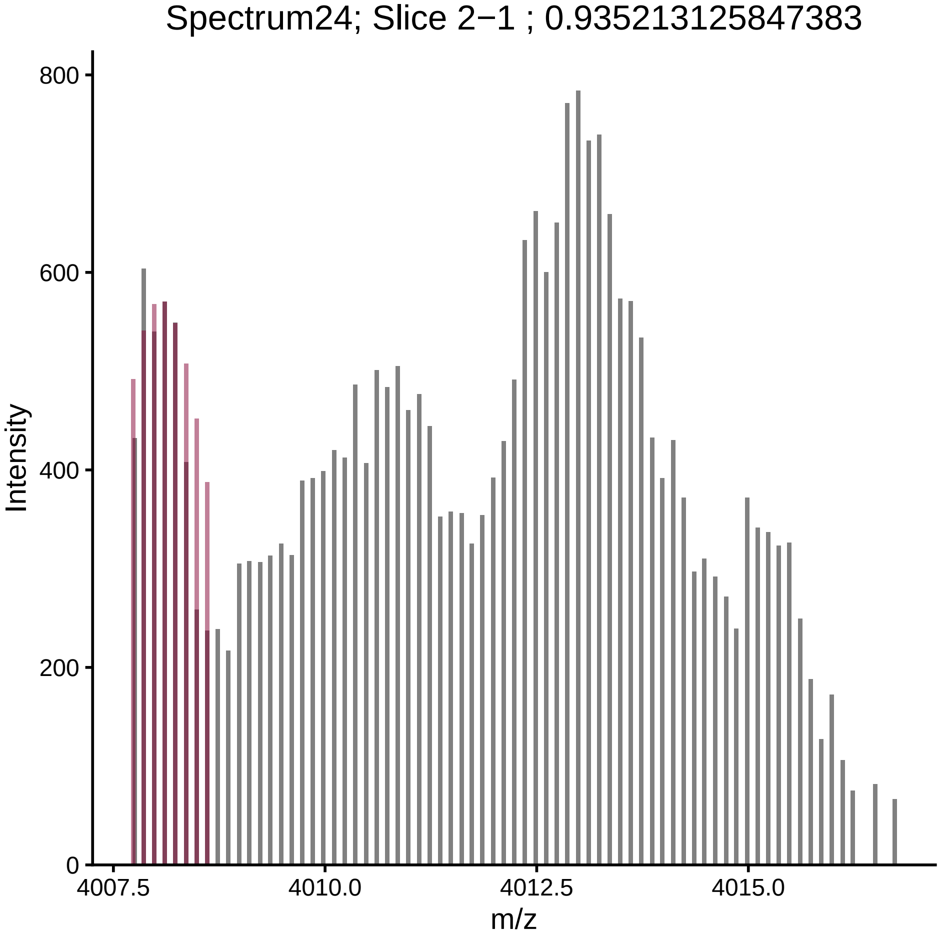

Spectrum24; Slice 2−1 ; 0.935213125847383
Intensity
800
600
400
200
0
4007.5
4010.0
4012.5
4015.0
m/z

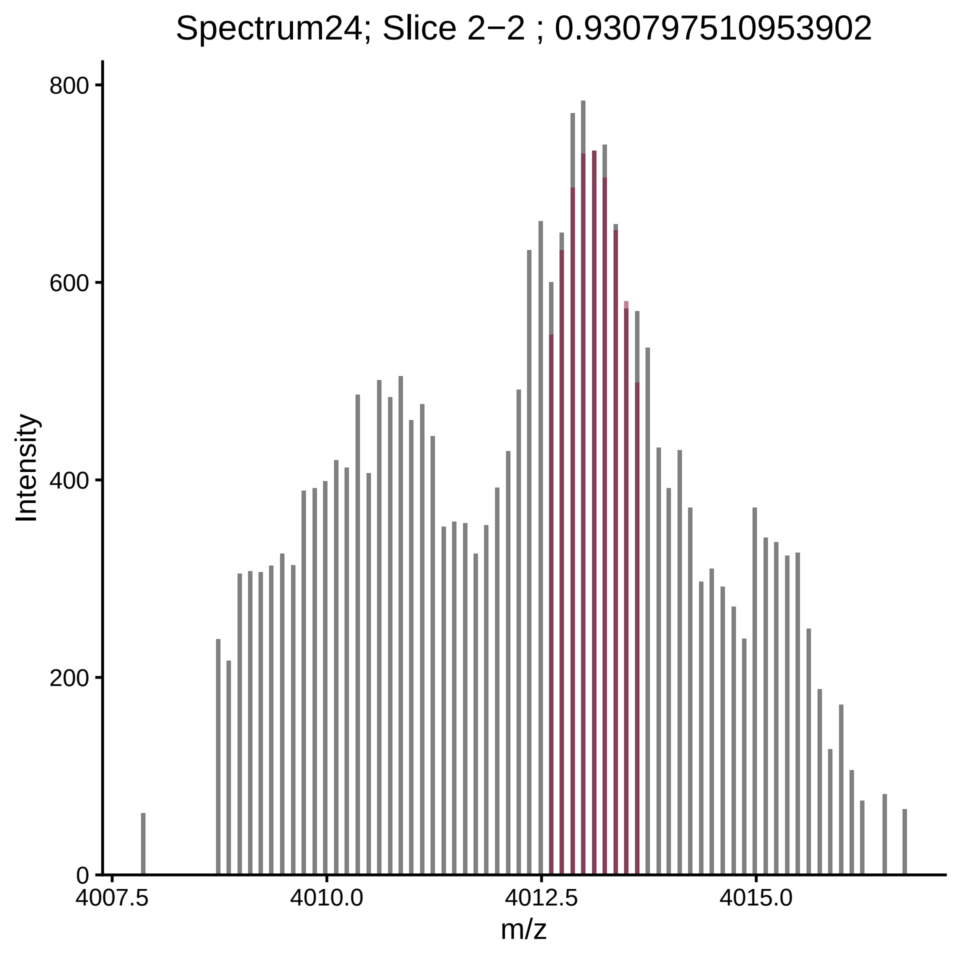

Spectrum24; Slice 2−2 ; 0.930797510953902
Intensity
800
600
400
200
0
4007.5
4010.0
4012.5
4015.0
m/z

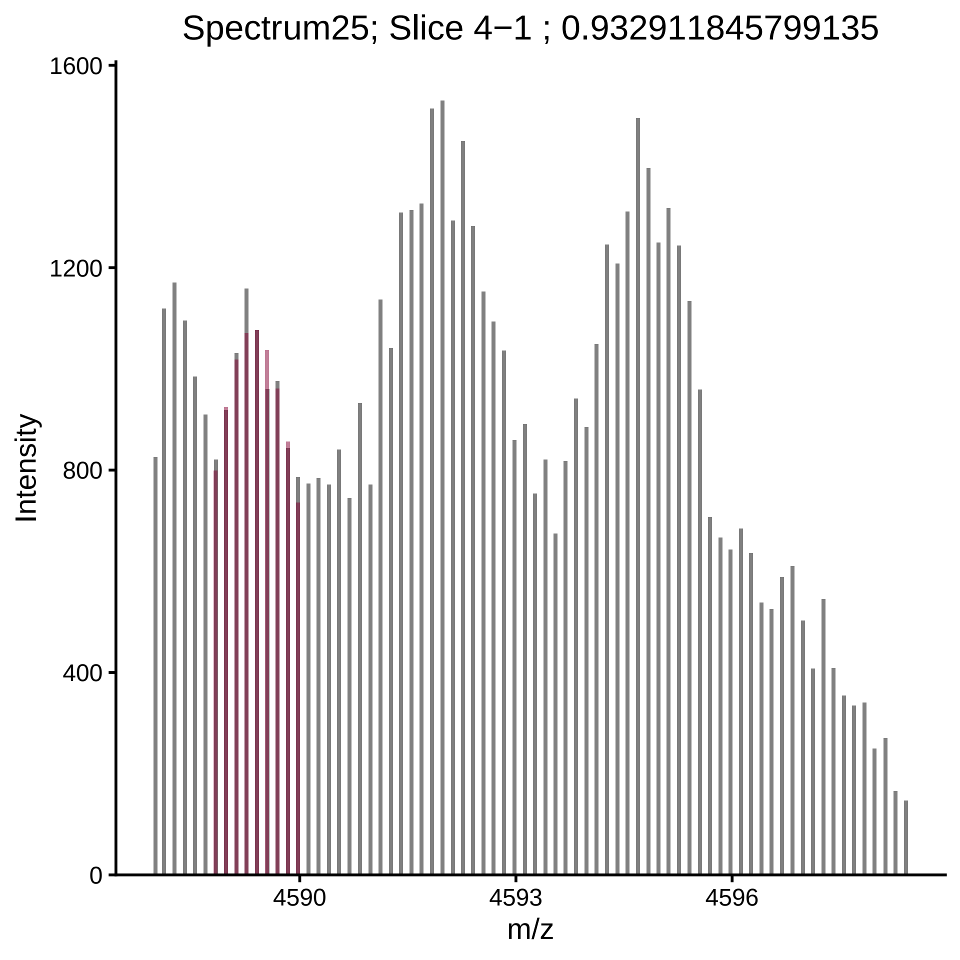

Spectrum25; Slice 4−1 ; 0.932911845799135
Intensity
1600
1200
800
400
0
4590
4593
4596
m/z

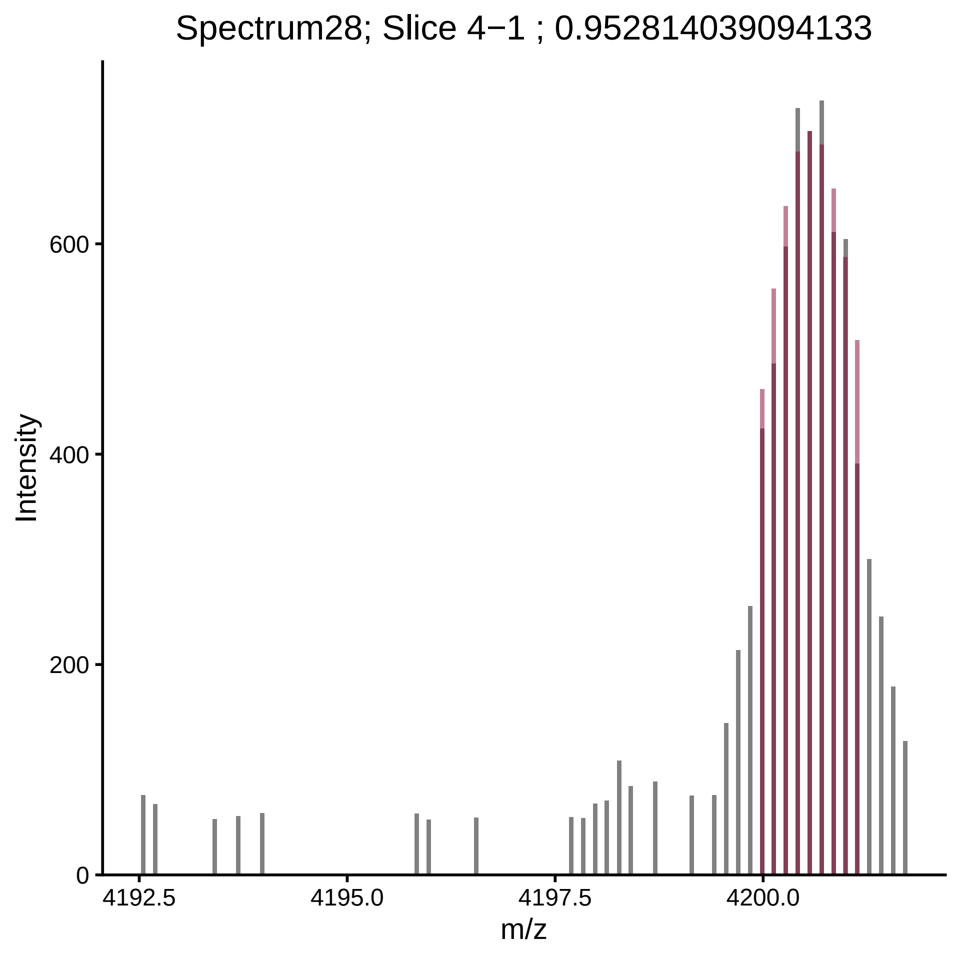
Spectrum28; Slice 4−1 ; 0.952814039094133
Intensity
600
400
200
0
4192.5
4195.0
4197.5
4200.0
m/z

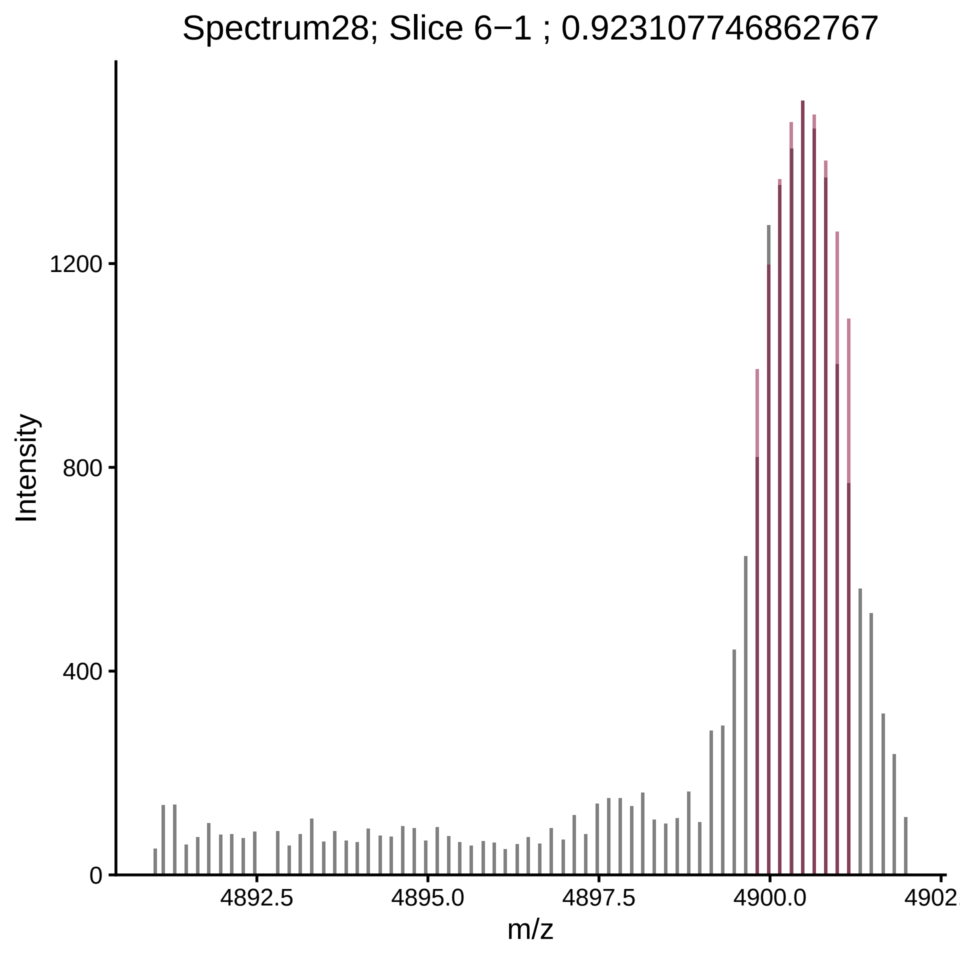

Spectrum28; Slice 6−1 ; 0.923107746862767
Intensity
1200
800
400
0
4892.5
4895.0
4897.5
4900.0
4902.
m/z

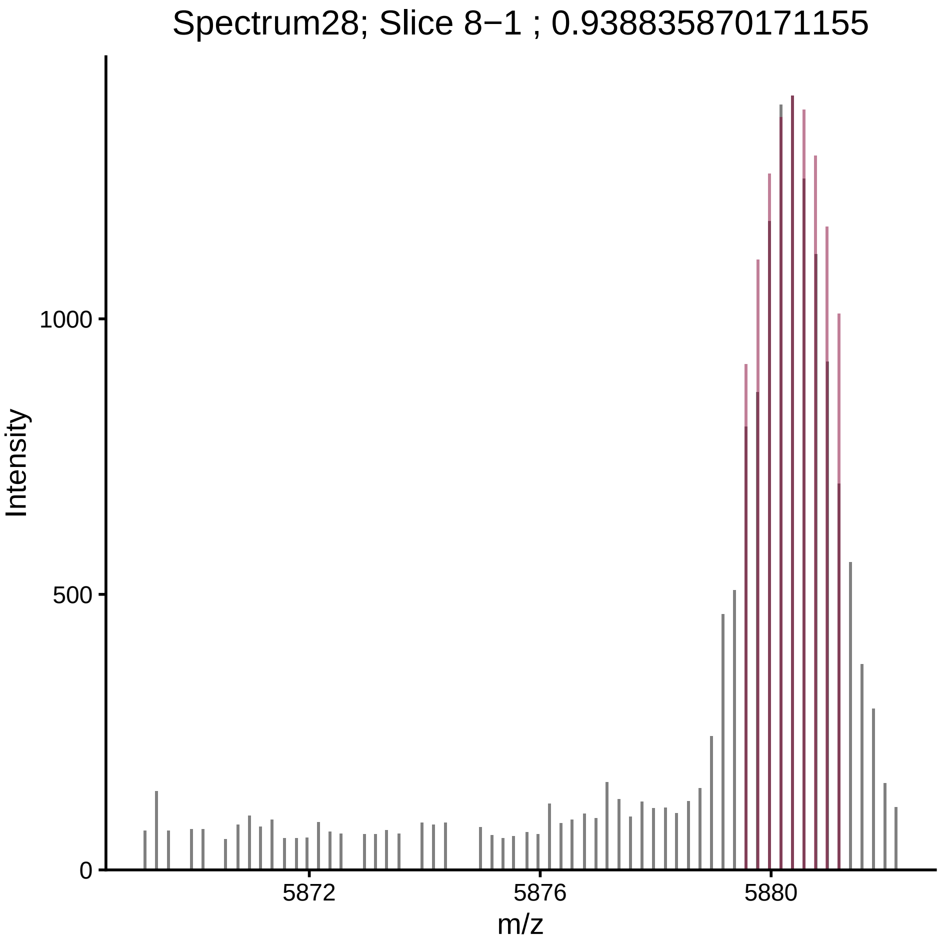
Spectrum28; Slice 8−1 ; 0.938835870171155
Intensity
1000
500
0
5872
5876
5880
m/z

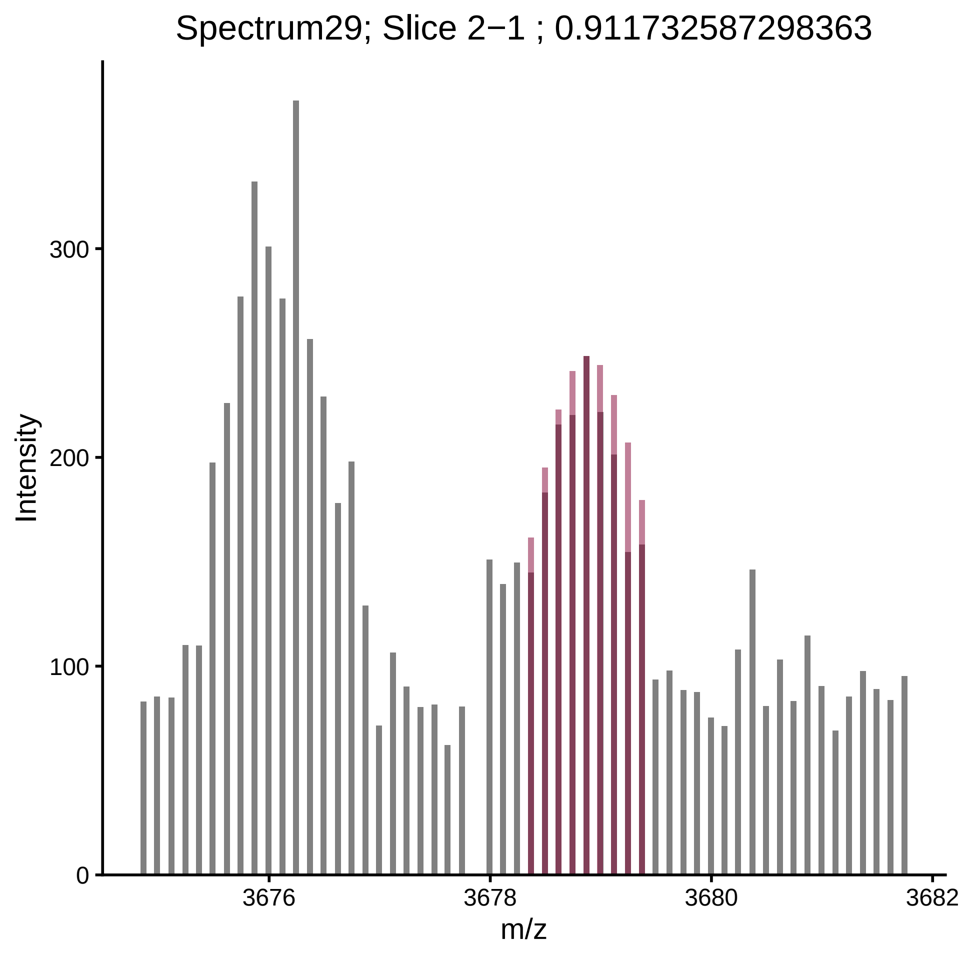
Spectrum29; Slice 2−1 ; 0.911732587298363
Intensity
m/z
0
100
200
300
3676
3678
3680
3682

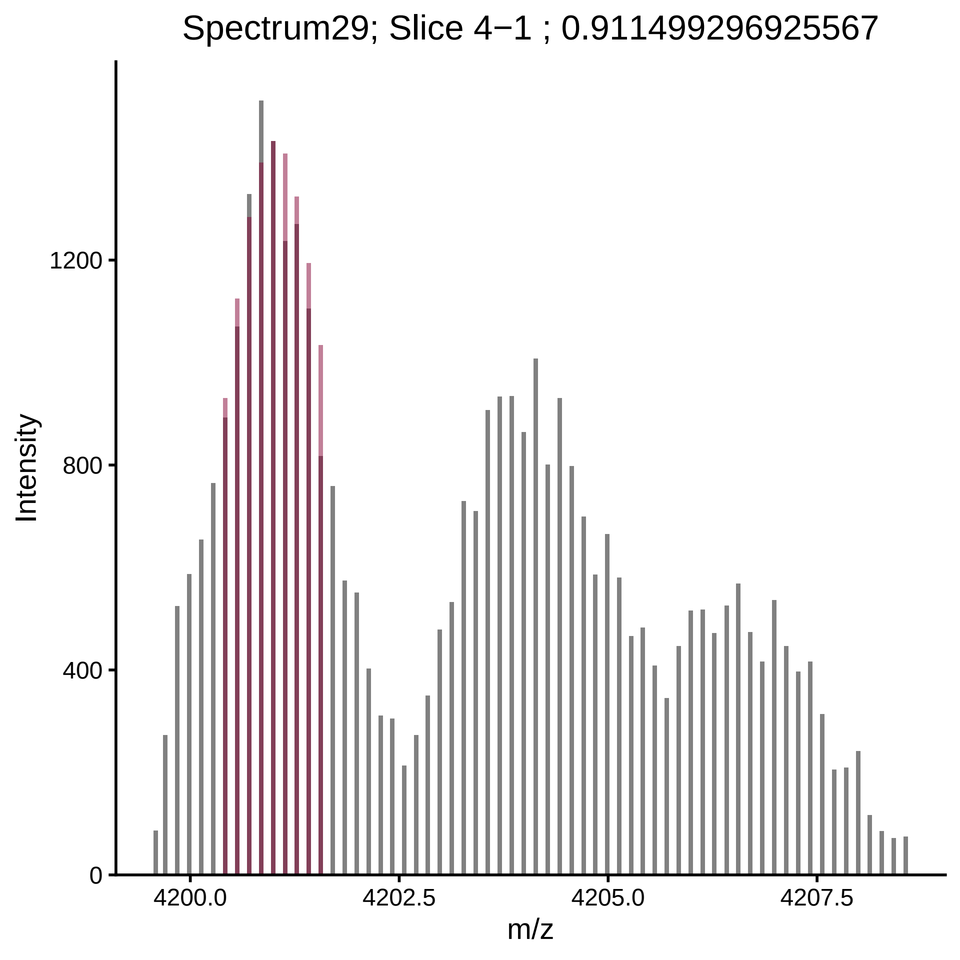
Spectrum29; Slice 4−1 ; 0.911499296925567
Intensity
1200
800
400
0
4200.0
4202.5
4205.0
4207.5
m/z

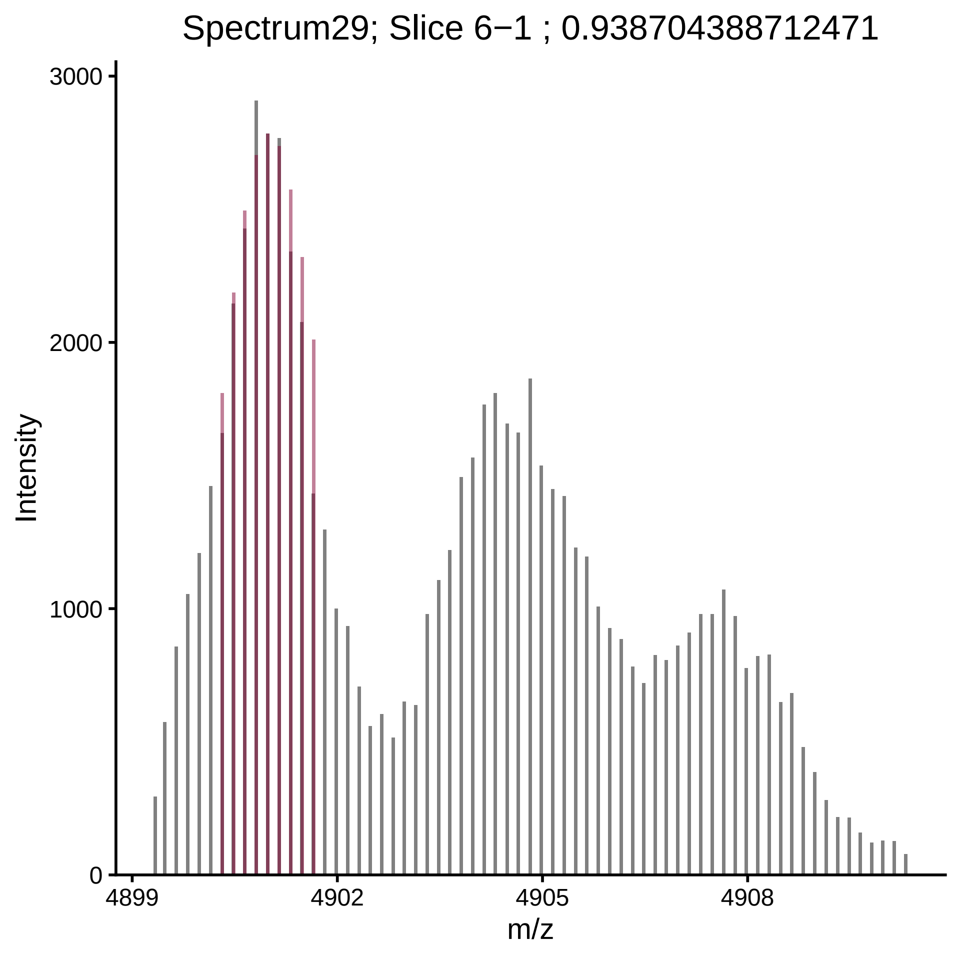

Spectrum29; Slice 6−1 ; 0.938704388712471
Intensity
3000
2000
1000
0
4899
4902
4905
4908
m/z

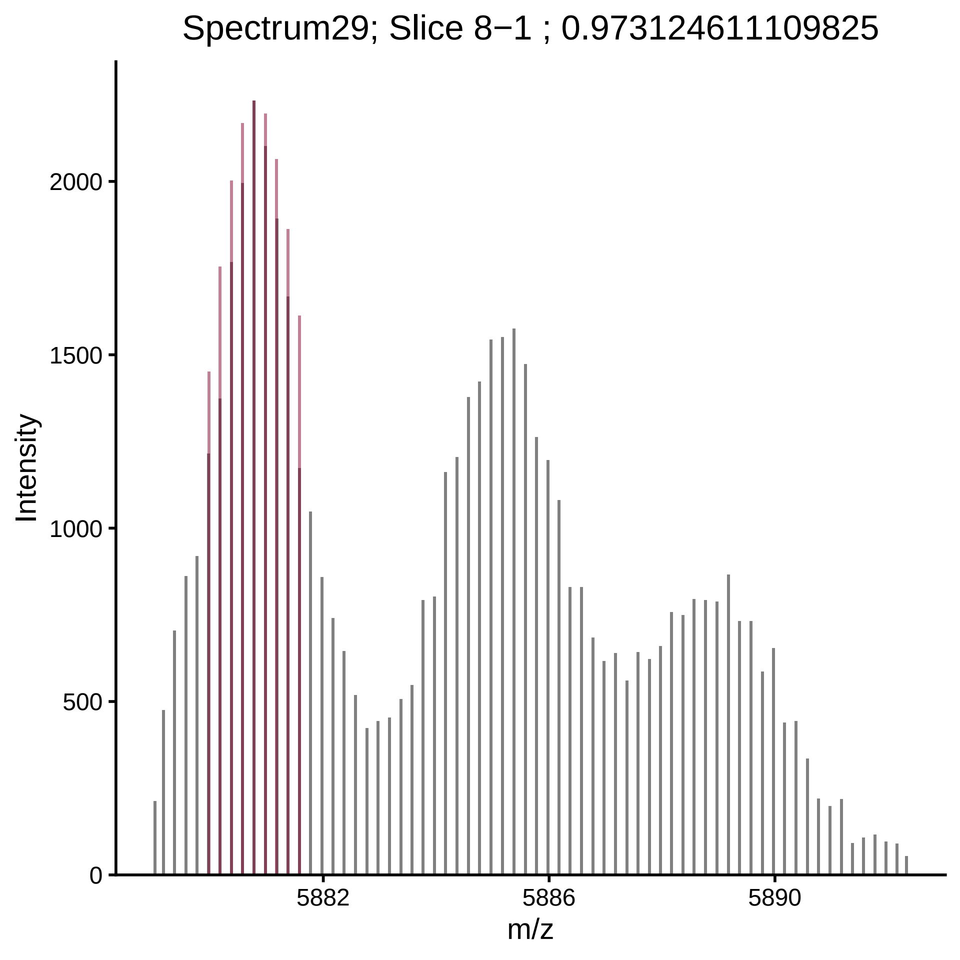

Spectrum29; Slice 8−1 ; 0.973124611109825
Intensity
m/z
0
500
1000
1500
2000
5882
5886
5890

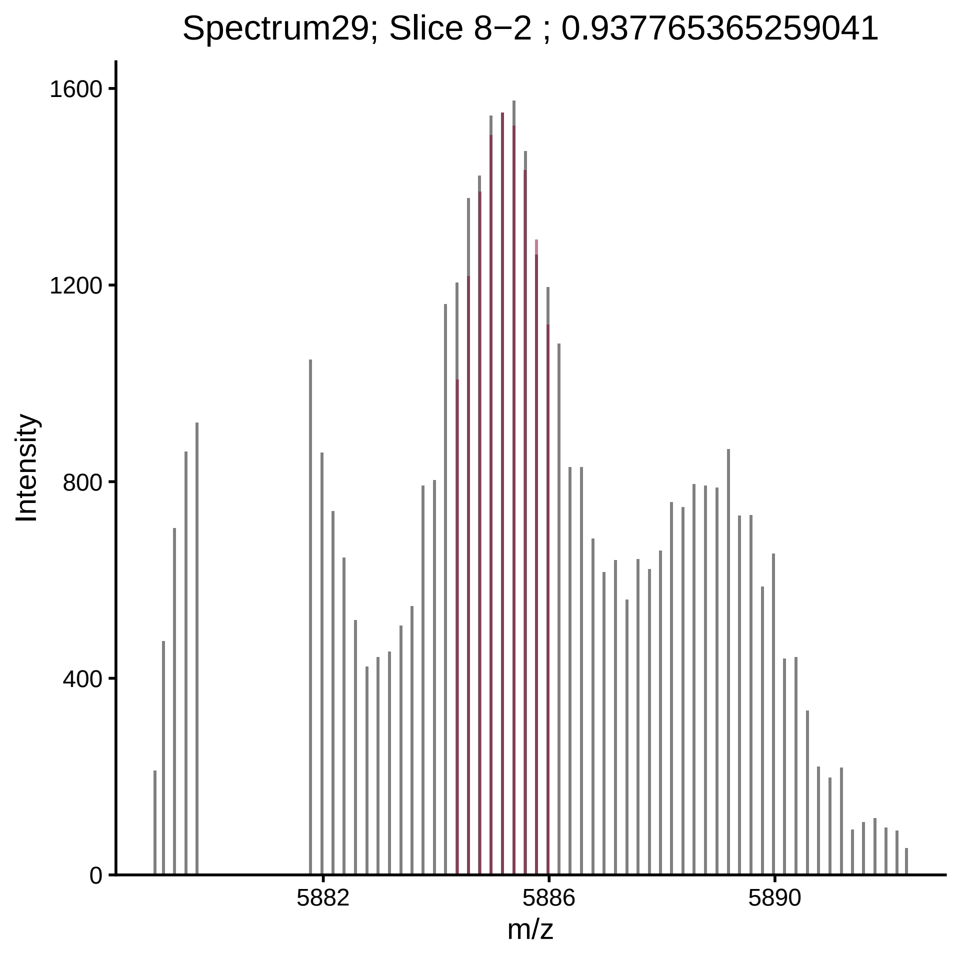

Spectrum29; Slice 8−2 ; 0.937765365259041
Intensity
1600
1200
800
400
0
5882
5886
5890
m/z

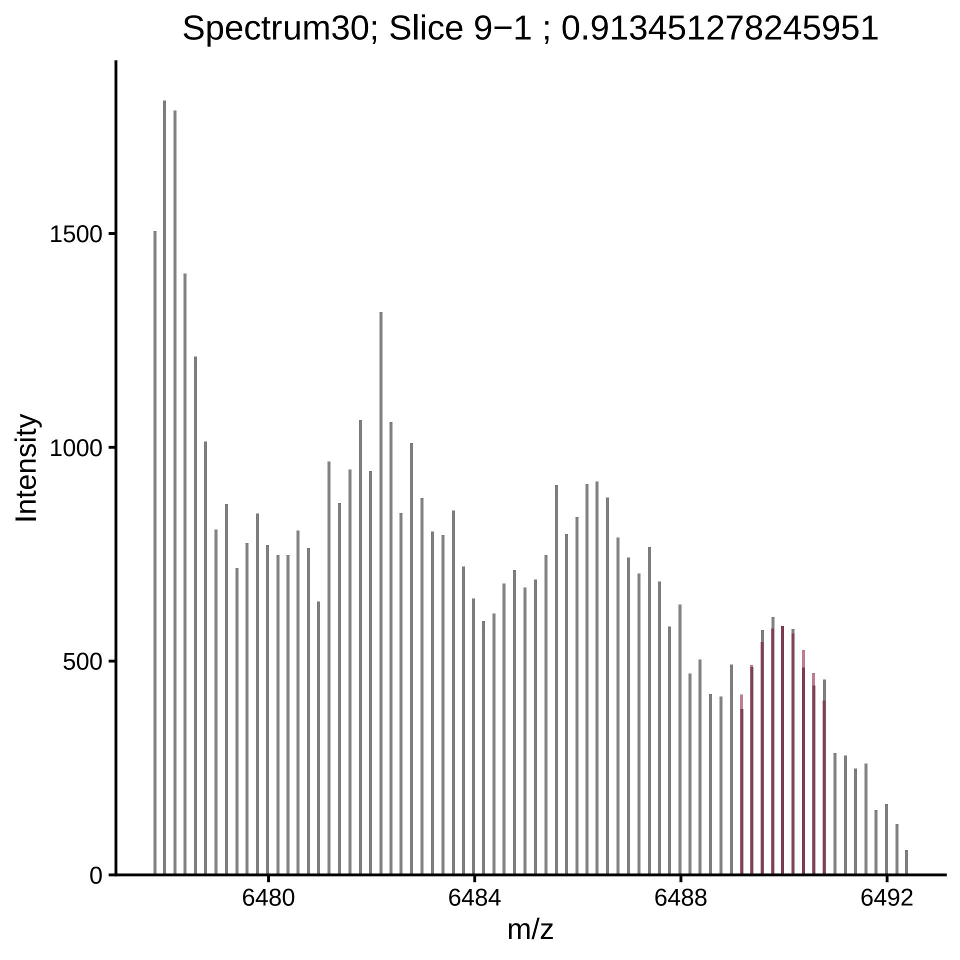

Spectrum30; Slice 9−1 ; 0.913451278245951
Intensity
1500
1000
500
0
6480
6484
6488
6492
m/z

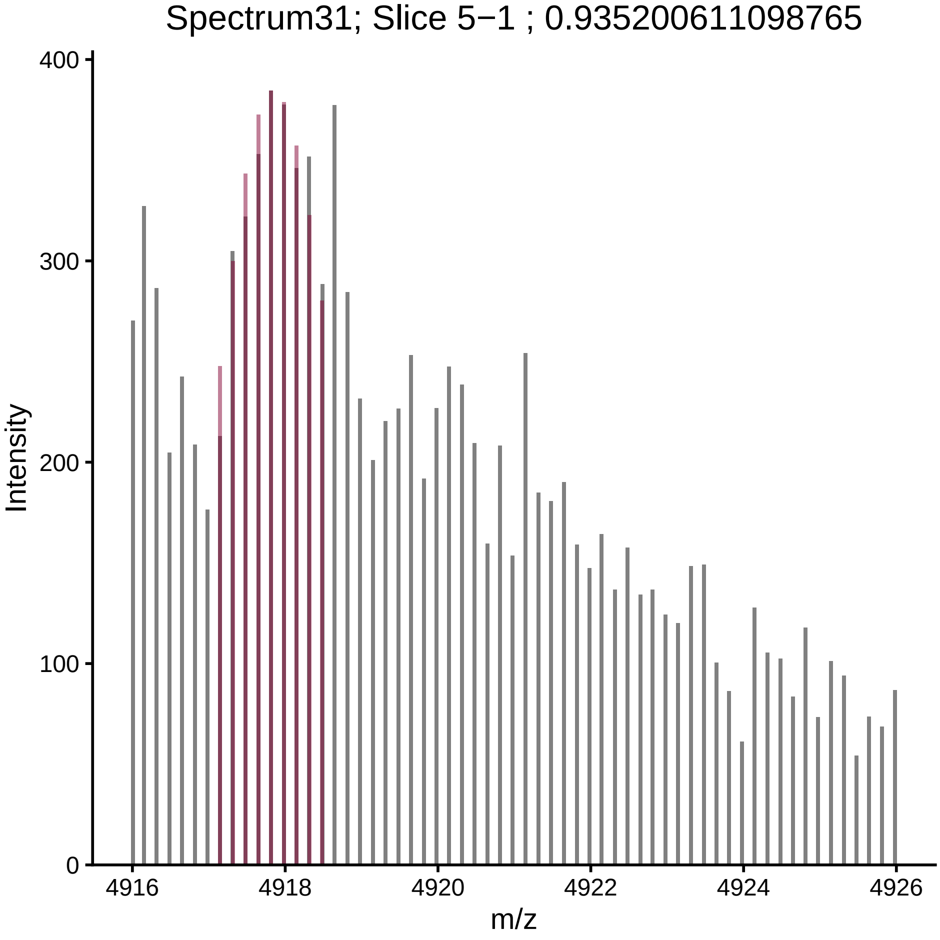

Spectrum31; Slice 5−1 ; 0.935200611098765
Intensity
400
300
200
100
0
4916
4918
4920
4922
4924
4926
m/z

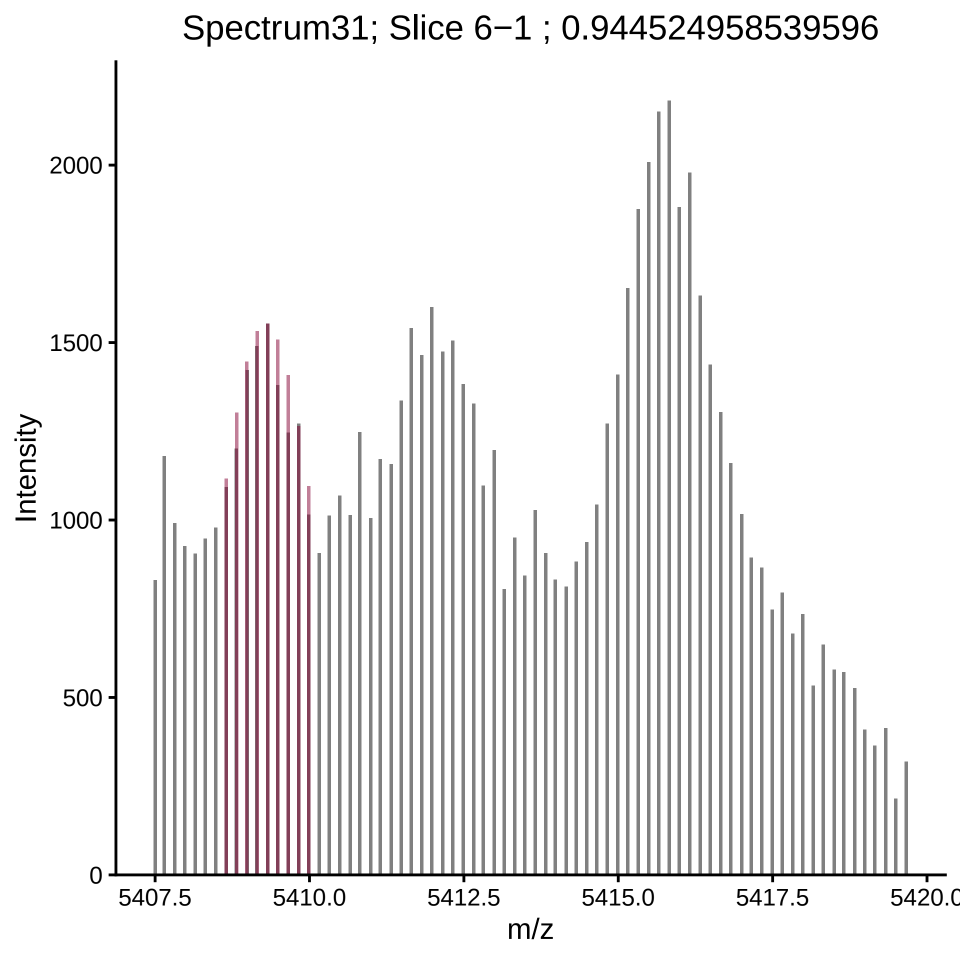

Spectrum31; Slice 6−1 ; 0.944524958539596
Intensity
2000
1500
1000
500
0
5407.5
5410.0
5412.5
5415.0
5417.5
5420.0
m/z

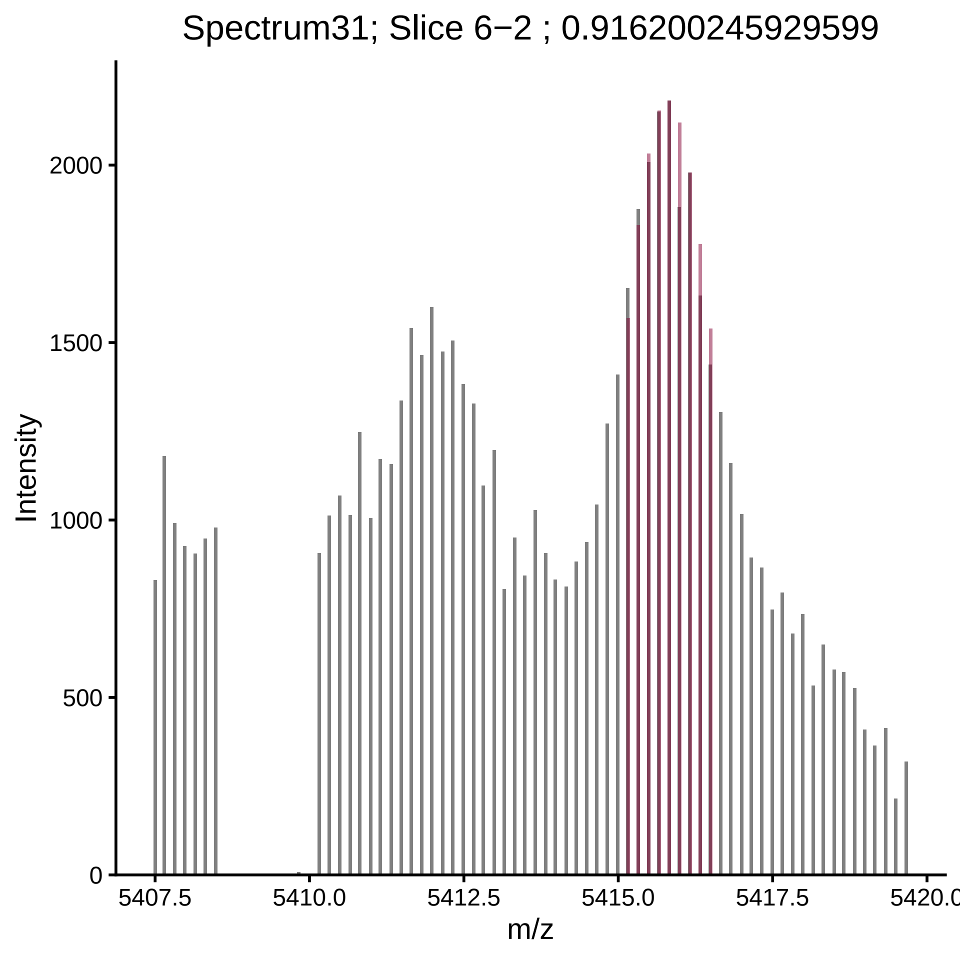

Spectrum31; Slice 6−2 ; 0.916200245929599
Intensity
2000
1500
1000
500
0
5407.5
5410.0
5412.5
5415.0
5417.5
5420.0
m/z

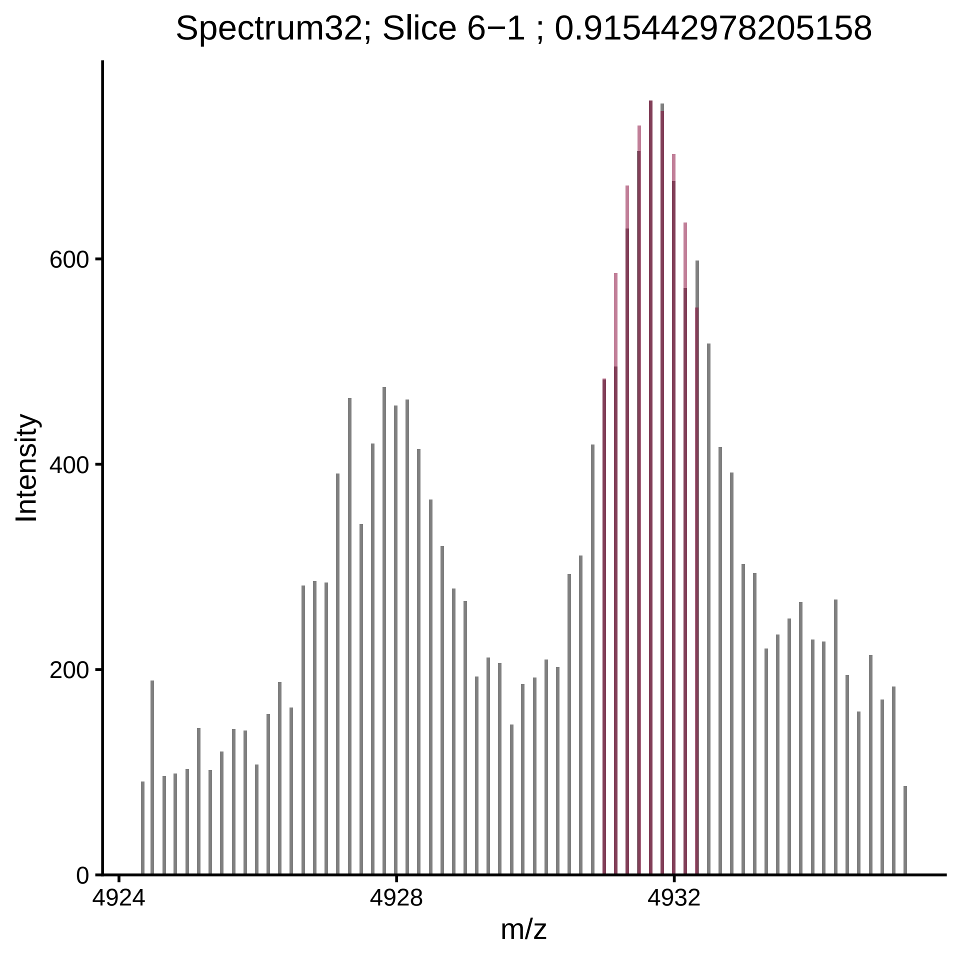

Spectrum32; Slice 6−1 ; 0.915442978205158
Intensity
0
200
400
600
4924
4928
4932
m/z

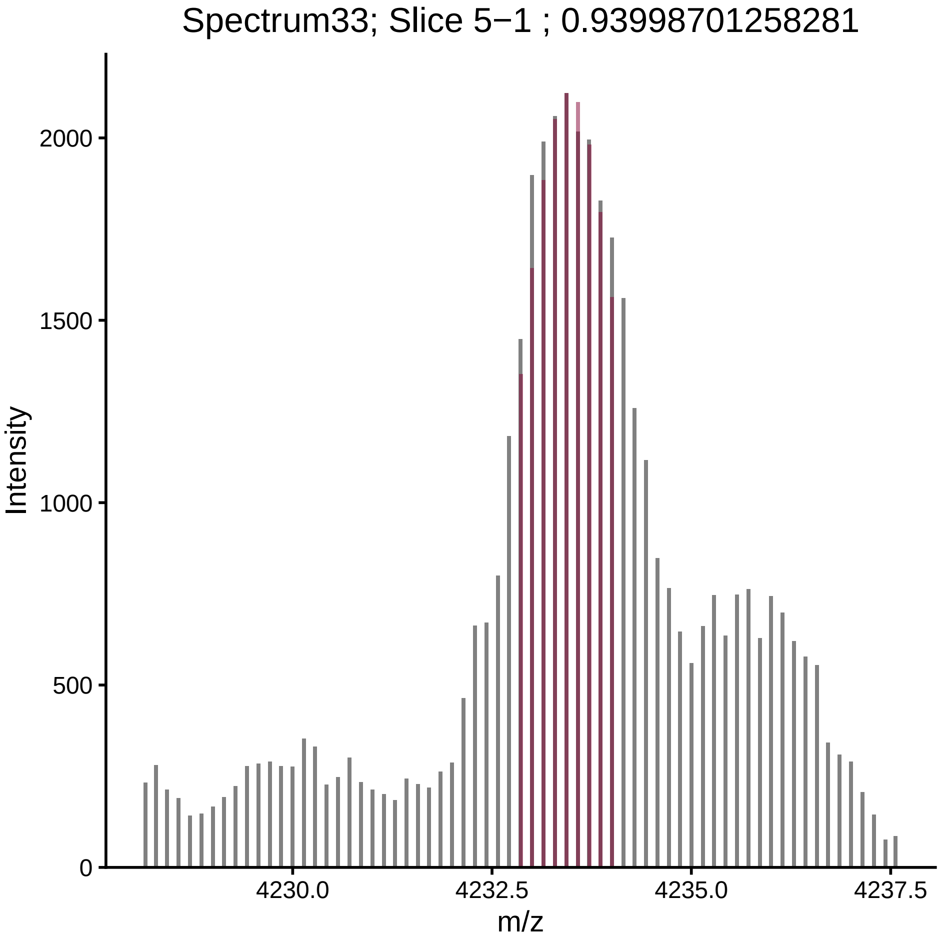
Spectrum33; Slice 5−1 ; 0.93998701258281
Intensity
2000
1500
1000
500
0
4230.0
4232.5
4235.0
4237.5
m/z

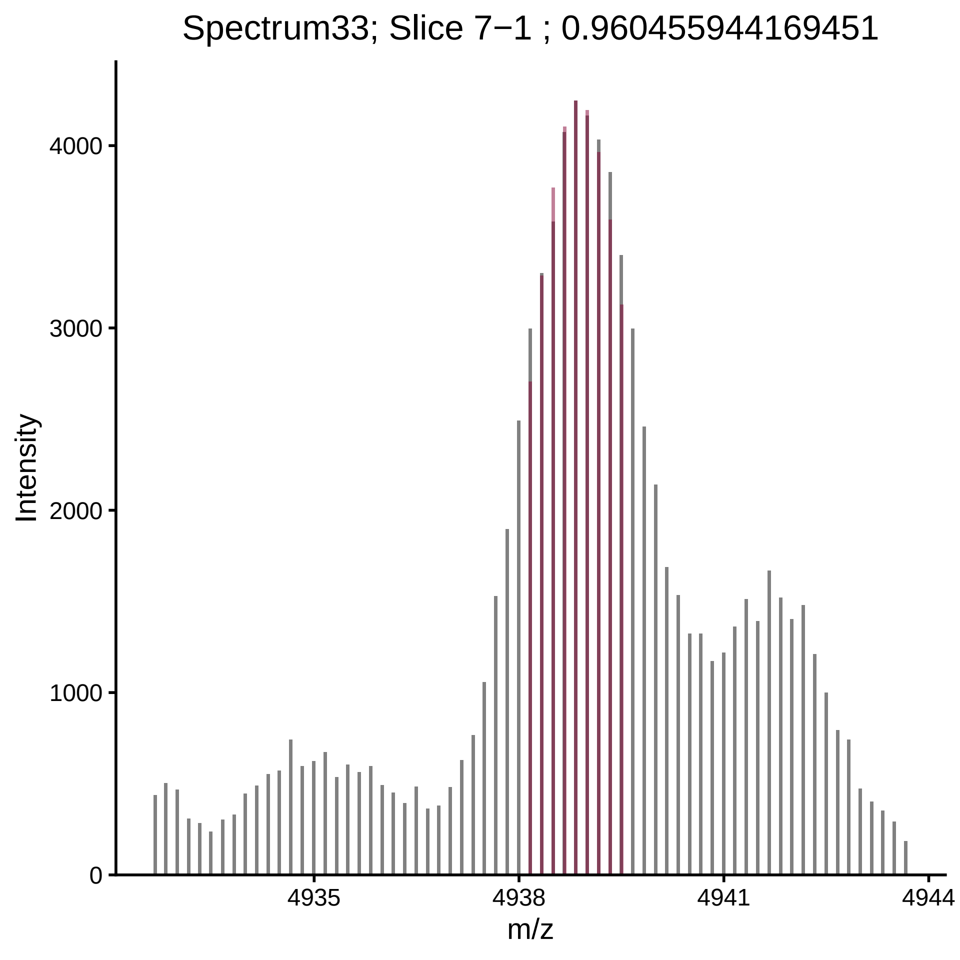

Spectrum33; Slice 7−1 ; 0.960455944169451
Intensity
0
1000
2000
3000
4000
4935
4938
4941
4944
m/z

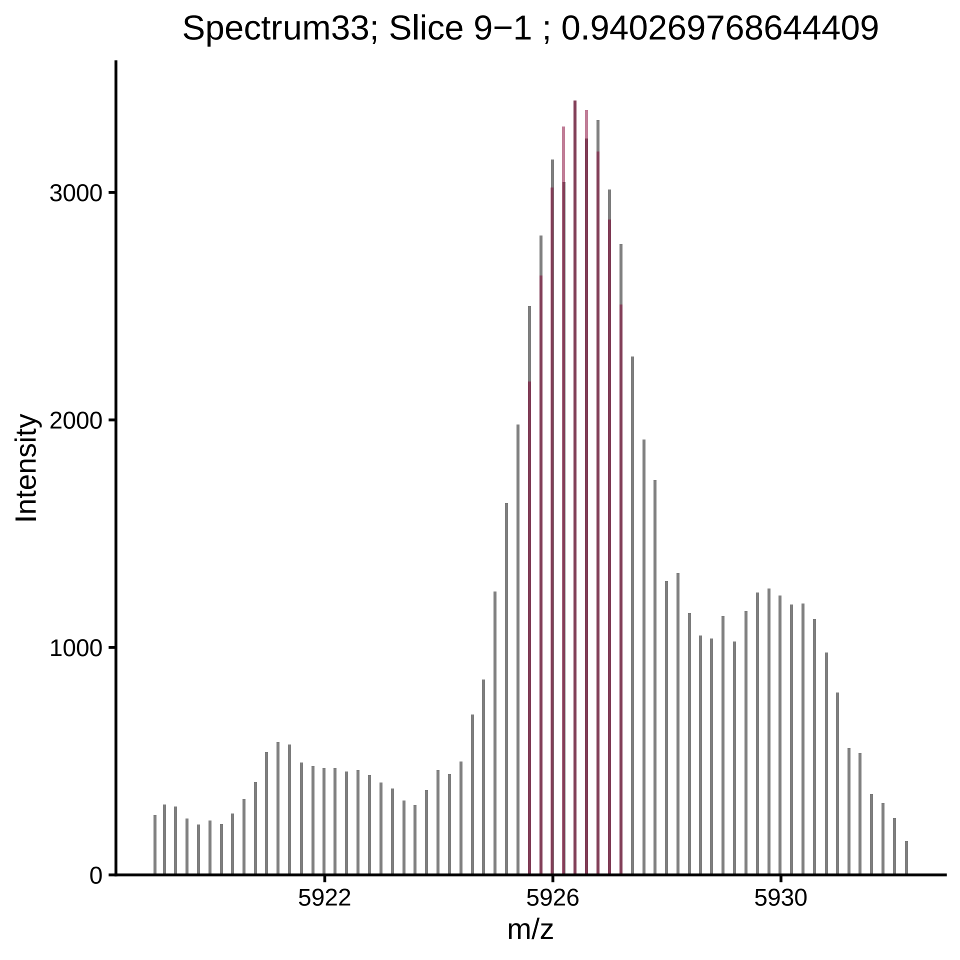

Spectrum33; Slice 9−1 ; 0.940269768644409
Intensity
0
1000
2000
3000
5922
5926
5930
m/z

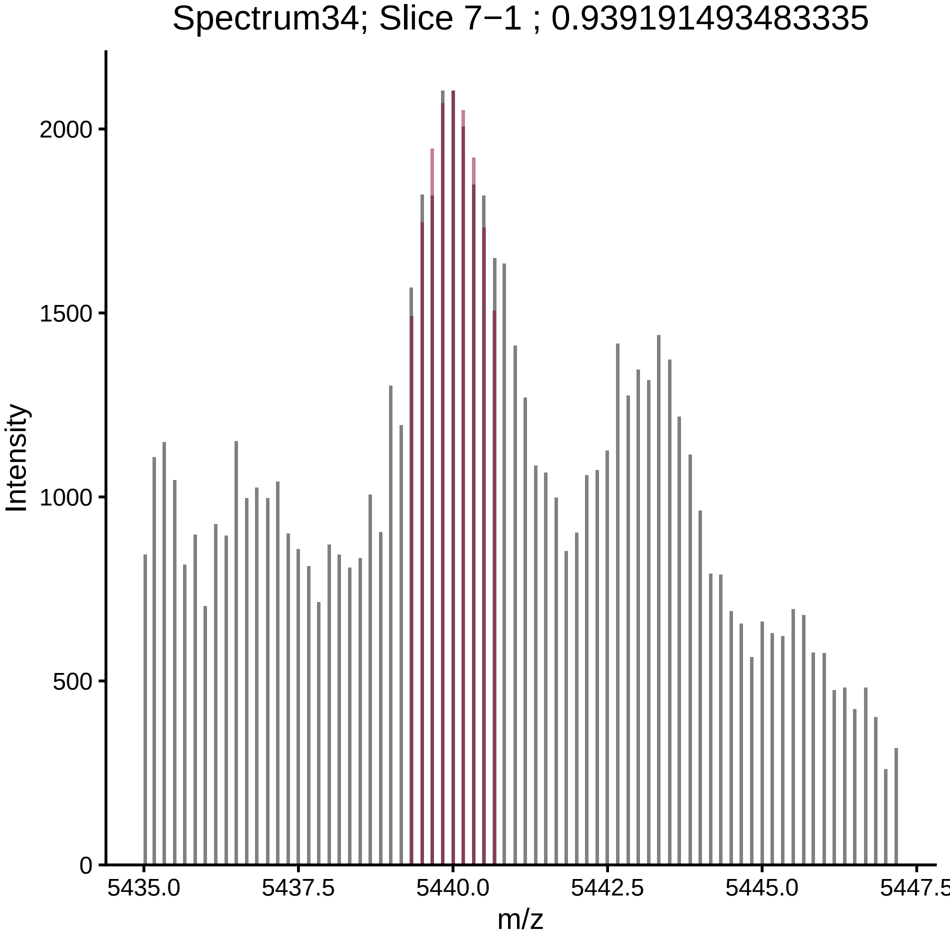

Spectrum34; Slice 7−1 ; 0.939191493483335
Intensity
2000
1500
1000
500
0
5435.0
5437.5
5440.0
5442.5
5445.0
5447.5
m/z

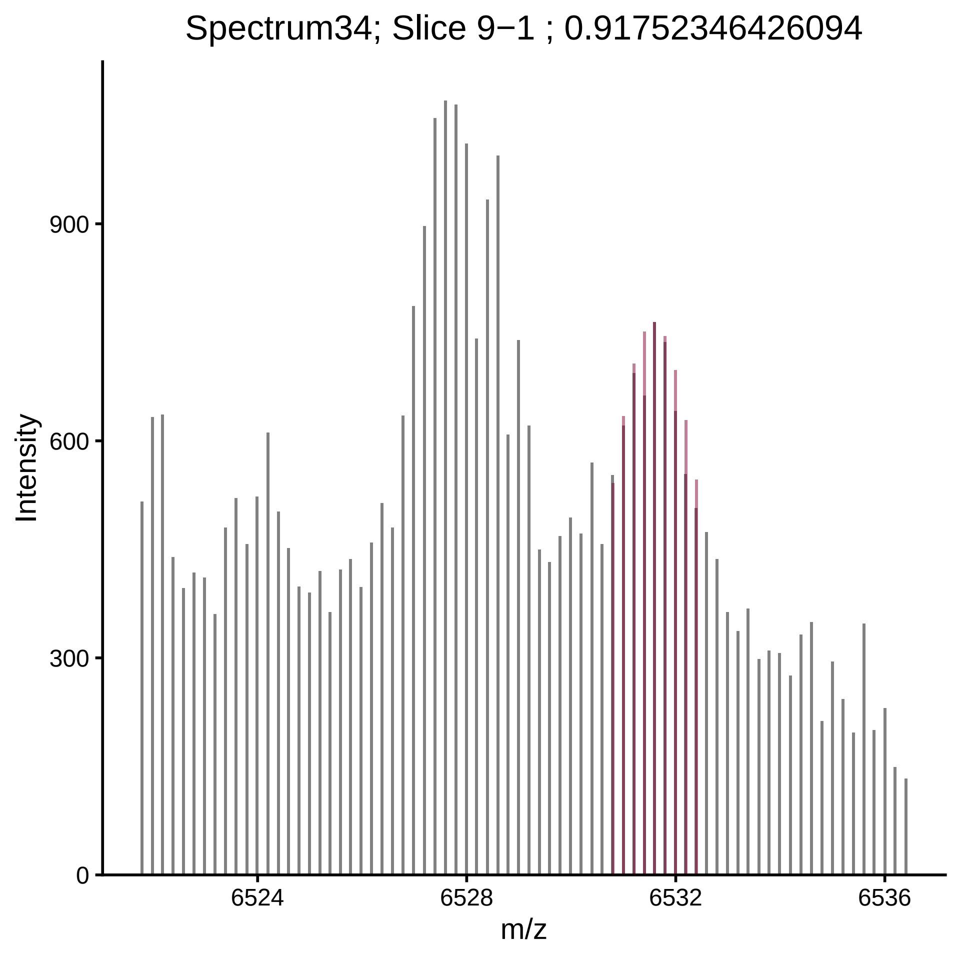
Spectrum34; Slice 9−1 ; 0.91752346426094
Intensity
900
600
300
0
6524
6528
6532
6536
m/z

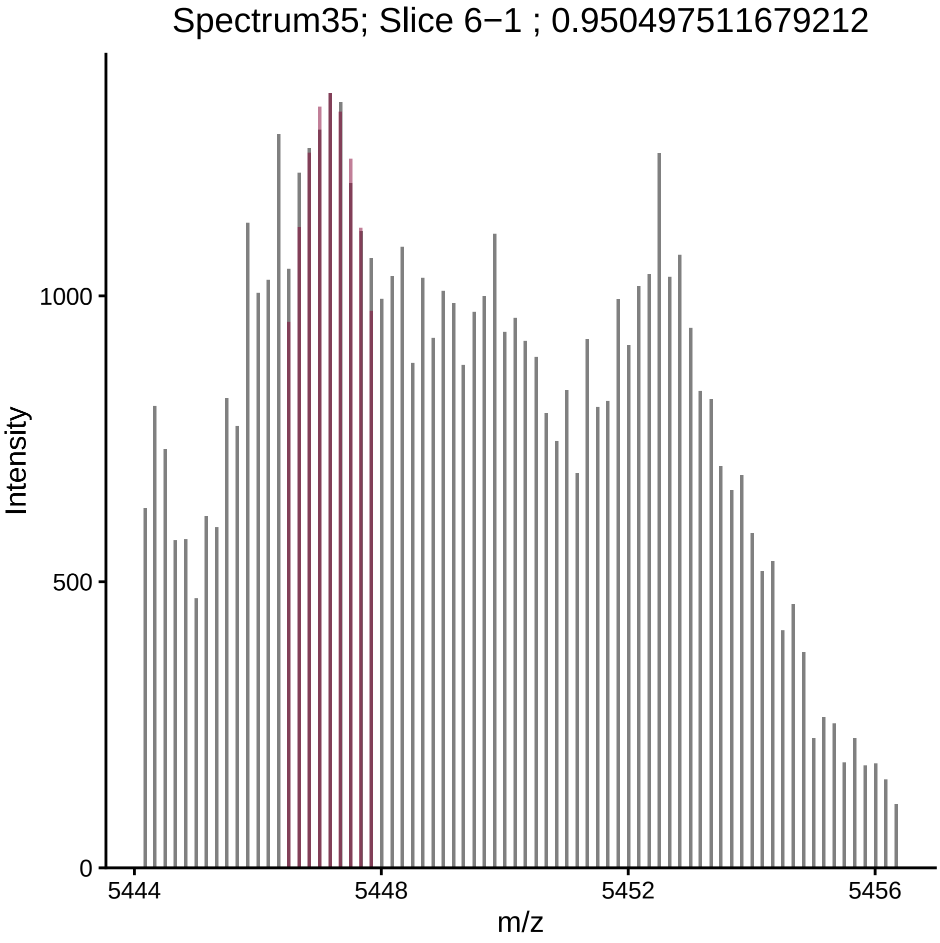

Spectrum35; Slice 6−1 ; 0.950497511679212
Intensity
1000
500
0
5444
5448
5452
5456
m/z

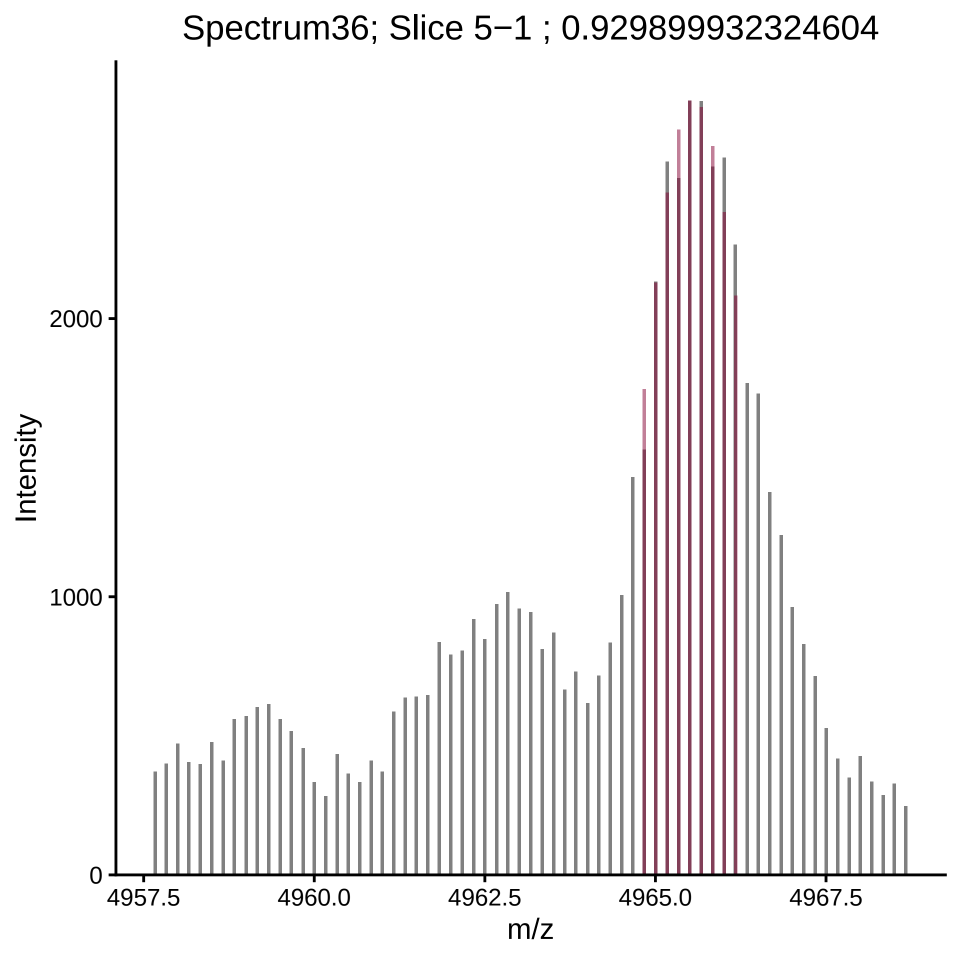

Spectrum36; Slice 5−1 ; 0.929899932324604
Intensity
2000
1000
0
4957.5
4960.0
4962.5
4965.0
4967.5
m/z

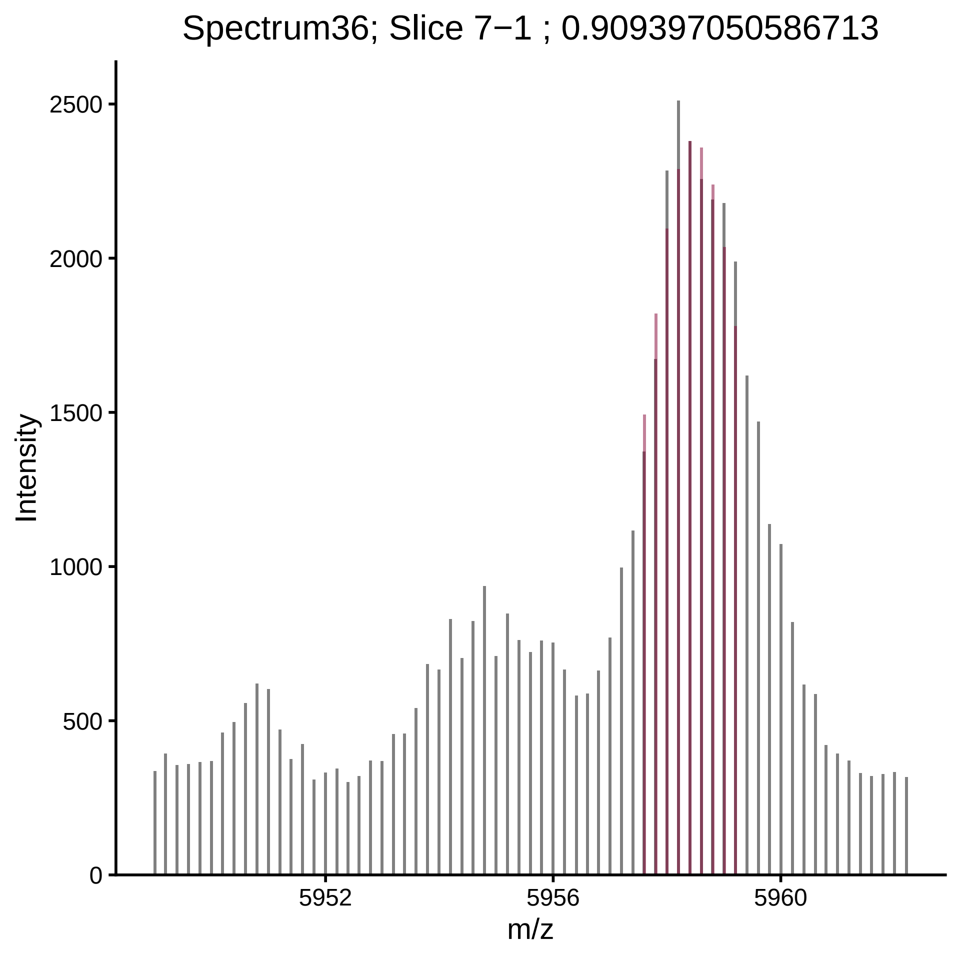
Spectrum36; Slice 7−1 ; 0.909397050586713
Intensity
2500
2000
1500
1000
500
0
5952
5956
5960
m/z

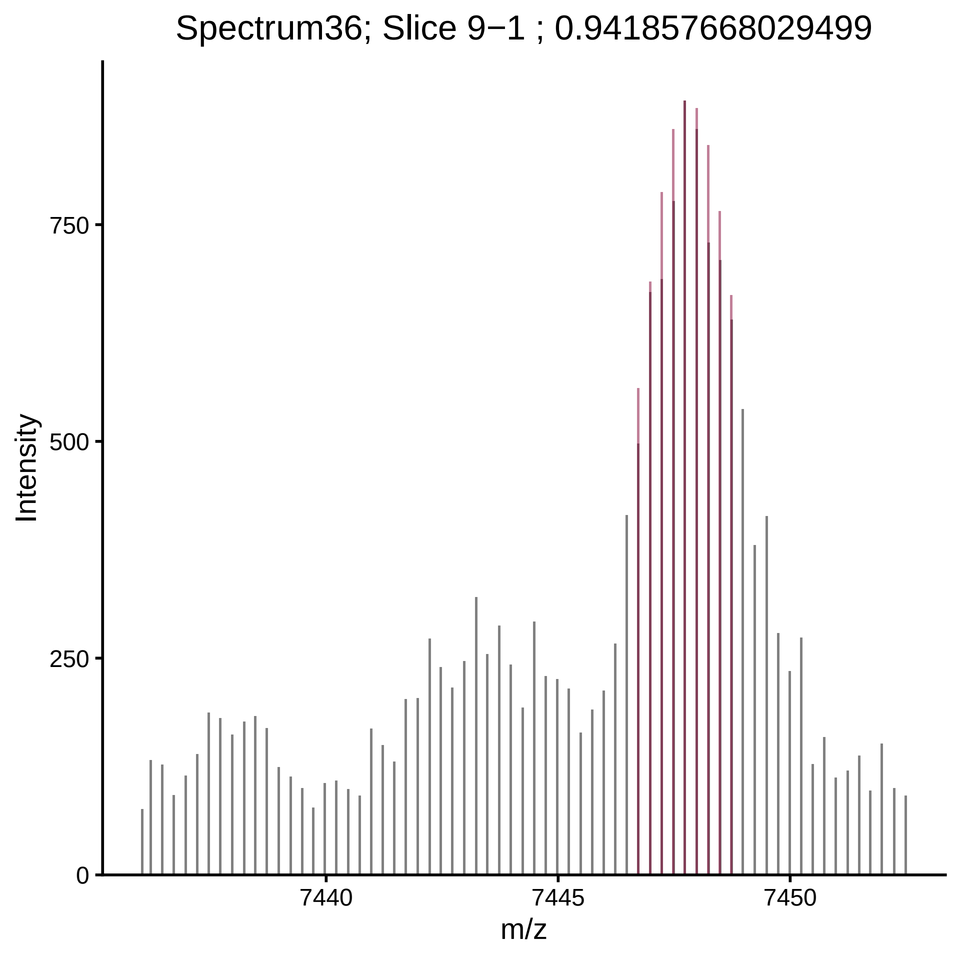

Spectrum36; Slice 9−1 ; 0.941857668029499
Intensity
750
500
250
0
7440
7445
7450
m/z

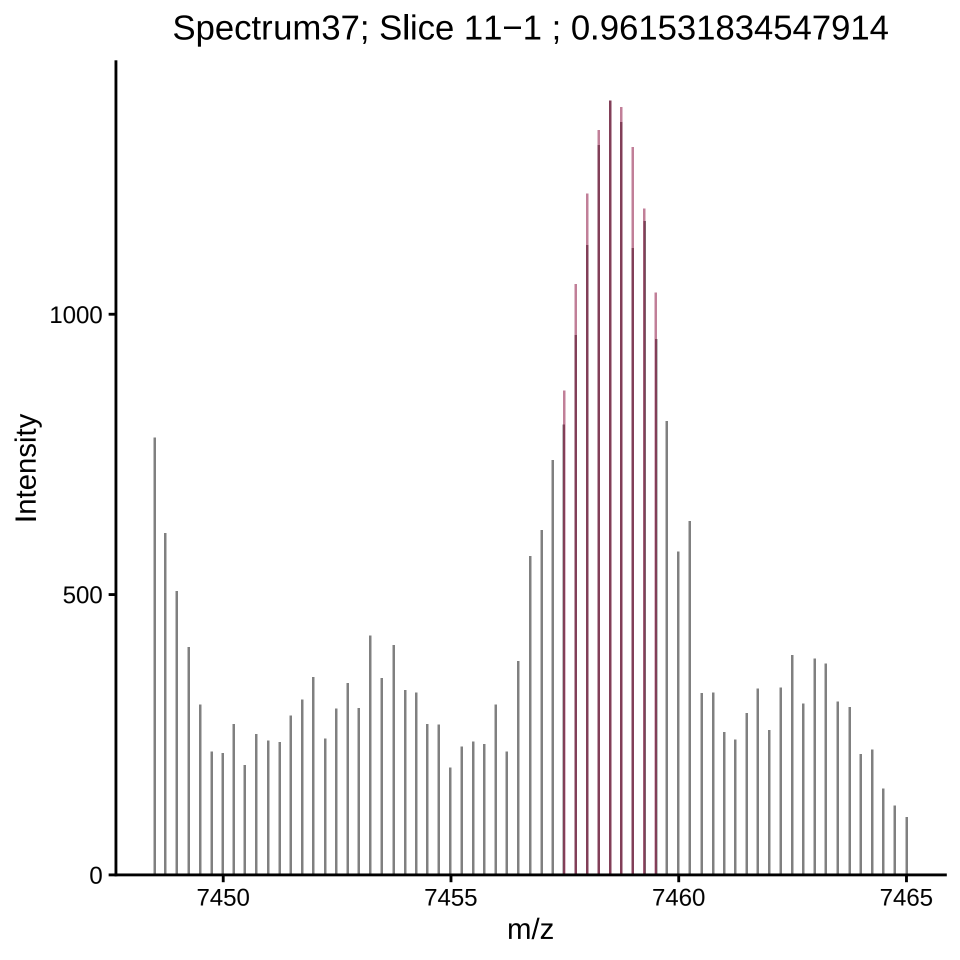

Spectrum37; Slice 11−1 ; 0.961531834547914
Intensity
1000
500
0
7450
7455
7460
7465
m/z

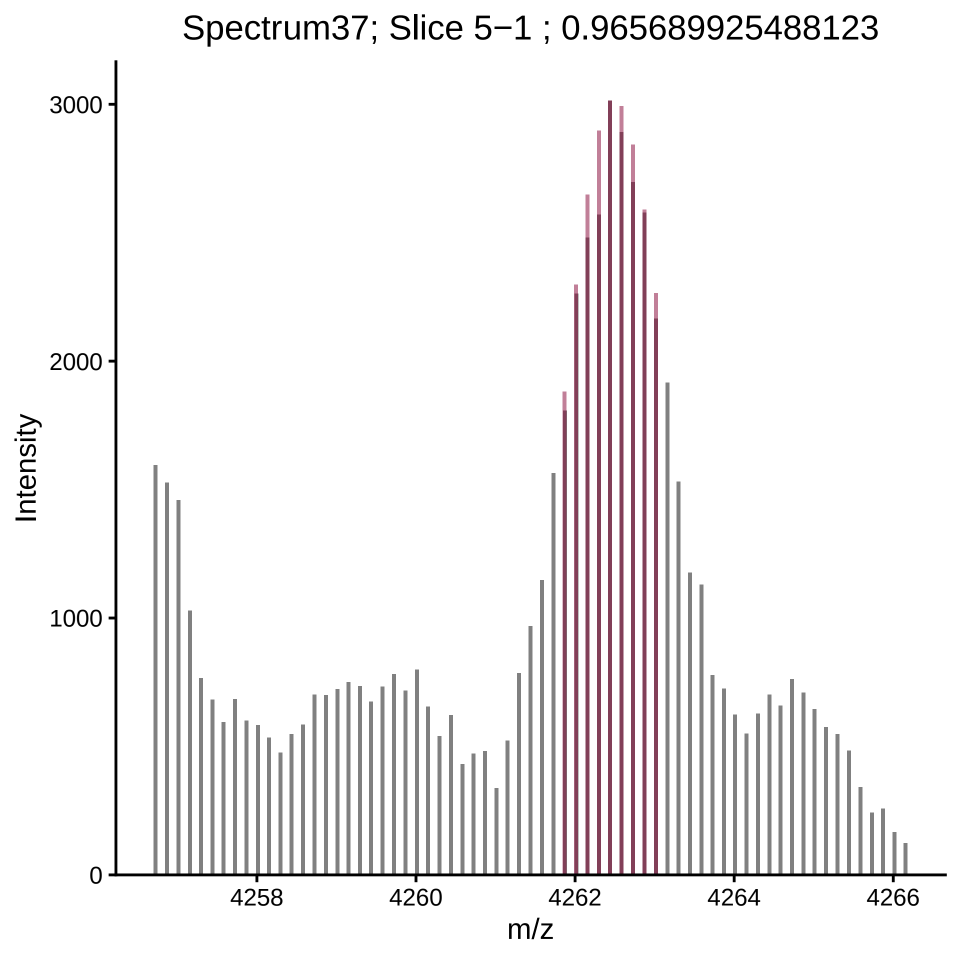
Spectrum37; Slice 5−1 ; 0.965689925488123
Intensity
3000
2000
1000
0
4258
4260
4262
4264
4266
m/z

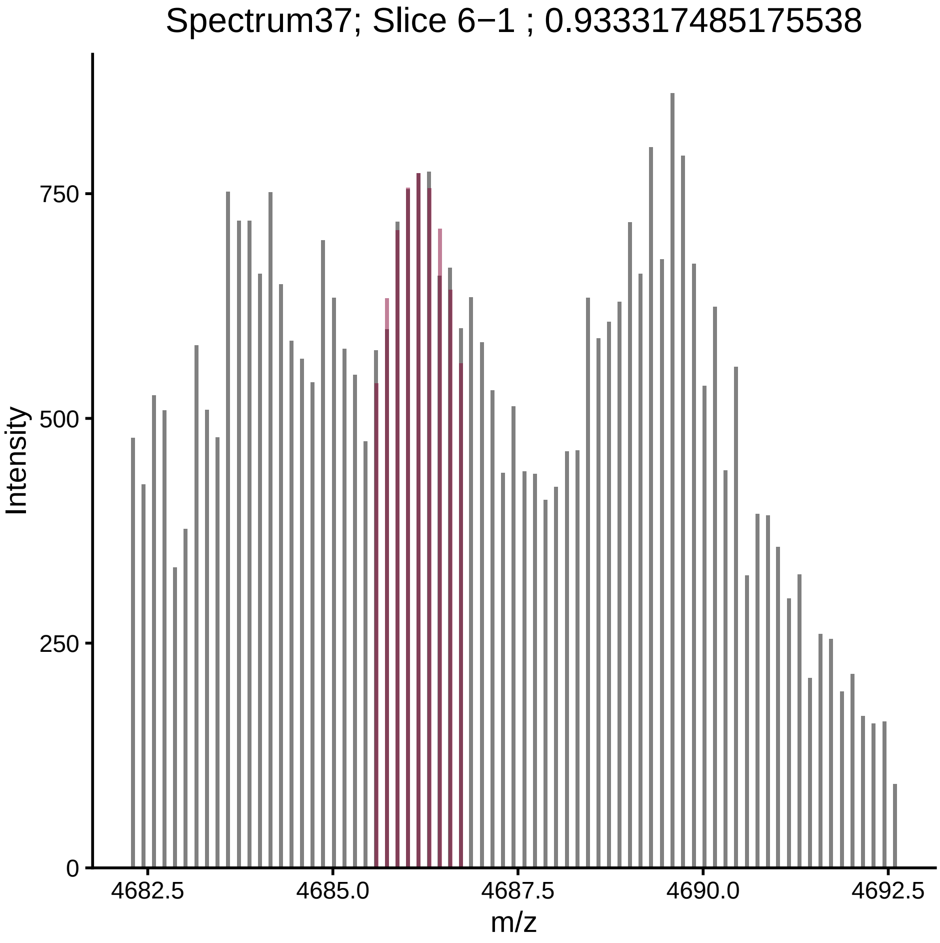
Spectrum37; Slice 6−1 ; 0.933317485175538
Intensity
0
250
500
750
m/z
4682.5
4685.0
4687.5
4690.0
4692.5

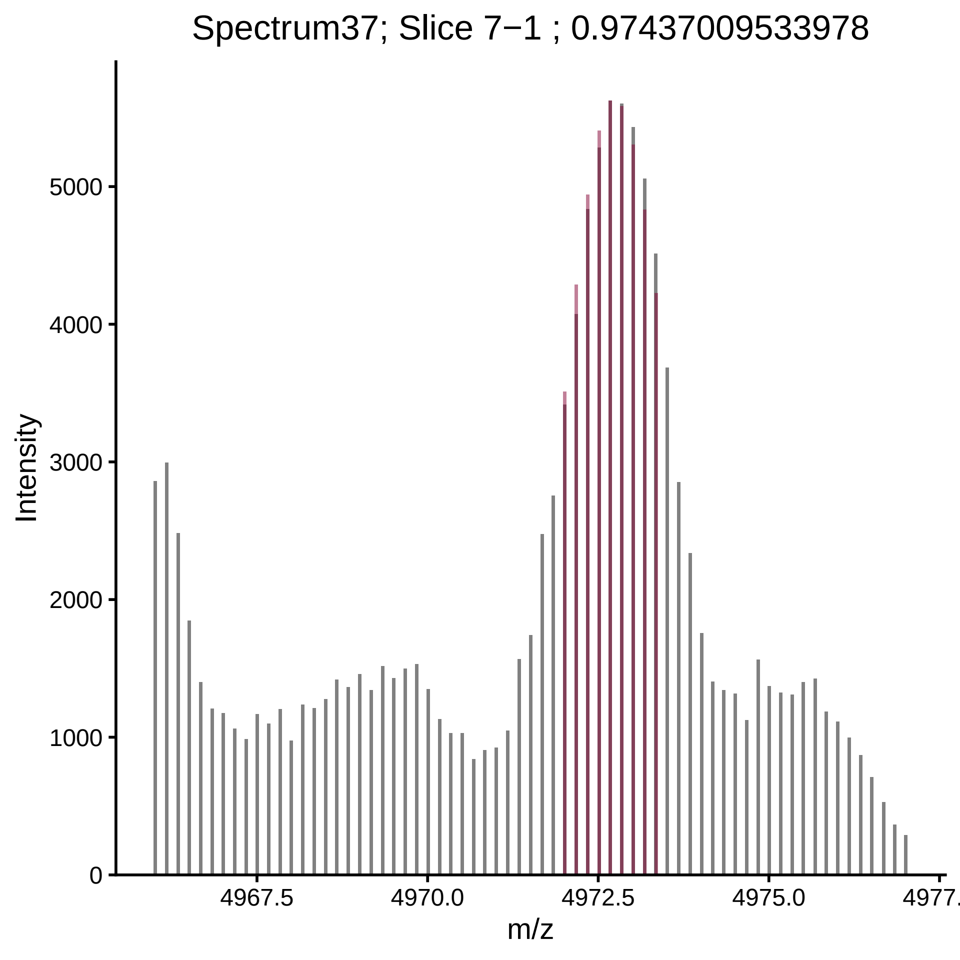

Spectrum37; Slice 7−1 ; 0.97437009533978
Intensity
0
1000
2000
3000
4000
5000
4967.5
4970.0
4972.5
4975.0
4977.
m/z

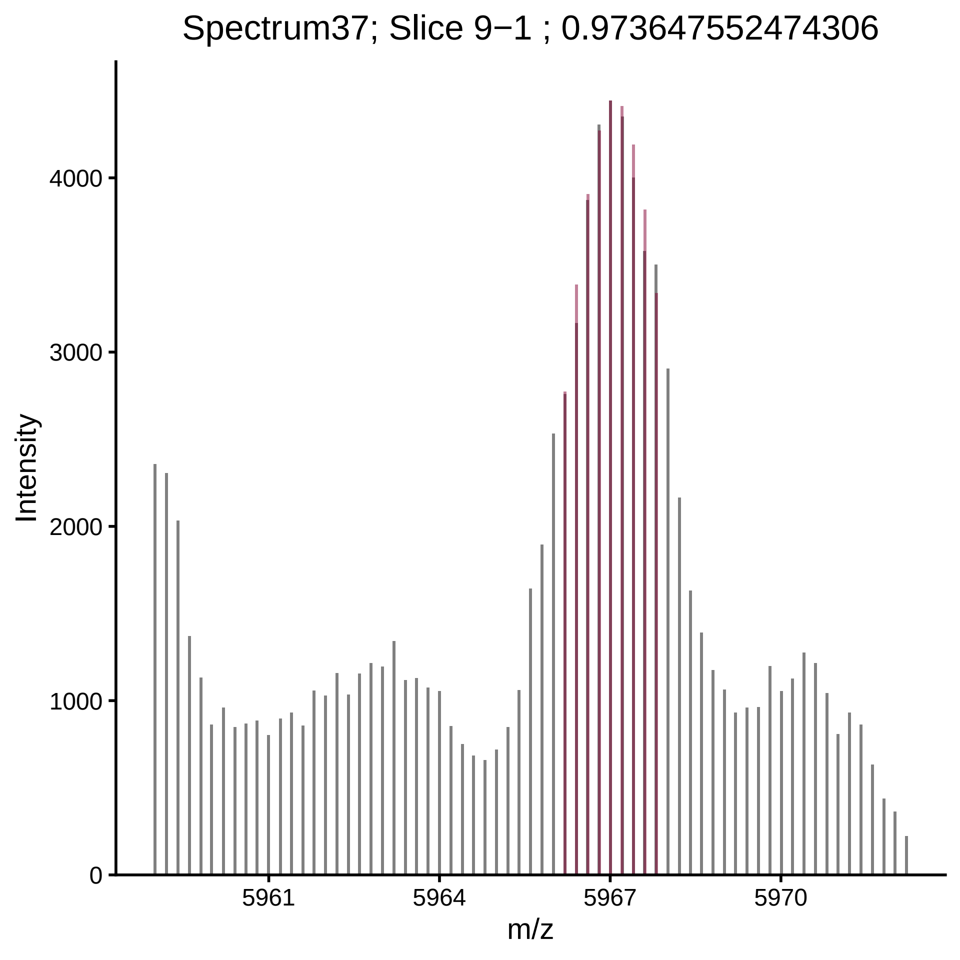

Spectrum37; Slice 9−1 ; 0.973647552474306
Intensity
4000
3000
2000
1000
0
5961
5964
5967
5970
m/z

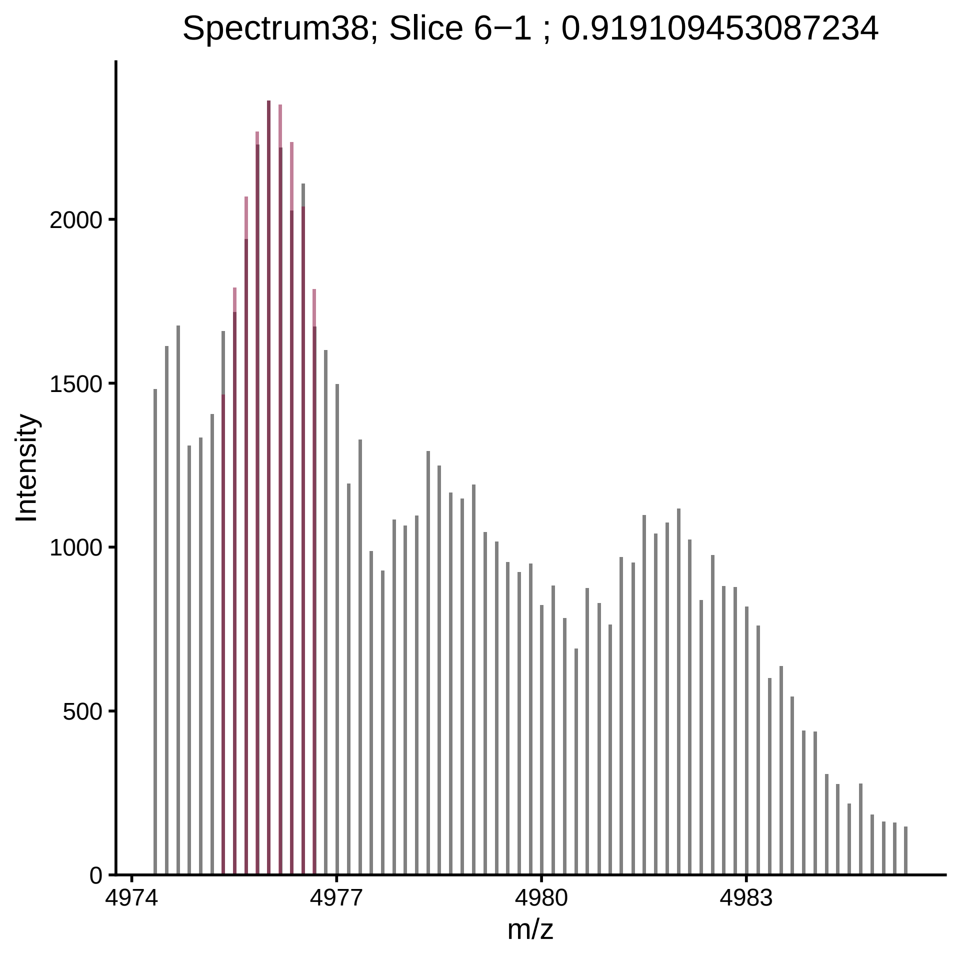

Spectrum38; Slice 6−1 ; 0.919109453087234
Intensity
0
500
1000
1500
2000
4974
4977
4980
4983
m/z

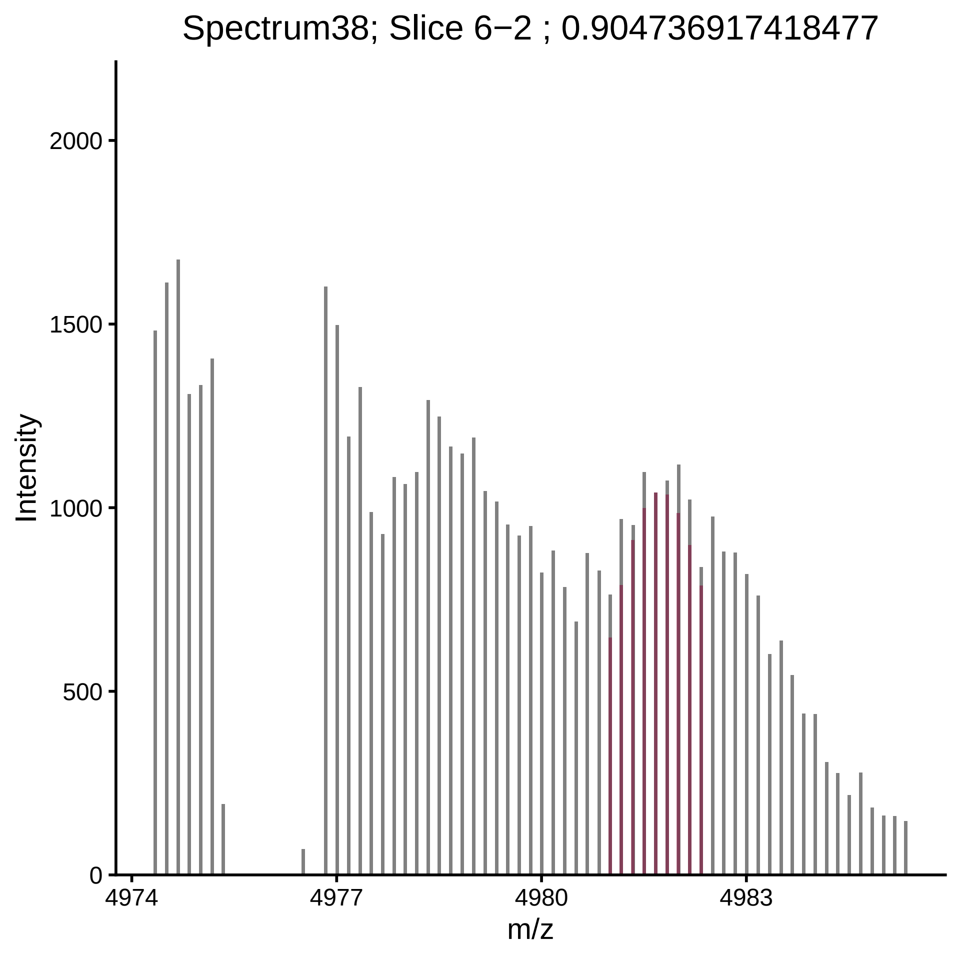
Spectrum38; Slice 6−2 ; 0.904736917418477
Intensity
2000
1500
1000
500
0
4974
4977
4980
4983
m/z

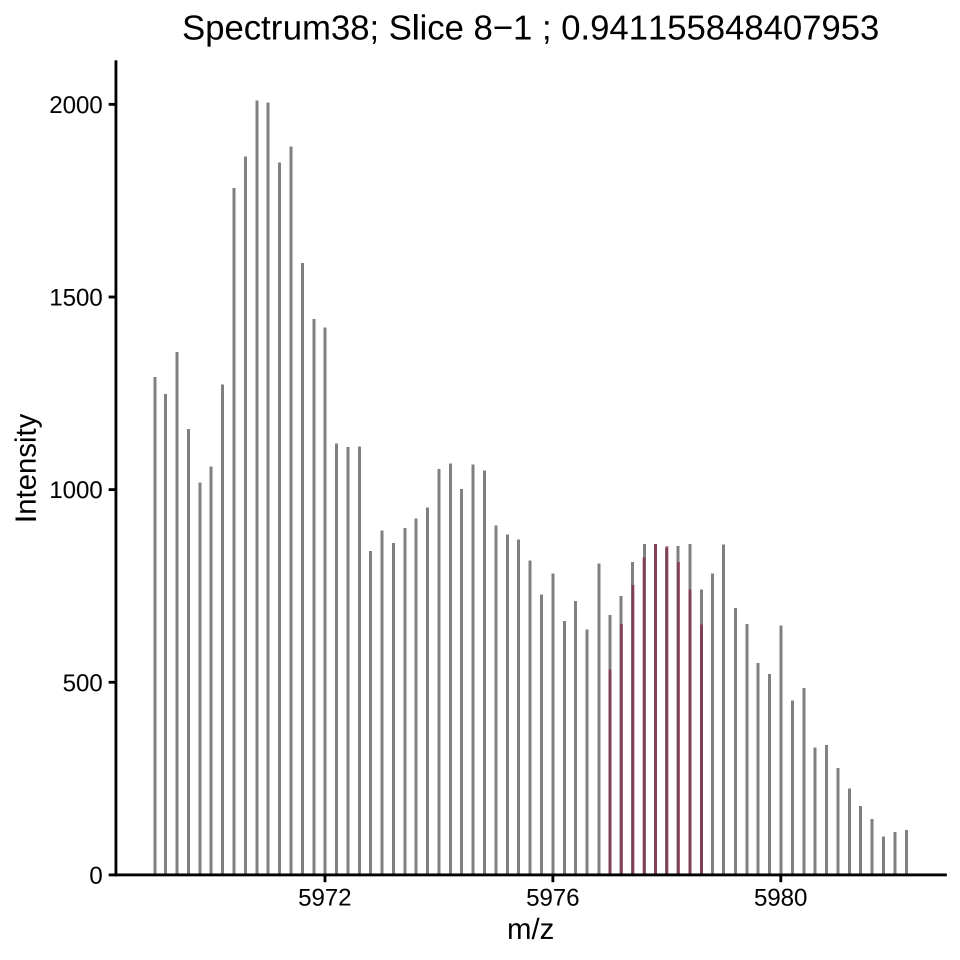
Spectrum38; Slice 8−1 ; 0.941155848407953
Intensity
2000
1500
1000
500
0
5972
5976
5980
m/z

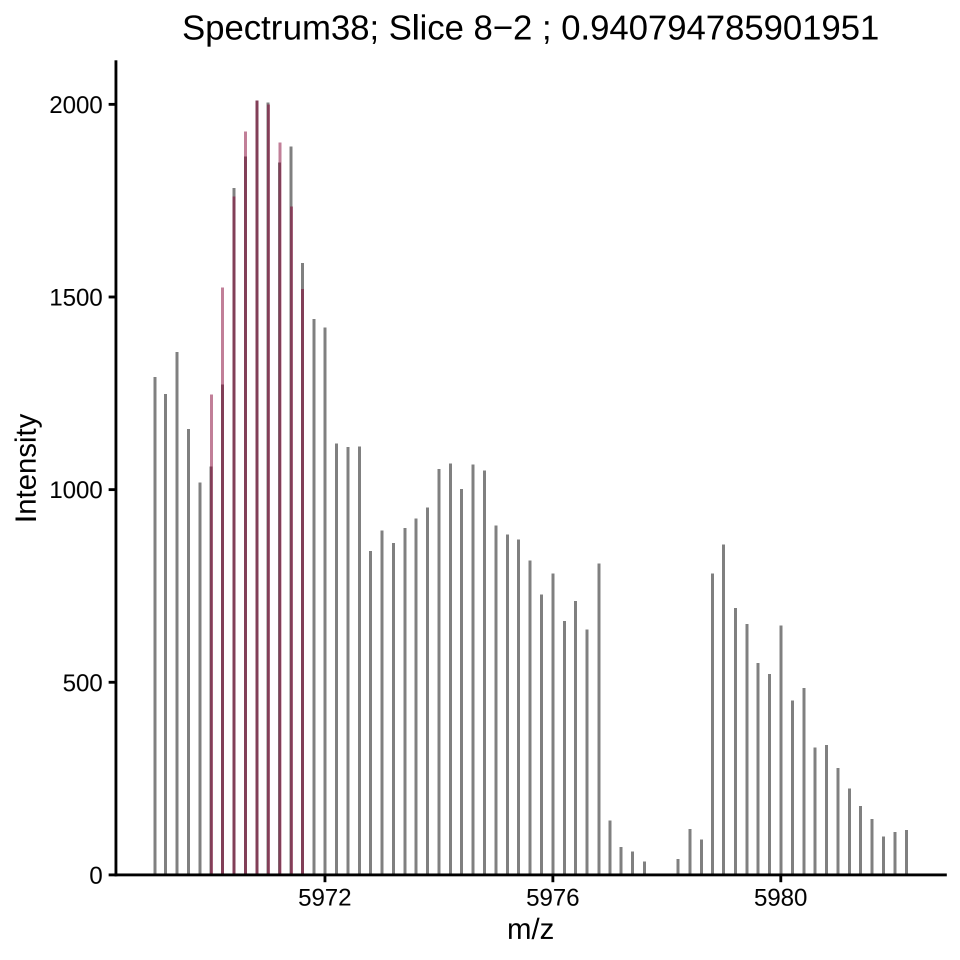

Spectrum38; Slice 8−2 ; 0.940794785901951
Intensity
2000
1500
1000
500
0
5972
5976
5980
m/z

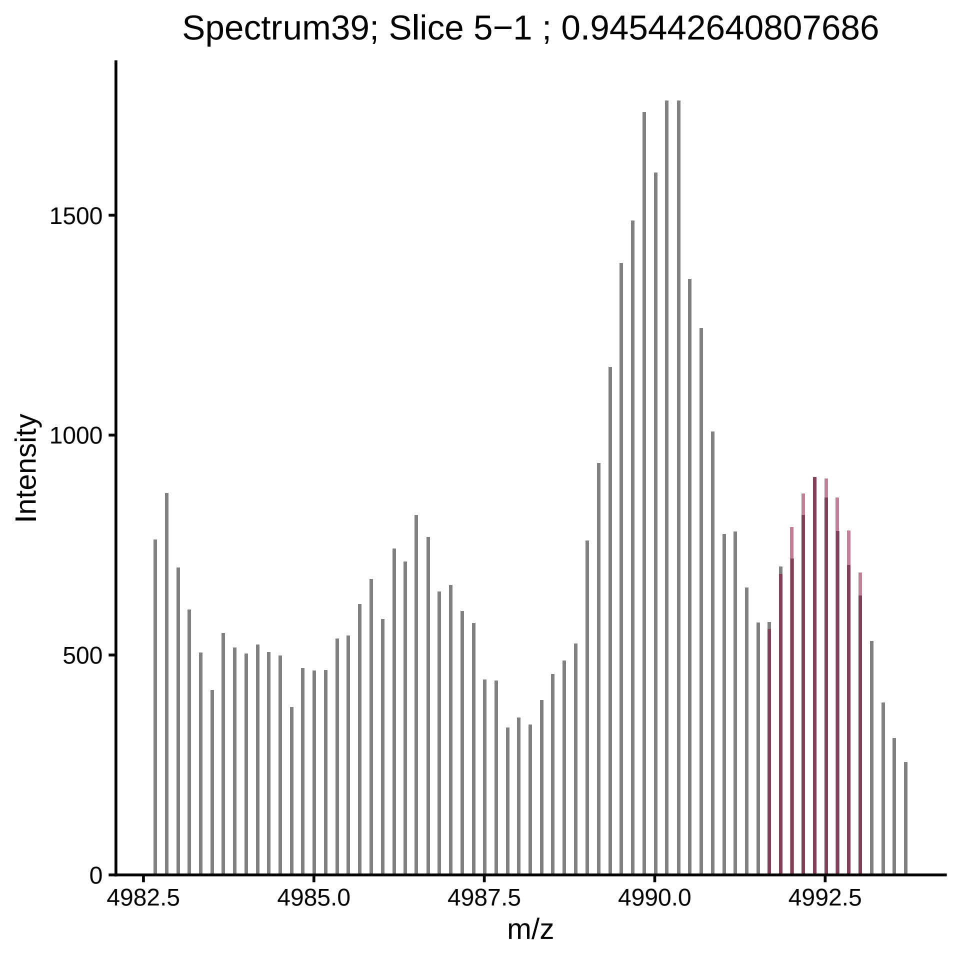

Spectrum39; Slice 5−1 ; 0.945442640807686
Intensity
1500
1000
500
0
4982.5
4985.0
4987.5
4990.0
4992.5
m/z

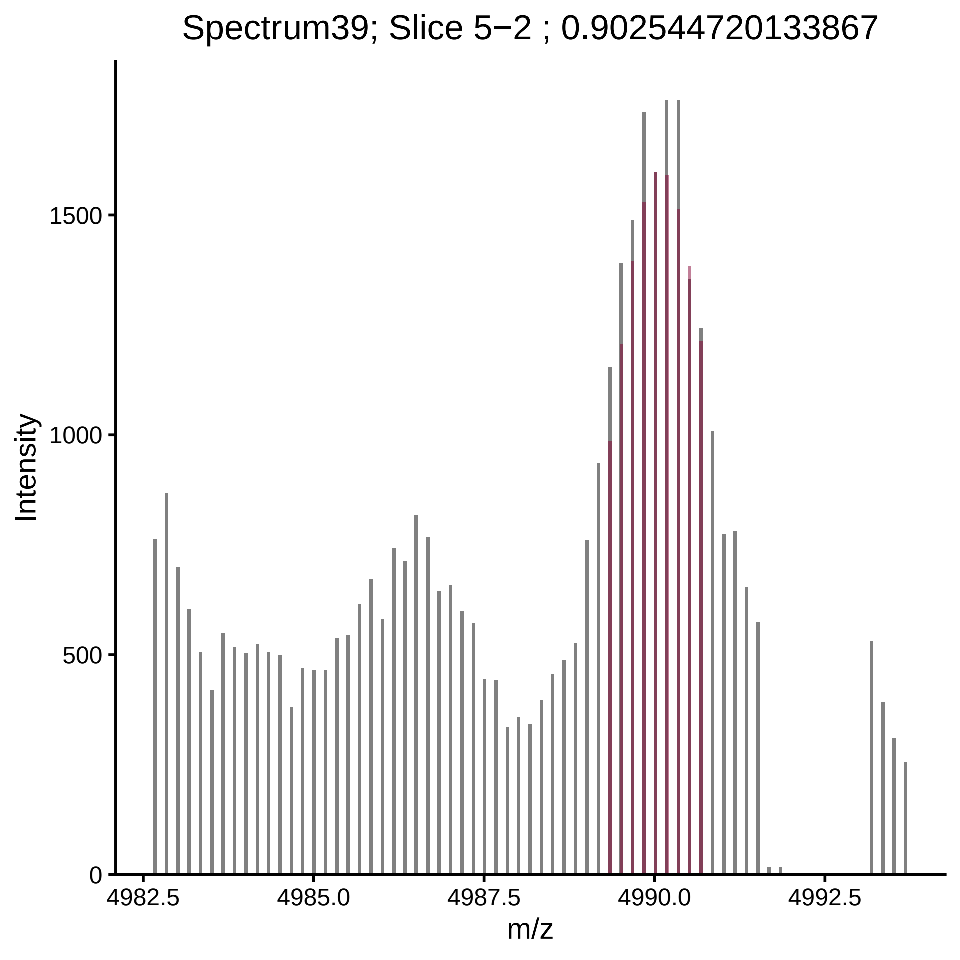

Spectrum39; Slice 5−2 ; 0.902544720133867
Intensity
1500
1000
500
0
4982.5
4985.0
4987.5
4990.0
4992.5
m/z

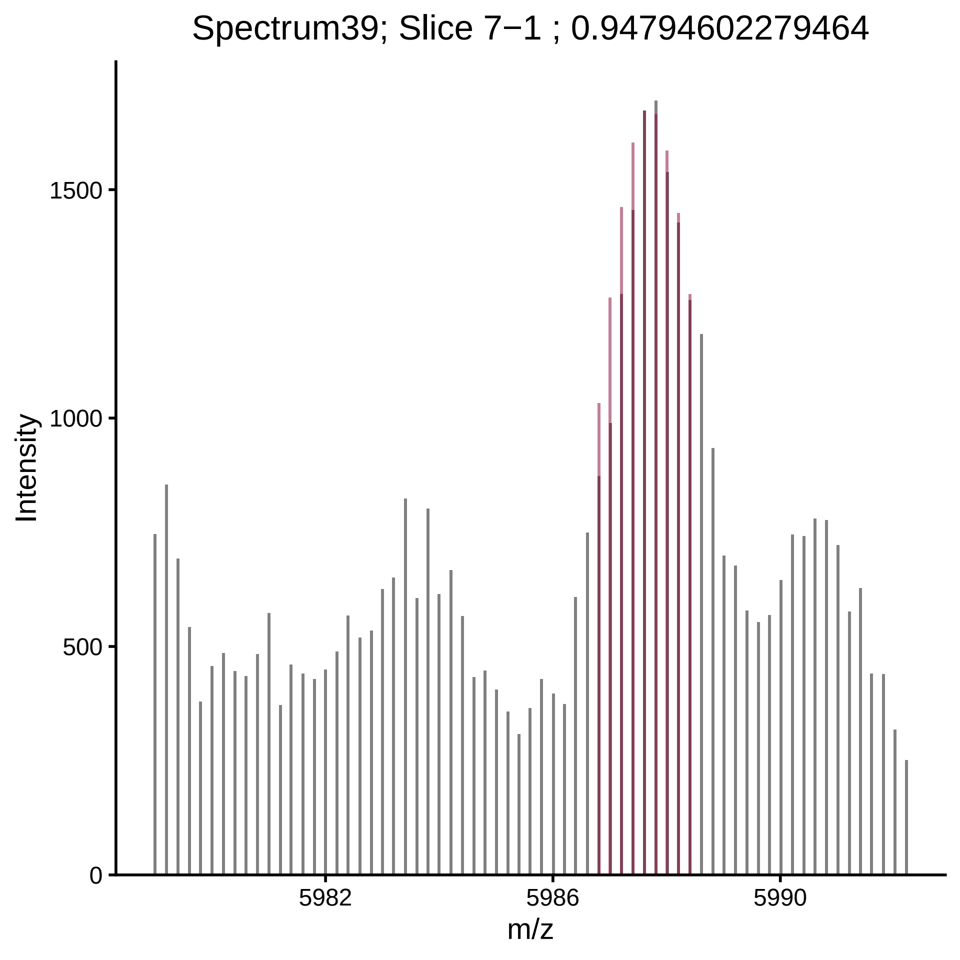

Spectrum39; Slice 7−1 ; 0.94794602279464
Intensity
1500
1000
500
0
5982
5986
5990
m/z

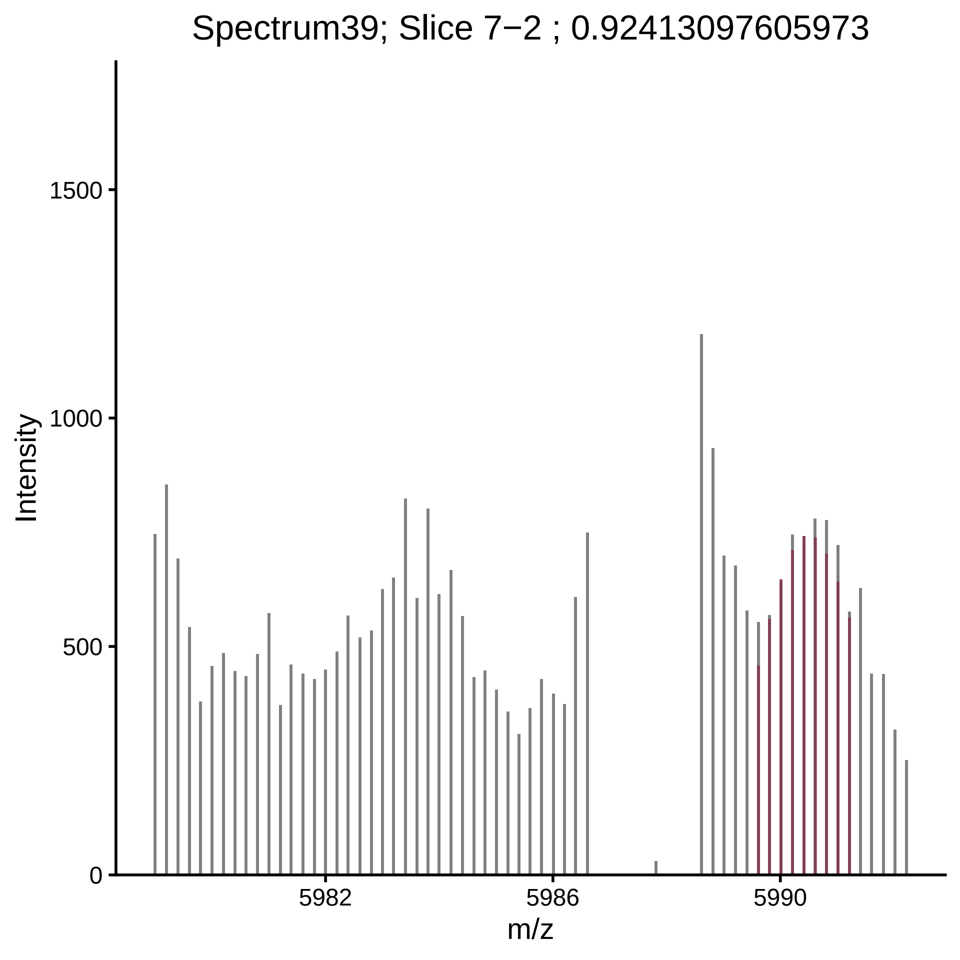

Spectrum39; Slice 7−2 ; 0.92413097605973
Intensity
1500
1000
500
0
5982
5986
5990
m/z

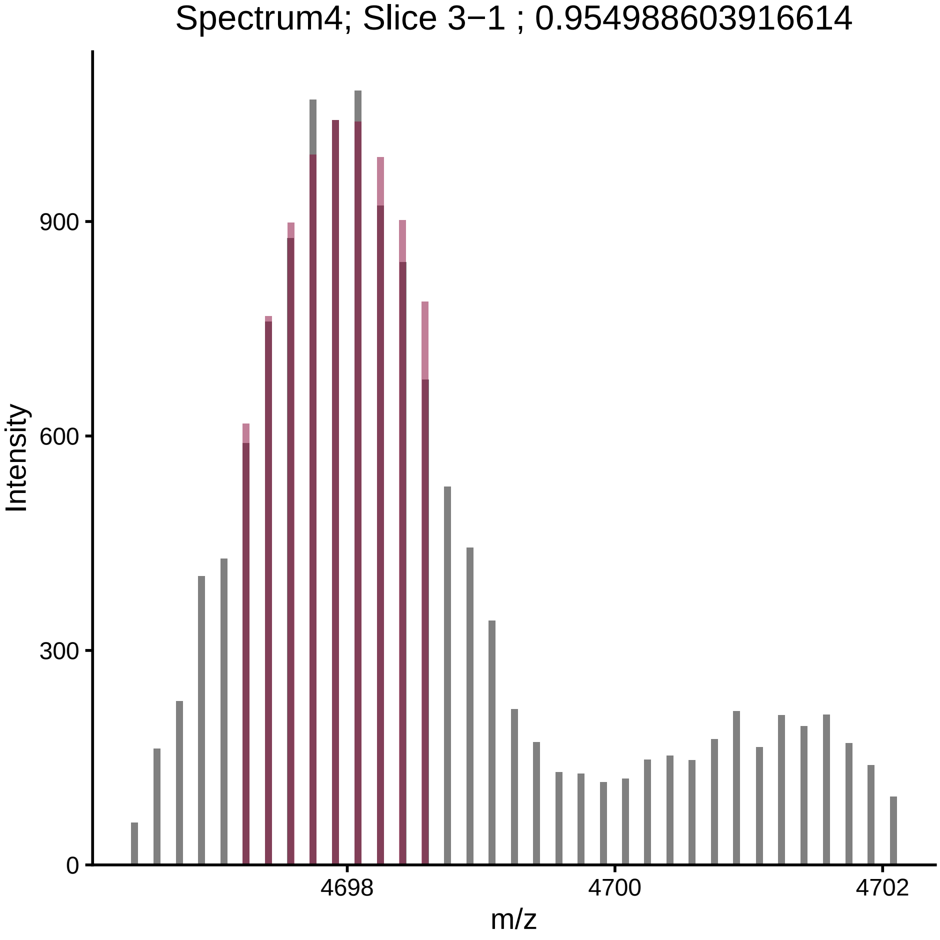
Spectrum4; Slice 3−1 ; 0.954988603916614
Intensity
900
600
300
0
4698
4700
4702
m/z

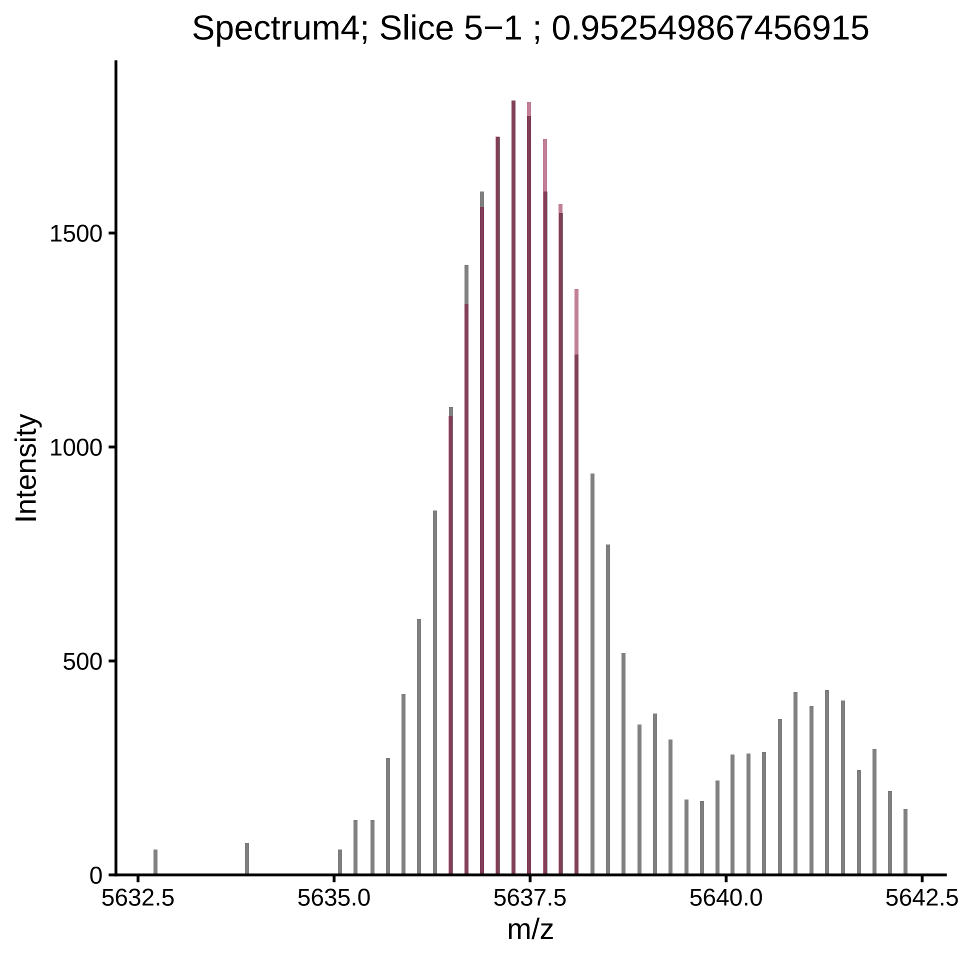
Spectrum4; Slice 5−1 ; 0.952549867456915
Intensity
0
500
1000
1500
5632.5
5635.0
5637.5
5640.0
5642.5
m/z

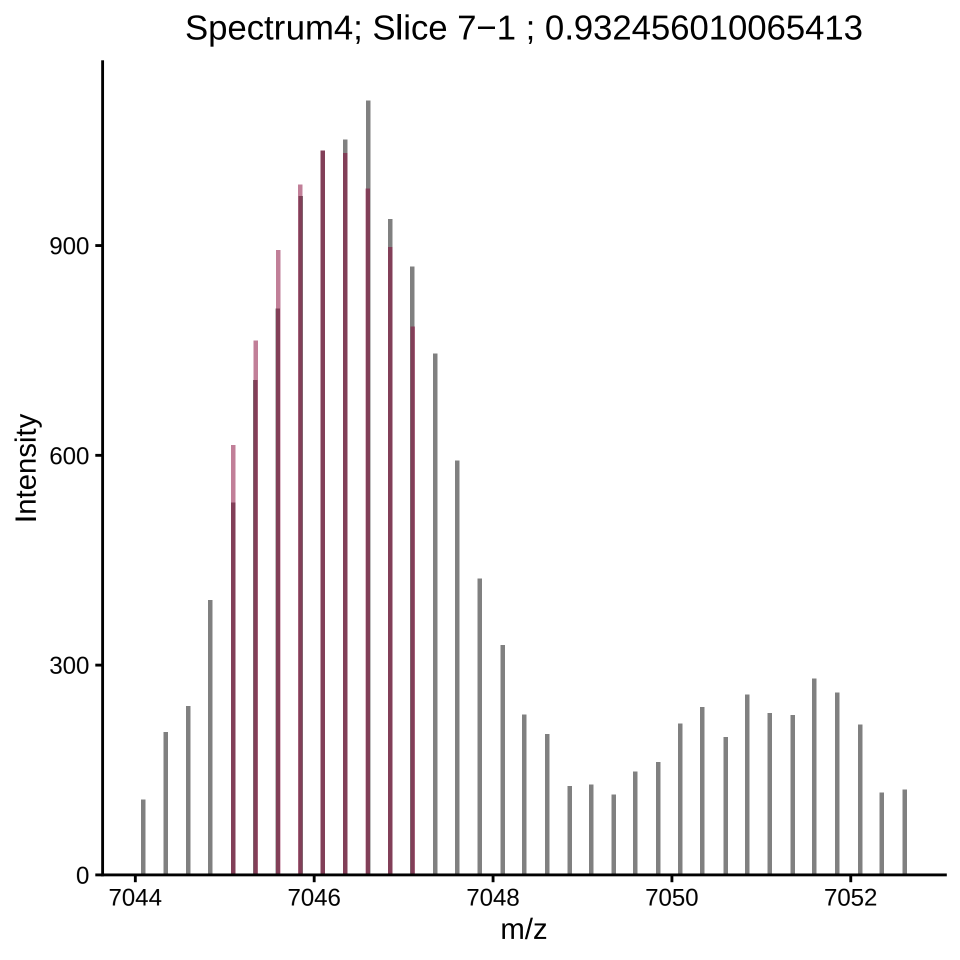

Spectrum4; Slice 7−1 ; 0.932456010065413
Intensity
900
600
300
0
7044
7046
7048
7050
7052
m/z

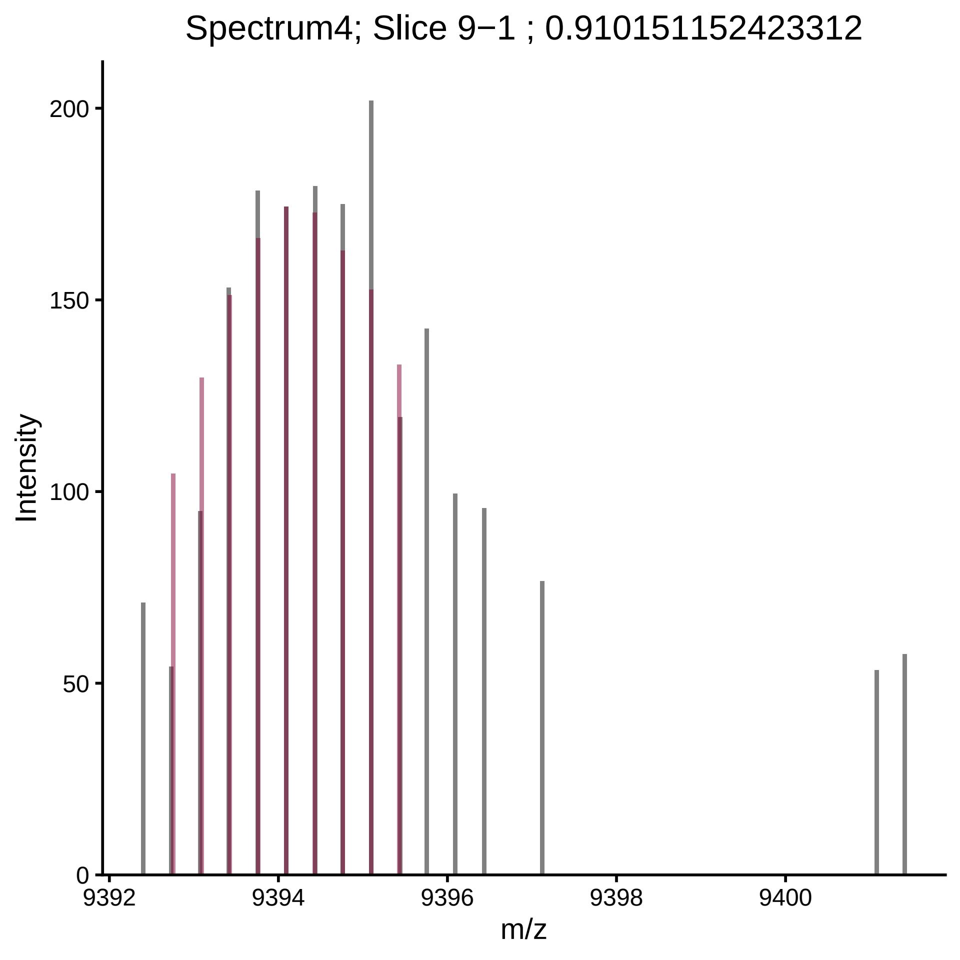

Spectrum4; Slice 9−1 ; 0.910151152423312
Intensity
200
150
100
50
0
9392
9394
9396
9398
9400
m/z

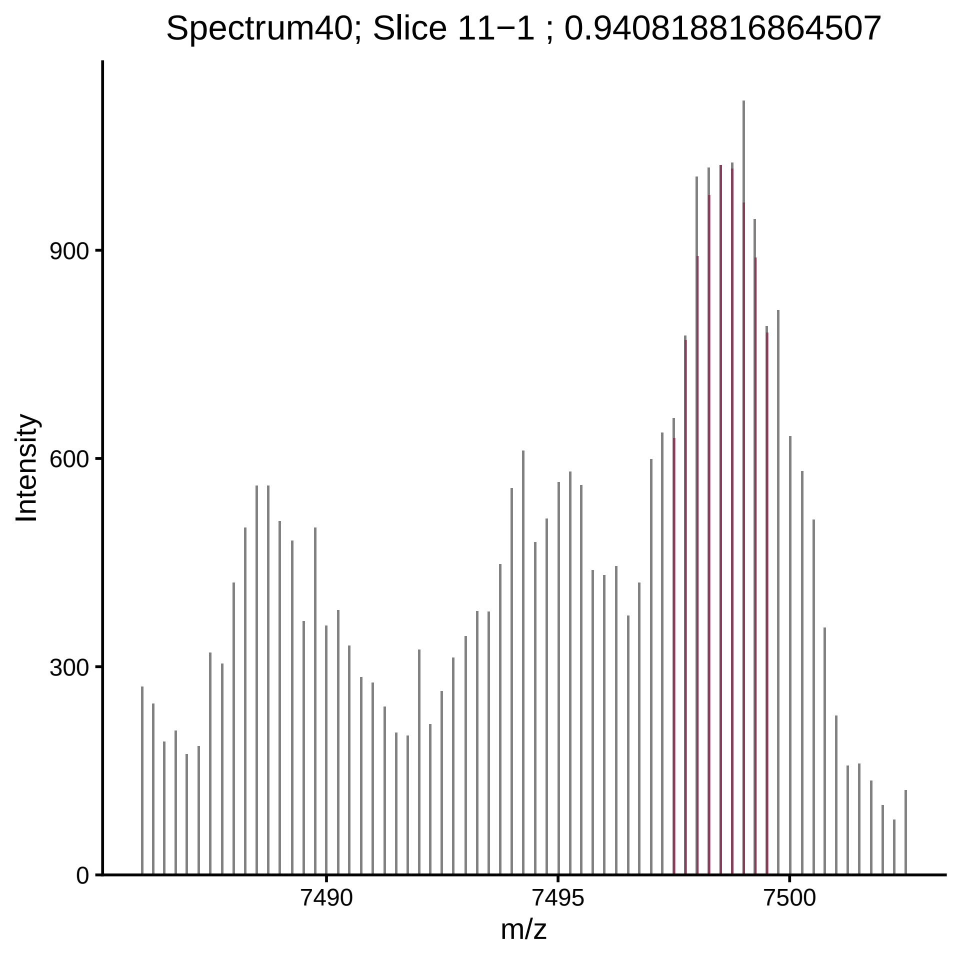

Spectrum40; Slice 11−1 ; 0.940818816864507
Intensity
900
600
300
0
7490
7495
7500
m/z

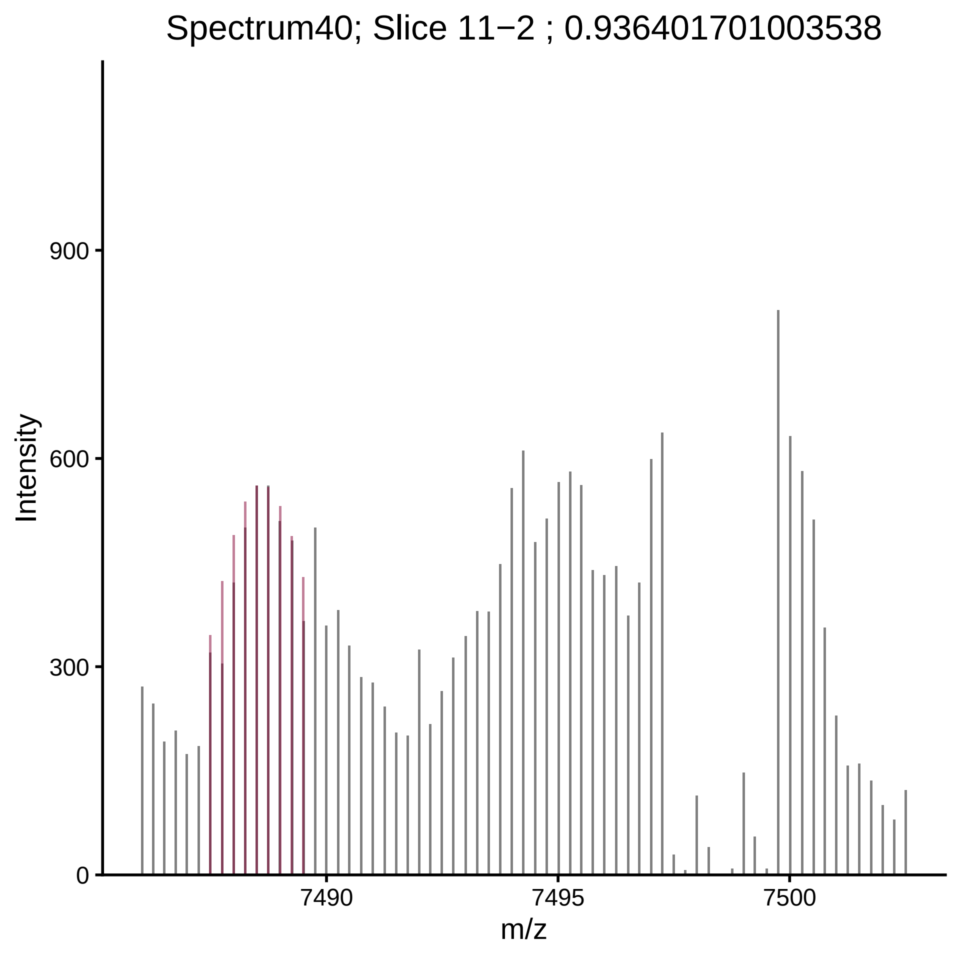

Spectrum40; Slice 11−2 ; 0.936401701003538
Intensity
900
600
300
0
7490
7495
7500
m/z

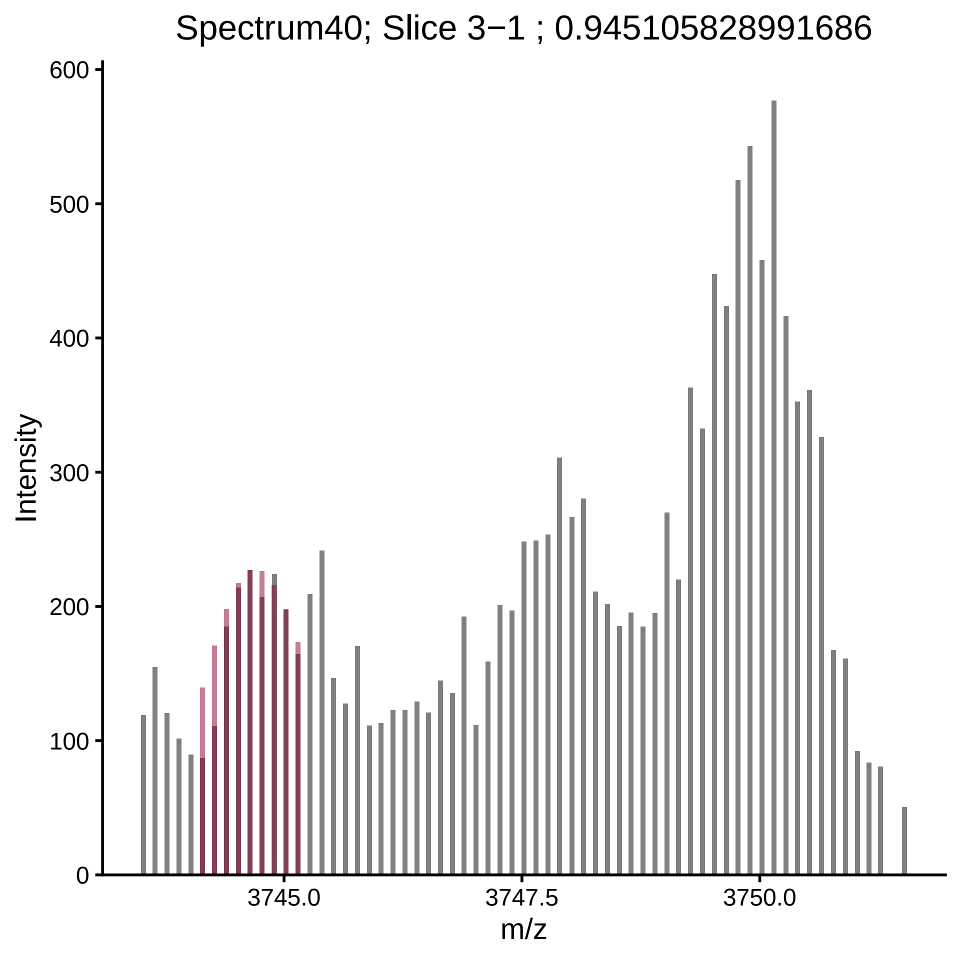

Spectrum40; Slice 3−1 ; 0.945105828991686
Intensity
600
500
400
300
200
100
0
3745.0
3747.5
3750.0
m/z

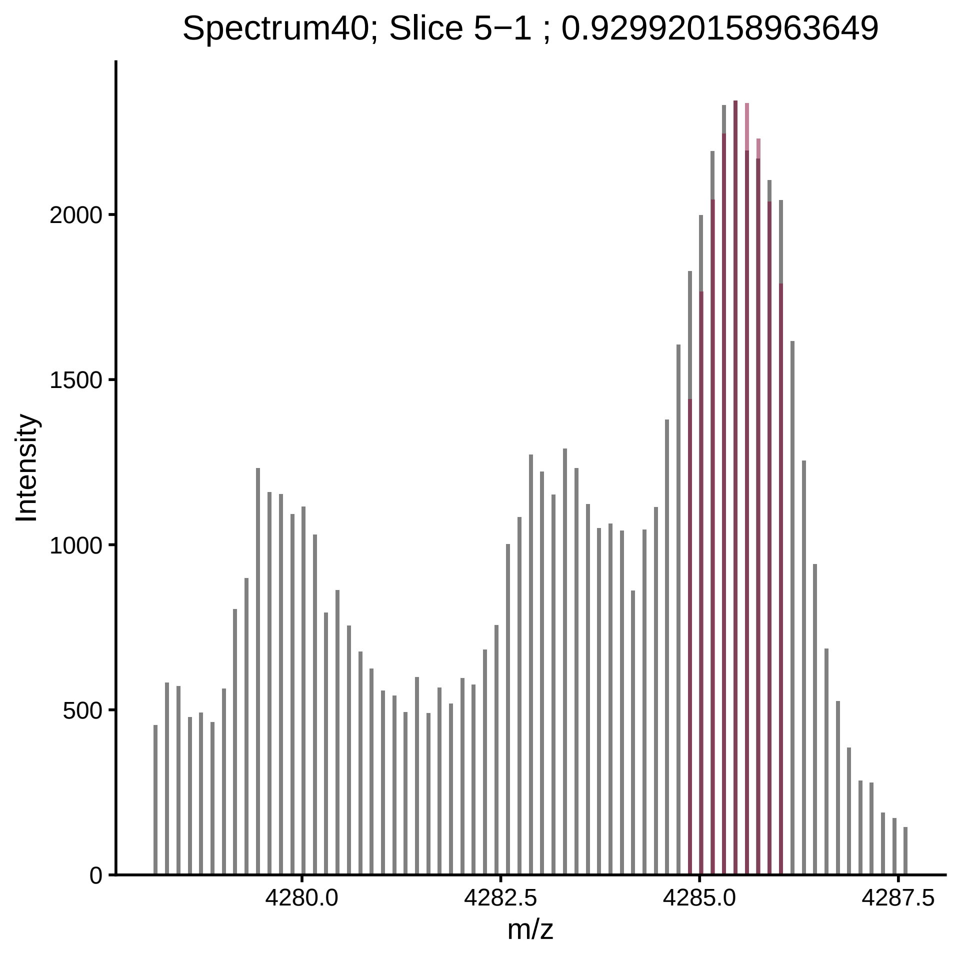

Spectrum40; Slice 5−1 ; 0.929920158963649
Intensity
2000
1500
1000
500
0
4280.0
4282.5
4285.0
4287.5
m/z

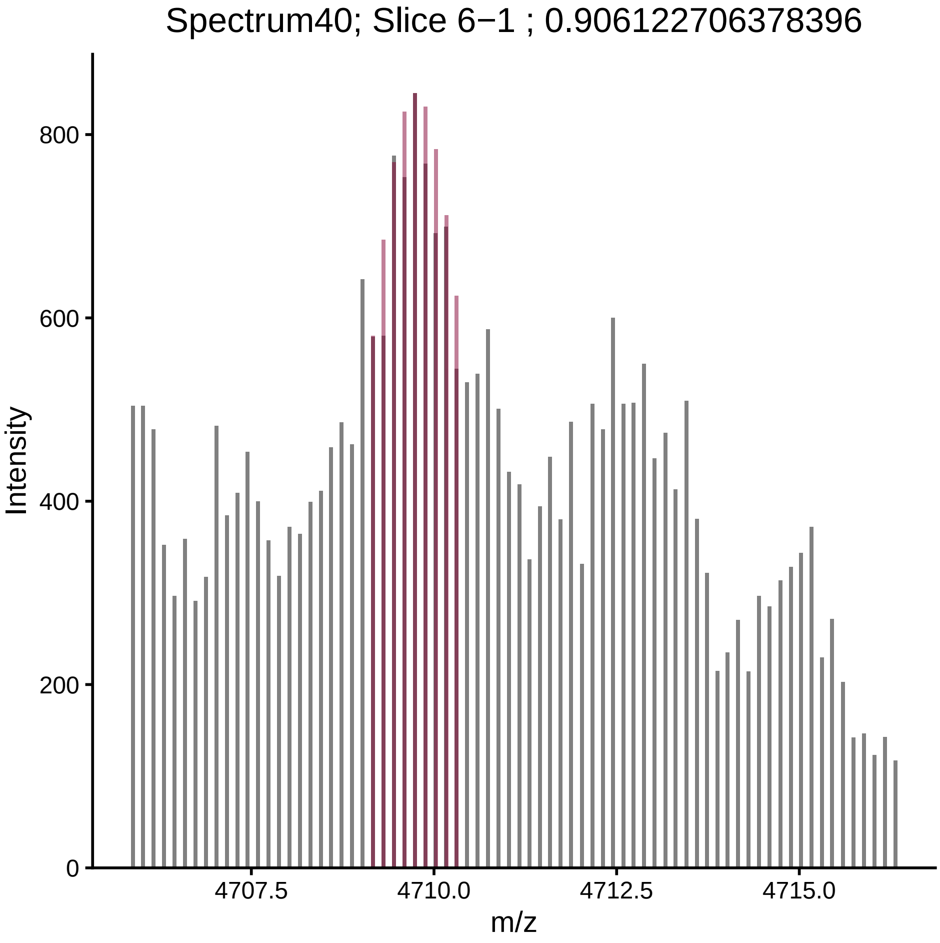

Spectrum40; Slice 6−1 ; 0.906122706378396
Intensity
800
600
400
200
0
4707.5
4710.0
4712.5
4715.0
m/z

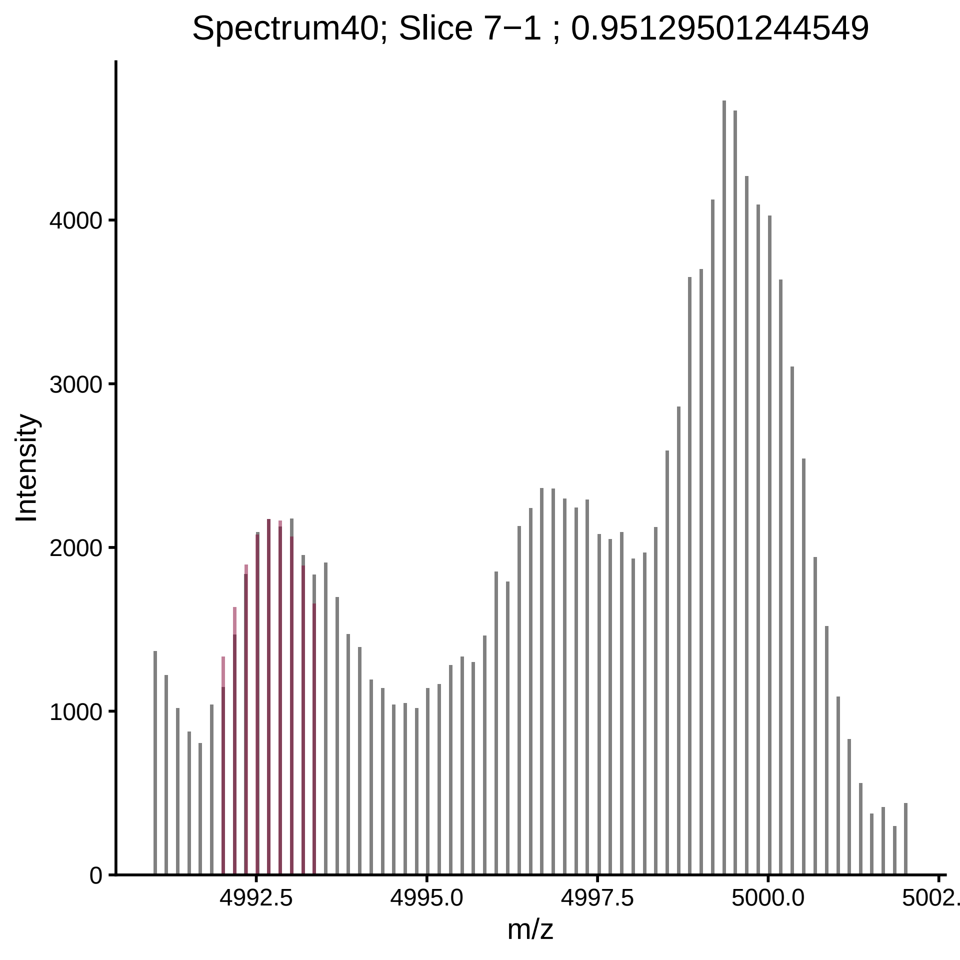
Spectrum40; Slice 7−1 ; 0.95129501244549
Intensity
0
1000
2000
3000
4000
4992.5
4995.0
4997.5
5000.0
5002.
m/z

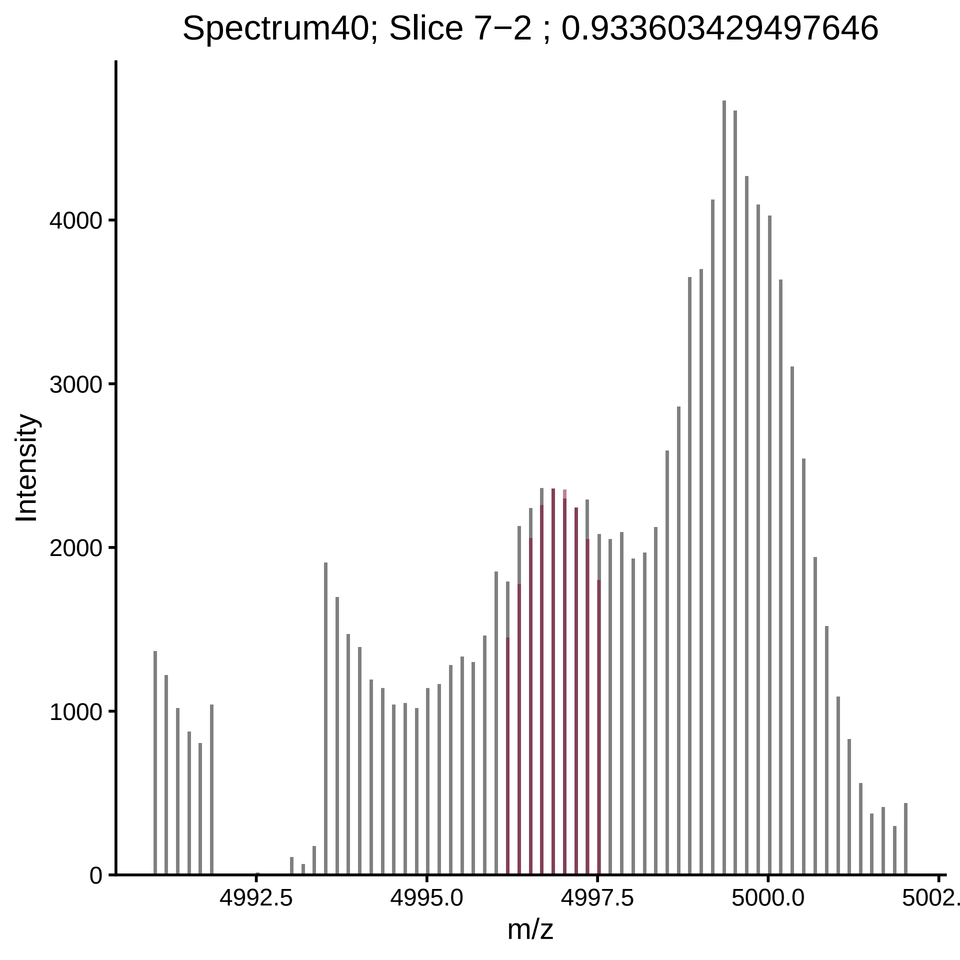

Spectrum40; Slice 7−2 ; 0.933603429497646
Intensity
0
1000
2000
3000
4000
4992.5
4995.0
4997.5
5000.0
5002.
m/z

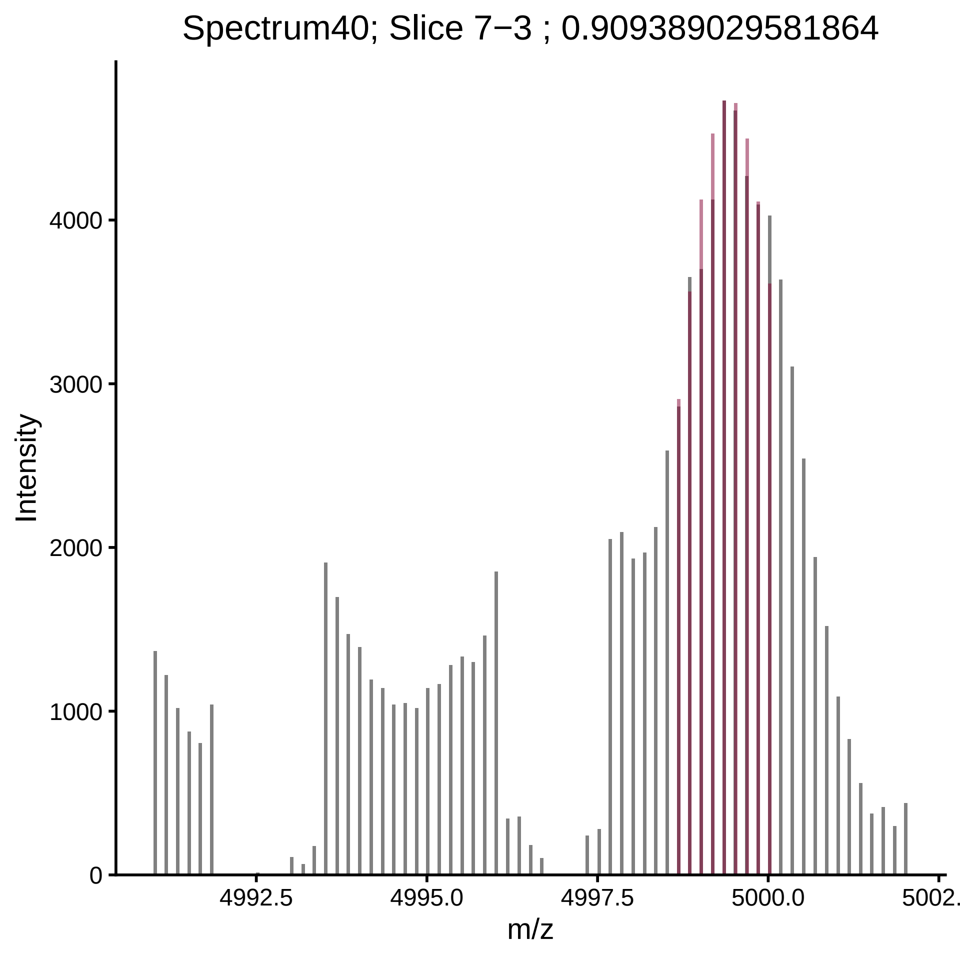

Spectrum40; Slice 7−3 ; 0.909389029581864
Intensity
m/z
0
1000
2000
3000
4000
4992.5
4995.0
4997.5
5000.0
5002.

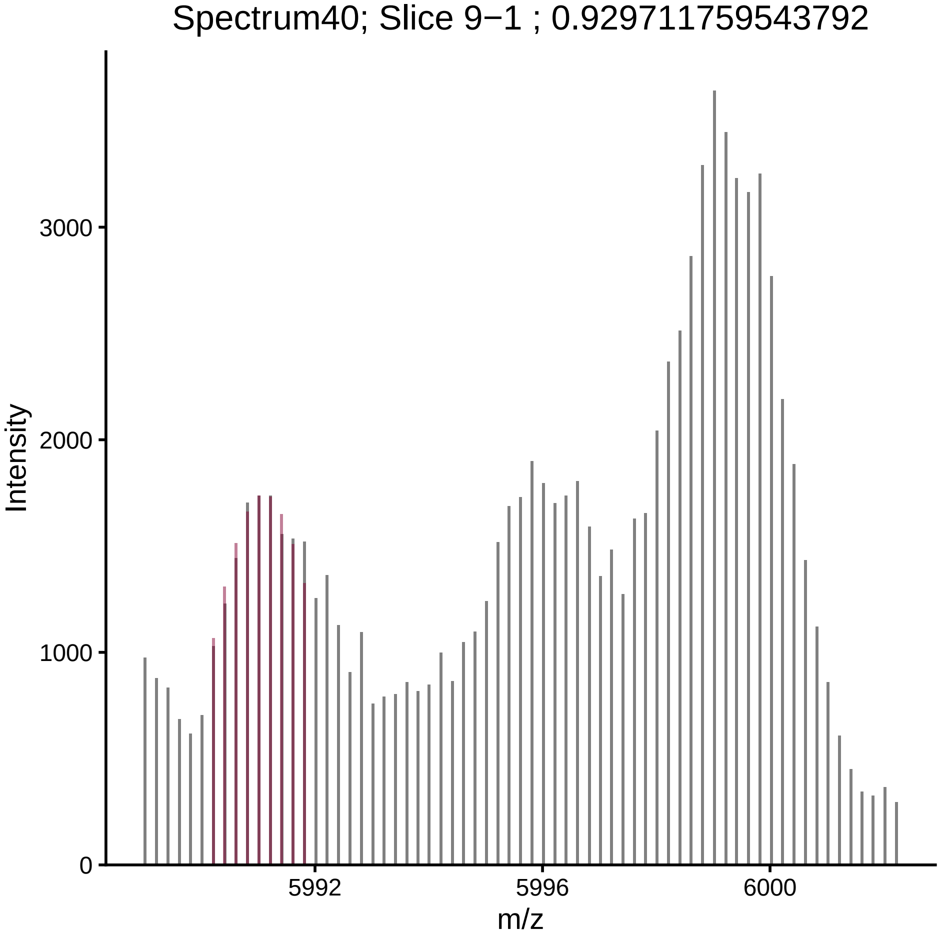

Spectrum40; Slice 9−1 ; 0.929711759543792
Intensity
3000
2000
1000
0
5992
5996
6000
m/z

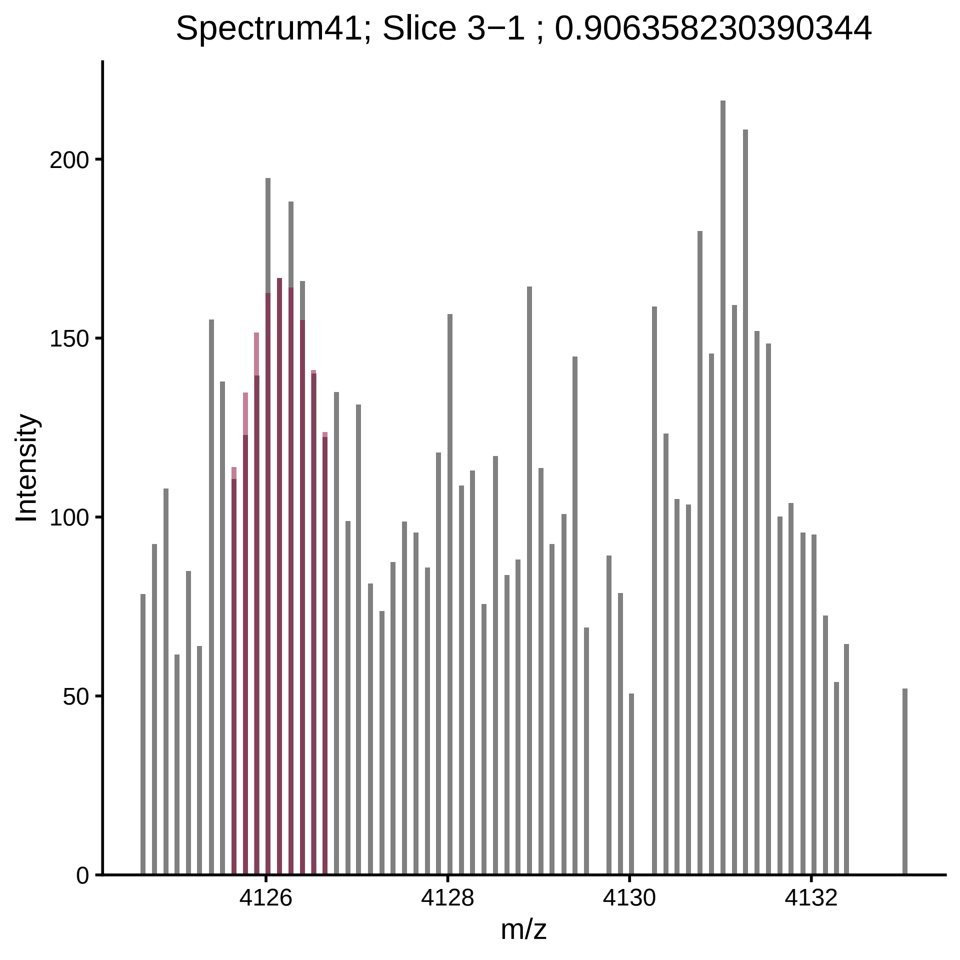

Spectrum41; Slice 3−1 ; 0.906358230390344
Intensity
0
50
100
150
200
4126
4128
4130
4132
m/z

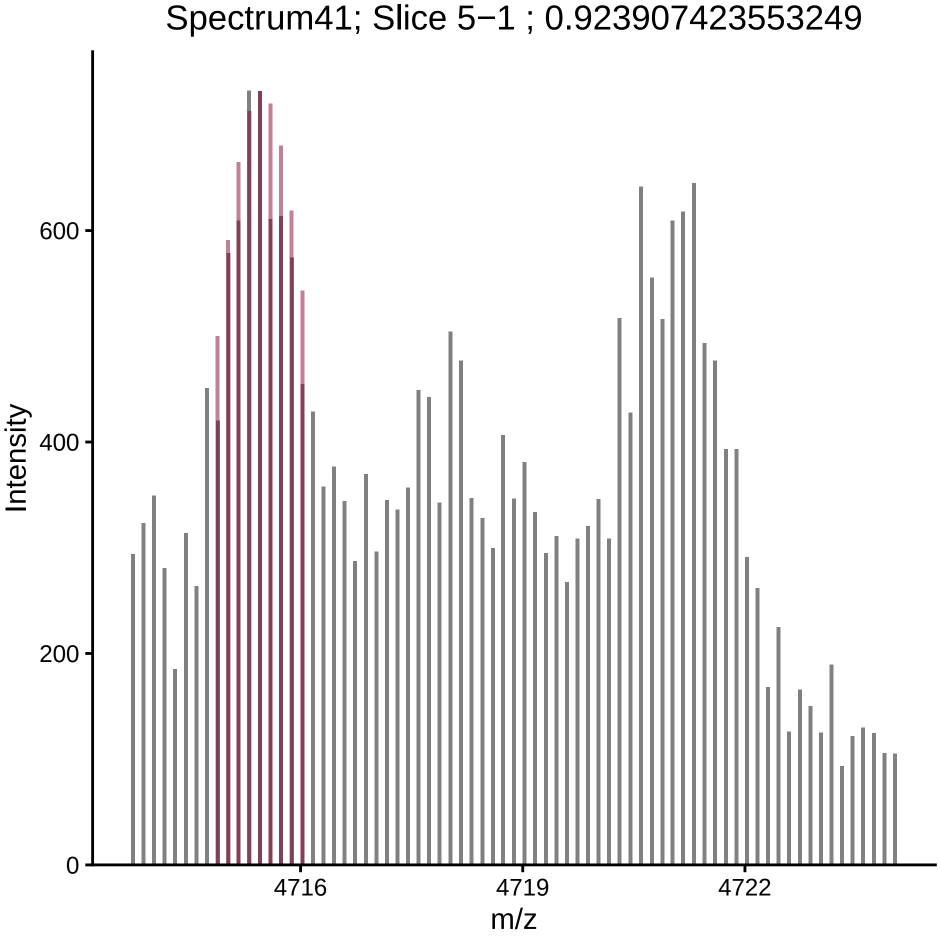

Spectrum41; Slice 5−1 ; 0.923907423553249
Intensity
600
400
200
0
4716
4719
4722
m/z

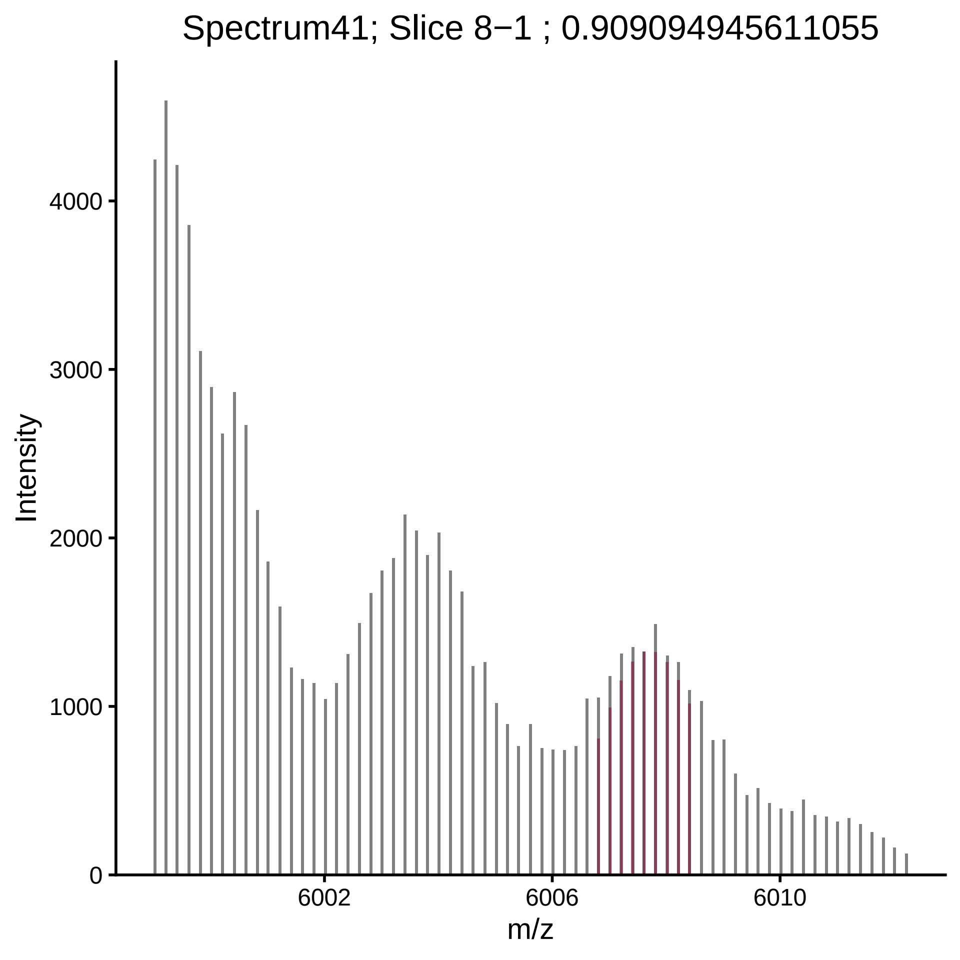

Spectrum41; Slice 8−1 ; 0.909094945611055
Intensity
4000
3000
2000
1000
0
6002
6006
6010
m/z

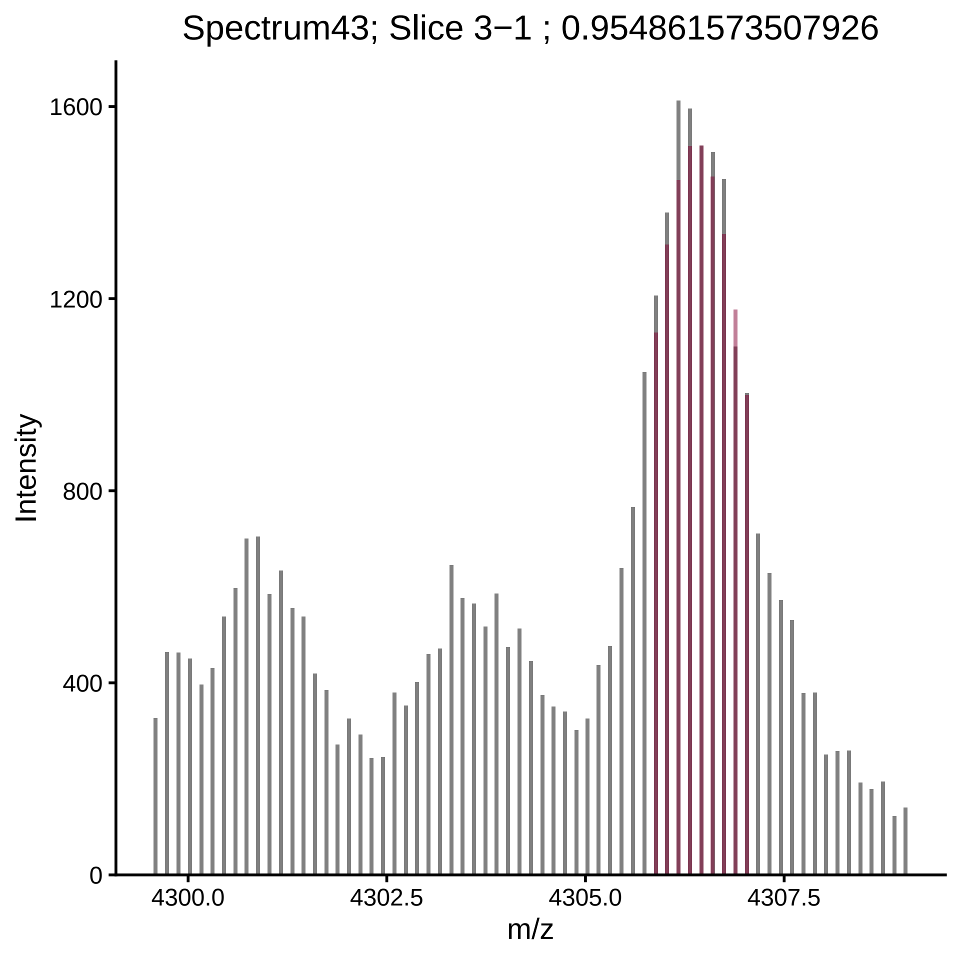

Spectrum43; Slice 3−1 ; 0.954861573507926
Intensity
1600
1200
800
400
0
4300.0
4302.5
4305.0
4307.5
m/z

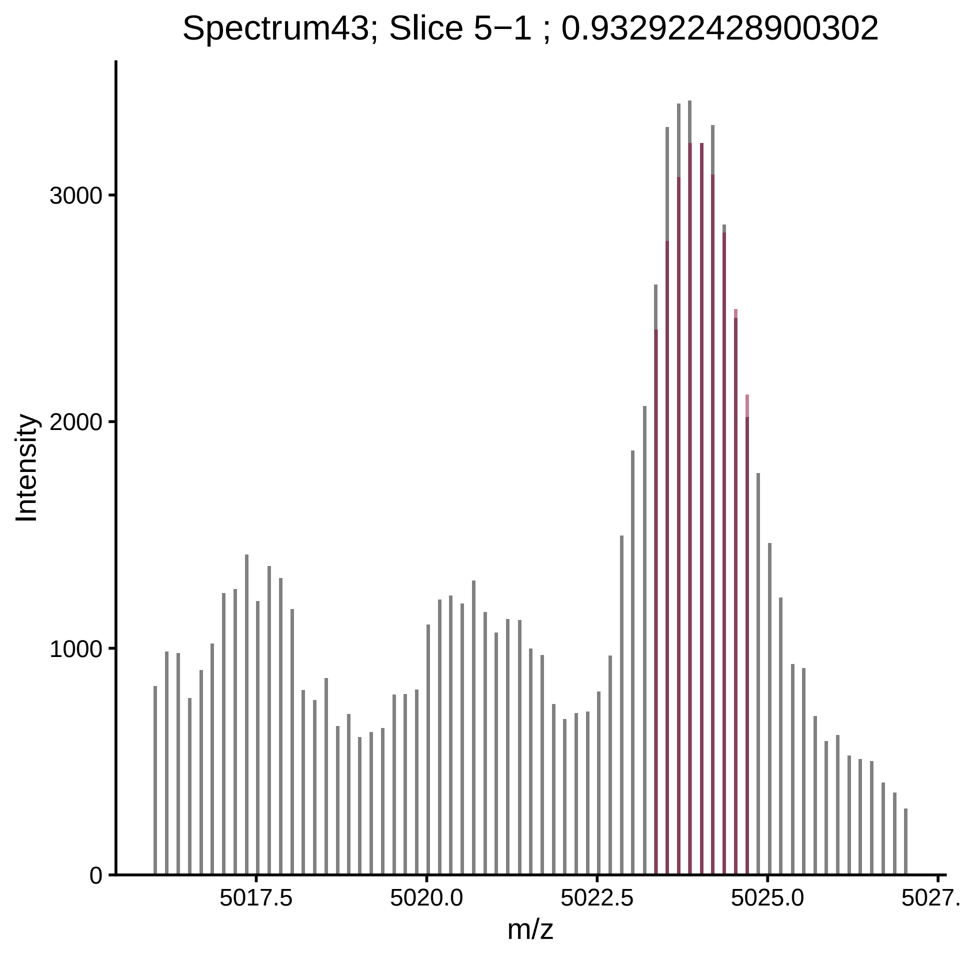
Spectrum43; Slice 5−1 ; 0.932922428900302
Intensity
3000
2000
1000
0
5017.5
5020.0
5022.5
5025.0
5027.
m/z

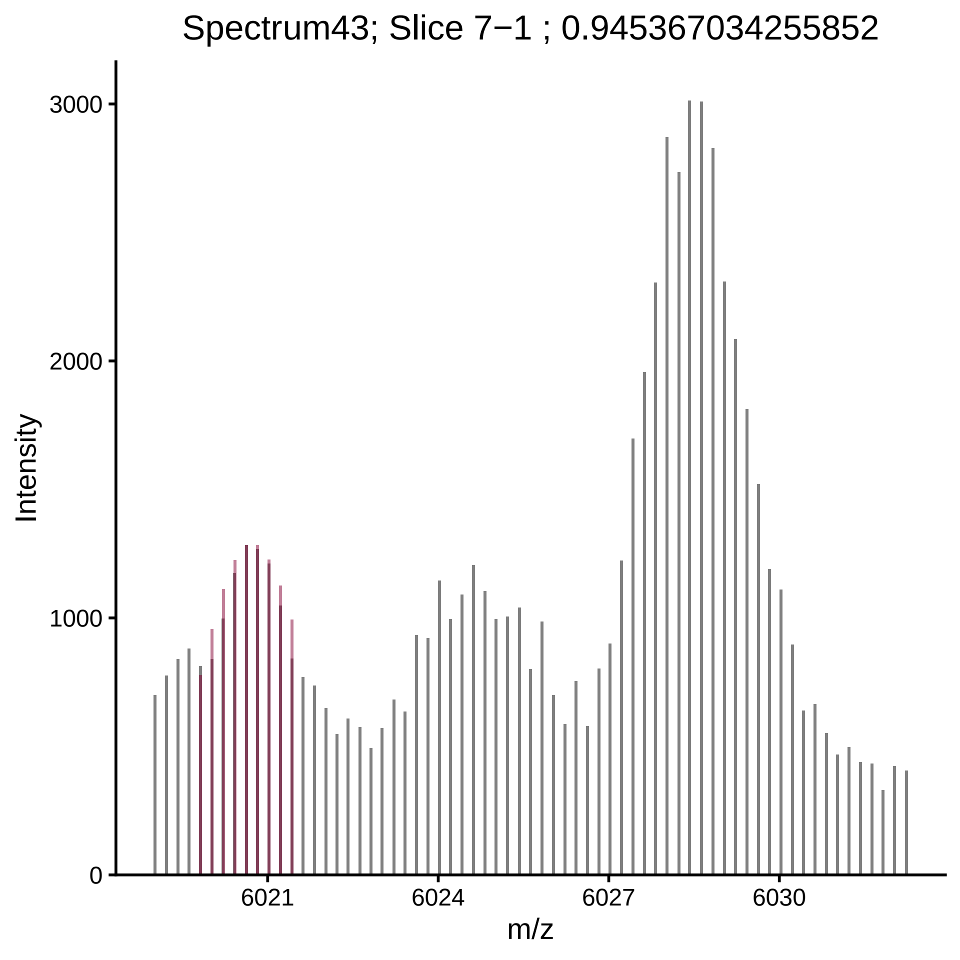

Spectrum43; Slice 7−1 ; 0.945367034255852
Intensity
3000
2000
1000
0
6021
6024
6027
6030
m/z

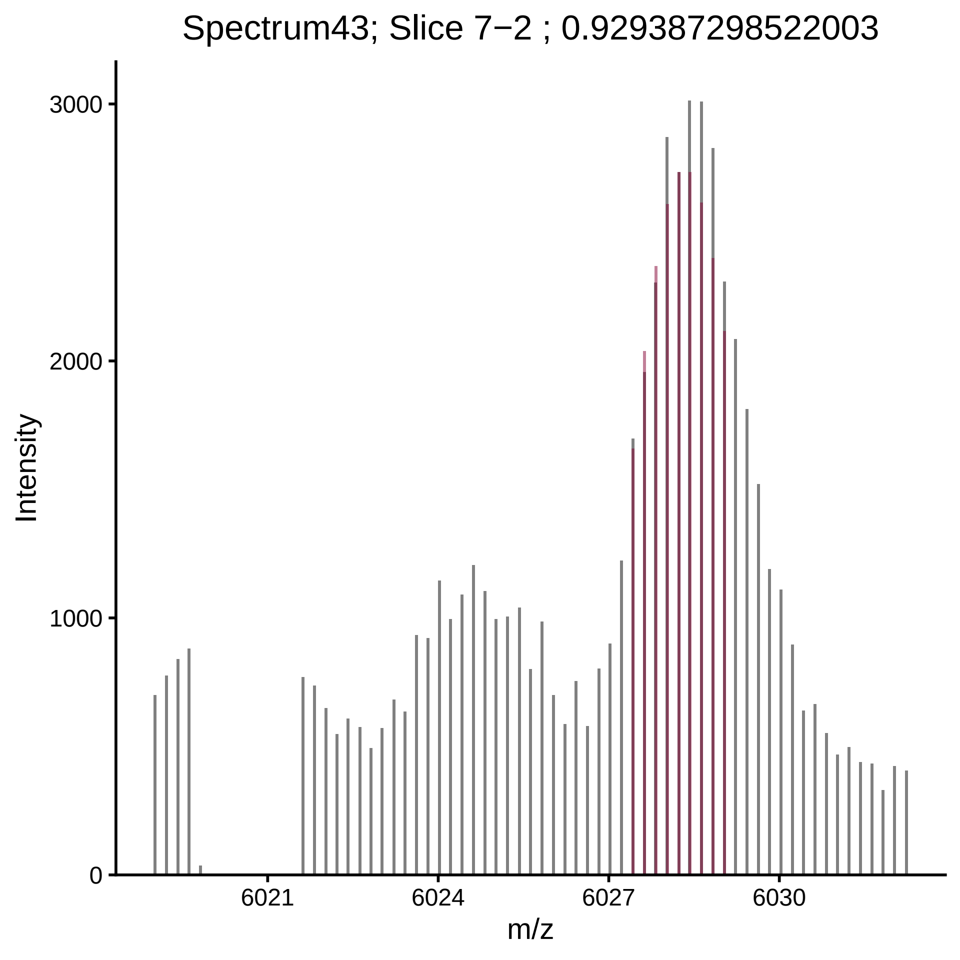

Spectrum43; Slice 7−2 ; 0.929387298522003
Intensity
3000
2000
1000
0
6021
6024
6027
6030
m/z

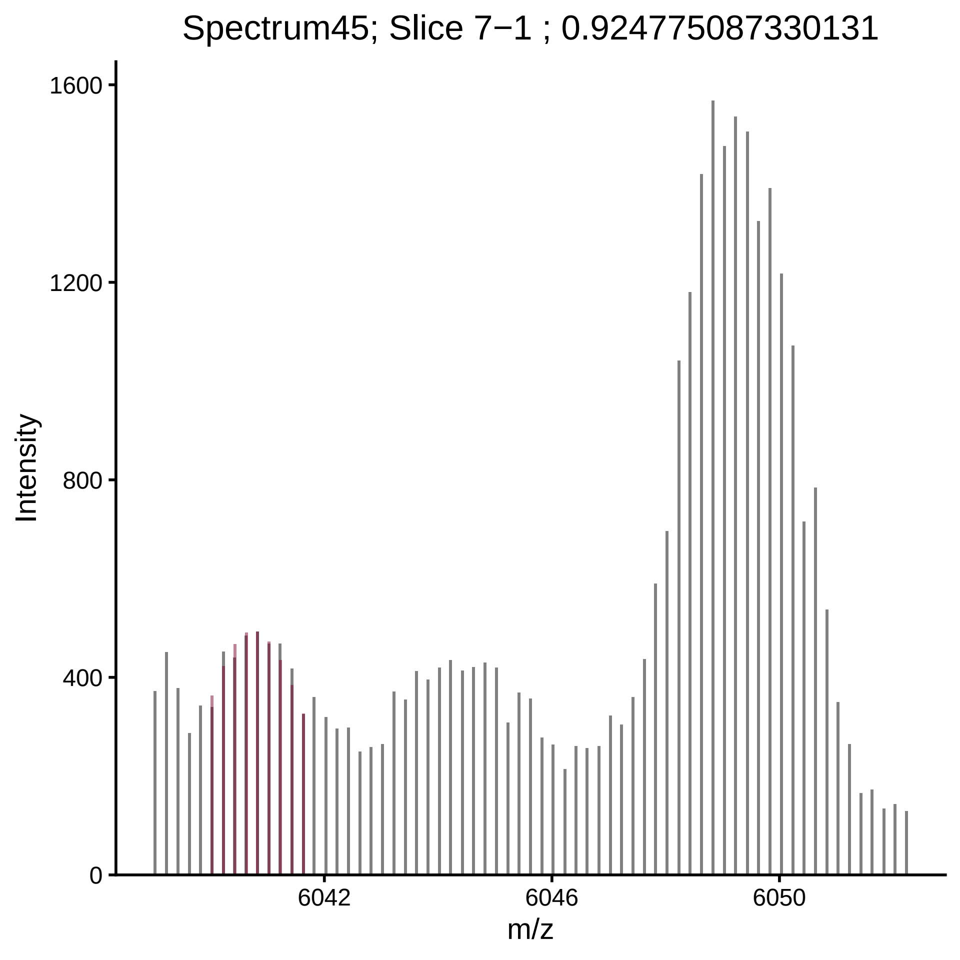
Spectrum45; Slice 7−1 ; 0.924775087330131
Intensity
0
400
800
1200
1600
6042
6046
6050
m/z

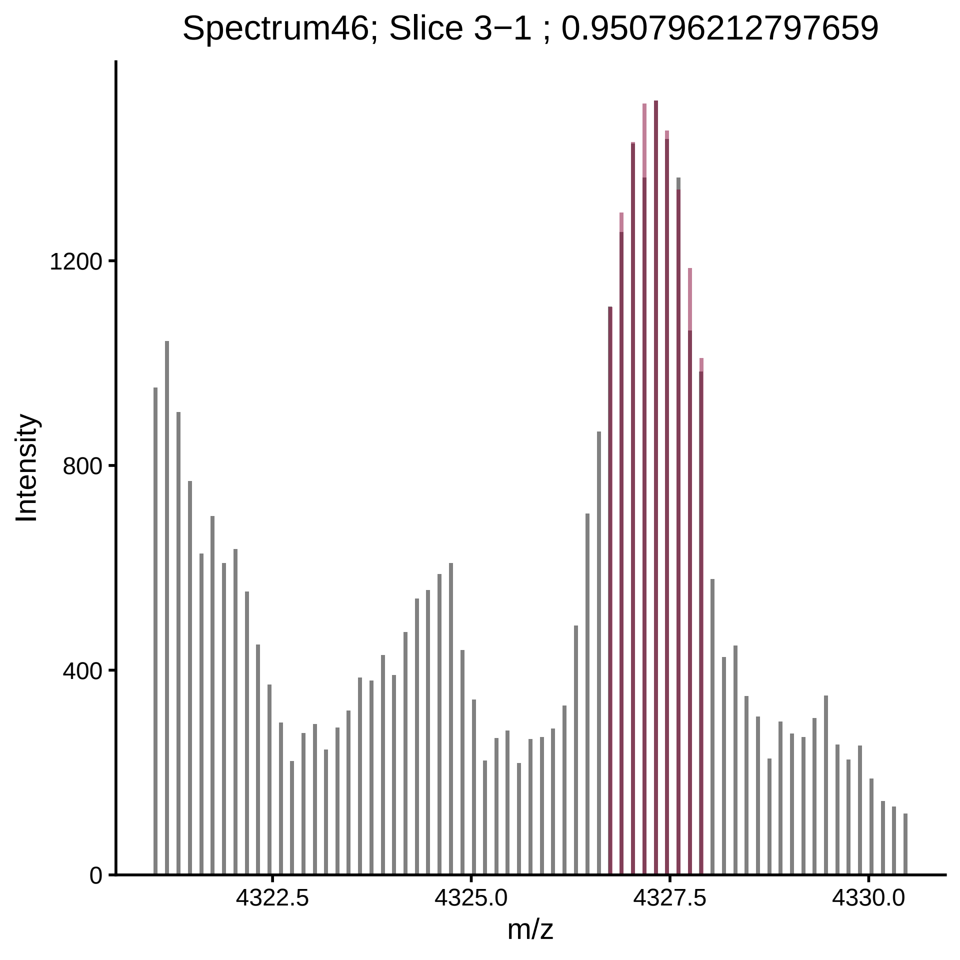
Spectrum46; Slice 3−1 ; 0.950796212797659
Intensity
1200
800
400
0
4322.5
4325.0
4327.5
4330.0
m/z

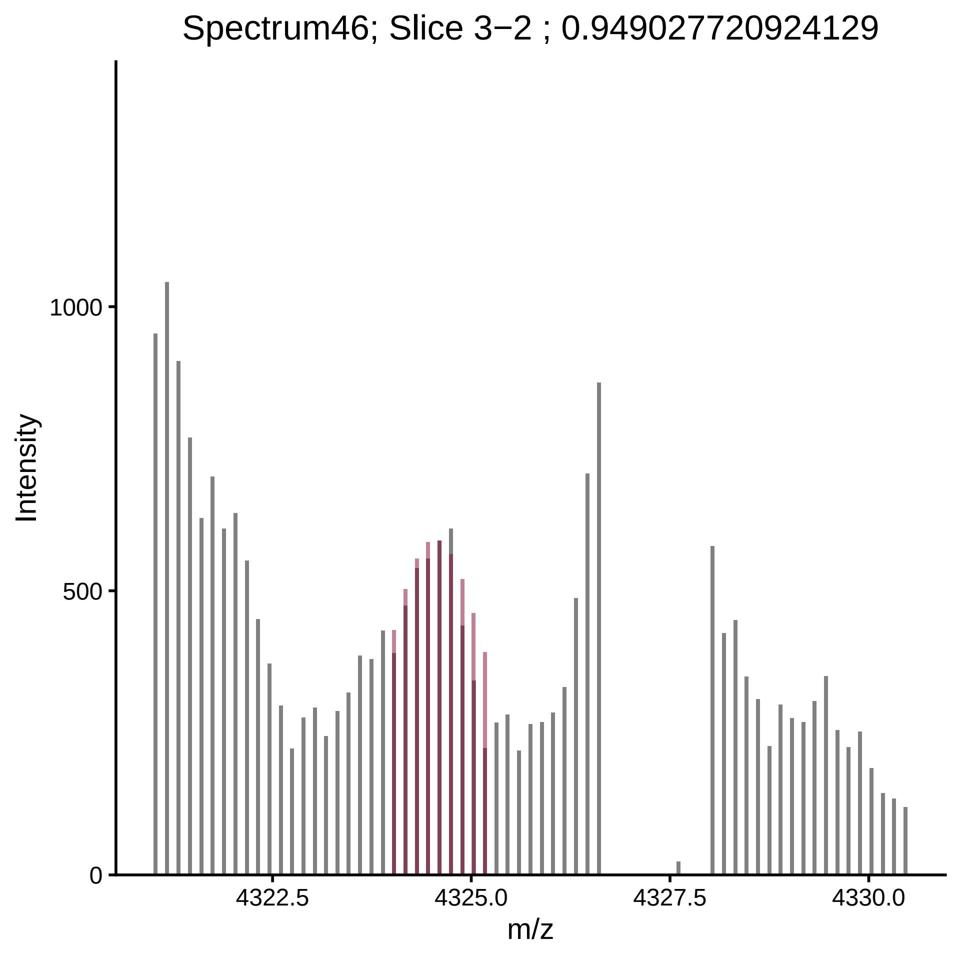

Spectrum46; Slice 3−2 ; 0.949027720924129
Intensity
1000
500
0
4322.5
4325.0
4327.5
4330.0
m/z

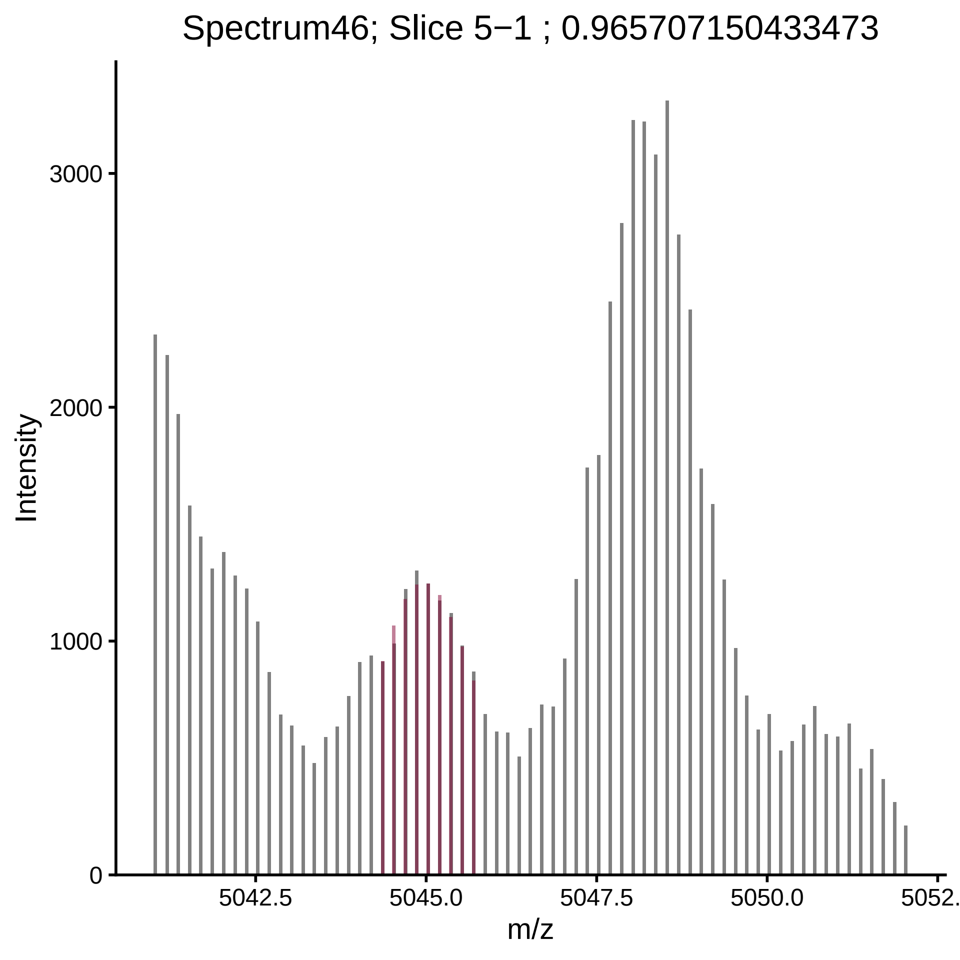

Spectrum46; Slice 5−1 ; 0.965707150433473
Intensity
3000
2000
1000
0
5042.5
5045.0
5047.5
5050.0
5052.
m/z

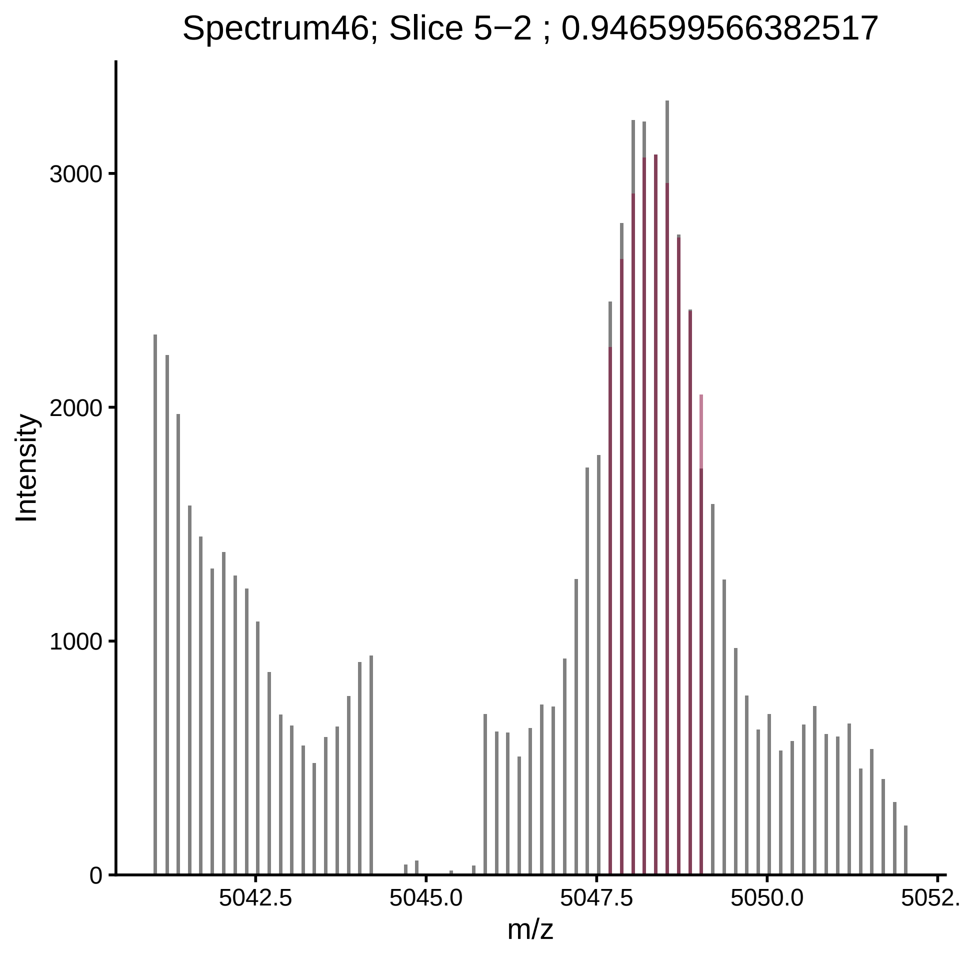

Spectrum46; Slice 5−2 ; 0.946599566382517
Intensity
3000
2000
1000
0
5042.5
5045.0
5047.5
5050.0
5052.
m/z

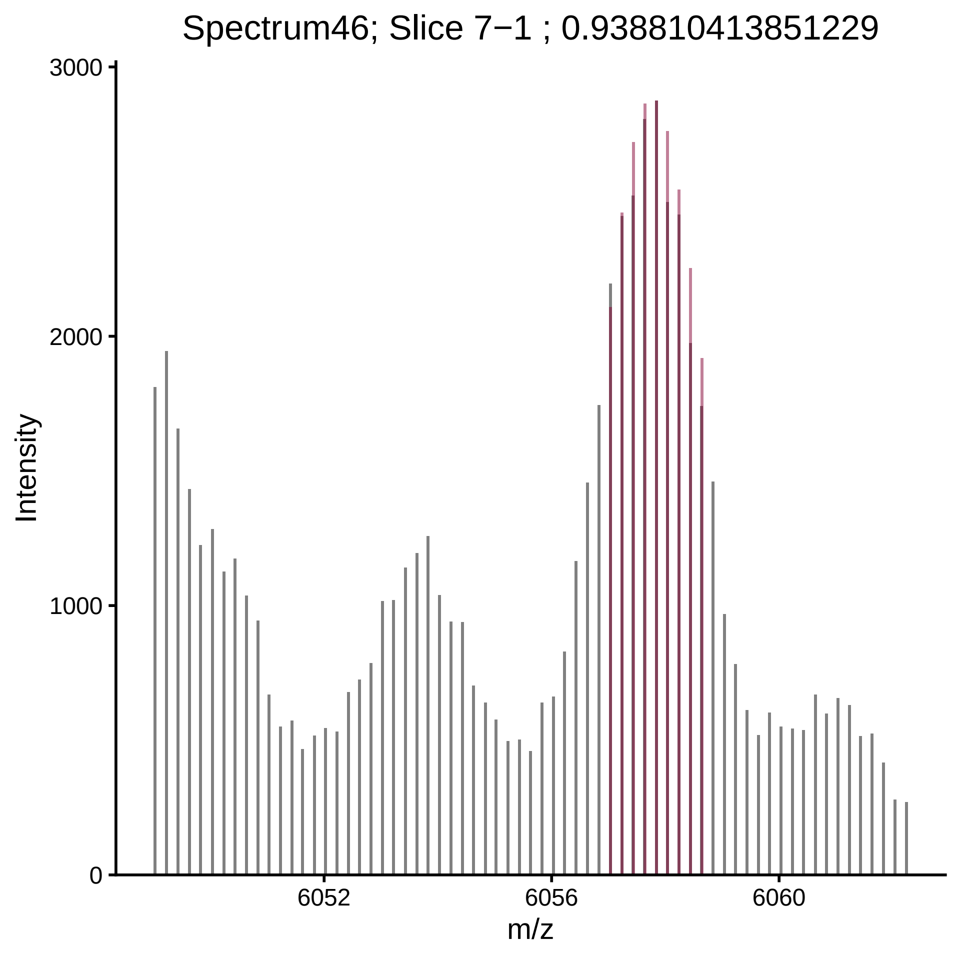

Spectrum46; Slice 7−1 ; 0.938810413851229
Intensity
3000
2000
1000
0
6052
6056
6060
m/z

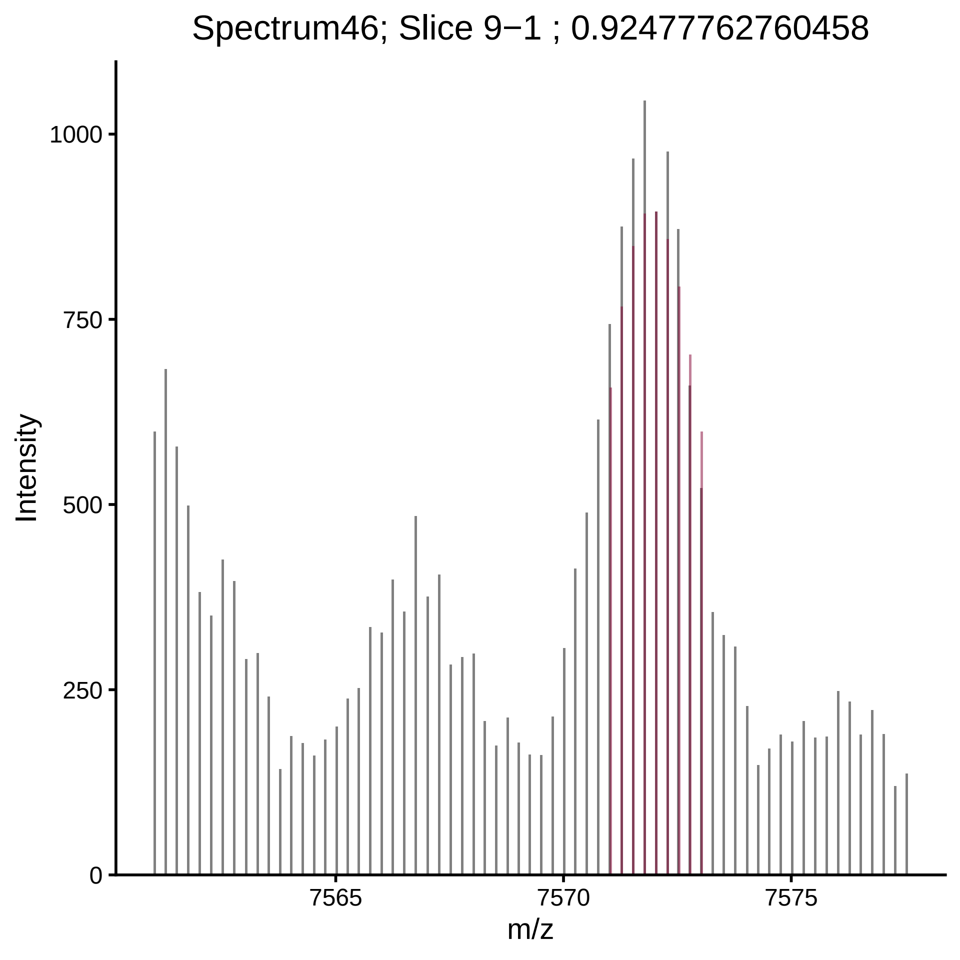

Spectrum46; Slice 9−1 ; 0.92477762760458
Intensity
1000
750
500
250
0
7565
7570
7575
m/z

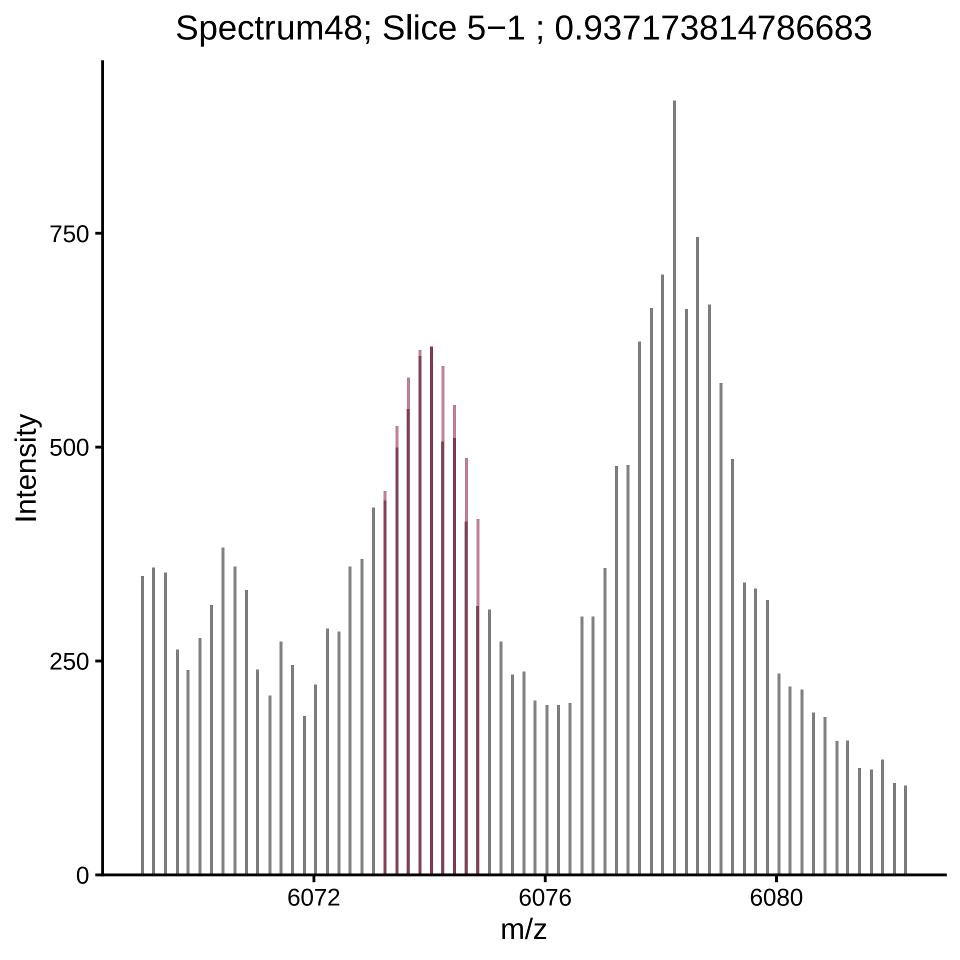
Spectrum48; Slice 5−1 ; 0.937173814786683
Intensity
750
500
250
0
6072
6076
6080
m/z

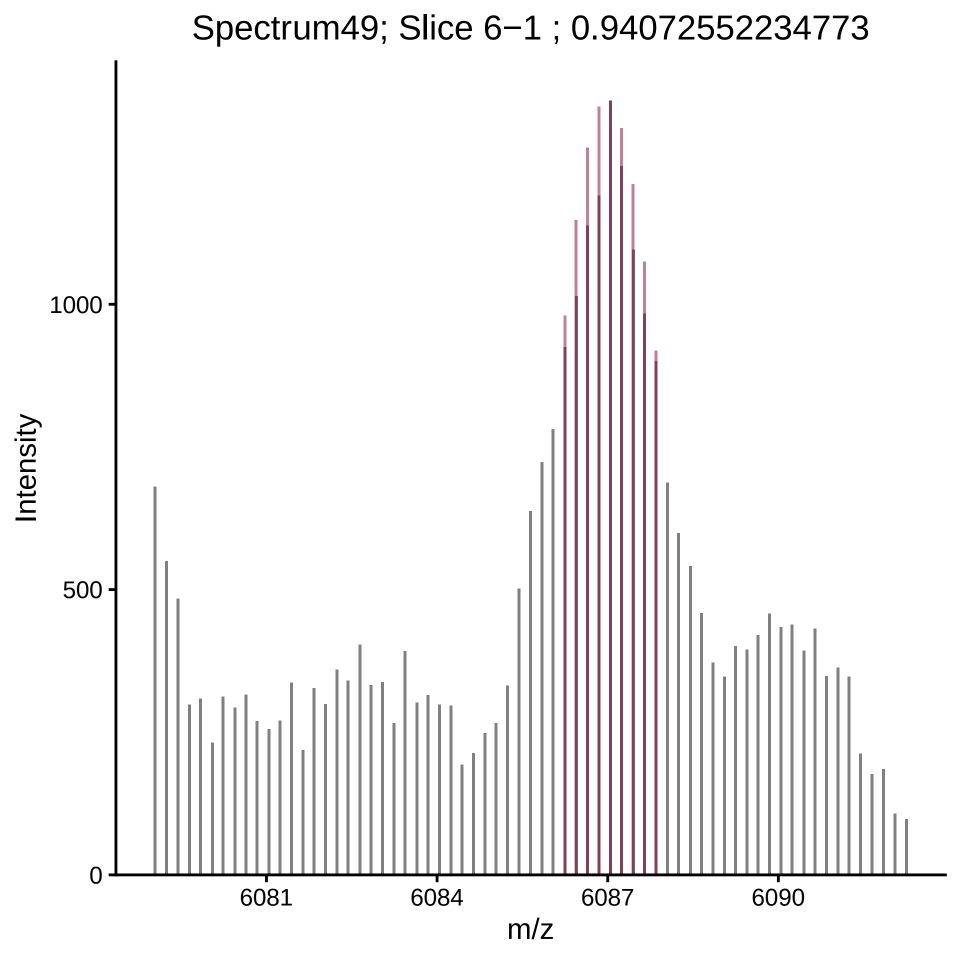

Spectrum49; Slice 6−1 ; 0.94072552234773
Intensity
1000
500
0
6081
6084
6087
6090
m/z

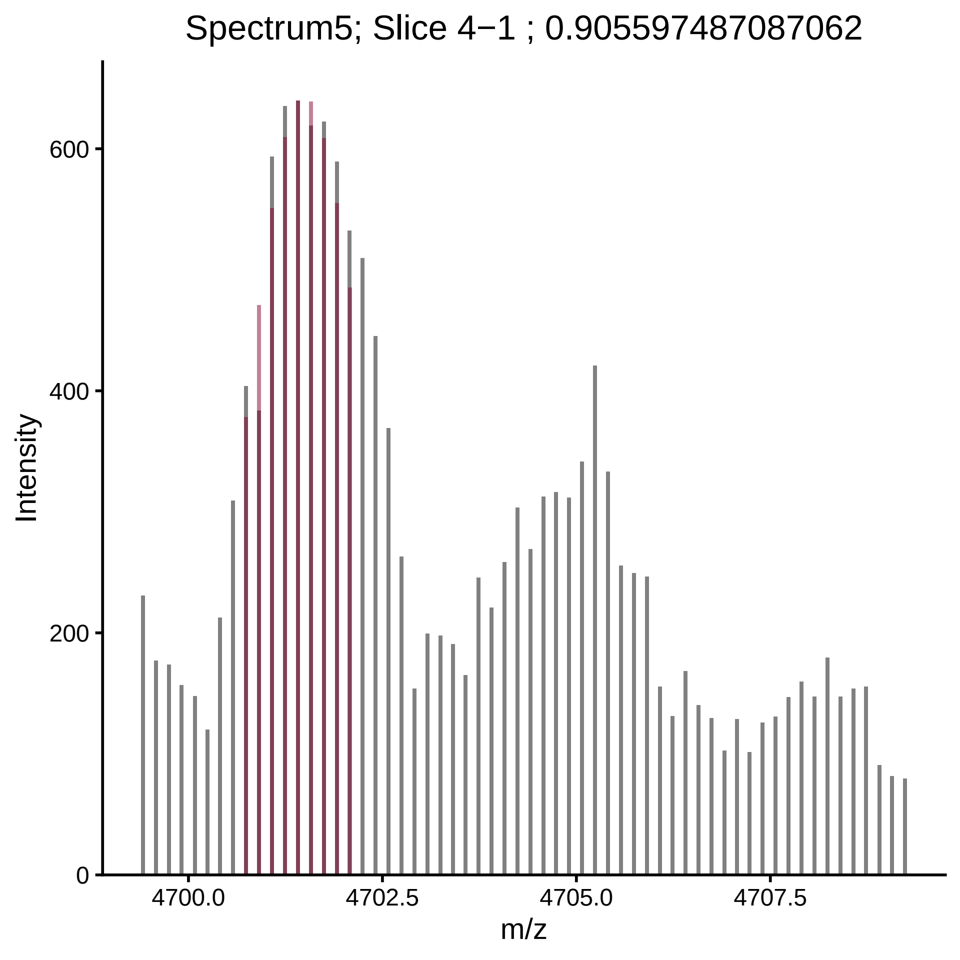

Spectrum5; Slice 4−1 ; 0.905597487087062
Intensity
0
200
400
600
4700.0
4702.5
4705.0
4707.5
m/z

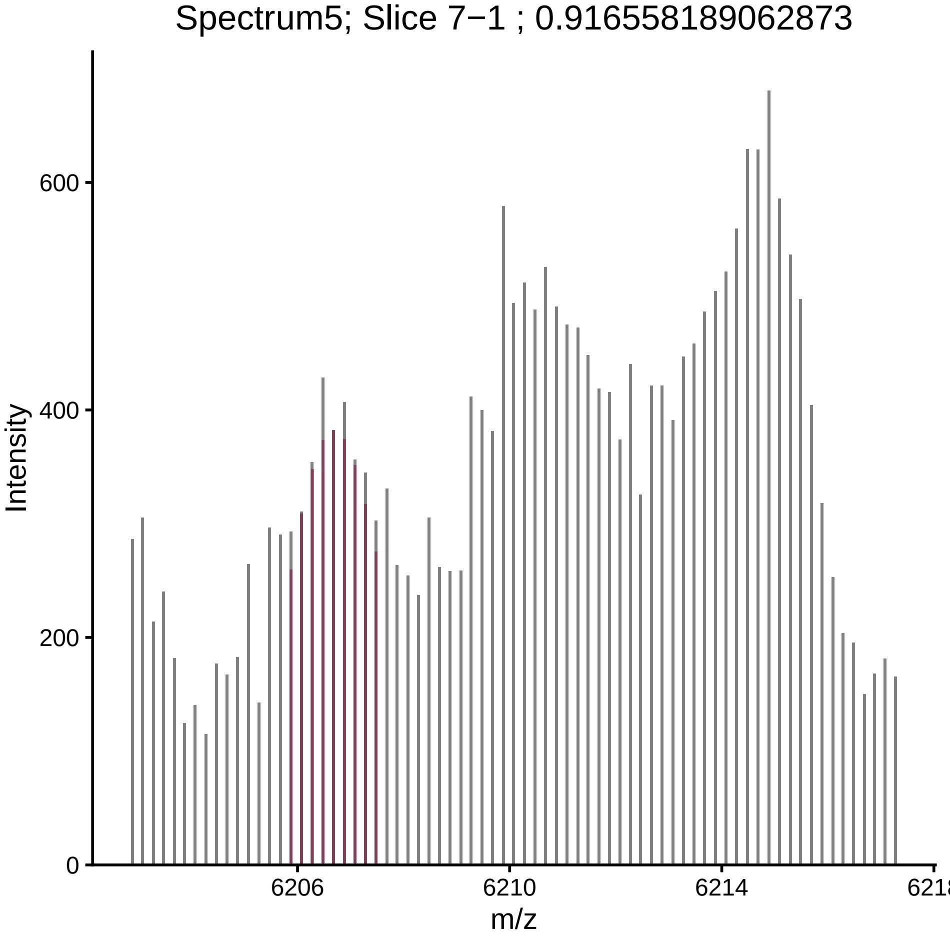

Spectrum5; Slice 7−1 ; 0.916558189062873
Intensity
0
200
400
600
6206
6210
6214
m/z

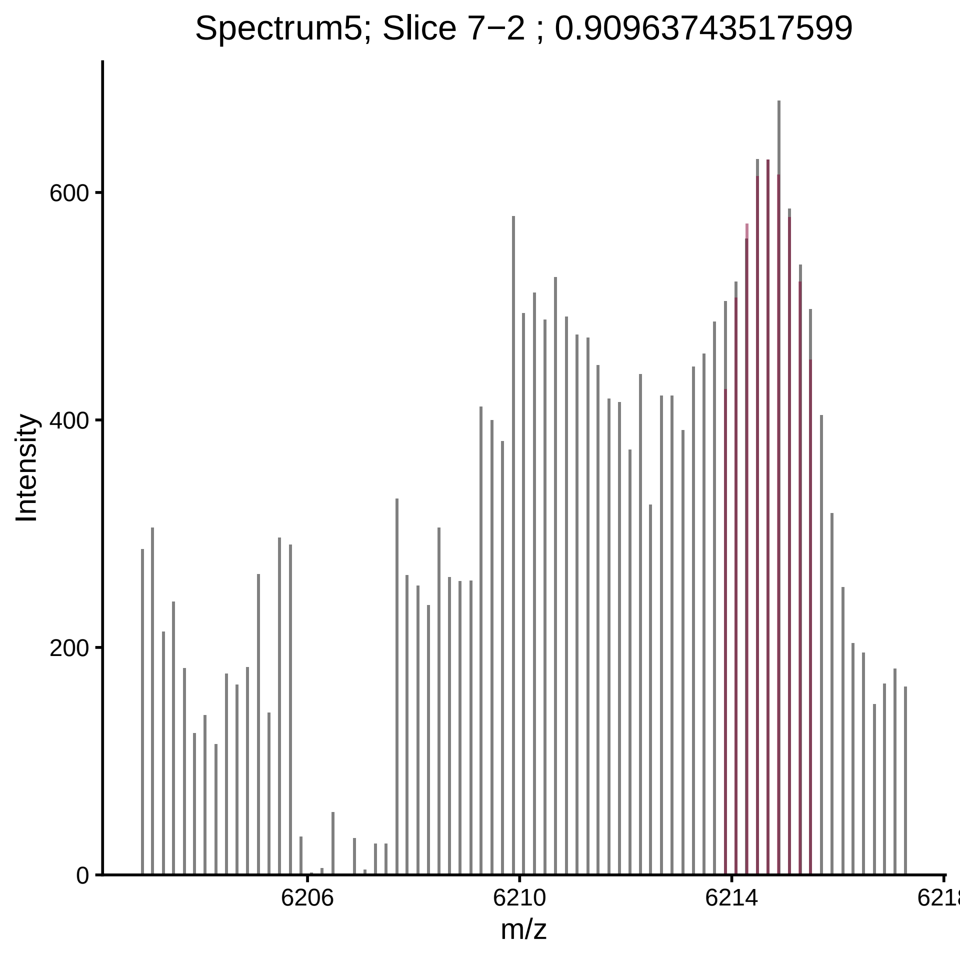
Spectrum5; Slice 7−2 ; 0.90963743517599
Intensity
m/z
0
200
400
600
6206
6210
6214
6218

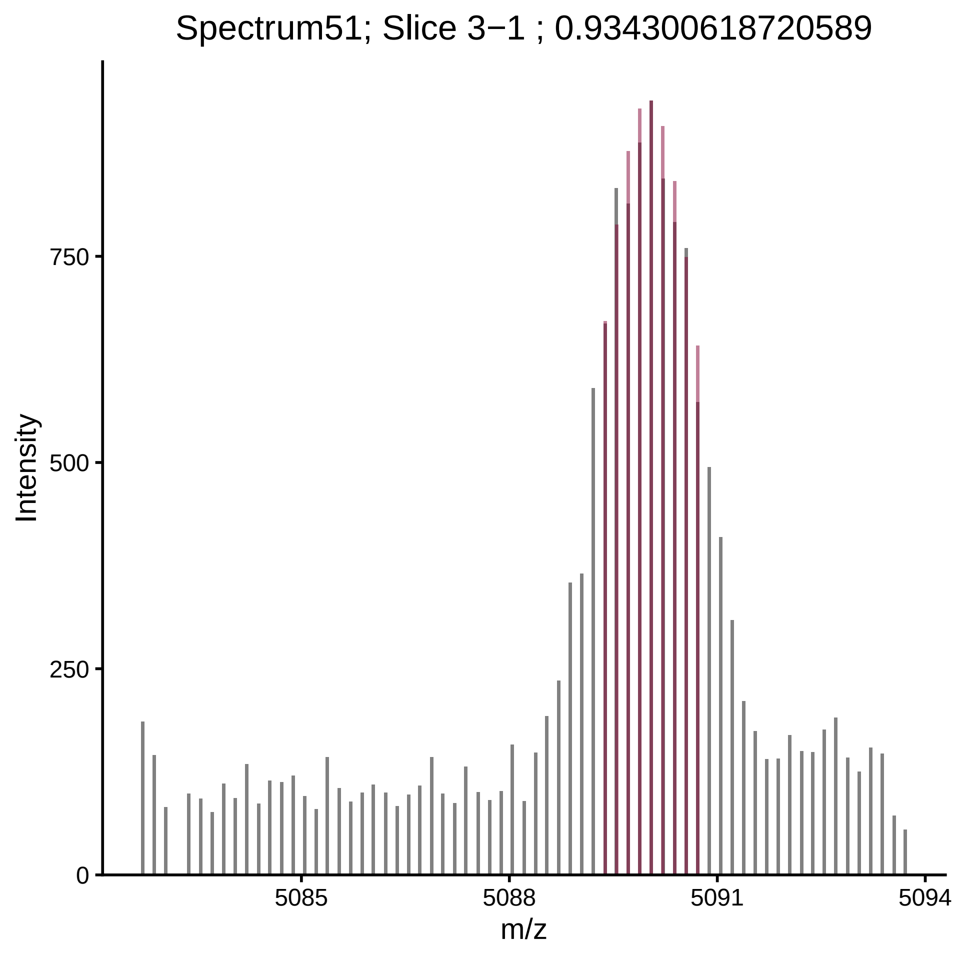

Spectrum51; Slice 3−1 ; 0.934300618720589
Intensity
750
500
250
0
5085
5088
5091
5094
m/z

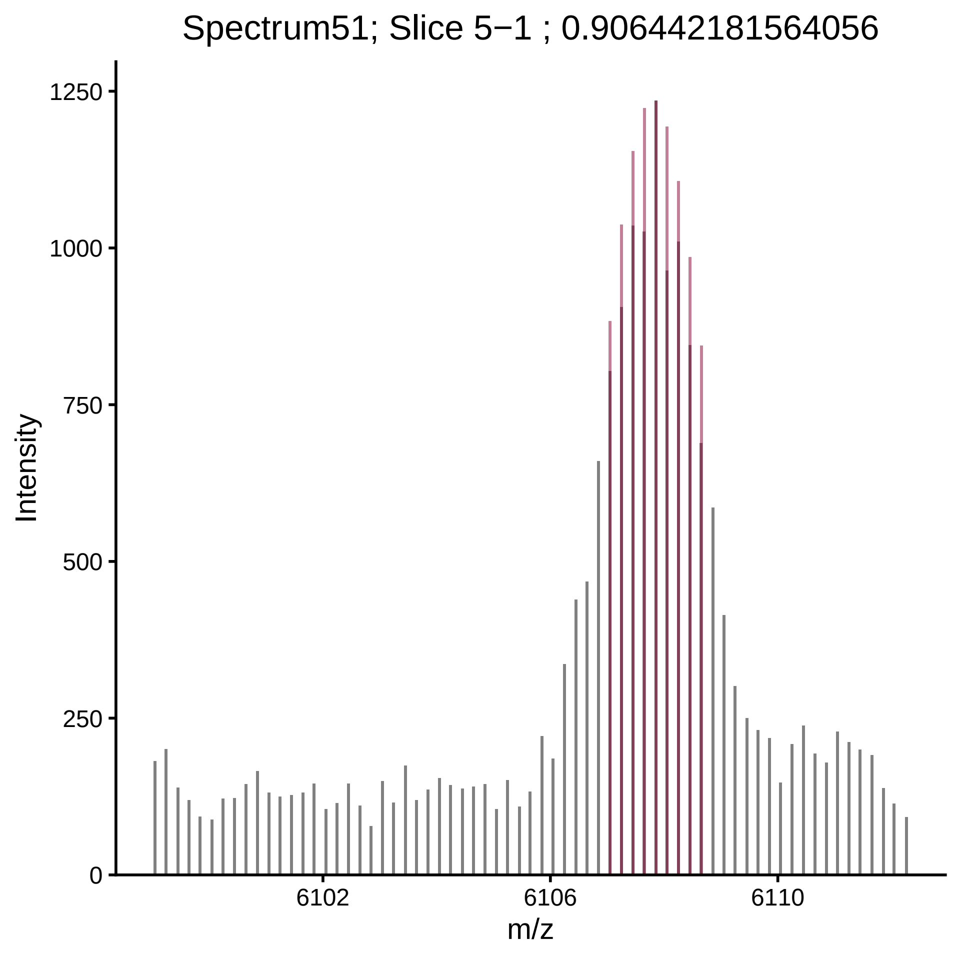
Spectrum51; Slice 5−1 ; 0.906442181564056
Intensity
1250
1000
750
500
250
0
6102
6106
6110
m/z

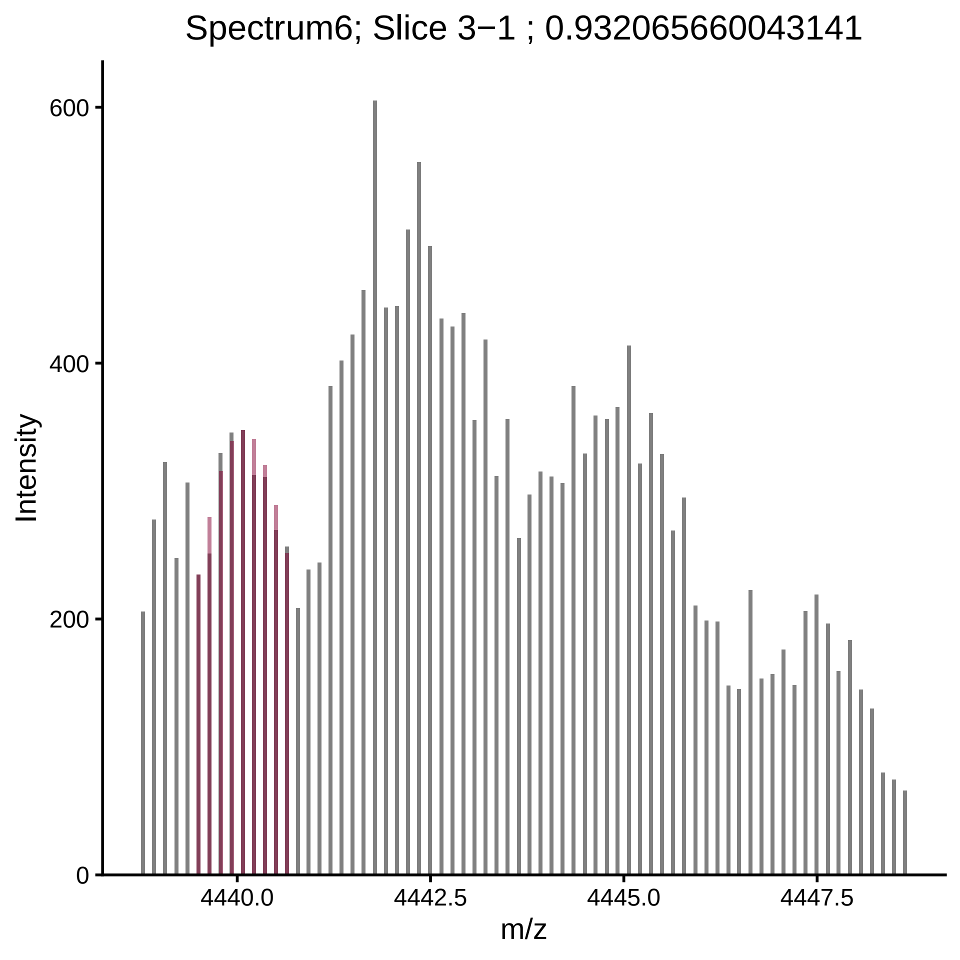
Spectrum6; Slice 3−1 ; 0.932065660043141
Intensity
600
400
200
0
4440.0
4442.5
4445.0
4447.5
m/z

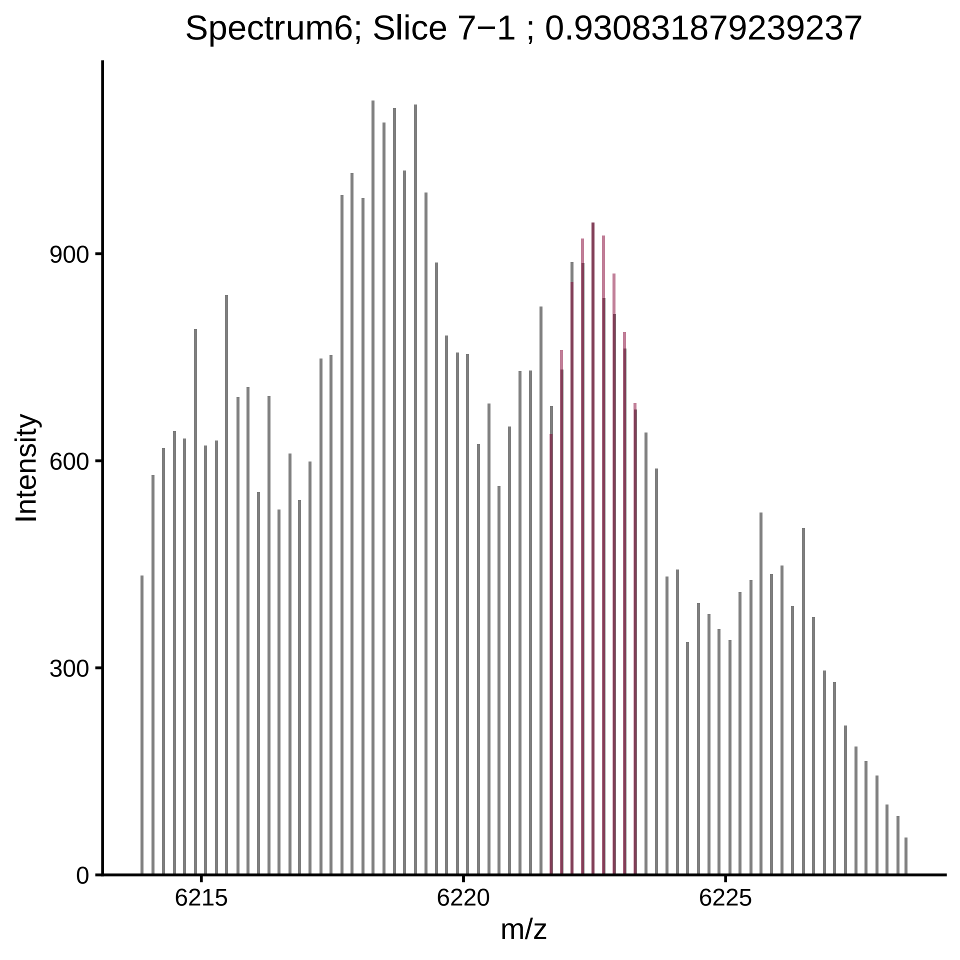
Spectrum6; Slice 7−1 ; 0.930831879239237
Intensity
0
300
600
900
6215
6220
6225
m/z

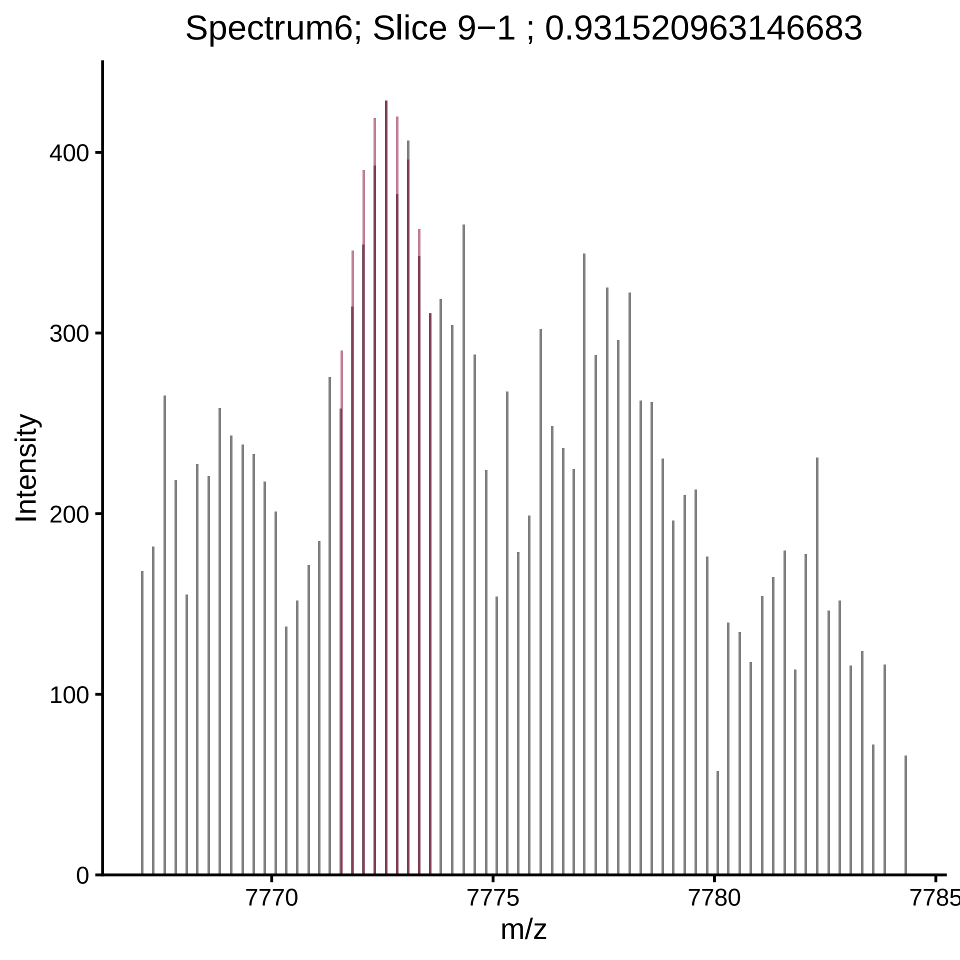

Spectrum6; Slice 9−1 ; 0.931520963146683
Intensity
0
100
200
300
400
7770
7775
7780
7785
m/z

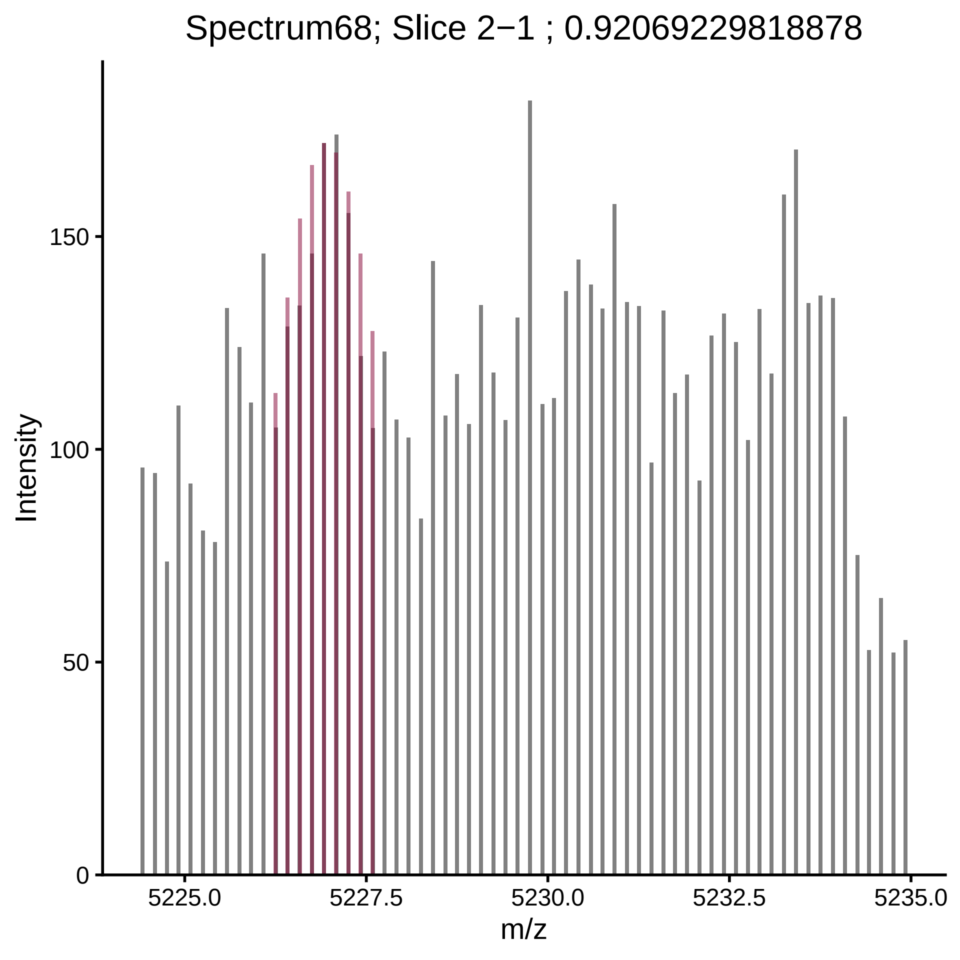

Spectrum68; Slice 2−1 ; 0.92069229818878
Intensity
150
100
50
0
5225.0
5227.5
5230.0
5232.5
5235.0
m/z

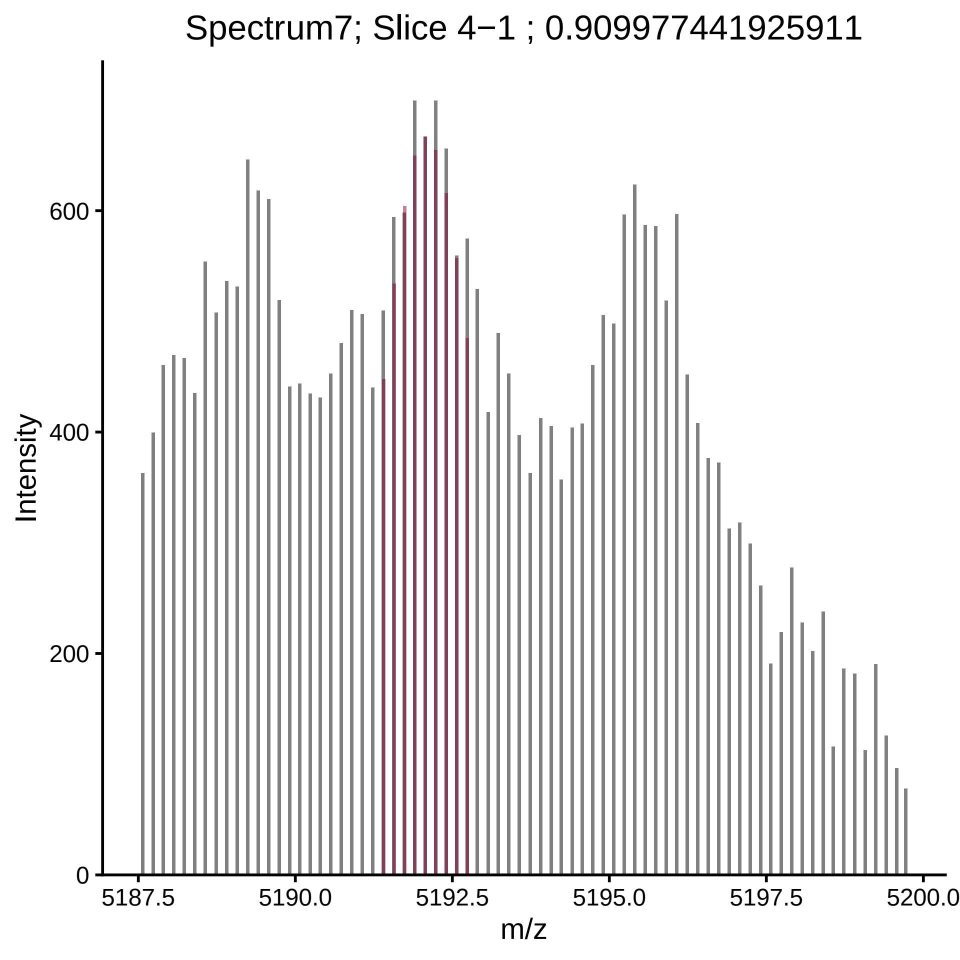

Spectrum7; Slice 4−1 ; 0.909977441925911
Intensity
600
400
200
0
5187.5
5190.0
5192.5
5195.0
5197.5
5200.0
m/z

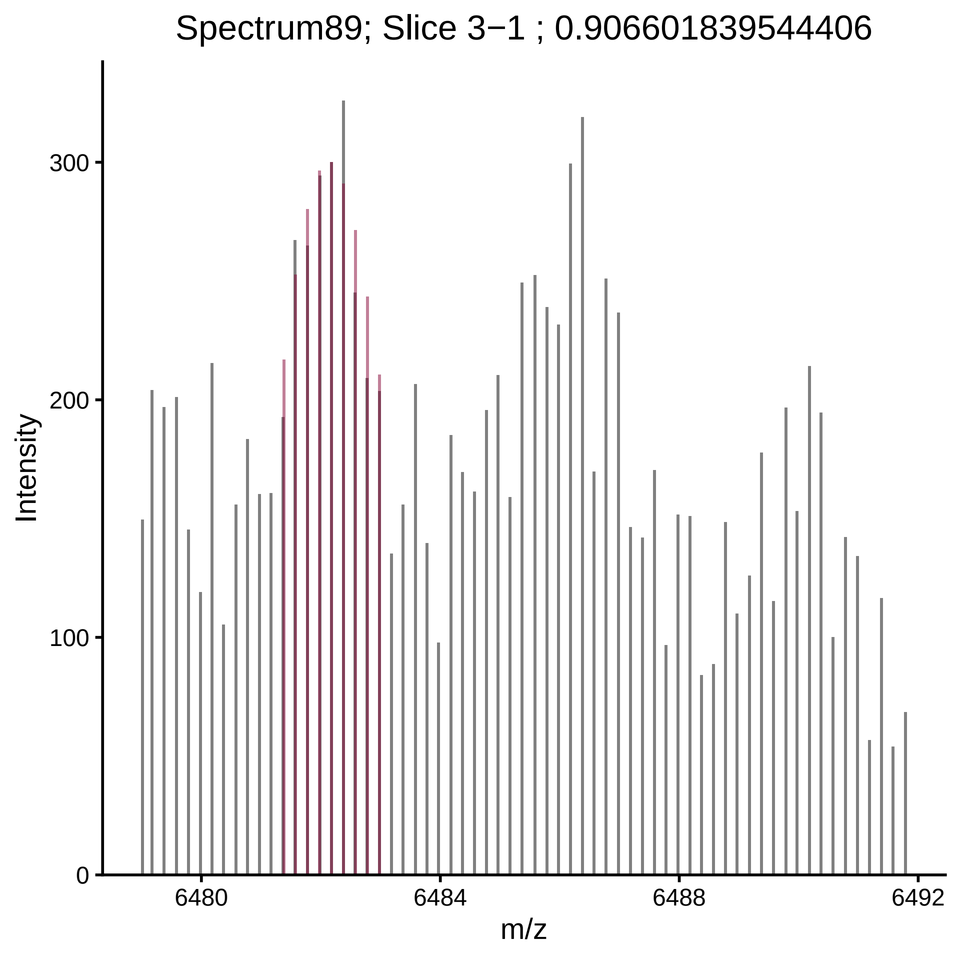

Spectrum89; Slice 3−1 ; 0.906601839544406
Intensity
300
200
100
0
6480
6484
6488
6492
m/z

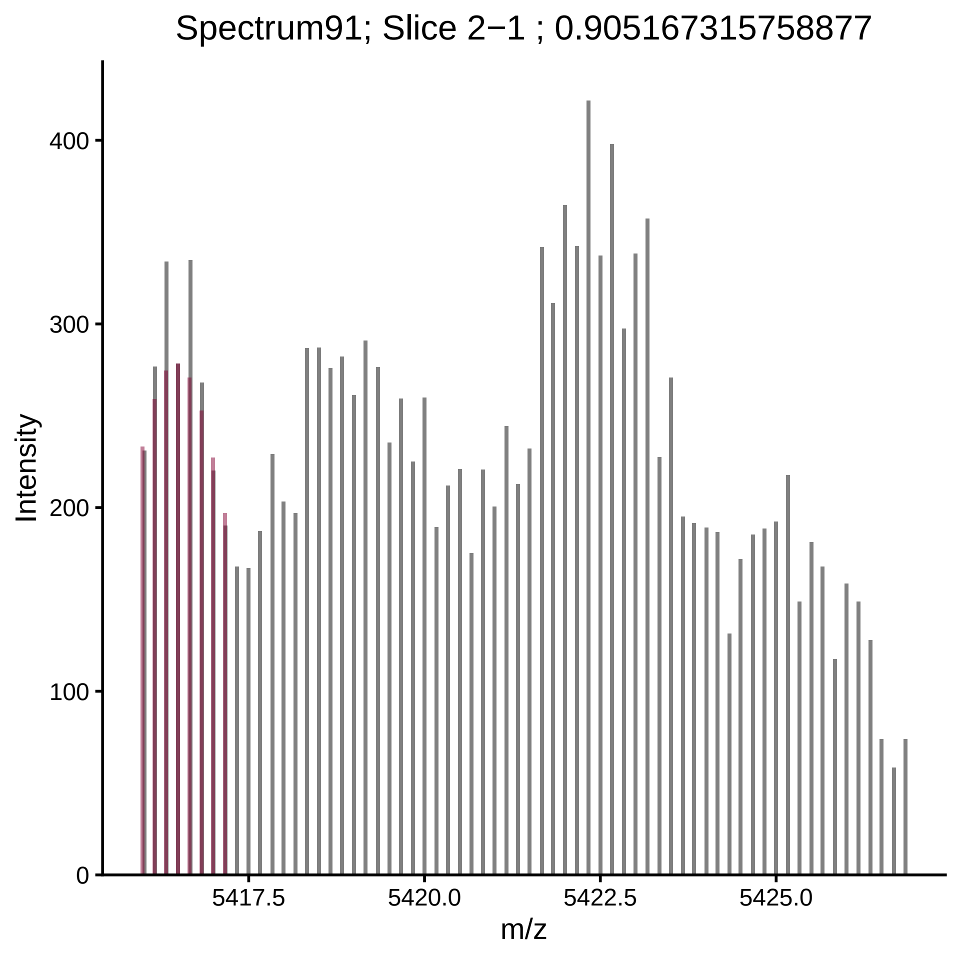
Spectrum91; Slice 2−1 ; 0.905167315758877
Intensity
0
100
200
300
400
5417.5
5420.0
5422.5
5425.0
m/z

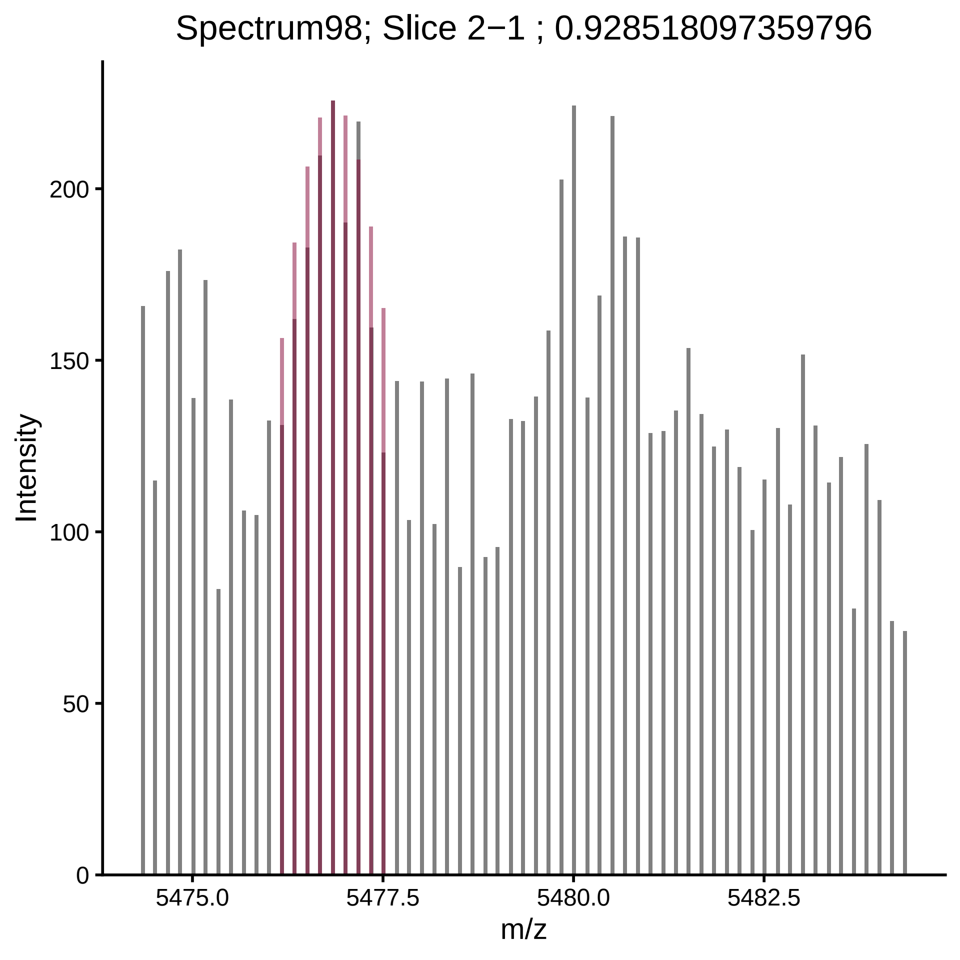

Spectrum98; Slice 2−1 ; 0.928518097359796
Intensity
200
150
100
50
0
5475.0
5477.5
5480.0
5482.5
m/z
