## Supplementary material for "Defining glycoproteoform landscapes through an integrated glycoproteomics approach enabled by high-resolving power proton transfer charge reduction tandem mass spectrometry": FileS2_cleanedSpectra

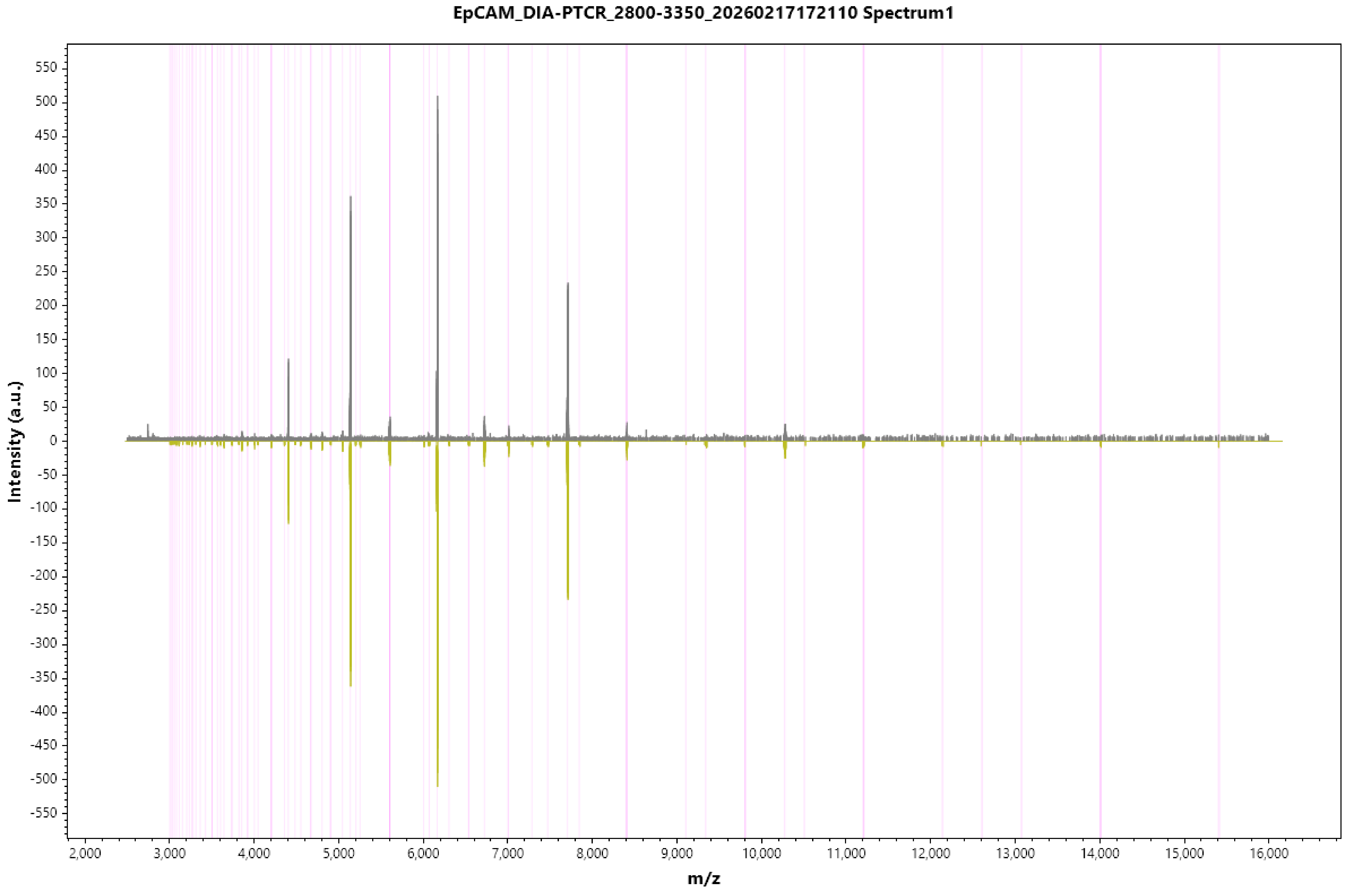
EpCAM_DIA-PTCR_2800-3350_20260217172110 Spectrum1
Intensity (a.u.)
m/z
550
500
450
400
350
300
250
200
150
100
50
0
-50
-100
-150
-200
-250
-300
-350
-400
-450
-500
-550
2,000
3,000
4,000
5,000
6,000
7,000
8,000
9,000
10,000
11,000
12,000
13,000
14,000
15,000
16,000

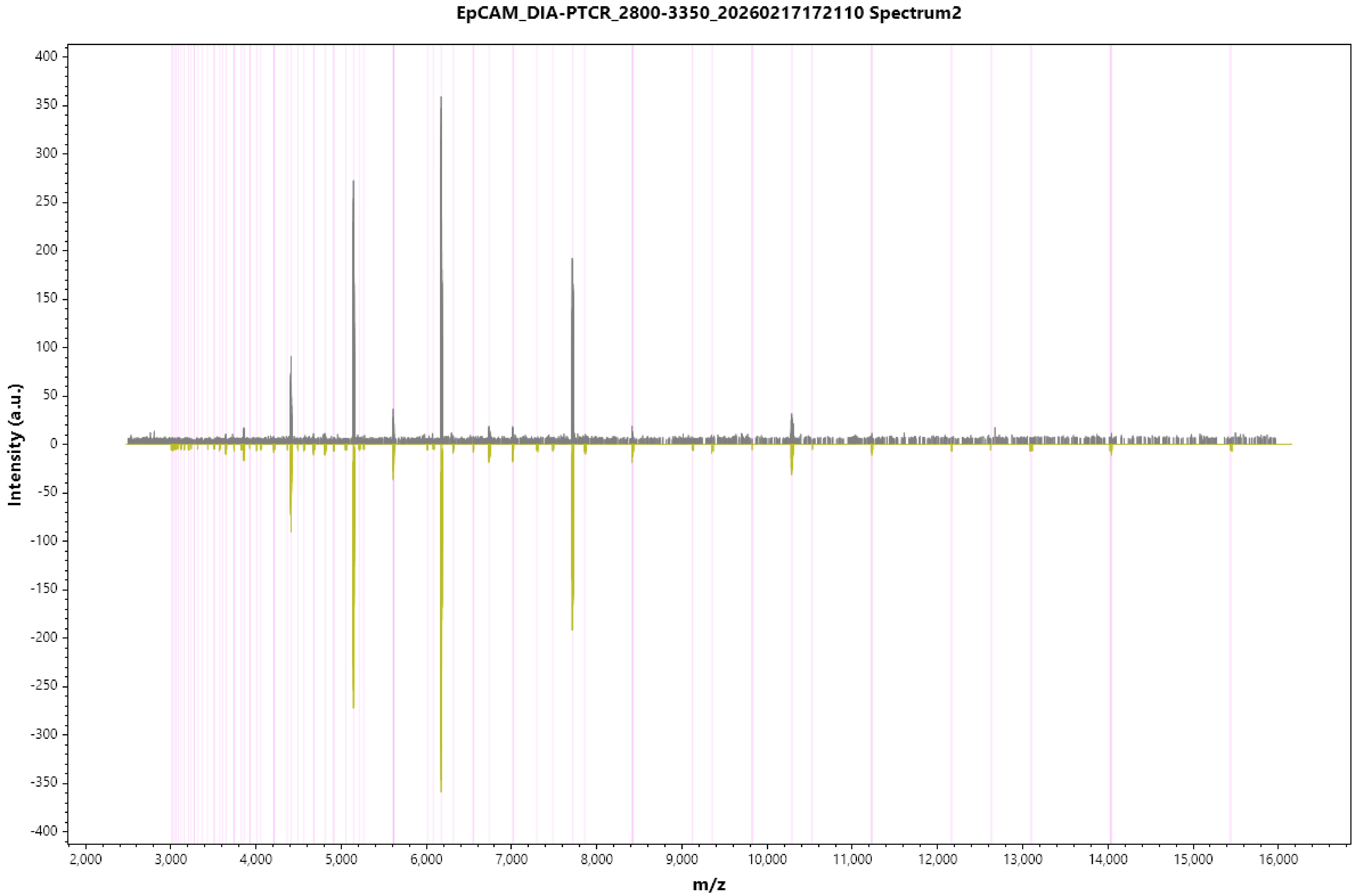
EpCAM_DIA-PTCR_2800-3350_20260217172110 Spectrum2
Intensity (a.u.)
m/z
400
350
300
250
200
150
100
50
0
-50
-100
-150
-200
-250
-300
-350
-400
2,000
3,000
4,000
5,000
6,000
7,000
8,000
9,000
10,000
11,000
12,000
13,000
14,000
15,000
16,000

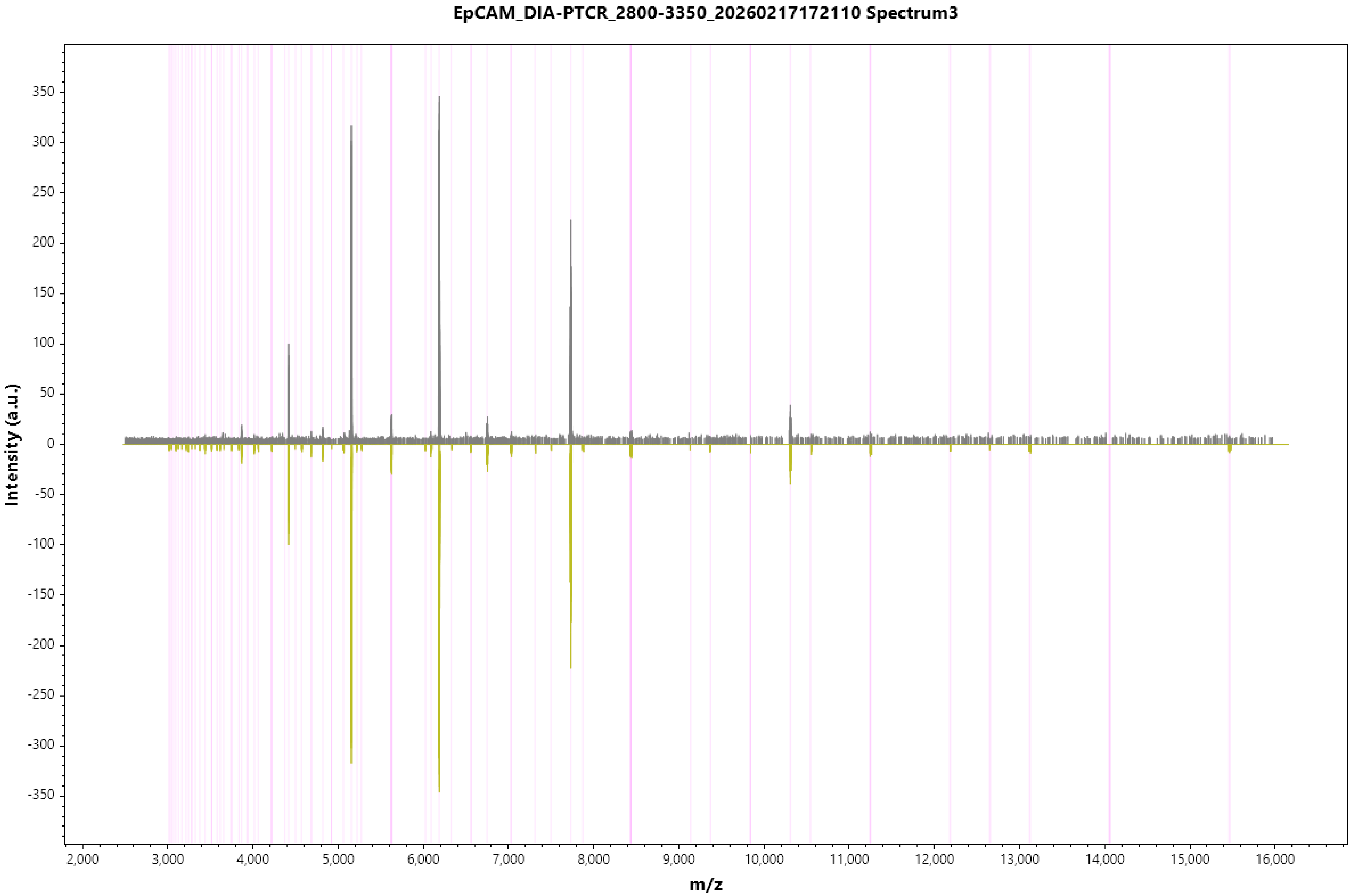
EpCAM_DIA-PTCR_2800-3350_20260217172110 Spectrum3
Intensity (a.u.)
350
300
250
200
150
100
50
0
-50
-100
-150
-200
-250
-300
-350
2,000
3,000
4,000
5,000
6,000
7,000
8,000
9,000
10,000
11,000
12,000
13,000
14,000
15,000
16,000
m/z

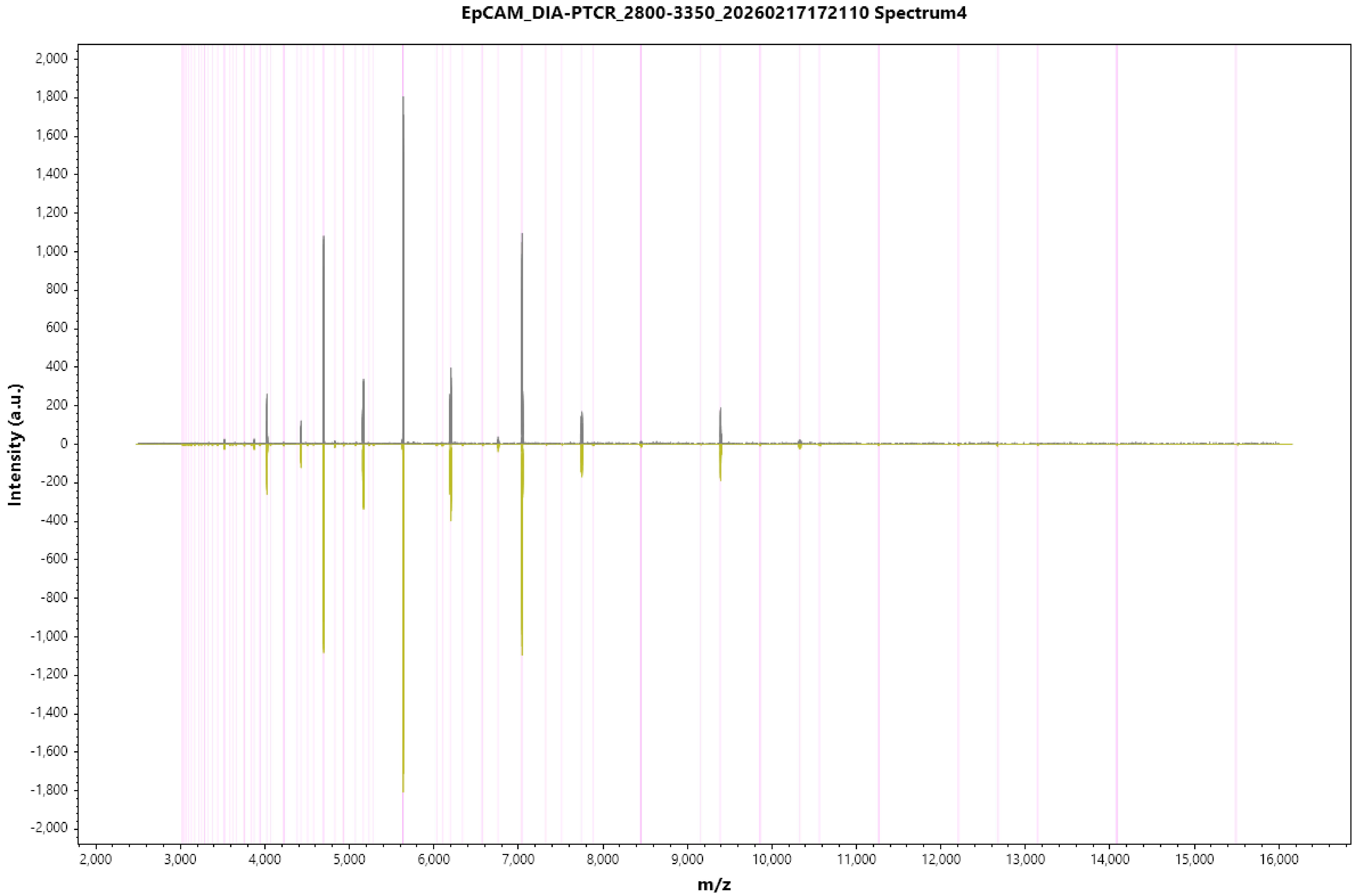

EpCAM_DIA-PTCR_2800-3350_20260217172110 Spectrum4
Intensity (a.u.)
m/z
2,000
1,800
1,600
1,400
1,200
1,000
800
600
400
200
0
-200
-400
-600
-800
-1,000
-1,200
-1,400
-1,600
-1,800
-2,000
2,000
3,000
4,000
5,000
6,000
7,000
8,000
9,000
10,000
11,000
12,000
13,000
14,000
15,000
16,000

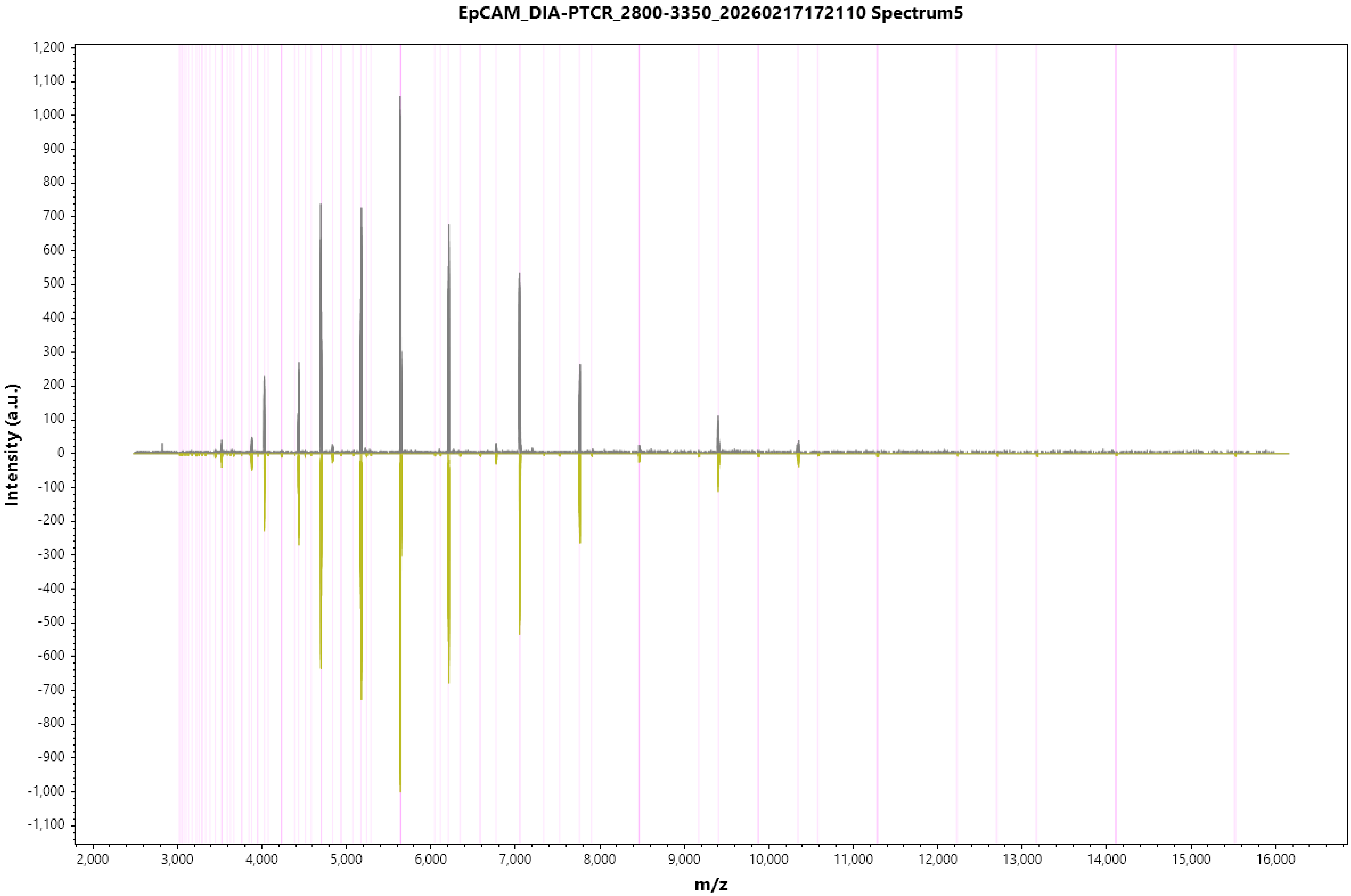
EpCAM_DIA-PTCR_2800-3350_20260217172110 Spectrum5
Intensity (a.u.)
m/z
1,200
1,100
1,000
900
800
700
600
500
400
300
200
100
0
-100
-200
-300
-400
-500
-600
-700
-800
-900
-1,000
-1,100
2,000
3,000
4,000
5,000
6,000
7,000
8,000
9,000
10,000
11,000
12,000
13,000
14,000
15,000
16,000

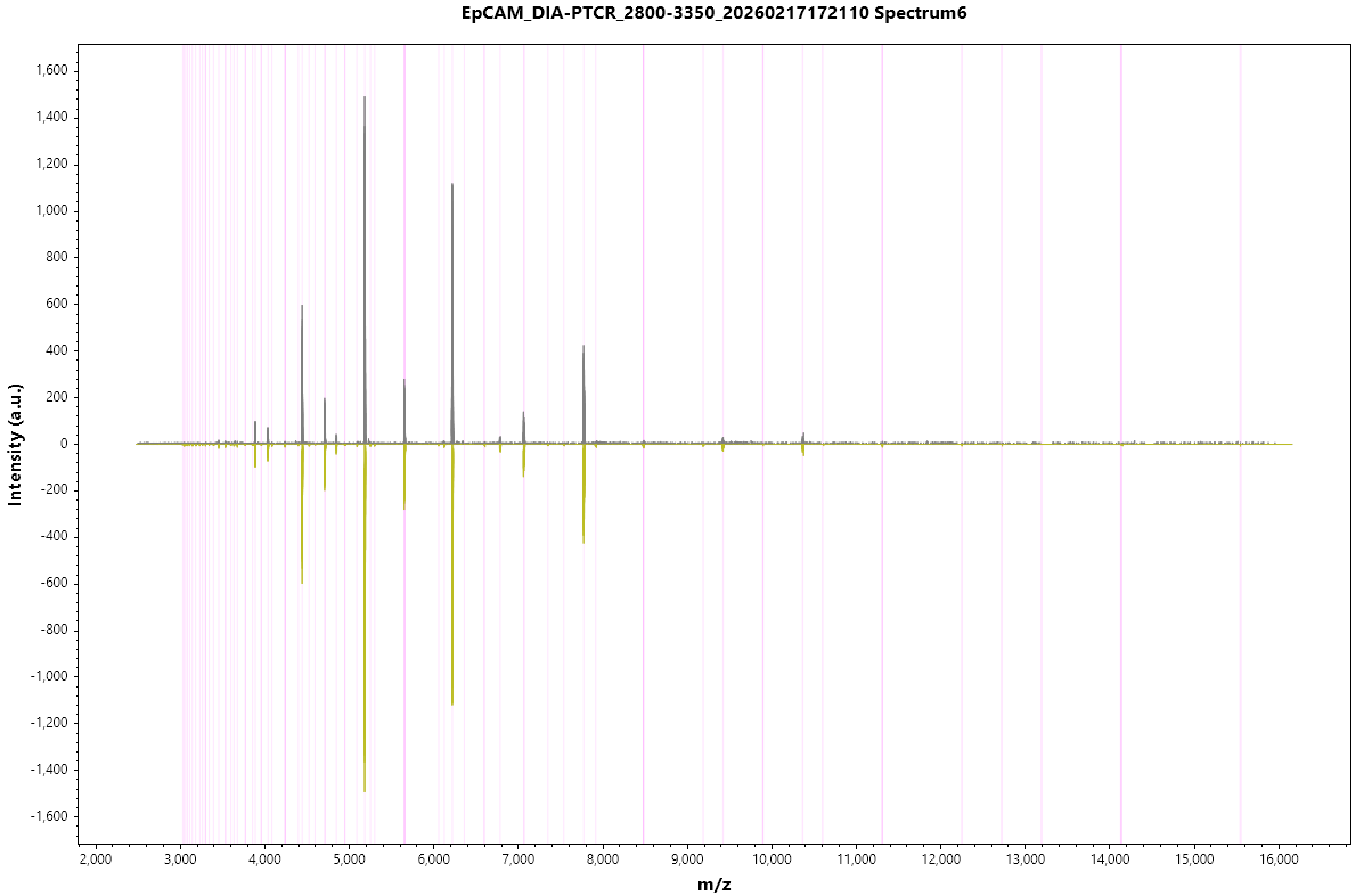
EpCAM_DIA-PTCR_2800-3350_20260217172110 Spectrum6
Intensity (a.u.)
m/z
1,600
1,400
1,200
1,000
800
600
400
200
0
-200
-400
-600
-800
-1,000
-1,200
-1,400
-1,600
2,000
3,000
4,000
5,000
6,000
7,000
8,000
9,000
10,000
11,000
12,000
13,000
14,000
15,000
16,000

EpCAM_DIA-PTCR_2800-3350_20260217172110 Spectrum7

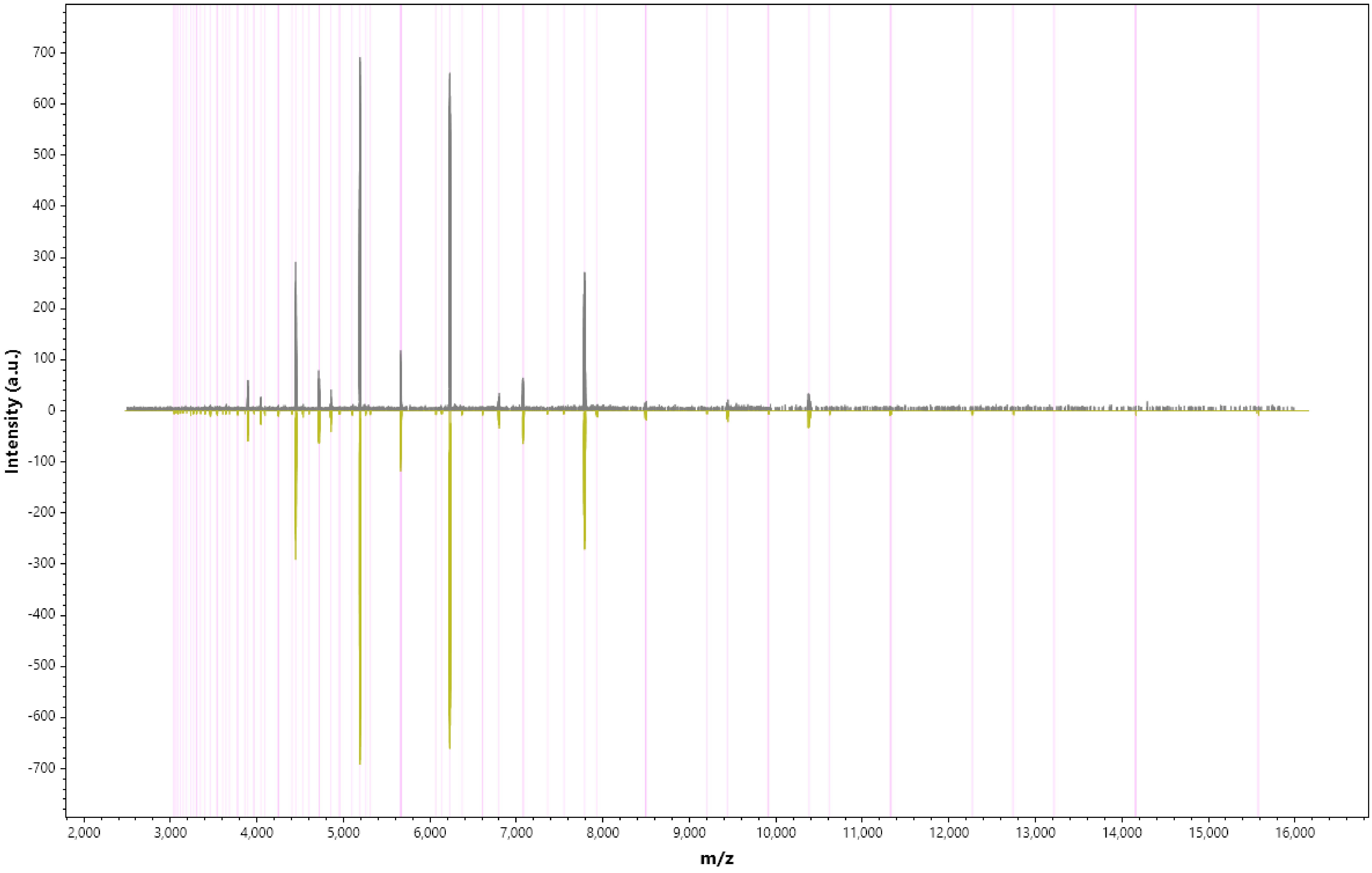

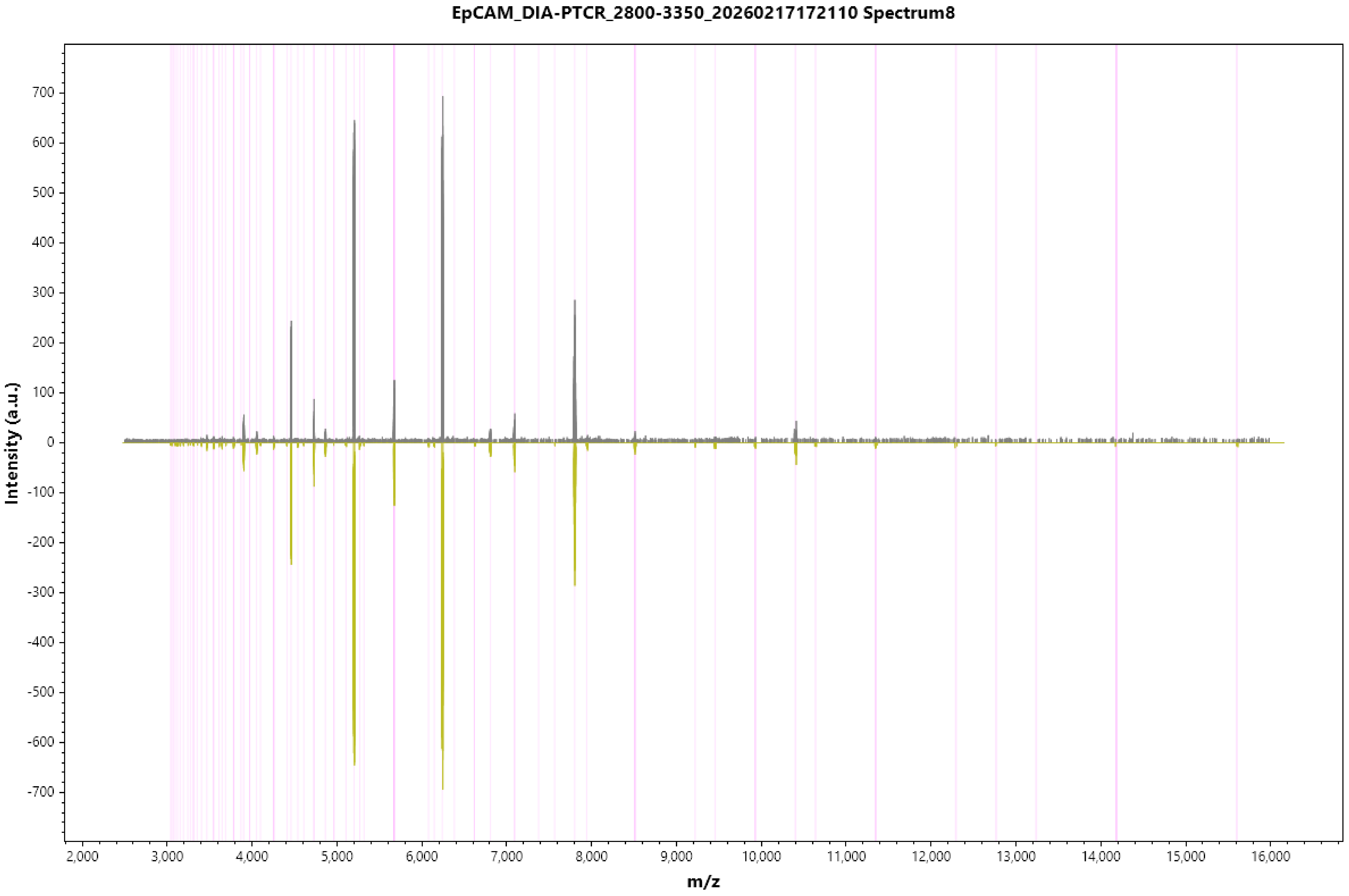
EpCAM_DIA-PTCR_2800-3350_20260217172110 Spectrum8
Intensity (a.u.)
m/z
700
600
500
400
300
200
100
0
-100
-200
-300
-400
-500
-600
-700
2,000
3,000
4,000
5,000
6,000
7,000
8,000
9,000
10,000
11,000
12,000
13,000
14,000
15,000
16,000

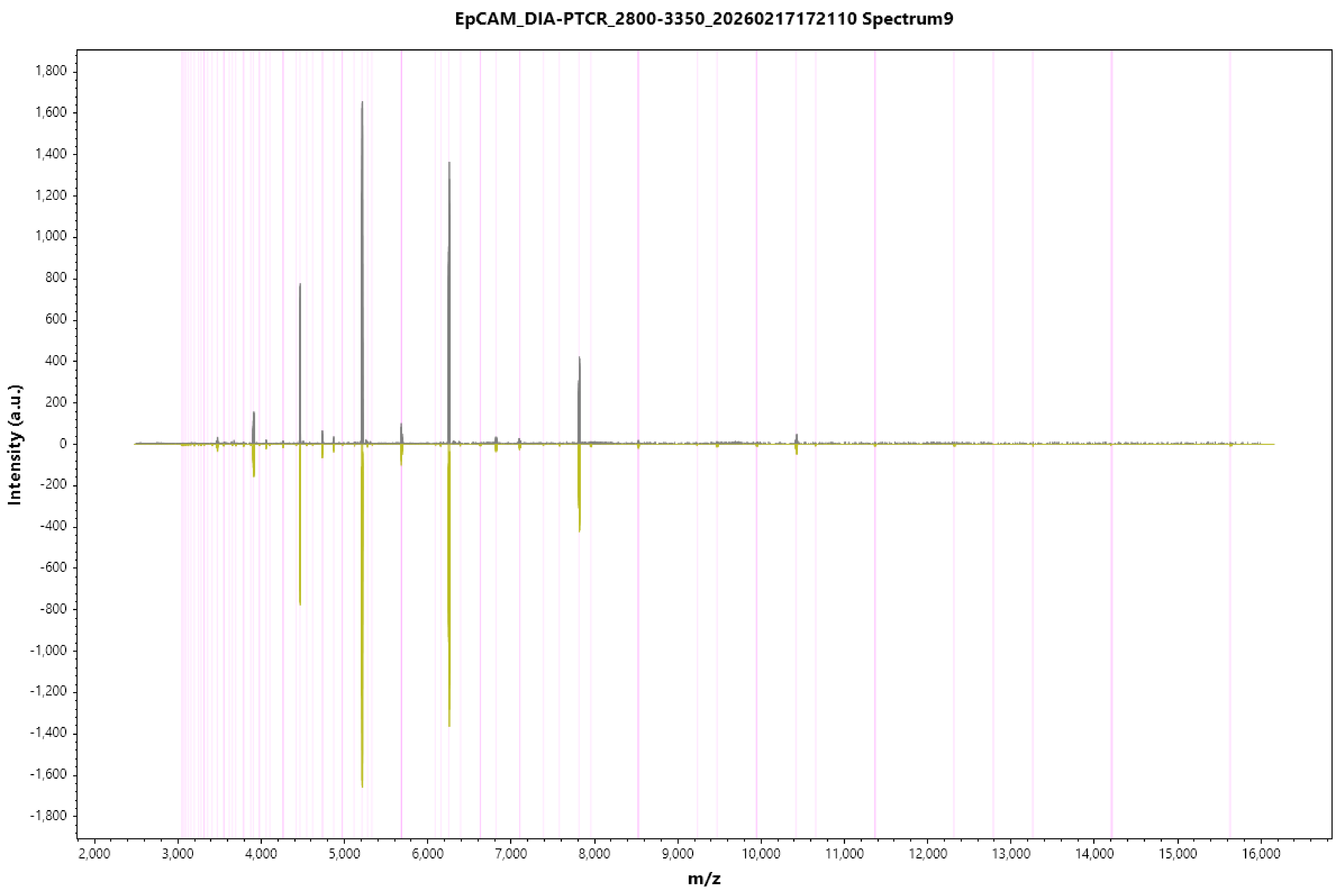
EpCAM_DIA-PTCR_2800-3350_20260217172110 Spectrum9
Intensity (a.u.)
m/z
1,800
1,600
1,400
1,200
1,000
800
600
400
200
0
-200
-400
-600
-800
-1,000
-1,200
-1,400
-1,600
-1,800
2,000
3,000
4,000
5,000
6,000
7,000
8,000
9,000
10,000
11,000
12,000
13,000
14,000
15,000
16,000

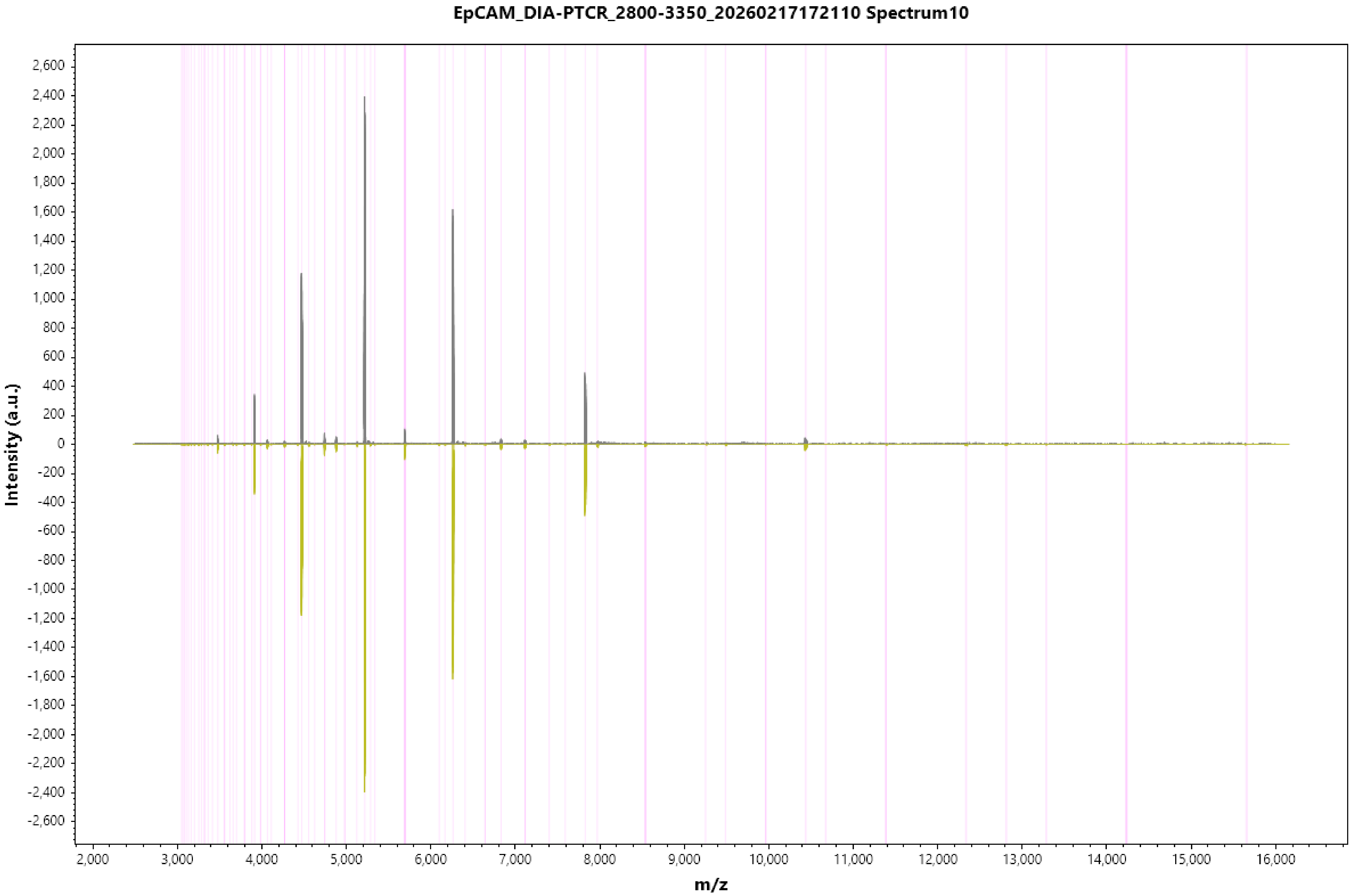

EpCAM_DIA-PTCR_2800-3350_20260217172110 Spectrum10
Intensity (a.u.)
m/z
2,600
2,400
2,200
2,000
1,800
1,600
1,400
1,200
1,000
800
600
400
200
0
-200
-400
-600
-800
-1,000
-1,200
-1,400
-1,600
-1,800
-2,000
-2,200
-2,400
-2,600
2,000
3,000
4,000
5,000
6,000
7,000
8,000
9,000
10,000
11,000
12,000
13,000
14,000
15,000
16,000

EpCAM_DIA-PTCR_2800-3350_20260217172110 Spectrum11

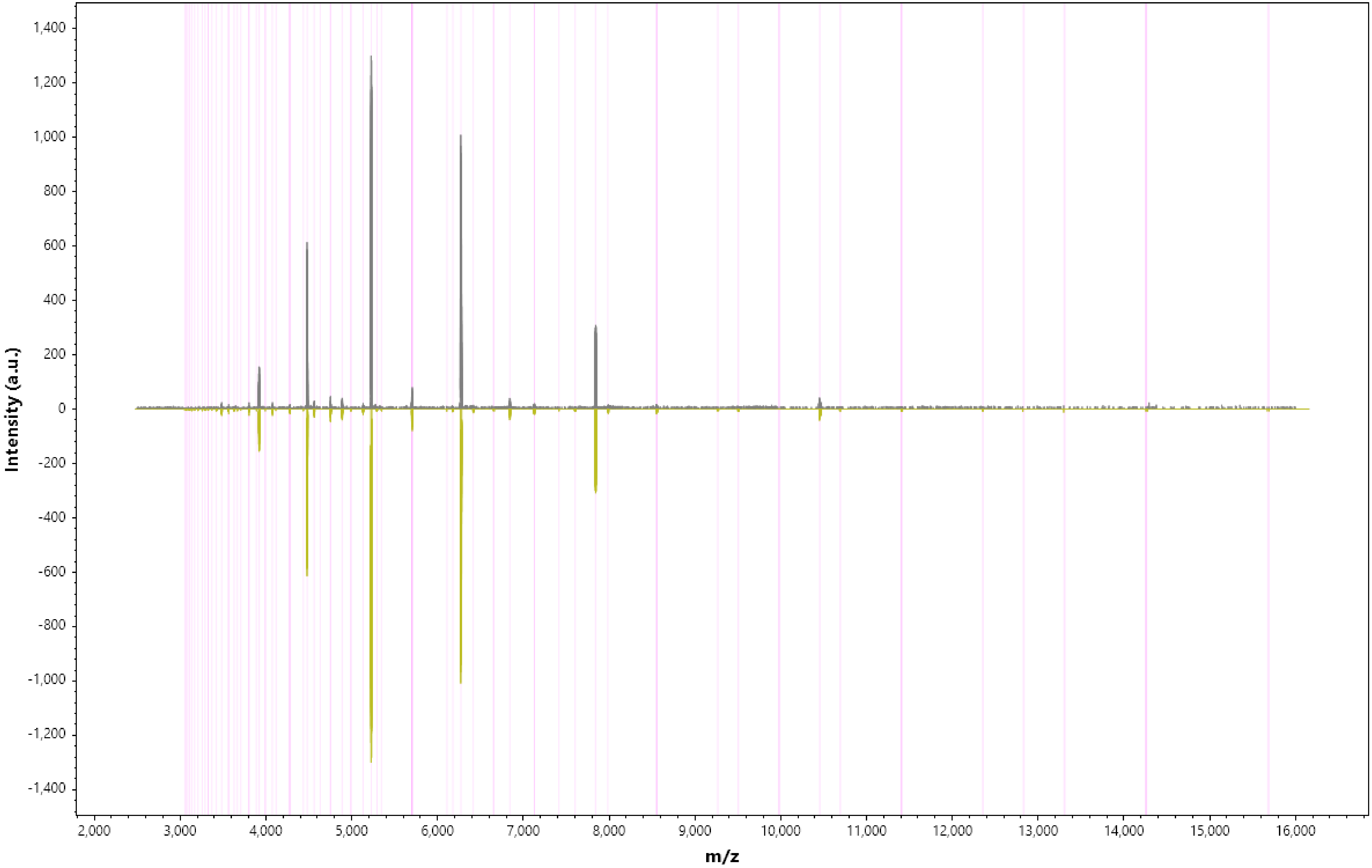

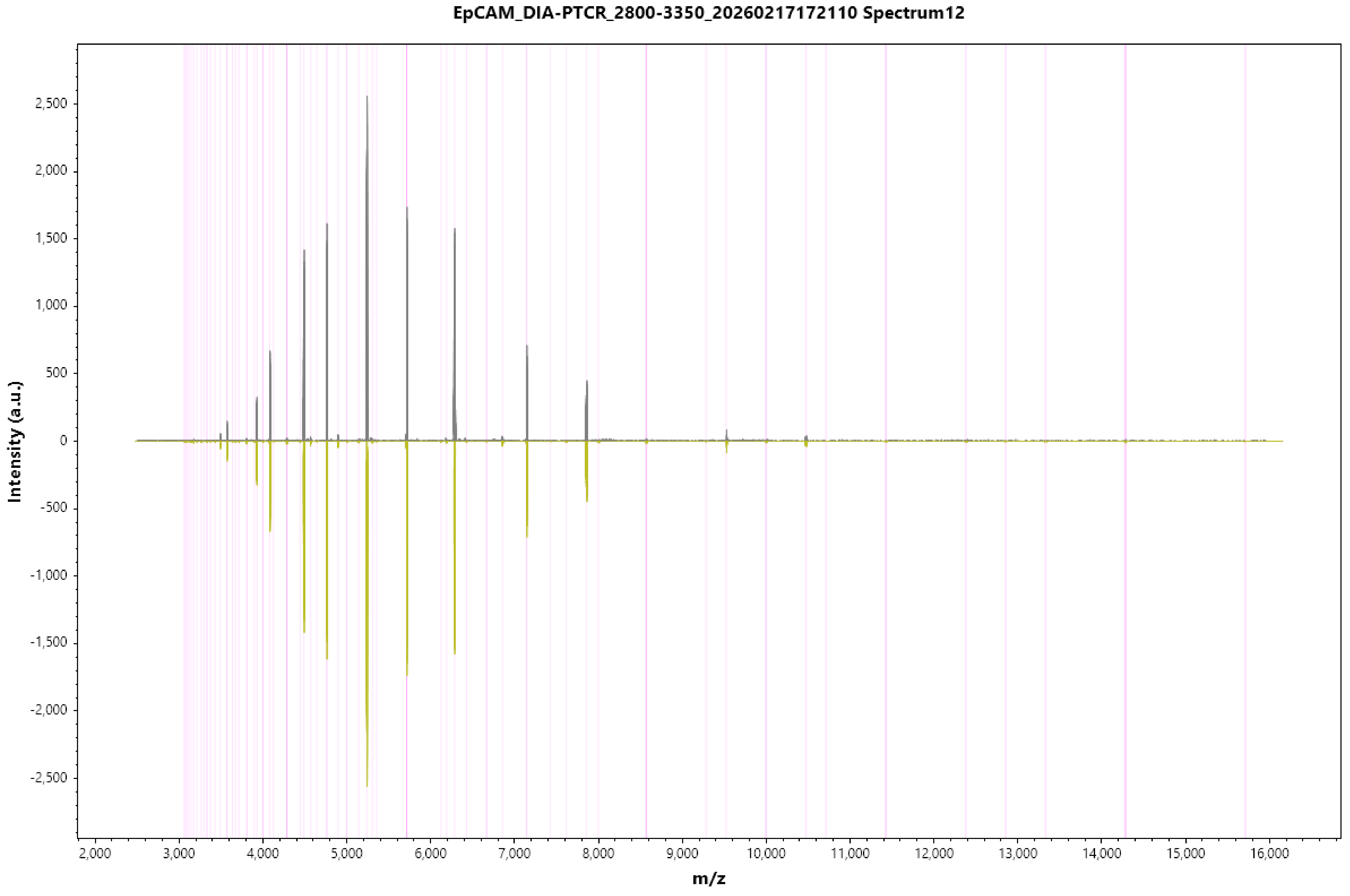
EpCAM_DIA-PTCR_2800-3350_20260217172110 Spectrum12
Intensity (a.u.)
2,500
2,000
1,500
1,000
500
0
-500
-1,000
-1,500
-2,000
-2,500
2,000
3,000
4,000
5,000
6,000
7,000
8,000
9,000
10,000
11,000
12,000
13,000
14,000
15,000
16,000
m/z

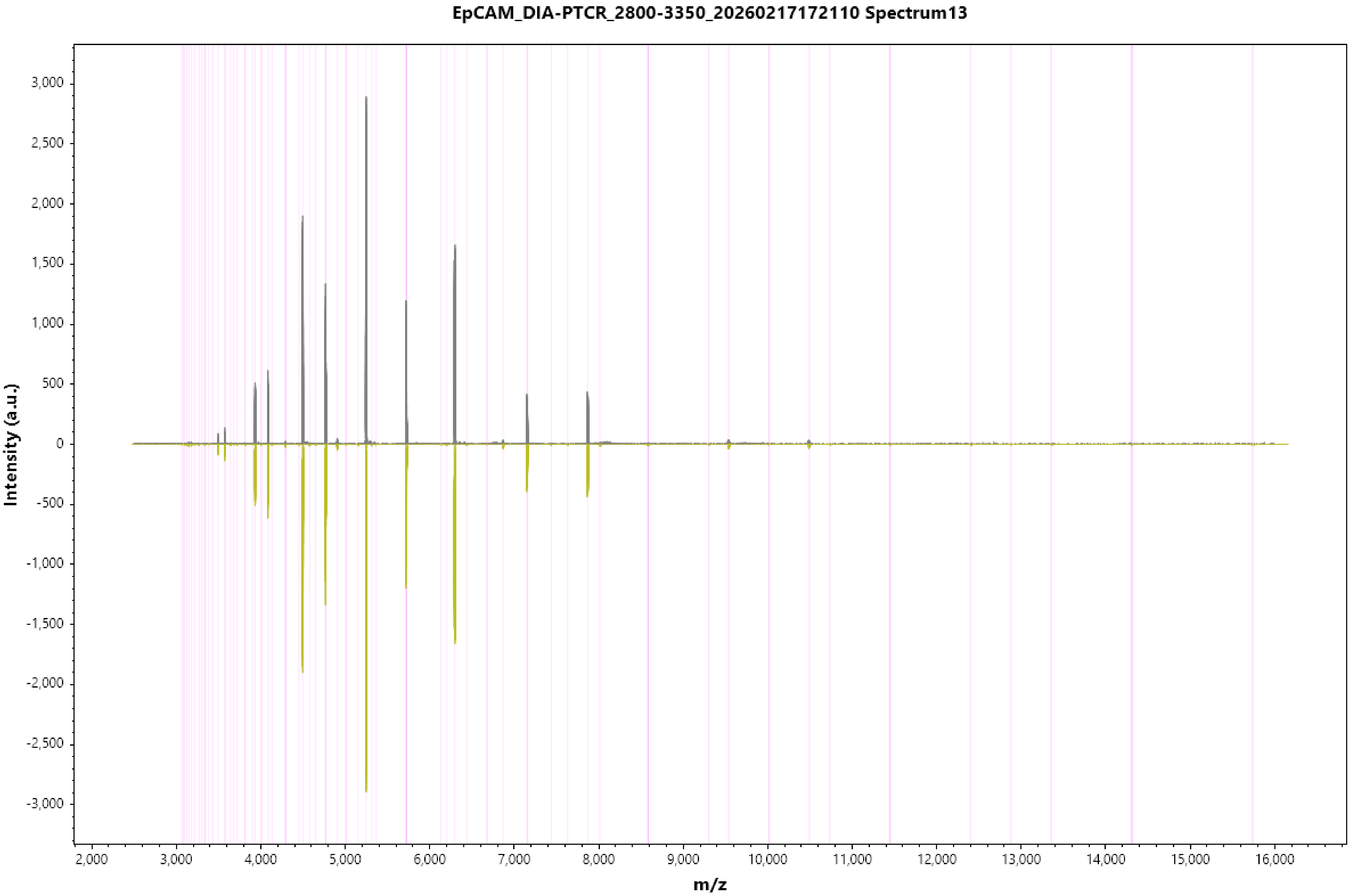
EpCAM_DIA-PTCR_2800-3350_20260217172110 Spectrum13
Intensity (a.u.)
m/z
3,000
2,500
2,000
1,500
1,000
500
0
-500
-1,000
-1,500
-2,000
-2,500
-3,000
2,000
3,000
4,000
5,000
6,000
7,000
8,000
9,000
10,000
11,000
12,000
13,000
14,000
15,000
16,000

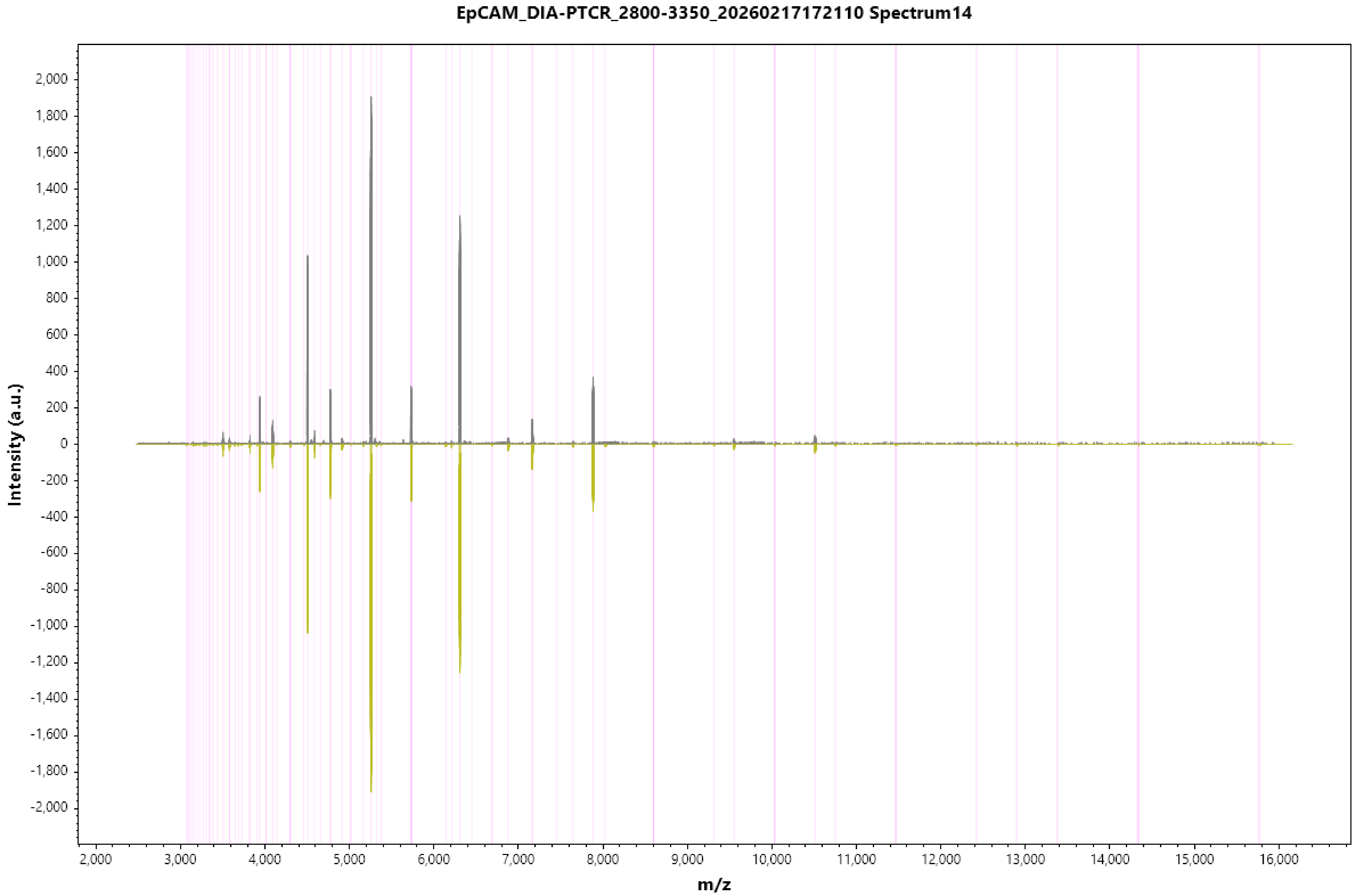

EpCAM_DIA-PTCR_2800-3350_20260217172110 Spectrum14
Intensity (a.u.)
m/z

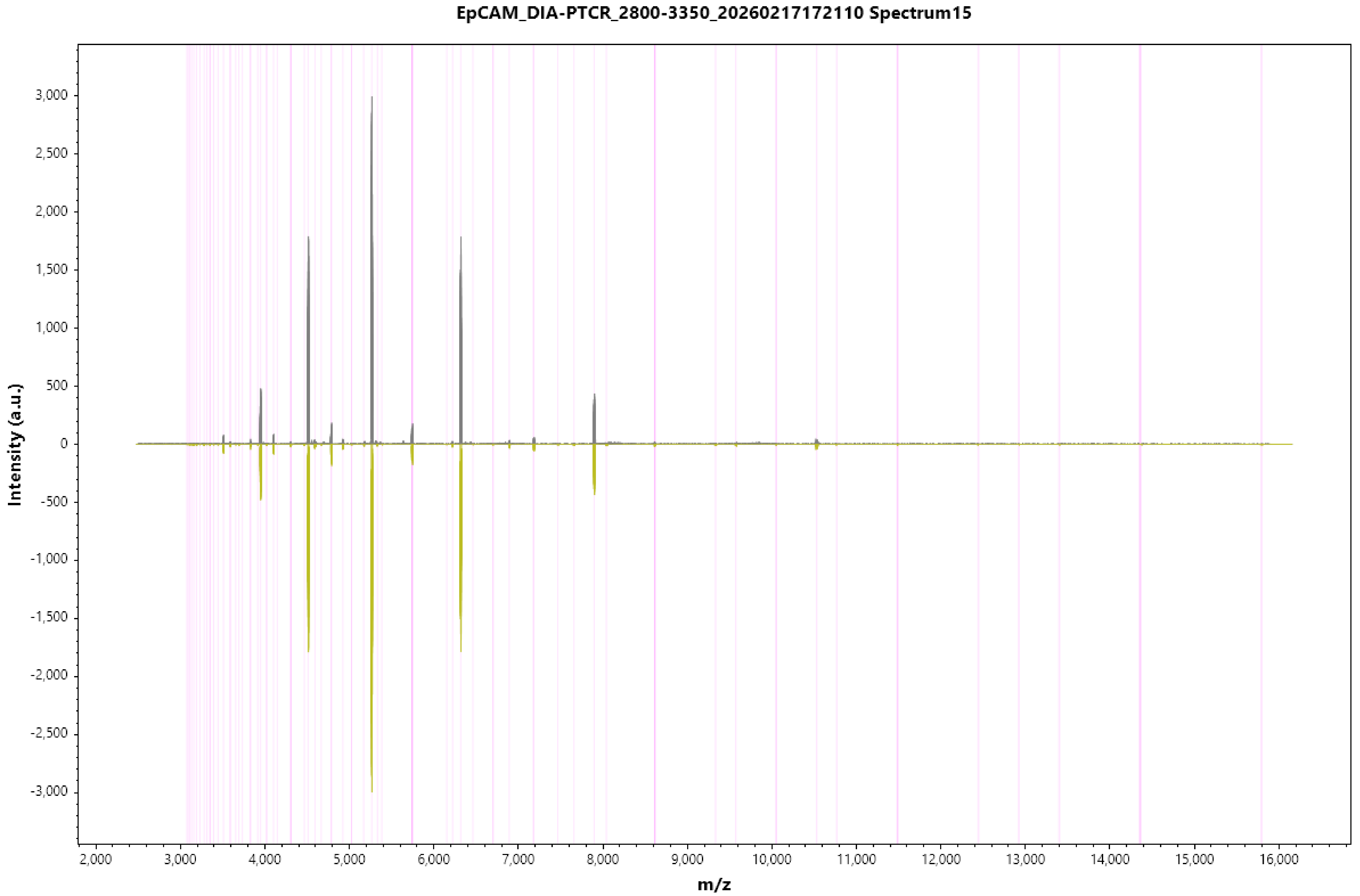
EpCAM_DIA-PTCR_2800-3350_20260217172110 Spectrum15
Intensity (a.u.)
m/z
3,000
2,500
2,000
1,500
1,000
500
0
-500
-1,000
-1,500
-2,000
-2,500
-3,000
2,000
3,000
4,000
5,000
6,000
7,000
8,000
9,000
10,000
11,000
12,000
13,000
14,000
15,000
16,000

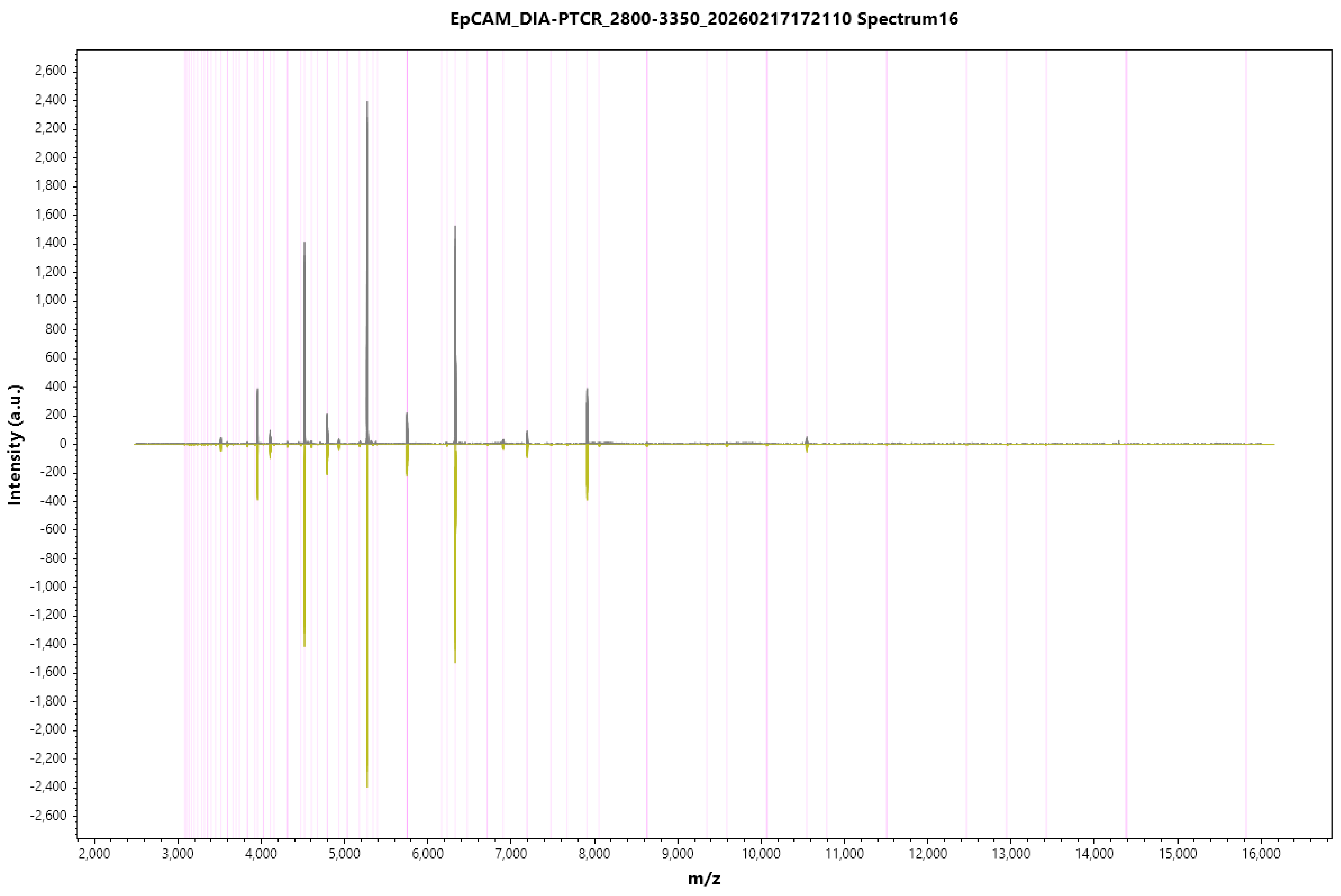
EpCAM_DIA-PTCR_2800-3350_20260217172110 Spectrum16
Intensity (a.u.)
m/z
2,600
2,400
2,200
2,000
1,800
1,600
1,400
1,200
1,000
800
600
400
200
0
-200
-400
-600
-800
-1,000
-1,200
-1,400
-1,600
-1,800
-2,000
-2,200
-2,400
-2,600
2,000
3,000
4,000
5,000
6,000
7,000
8,000
9,000
10,000
11,000
12,000
13,000
14,000
15,000
16,000

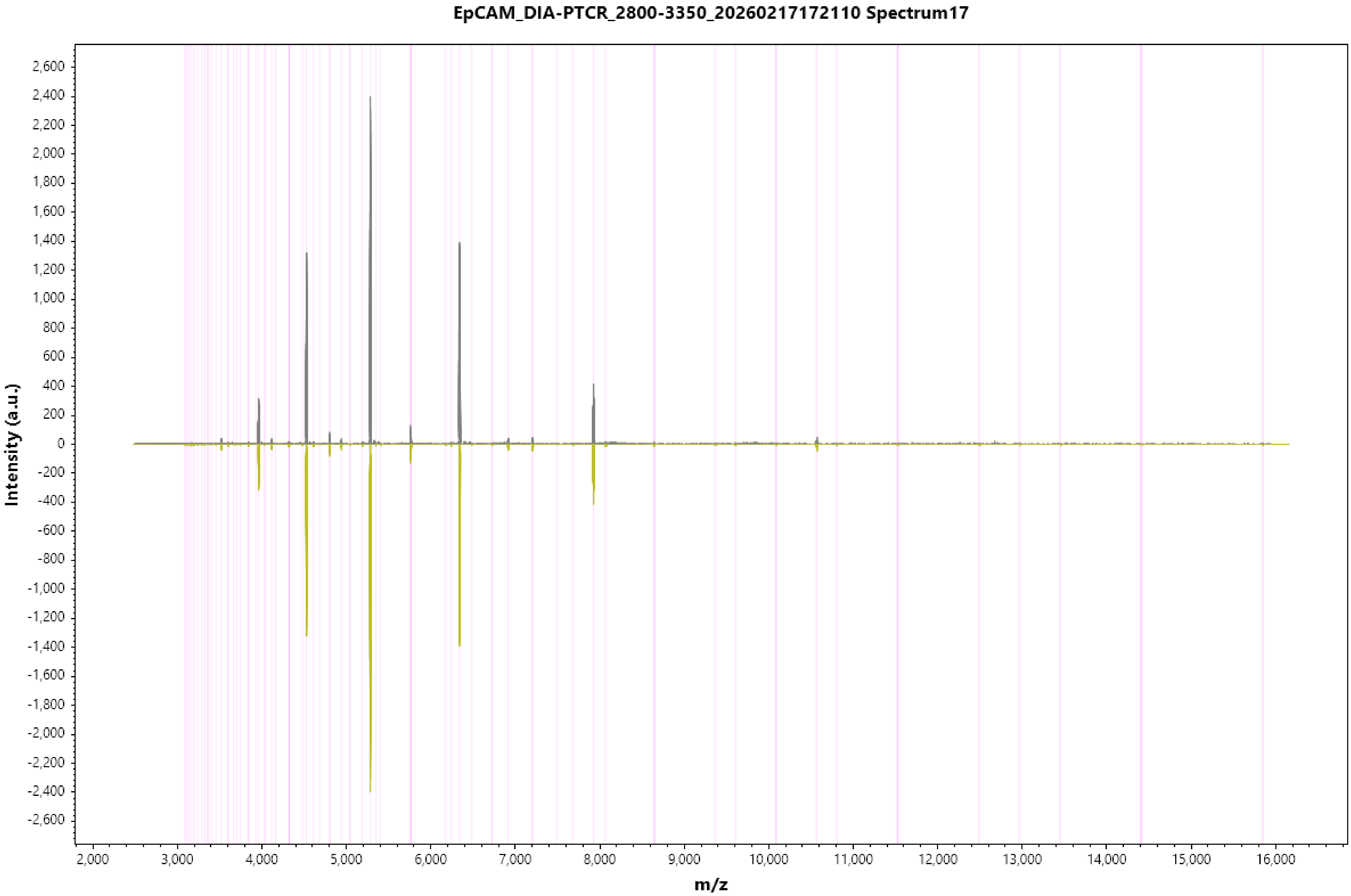
EpCAM_DIA-PTCR_2800-3350_20260217172110 Spectrum17
Intensity (a.u.)
m/z
2,600
2,400
2,200
2,000
1,800
1,600
1,400
1,200
1,000
800
600
400
200
0
-200
-400
-600
-800
-1,000
-1,200
-1,400
-1,600
-1,800
-2,000
-2,200
-2,400
-2,600
2,000
3,000
4,000
5,000
6,000
7,000
8,000
9,000
10,000
11,000
12,000
13,000
14,000
15,000
16,000

### EpCAM_DIA-PTCR_2800-3350_20260217172110 Spectrum18

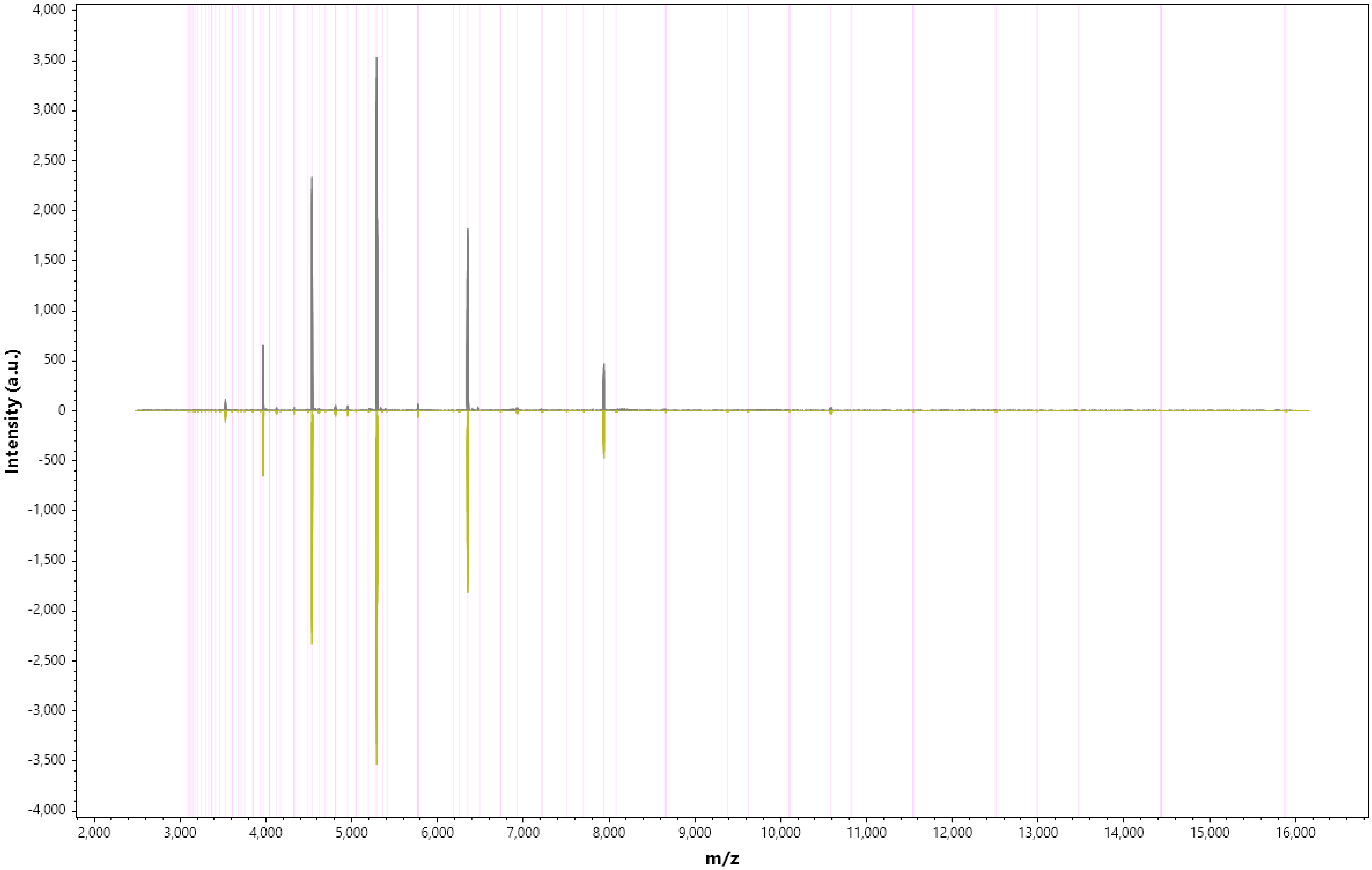

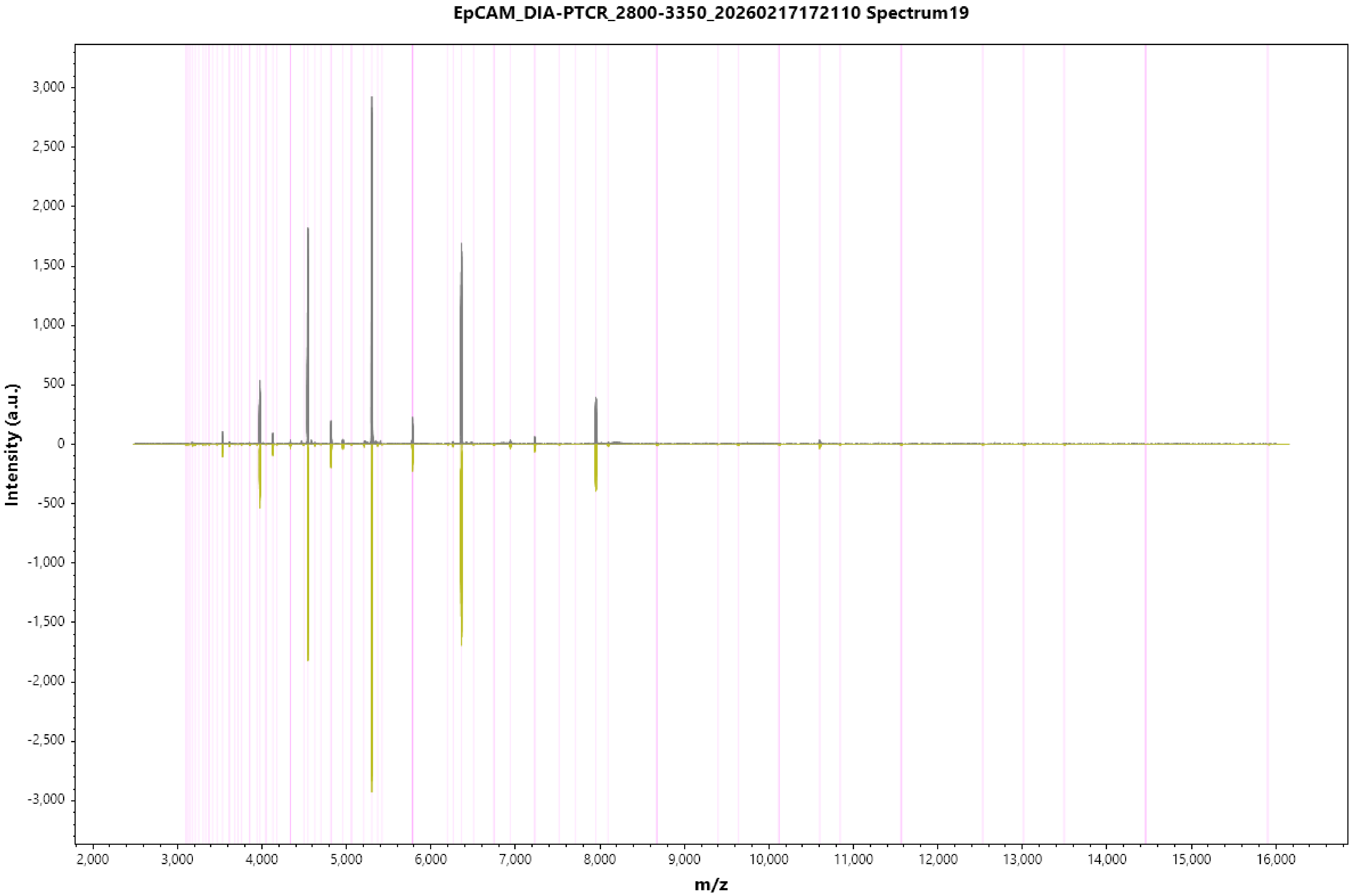
EpCAM_DIA-PTCR_2800-3350_20260217172110 Spectrum19
Intensity (a.u.)
3,000
2,500
2,000
1,500
1,000
500
0
-500
-1,000
-1,500
-2,000
-2,500
-3,000
2,000
3,000
4,000
5,000
6,000
7,000
8,000
9,000
10,000
11,000
12,000
13,000
14,000
15,000
16,000
m/z

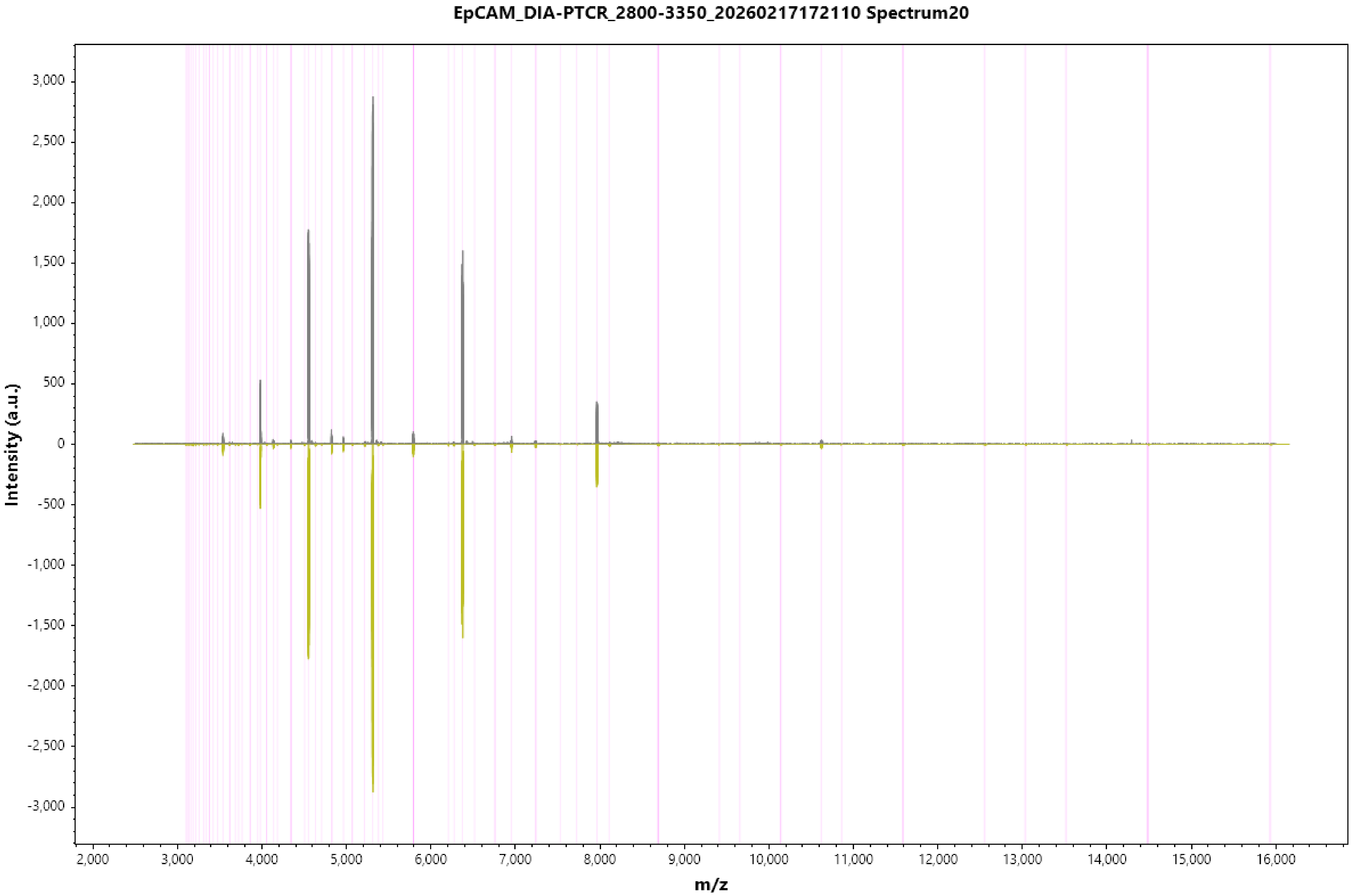

EpCAM_DIA-PTCR_2800-3350_20260217172110 Spectrum20
Intensity (a.u.)
3,000
2,500
2,000
1,500
1,000
500
0
-500
-1,000
-1,500
-2,000
-2,500
-3,000
m/z
2,000
3,000
4,000
5,000
6,000
7,000
8,000
9,000
10,000
11,000
12,000
13,000
14,000
15,000
16,000

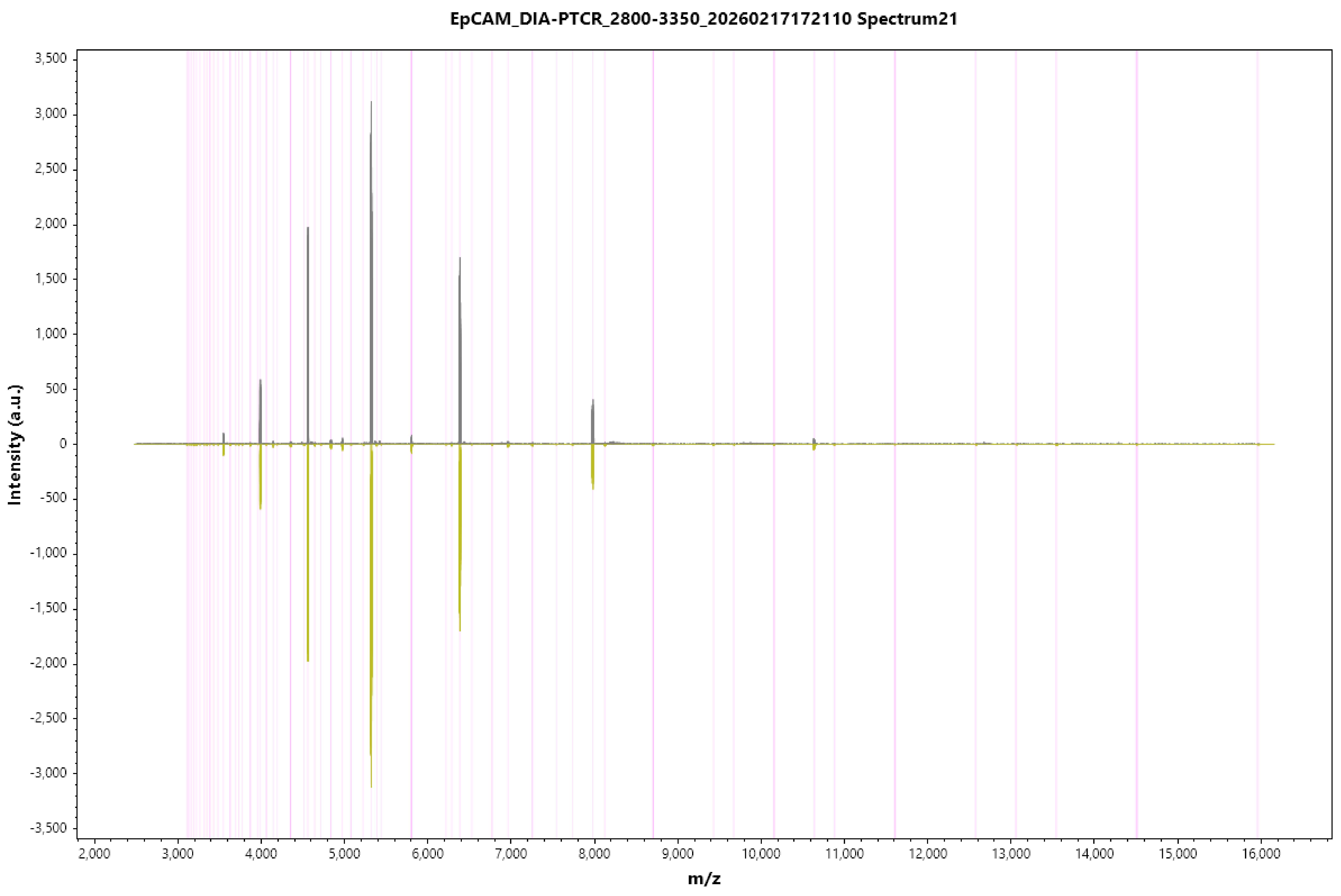
EpCAM_DIA-PTCR_2800-3350_20260217172110 Spectrum21
Intensity (a.u.)
3,500
3,000
2,500
2,000
1,500
1,000
500
0
-500
-1,000
-1,500
-2,000
-2,500
-3,000
-3,500
2,000
3,000
4,000
5,000
6,000
7,000
8,000
9,000
10,000
11,000
12,000
13,000
14,000
15,000
16,000
m/z

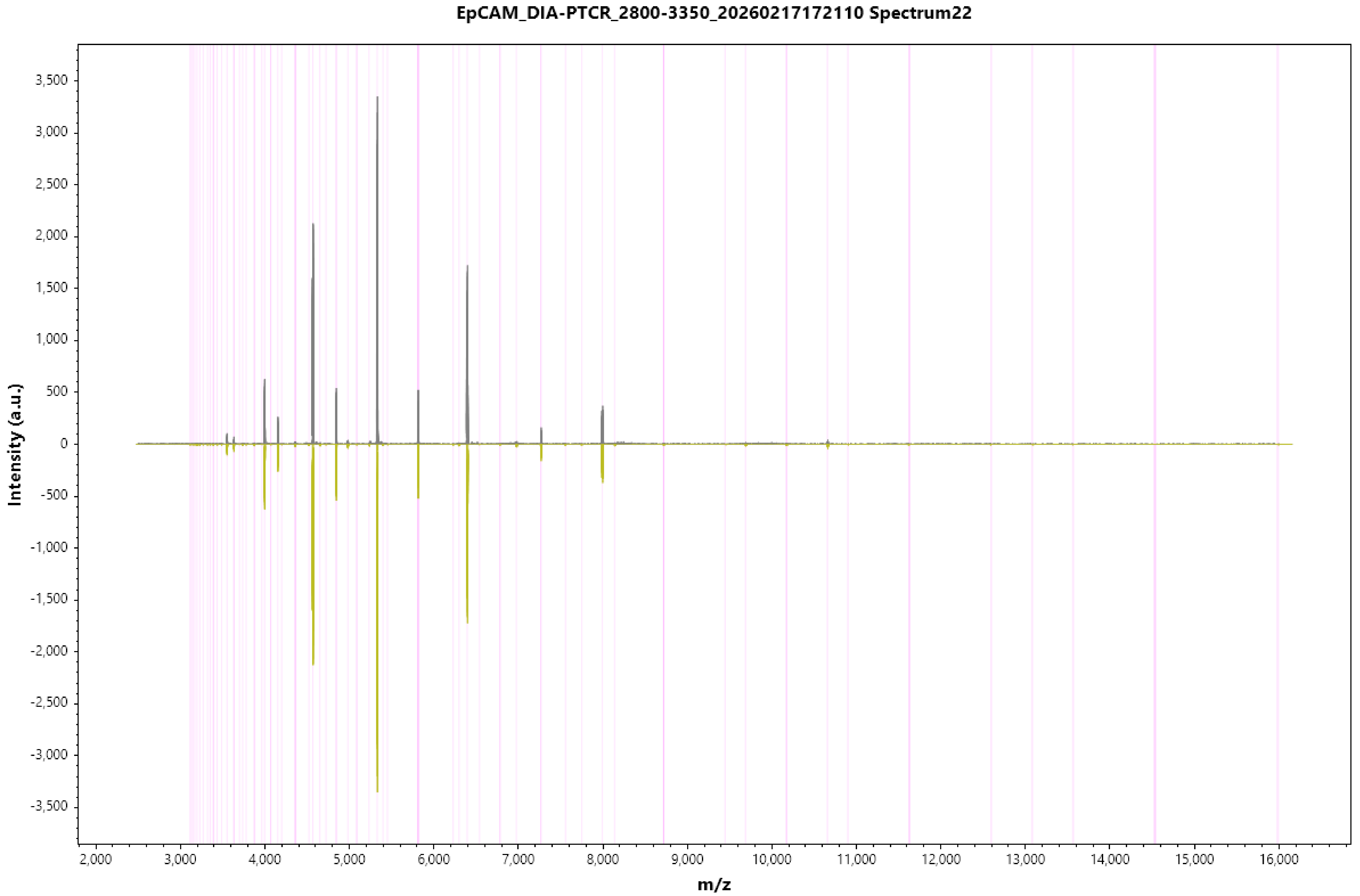

EpCAM_DIA-PTCR_2800-3350_20260217172110 Spectrum22
Intensity (a.u.)
m/z
3,500
3,000
2,500
2,000
1,500
1,000
500
0
-500
-1,000
-1,500
-2,000
-2,500
-3,000
-3,500
2,000
3,000
4,000
5,000
6,000
7,000
8,000
9,000
10,000
11,000
12,000
13,000
14,000
15,000
16,000

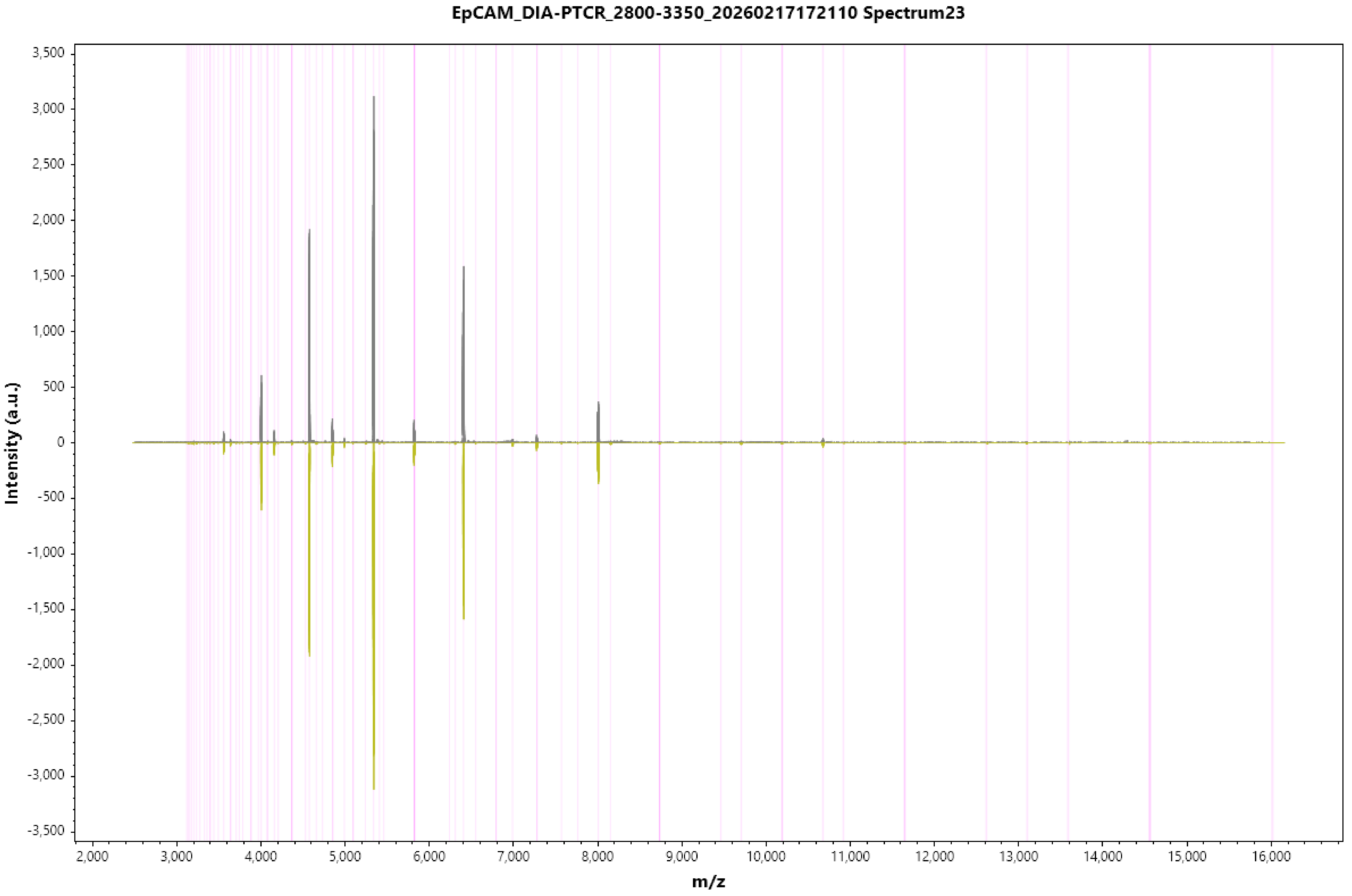
EpCAM_DIA-PTCR_2800-3350_20260217172110 Spectrum23
Intensity (a.u.)
3,500
3,000
2,500
2,000
1,500
1,000
500
0
-500
-1,000
-1,500
-2,000
-2,500
-3,000
-3,500
2,000
3,000
4,000
5,000
6,000
7,000
8,000
9,000
10,000
11,000
12,000
13,000
14,000
15,000
16,000
m/z

EpCAM_DIA-PTCR_2800-3350_20260217172110 Spectrum24

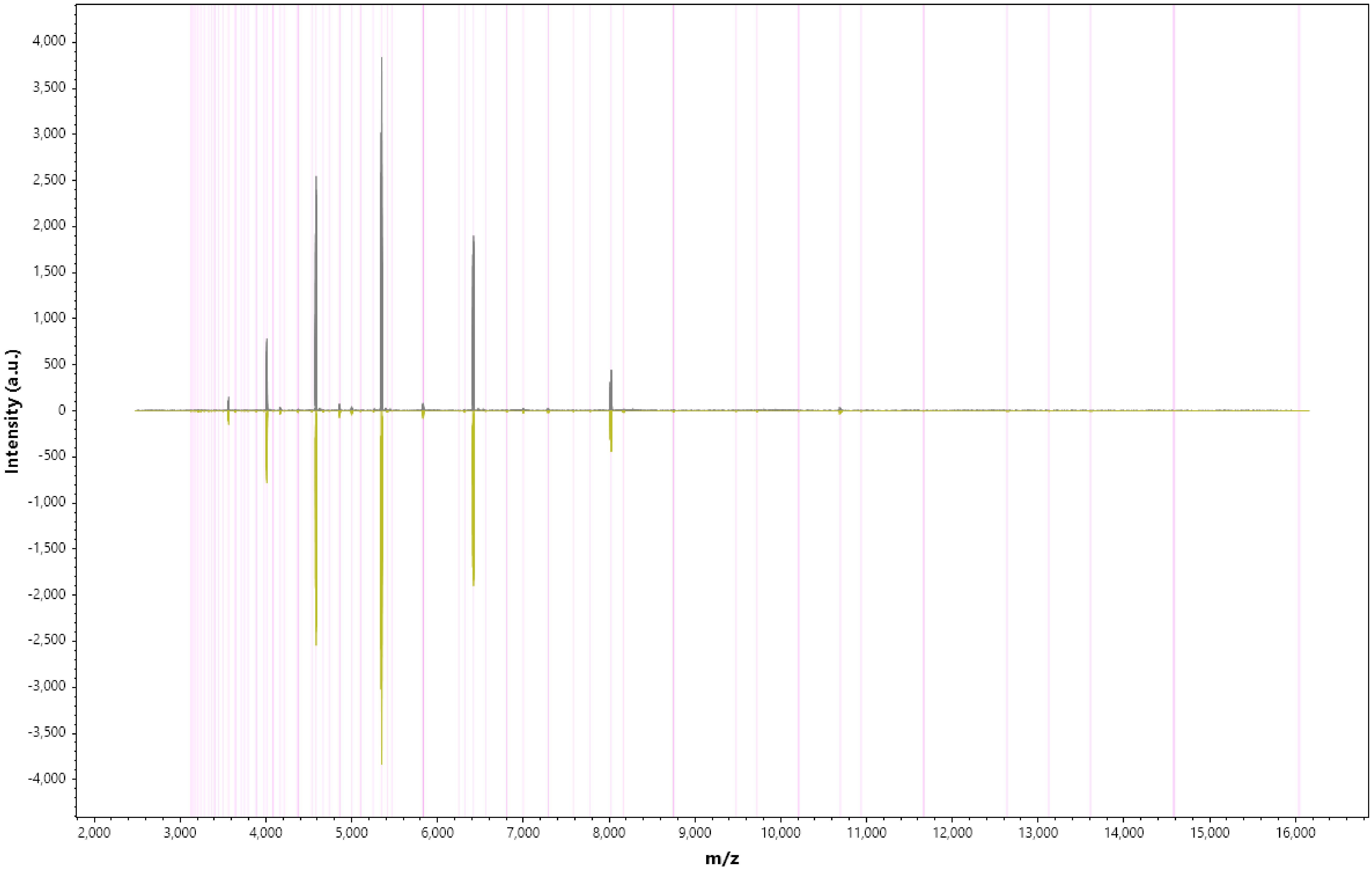

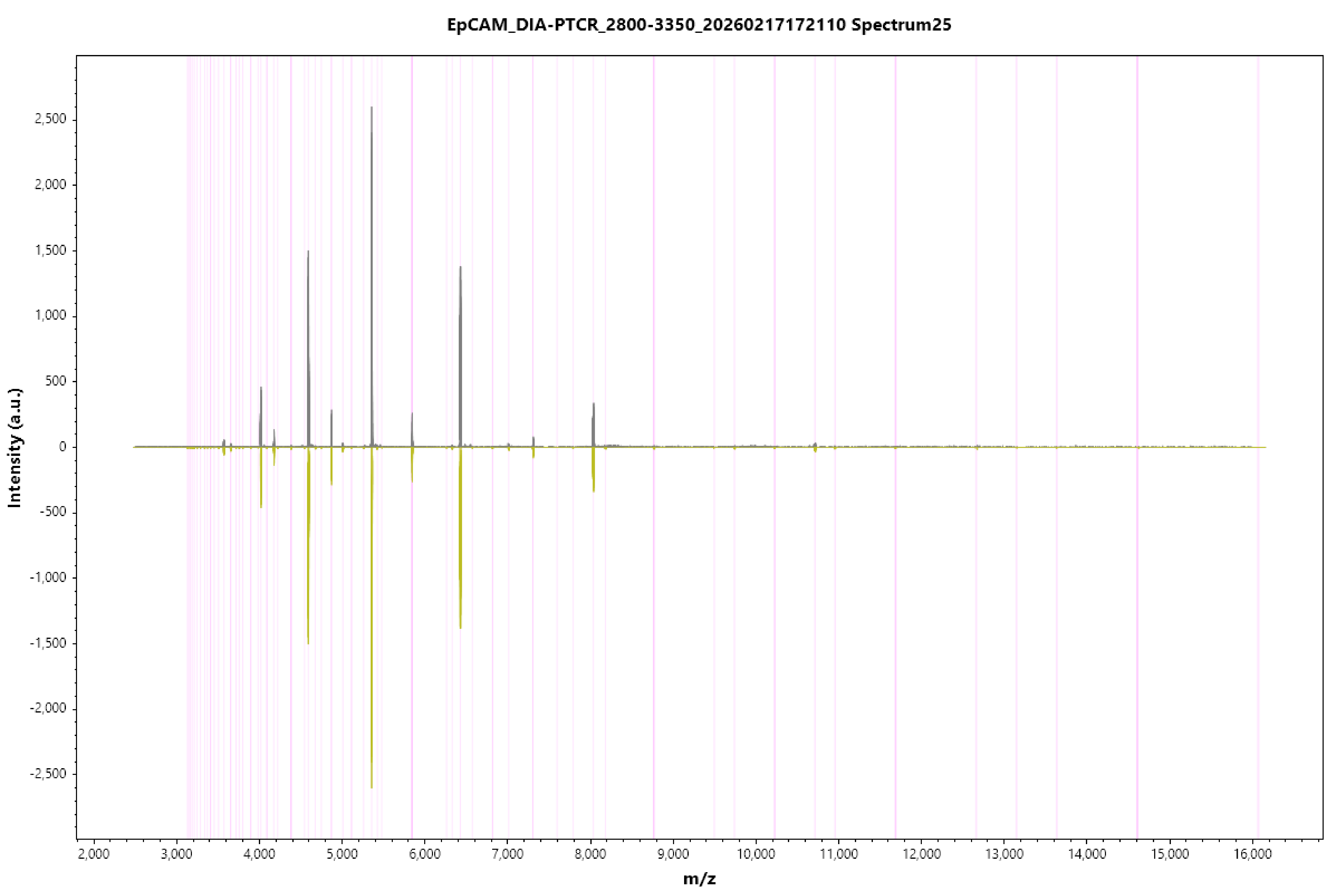

EpCAM_DIA-PTCR_2800-3350_20260217172110 Spectrum25
Intensity (a.u.)
2,500
2,000
1,500
1,000
500
0
-500
-1,000
-1,500
-2,000
-2,500
2,000
3,000
4,000
5,000
6,000
7,000
8,000
9,000
10,000
11,000
12,000
13,000
14,000
15,000
16,000
m/z

EpCAM_DIA-PTCR_2800-3350_20260217172110 Spectrum26

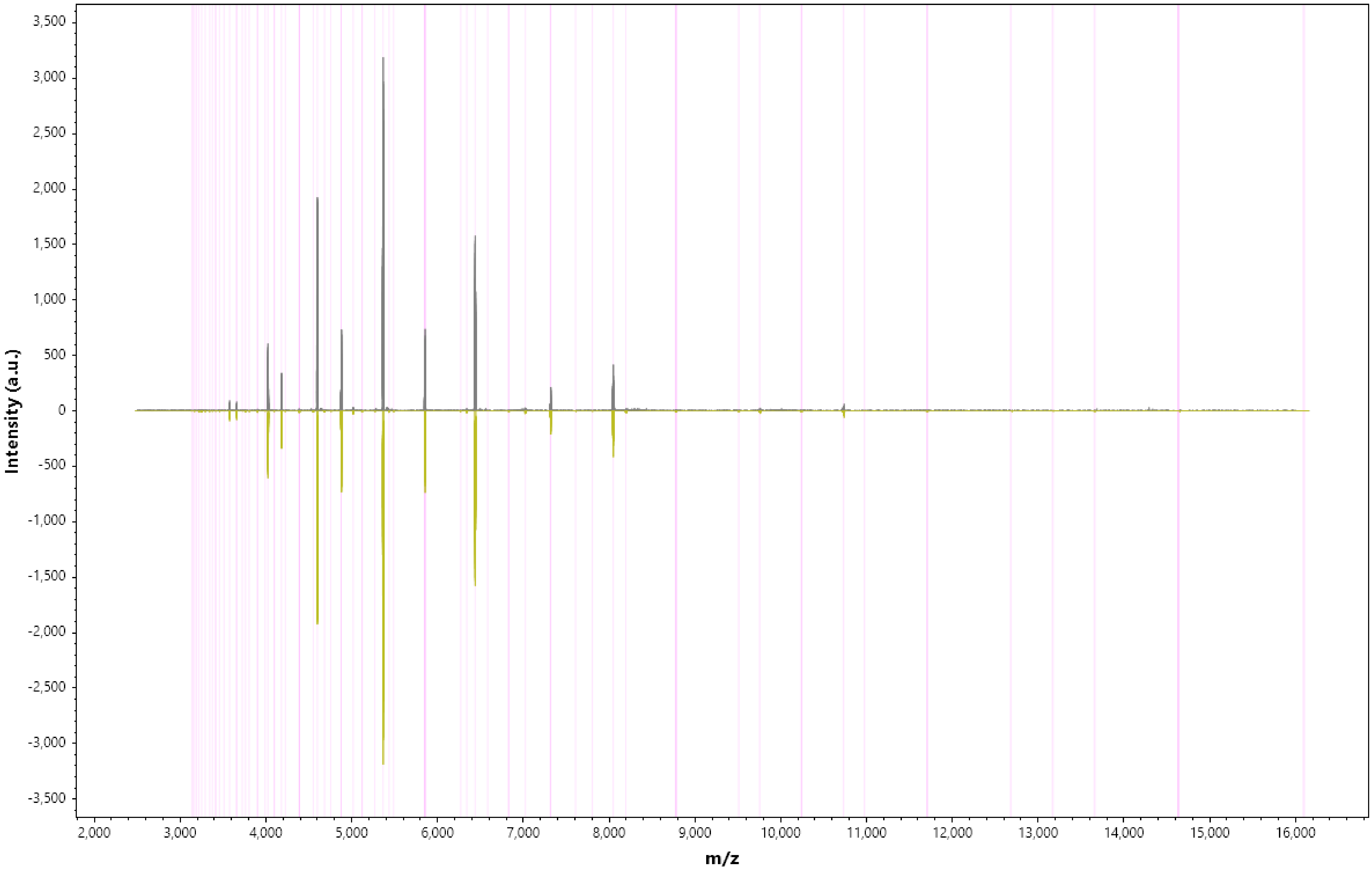

EpCAM_DIA-PTCR_2800-3350_20260217172110 Spectrum27

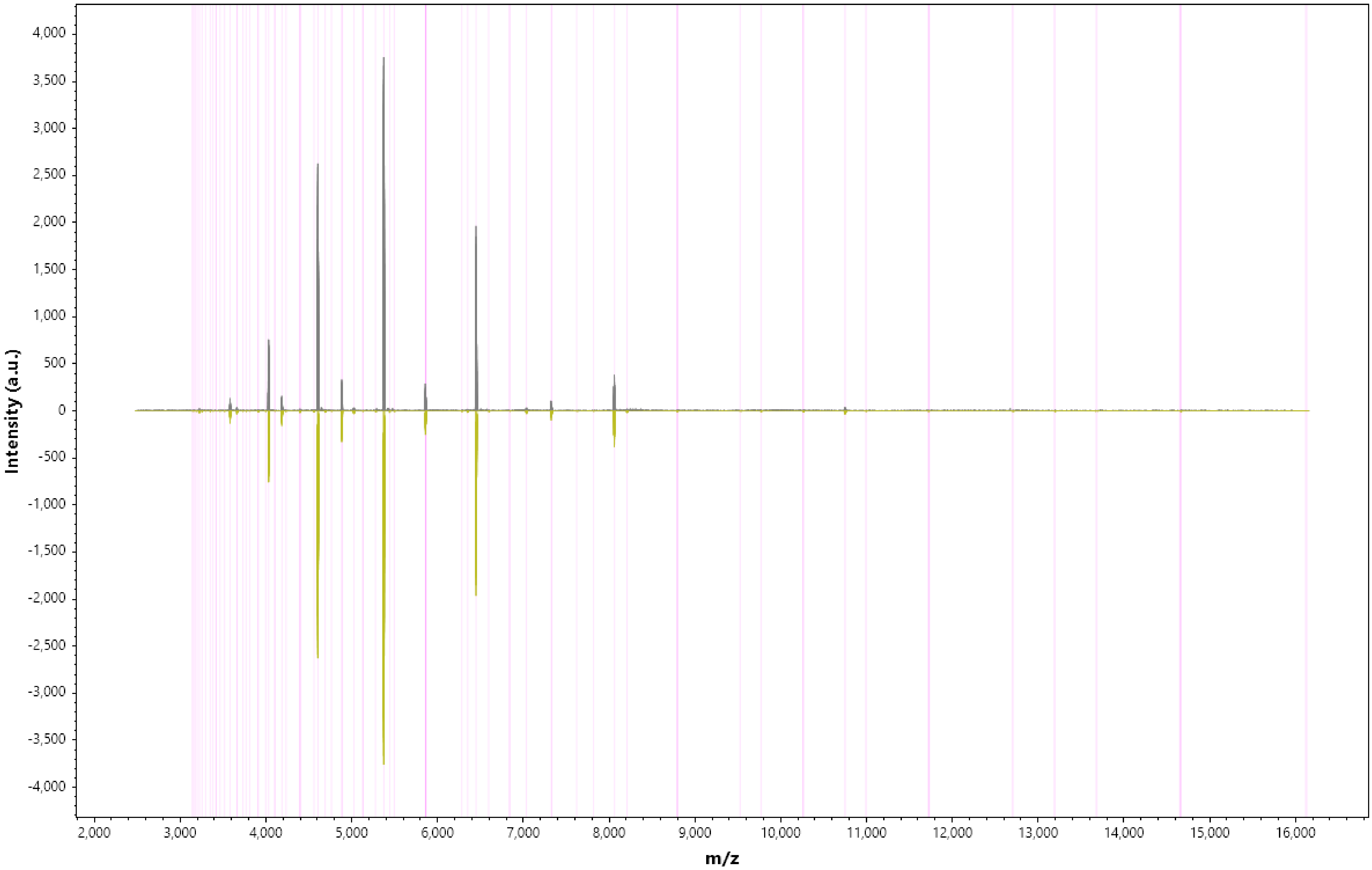

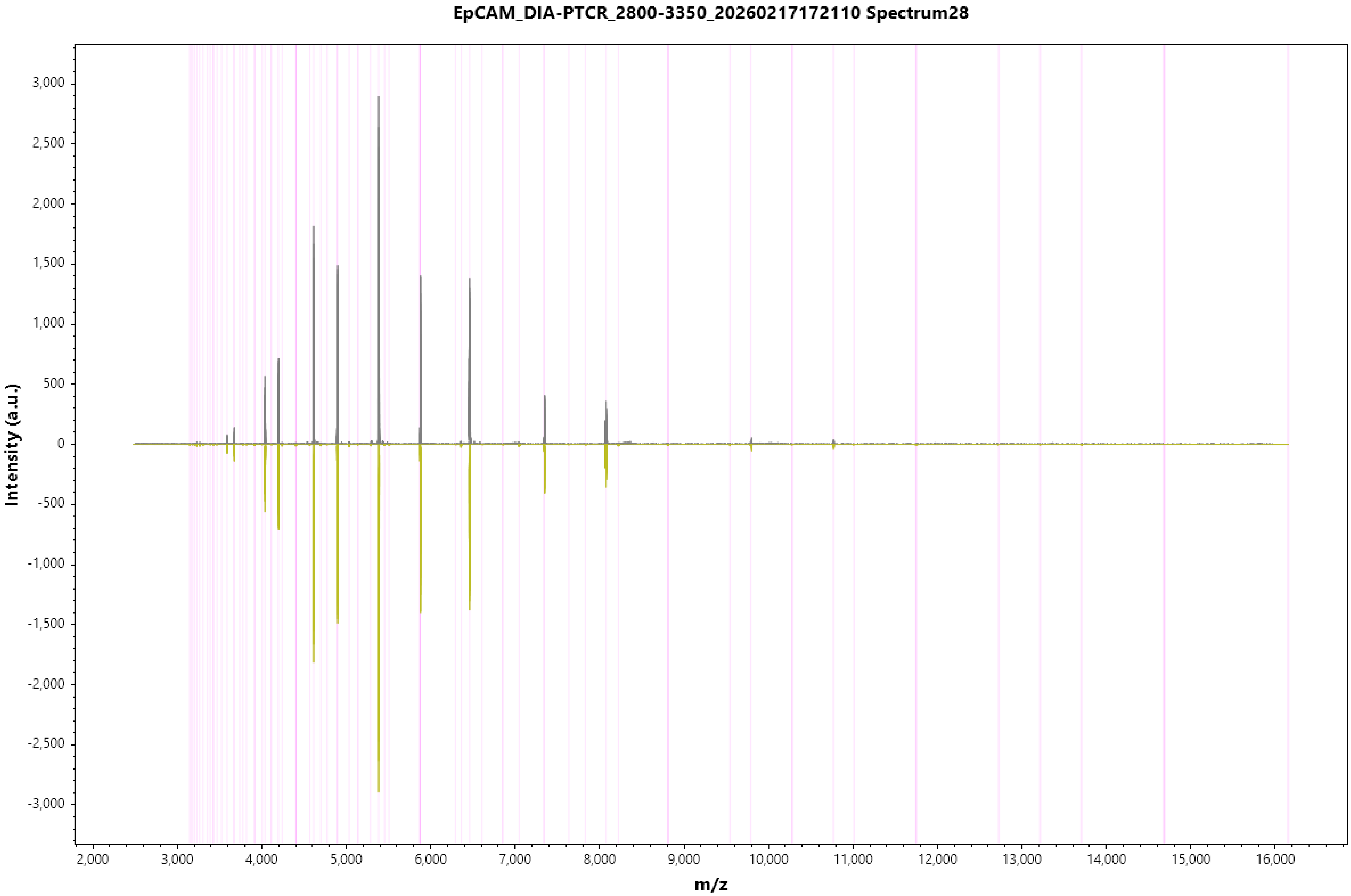
EpCAM_DIA-PTCR_2800-3350_20260217172110 Spectrum28
Intensity (a.u.)
3,000
2,500
2,000
1,500
1,000
500
0
-500
-1,000
-1,500
-2,000
-2,500
-3,000
2,000
3,000
4,000
5,000
6,000
7,000
8,000
9,000
10,000
11,000
12,000
13,000
14,000
15,000
16,000
m/z

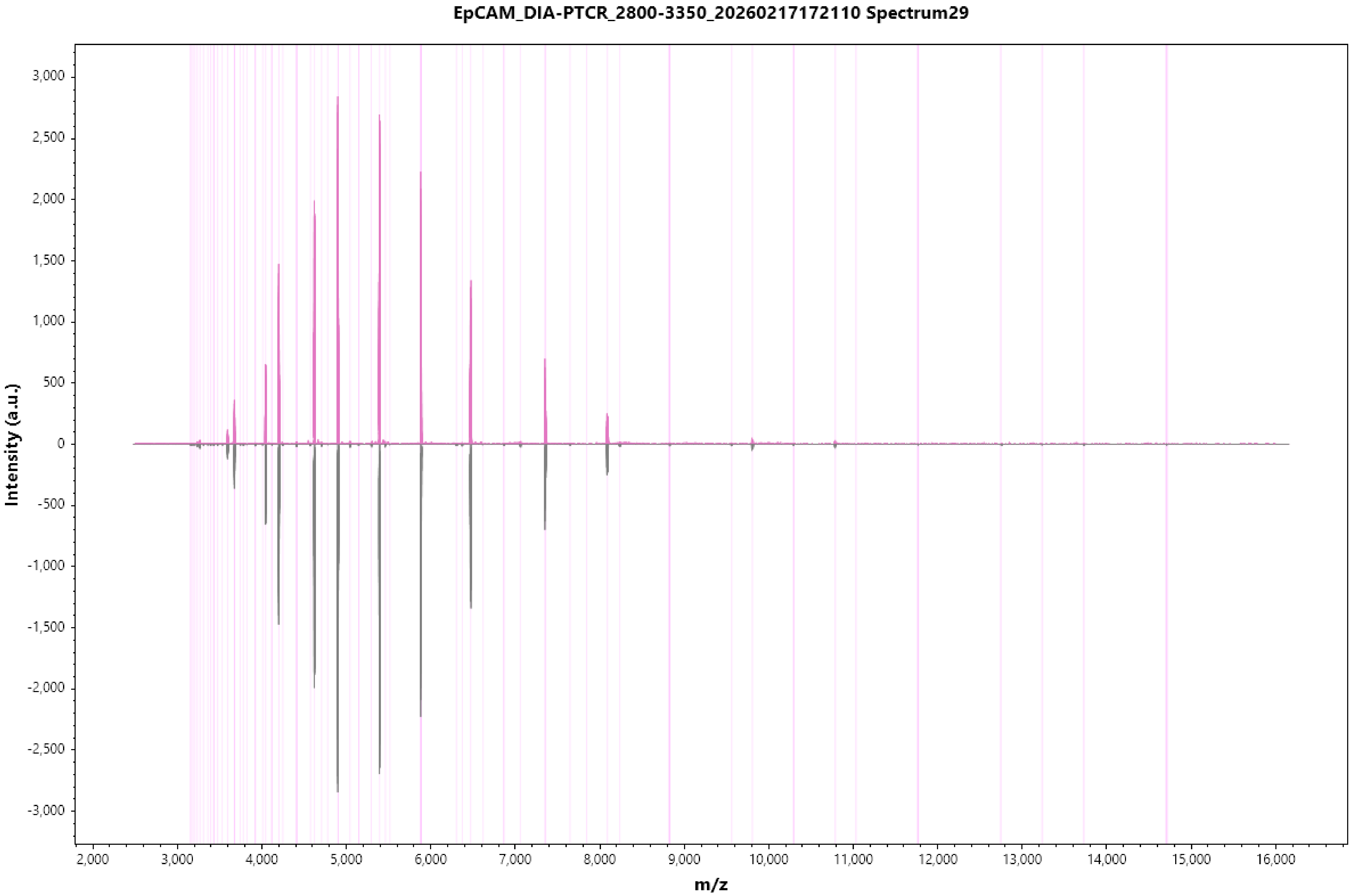

EpCAM_DIA-PTCR_2800-3350_20260217172110 Spectrum29
Intensity (a.u.)
3,000
2,500
2,000
1,500
1,000
500
0
-500
-1,000
-1,500
-2,000
-2,500
-3,000
2,000
3,000
4,000
5,000
6,000
7,000
8,000
9,000
10,000
11,000
12,000
13,000
14,000
15,000
16,000
m/z

#### EpCAM_DIA-PTCR_2800-3350_20260217172110 Spectrum30

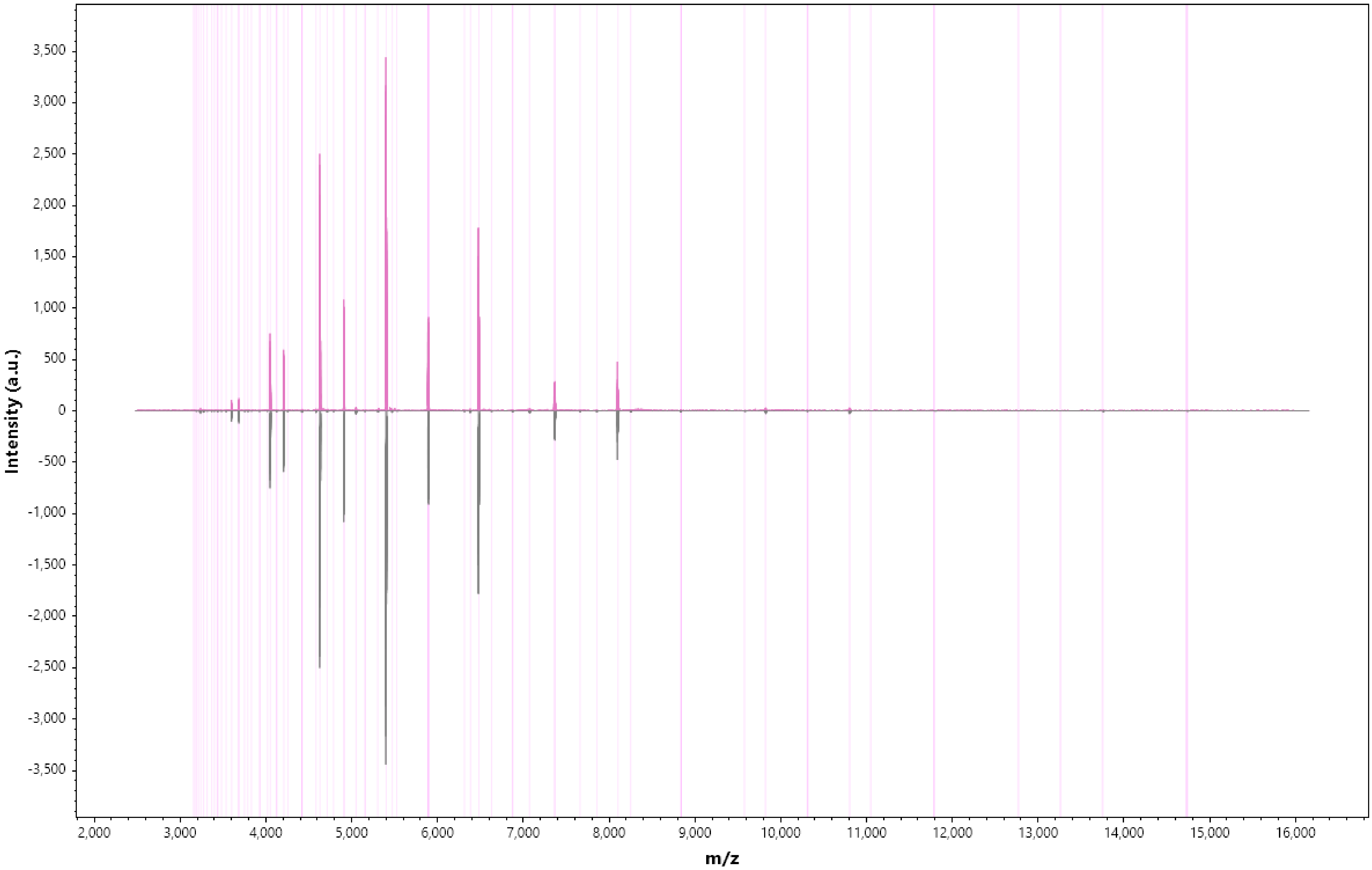

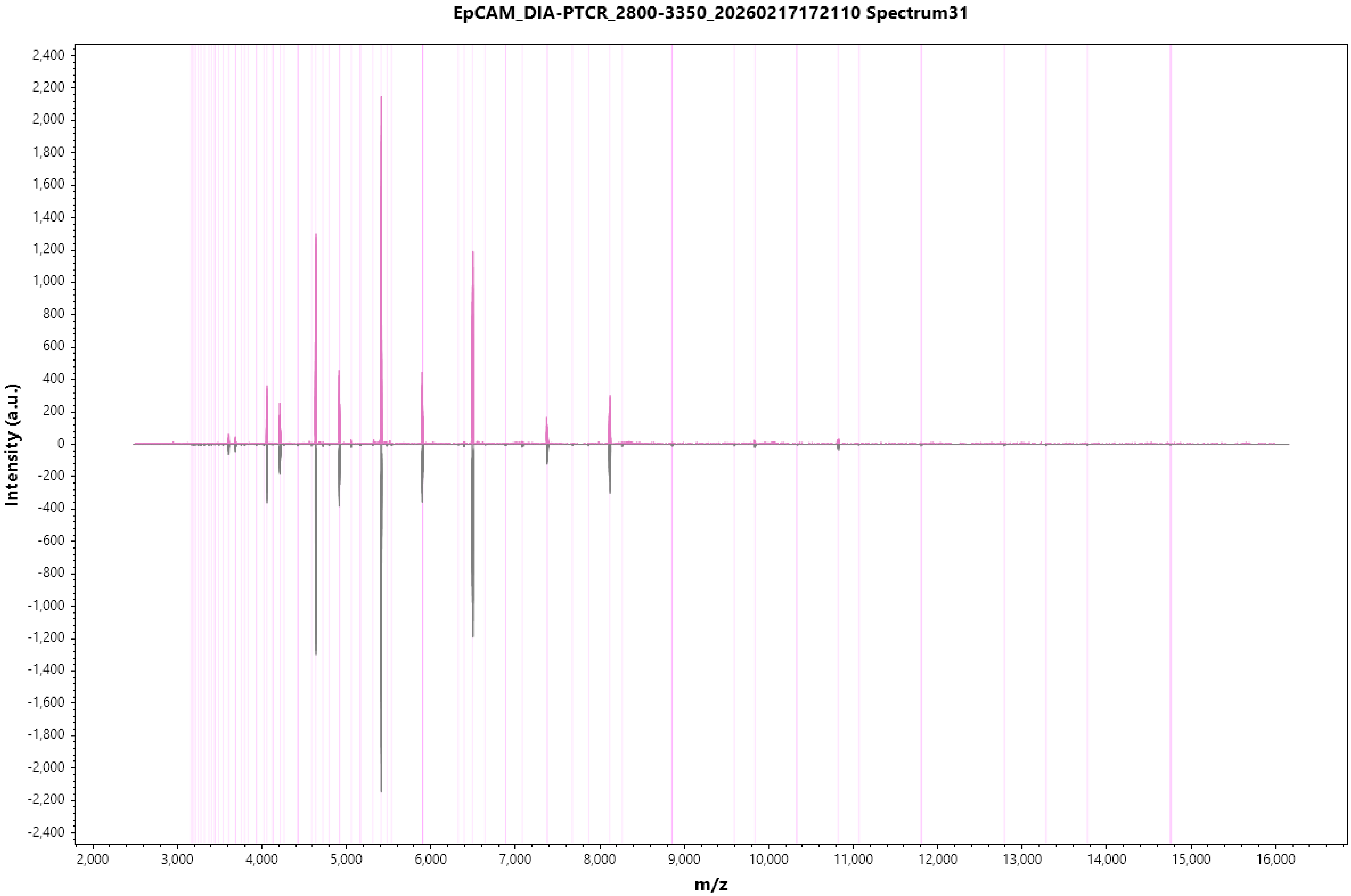
EpCAM_DIA-PTCR_2800-3350_20260217172110 Spectrum31
Intensity (a.u.)
m/z
2,400
2,200
2,000
1,800
1,600
1,400
1,200
1,000
800
600
400
200
0
-200
-400
-600
-800
-1,000
-1,200
-1,400
-1,600
-1,800
-2,000
-2,200
-2,400
2,000
3,000
4,000
5,000
6,000
7,000
8,000
9,000
10,000
11,000
12,000
13,000
14,000
15,000
16,000

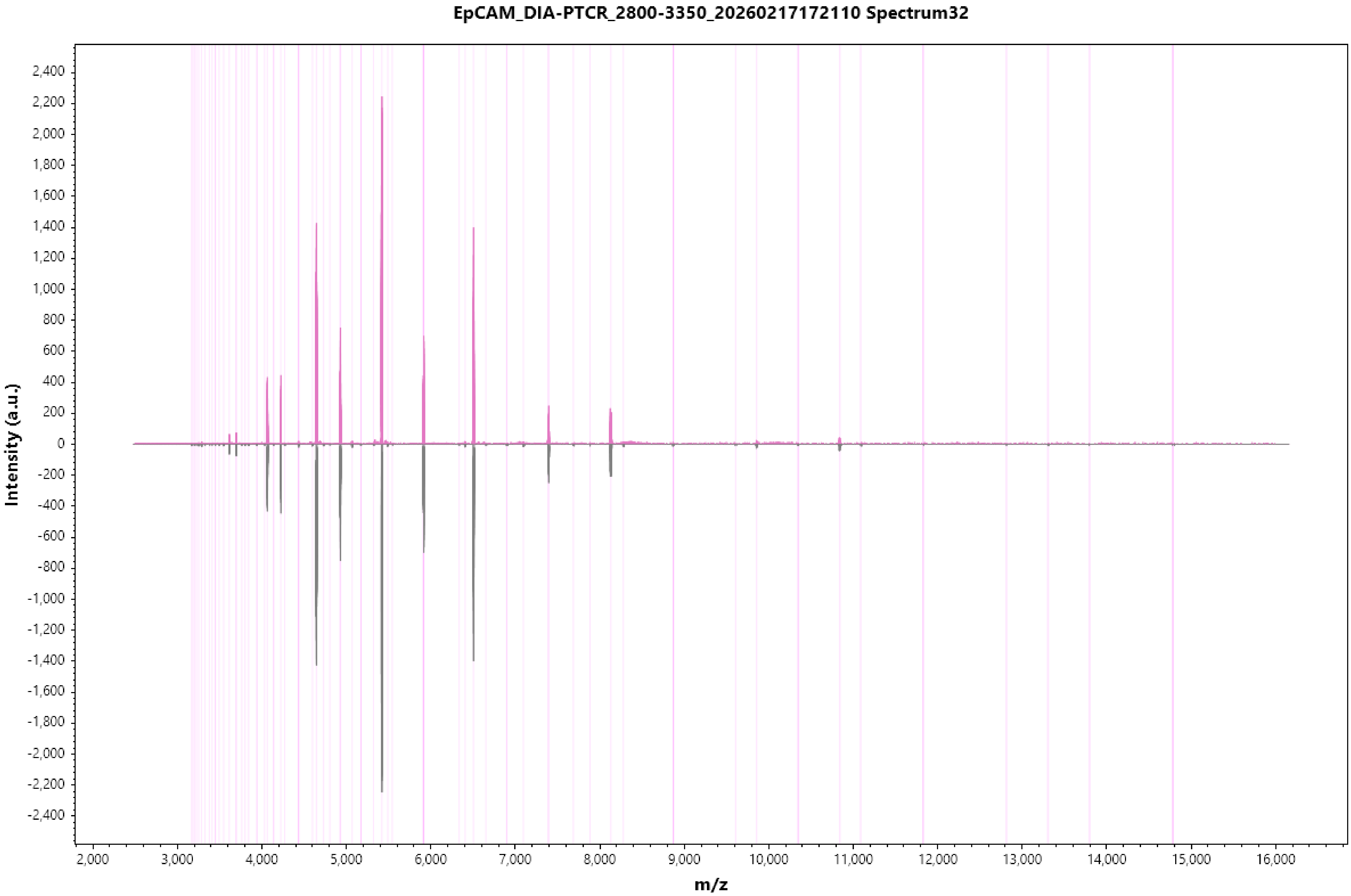
EpCAM_DIA-PTCR_2800-3350_20260217172110 Spectrum32
Intensity (a.u.)
m/z
2,400
2,200
2,000
1,800
1,600
1,400
1,200
1,000
800
600
400
200
0
-200
-400
-600
-800
-1,000
-1,200
-1,400
-1,600
-1,800
-2,000
-2,200
-2,400
2,000
3,000
4,000
5,000
6,000
7,000
8,000
9,000
10,000
11,000
12,000
13,000
14,000
15,000
16,000

EpCAM_DIA-PTCR_2800-3350_20260217172110 Spectrum33

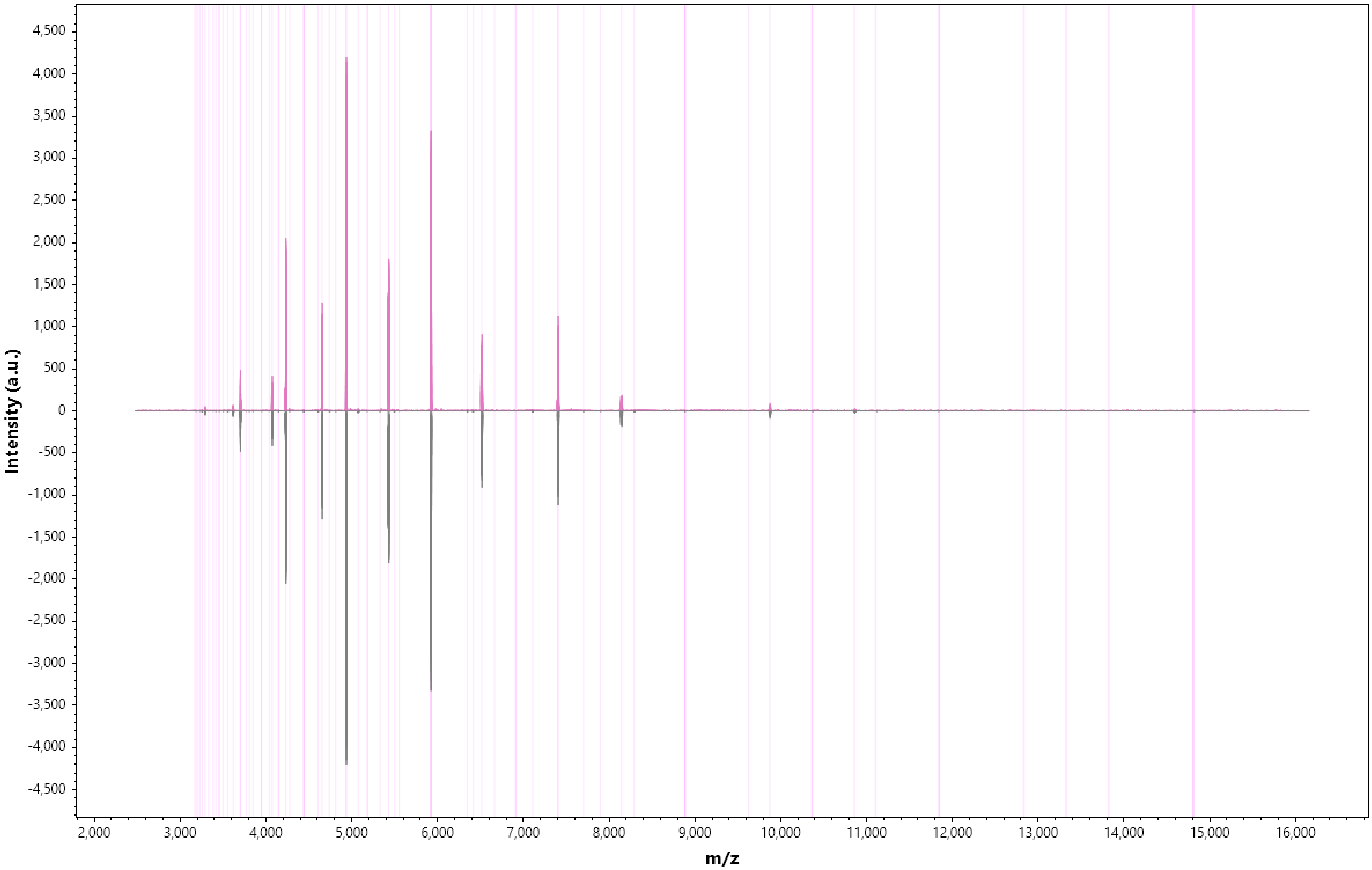

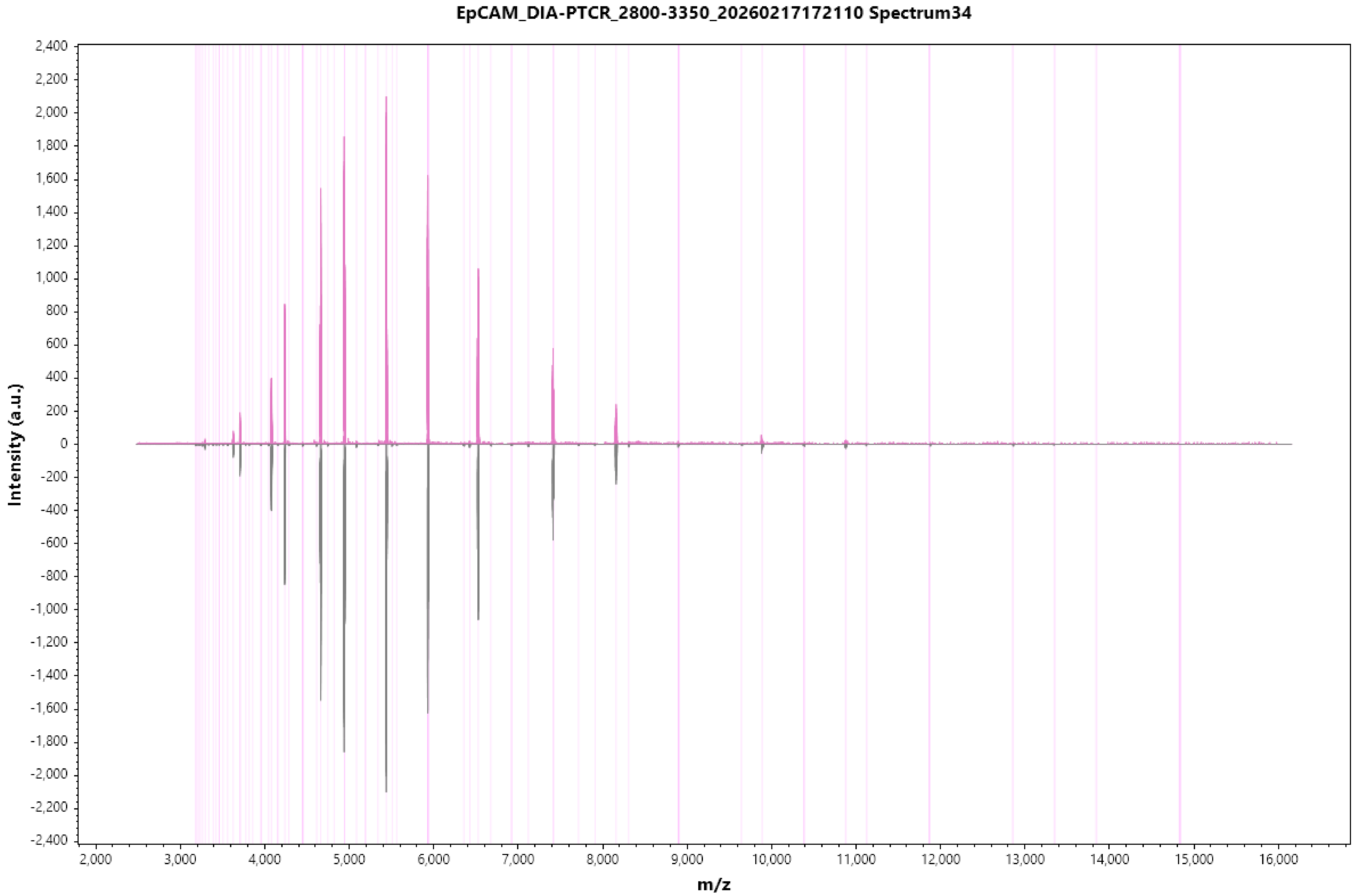
EpCAM_DIA-PTCR_2800-3350_20260217172110 Spectrum34
Intensity (a.u.)
m/z
2,400
2,200
2,000
1,800
1,600
1,400
1,200
1,000
800
600
400
200
0
-200
-400
-600
-800
-1,000
-1,200
-1,400
-1,600
-1,800
-2,000
-2,200
-2,400
2,000
3,000
4,000
5,000
6,000
7,000
8,000
9,000
10,000
11,000
12,000
13,000
14,000
15,000
16,000

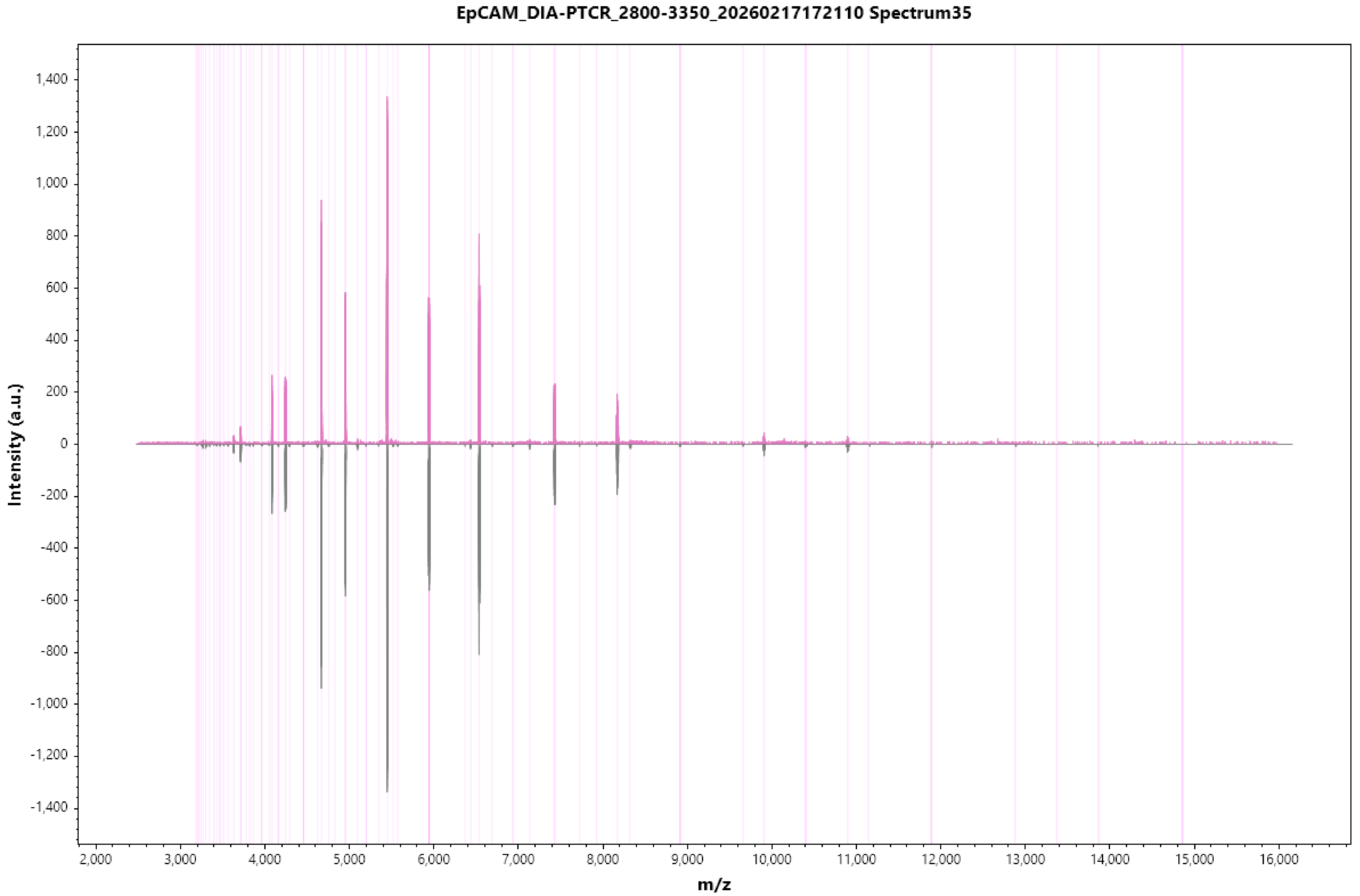

EpCAM_DIA-PTCR_2800-3350_20260217172110 Spectrum35
Intensity (a.u.)
m/z
1,400
1,200
1,000
800
600
400
200
0
-200
-400
-600
-800
-1,000
-1,200
-1,400
2,000
3,000
4,000
5,000
6,000
7,000
8,000
9,000
10,000
11,000
12,000
13,000
14,000
15,000
16,000

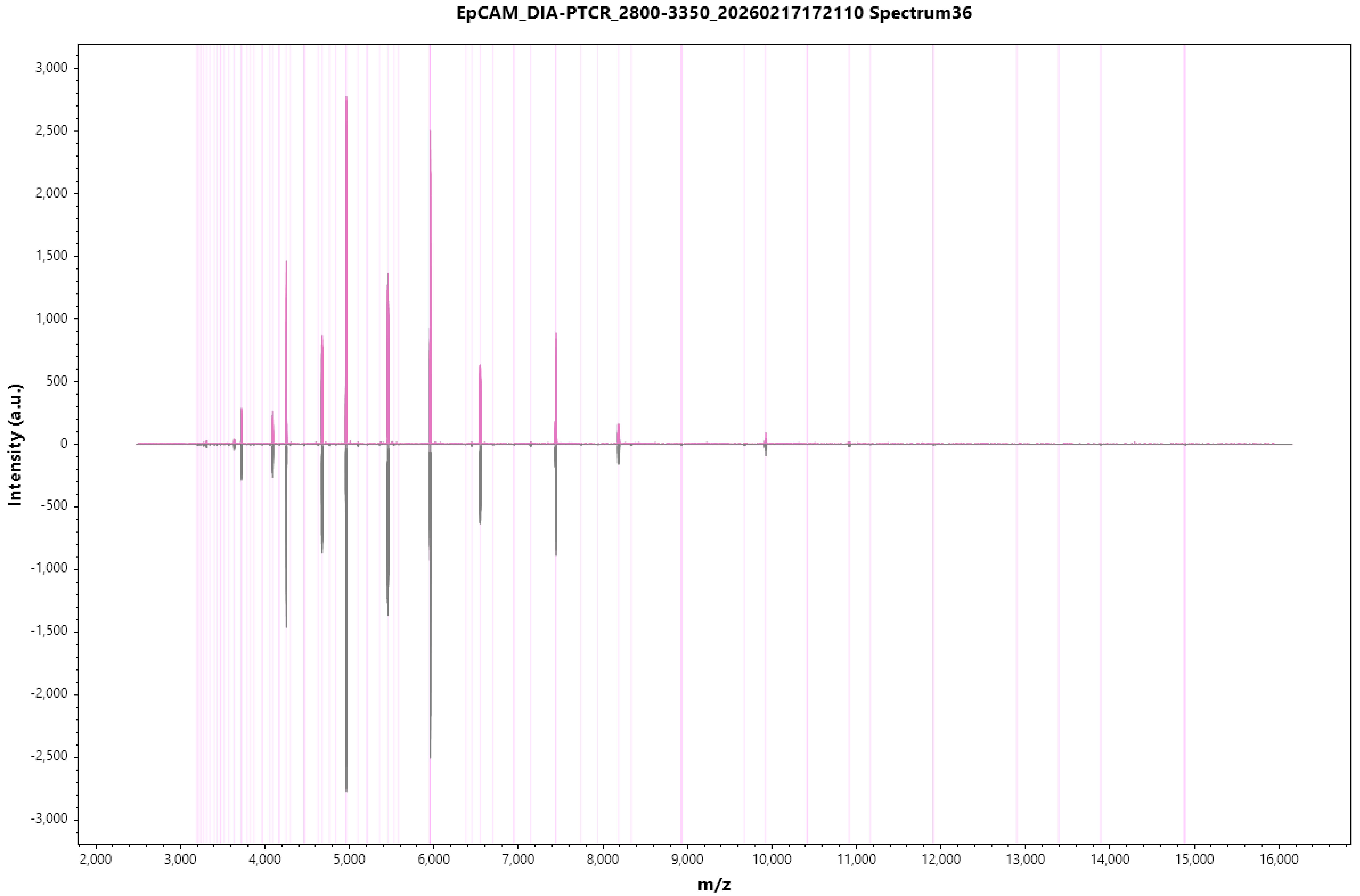
EpCAM_DIA-PTCR_2800-3350_20260217172110 Spectrum36
Intensity (a.u.)
3,000
2,500
2,000
1,500
1,000
500
0
-500
-1,000
-1,500
-2,000
-2,500
-3,000
2,000
3,000
4,000
5,000
6,000
7,000
8,000
9,000
10,000
11,000
12,000
13,000
14,000
15,000
16,000
m/z

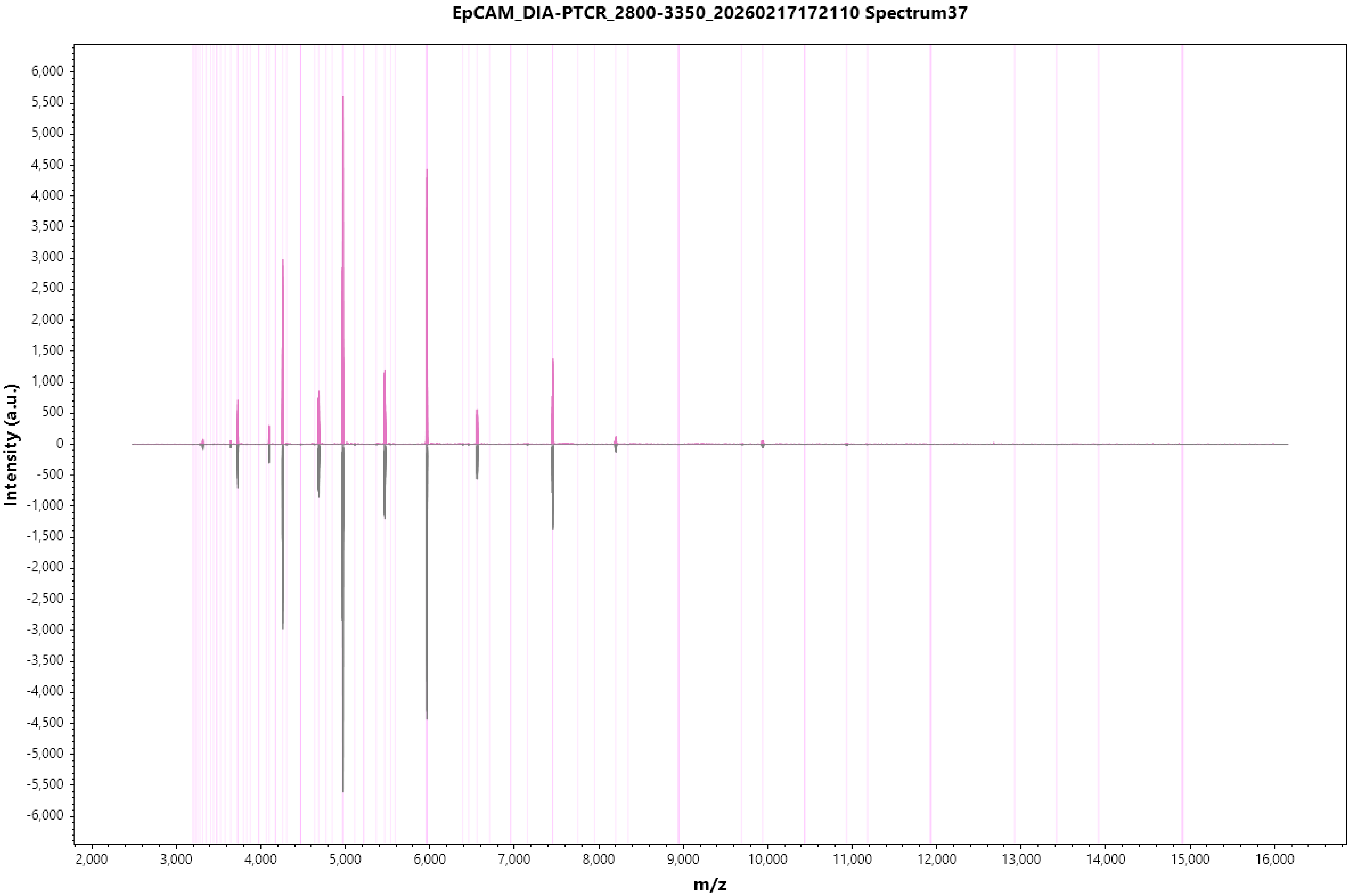
EpCAM_DIA-PTCR_2800-3350_20260217172110 Spectrum37
Intensity (a.u.)
m/z
6,000
5,500
5,000
4,500
4,000
3,500
3,000
2,500
2,000
1,500
1,000
500
0
-500
-1,000
-1,500
-2,000
-2,500
-3,000
-3,500
-4,000
-4,500
-5,000
-5,500
-6,000
2,000
3,000
4,000
5,000
6,000
7,000
8,000
9,000
10,000
11,000
12,000
13,000
14,000
15,000
16,000

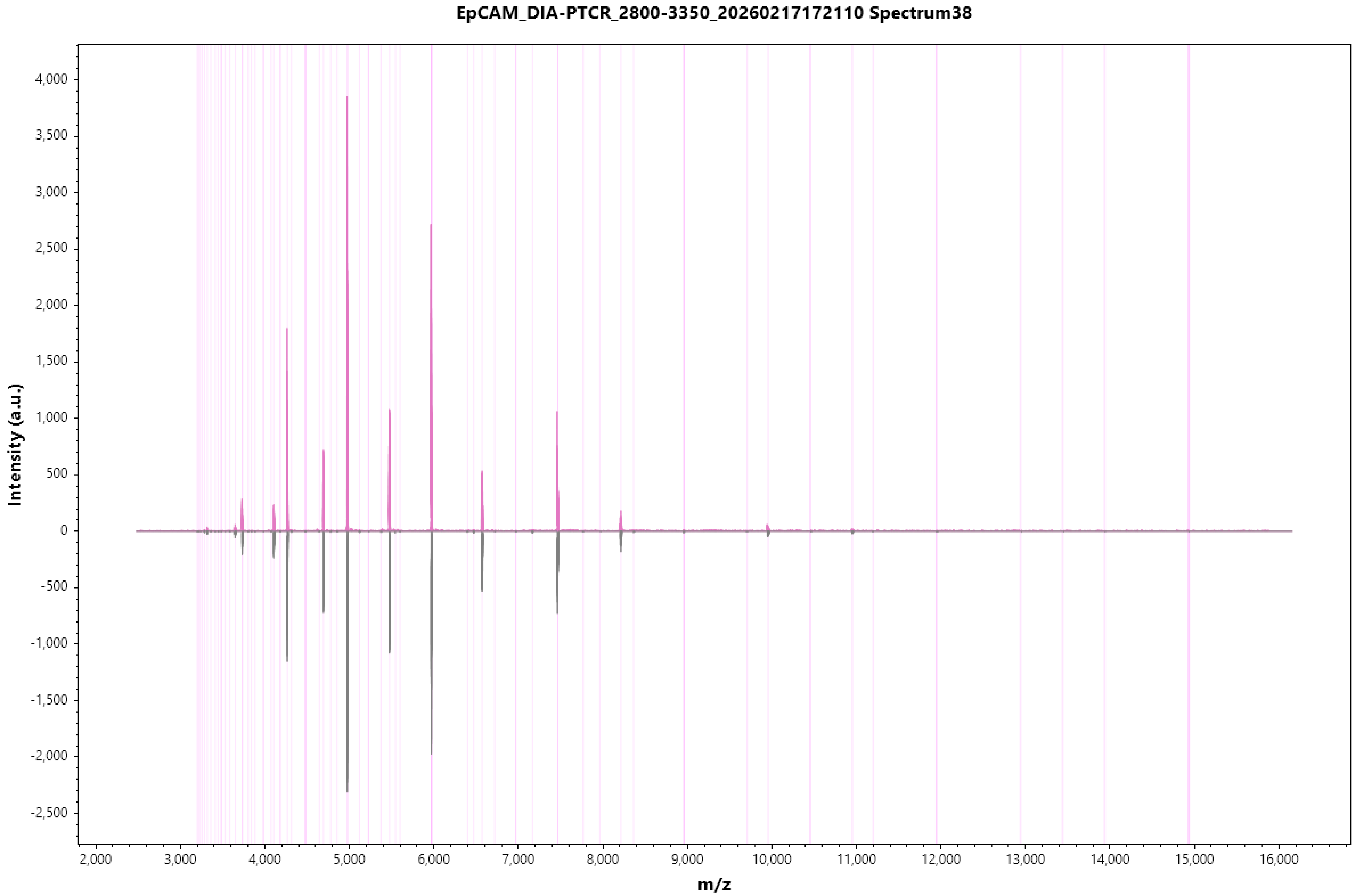

EpCAM_DIA-PTCR_2800-3350_20260217172110 Spectrum38
Intensity (a.u.)
4,000
3,500
3,000
2,500
2,000
1,500
1,000
500
0
-500
-1,000
-1,500
-2,000
-2,500
2,000
3,000
4,000
5,000
6,000
7,000
8,000
9,000
10,000
11,000
12,000
13,000
14,000
15,000
16,000
m/z

EpCAM_DIA-PTCR_2800-3350_20260217172110 Spectrum39

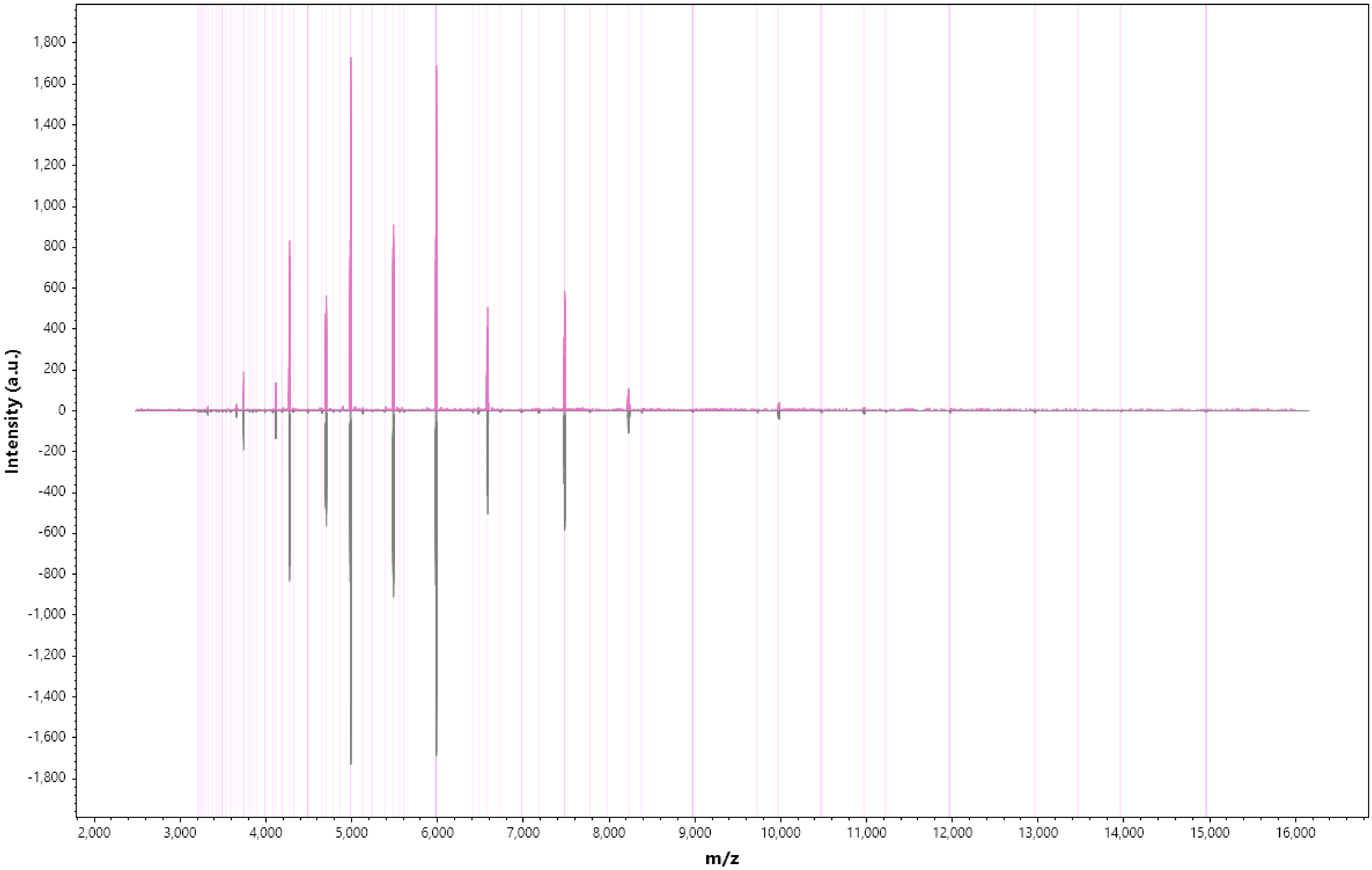

#### EpCAM_DIA-PTCR_2800-3350_20260217172110 Spectrum40

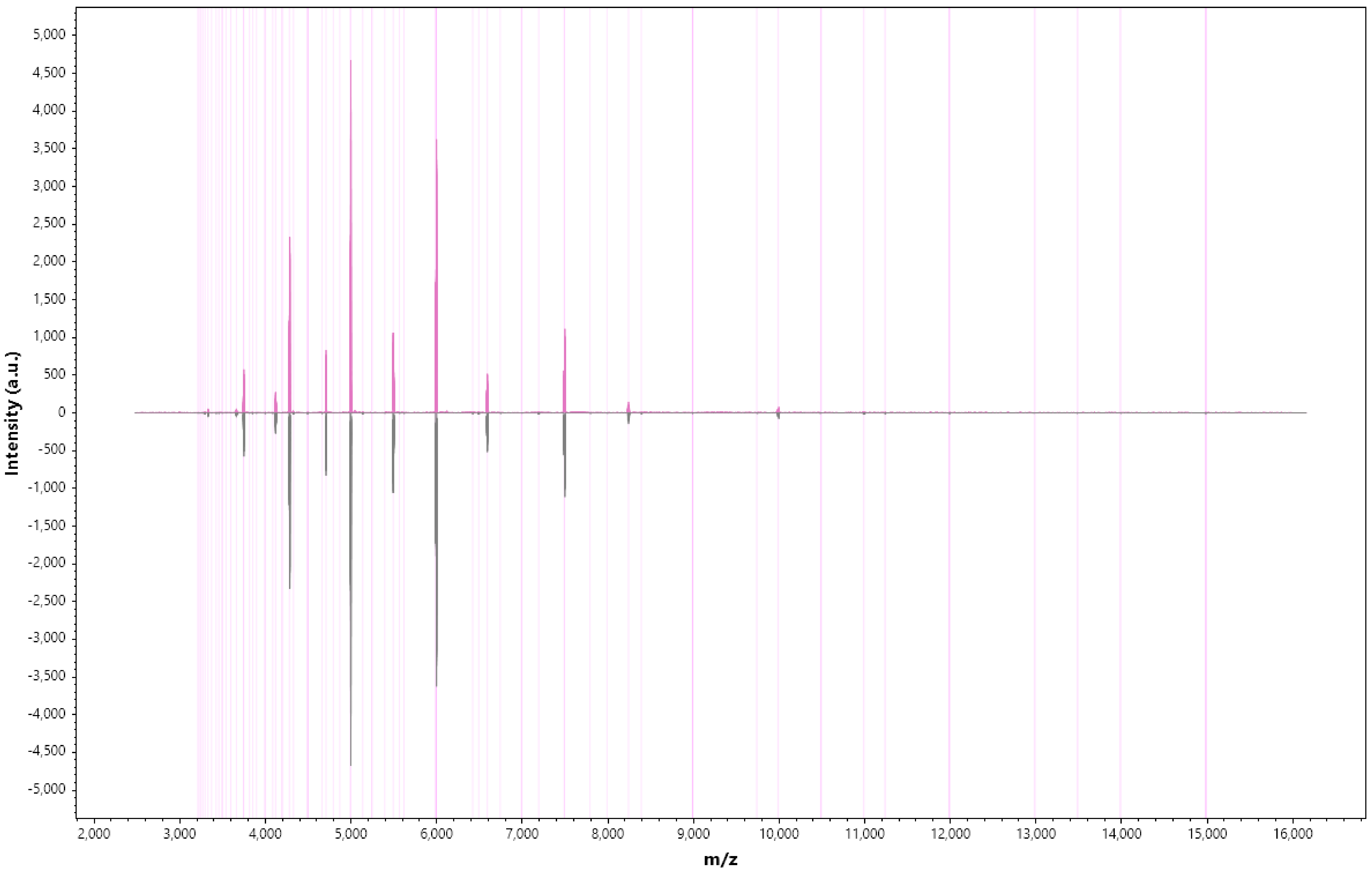

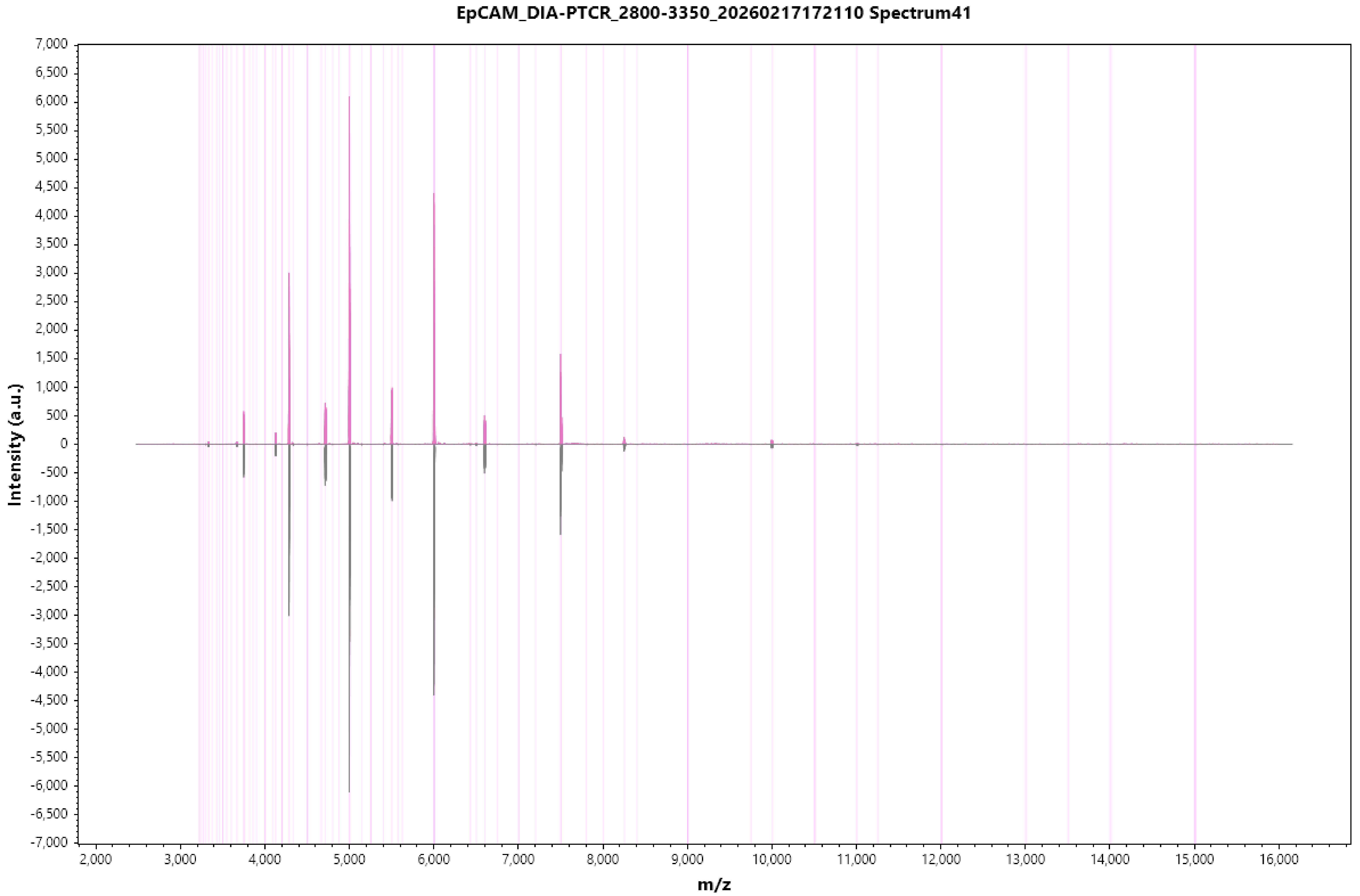

EpCAM_DIA-PTCR_2800-3350_20260217172110 Spectrum41
Intensity (a.u.)
m/z

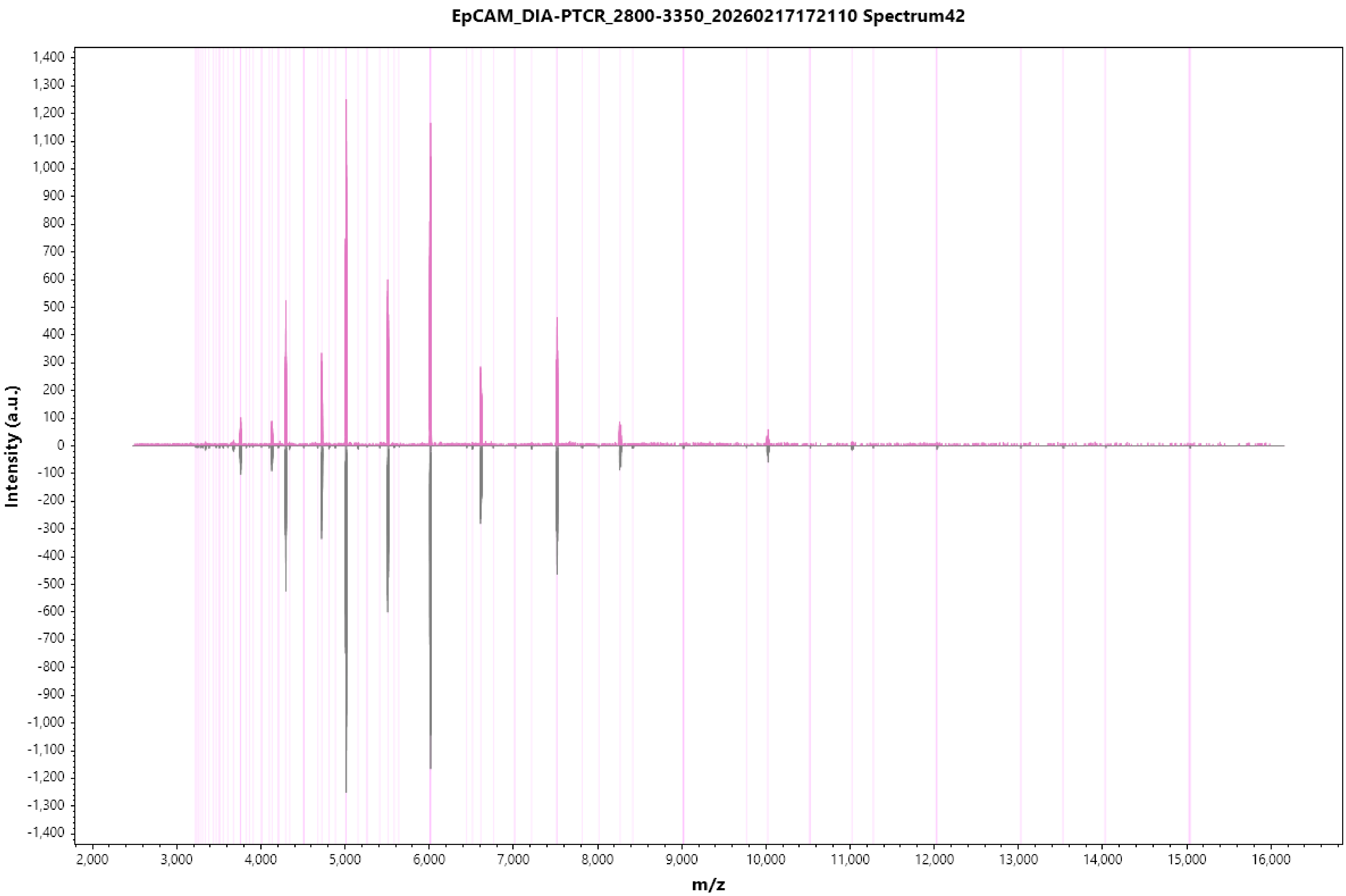
EpCAM_DIA-PTCR_2800-3350_20260217172110 Spectrum42
Intensity (a.u.)
m/z
1,400
1,300
1,200
1,100
1,000
900
800
700
600
500
400
300
200
100
0
-100
-200
-300
-400
-500
-600
-700
-800
-900
-1,000
-1,100
-1,200
-1,300
-1,400
2,000
3,000
4,000
5,000
6,000
7,000
8,000
9,000
10,000
11,000
12,000
13,000
14,000
15,000
16,000

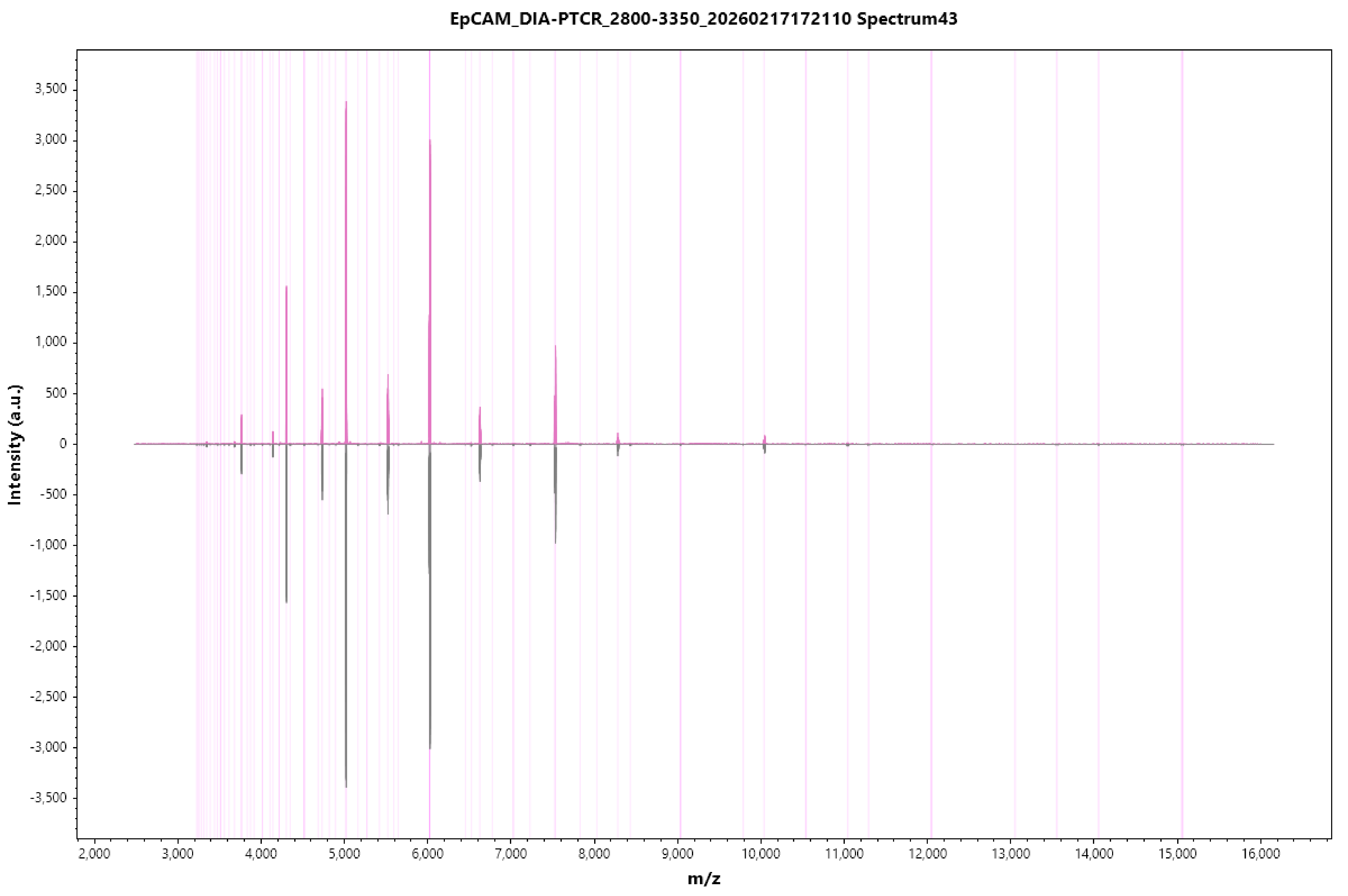
EpCAM_DIA-PTCR_2800-3350_20260217172110 Spectrum43
Intensity (a.u.)
m/z
3,500
3,000
2,500
2,000
1,500
1,000
500
0
-500
-1,000
-1,500
-2,000
-2,500
-3,000
-3,500
2,000
3,000
4,000
5,000
6,000
7,000
8,000
9,000
10,000
11,000
12,000
13,000
14,000
15,000
16,000

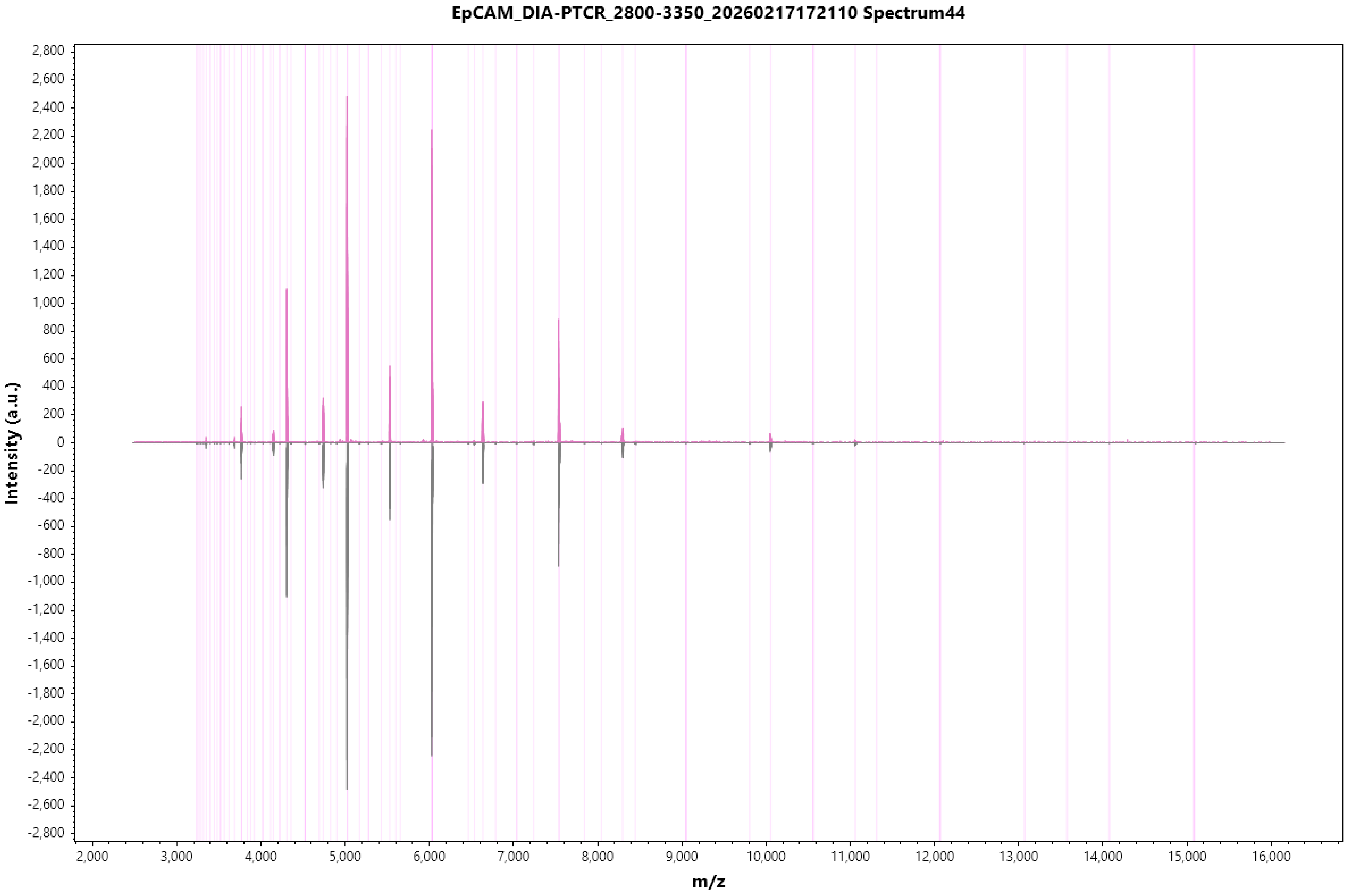
EpCAM_DIA-PTCR_2800-3350_20260217172110 Spectrum44
Intensity (a.u.)
m/z
2,800
2,600
2,400
2,200
2,000
1,800
1,600
1,400
1,200
1,000
800
600
400
200
0
-200
-400
-600
-800
-1,000
-1,200
-1,400
-1,600
-1,800
-2,000
-2,200
-2,400
-2,600
-2,800
2,000
3,000
4,000
5,000
6,000
7,000
8,000
9,000
10,000
11,000
12,000
13,000
14,000
15,000
16,000

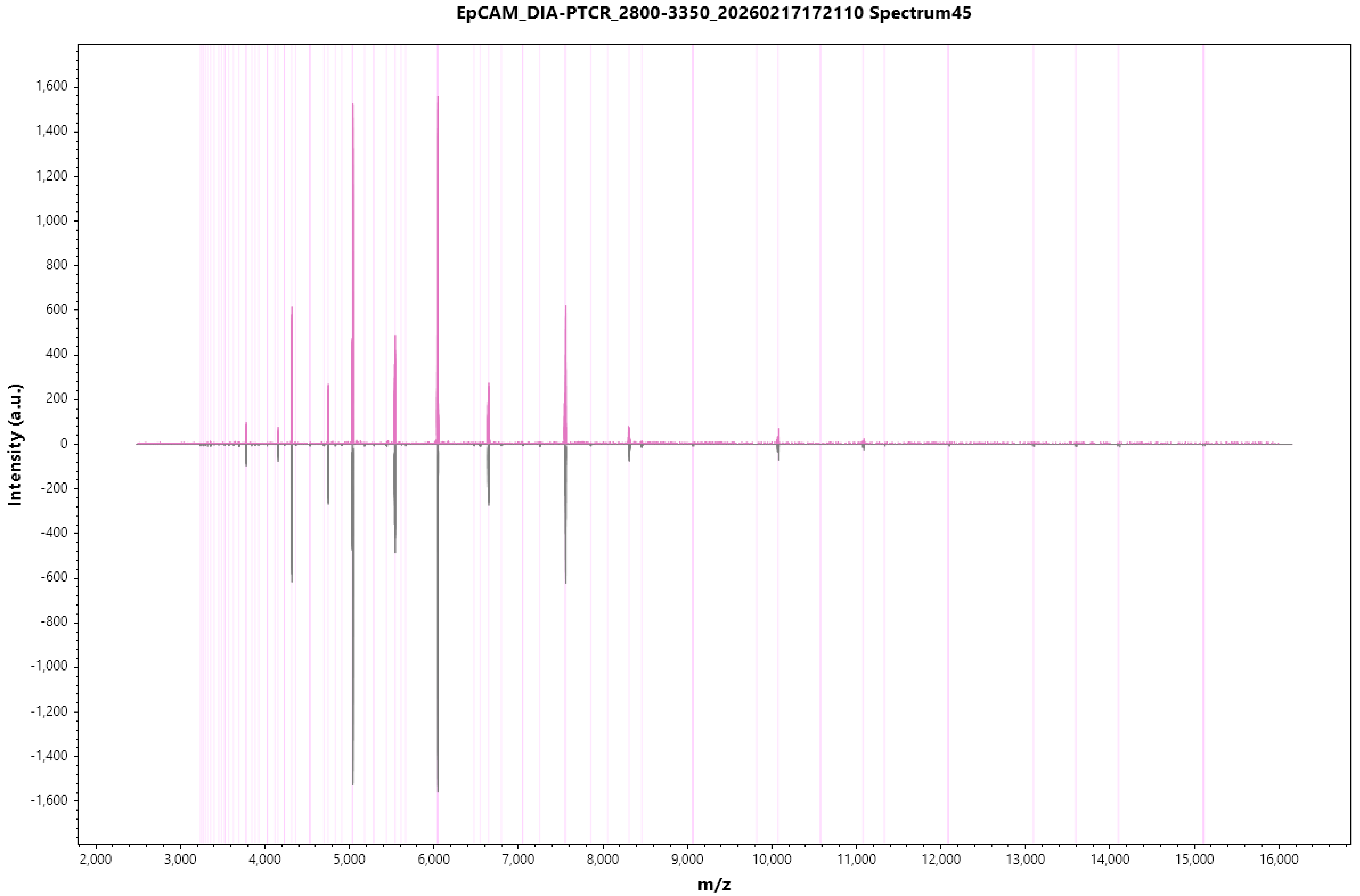
EpCAM_DIA-PTCR_2800-3350_20260217172110 Spectrum45
Intensity (a.u.)
m/z
1,600
1,400
1,200
1,000
800
600
400
200
0
-200
-400
-600
-800
-1,000
-1,200
-1,400
-1,600
2,000
3,000
4,000
5,000
6,000
7,000
8,000
9,000
10,000
11,000
12,000
13,000
14,000
15,000
16,000

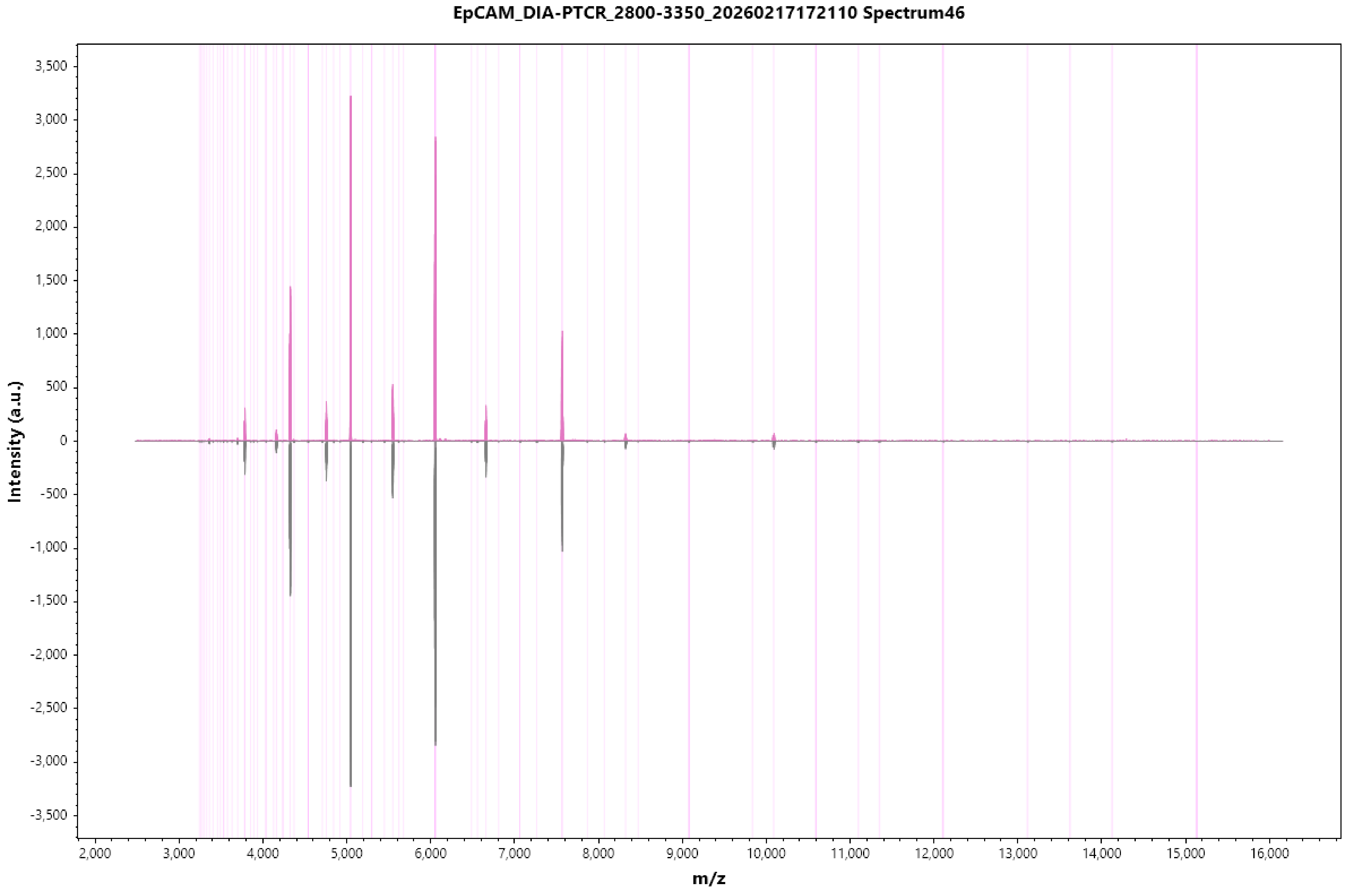
EpCAM_DIA-PTCR_2800-3350_20260217172110 Spectrum46
Intensity (a.u.)
3,500
3,000
2,500
2,000
1,500
1,000
500
0
-500
-1,000
-1,500
-2,000
-2,500
-3,000
-3,500
2,000
3,000
4,000
5,000
6,000
7,000
8,000
9,000
10,000
11,000
12,000
13,000
14,000
15,000
16,000
m/z

EpCAM_DIA-PTCR_2800-3350_20260217172110 Spectrum47

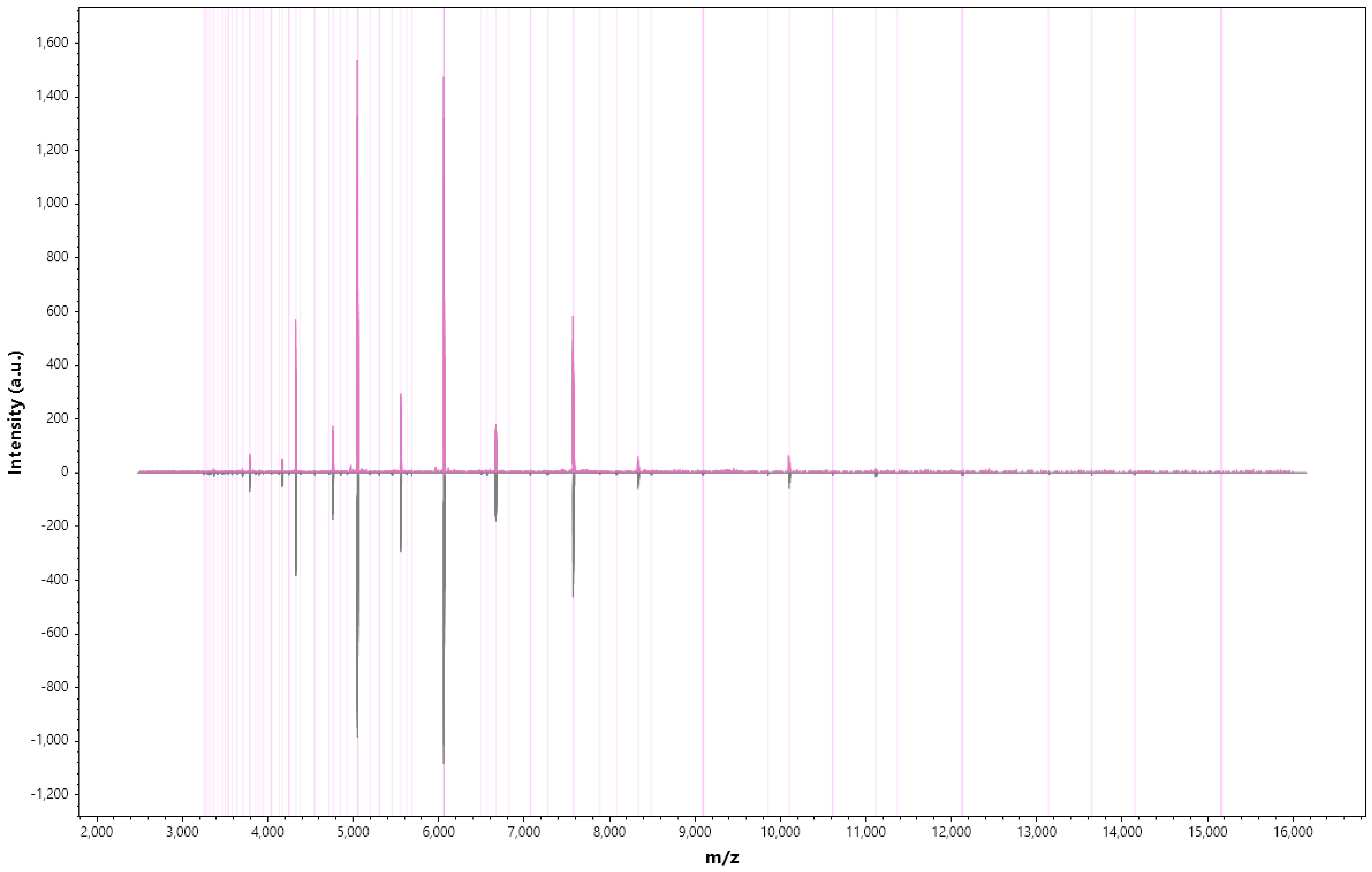

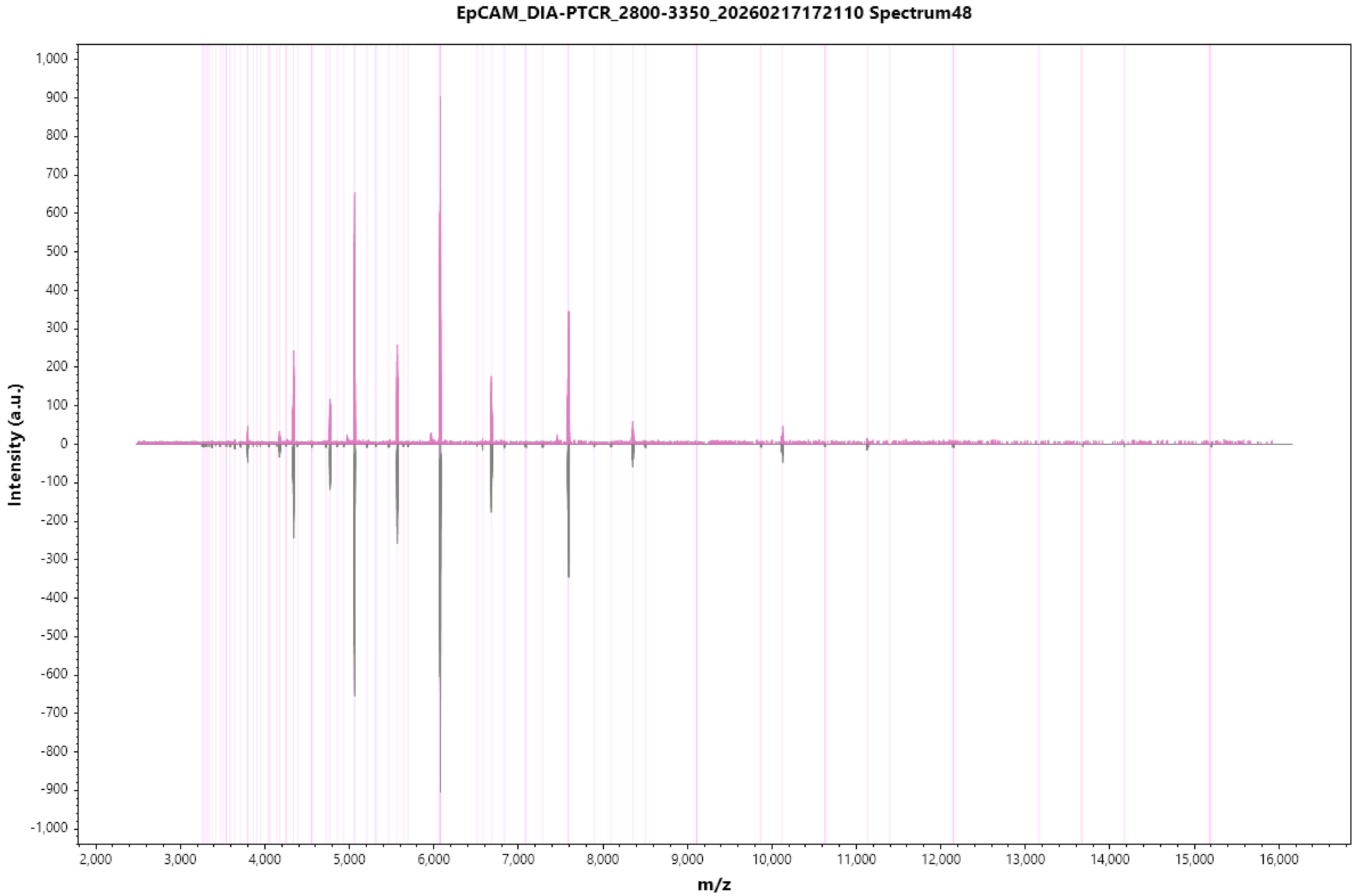

EpCAM_DIA-PTCR_2800-3350_20260217172110 Spectrum48
Intensity (a.u.)
1,000
900
800
700
600
500
400
300
200
100
0
-100
-200
-300
-400
-500
-600
-700
-800
-900
-1,000
2,000
3,000
4,000
5,000
6,000
7,000
8,000
9,000
10,000
11,000
12,000
13,000
14,000
15,000
16,000
m/z

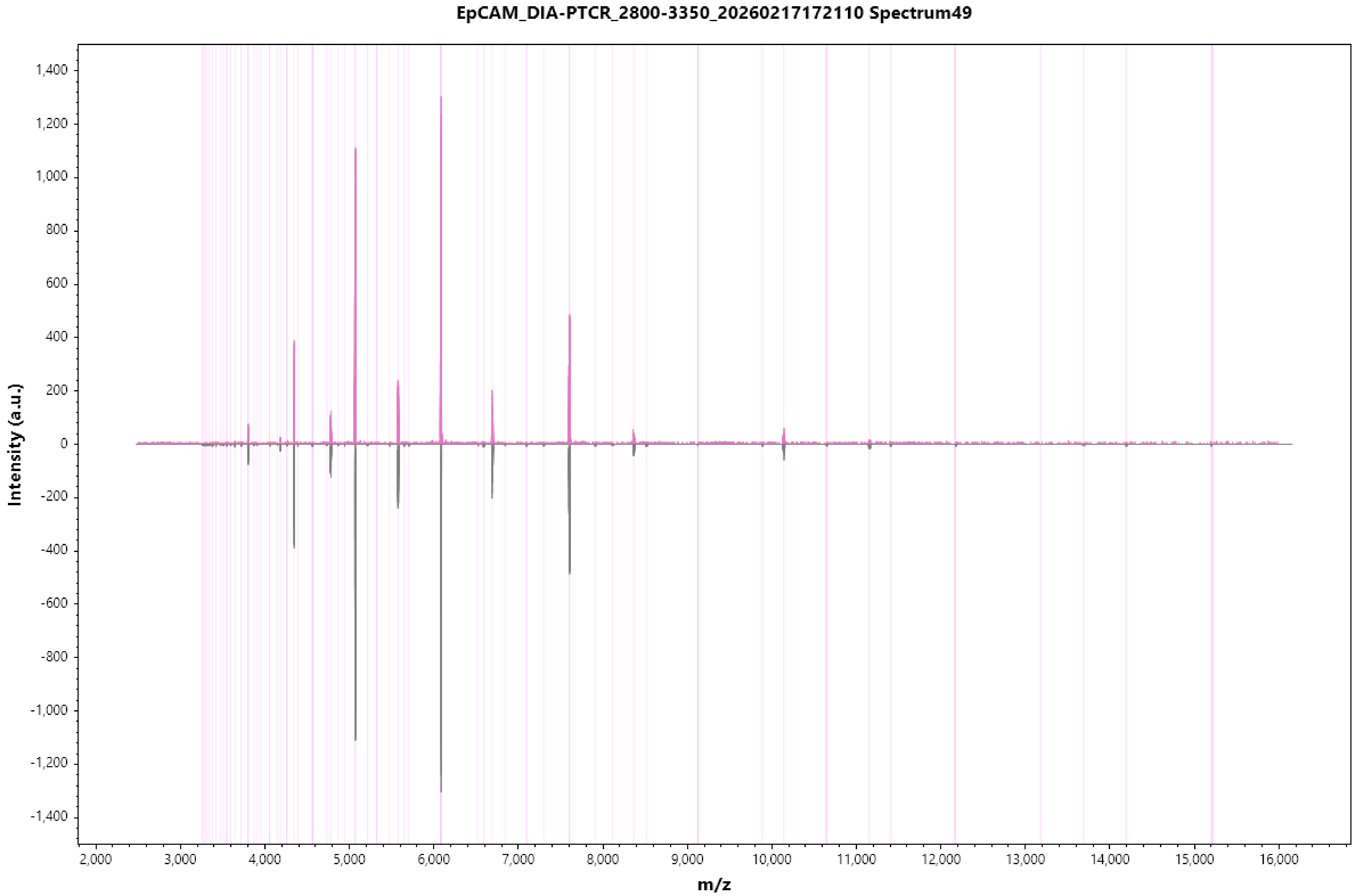
EpCAM_DIA-PTCR_2800-3350_20260217172110 Spectrum49
Intensity (a.u.)
1,400
1,200
1,000
800
600
400
200
0
-200
-400
-600
-800
-1,000
-1,200
-1,400
2,000
3,000
4,000
5,000
6,000
7,000
8,000
9,000
10,000
11,000
12,000
13,000
14,000
15,000
16,000
m/z

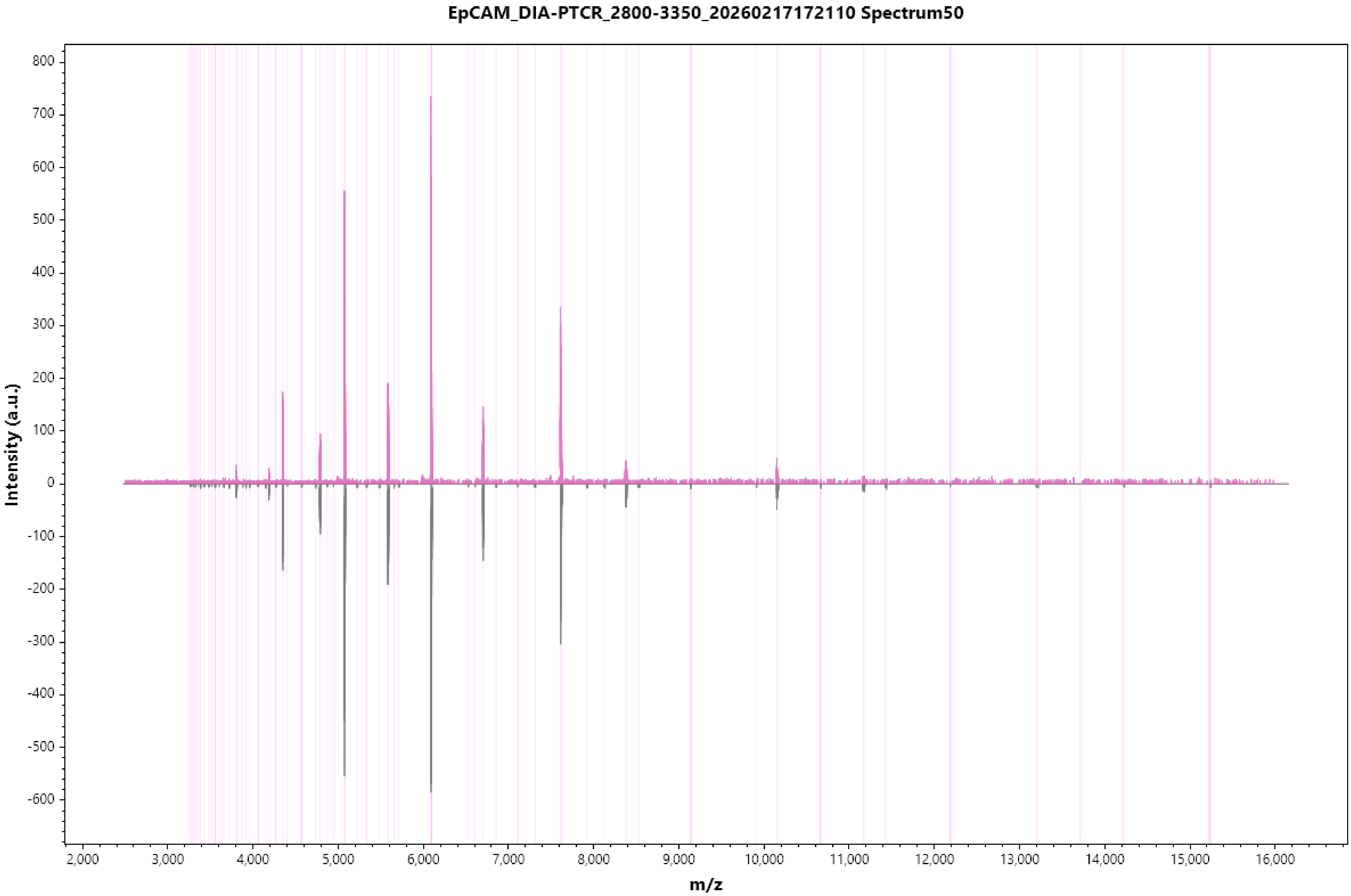
EpCAM_DIA-PTCR_2800-3350_20260217172110 Spectrum50
Intensity (a.u.)
m/z
800
700
600
500
400
300
200
100
0
-100
-200
-300
-400
-500
-600
2,000
3,000
4,000
5,000
6,000
7,000
8,000
9,000
10,000
11,000
12,000
13,000
14,000
15,000
16,000

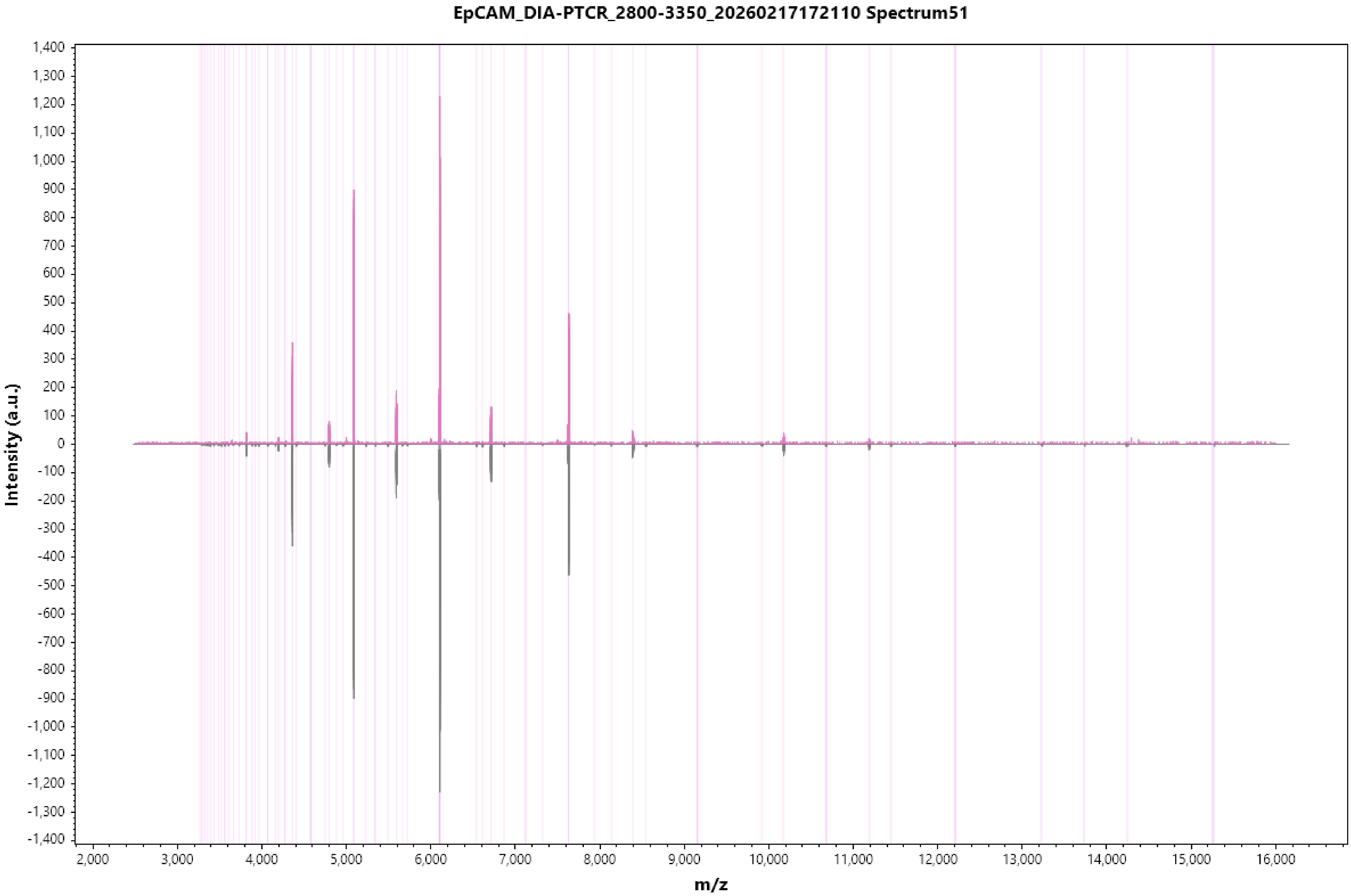
EpCAM_DIA-PTCR_2800-3350_20260217172110 Spectrum51
Intensity (a.u.)
m/z
1,400
1,300
1,200
1,100
1,000
900
800
700
600
500
400
300
200
100
0
-100
-200
-300
-400
-500
-600
-700
-800
-900
-1,000
-1,100
-1,200
-1,300
-1,400
2,000
3,000
4,000
5,000
6,000
7,000
8,000
9,000
10,000
11,000
12,000
13,000
14,000
15,000
16,000

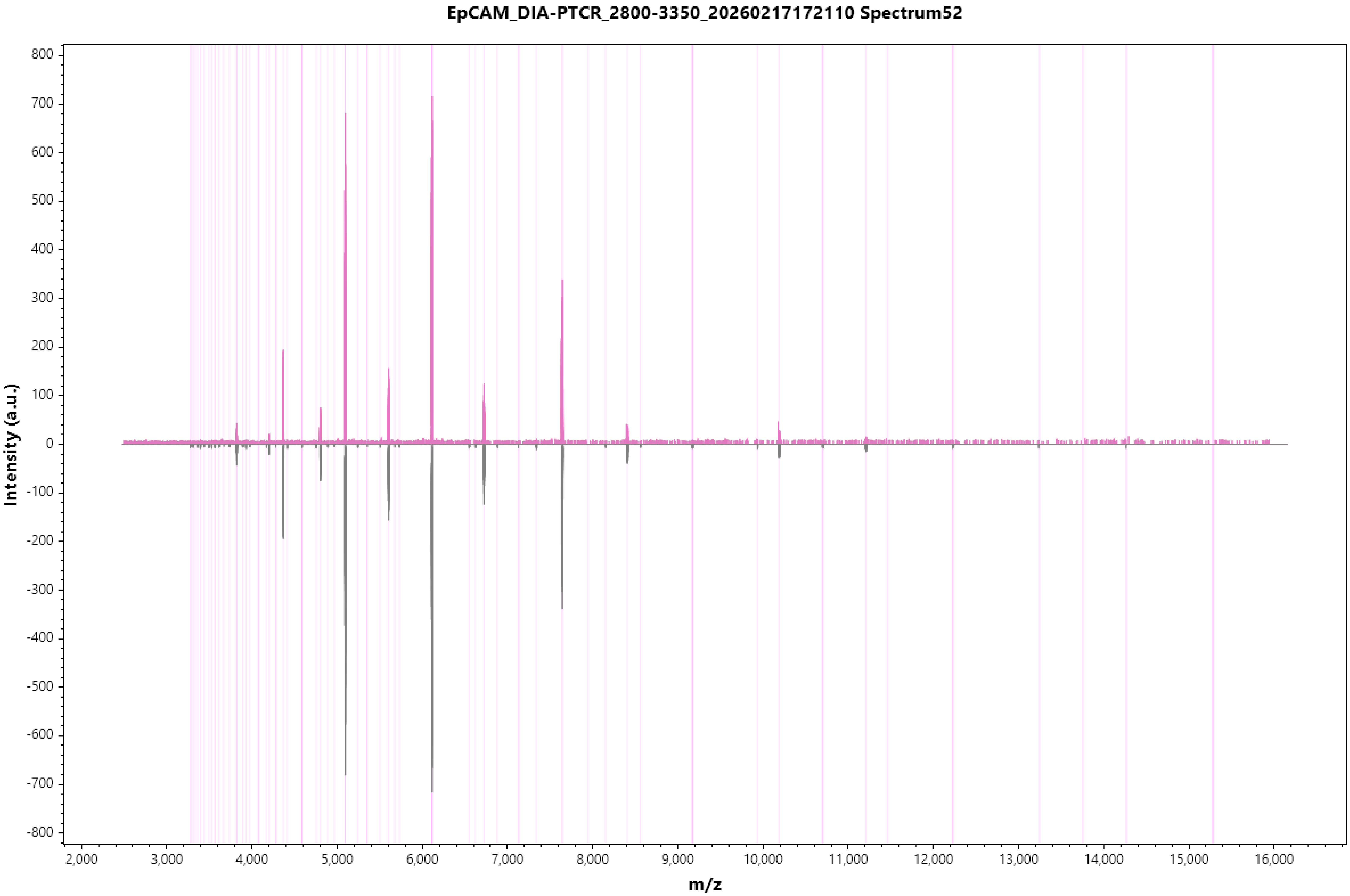
EpCAM_DIA-PTCR_2800-3350_20260217172110 Spectrum52
Intensity (a.u.)
800
700
600
500
400
300
200
100
0
-100
-200
-300
-400
-500
-600
-700
-800
2,000
3,000
4,000
5,000
6,000
7,000
8,000
9,000
10,000
11,000
12,000
13,000
14,000
15,000
16,000
m/z

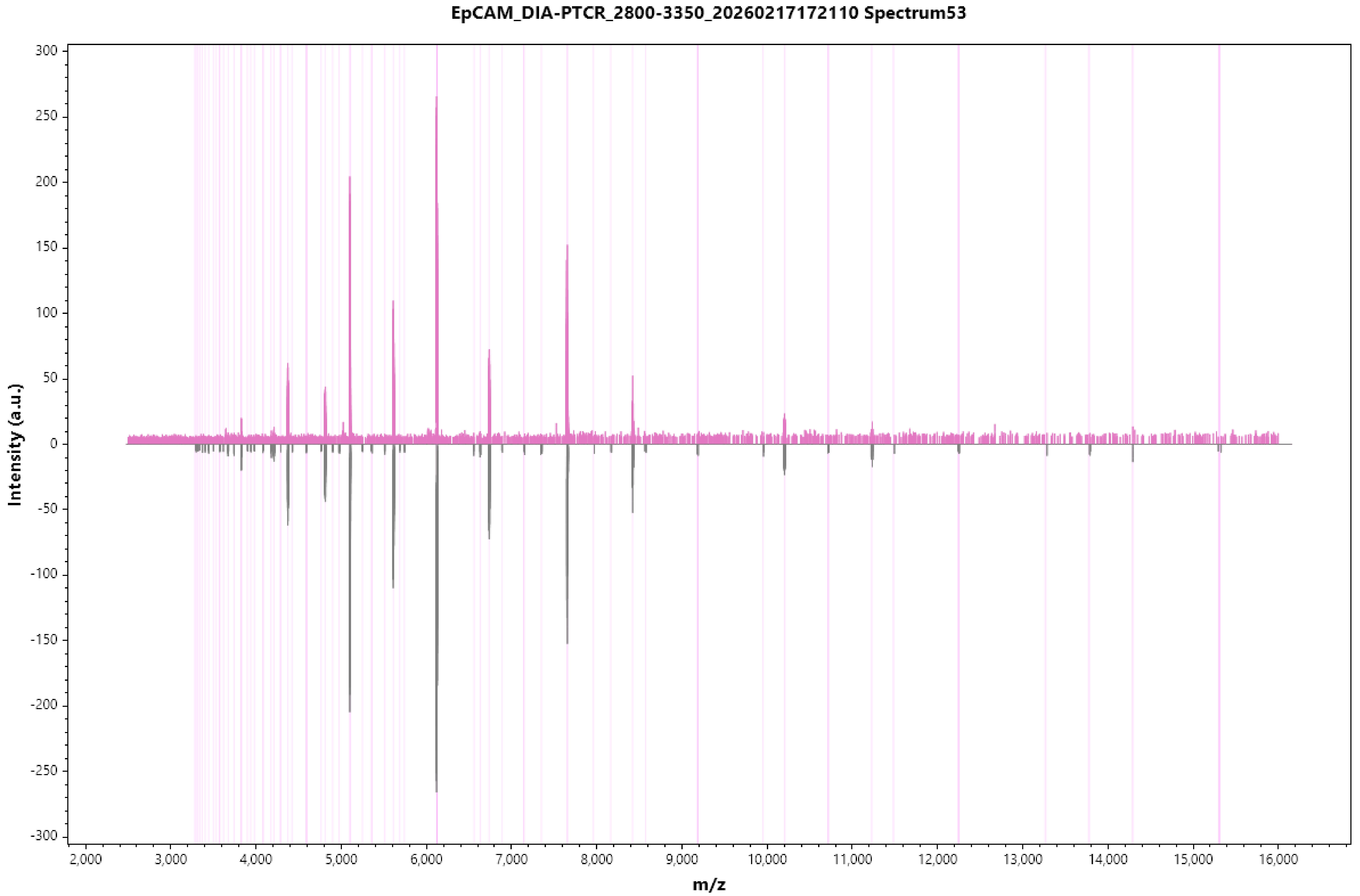
EpCAM_DIA-PTCR_2800-3350_20260217172110 Spectrum53
Intensity (a.u.)
300
250
200
150
100
50
0
-50
-100
-150
-200
-250
-300
2,000
3,000
4,000
5,000
6,000
7,000
8,000
9,000
10,000
11,000
12,000
13,000
14,000
15,000
16,000
m/z

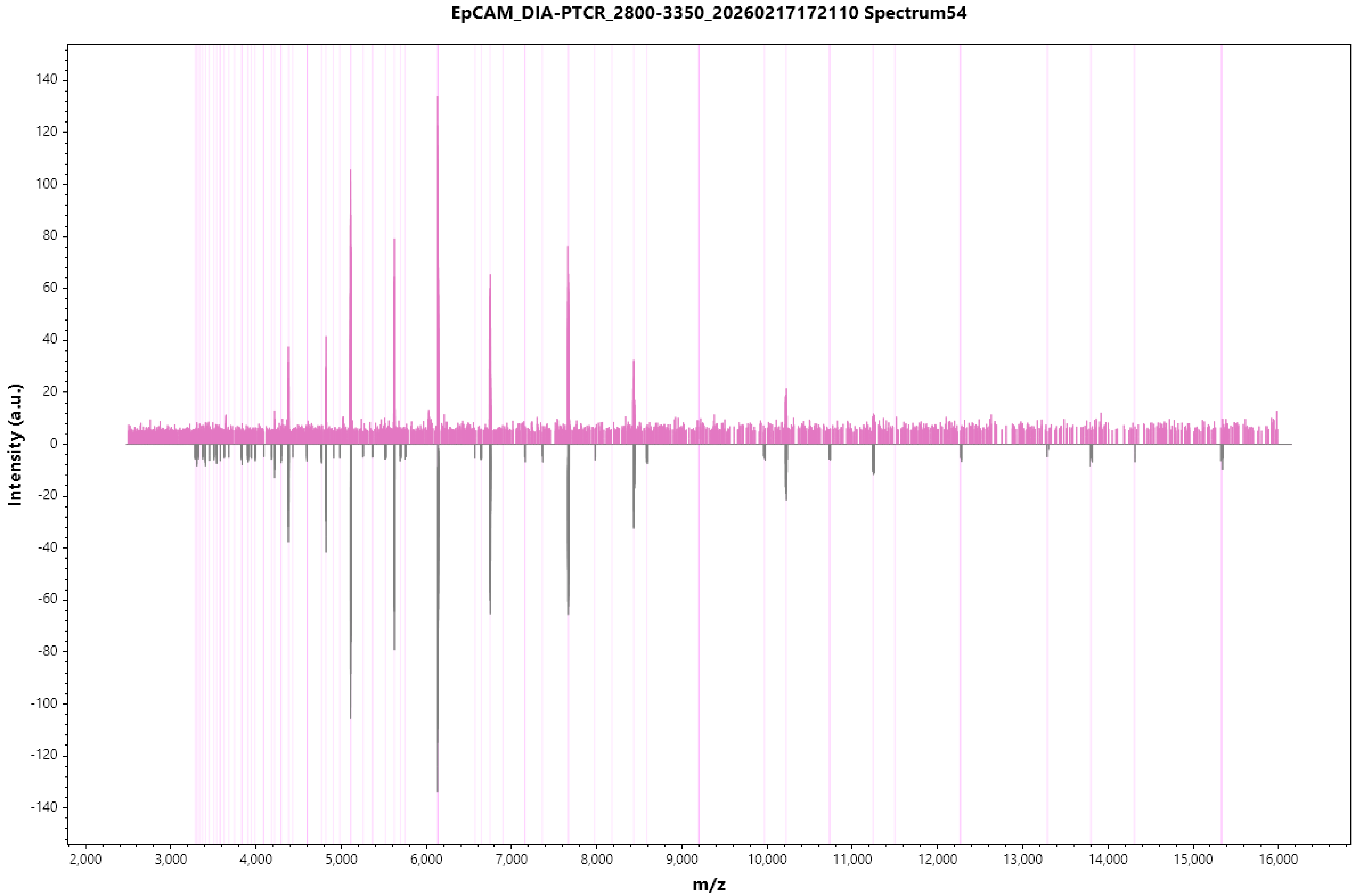
EpCAM_DIA-PTCR_2800-3350_20260217172110 Spectrum54
Intensity (a.u.)
m/z
140
120
100
80
60
40
20
0
-20
-40
-60
-80
-100
-120
-140
2,000
3,000
4,000
5,000
6,000
7,000
8,000
9,000
10,000
11,000
12,000
13,000
14,000
15,000
16,000

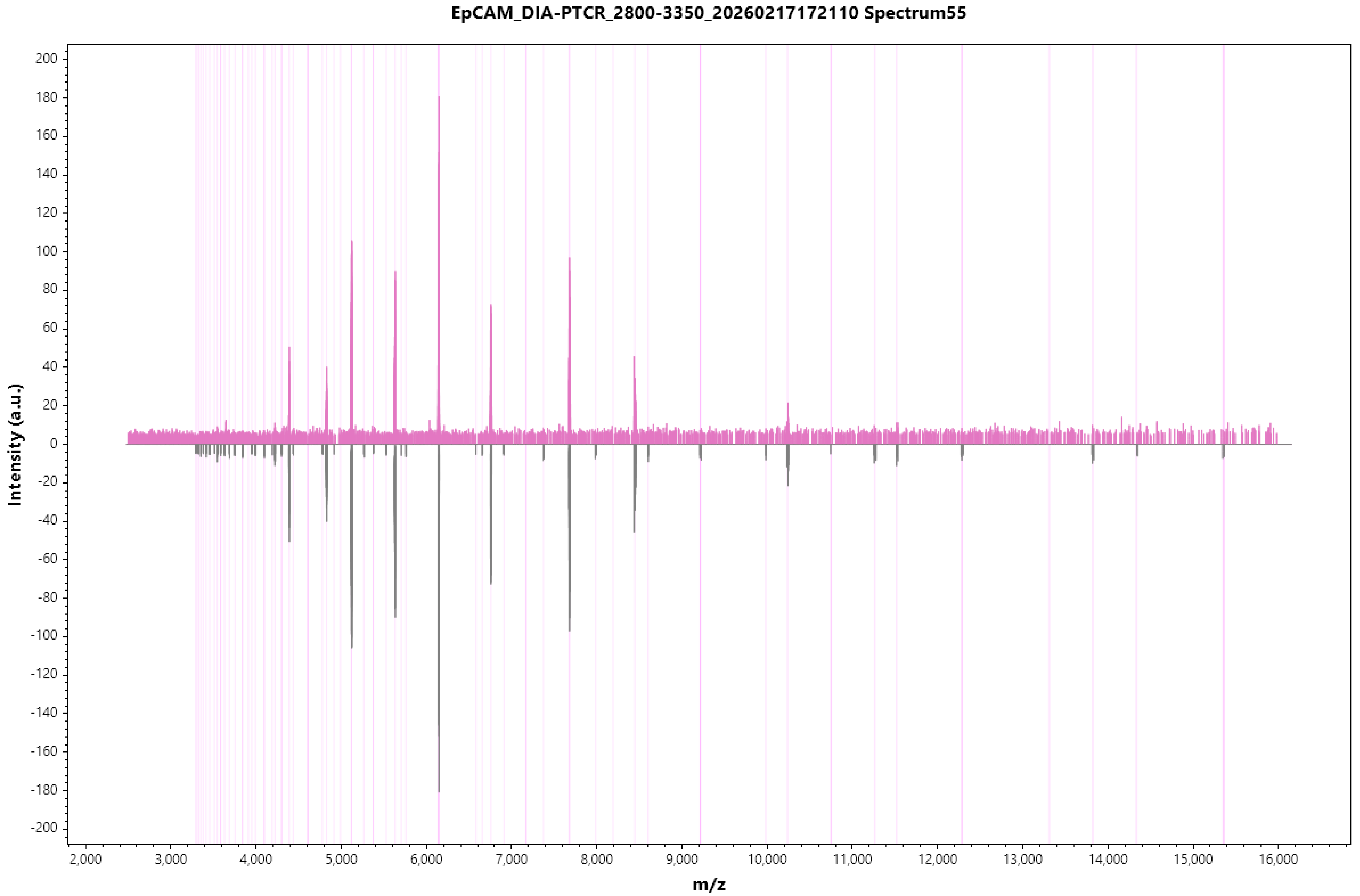
EpCAM_DIA-PTCR_2800-3350_20260217172110 Spectrum55
Intensity (a.u.)
m/z

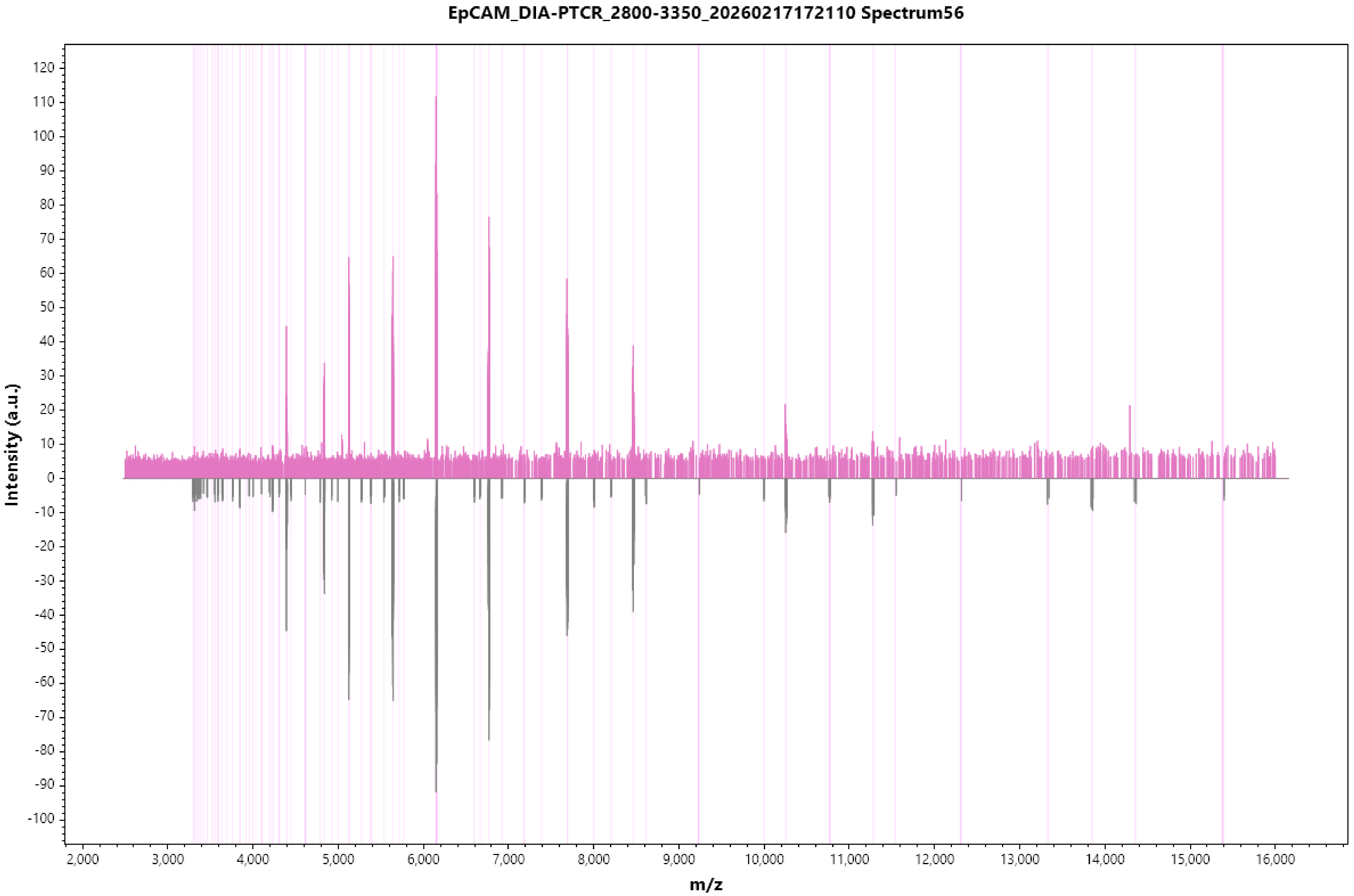
EpCAM_DIA-PTCR_2800-3350_20260217172110 Spectrum56
Intensity (a.u.)
m/z
120
110
100
90
80
70
60
50
40
30
20
10
0
-10
-20
-30
-40
-50
-60
-70
-80
-90
-100
2,000
3,000
4,000
5,000
6,000
7,000
8,000
9,000
10,000
11,000
12,000
13,000
14,000
15,000
16,000

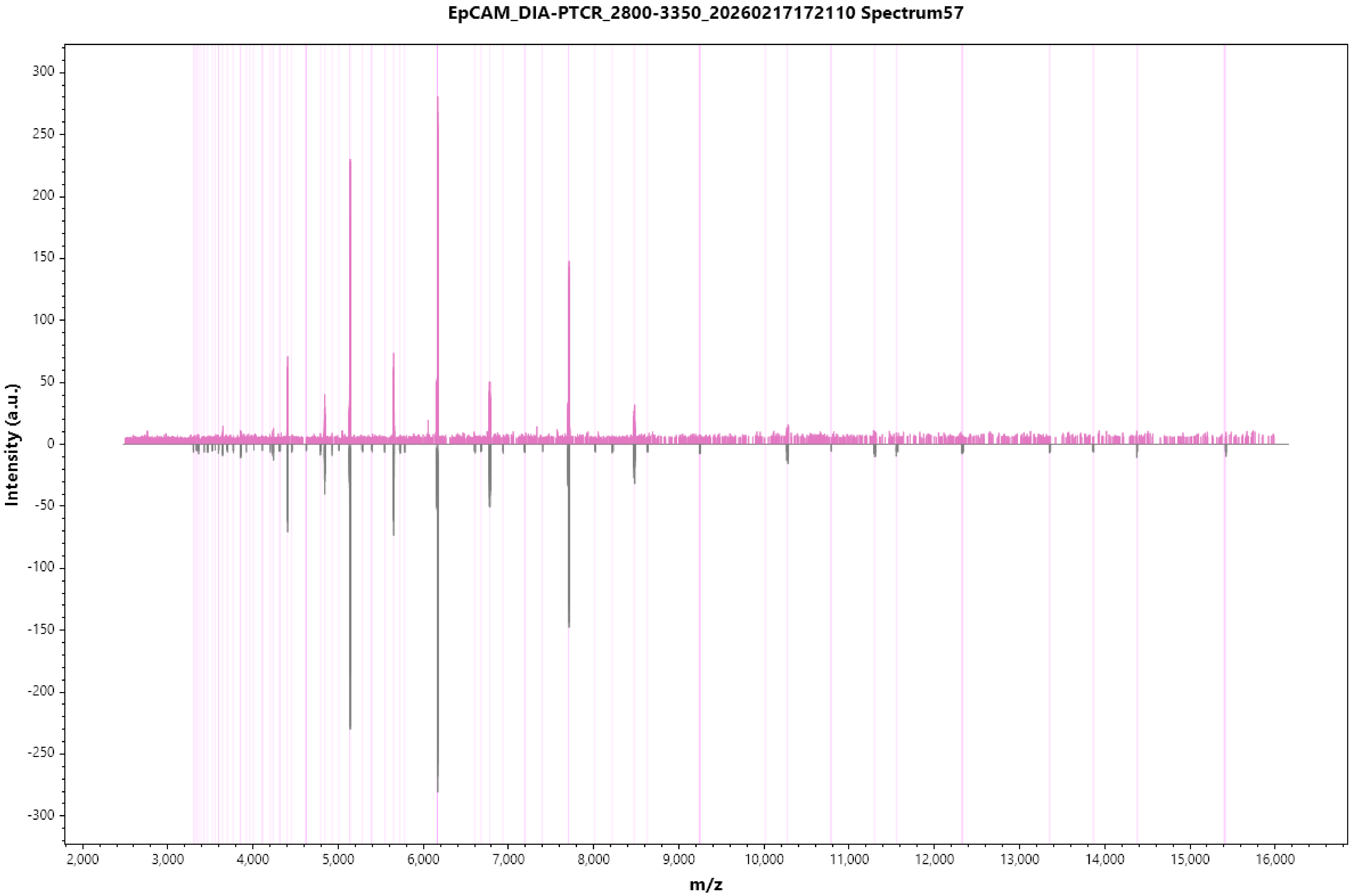
EpCAM_DIA-PTCR_2800-3350_20260217172110 Spectrum57
Intensity (a.u.)
300
250
200
150
100
50
0
-50
-100
-150
-200
-250
-300
2,000
3,000
4,000
5,000
6,000
7,000
8,000
9,000
10,000
11,000
12,000
13,000
14,000
15,000
16,000
m/z

EpCAM_DIA-PTCR_2800-3350_20260217172110 Spectrum58

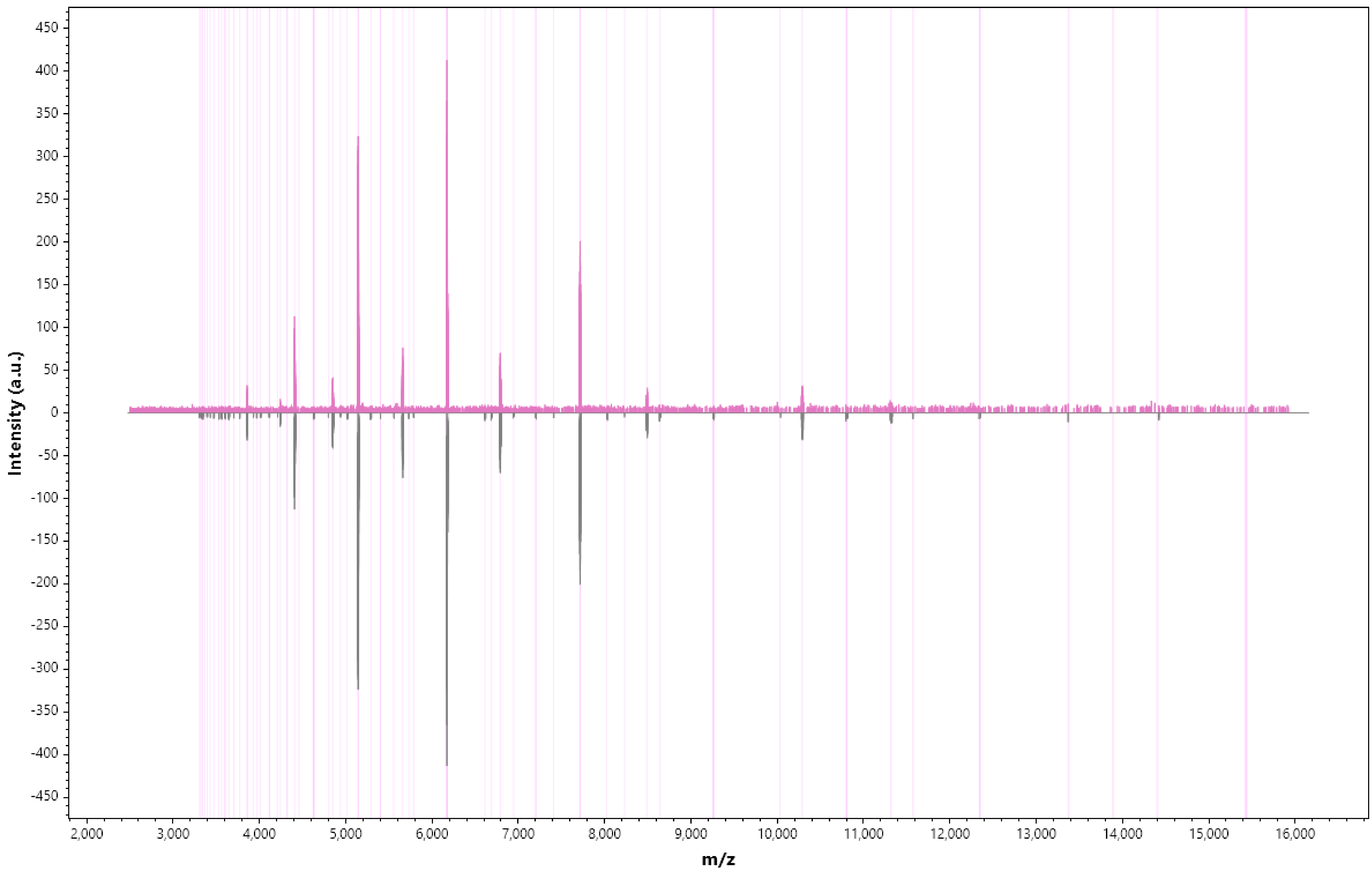

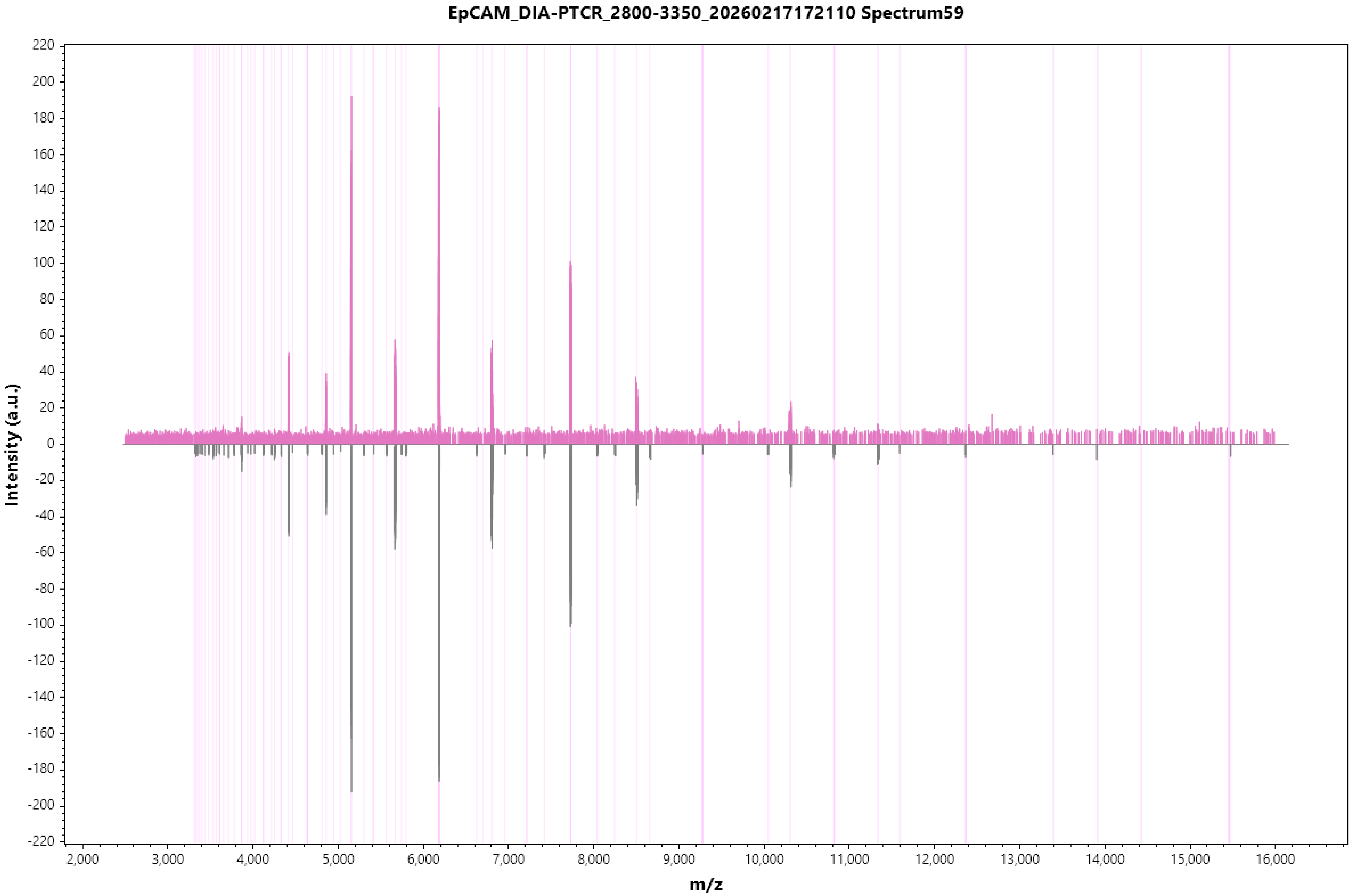

EpCAM_DIA-PTCR_2800-3350_20260217172110 Spectrum59
Intensity (a.u.)
m/z
220
200
180
160
140
120
100
80
60
40
20
0
-20
-40
-60
-80
-100
-120
-140
-160
-180
-200
-220
2,000
3,000
4,000
5,000
6,000
7,000
8,000
9,000
10,000
11,000
12,000
13,000
14,000
15,000
16,000

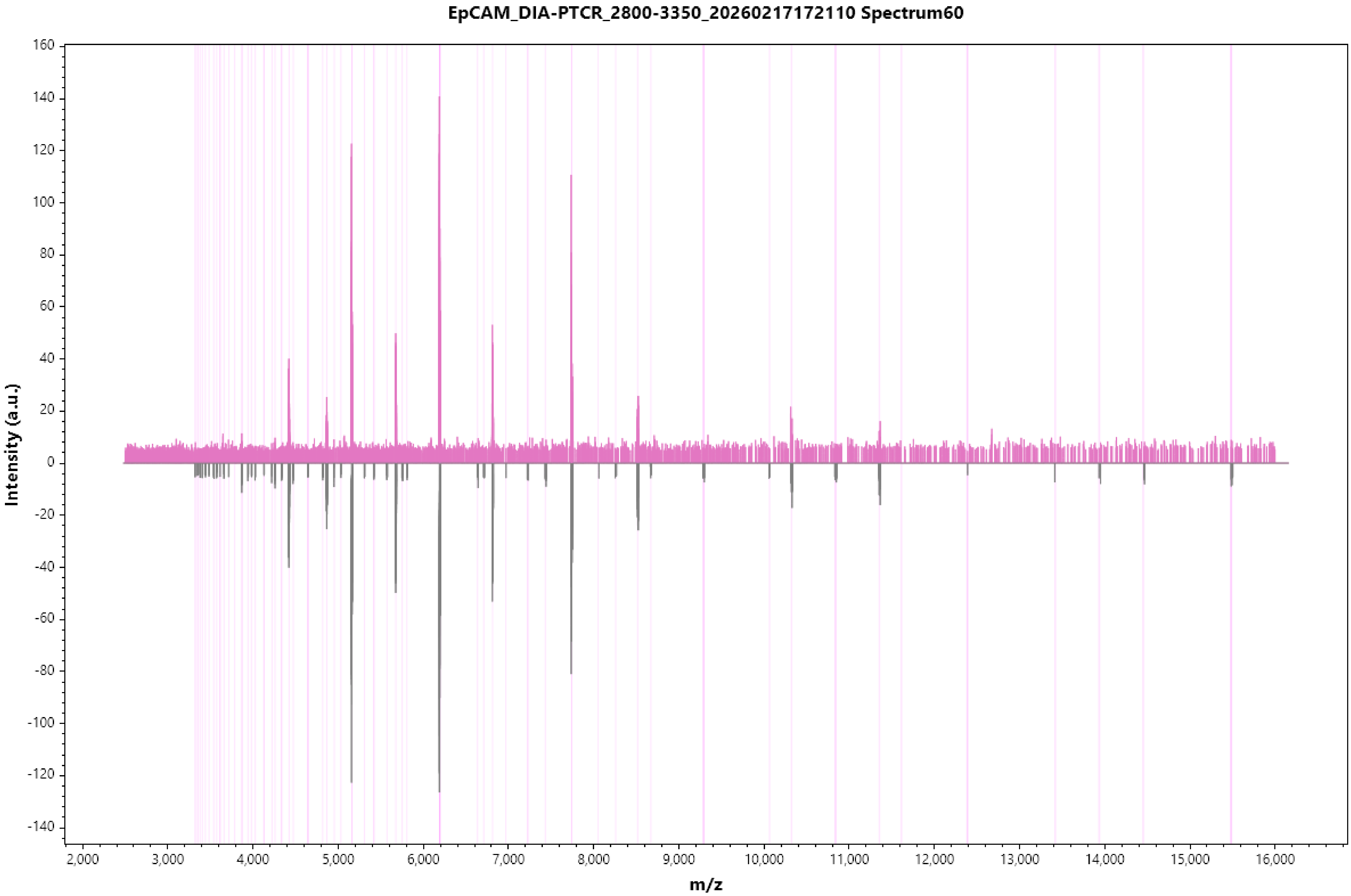

EpCAM_DIA-PTCR_2800-3350_20260217172110 Spectrum60
Intensity (a.u.)
160
140
120
100
80
60
40
20
0
-20
-40
-60
-80
-100
-120
-140
m/z
2,000
3,000
4,000
5,000
6,000
7,000
8,000
9,000
10,000
11,000
12,000
13,000
14,000
15,000
16,000

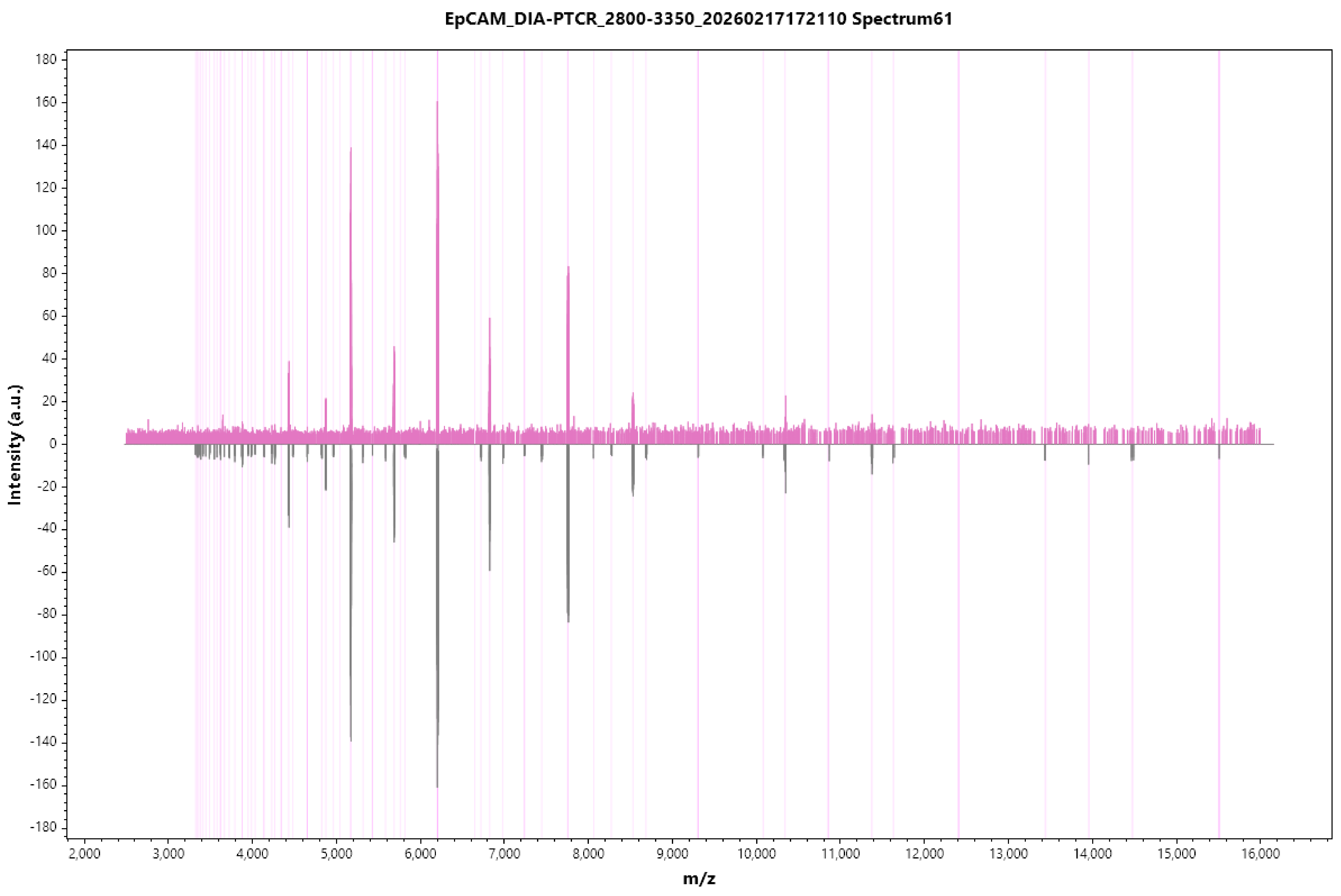
EpCAM_DIA-PTCR_2800-3350_20260217172110 Spectrum61
Intensity (a.u.)
m/z
180
160
140
120
100
80
60
40
20
0
-20
-40
-60
-80
-100
-120
-140
-160
-180
2,000
3,000
4,000
5,000
6,000
7,000
8,000
9,000
10,000
11,000
12,000
13,000
14,000
15,000
16,000

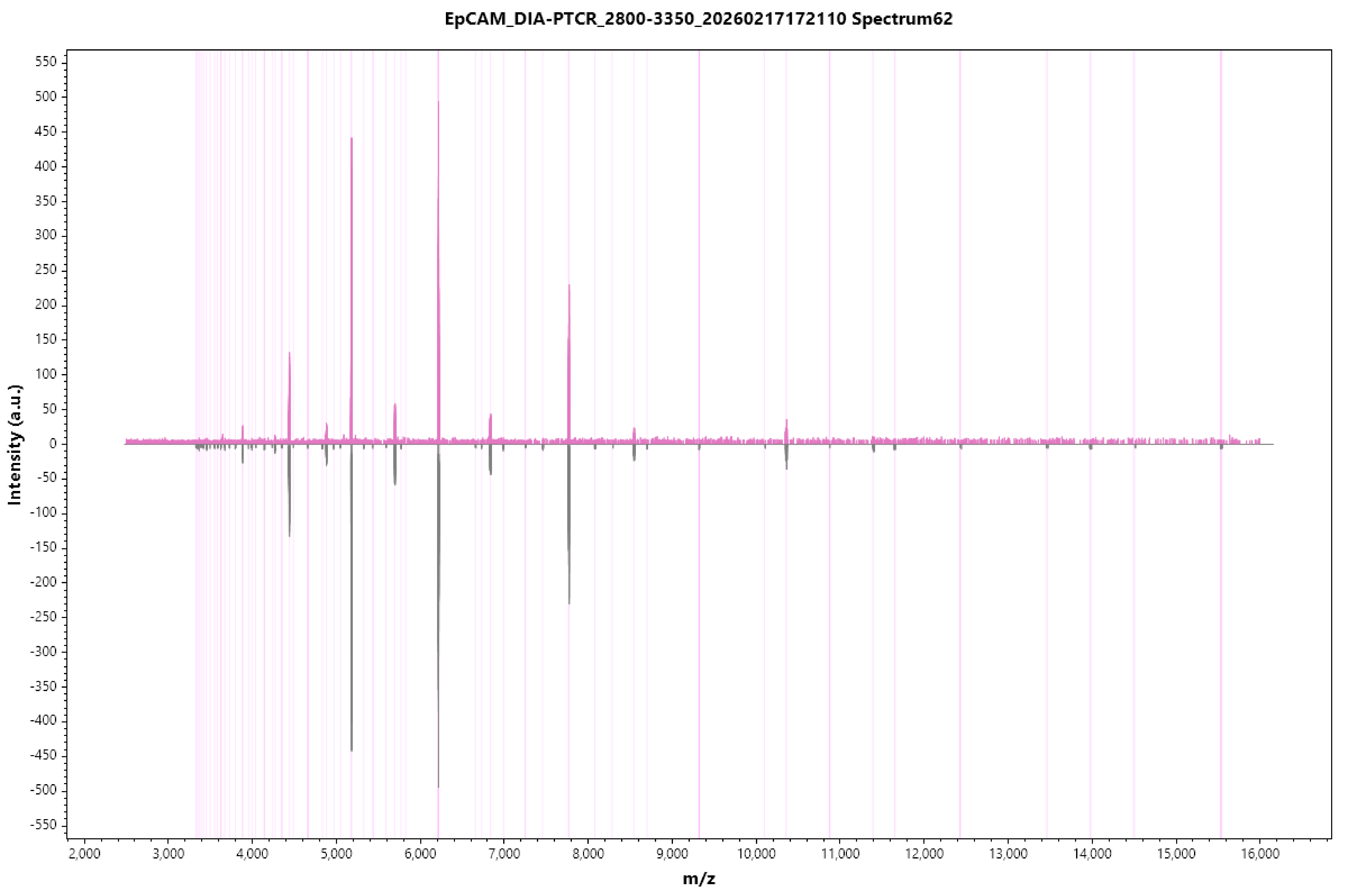

EpCAM_DIA-PTCR_2800-3350_20260217172110 Spectrum62
Intensity (a.u.)
m/z
550
500
450
400
350
300
250
200
150
100
50
0
-50
-100
-150
-200
-250
-300
-350
-400
-450
-500
-550
2,000
3,000
4,000
5,000
6,000
7,000
8,000
9,000
10,000
11,000
12,000
13,000
14,000
15,000
16,000

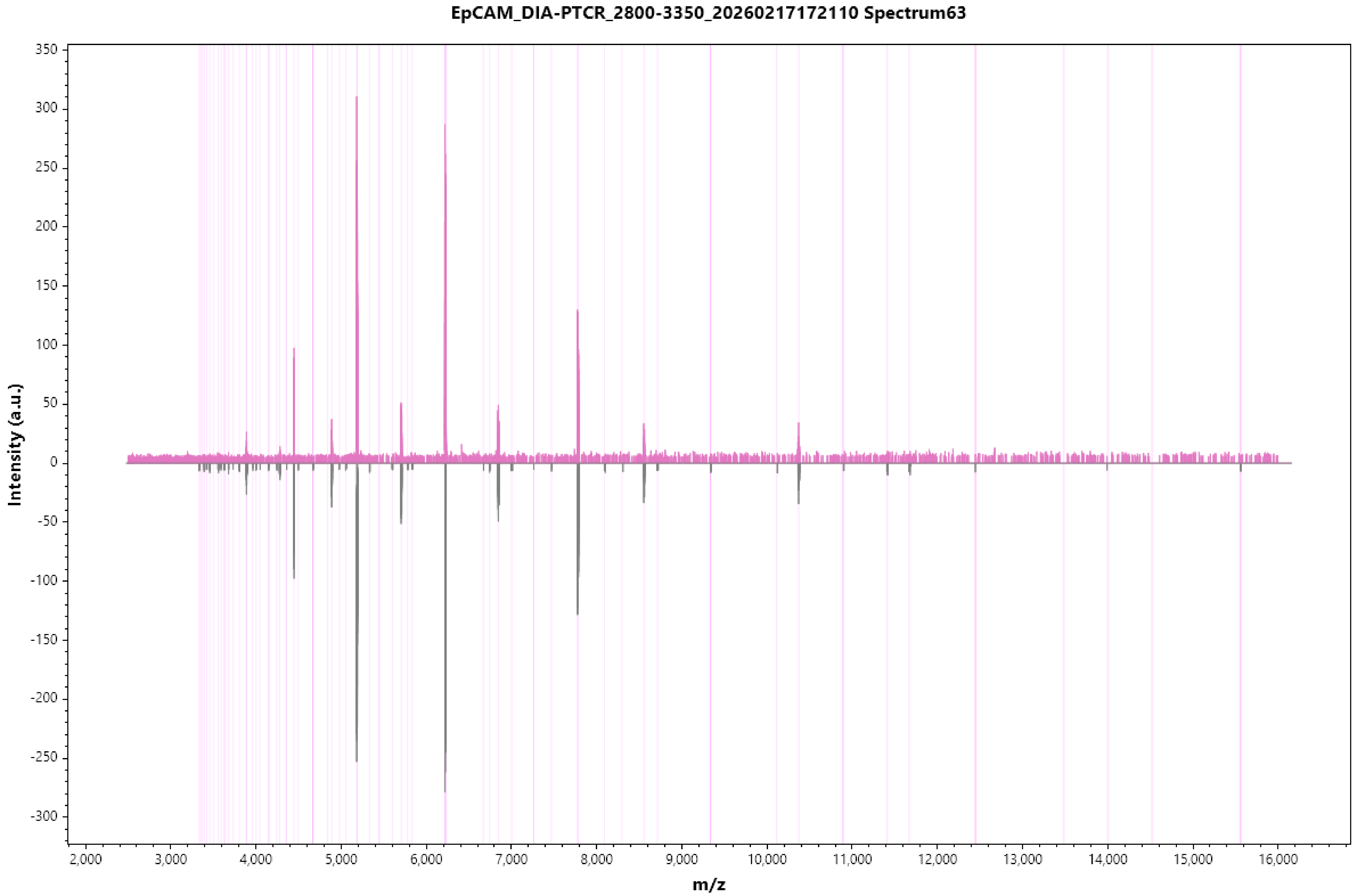
EpCAM_DIA-PTCR_2800-3350_20260217172110 Spectrum63
Intensity (a.u.)
m/z
350
300
250
200
150
100
50
0
-50
-100
-150
-200
-250
-300
2,000
3,000
4,000
5,000
6,000
7,000
8,000
9,000
10,000
11,000
12,000
13,000
14,000
15,000
16,000

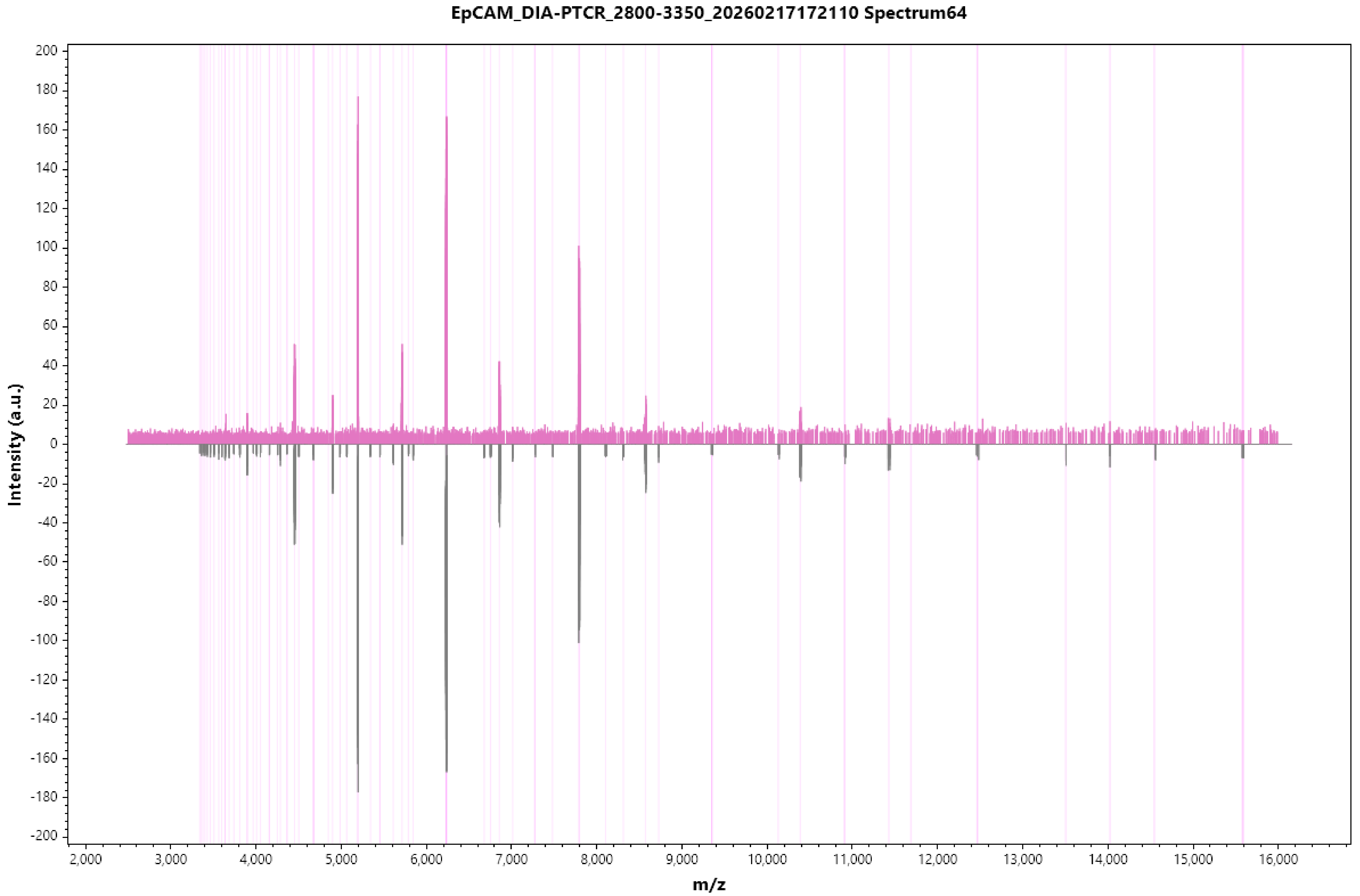
EpCAM_DIA-PTCR_2800-3350_20260217172110 Spectrum64
Intensity (a.u.)
m/z
200
180
160
140
120
100
80
60
40
20
0
-20
-40
-60
-80
-100
-120
-140
-160
-180
-200
2,000
3,000
4,000
5,000
6,000
7,000
8,000
9,000
10,000
11,000
12,000
13,000
14,000
15,000
16,000

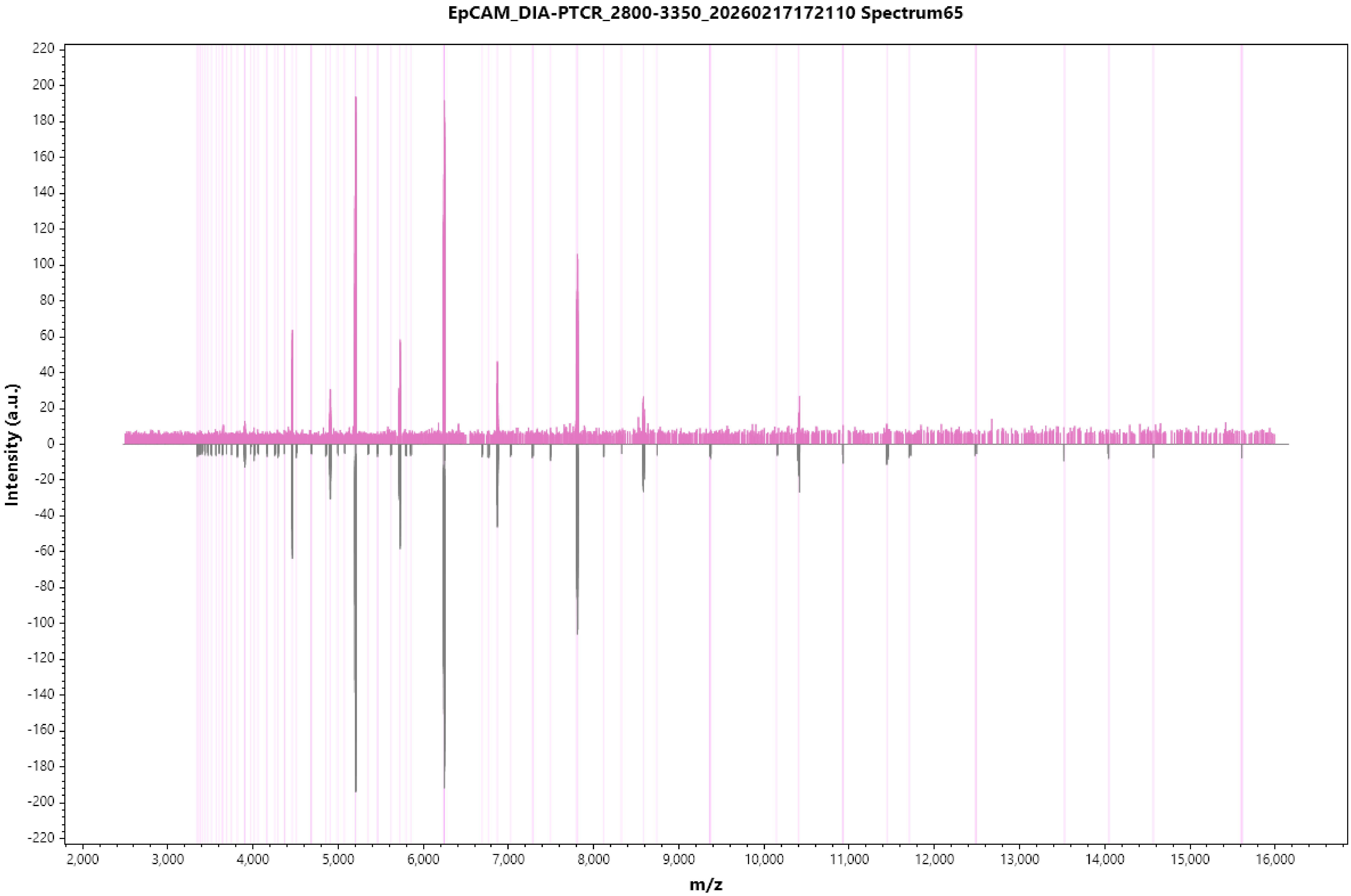

EpCAM_DIA-PTCR_2800-3350_20260217172110 Spectrum65
Intensity (a.u.)
m/z

#### EpCAM_DIA-PTCR_2800-3350_20260217172110 Spectrum66

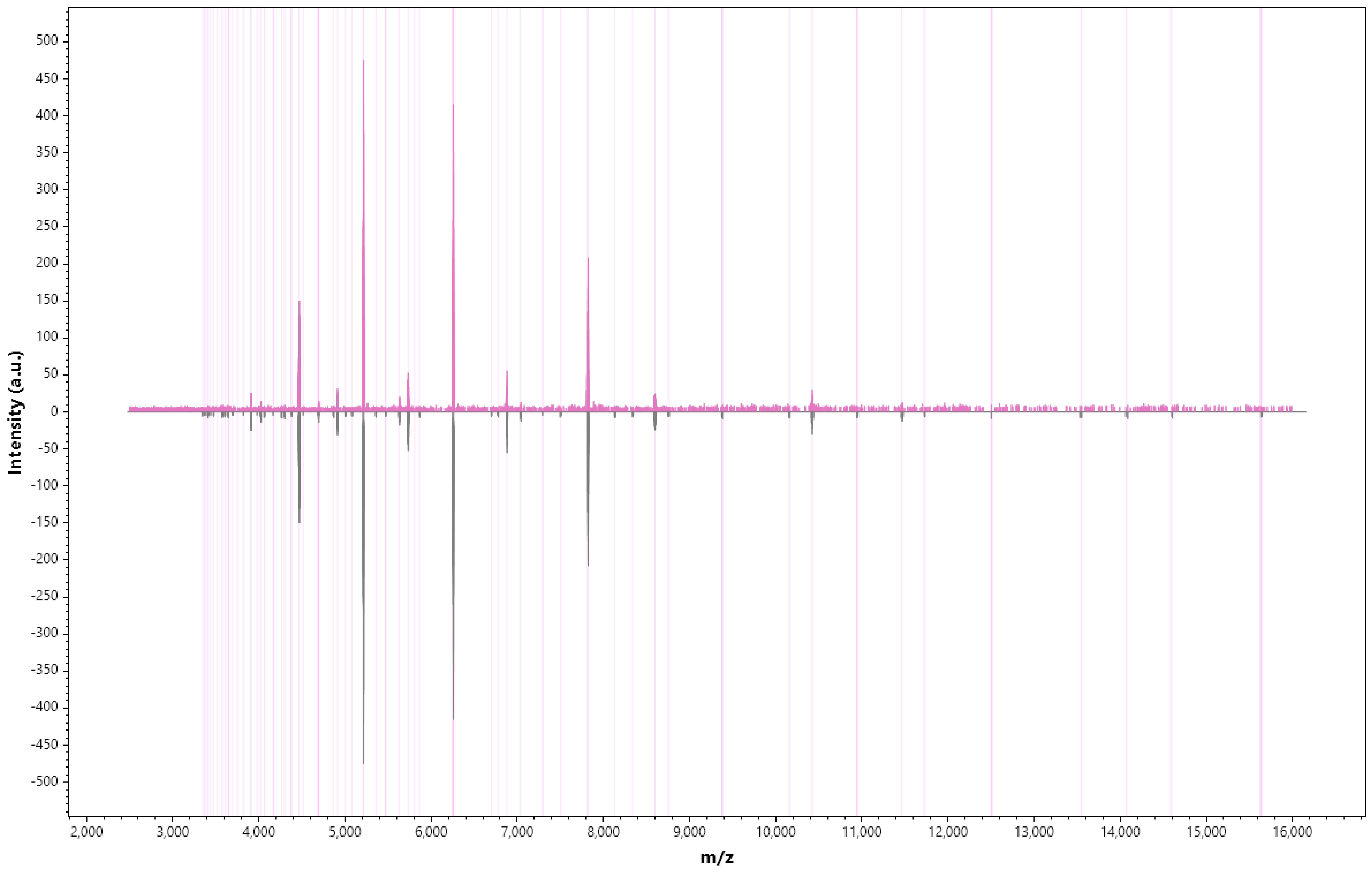

EpCAM_DIA-PTCR_2800-3350_20260217172110 Spectrum67

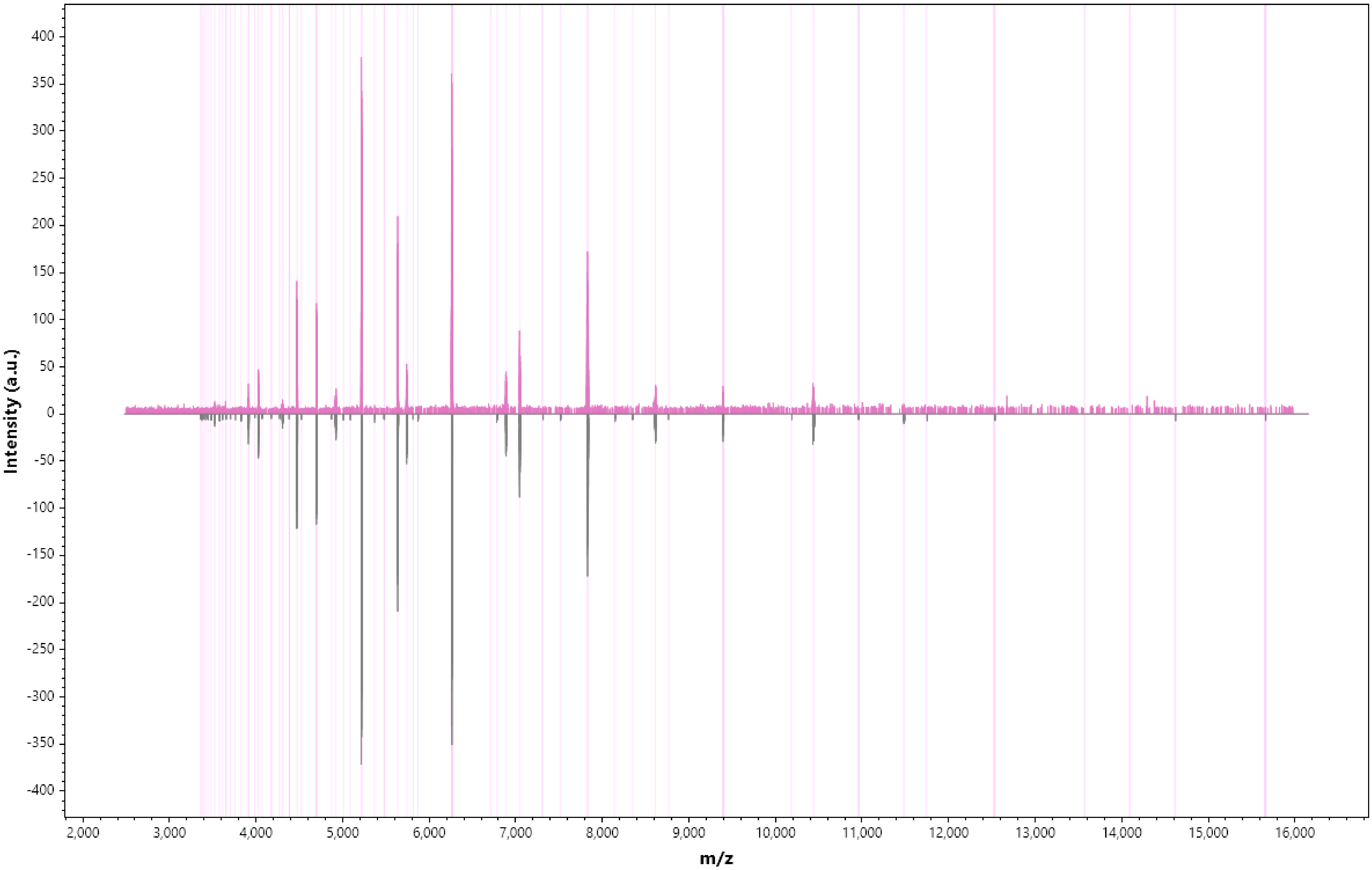

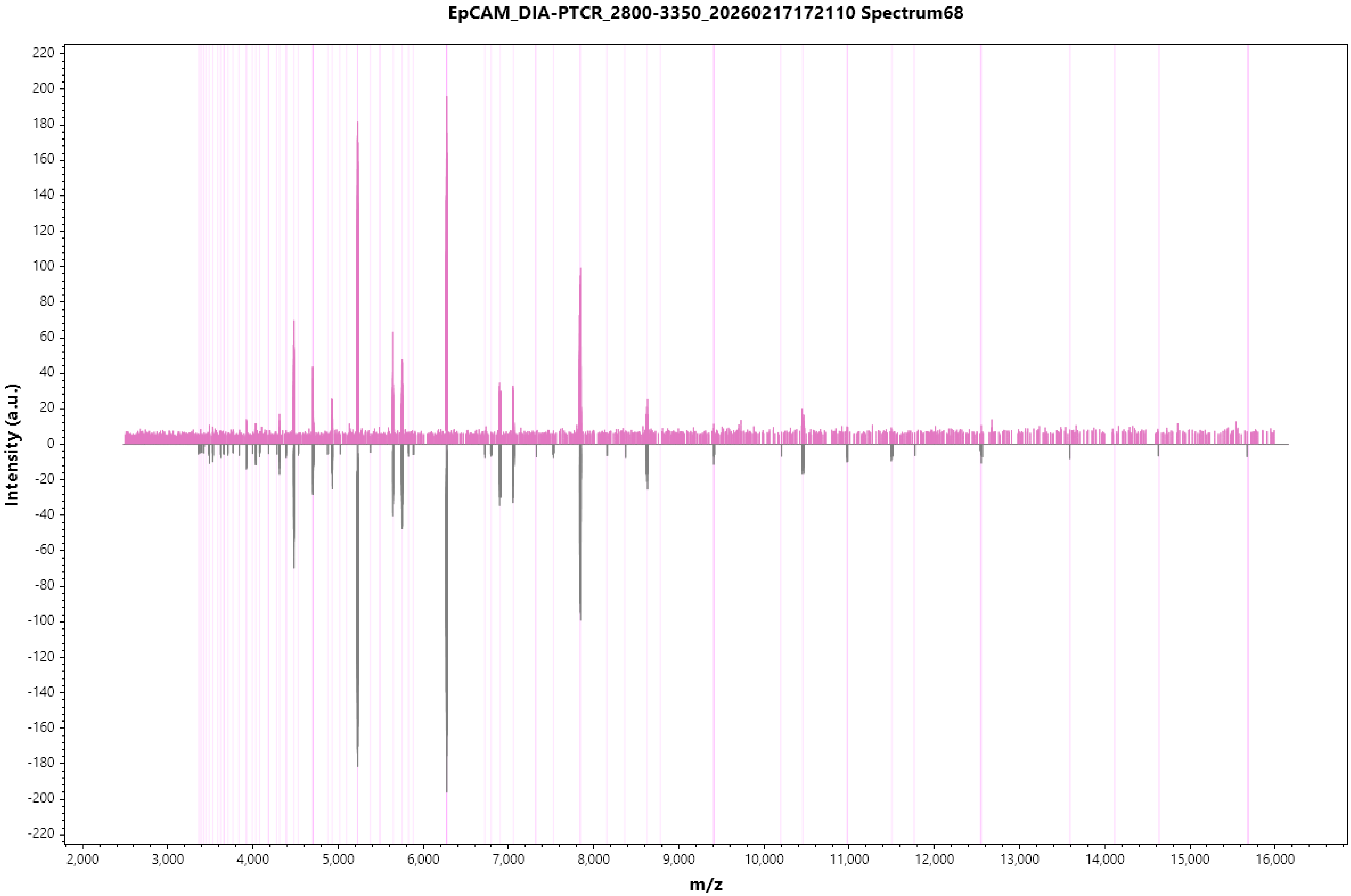
EpCAM_DIA-PTCR_2800-3350_20260217172110 Spectrum68
Intensity (a.u.)
m/z
220
200
180
160
140
120
100
80
60
40
20
0
-20
-40
-60
-80
-100
-120
-140
-160
-180
-200
-220
2,000
3,000
4,000
5,000
6,000
7,000
8,000
9,000
10,000
11,000
12,000
13,000
14,000
15,000
16,000

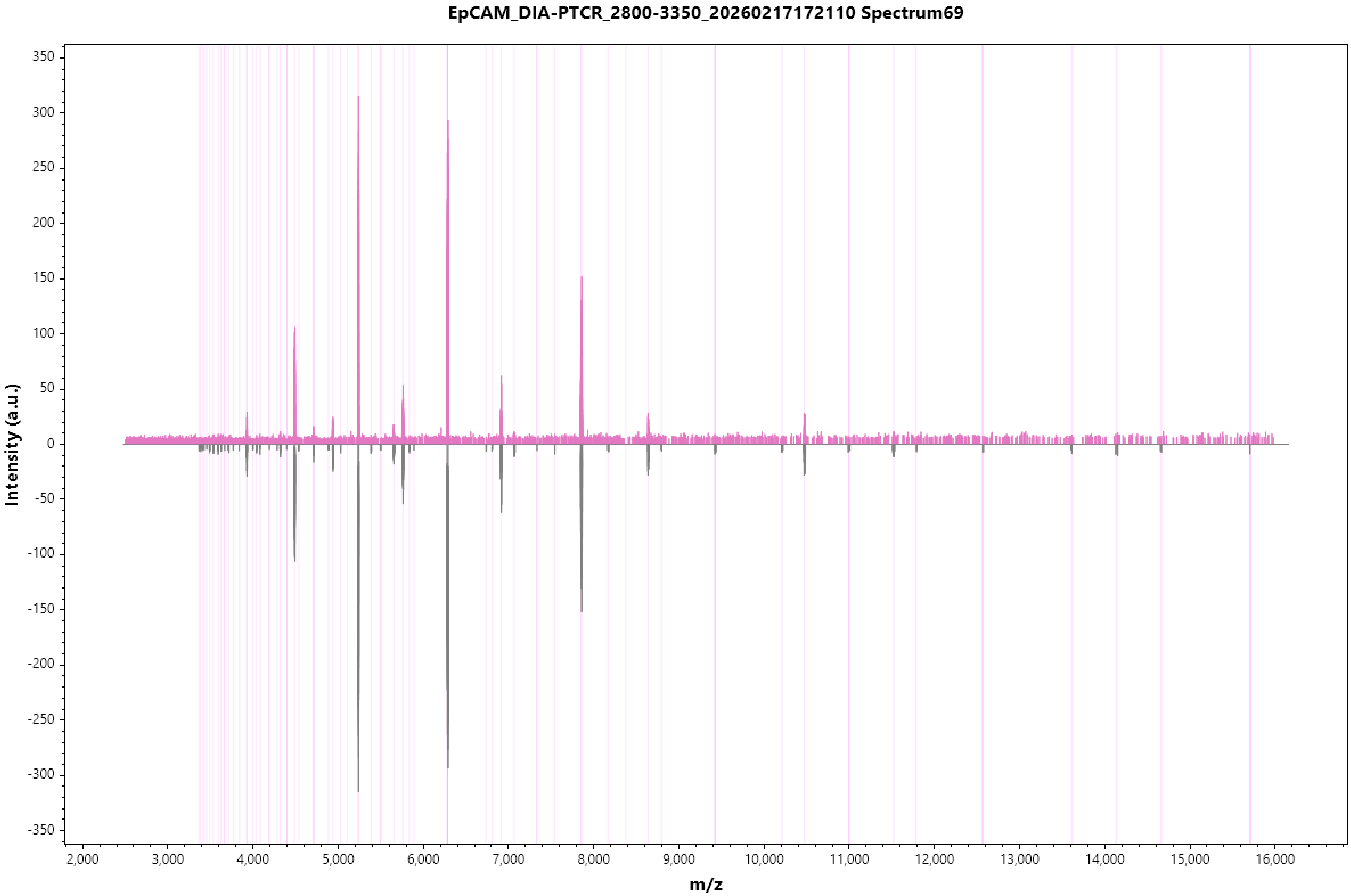

EpCAM_DIA-PTCR_2800-3350_20260217172110 Spectrum69
Intensity (a.u.)
350
300
250
200
150
100
50
0
-50
-100
-150
-200
-250
-300
-350
2,000
3,000
4,000
5,000
6,000
7,000
8,000
9,000
10,000
11,000
12,000
13,000
14,000
15,000
16,000
m/z

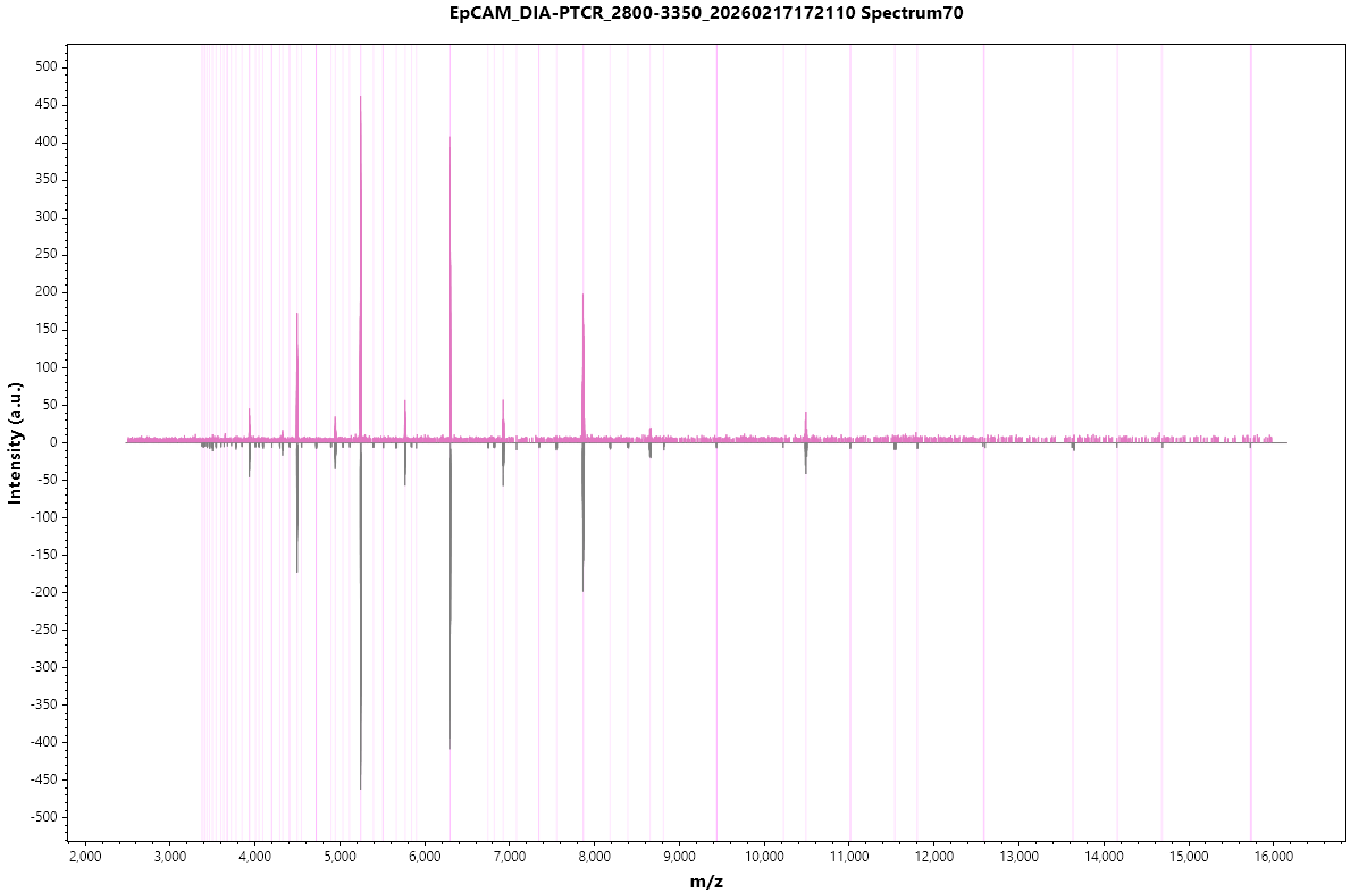
EpCAM_DIA-PTCR_2800-3350_20260217172110 Spectrum70
Intensity (a.u.)
m/z

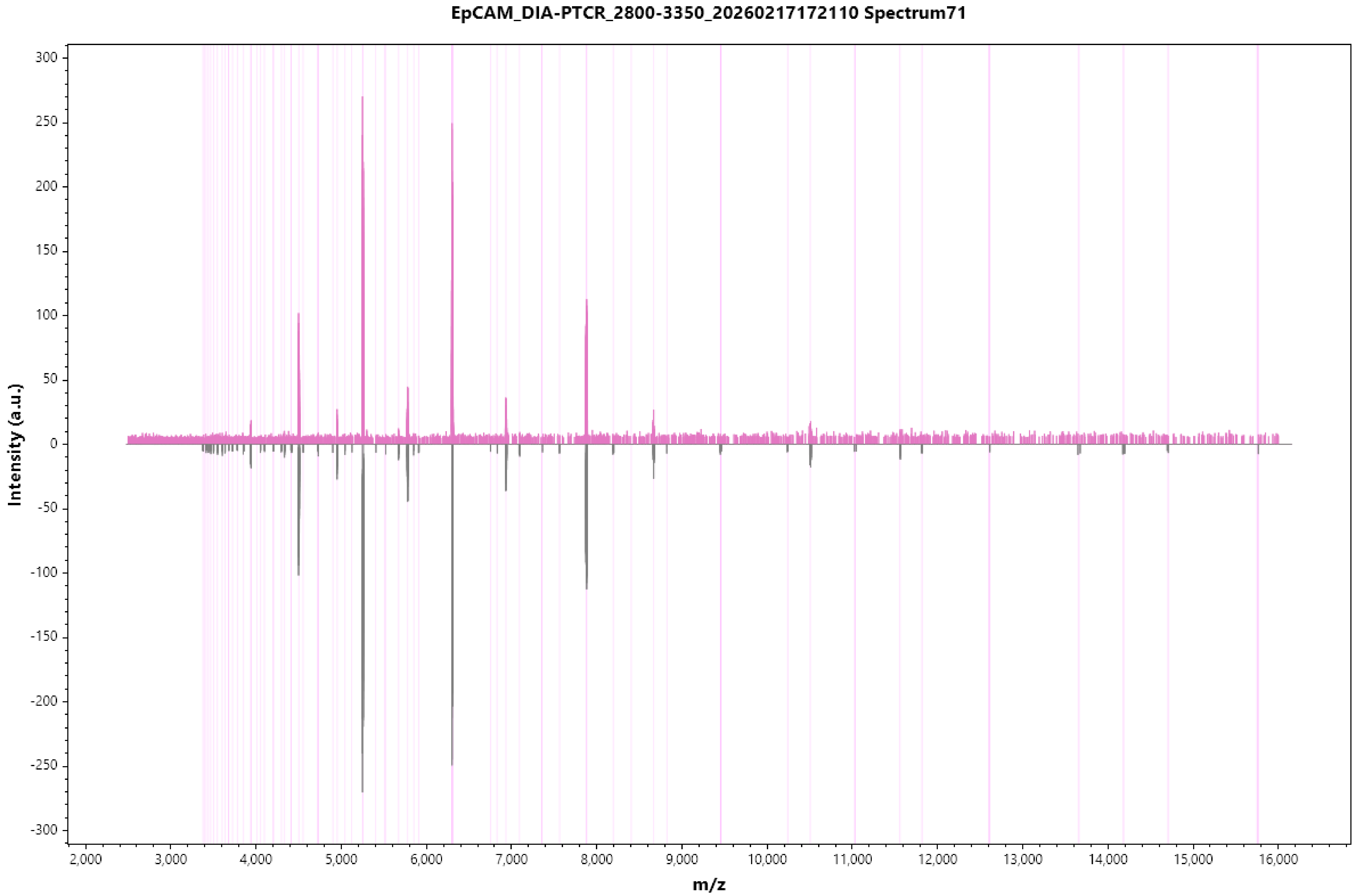
EpCAM_DIA-PTCR_2800-3350_20260217172110 Spectrum71
Intensity (a.u.)
m/z
300
250
200
150
100
50
0
-50
-100
-150
-200
-250
-300
2,000
3,000
4,000
5,000
6,000
7,000
8,000
9,000
10,000
11,000
12,000
13,000
14,000
15,000
16,000

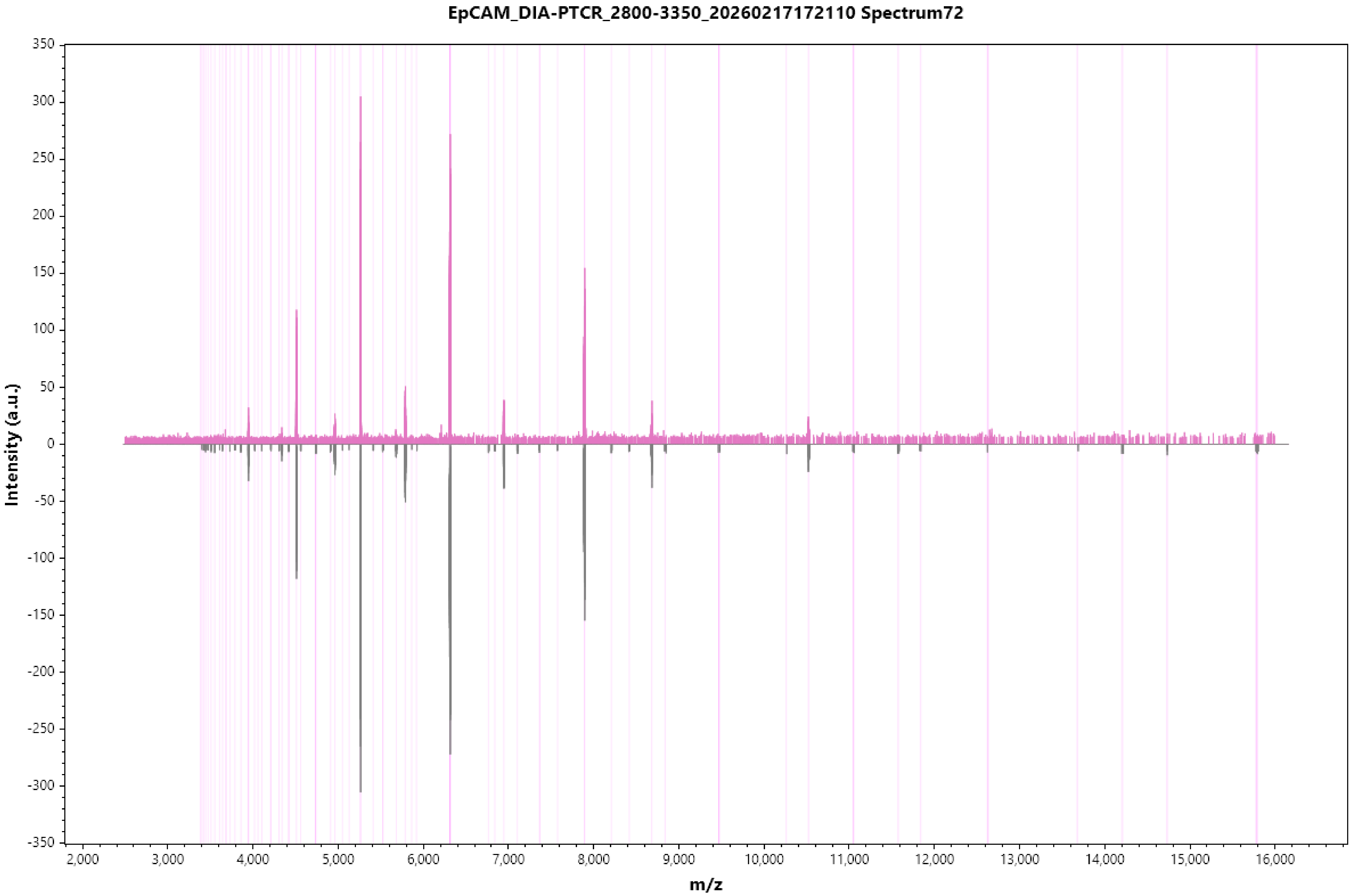
EpCAM_DIA-PTCR_2800-3350_20260217172110 Spectrum72
Intensity (a.u.)
m/z
350
300
250
200
150
100
50
0
-50
-100
-150
-200
-250
-300
-350
2,000
3,000
4,000
5,000
6,000
7,000
8,000
9,000
10,000
11,000
12,000
13,000
14,000
15,000
16,000

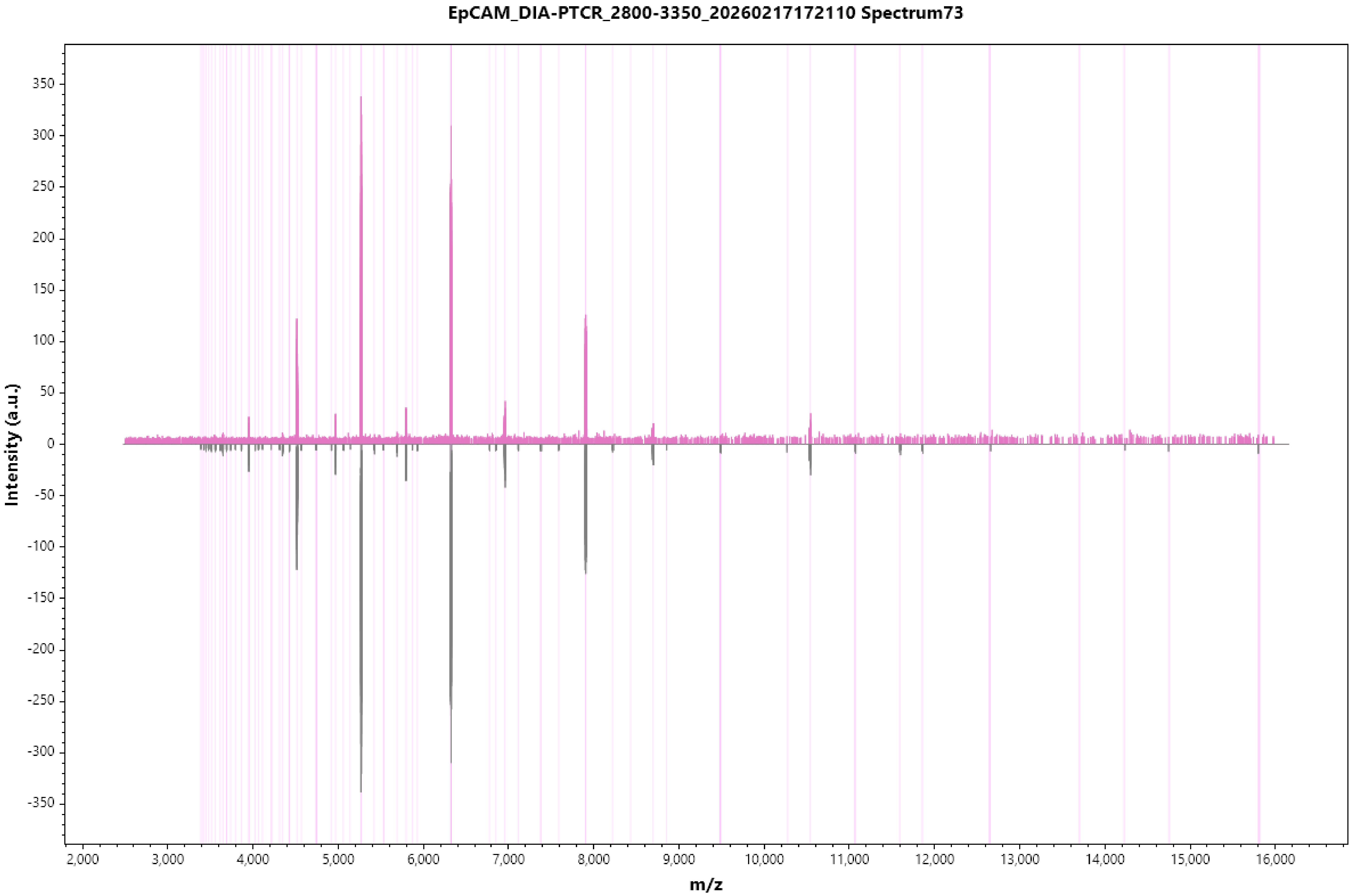
EpCAM_DIA-PTCR_2800-3350_20260217172110 Spectrum73
Intensity (a.u.)
m/z

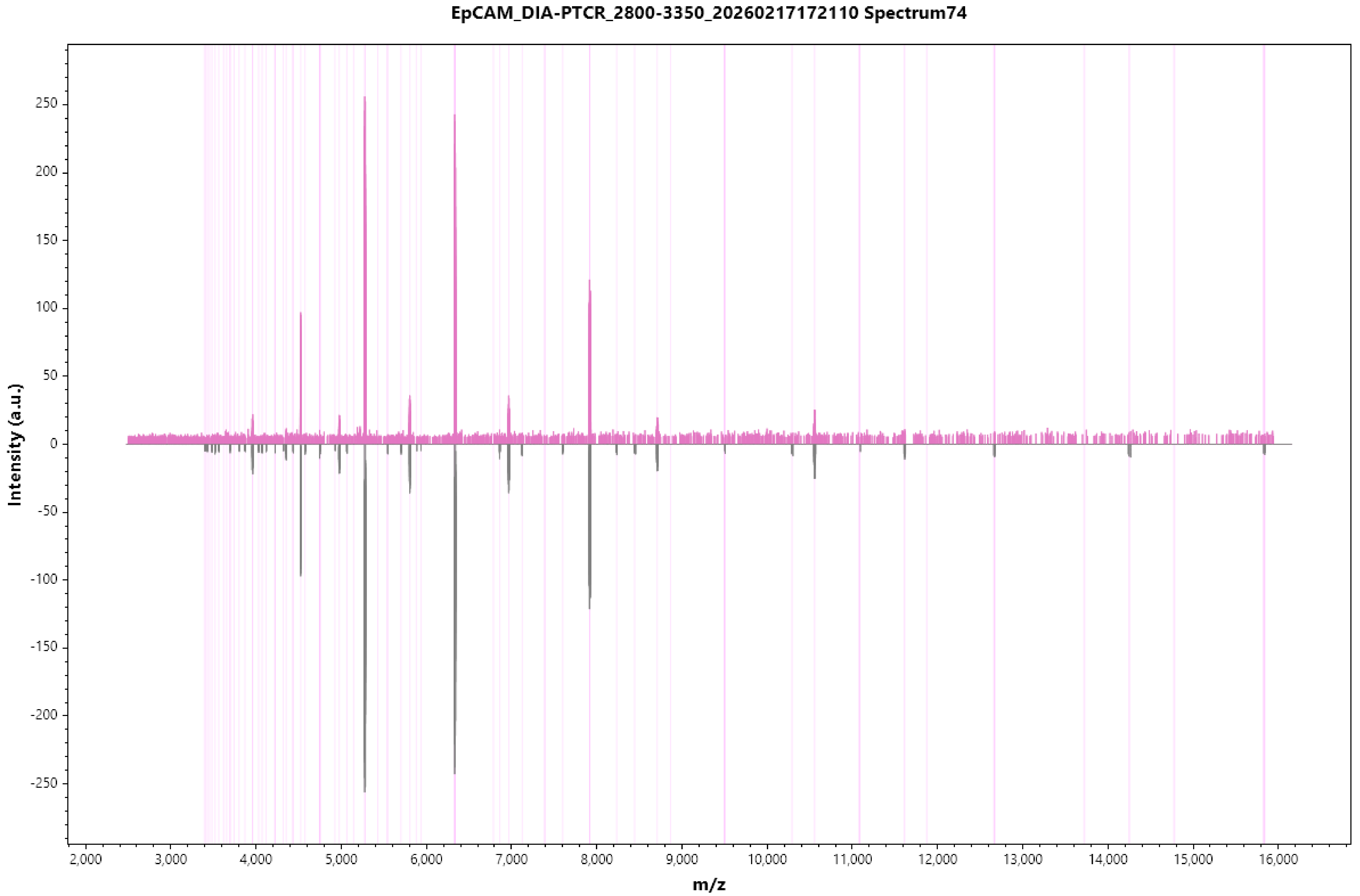
EpCAM_DIA-PTCR_2800-3350_20260217172110 Spectrum74
Intensity (a.u.)
m/z
250
200
150
100
50
0
-50
-100
-150
-200
-250
2,000
3,000
4,000
5,000
6,000
7,000
8,000
9,000
10,000
11,000
12,000
13,000
14,000
15,000
16,000

EpCAM_DIA-PTCR_2800-3350_20260217172110 Spectrum75

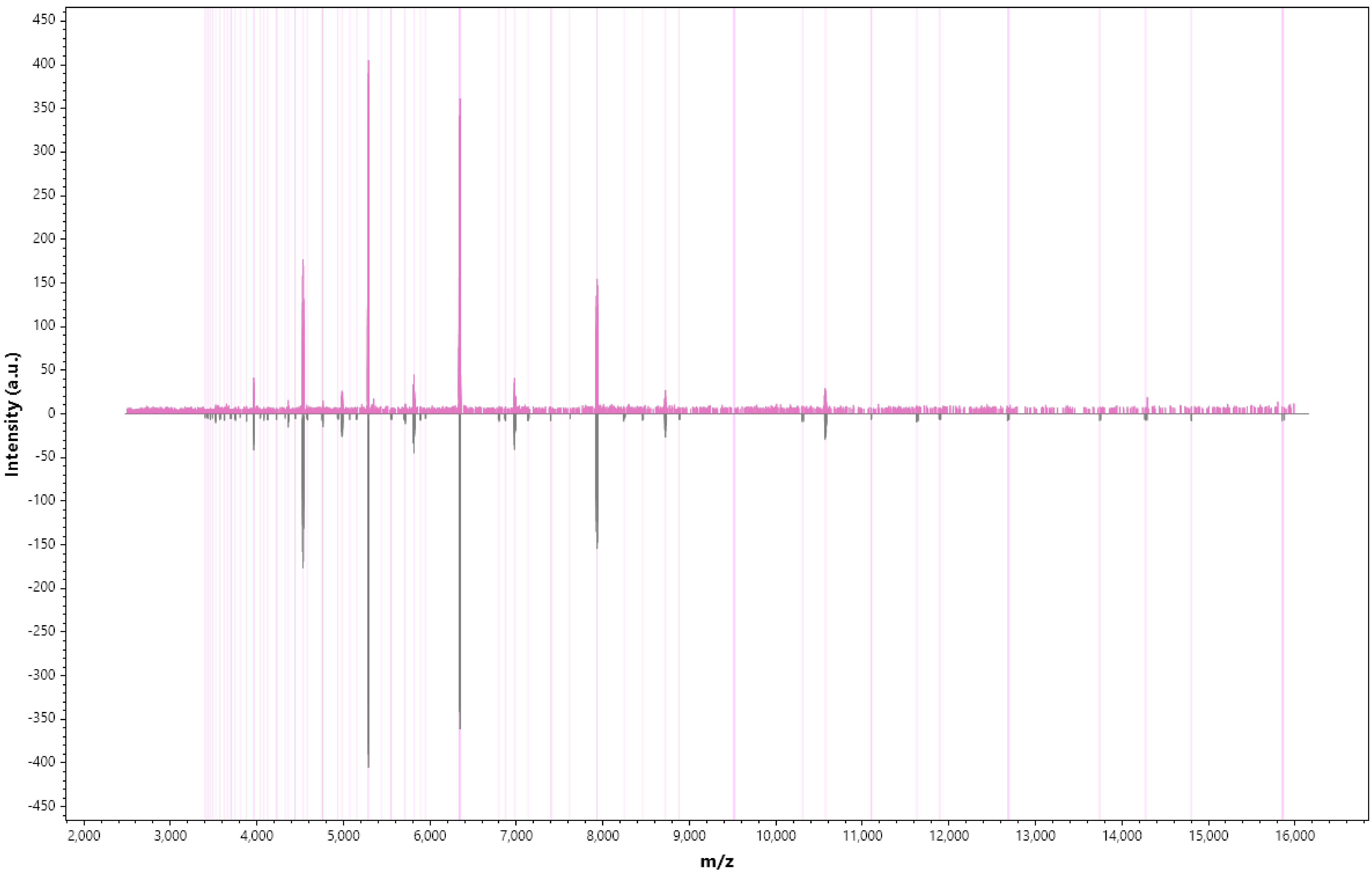

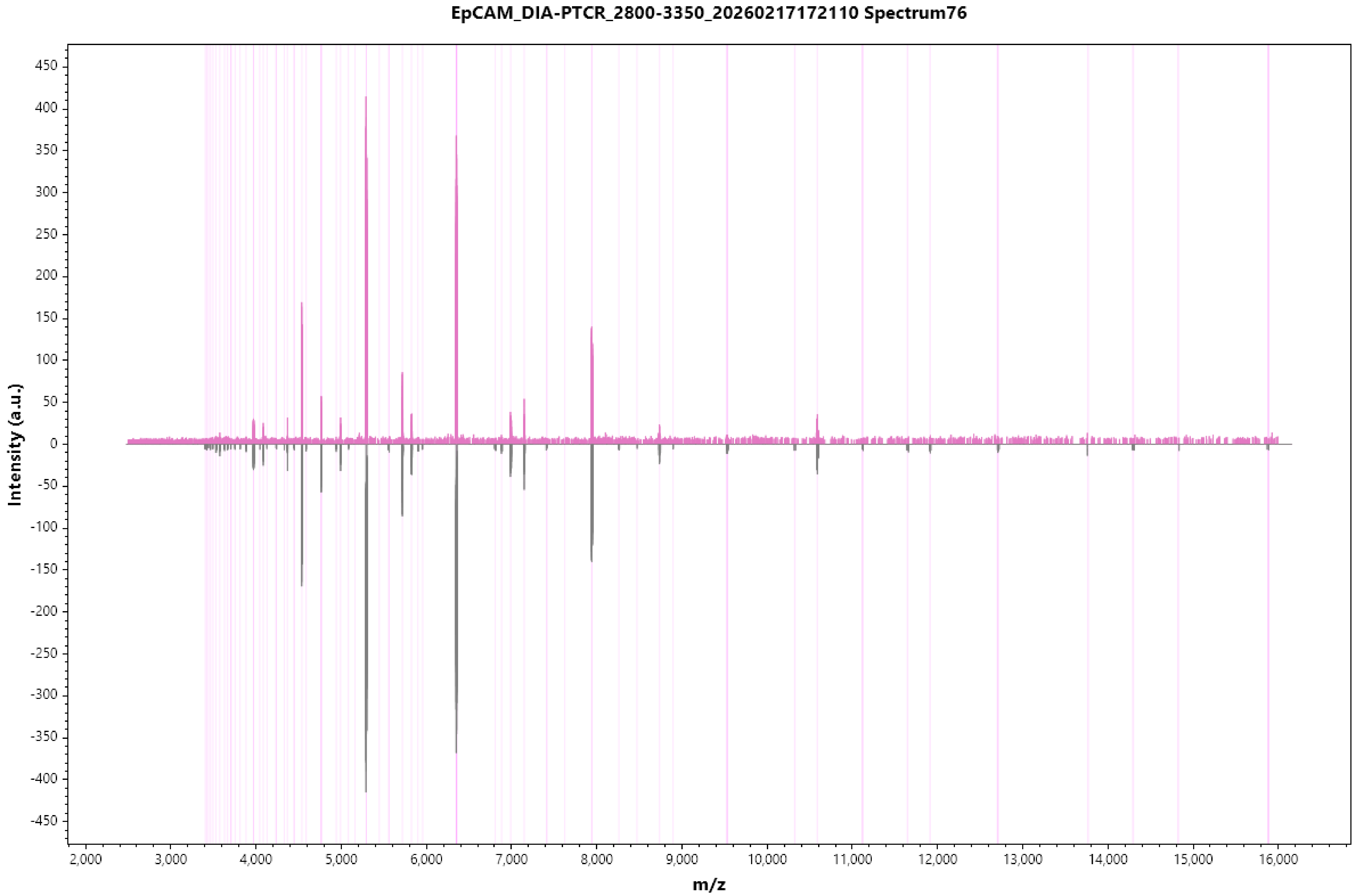

EpCAM_DIA-PTCR_2800-3350_20260217172110 Spectrum76
Intensity (a.u.)
m/z
450
400
350
300
250
200
150
100
50
0
-50
-100
-150
-200
-250
-300
-350
-400
-450
2,000
3,000
4,000
5,000
6,000
7,000
8,000
9,000
10,000
11,000
12,000
13,000
14,000
15,000
16,000

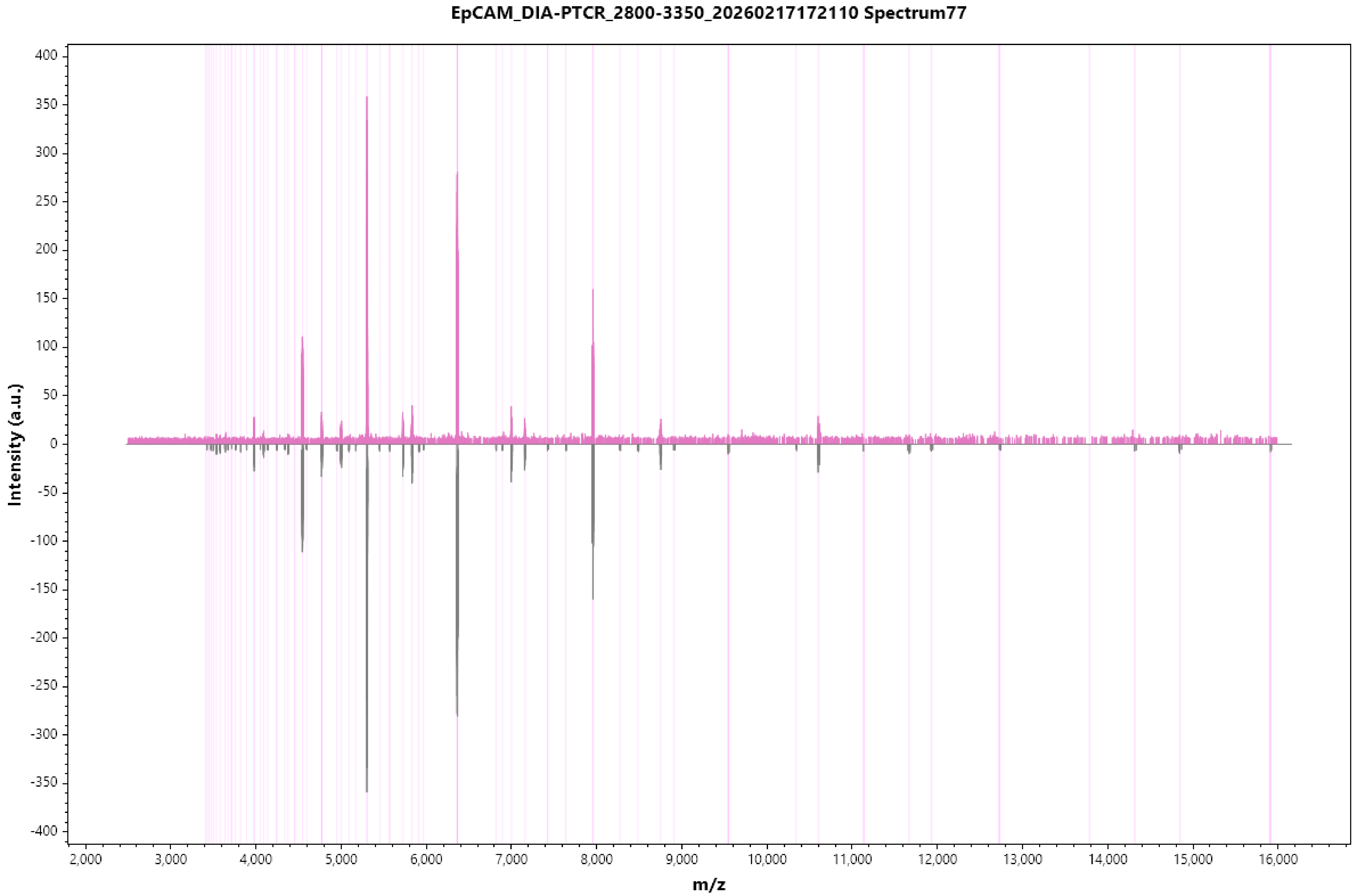
EpCAM_DIA-PTCR_2800-3350_20260217172110 Spectrum77
Intensity (a.u.)
m/z
400
350
300
250
200
150
100
50
0
-50
-100
-150
-200
-250
-300
-350
-400
2,000
3,000
4,000
5,000
6,000
7,000
8,000
9,000
10,000
11,000
12,000
13,000
14,000
15,000
16,000

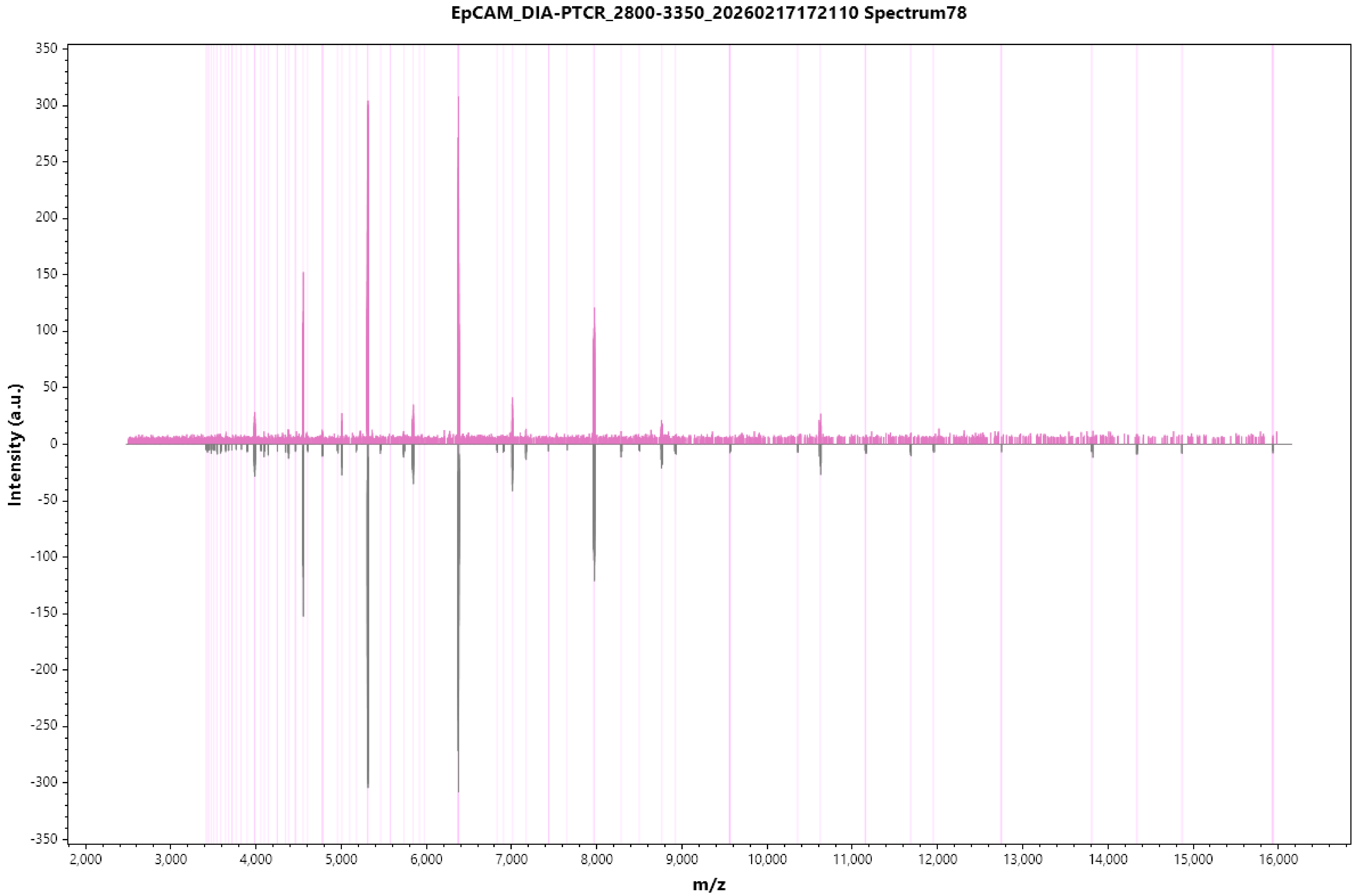
EpCAM_DIA-PTCR_2800-3350_20260217172110 Spectrum78
Intensity (a.u.)
350
300
250
200
150
100
50
0
-50
-100
-150
-200
-250
-300
-350
2,000
3,000
4,000
5,000
6,000
7,000
8,000
9,000
10,000
11,000
12,000
13,000
14,000
15,000
16,000
m/z

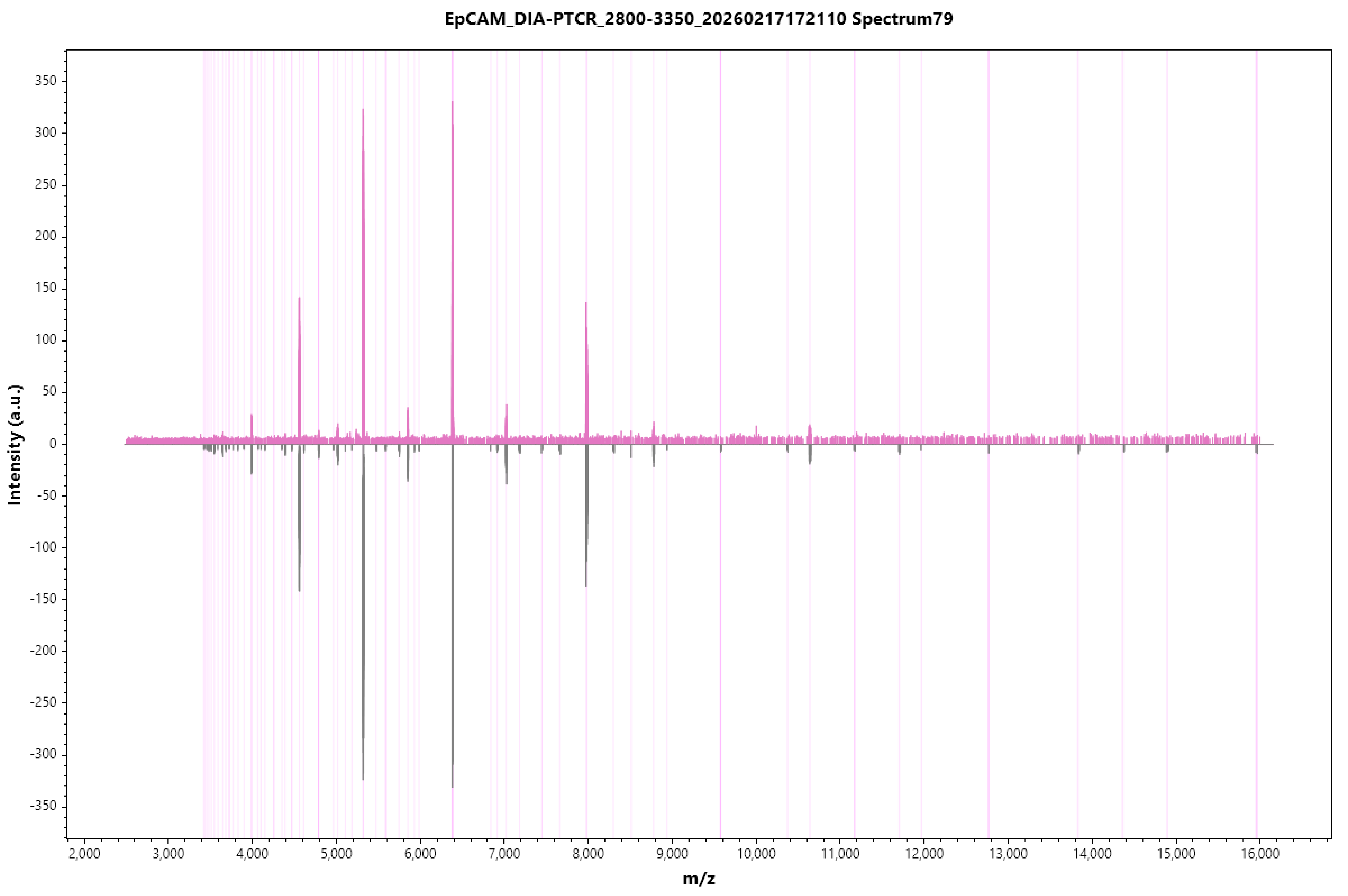
EpCAM_DIA-PTCR_2800-3350_20260217172110 Spectrum79
Intensity (a.u.)
m/z
350
300
250
200
150
100
50
0
-50
-100
-150
-200
-250
-300
-350
2,000
3,000
4,000
5,000
6,000
7,000
8,000
9,000
10,000
11,000
12,000
13,000
14,000
15,000
16,000

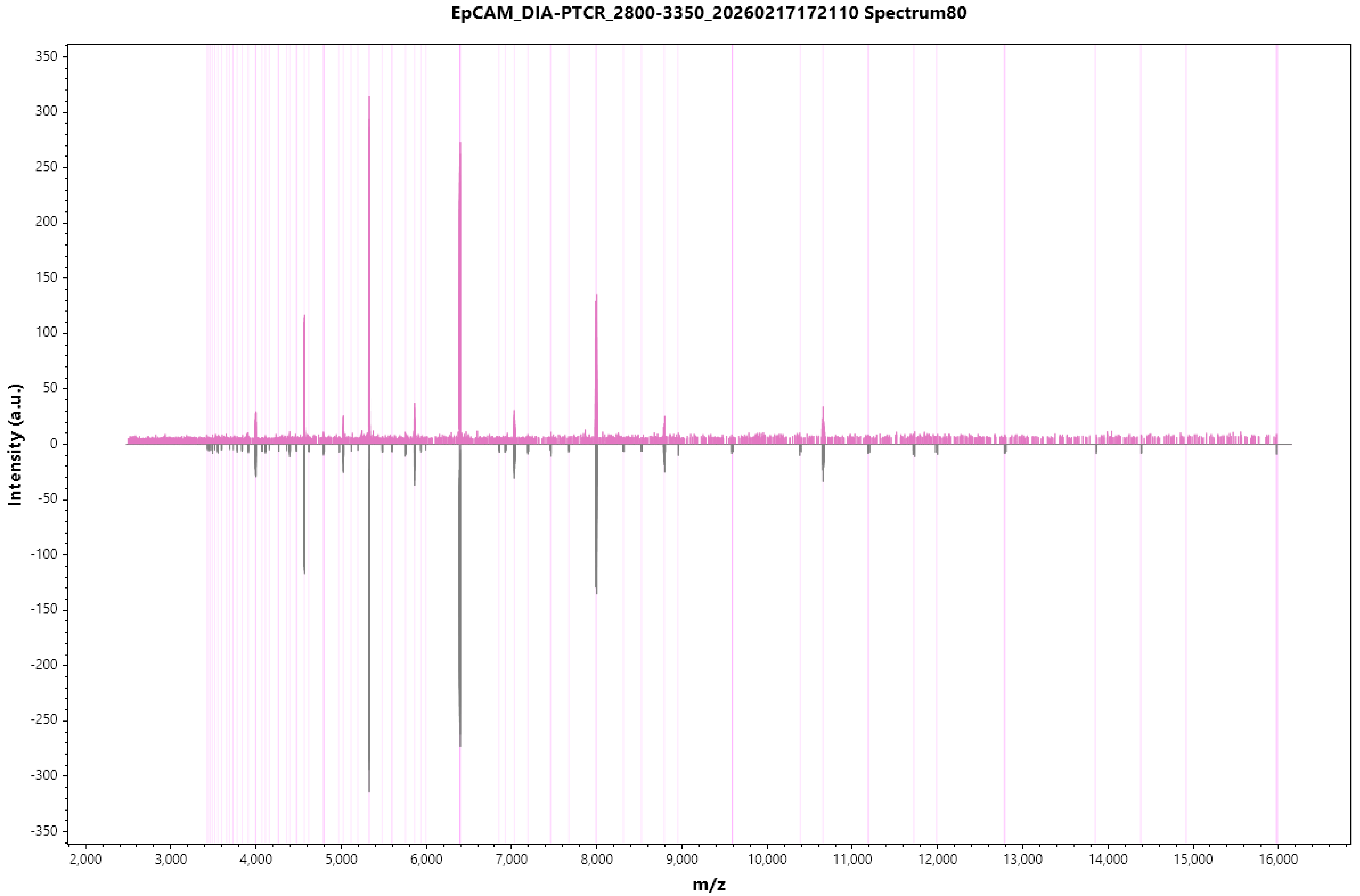
EpCAM_DIA-PTCR_2800-3350_20260217172110 Spectrum80
Intensity (a.u.)
m/z
350
300
250
200
150
100
50
0
-50
-100
-150
-200
-250
-300
-350
2,000
3,000
4,000
5,000
6,000
7,000
8,000
9,000
10,000
11,000
12,000
13,000
14,000
15,000
16,000

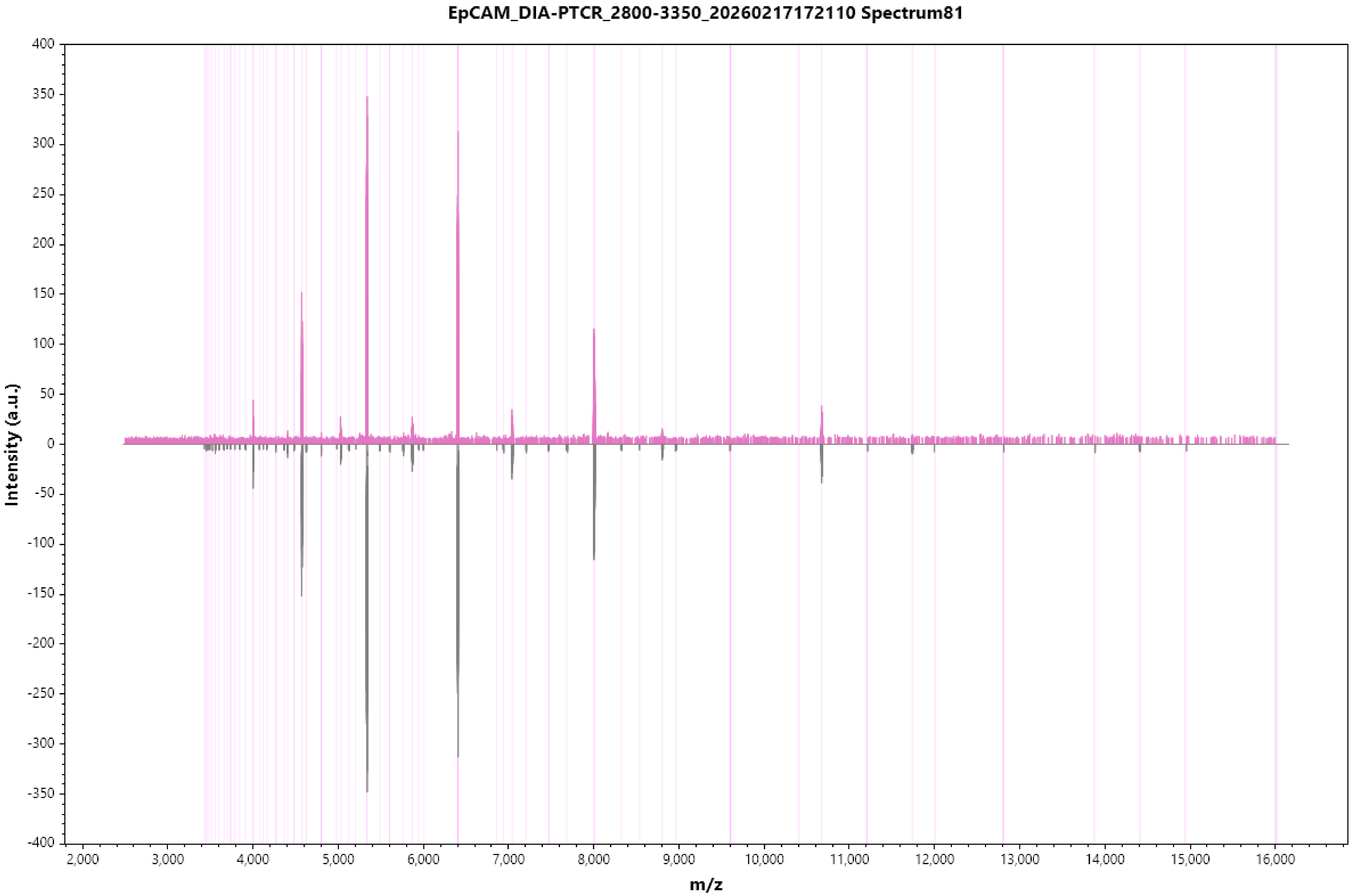
EpCAM_DIA-PTCR_2800-3350_20260217172110 Spectrum81
Intensity (a.u.)
m/z
400
350
300
250
200
150
100
50
0
-50
-100
-150
-200
-250
-300
-350
-400
2,000
3,000
4,000
5,000
6,000
7,000
8,000
9,000
10,000
11,000
12,000
13,000
14,000
15,000
16,000

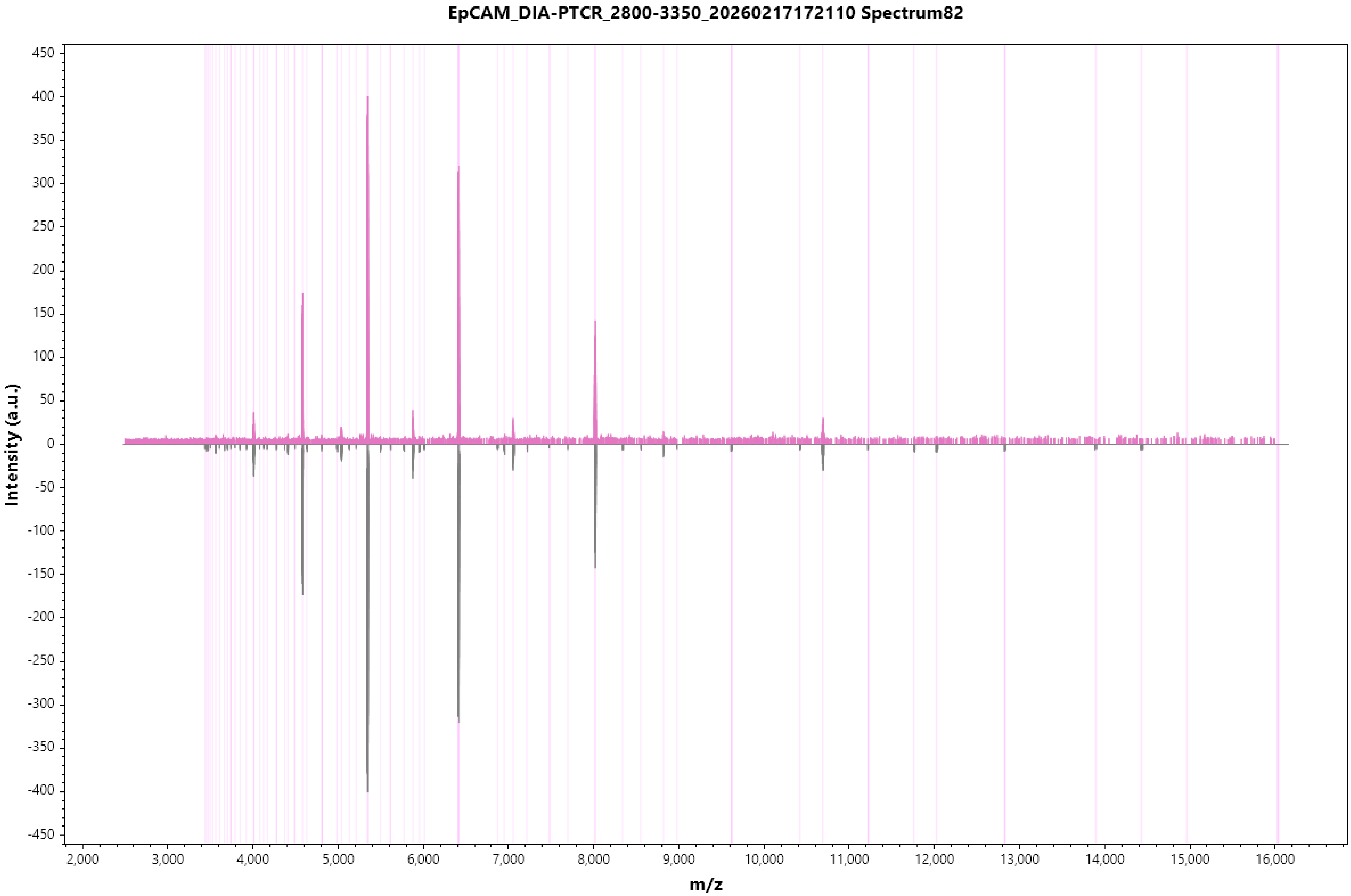

EpCAM_DIA-PTCR_2800-3350_20260217172110 Spectrum82
Intensity (a.u.)
m/z
450
400
350
300
250
200
150
100
50
0
-50
-100
-150
-200
-250
-300
-350
-400
-450
2,000
3,000
4,000
5,000
6,000
7,000
8,000
9,000
10,000
11,000
12,000
13,000
14,000
15,000
16,000

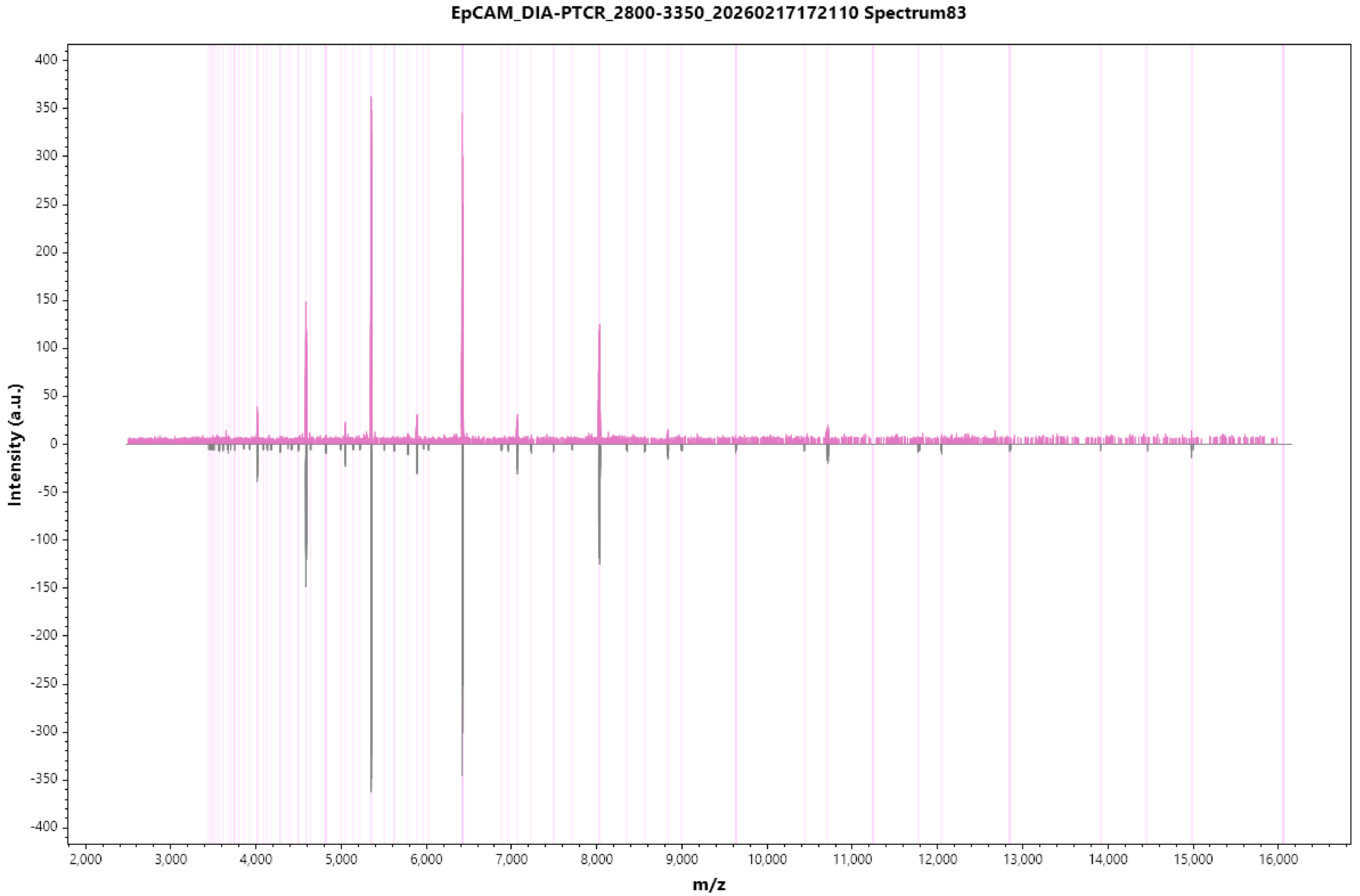
EpCAM_DIA-PTCR_2800-3350_20260217172110 Spectrum83
Intensity (a.u.)
400
350
300
250
200
150
100
50
0
-50
-100
-150
-200
-250
-300
-350
-400
2,000
3,000
4,000
5,000
6,000
7,000
8,000
9,000
10,000
11,000
12,000
13,000
14,000
15,000
16,000
m/z

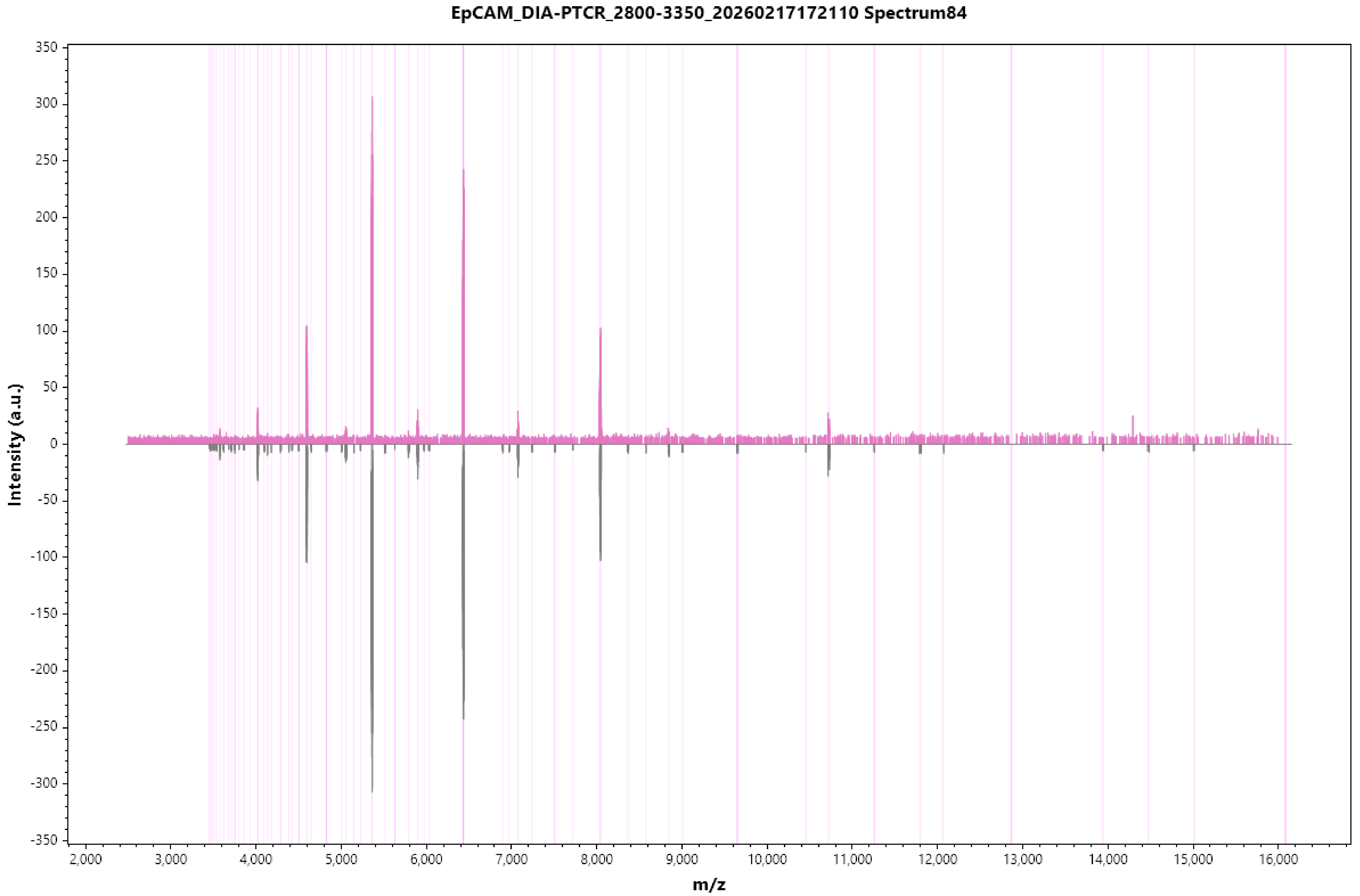
EpCAM_DIA-PTCR_2800-3350_20260217172110 Spectrum84
Intensity (a.u.)
m/z
350
300
250
200
150
100
50
0
-50
-100
-150
-200
-250
-300
-350
2,000
3,000
4,000
5,000
6,000
7,000
8,000
9,000
10,000
11,000
12,000
13,000
14,000
15,000
16,000

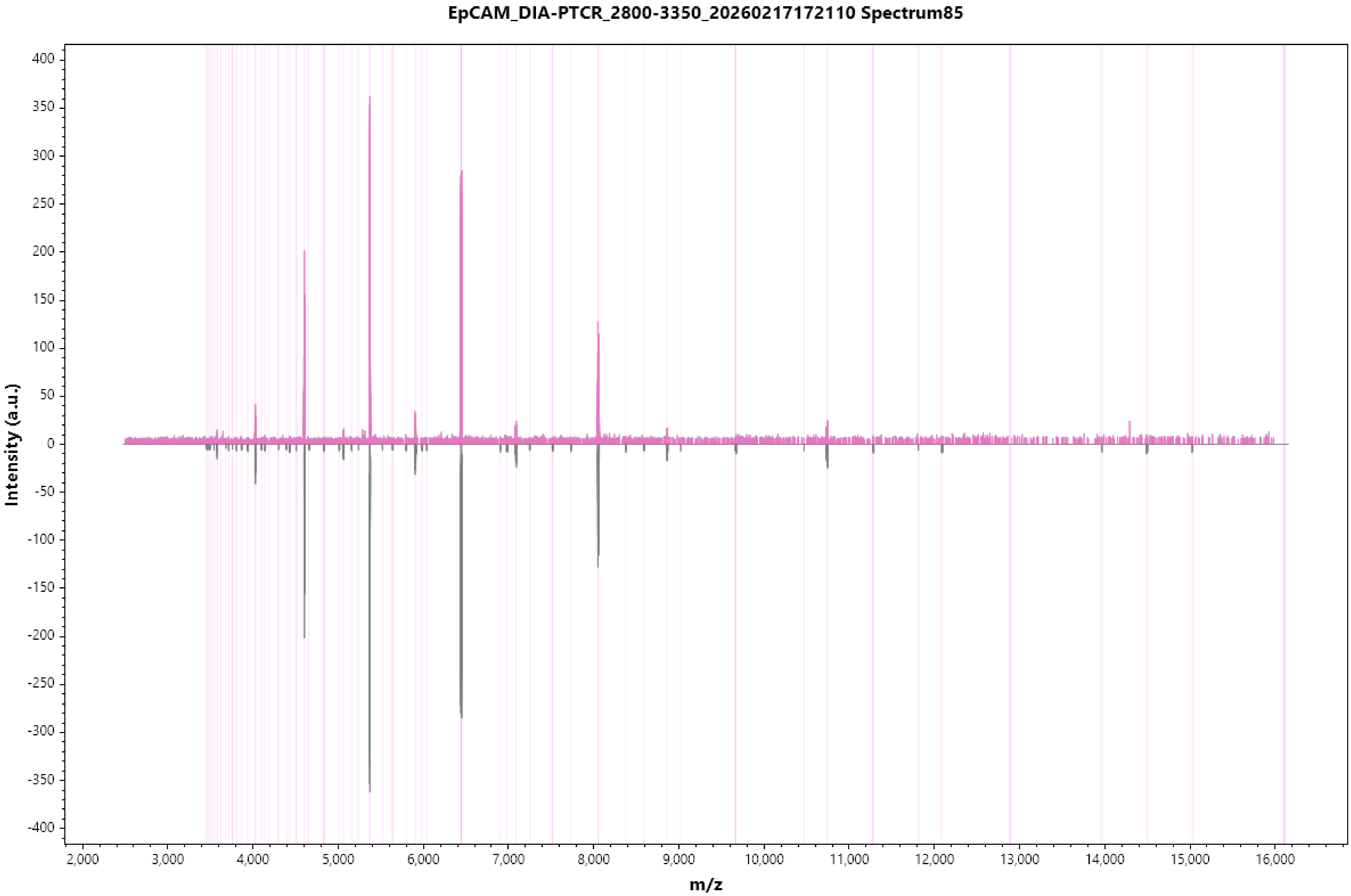
EpCAM_DIA-PTCR_2800-3350_20260217172110 Spectrum85
Intensity (a.u.)
400
350
300
250
200
150
100
50
0
-50
-100
-150
-200
-250
-300
-350
-400
2,000
3,000
4,000
5,000
6,000
7,000
8,000
9,000
10,000
11,000
12,000
13,000
14,000
15,000
16,000
m/z

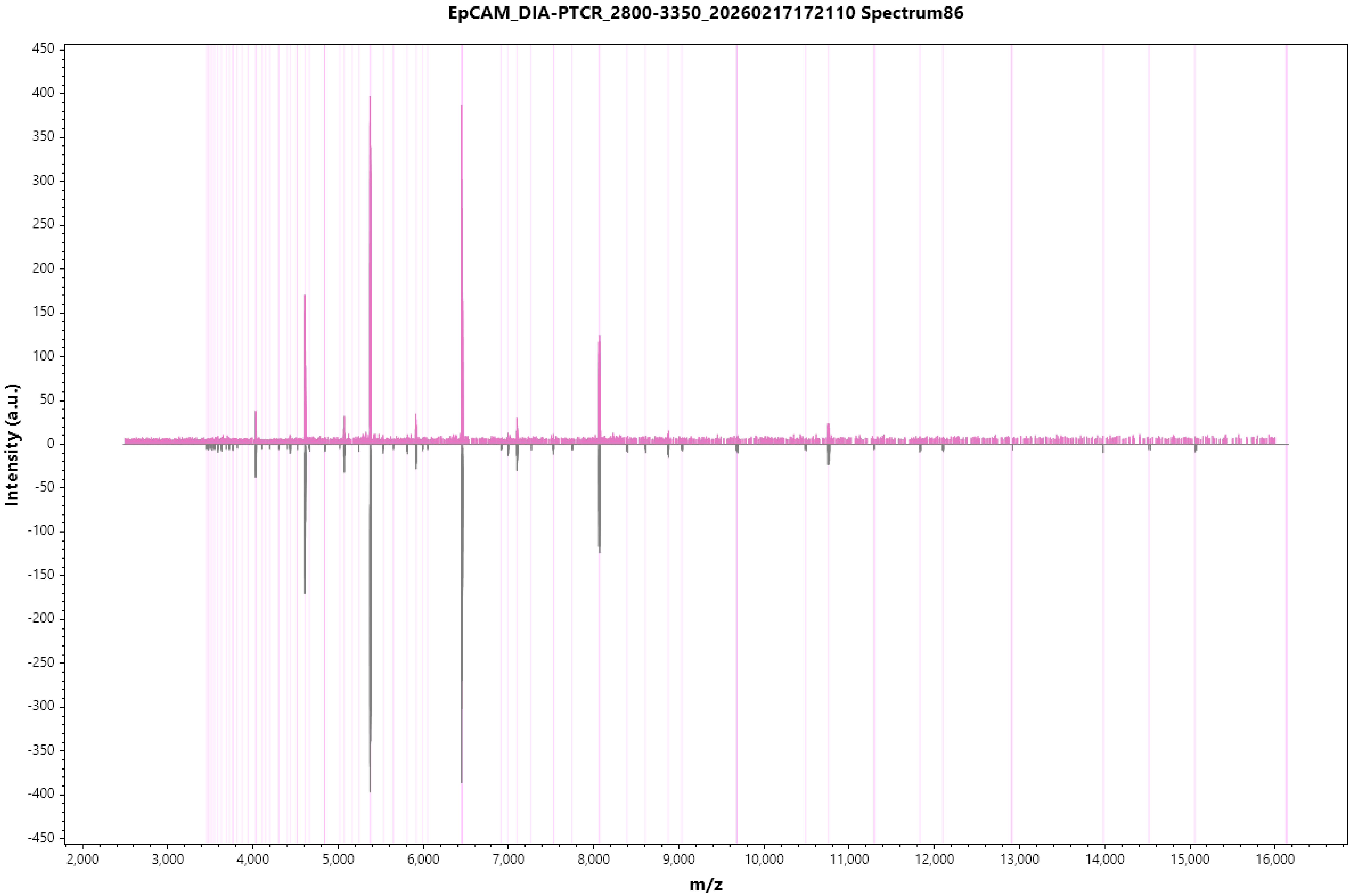
EpCAM_DIA-PTCR_2800-3350_20260217172110 Spectrum86
Intensity (a.u.)
450
400
350
300
250
200
150
100
50
0
-50
-100
-150
-200
-250
-300
-350
-400
-450
2,000
3,000
4,000
5,000
6,000
7,000
8,000
9,000
10,000
11,000
12,000
13,000
14,000
15,000
16,000
m/z

EpCAM_DIA-PTCR_2800-3350_20260217172110 Spectrum87

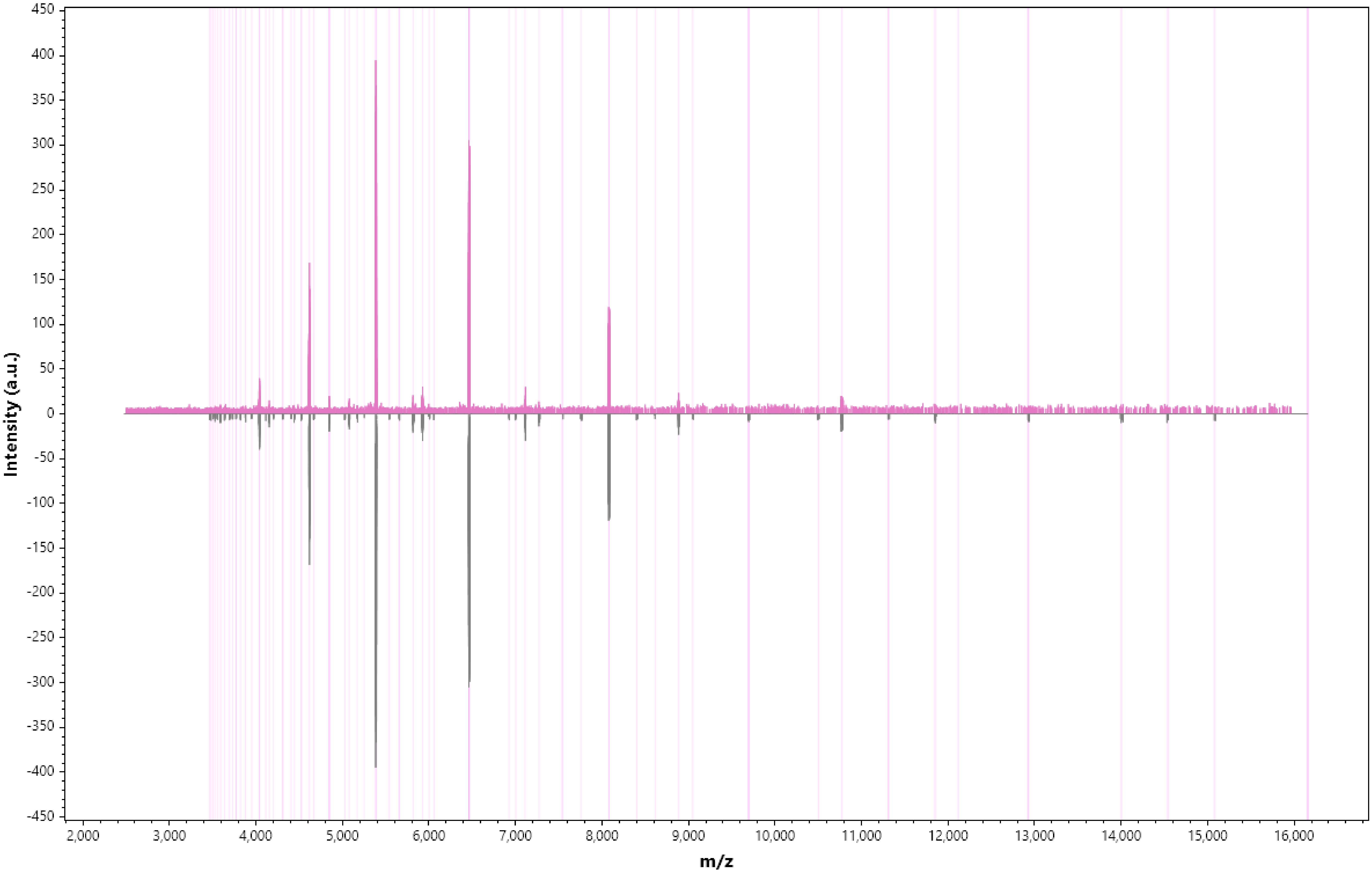

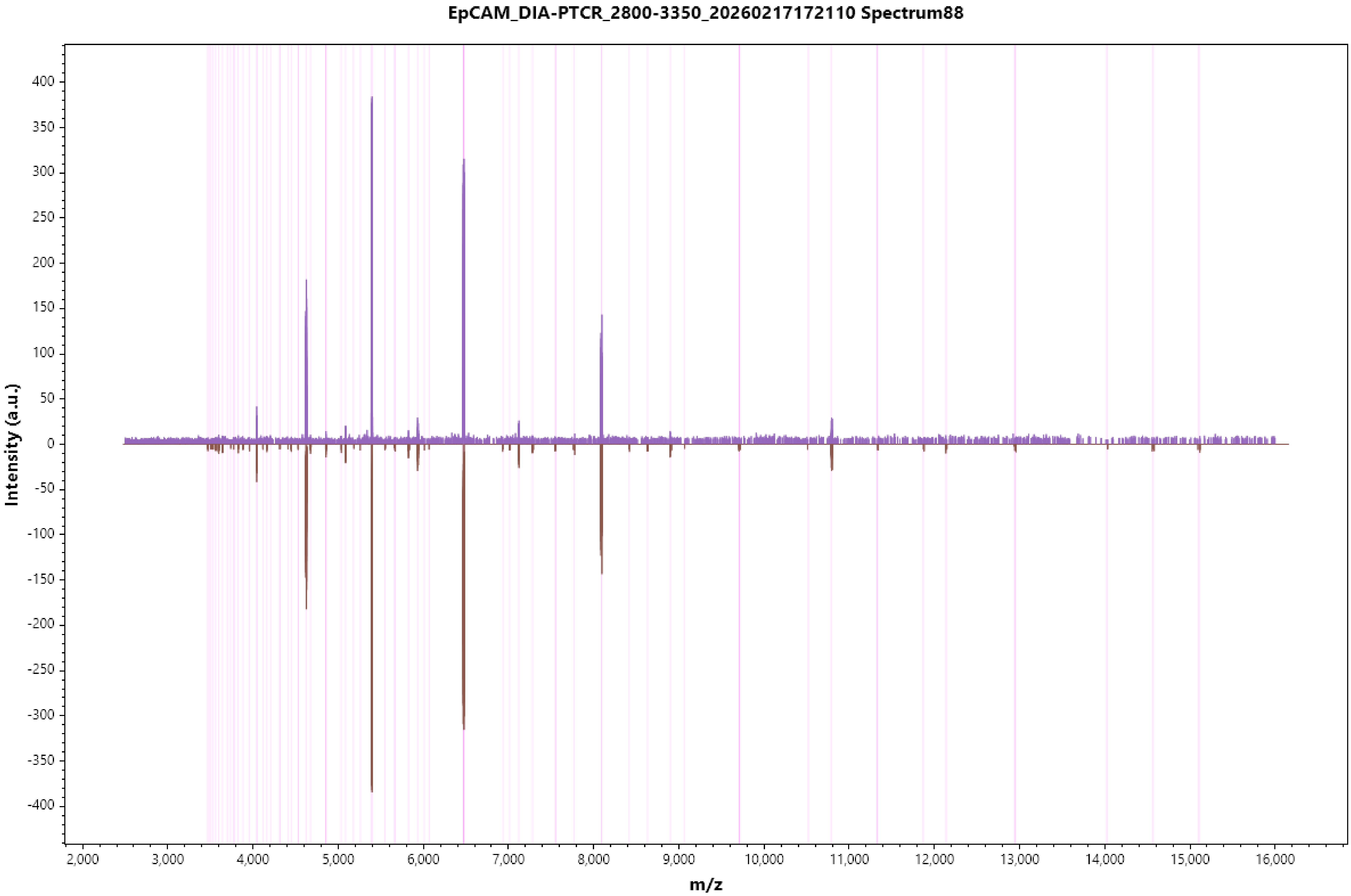

EpCAM_DIA-PTCR_2800-3350_20260217172110 Spectrum88
Intensity (a.u.)
400
350
300
250
200
150
100
50
0
-50
-100
-150
-200
-250
-300
-350
-400
2,000
3,000
4,000
5,000
6,000
7,000
8,000
9,000
10,000
11,000
12,000
13,000
14,000
15,000
16,000
m/z

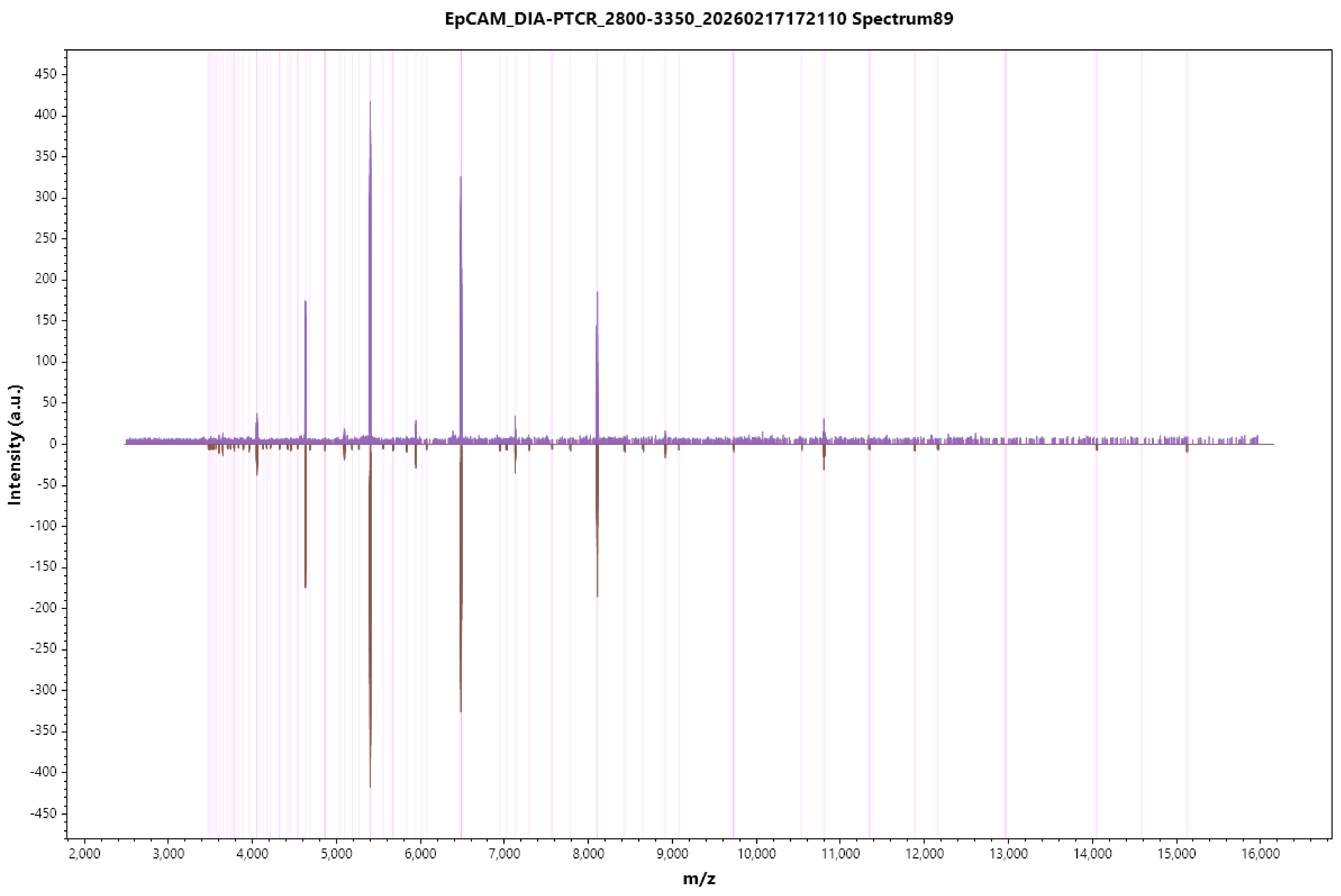
EpCAM_DIA-PTCR_2800-3350_20260217172110 Spectrum89
Intensity (a.u.)
450
400
350
300
250
200
150
100
50
0
-50
-100
-150
-200
-250
-300
-350
-400
-450
2,000
3,000
4,000
5,000
6,000
7,000
8,000
9,000
10,000
11,000
12,000
13,000
14,000
15,000
16,000
m/z

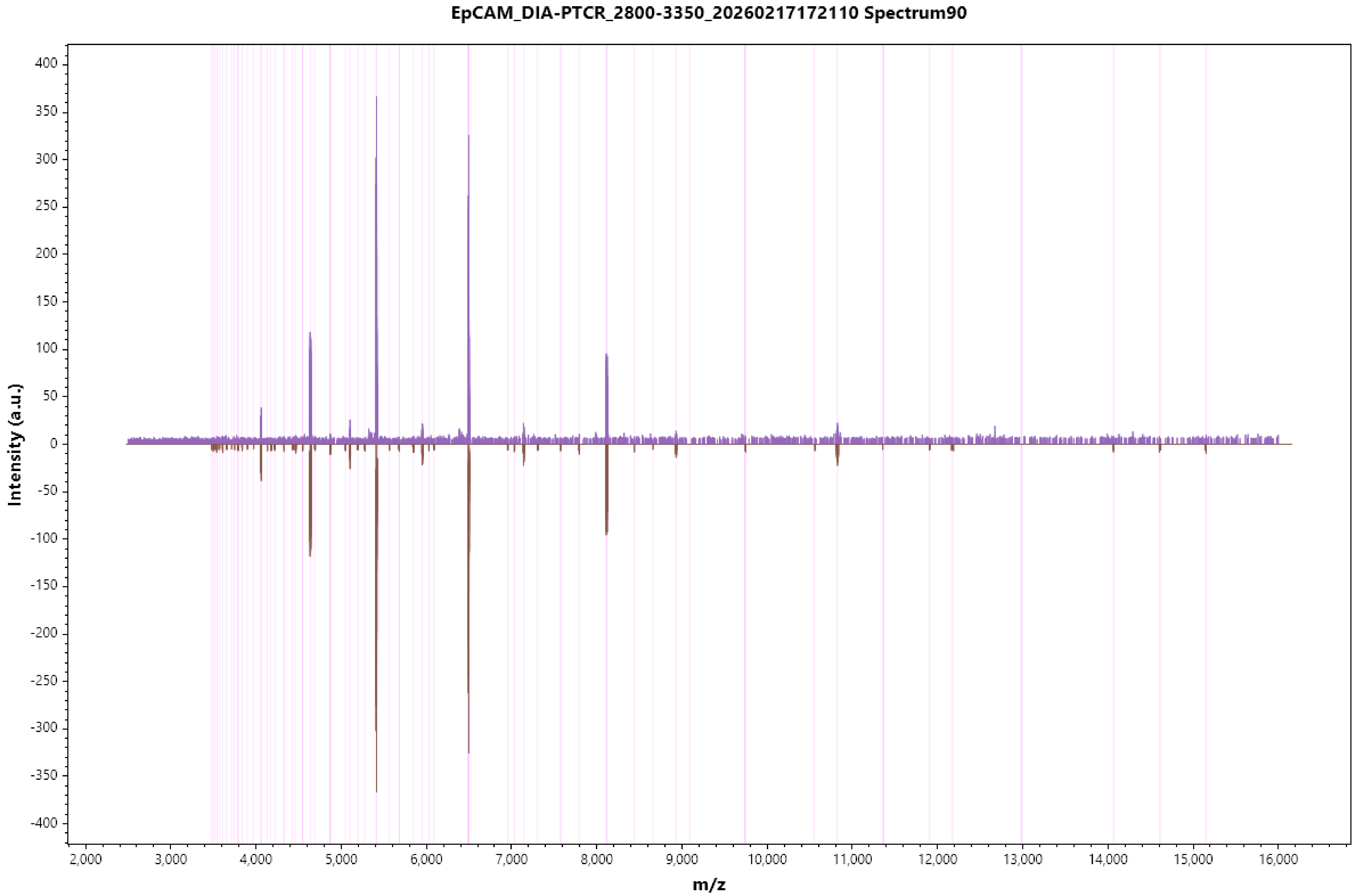
EpCAM_DIA-PTCR_2800-3350_20260217172110 Spectrum90
Intensity (a.u.)
m/z
400
350
300
250
200
150
100
50
0
-50
-100
-150
-200
-250
-300
-350
-400
2,000
3,000
4,000
5,000
6,000
7,000
8,000
9,000
10,000
11,000
12,000
13,000
14,000
15,000
16,000

EpCAM_DIA-PTCR_2800-3350_20260217172110 Spectrum91

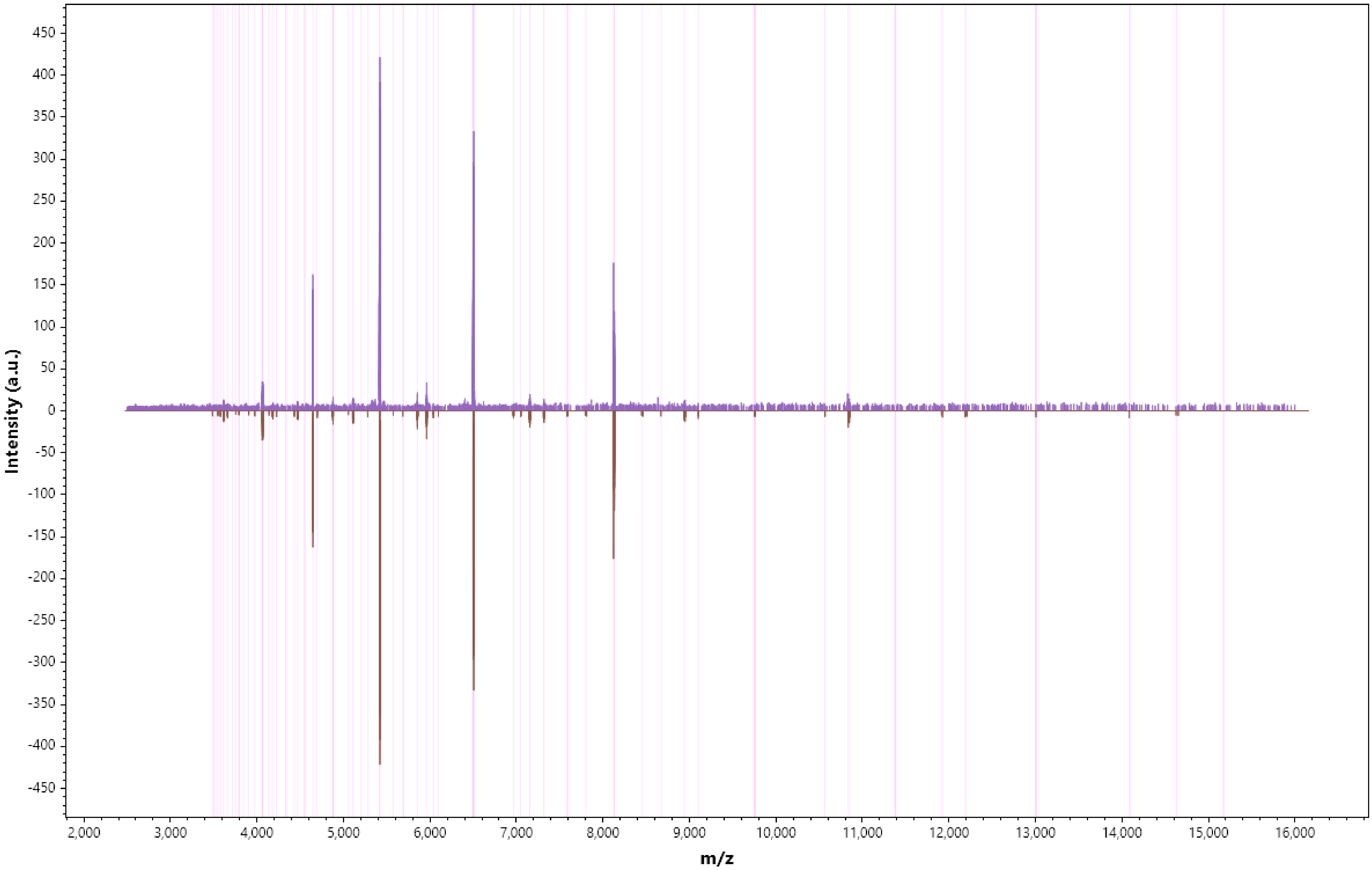

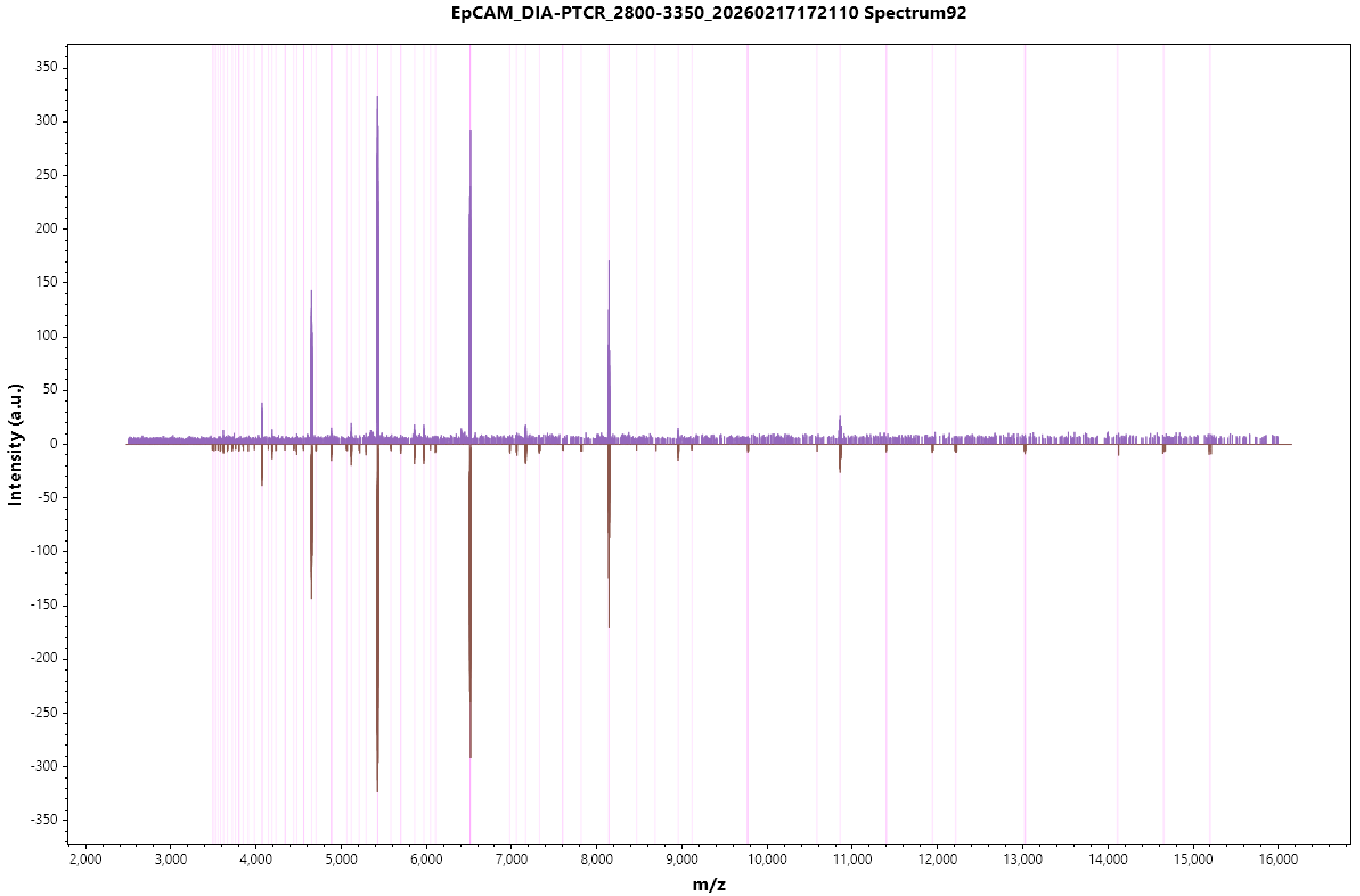
EpCAM_DIA-PTCR_2800-3350_20260217172110 Spectrum92
Intensity (a.u.)
350
300
250
200
150
100
50
0
-50
-100
-150
-200
-250
-300
-350
2,000
3,000
4,000
5,000
6,000
7,000
8,000
9,000
10,000
11,000
12,000
13,000
14,000
15,000
16,000
m/z

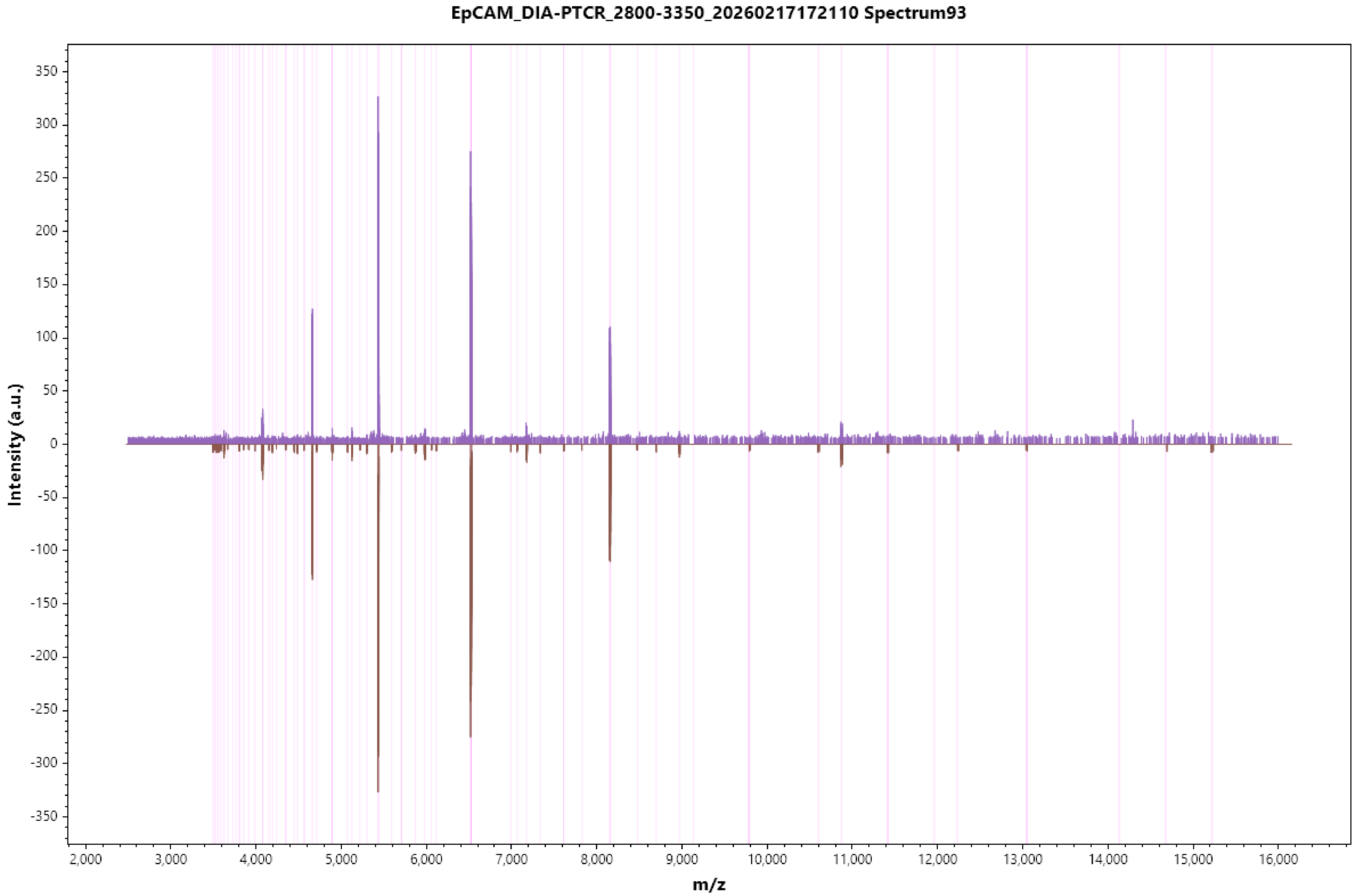

EpCAM_DIA-PTCR_2800-3350_20260217172110 Spectrum93
Intensity (a.u.)
m/z
350
300
250
200
150
100
50
0
-50
-100
-150
-200
-250
-300
-350
2,000
3,000
4,000
5,000
6,000
7,000
8,000
9,000
10,000
11,000
12,000
13,000
14,000
15,000
16,000

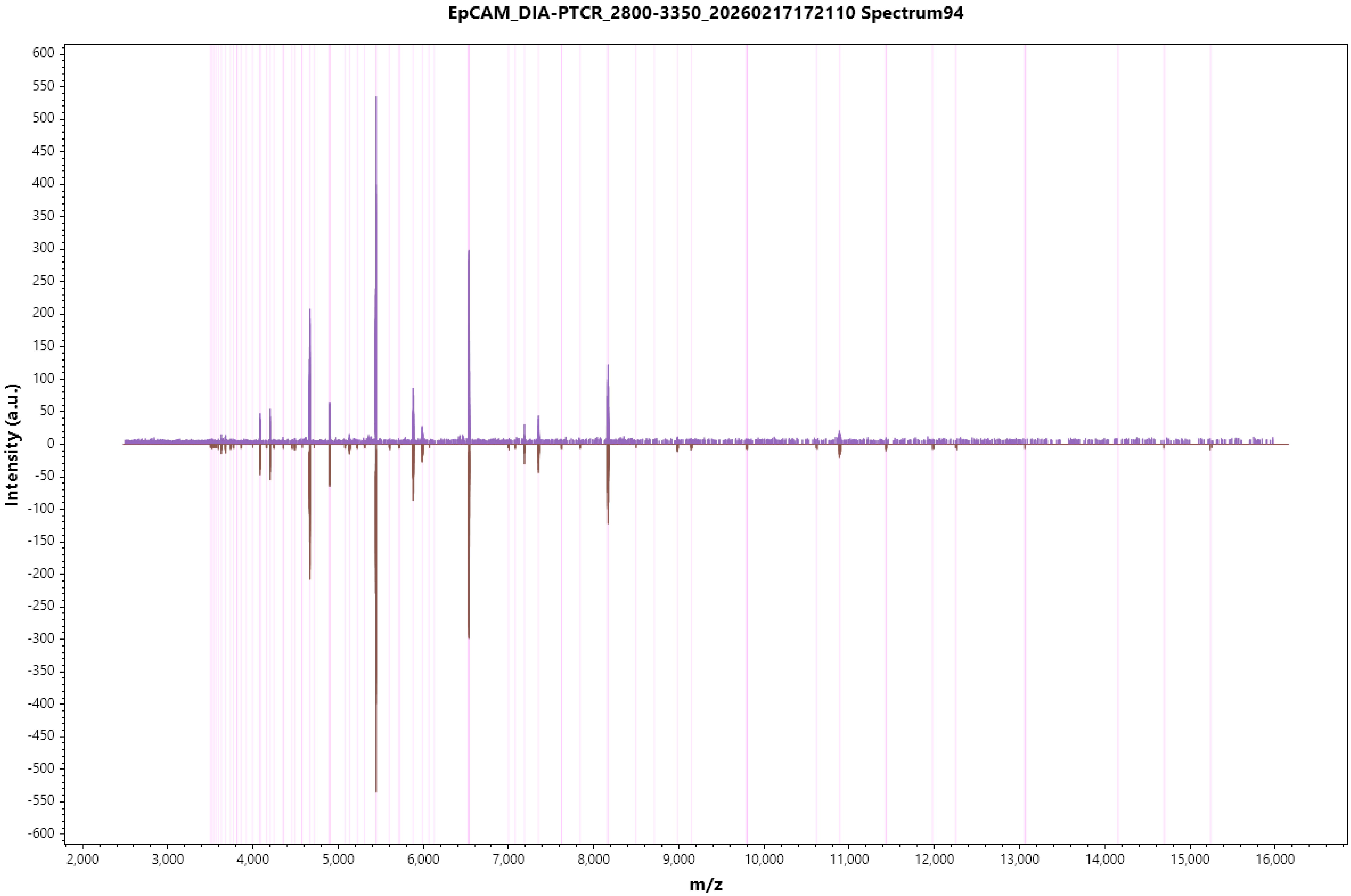

EpCAM_DIA-PTCR_2800-3350_20260217172110 Spectrum94
Intensity (a.u.)
m/z
600
550
500
450
400
350
300
250
200
150
100
50
0
-50
-100
-150
-200
-250
-300
-350
-400
-450
-500
-550
-600
2,000
3,000
4,000
5,000
6,000
7,000
8,000
9,000
10,000
11,000
12,000
13,000
14,000
15,000
16,000

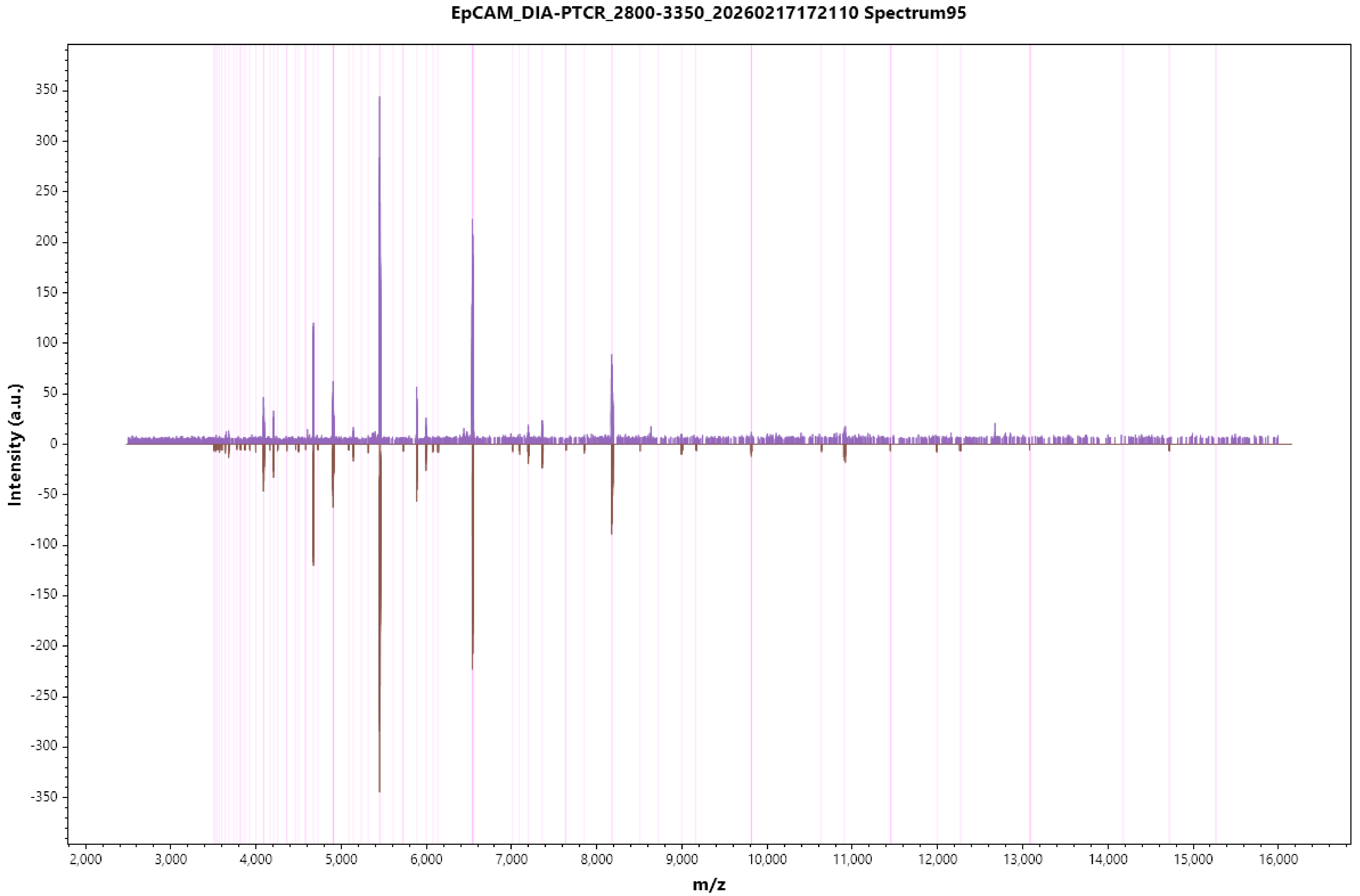

EpCAM_DIA-PTCR_2800-3350_20260217172110 Spectrum95
Intensity (a.u.)
m/z
350
300
250
200
150
100
50
0
-50
-100
-150
-200
-250
-300
-350
2,000
3,000
4,000
5,000
6,000
7,000
8,000
9,000
10,000
11,000
12,000
13,000
14,000
15,000
16,000

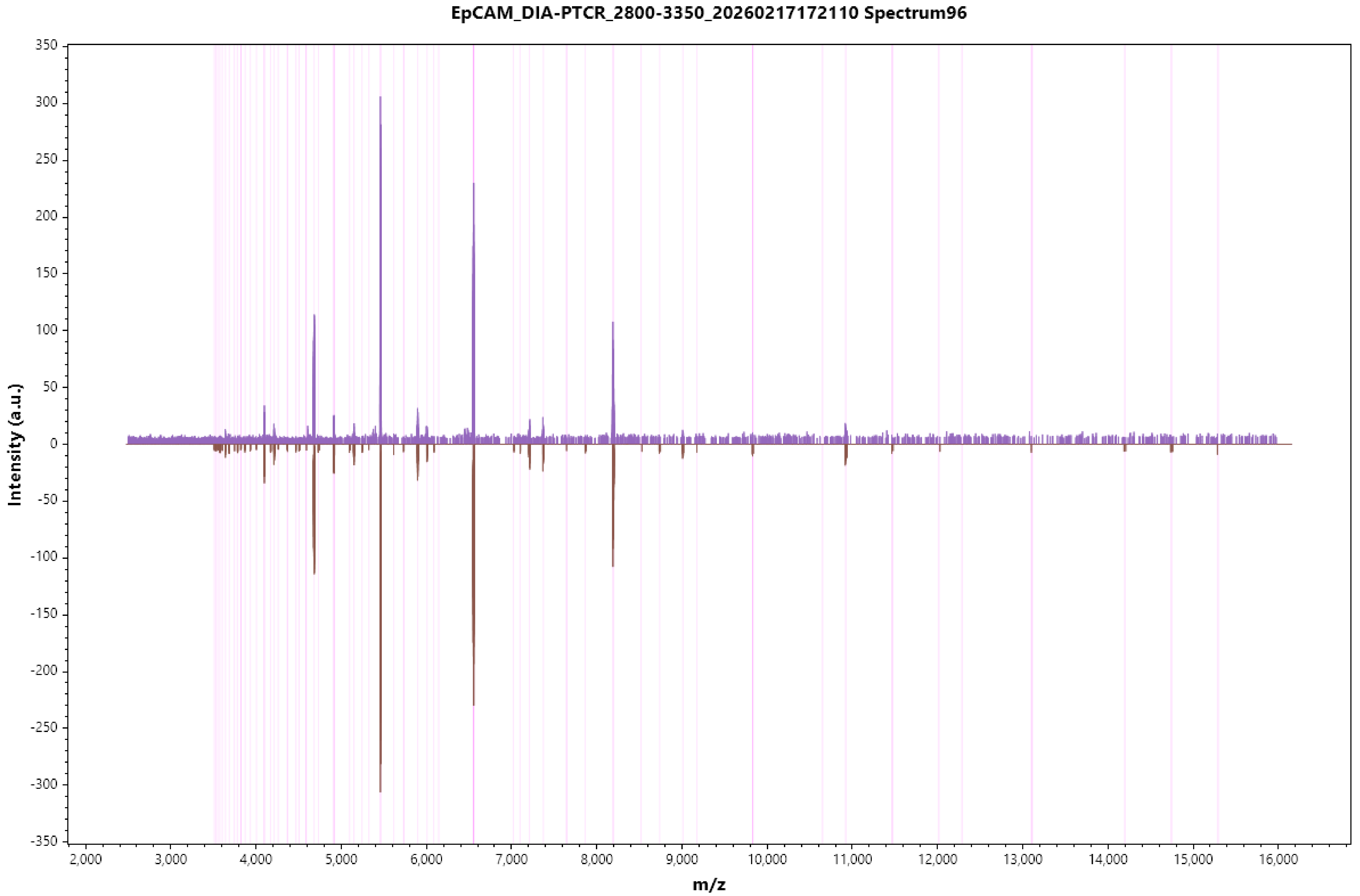

EpCAM_DIA-PTCR_2800-3350_20260217172110 Spectrum96
Intensity (a.u.)
350
300
250
200
150
100
50
0
-50
-100
-150
-200
-250
-300
-350
2,000
3,000
4,000
5,000
6,000
7,000
8,000
9,000
10,000
11,000
12,000
13,000
14,000
15,000
16,000
m/z

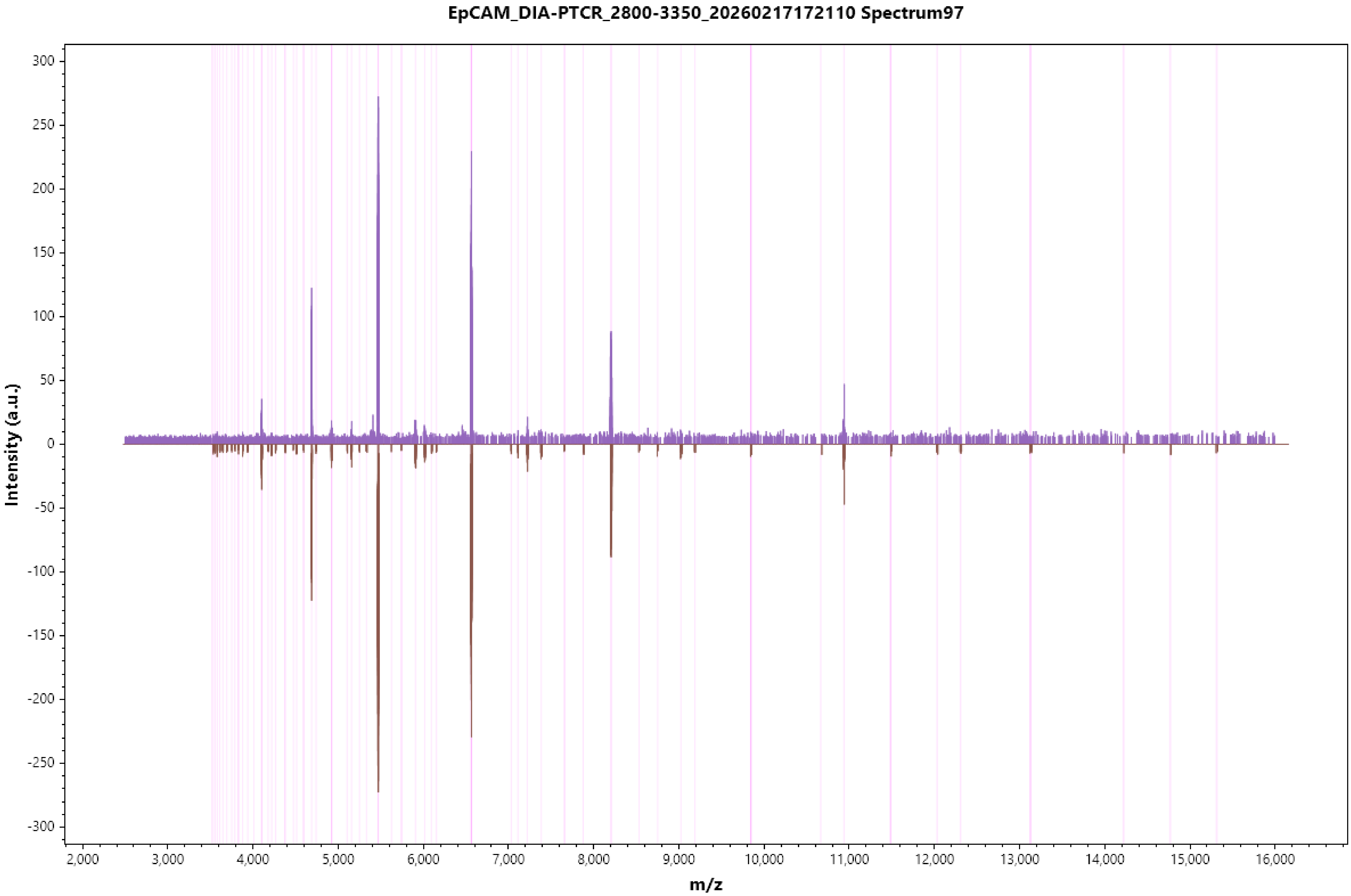

EpCAM_DIA-PTCR_2800-3350_20260217172110 Spectrum97
Intensity (a.u.)
m/z
300
250
200
150
100
50
0
-50
-100
-150
-200
-250
-300
2,000
3,000
4,000
5,000
6,000
7,000
8,000
9,000
10,000
11,000
12,000
13,000
14,000
15,000
16,000

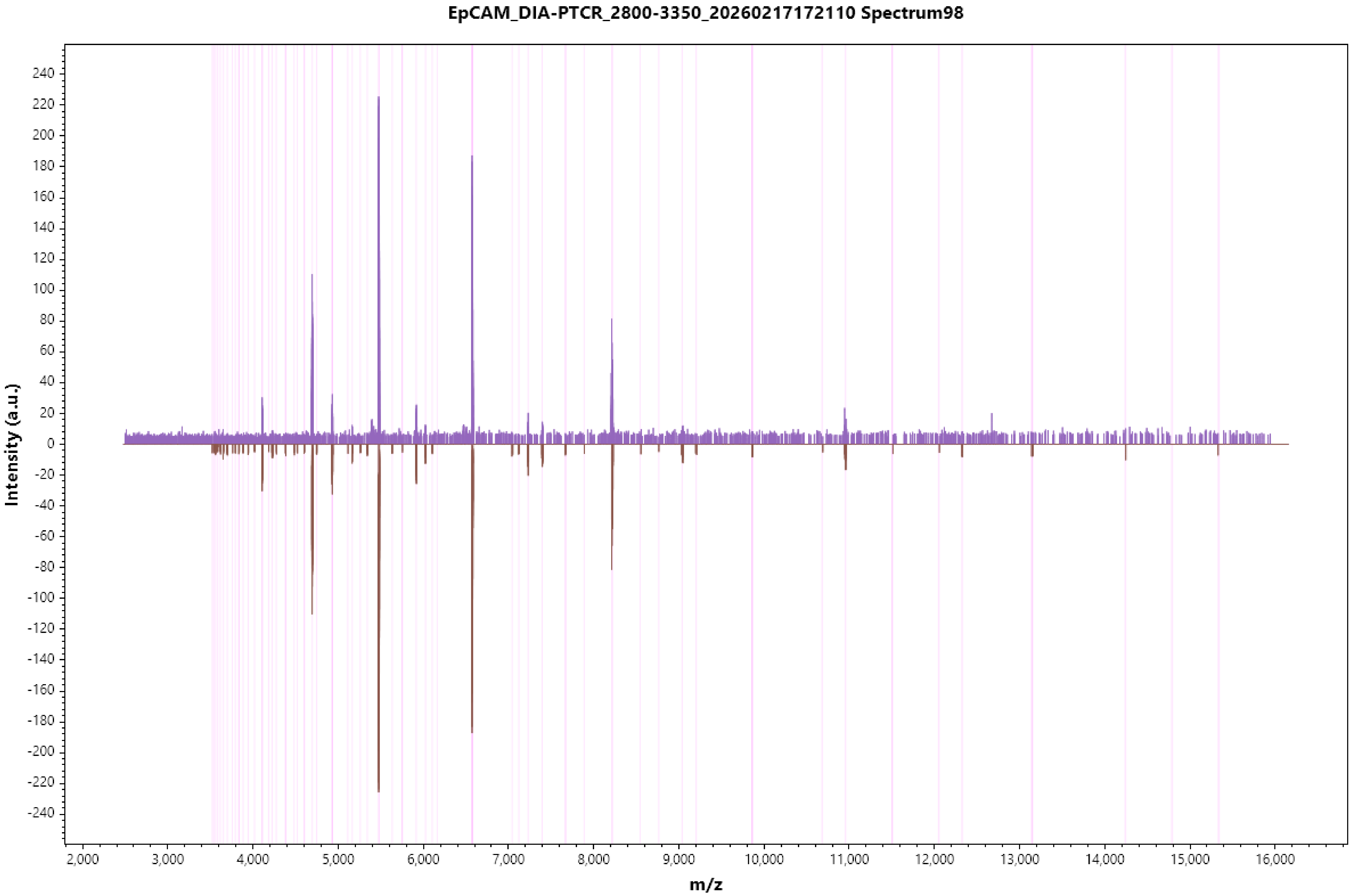
EpCAM_DIA-PTCR_2800-3350_20260217172110 Spectrum98
Intensity (a.u.)
m/z
240
220
200
180
160
140
120
100
80
60
40
20
0
-20
-40
-60
-80
-100
-120
-140
-160
-180
-200
-220
-240
2,000
3,000
4,000
5,000
6,000
7,000
8,000
9,000
10,000
11,000
12,000
13,000
14,000
15,000
16,000

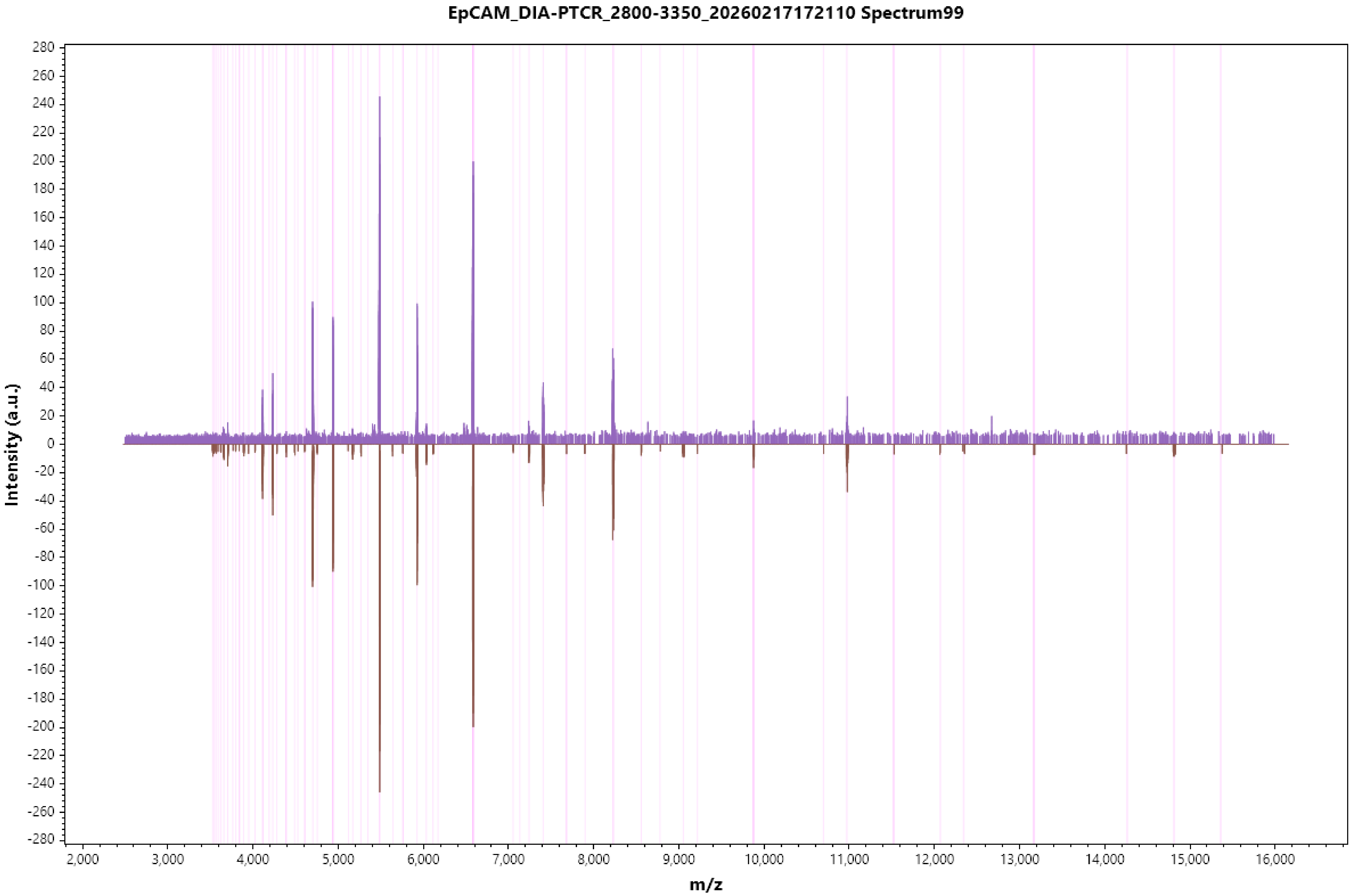
EpCAM_DIA-PTCR_2800-3350_20260217172110 Spectrum99
Intensity (a.u.)
m/z
280
260
240
220
200
180
160
140
120
100
80
60
40
20
0
-20
-40
-60
-80
-100
-120
-140
-160
-180
-200
-220
-240
-260
-280
2,000
3,000
4,000
5,000
6,000
7,000
8,000
9,000
10,000
11,000
12,000
13,000
14,000
15,000
16,000

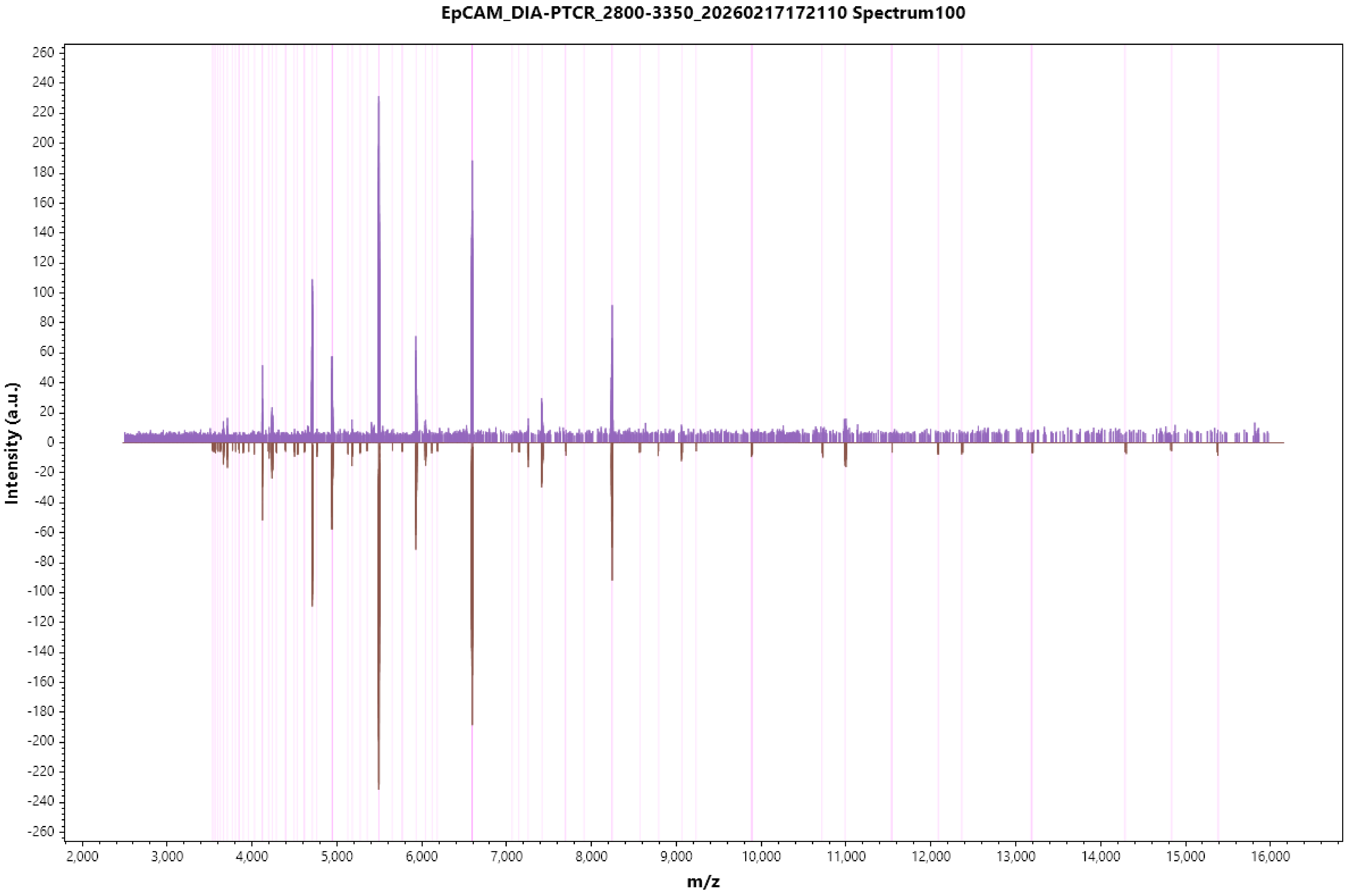
EpCAM_DIA-PTCR_2800-3350_20260217172110 Spectrum100
Intensity (a.u.)
m/z

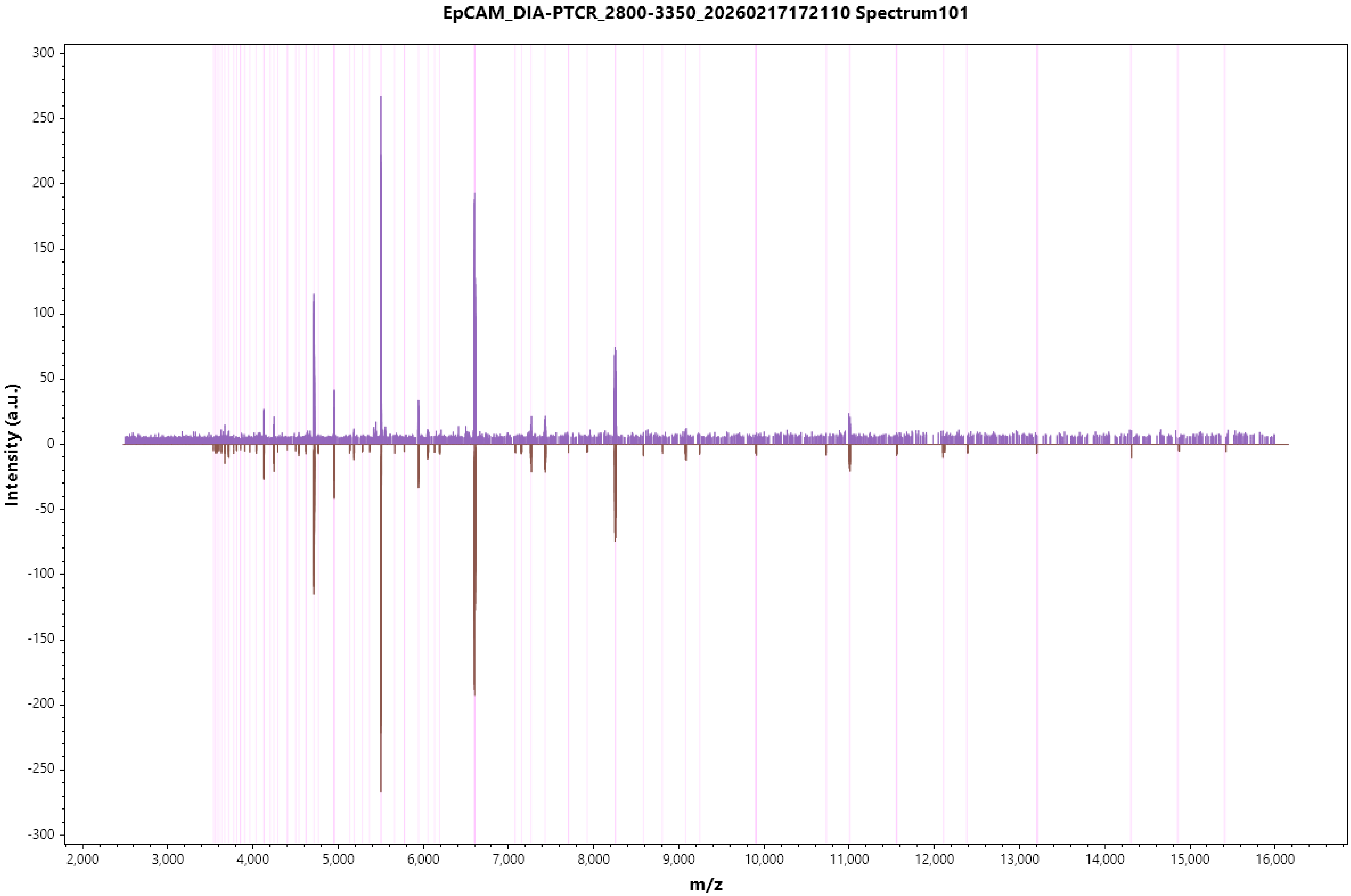
EpCAM_DIA-PTCR_2800-3350_20260217172110 Spectrum101
Intensity (a.u.)
m/z
300
250
200
150
100
50
0
-50
-100
-150
-200
-250
-300
2,000
3,000
4,000
5,000
6,000
7,000
8,000
9,000
10,000
11,000
12,000
13,000
14,000
15,000
16,000

### EpCAM_DIA-PTCR_2800-3350_20260217172110 Spectrum102

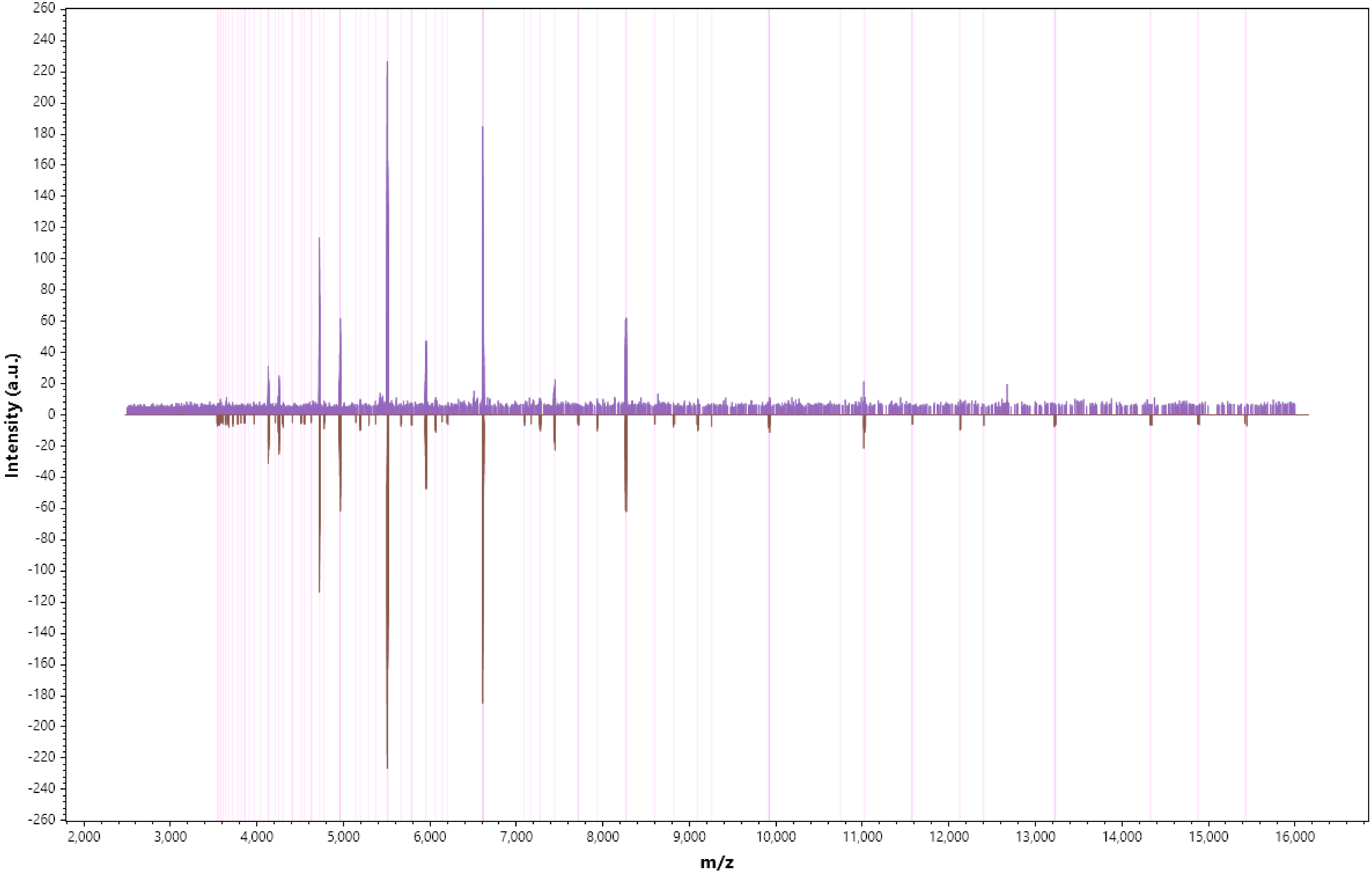

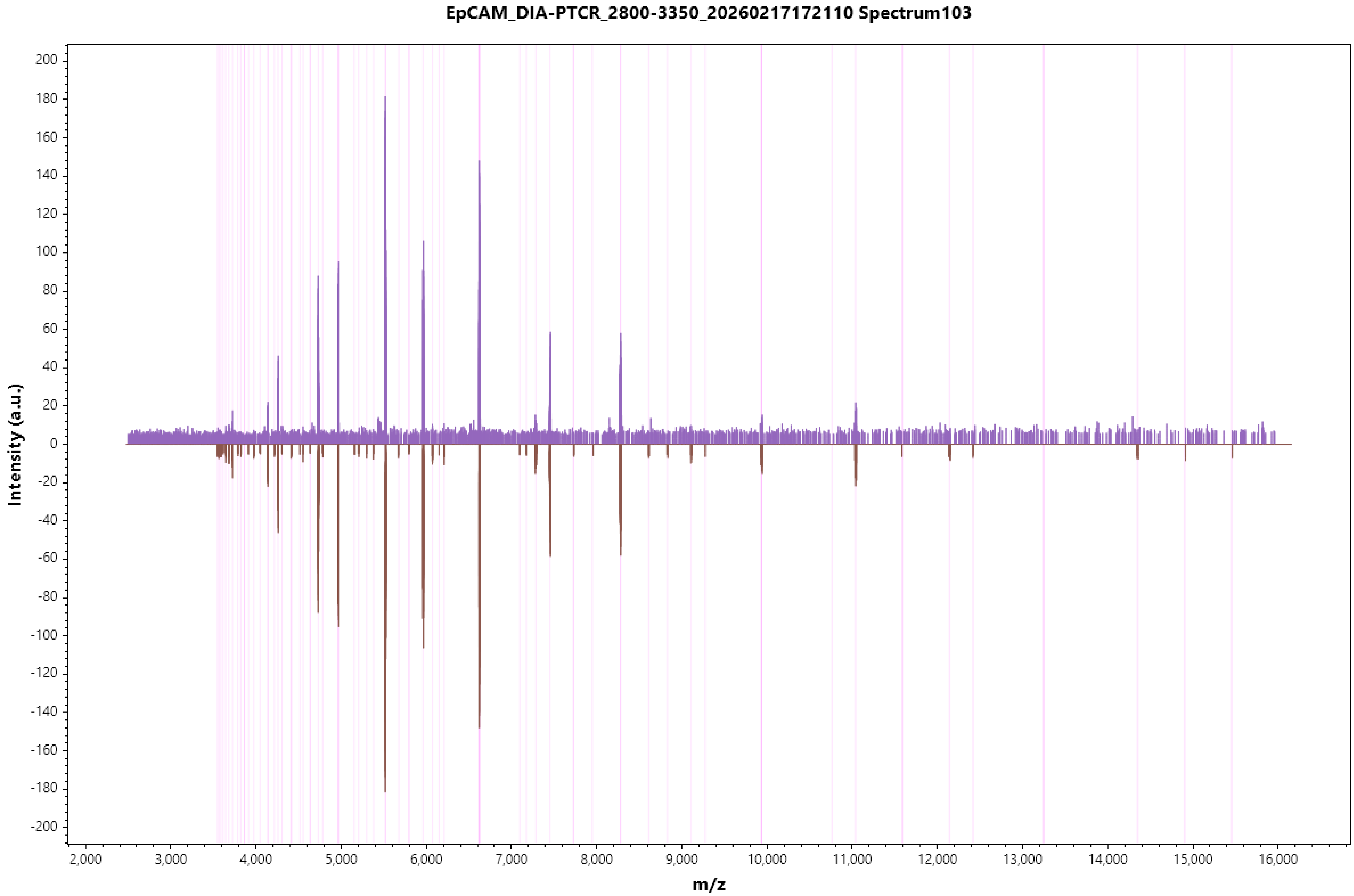
EpCAM_DIA-PTCR_2800-3350_20260217172110 Spectrum103
Intensity (a.u.)
m/z
200
180
160
140
120
100
80
60
40
20
0
-20
-40
-60
-80
-100
-120
-140
-160
-180
-200
2,000
3,000
4,000
5,000
6,000
7,000
8,000
9,000
10,000
11,000
12,000
13,000
14,000
15,000
16,000

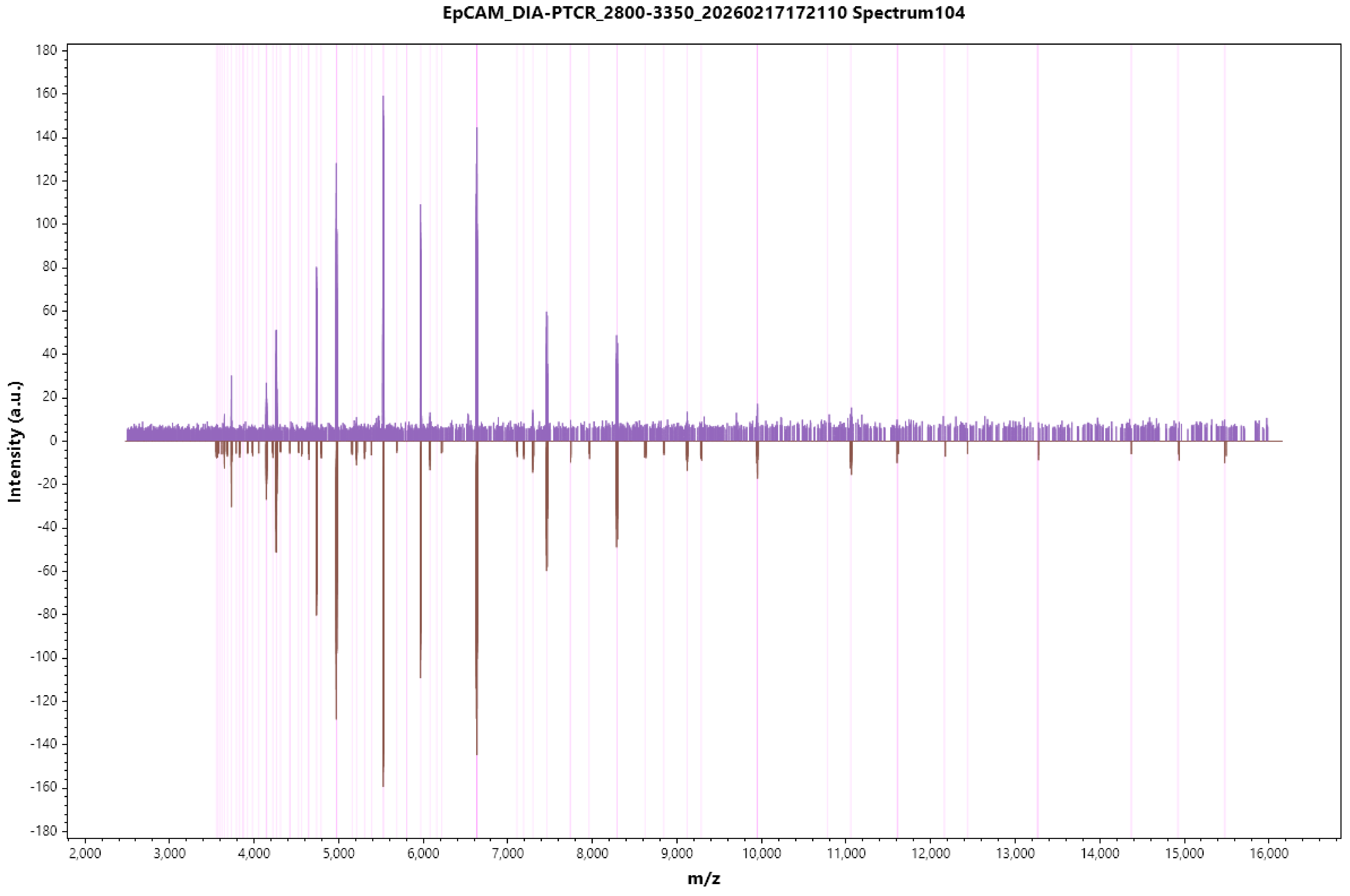
EpCAM_DIA-PTCR_2800-3350_20260217172110 Spectrum104
Intensity (a.u.)
m/z
180
160
140
120
100
80
60
40
20
0
-20
-40
-60
-80
-100
-120
-140
-160
-180
2,000
3,000
4,000
5,000
6,000
7,000
8,000
9,000
10,000
11,000
12,000
13,000
14,000
15,000
16,000

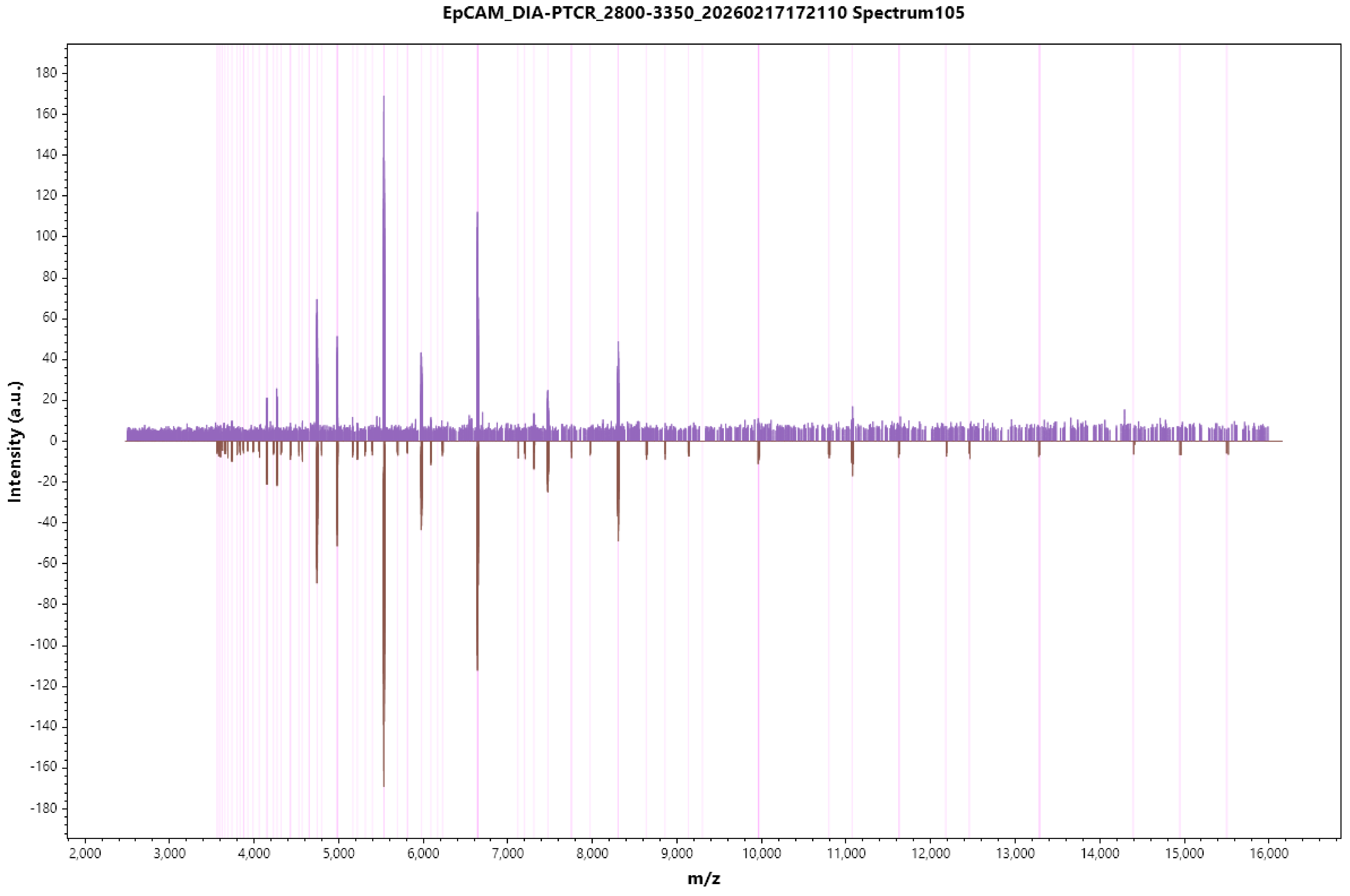

EpCAM_DIA-PTCR_2800-3350_20260217172110 Spectrum105
Intensity (a.u.)
m/z
180
160
140
120
100
80
60
40
20
0
-20
-40
-60
-80
-100
-120
-140
-160
-180
2,000
3,000
4,000
5,000
6,000
7,000
8,000
9,000
10,000
11,000
12,000
13,000
14,000
15,000
16,000

#### EpCAM_DIA-PTCR_2800-3350_20260217172110 Spectrum106

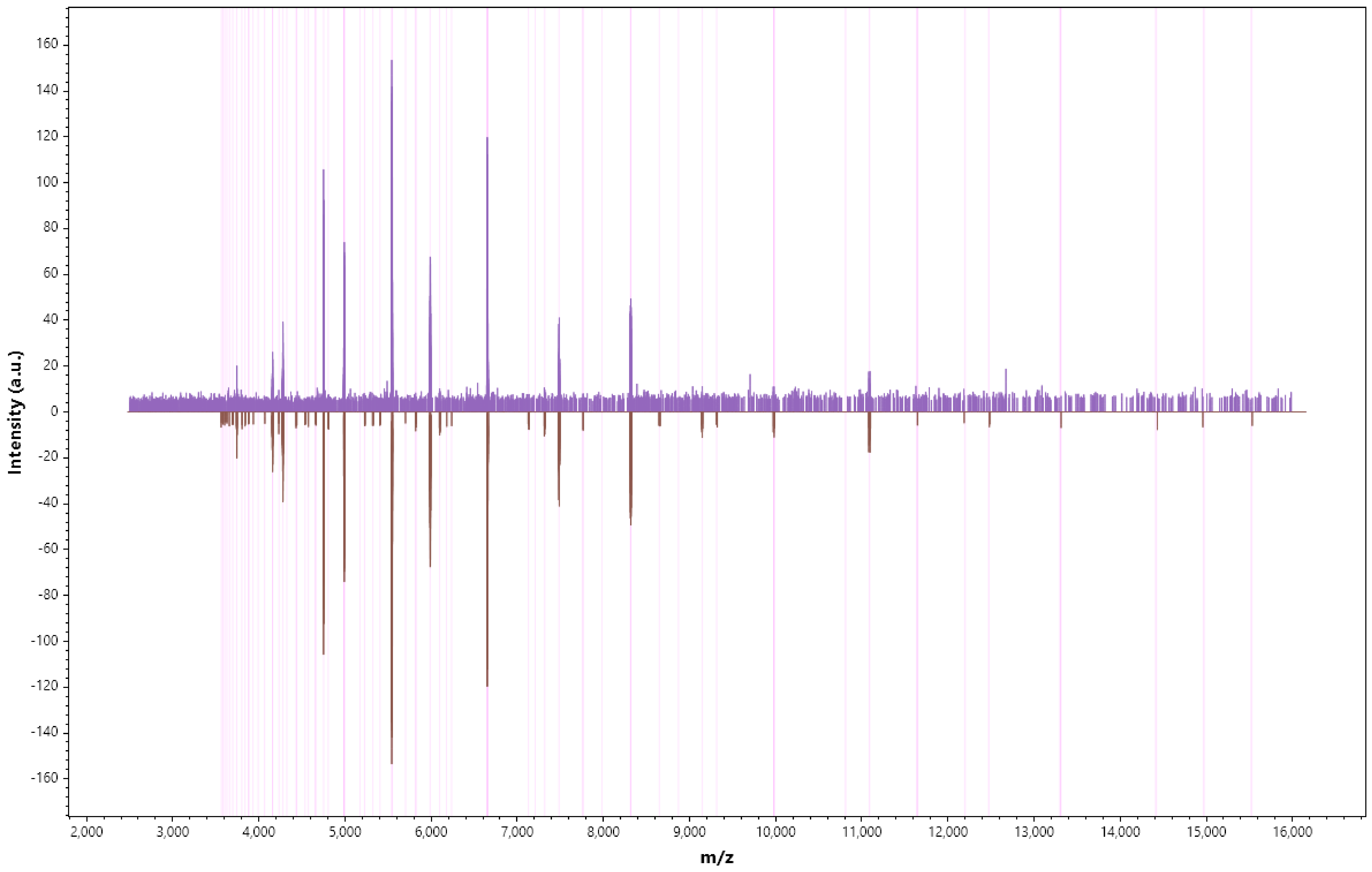

EpCAM_DIA-PTCR_2800-3350_20260217172110 Spectrum107

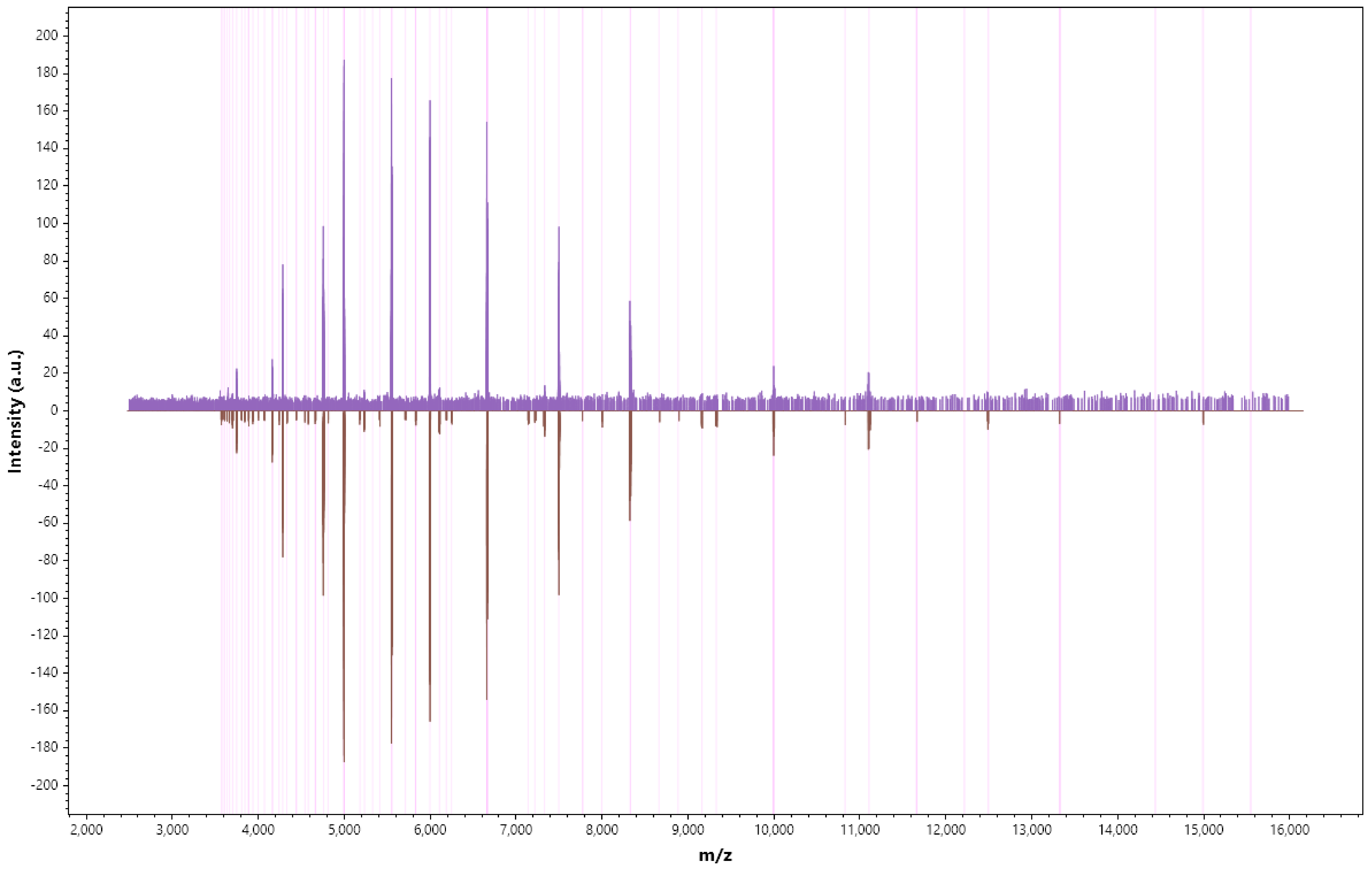

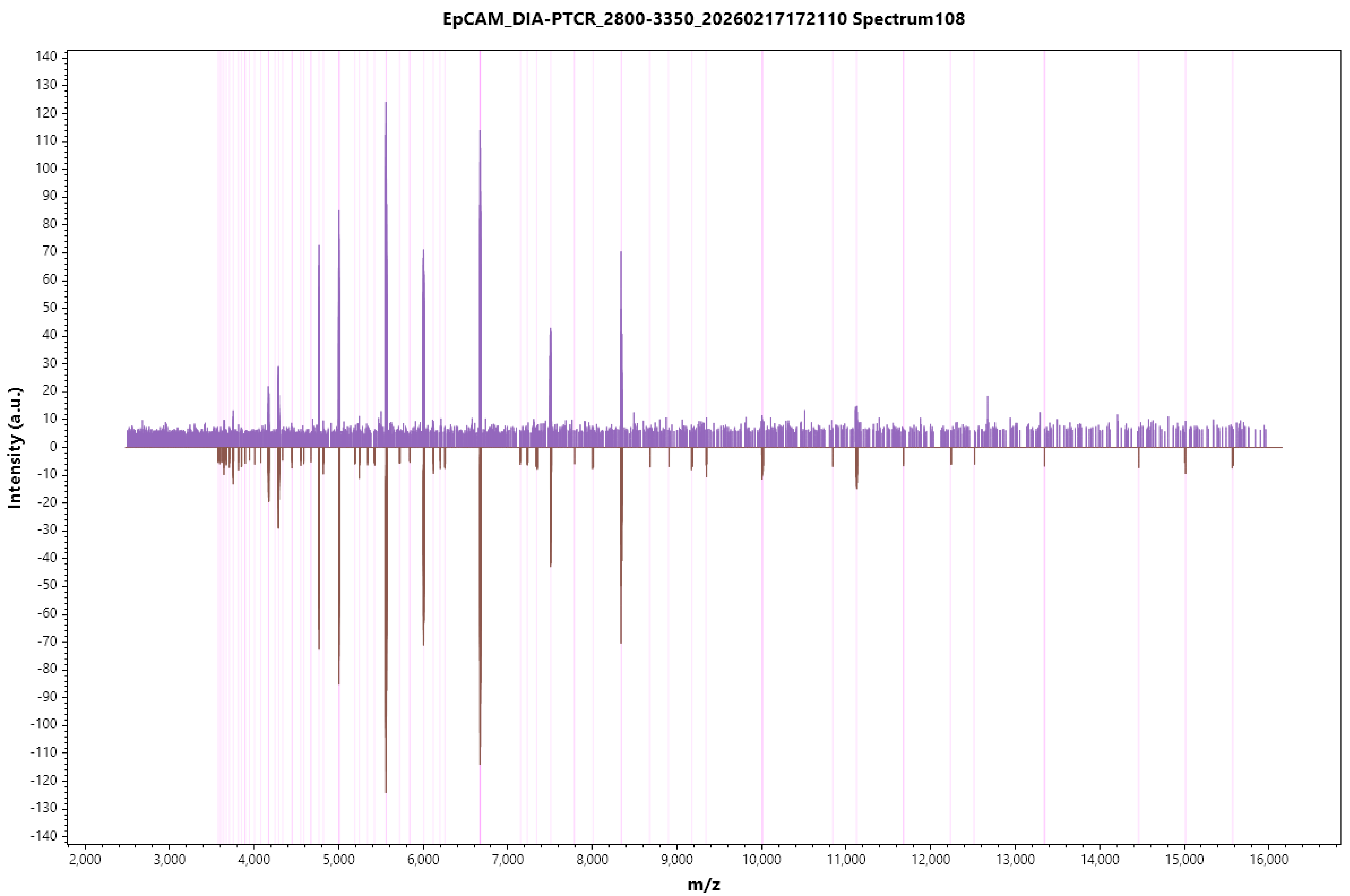
EpCAM_DIA-PTCR_2800-3350_20260217172110 Spectrum108
Intensity (a.u.)
m/z
140
130
120
110
100
90
80
70
60
50
40
30
20
10
0
-10
-20
-30
-40
-50
-60
-70
-80
-90
-100
-110
-120
-130
-140
2,000
3,000
4,000
5,000
6,000
7,000
8,000
9,000
10,000
11,000
12,000
13,000
14,000
15,000
16,000

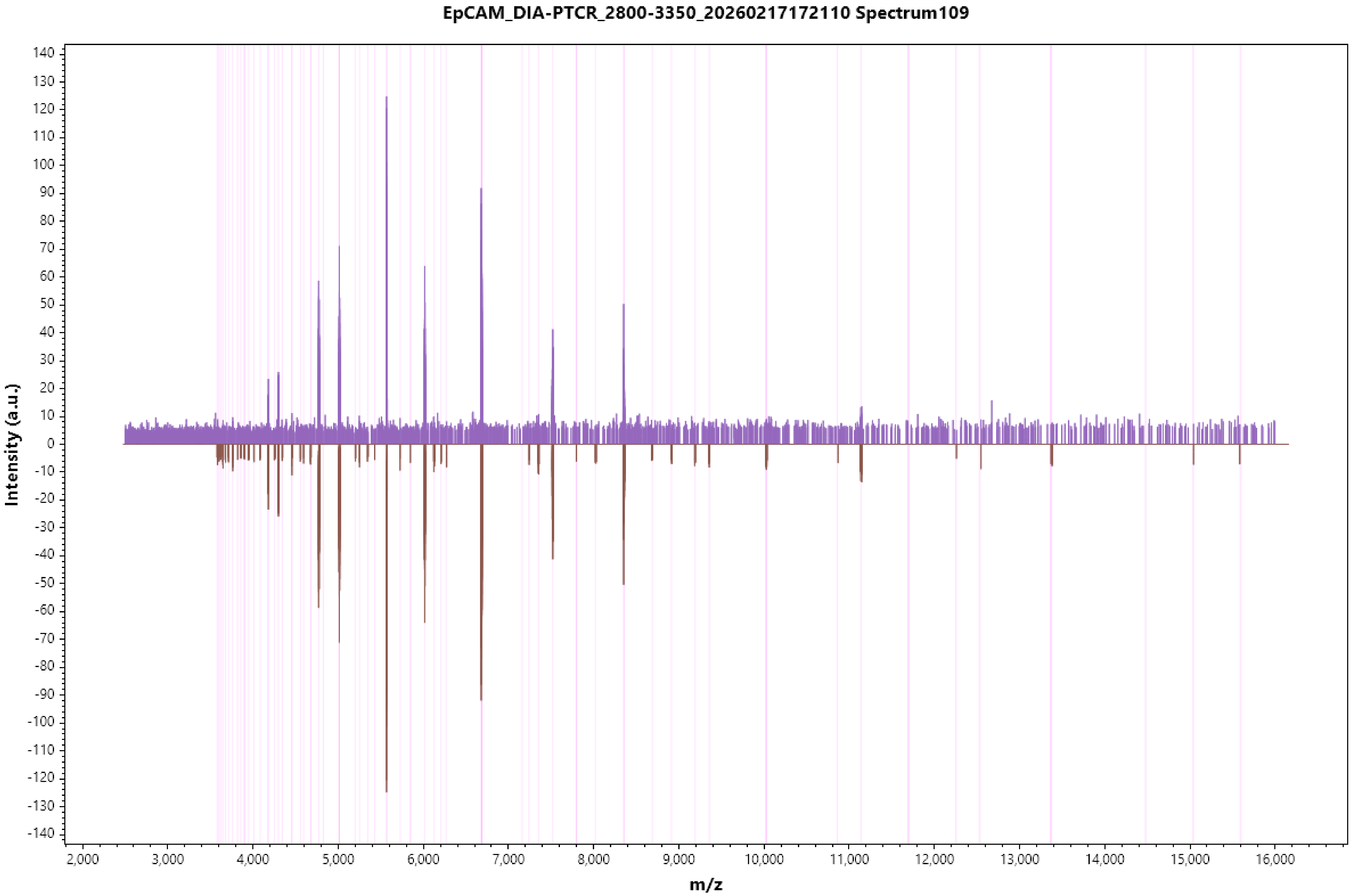

EpCAM_DIA-PTCR_2800-3350_20260217172110 Spectrum109
Intensity (a.u.)
m/z

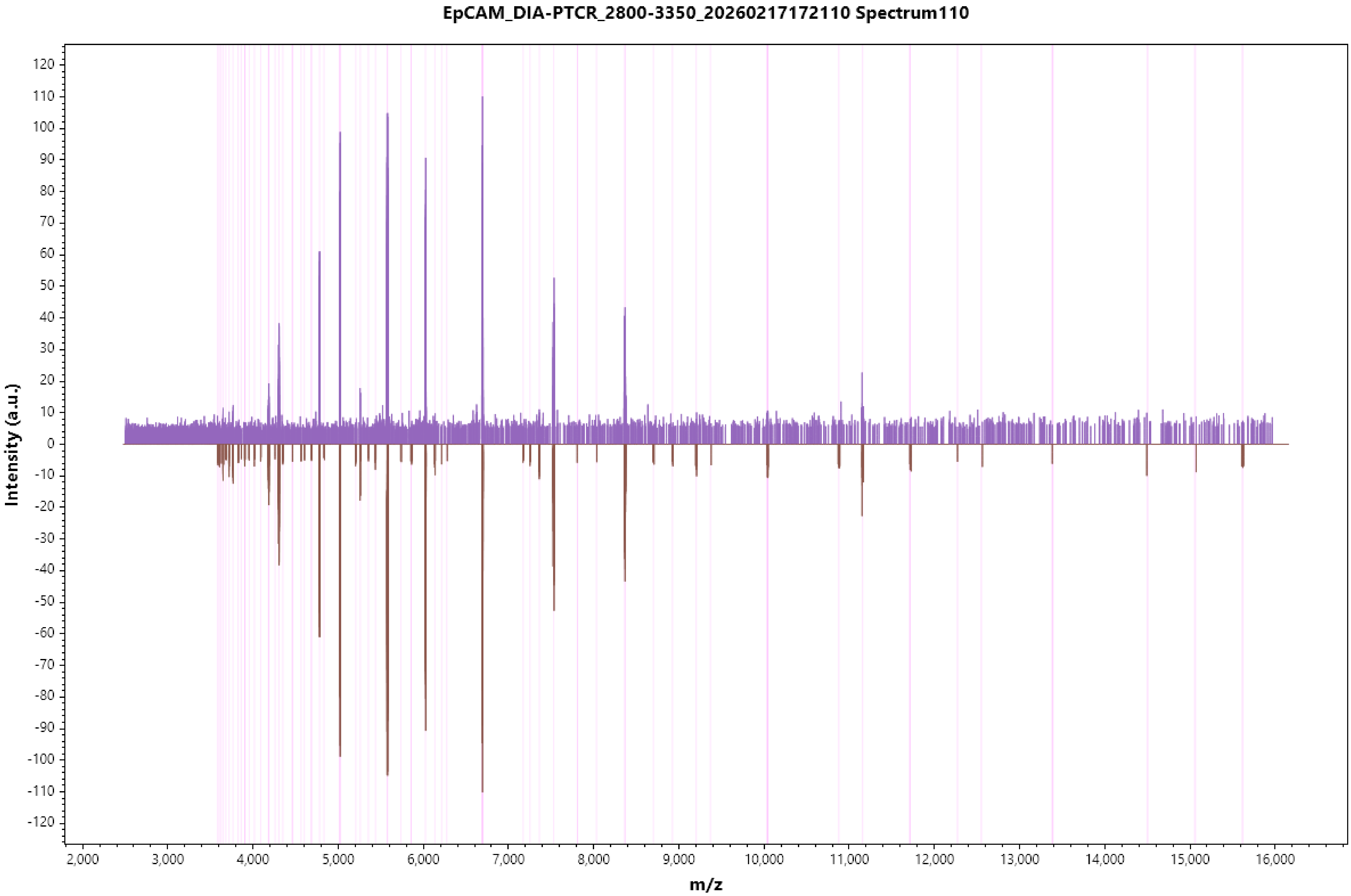

EpCAM_DIA-PTCR_2800-3350_20260217172110 Spectrum110
Intensity (a.u.)
120
110
100
90
80
70
60
50
40
30
20
10
0
-10
-20
-30
-40
-50
-60
-70
-80
-90
-100
-110
-120
2,000
3,000
4,000
5,000
6,000
7,000
8,000
9,000
10,000
11,000
12,000
13,000
14,000
15,000
16,000
m/z

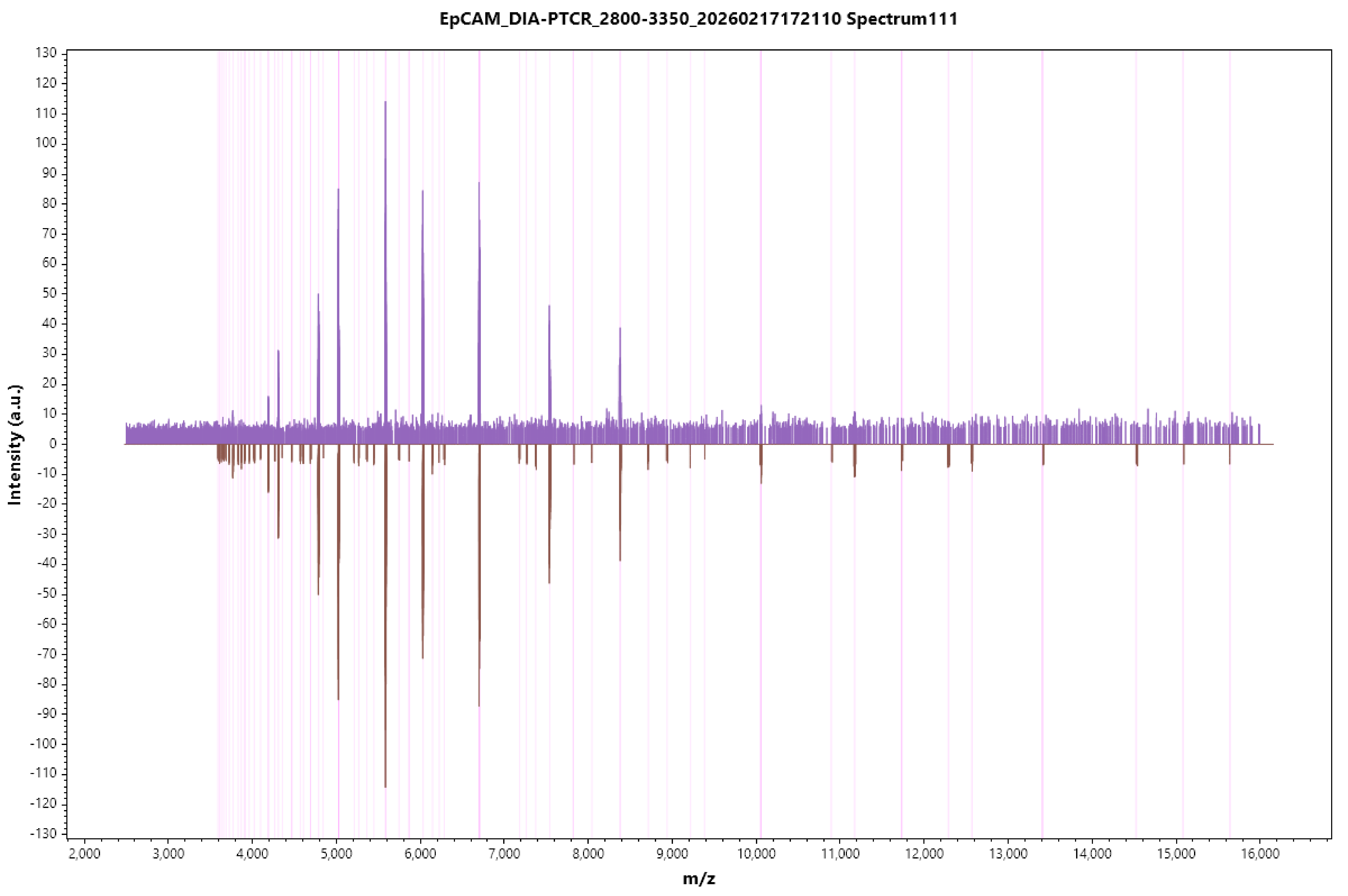
EpCAM_DIA-PTCR_2800-3350_20260217172110 Spectrum111
Intensity (a.u.)
m/z

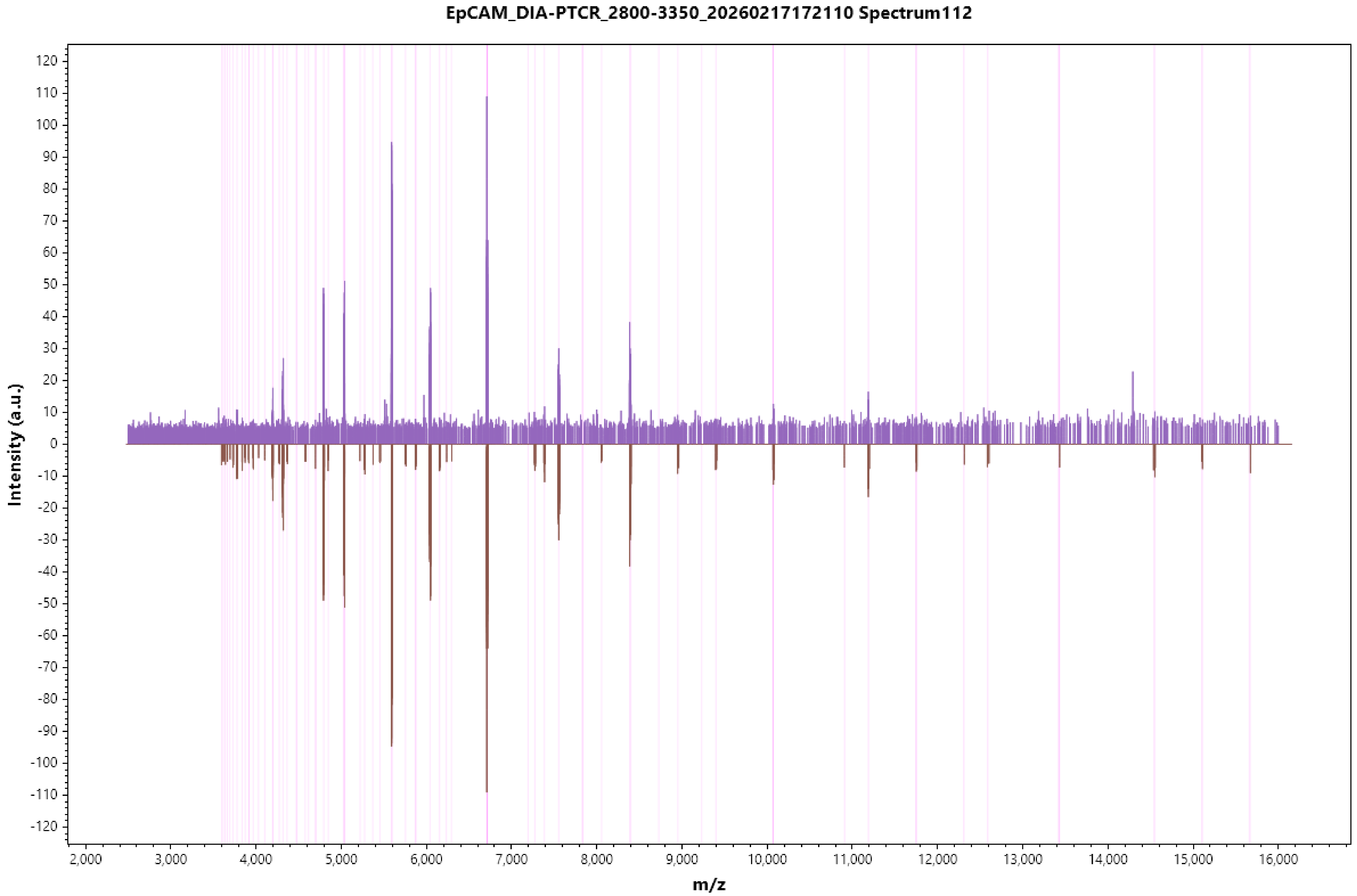
EpCAM_DIA-PTCR_2800-3350_20260217172110 Spectrum112
Intensity (a.u.)
m/z
120
110
100
90
80
70
60
50
40
30
20
10
0
-10
-20
-30
-40
-50
-60
-70
-80
-90
-100
-110
-120
2,000
3,000
4,000
5,000
6,000
7,000
8,000
9,000
10,000
11,000
12,000
13,000
14,000
15,000
16,000

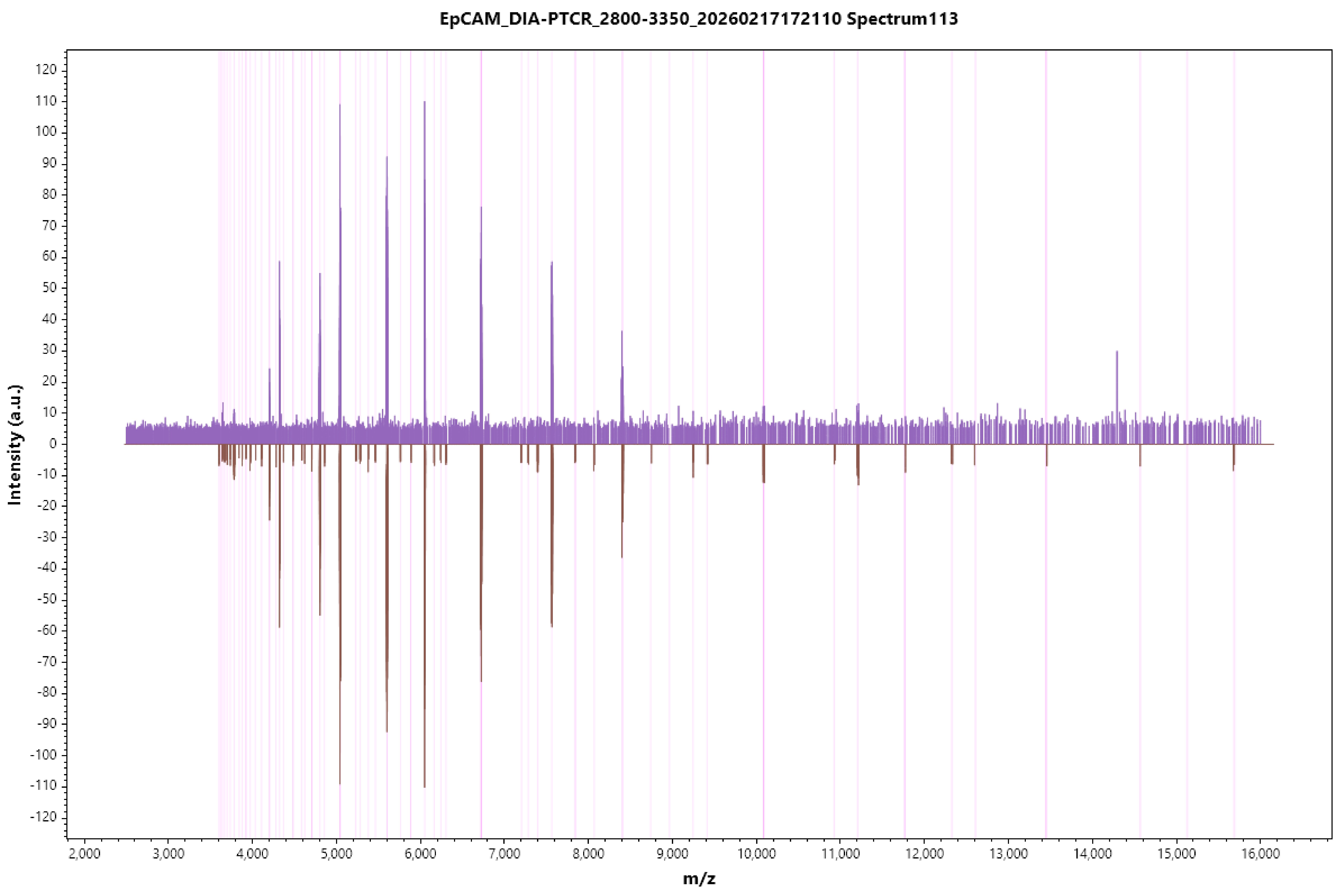

EpCAM_DIA-PTCR_2800-3350_20260217172110 Spectrum113
Intensity (a.u.)
m/z
120
110
100
90
80
70
60
50
40
30
20
10
0
-10
-20
-30
-40
-50
-60
-70
-80
-90
-100
-110
-120
2,000
3,000
4,000
5,000
6,000
7,000
8,000
9,000
10,000
11,000
12,000
13,000
14,000
15,000
16,000

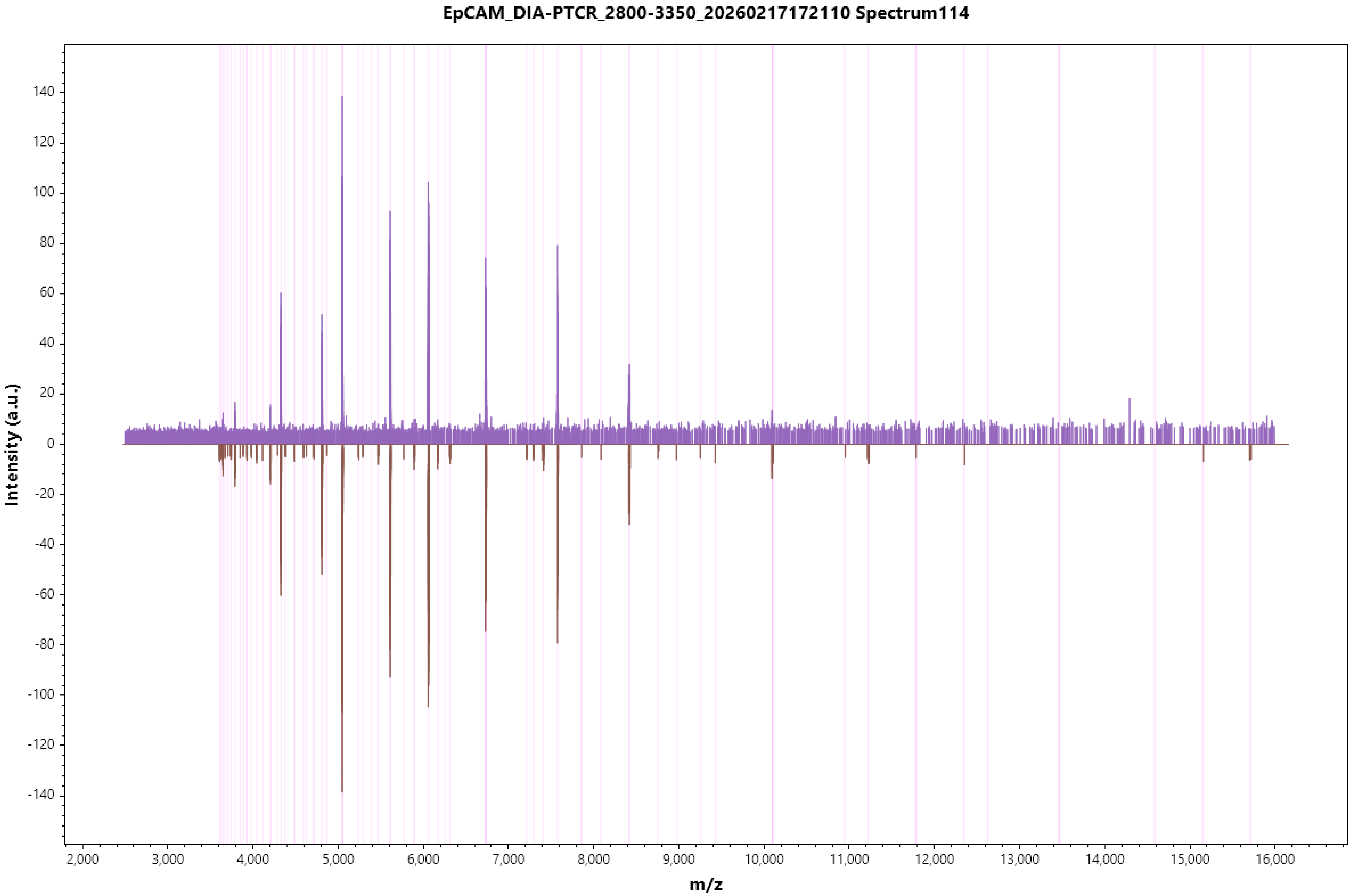

EpCAM_DIA-PTCR_2800-3350_20260217172110 Spectrum114
Intensity (a.u.)
m/z
140
120
100
80
60
40
20
0
-20
-40
-60
-80
-100
-120
-140
2,000
3,000
4,000
5,000
6,000
7,000
8,000
9,000
10,000
11,000
12,000
13,000
14,000
15,000
16,000

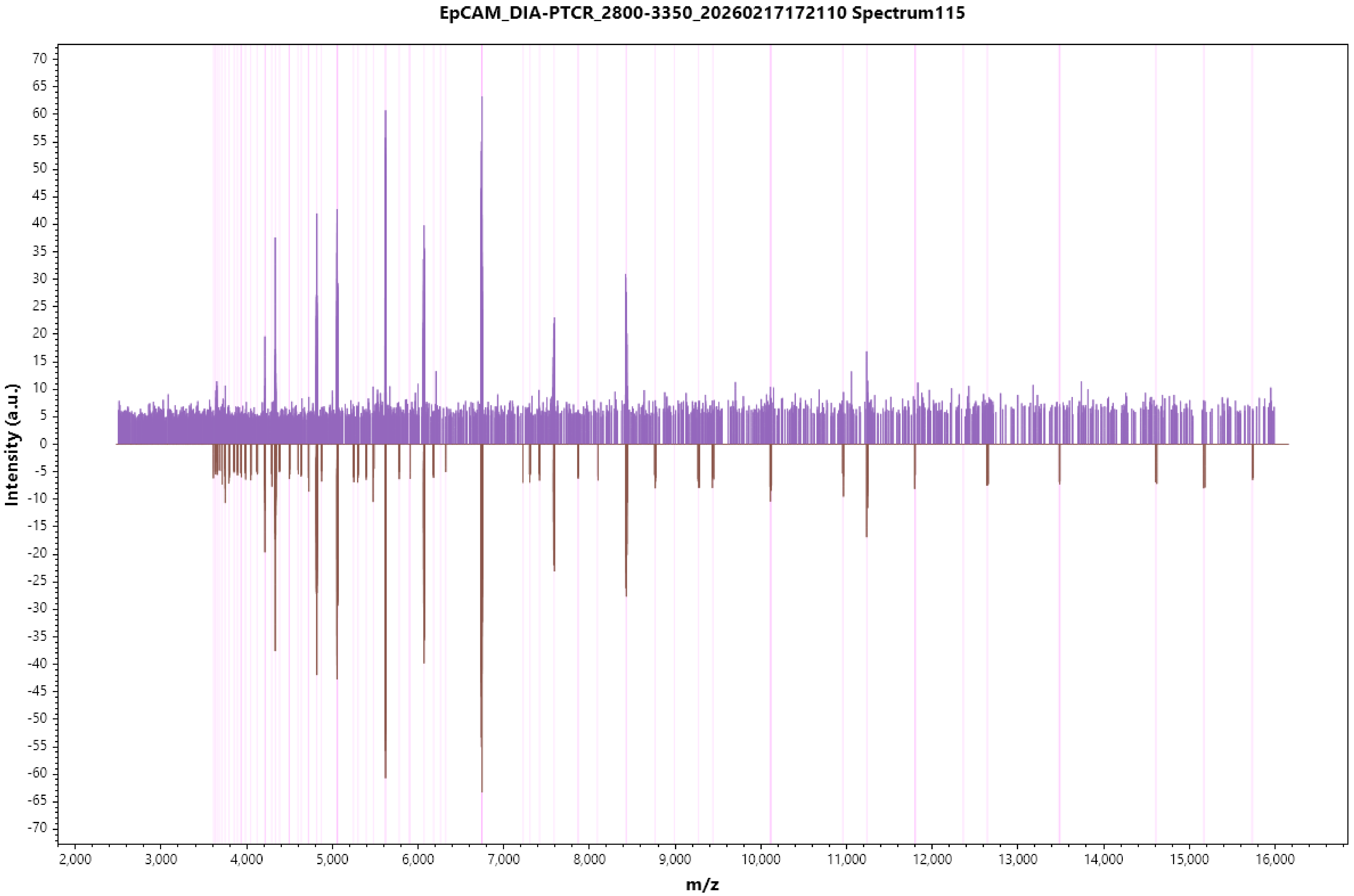
EpCAM_DIA-PTCR_2800-3350_20260217172110 Spectrum115
Intensity (a.u.)
m/z
70
65
60
55
50
45
40
35
30
25
20
15
10
5
0
-5
-10
-15
-20
-25
-30
-35
-40
-45
-50
-55
-60
-65
-70
2,000
3,000
4,000
5,000
6,000
7,000
8,000
9,000
10,000
11,000
12,000
13,000
14,000
15,000
16,000

#### EpCAM_DIA-PTCR_2800-3350_20260217172110 Spectrum116

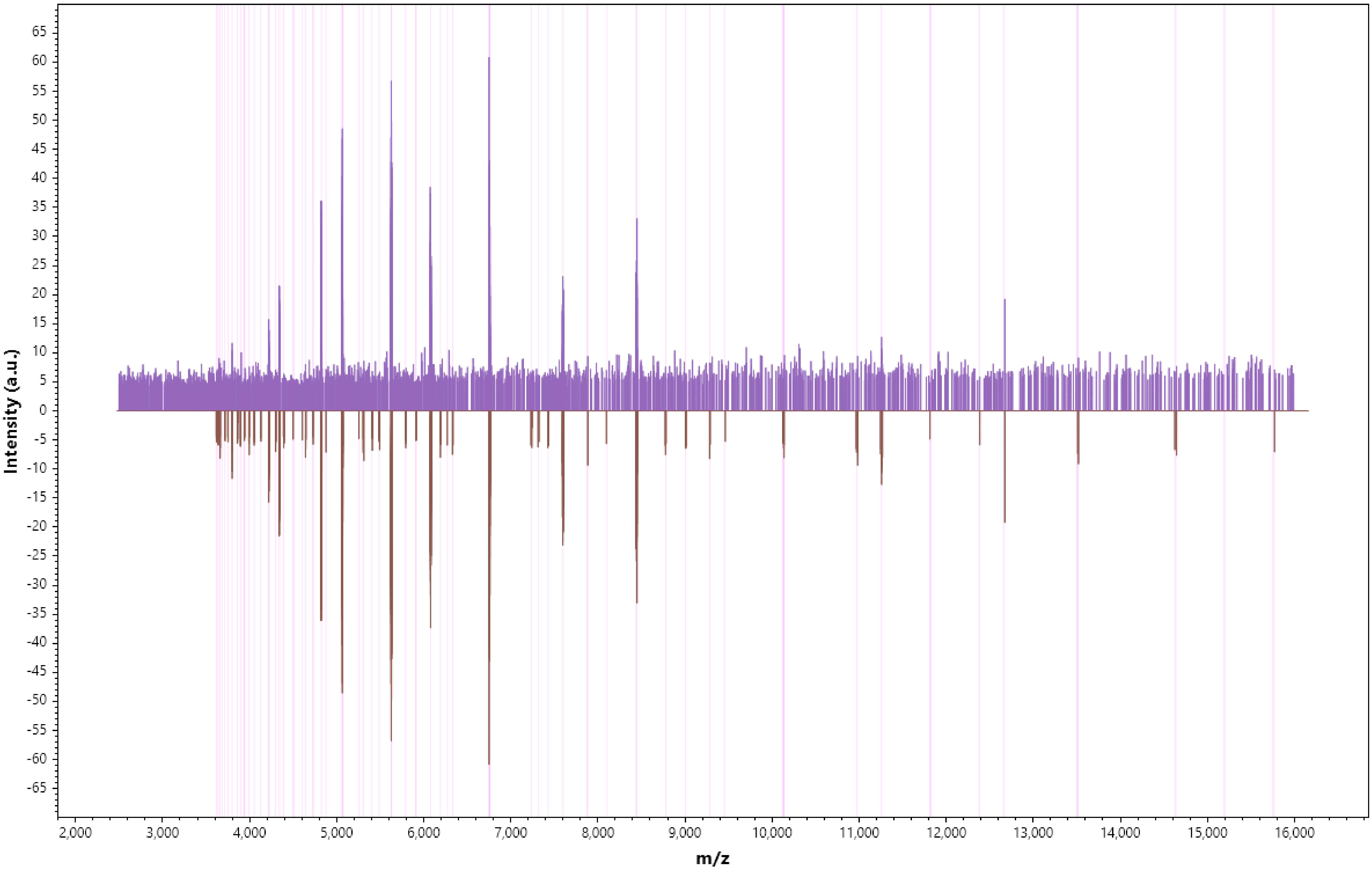

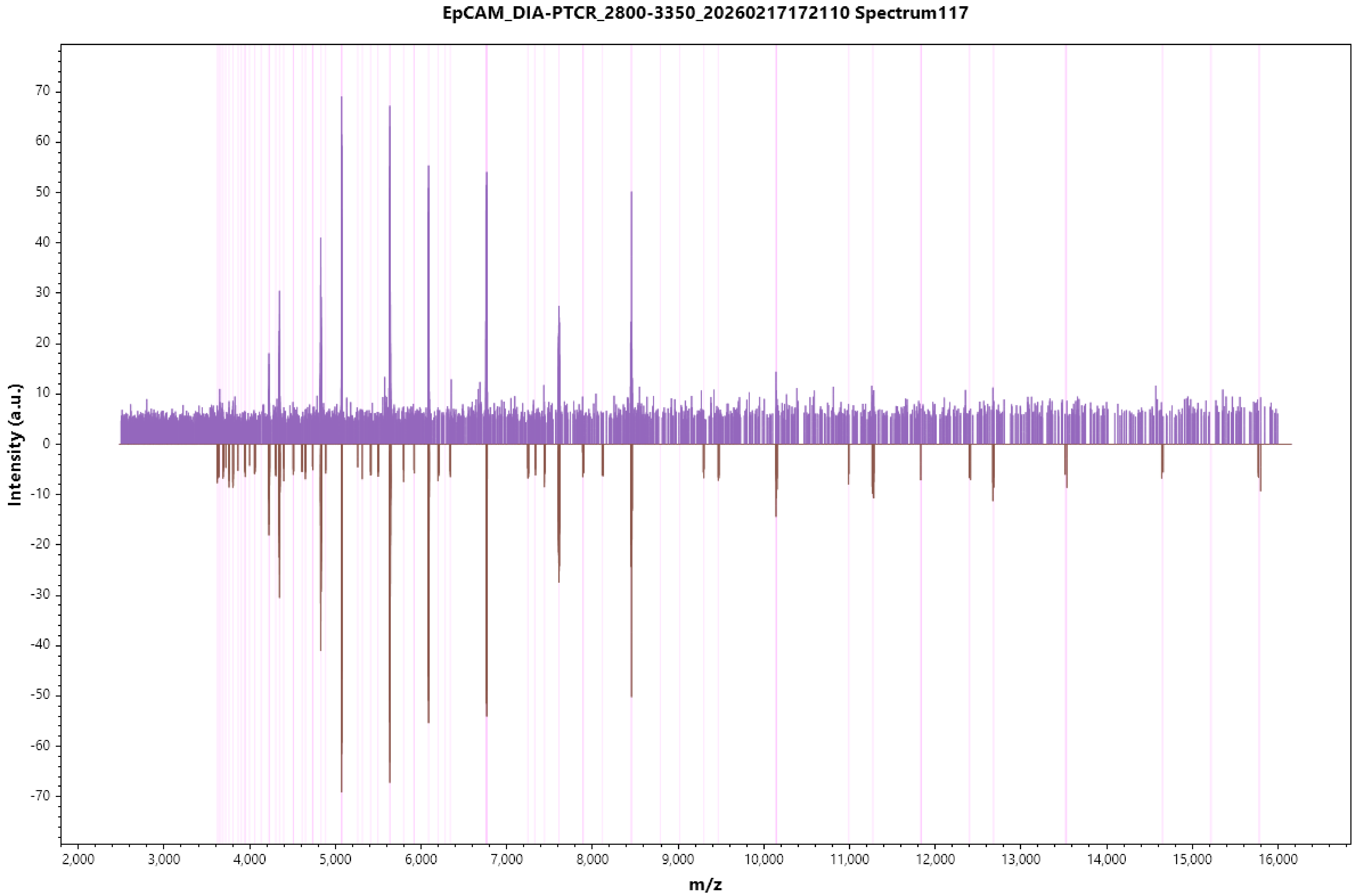
EpCAM_DIA-PTCR_2800-3350_20260217172110 Spectrum117
Intensity (a.u.)
m/z
70
60
50
40
30
20
10
0
-10
-20
-30
-40
-50
-60
-70
2,000
3,000
4,000
5,000
6,000
7,000
8,000
9,000
10,000
11,000
12,000
13,000
14,000
15,000
16,000

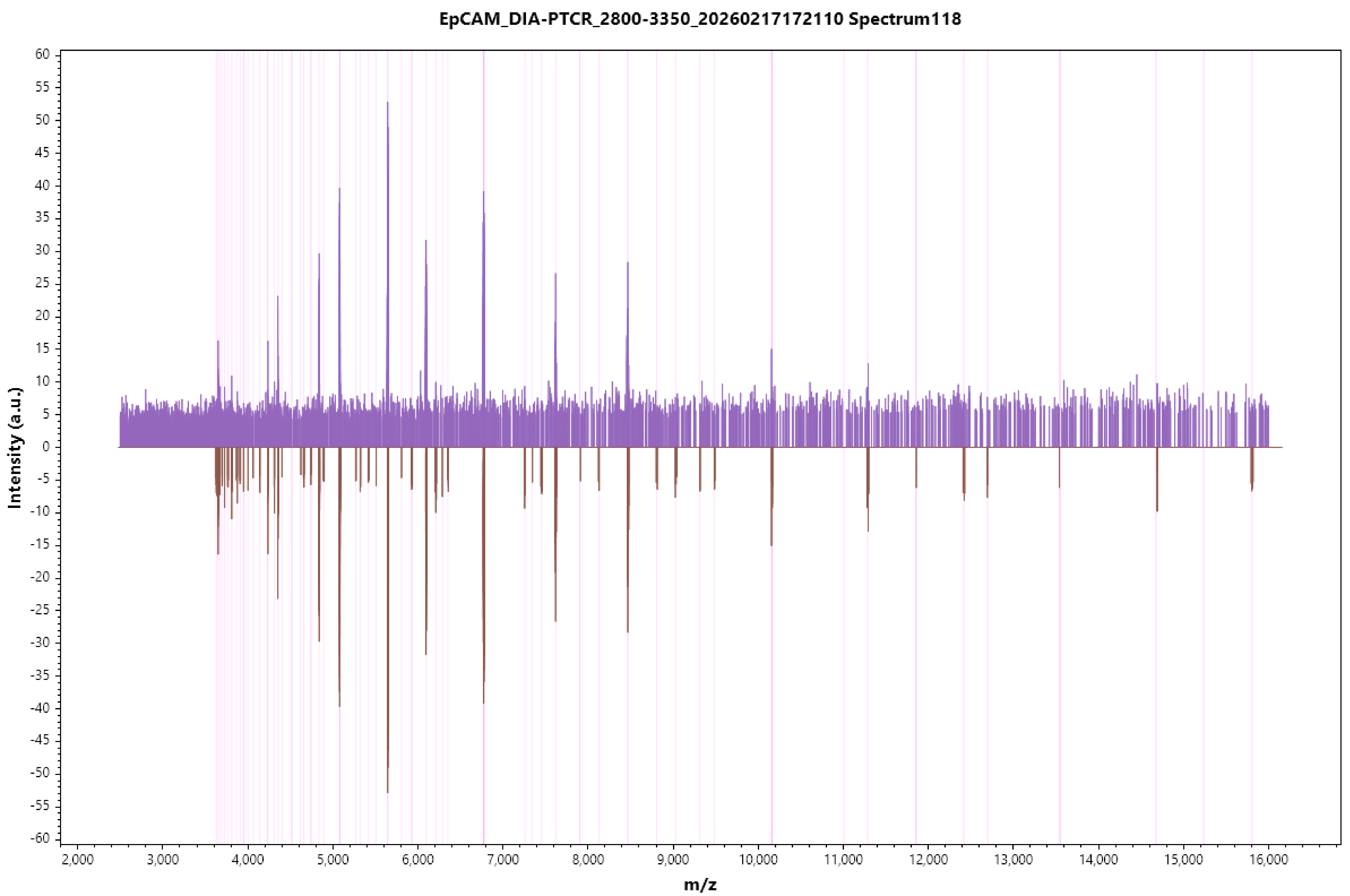

EpCAM_DIA-PTCR_2800-3350_20260217172110 Spectrum118
Intensity (a.u.)
m/z
60
55
50
45
40
35
30
25
20
15
10
5
0
-5
-10
-15
-20
-25
-30
-35
-40
-45
-50
-55
-60
2,000
3,000
4,000
5,000
6,000
7,000
8,000
9,000
10,000
11,000
12,000
13,000
14,000
15,000
16,000

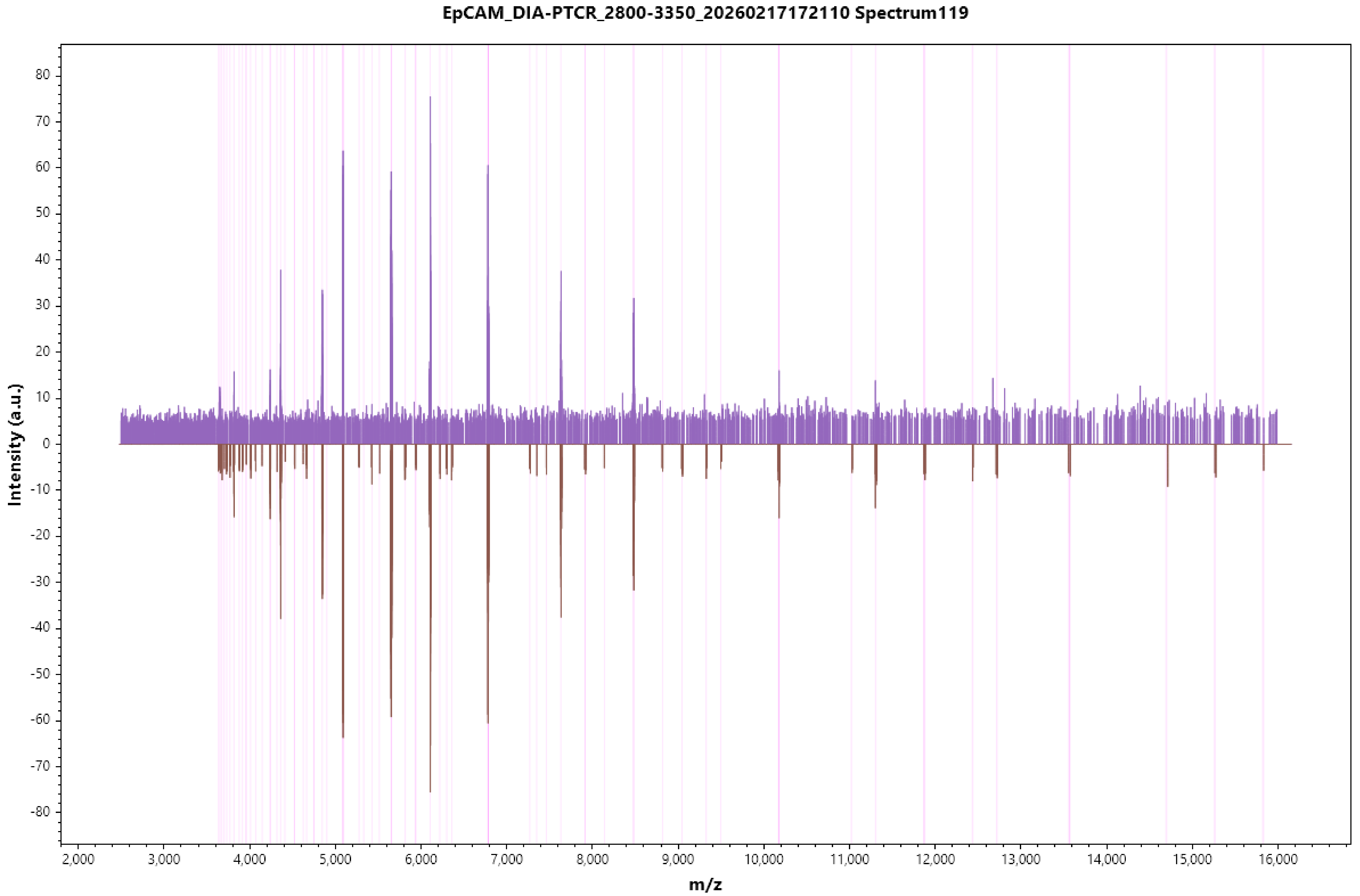
EpCAM_DIA-PTCR_2800-3350_20260217172110 Spectrum119
Intensity (a.u.)
m/z
80
70
60
50
40
30
20
10
0
-10
-20
-30
-40
-50
-60
-70
-80
2,000
3,000
4,000
5,000
6,000
7,000
8,000
9,000
10,000
11,000
12,000
13,000
14,000
15,000
16,000

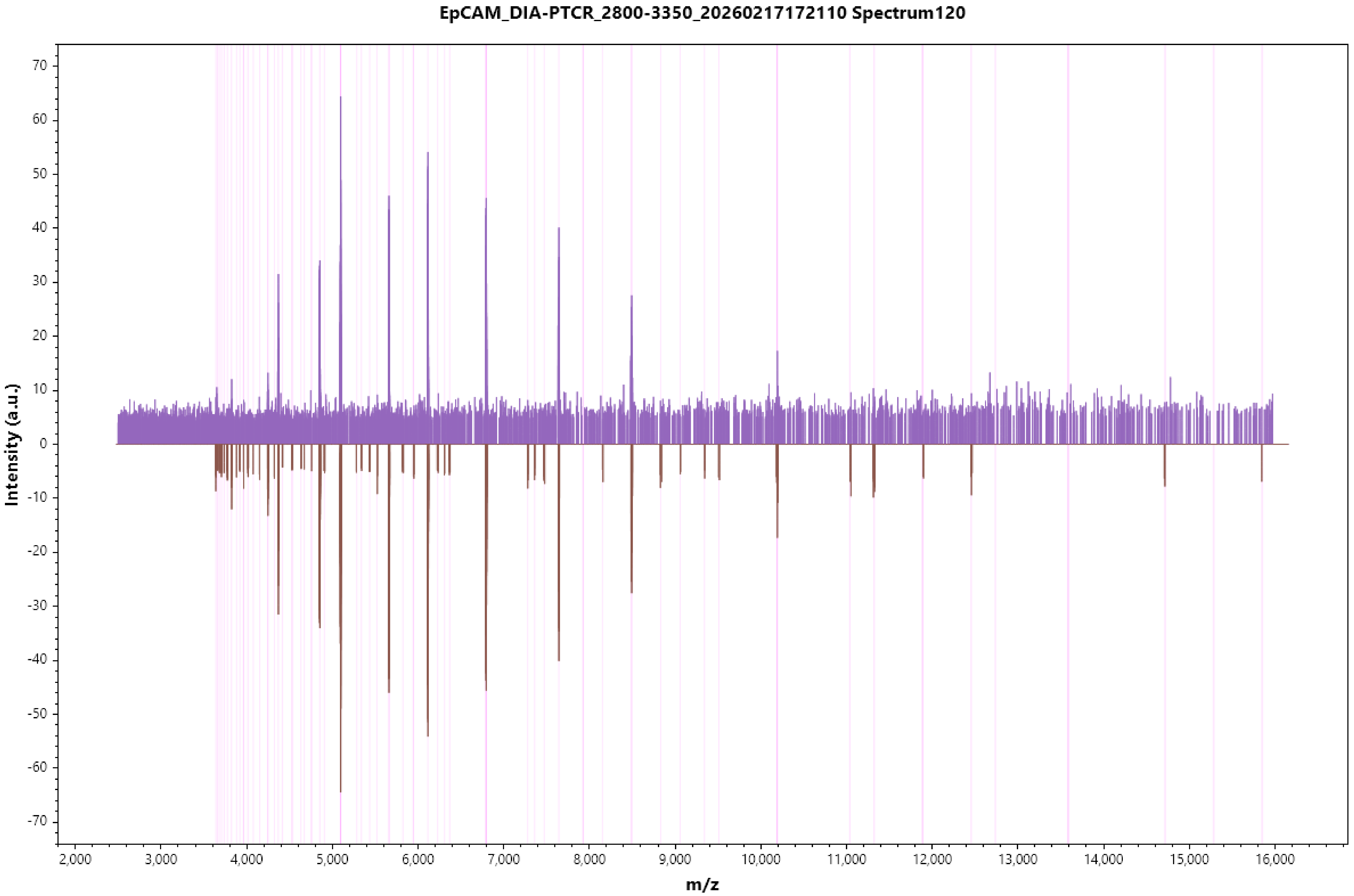

EpCAM_DIA-PTCR_2800-3350_20260217172110 Spectrum120
Intensity (a.u.)
70
60
50
40
30
20
10
0
-10
-20
-30
-40
-50
-60
-70
2,000
3,000
4,000
5,000
6,000
7,000
8,000
9,000
10,000
11,000
12,000
13,000
14,000
15,000
16,000
m/z

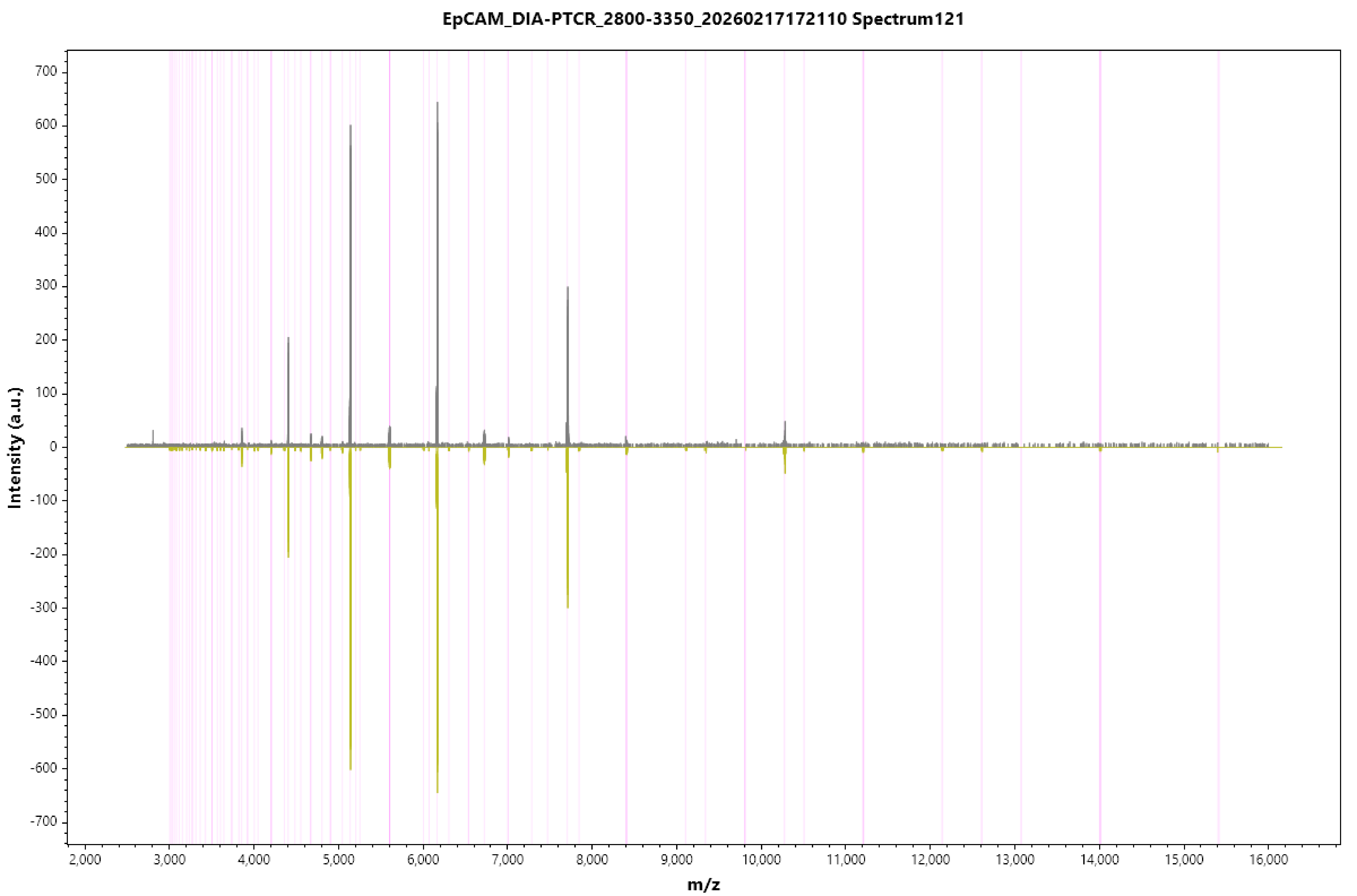
EpCAM_DIA-PTCR_2800-3350_20260217172110 Spectrum121
Intensity (a.u.)
700
600
500
400
300
200
100
0
-100
-200
-300
-400
-500
-600
-700
2,000
3,000
4,000
5,000
6,000
7,000
8,000
9,000
10,000
11,000
12,000
13,000
14,000
15,000
16,000
m/z

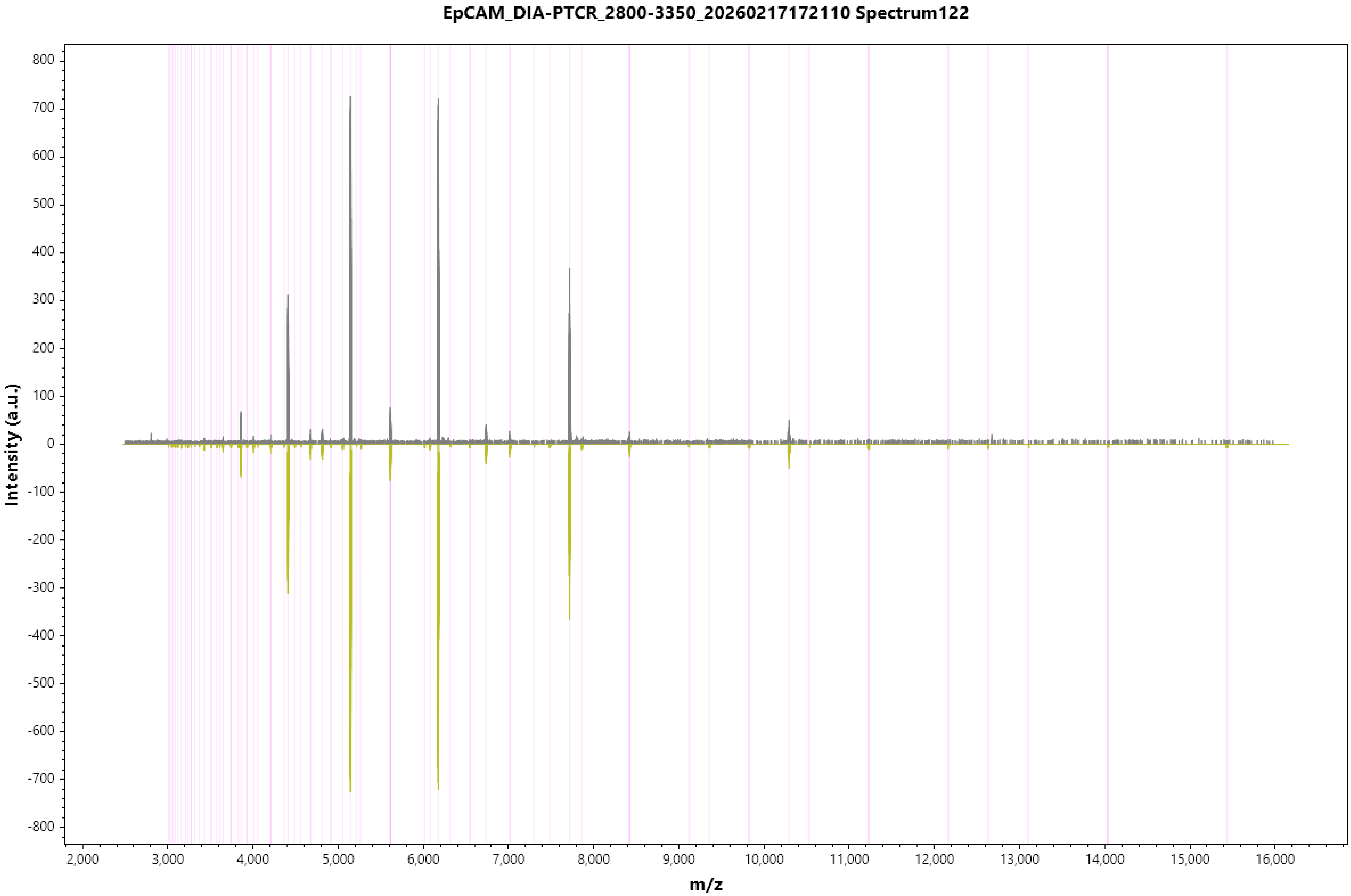
EpCAM_DIA-PTCR_2800-3350_20260217172110 Spectrum122
Intensity (a.u.)
m/z
800
700
600
500
400
300
200
100
0
-100
-200
-300
-400
-500
-600
-700
-800
2,000
3,000
4,000
5,000
6,000
7,000
8,000
9,000
10,000
11,000
12,000
13,000
14,000
15,000
16,000

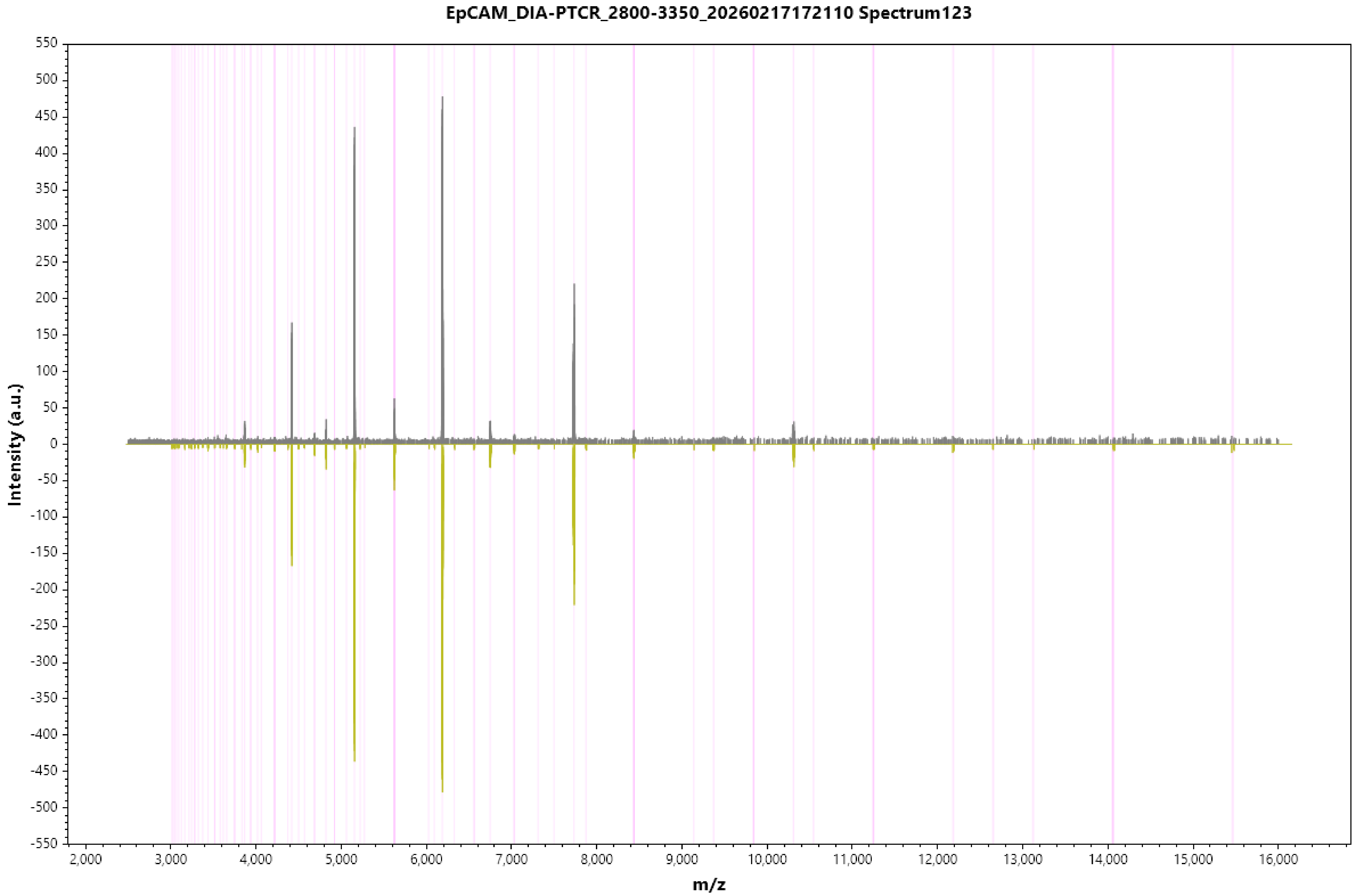
EpCAM_DIA-PTCR_2800-3350_20260217172110 Spectrum123
Intensity (a.u.)
m/z
550
500
450
400
350
300
250
200
150
100
50
0
-50
-100
-150
-200
-250
-300
-350
-400
-450
-500
-550
2,000
3,000
4,000
5,000
6,000
7,000
8,000
9,000
10,000
11,000
12,000
13,000
14,000
15,000
16,000

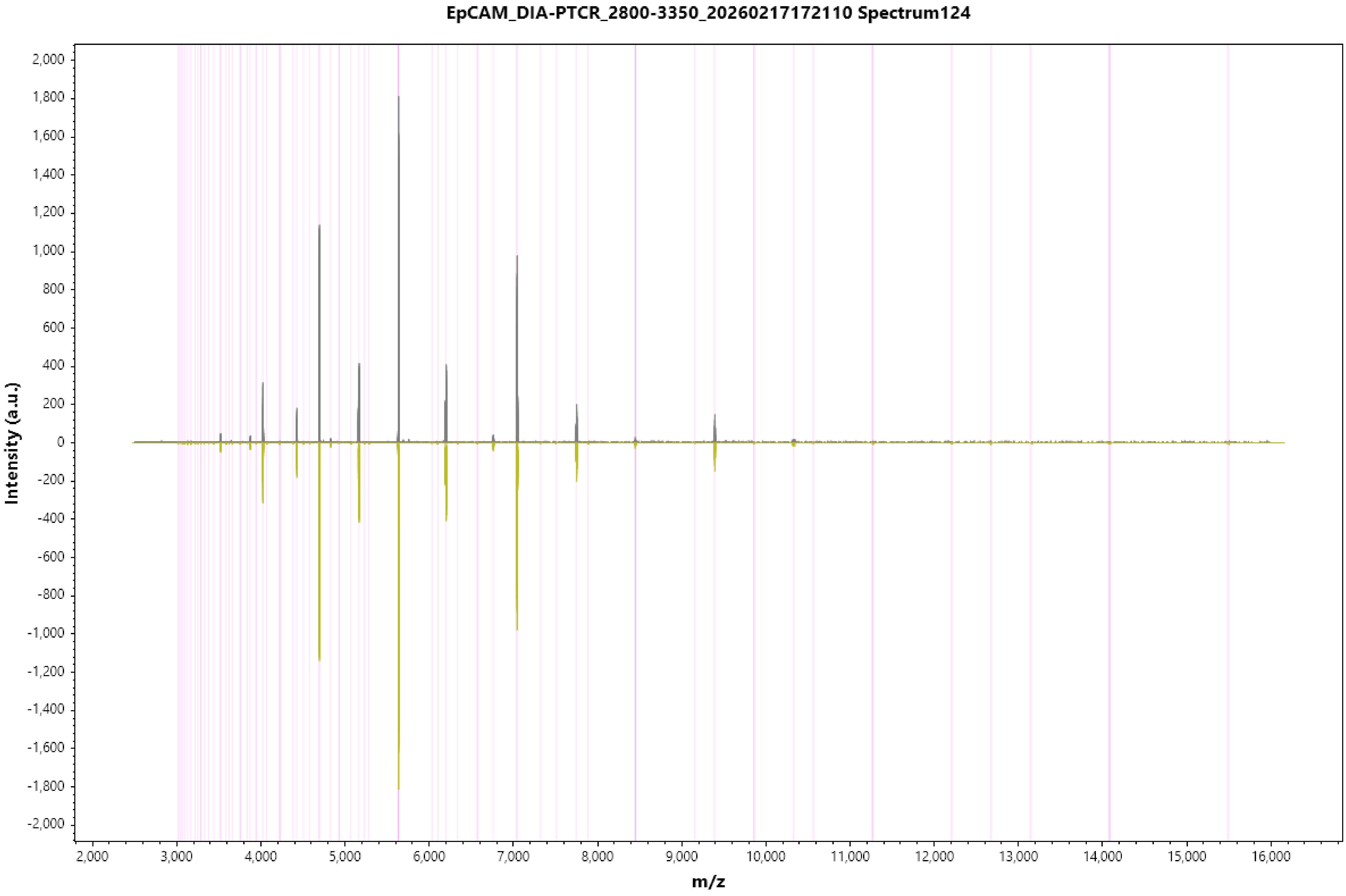
EpCAM_DIA-PTCR_2800-3350_20260217172110 Spectrum124
Intensity (a.u.)
m/z
2,000
1,800
1,600
1,400
1,200
1,000
800
600
400
200
0
-200
-400
-600
-800
-1,000
-1,200
-1,400
-1,600
-1,800
-2,000
2,000
3,000
4,000
5,000
6,000
7,000
8,000
9,000
10,000
11,000
12,000
13,000
14,000
15,000
16,000

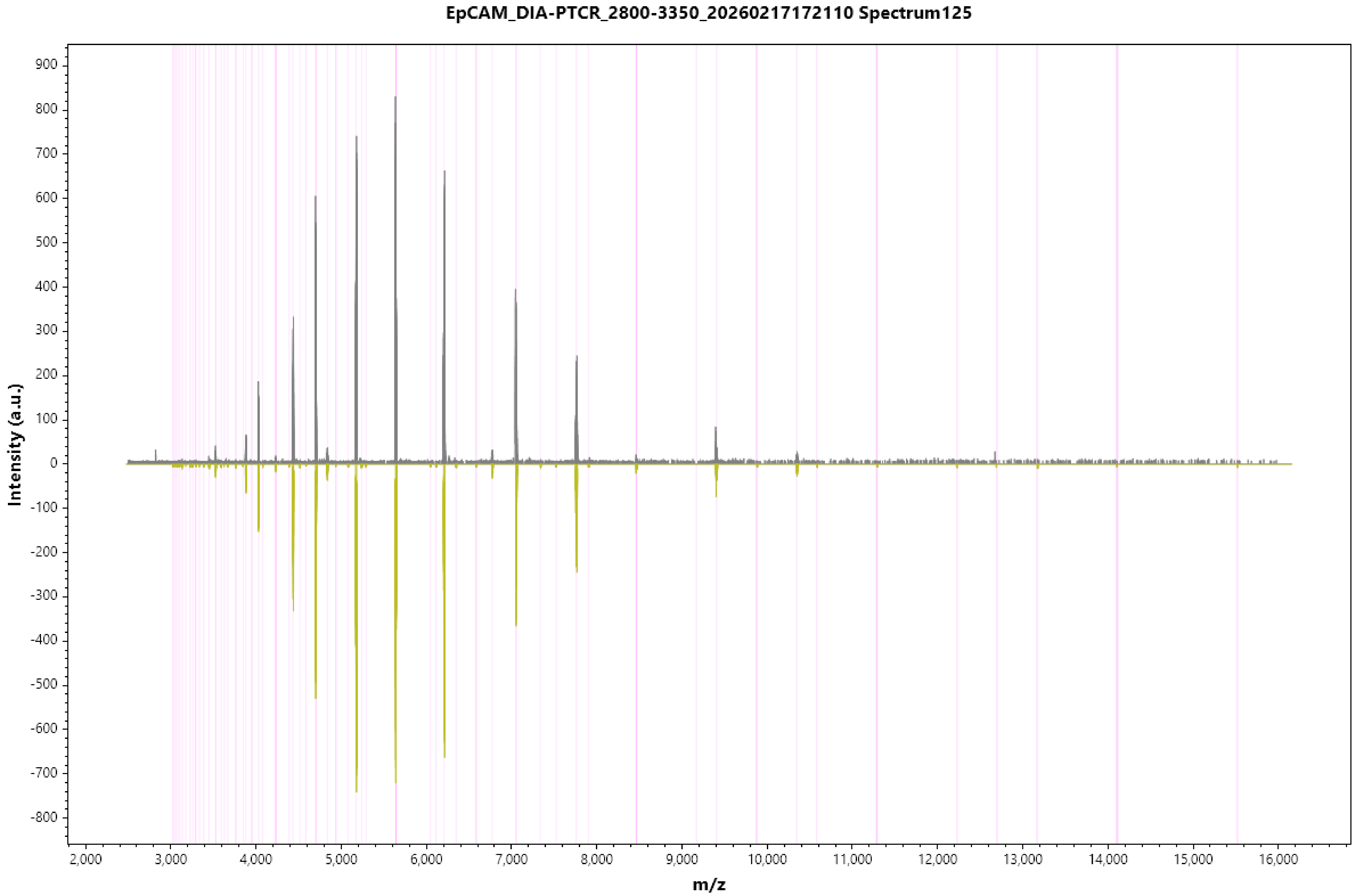
EpCAM_DIA-PTCR_2800-3350_20260217172110 Spectrum125
Intensity (a.u.)
m/z
900
800
700
600
500
400
300
200
100
0
-100
-200
-300
-400
-500
-600
-700
-800
2,000
3,000
4,000
5,000
6,000
7,000
8,000
9,000
10,000
11,000
12,000
13,000
14,000
15,000
16,000

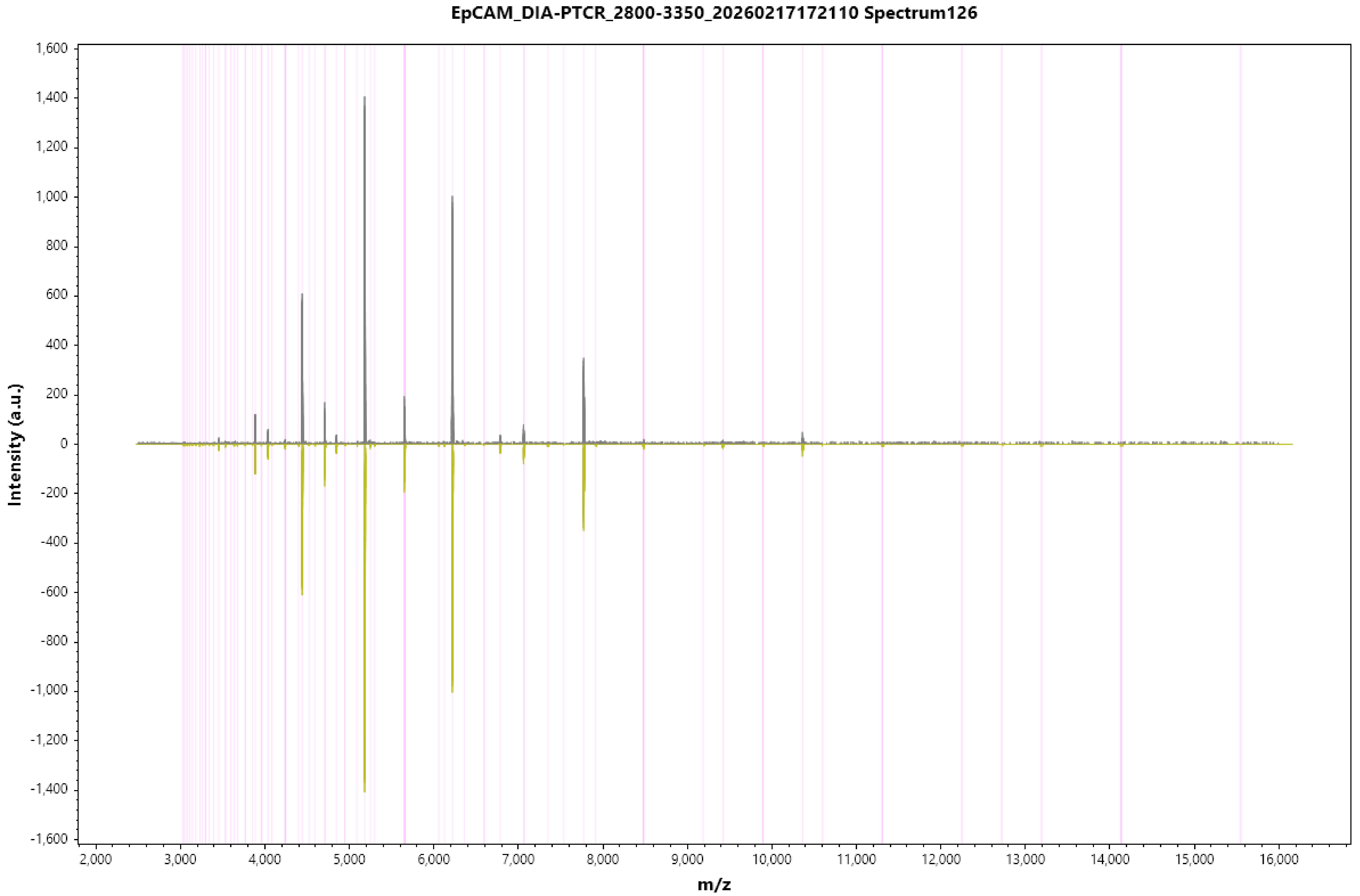
EpCAM_DIA-PTCR_2800-3350_20260217172110 Spectrum126
Intensity (a.u.)
1,600
1,400
1,200
1,000
800
600
400
200
0
-200
-400
-600
-800
-1,000
-1,200
-1,400
-1,600
2,000
3,000
4,000
5,000
6,000
7,000
8,000
9,000
10,000
11,000
12,000
13,000
14,000
15,000
16,000
m/z

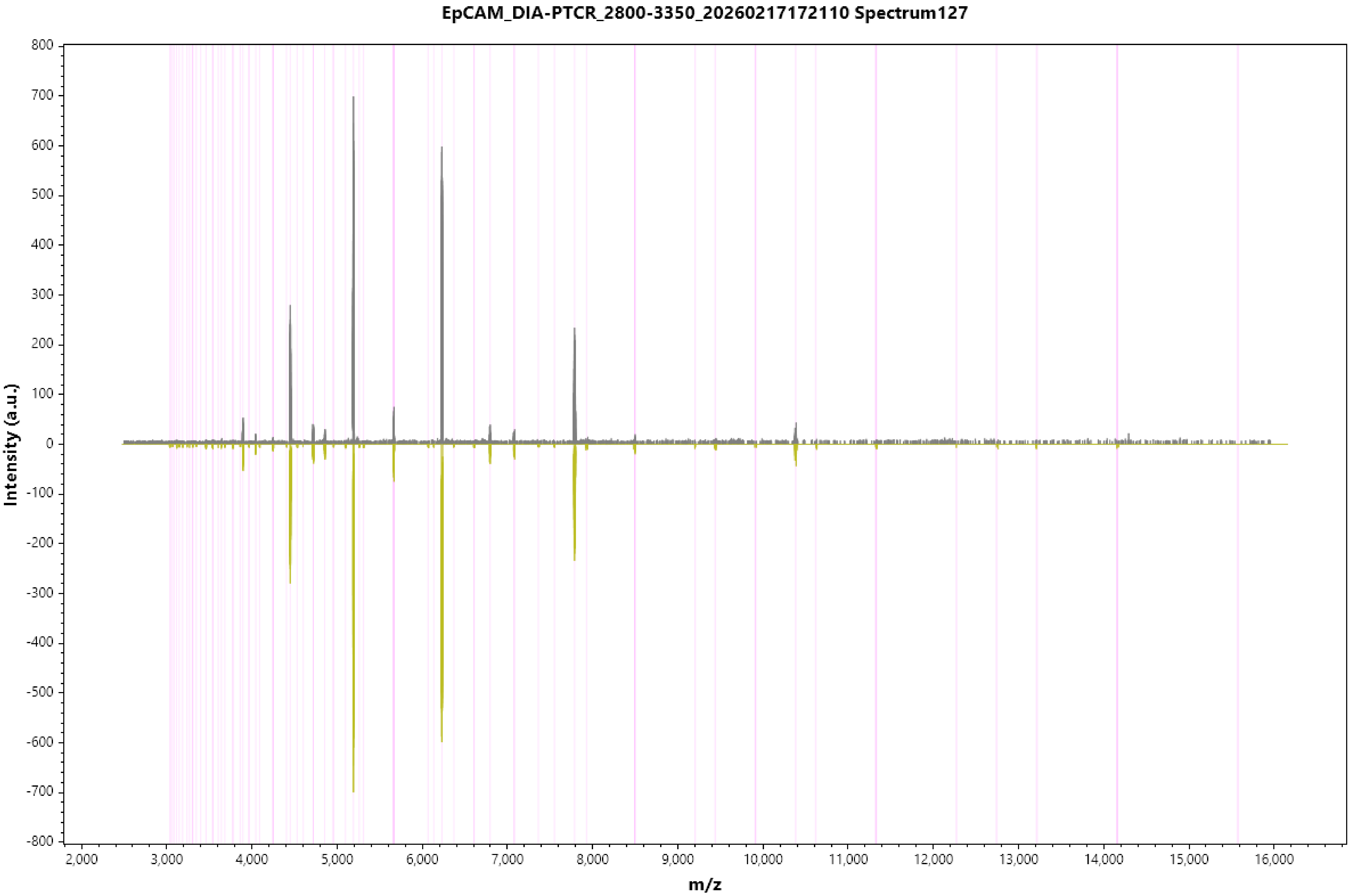
EpCAM_DIA-PTCR_2800-3350_20260217172110 Spectrum127
Intensity (a.u.)
m/z
800
700
600
500
400
300
200
100
0
-100
-200
-300
-400
-500
-600
-700
-800
2,000
3,000
4,000
5,000
6,000
7,000
8,000
9,000
10,000
11,000
12,000
13,000
14,000
15,000
16,000

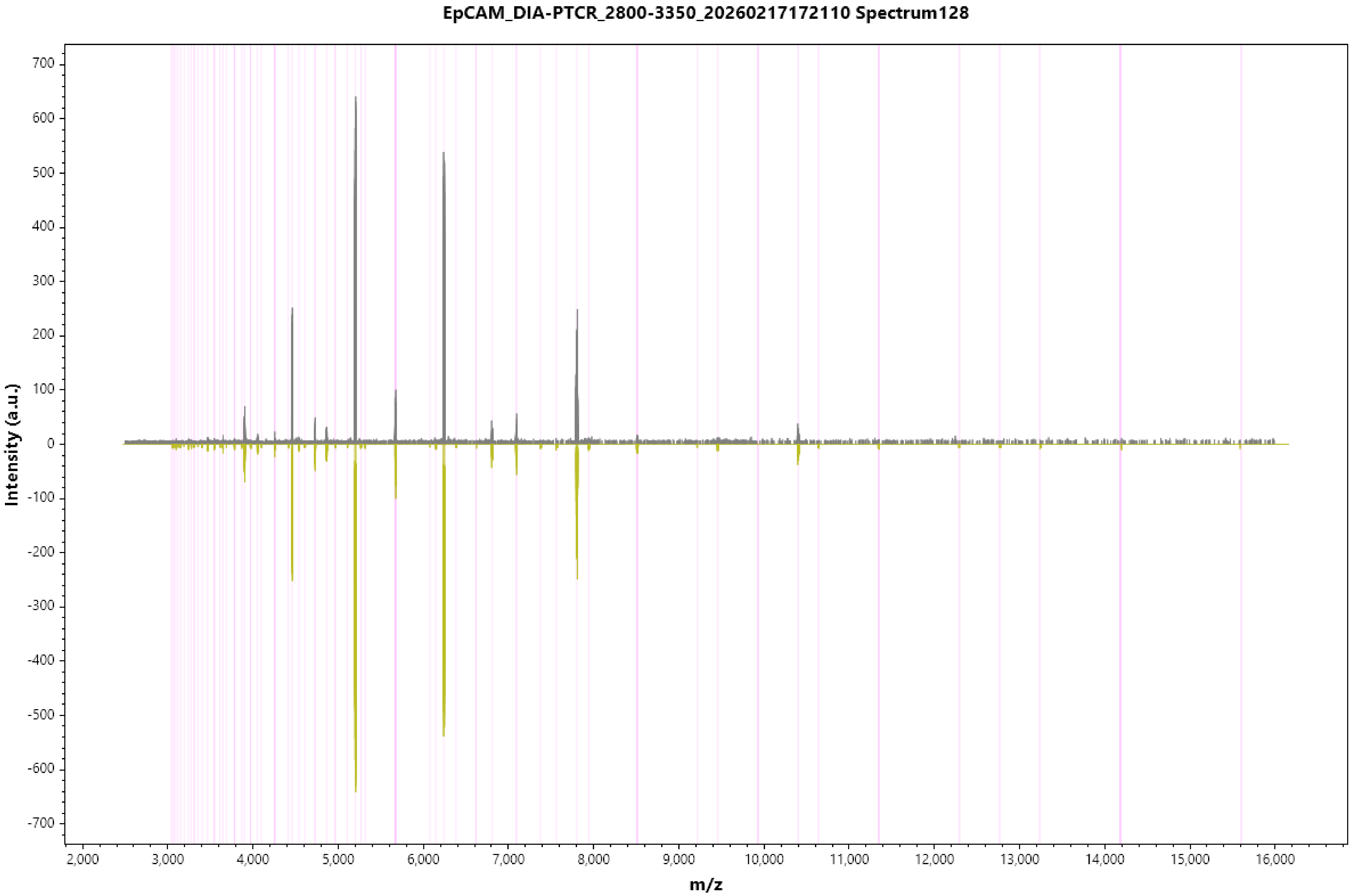
EpCAM_DIA-PTCR_2800-3350_20260217172110 Spectrum128
Intensity (a.u.)
m/z
700
600
500
400
300
200
100
0
-100
-200
-300
-400
-500
-600
-700
2,000
3,000
4,000
5,000
6,000
7,000
8,000
9,000
10,000
11,000
12,000
13,000
14,000
15,000
16,000

#### EpCAM_DIA-PTCR_2800-3350_20260217172110 Spectrum129

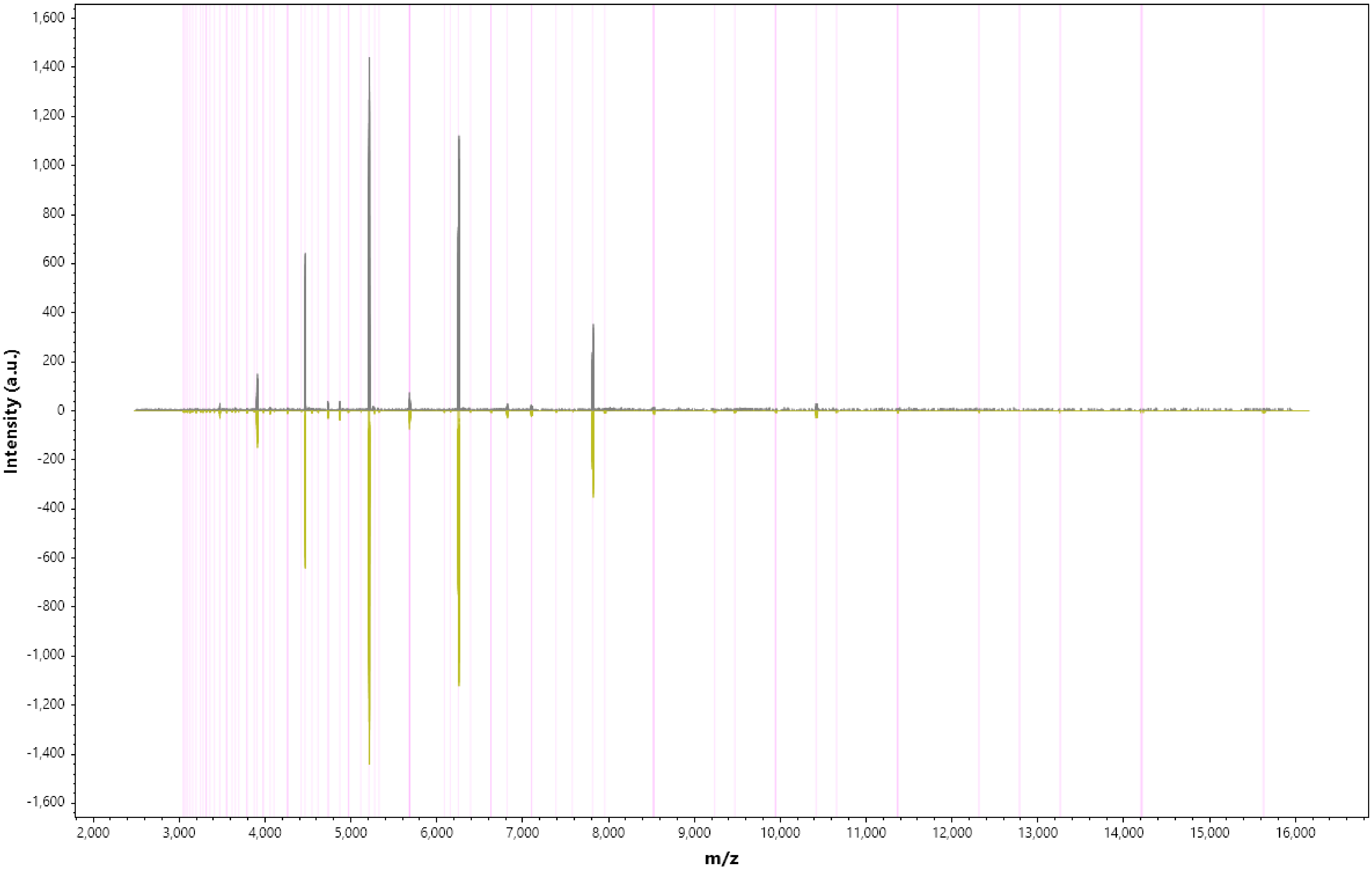

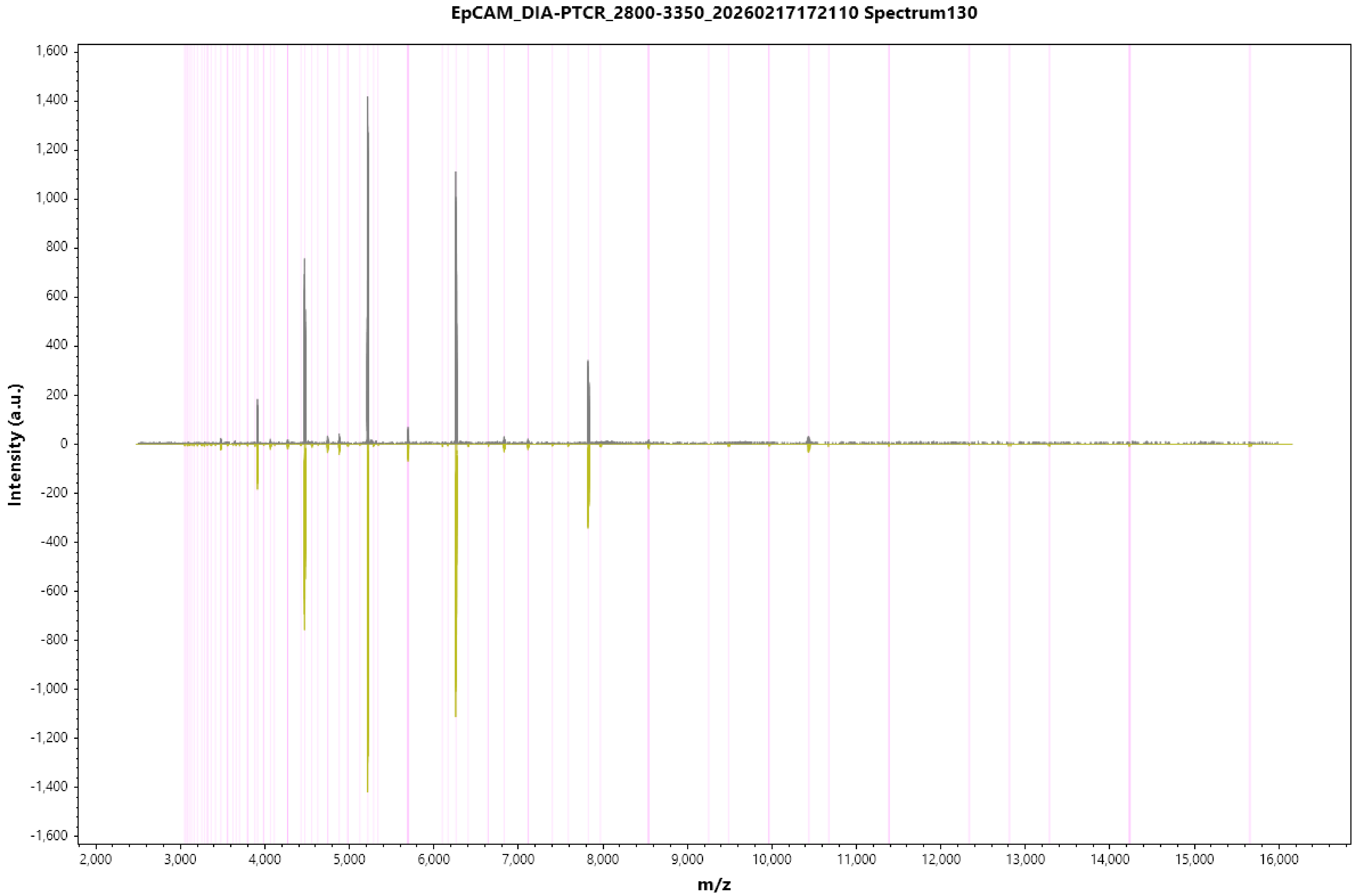
EpCAM_DIA-PTCR_2800-3350_20260217172110 Spectrum130
Intensity (a.u.)
m/z
1,600
1,400
1,200
1,000
800
600
400
200
0
-200
-400
-600
-800
-1,000
-1,200
-1,400
-1,600
2,000
3,000
4,000
5,000
6,000
7,000
8,000
9,000
10,000
11,000
12,000
13,000
14,000
15,000
16,000

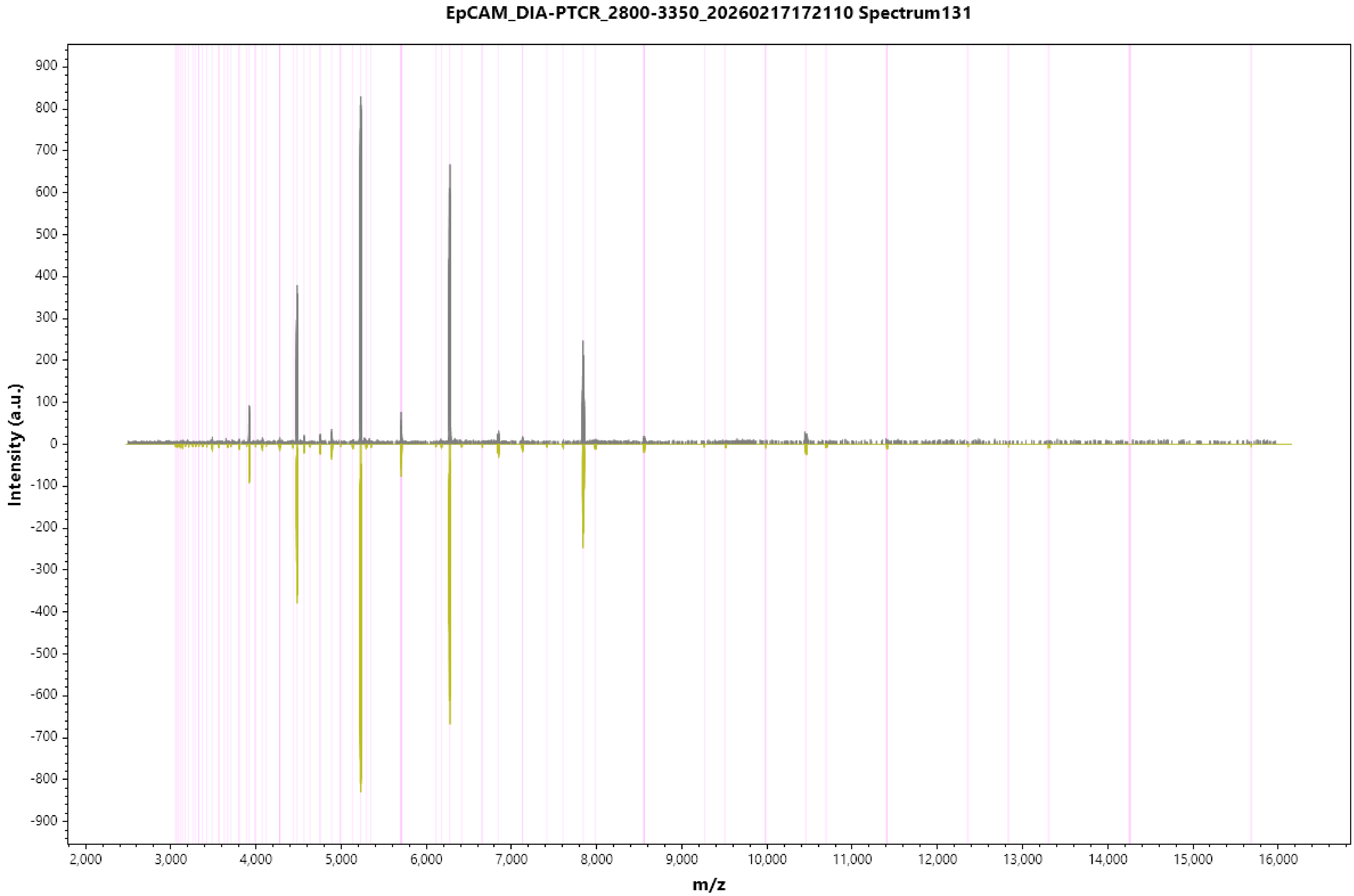

EpCAM_DIA-PTCR_2800-3350_20260217172110 Spectrum131
Intensity (a.u.)
900
800
700
600
500
400
300
200
100
0
-100
-200
-300
-400
-500
-600
-700
-800
-900
2,000
3,000
4,000
5,000
6,000
7,000
8,000
9,000
10,000
11,000
12,000
13,000
14,000
15,000
16,000
m/z

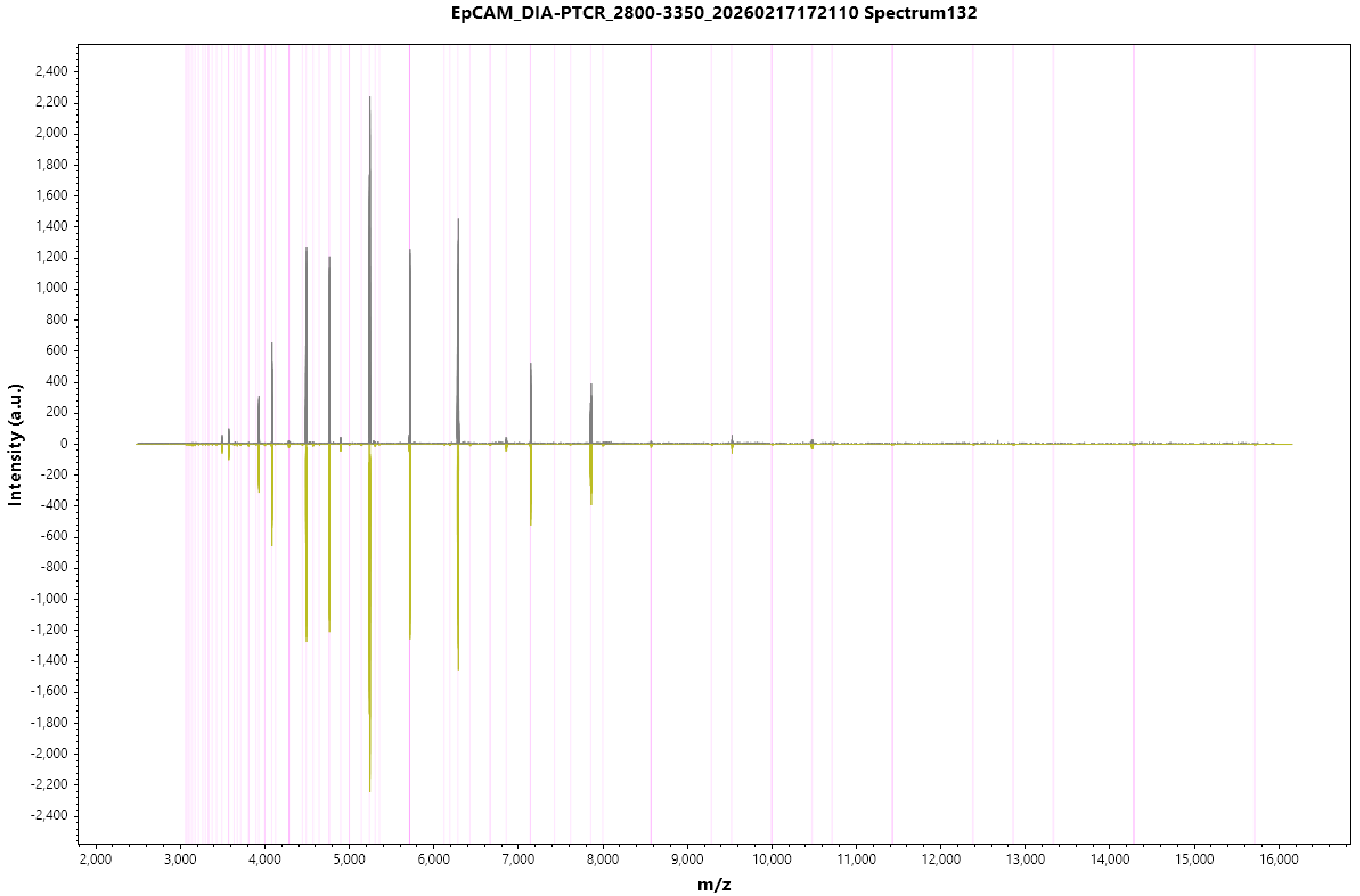
EpCAM_DIA-PTCR_2800-3350_20260217172110 Spectrum132
Intensity (a.u.)
2,400
2,200
2,000
1,800
1,600
1,400
1,200
1,000
800
600
400
200
0
-200
-400
-600
-800
-1,000
-1,200
-1,400
-1,600
-1,800
-2,000
-2,200
-2,400
2,000
3,000
4,000
5,000
6,000
7,000
8,000
9,000
10,000
11,000
12,000
13,000
14,000
15,000
16,000
m/z

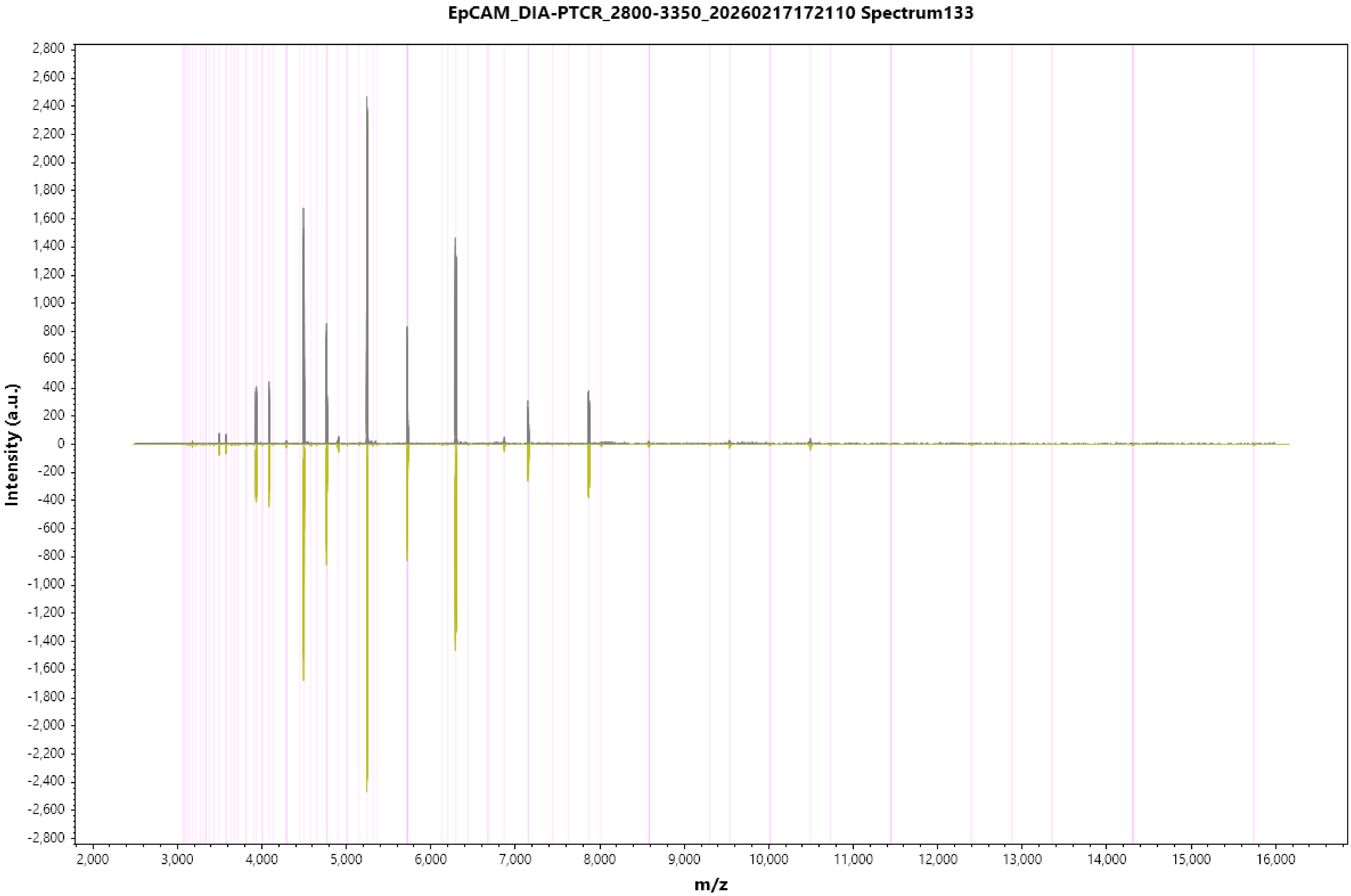
EpCAM_DIA-PTCR_2800-3350_20260217172110 Spectrum133
Intensity (a.u.)
m/z
2,800
2,600
2,400
2,200
2,000
1,800
1,600
1,400
1,200
1,000
800
600
400
200
0
-200
-400
-600
-800
-1,000
-1,200
-1,400
-1,600
-1,800
-2,000
-2,200
-2,400
-2,600
-2,800
2,000
3,000
4,000
5,000
6,000
7,000
8,000
9,000
10,000
11,000
12,000
13,000
14,000
15,000
16,000

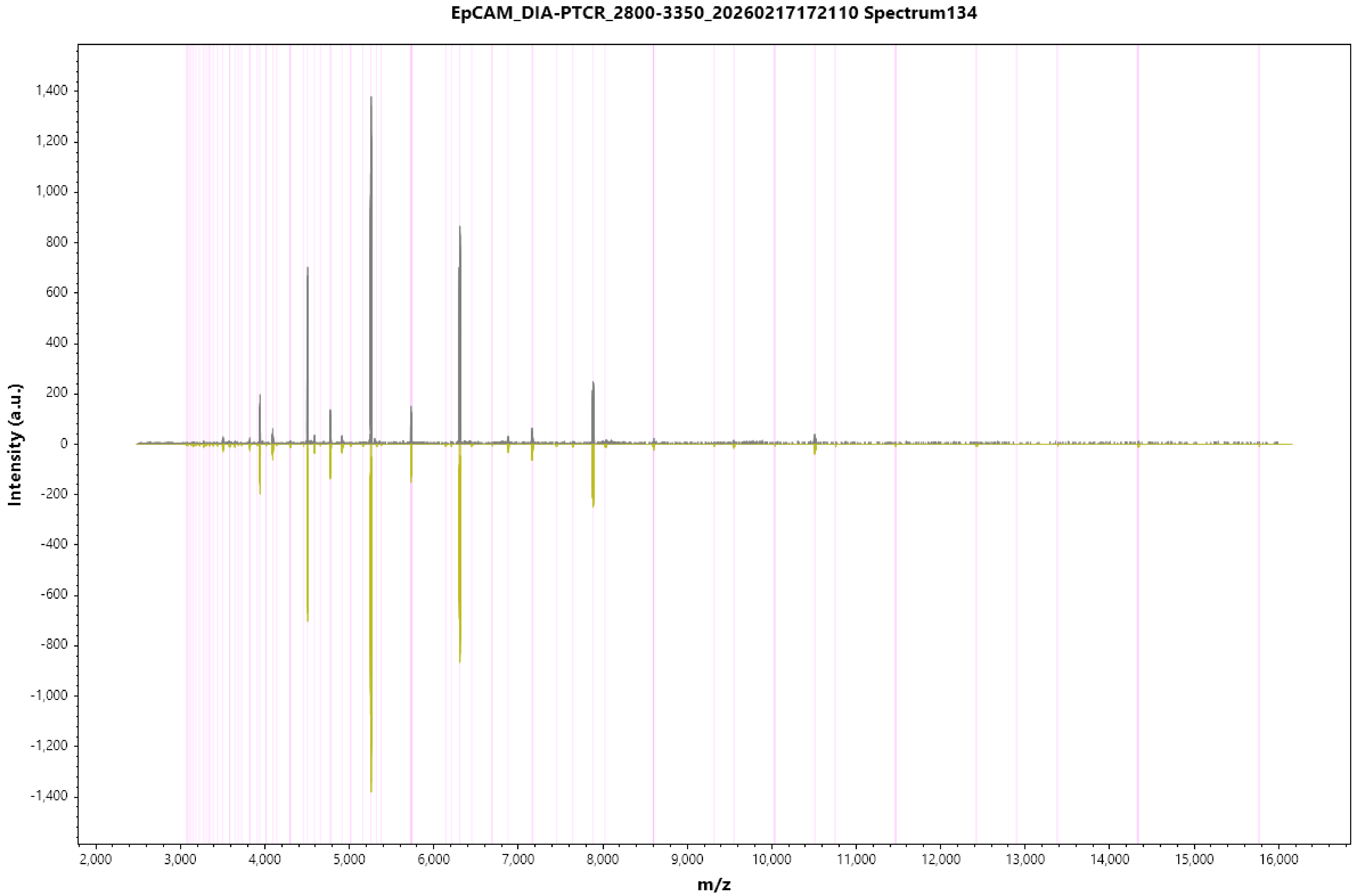

EpCAM_DIA-PTCR_2800-3350_20260217172110 Spectrum134
Intensity (a.u.)
1,400
1,200
1,000
800
600
400
200
0
-200
-400
-600
-800
-1,000
-1,200
-1,400
2,000
3,000
4,000
5,000
6,000
7,000
8,000
9,000
10,000
11,000
12,000
13,000
14,000
15,000
16,000
m/z

EpCAM_DIA-PTCR_2800-3350_20260217172110 Spectrum135

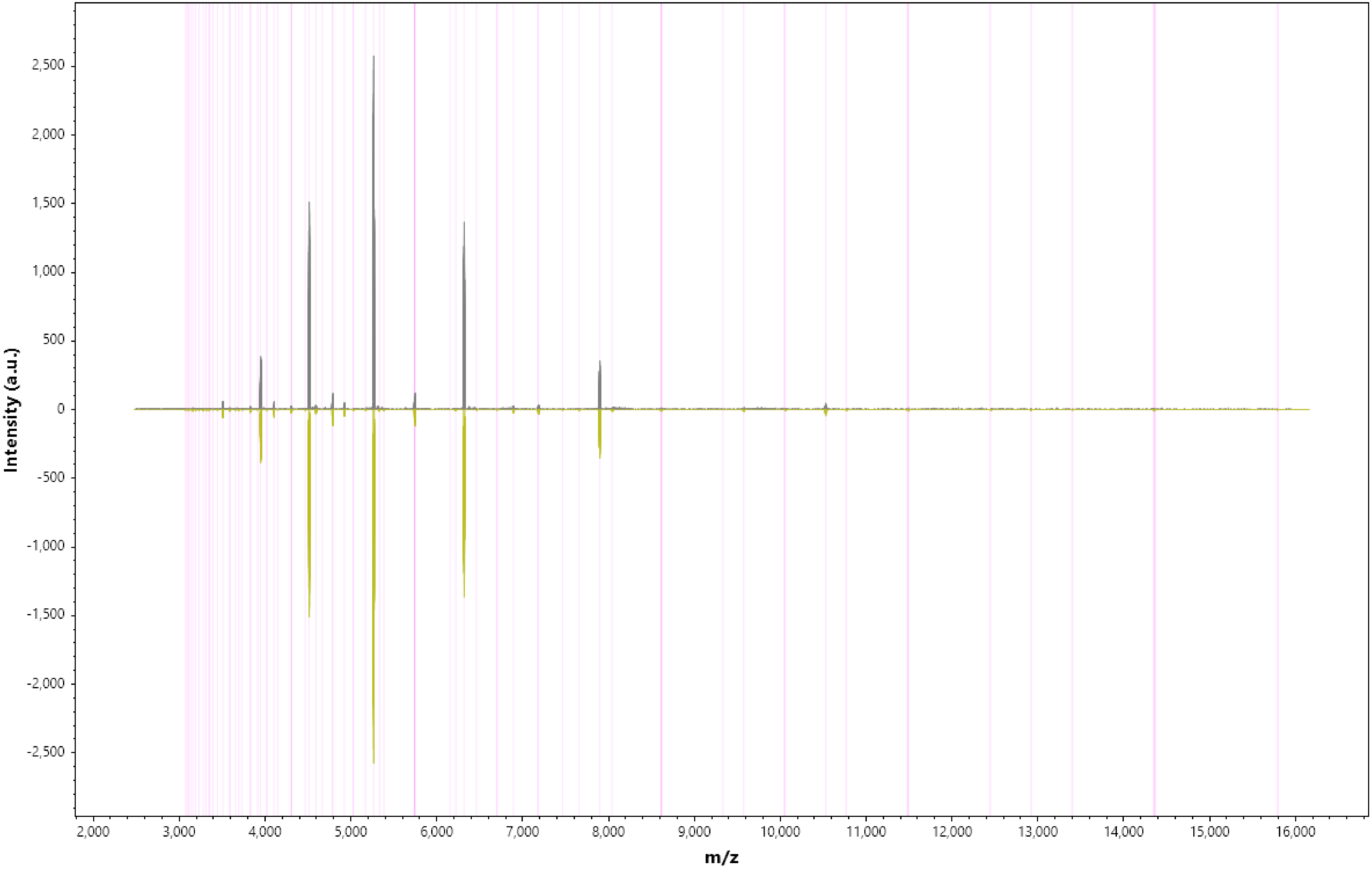

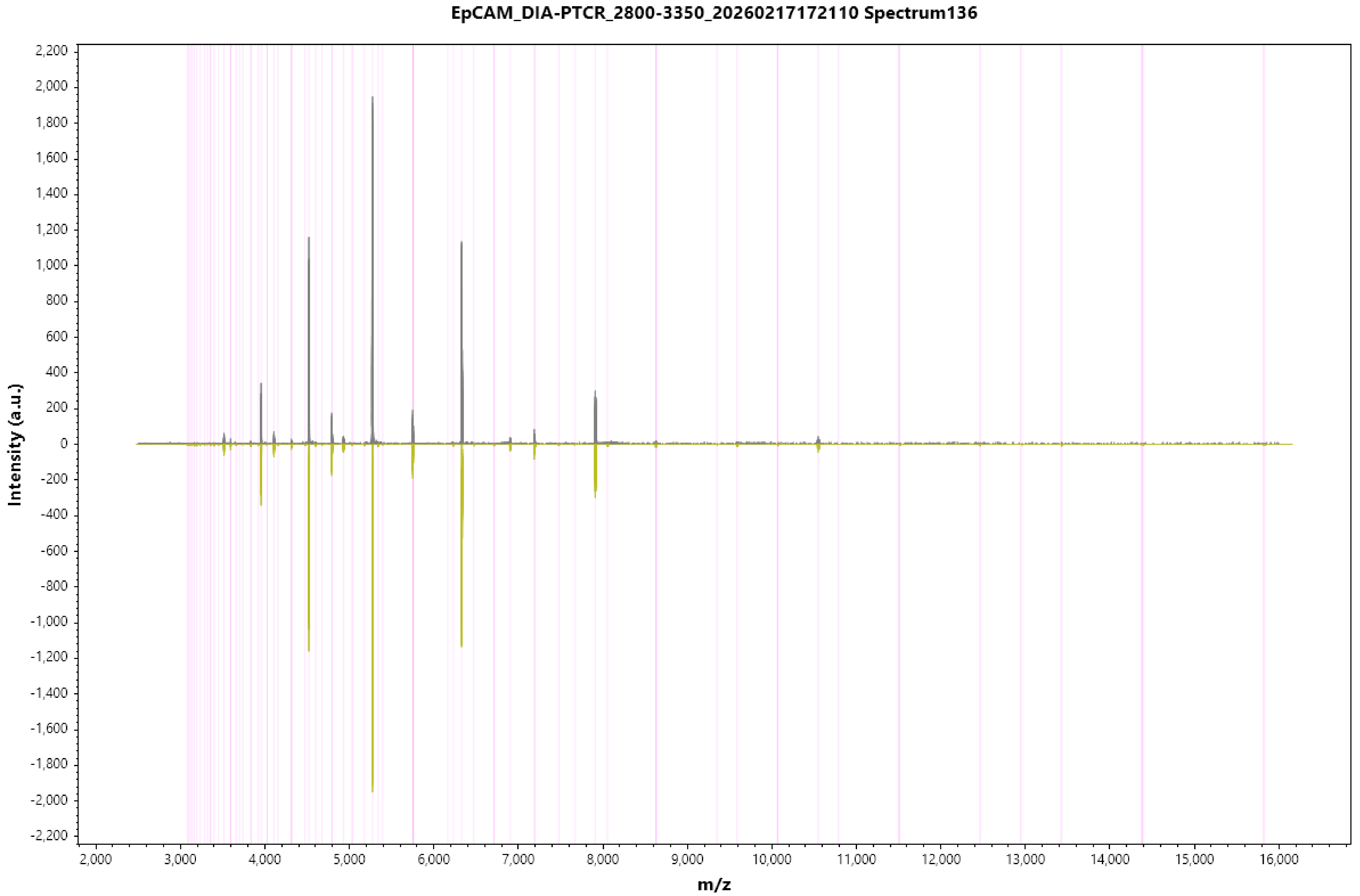
EpCAM_DIA-PTCR_2800-3350_20260217172110 Spectrum136
Intensity (a.u.)
2,200
2,000
1,800
1,600
1,400
1,200
1,000
800
600
400
200
0
-200
-400
-600
-800
-1,000
-1,200
-1,400
-1,600
-1,800
-2,000
-2,200
2,000
3,000
4,000
5,000
6,000
7,000
8,000
9,000
10,000
11,000
12,000
13,000
14,000
15,000
16,000
m/z

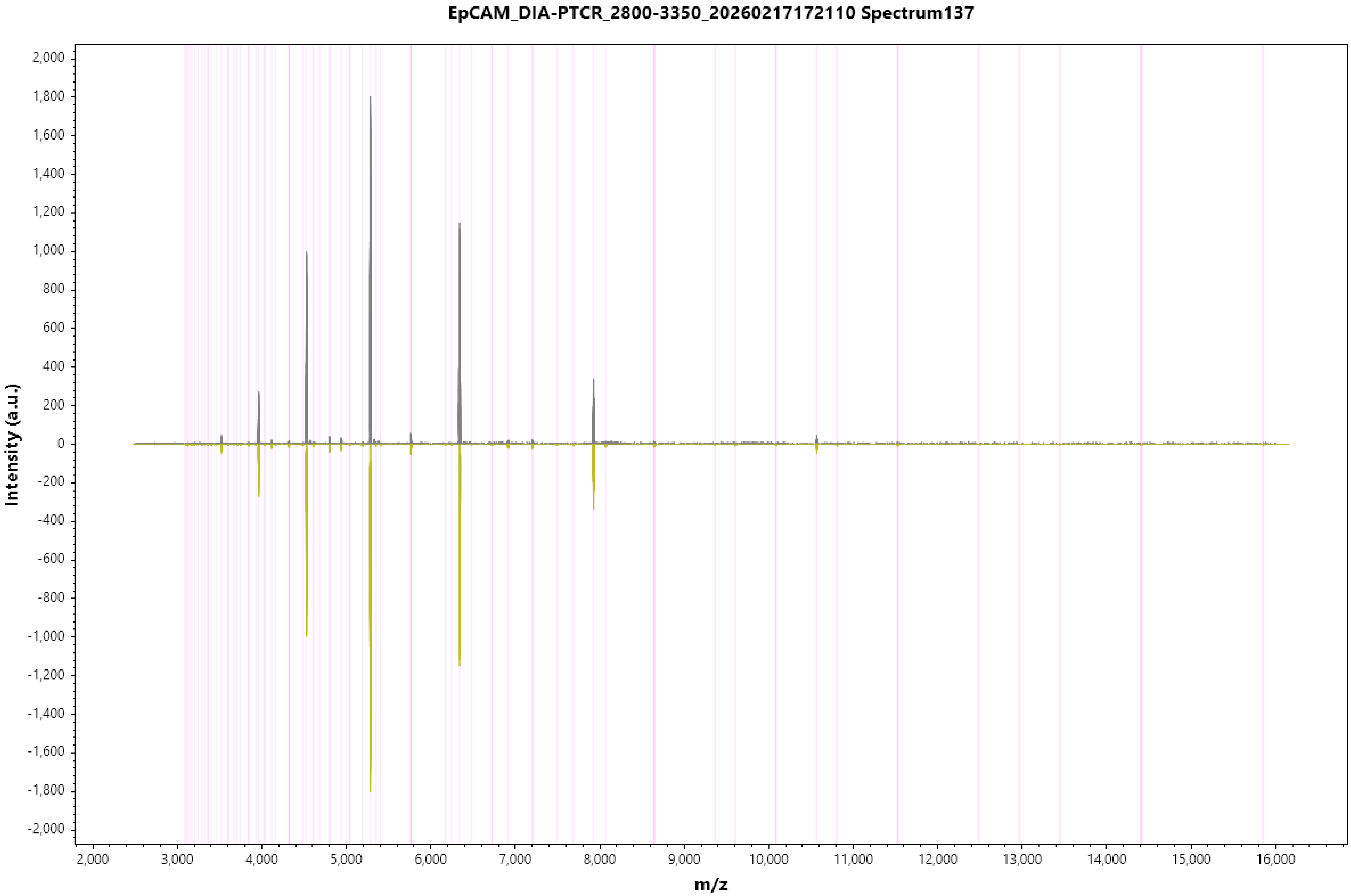
EpCAM_DIA-PTCR_2800-3350_20260217172110 Spectrum137
Intensity (a.u.)
m/z
2,000
1,800
1,600
1,400
1,200
1,000
800
600
400
200
0
-200
-400
-600
-800
-1,000
-1,200
-1,400
-1,600
-1,800
-2,000
2,000
3,000
4,000
5,000
6,000
7,000
8,000
9,000
10,000
11,000
12,000
13,000
14,000
15,000
16,000

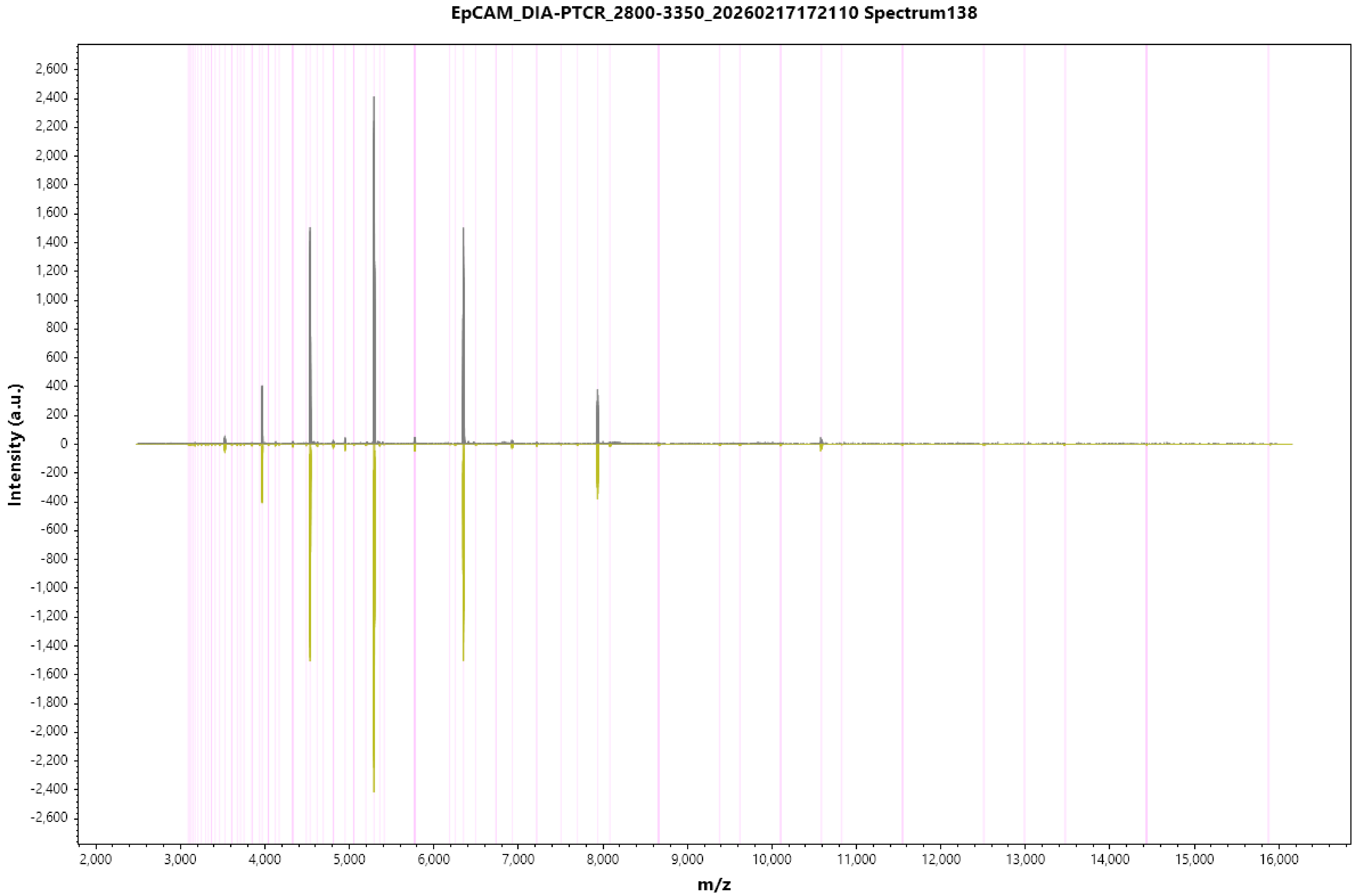
EpCAM_DIA-PTCR_2800-3350_20260217172110 Spectrum138
Intensity (a.u.)
m/z
2,600
2,400
2,200
2,000
1,800
1,600
1,400
1,200
1,000
800
600
400
200
0
-200
-400
-600
-800
-1,000
-1,200
-1,400
-1,600
-1,800
-2,000
-2,200
-2,400
-2,600
2,000
3,000
4,000
5,000
6,000
7,000
8,000
9,000
10,000
11,000
12,000
13,000
14,000
15,000
16,000

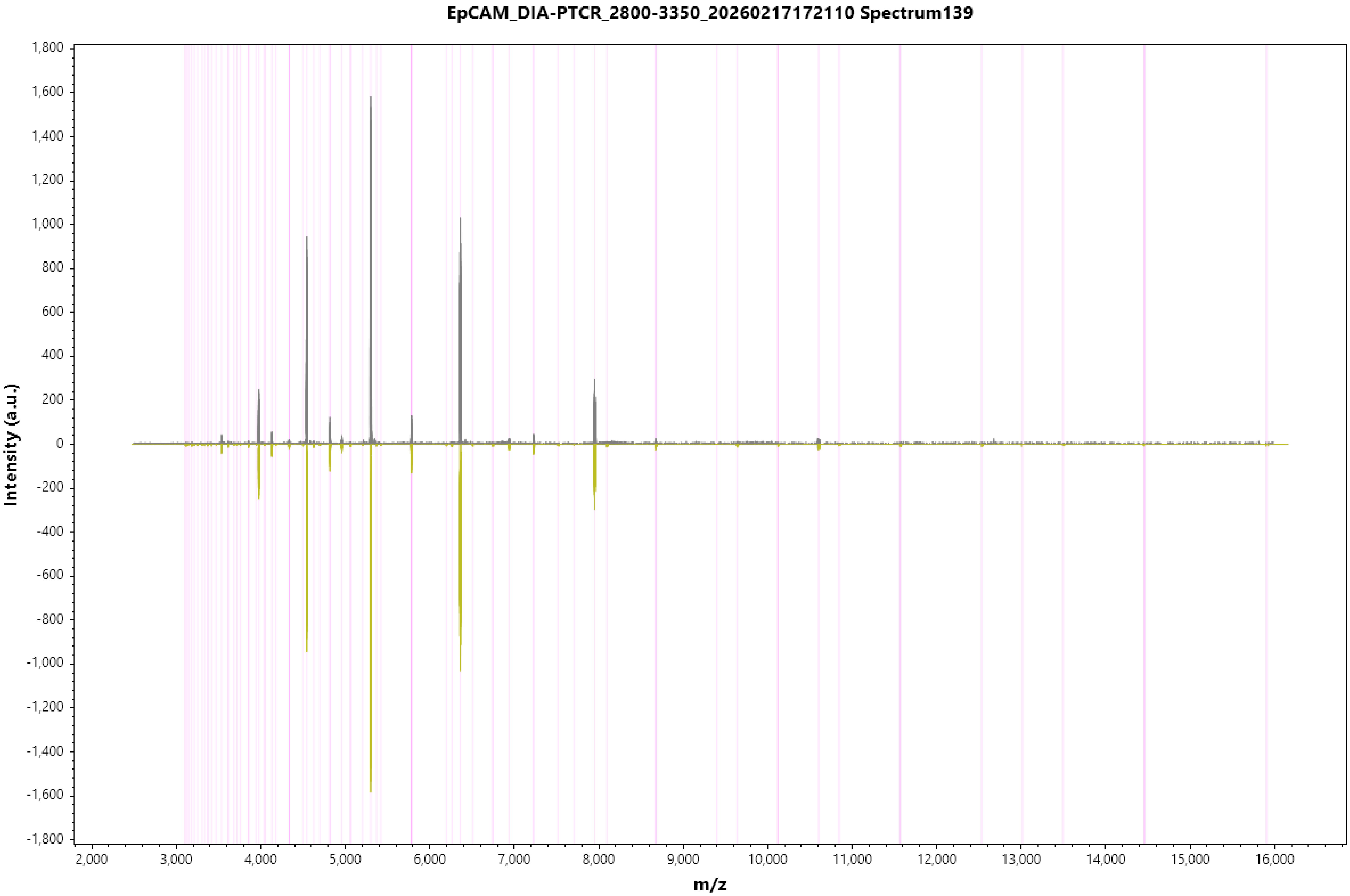
EpCAM_DIA-PTCR_2800-3350_20260217172110 Spectrum139
Intensity (a.u.)
1,800
1,600
1,400
1,200
1,000
800
600
400
200
0
-200
-400
-600
-800
-1,000
-1,200
-1,400
-1,600
-1,800
2,000
3,000
4,000
5,000
6,000
7,000
8,000
9,000
10,000
11,000
12,000
13,000
14,000
15,000
16,000
m/z

#### EpCAM_DIA-PTCR_2800-3350_20260217172110 Spectrum140

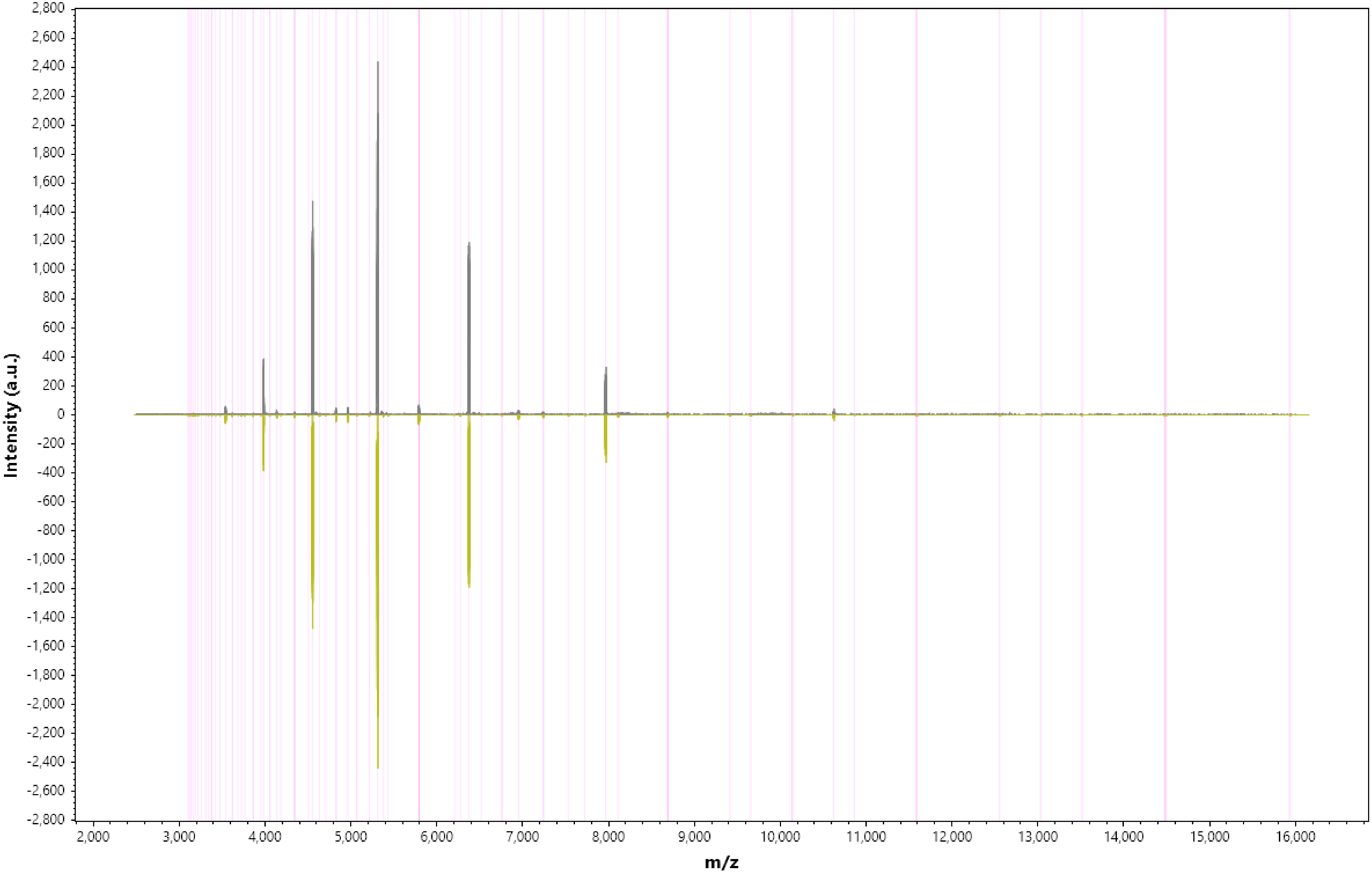

EpCAM_DIA-PTCR_2800-3350_20260217172110 Spectrum141

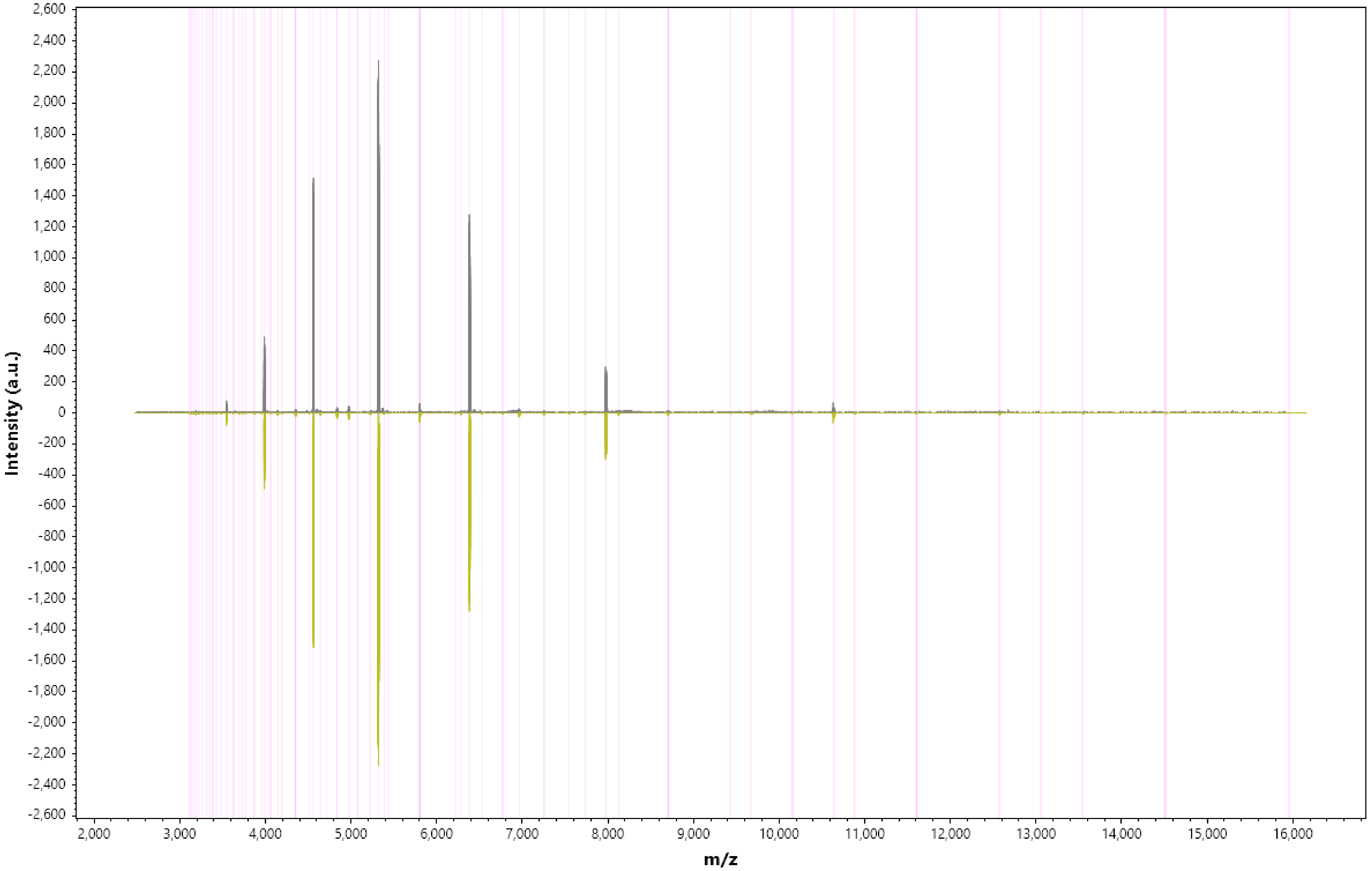

EpCAM_DIA-PTCR_2800-3350_20260217172110 Spectrum142

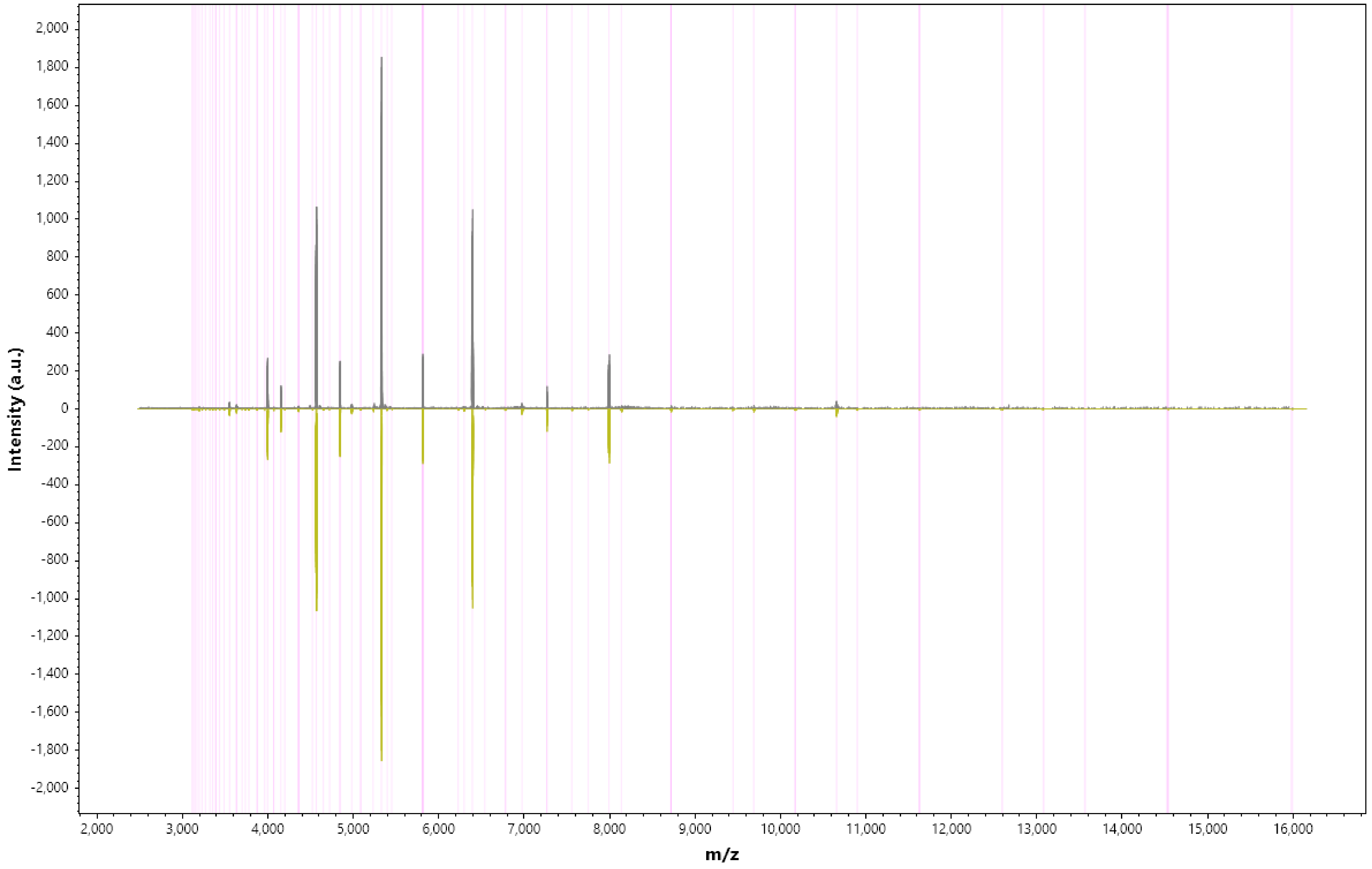

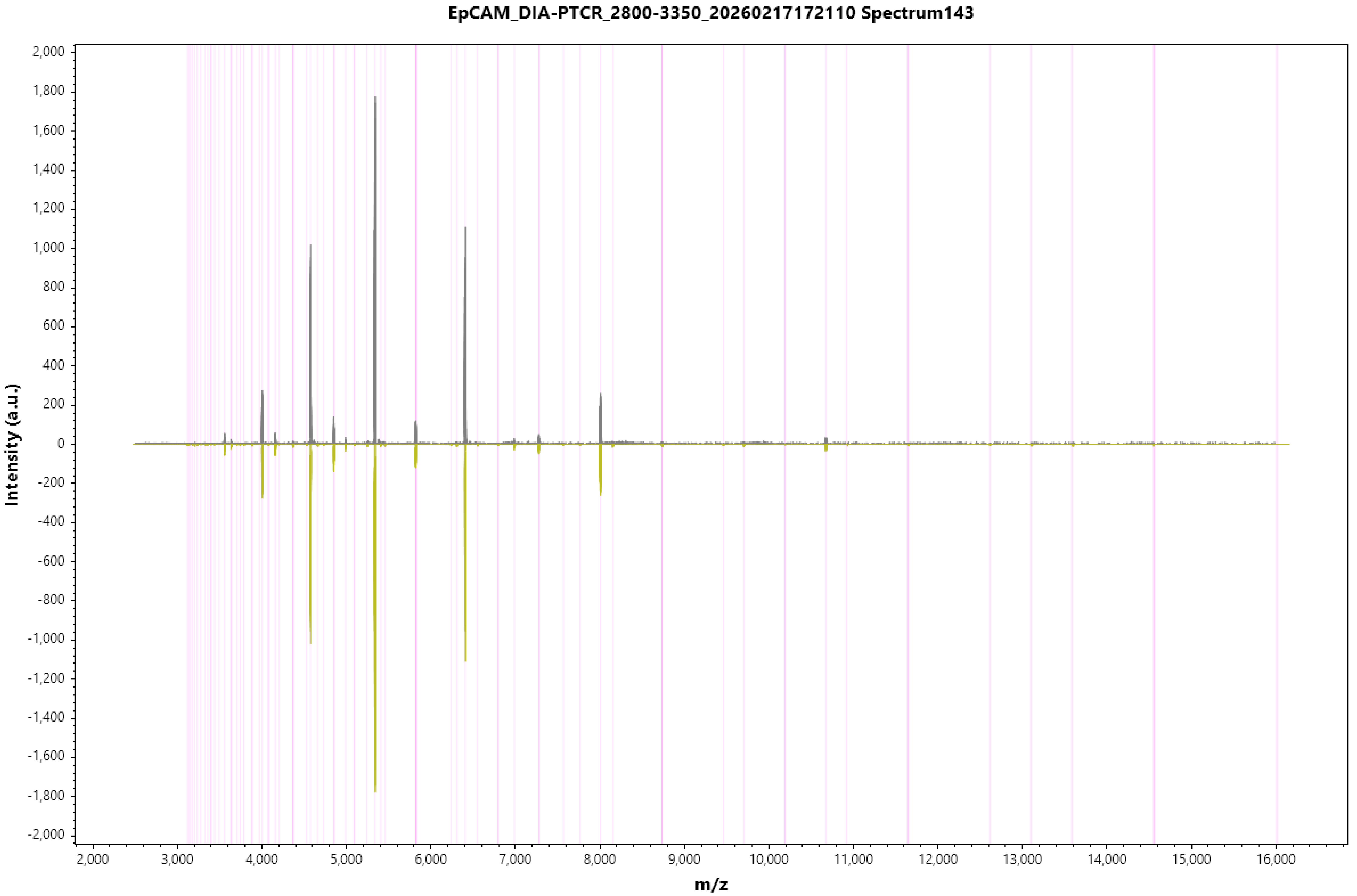
EpCAM_DIA-PTCR_2800-3350_20260217172110 Spectrum143
Intensity (a.u.)
2,000
1,800
1,600
1,400
1,200
1,000
800
600
400
200
0
-200
-400
-600
-800
-1,000
-1,200
-1,400
-1,600
-1,800
-2,000
2,000
3,000
4,000
5,000
6,000
7,000
8,000
9,000
10,000
11,000
12,000
13,000
14,000
15,000
16,000
m/z

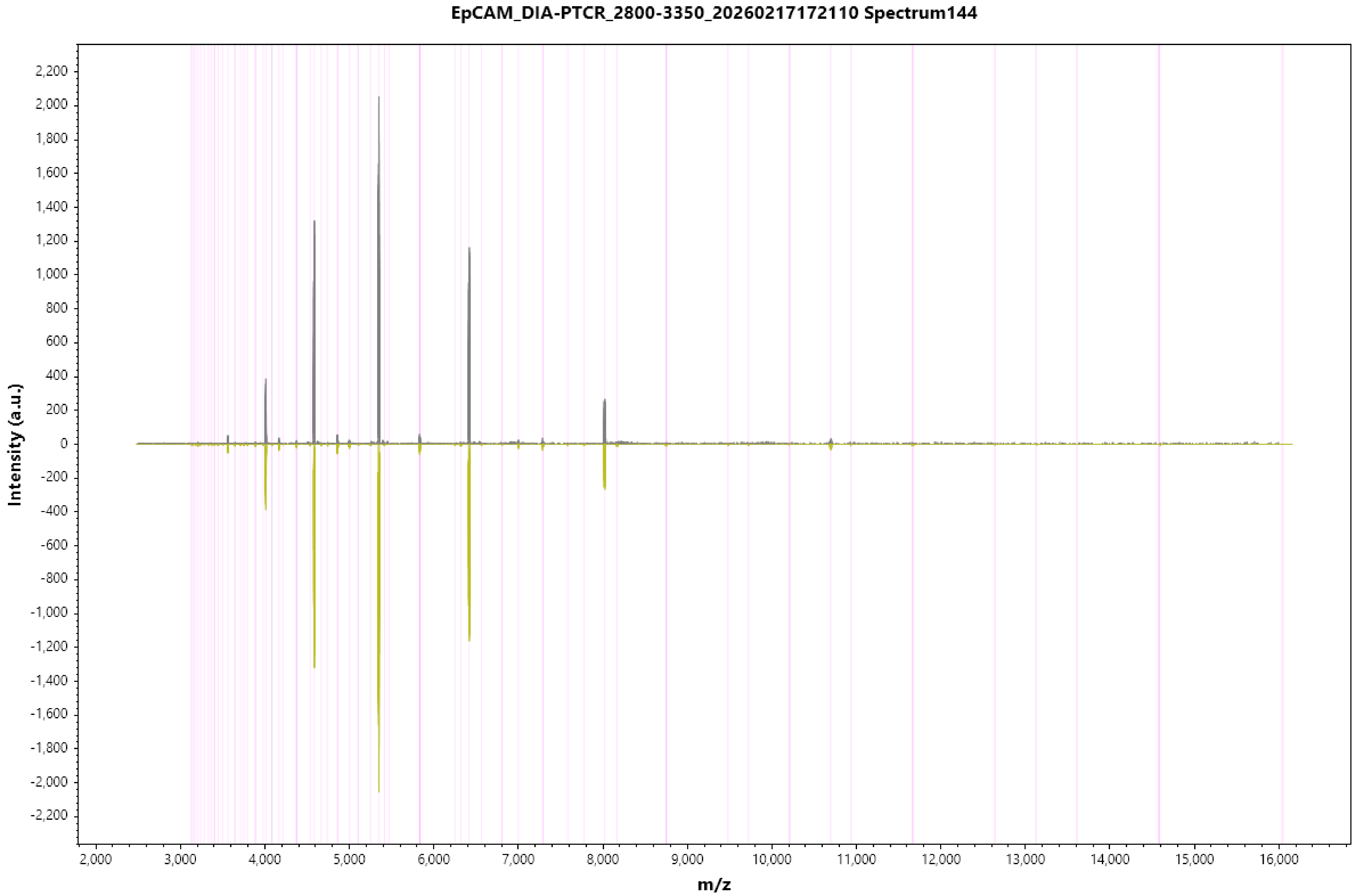
EpCAM_DIA-PTCR_2800-3350_20260217172110 Spectrum144
Intensity (a.u.)
m/z
2,200
2,000
1,800
1,600
1,400
1,200
1,000
800
600
400
200
0
-200
-400
-600
-800
-1,000
-1,200
-1,400
-1,600
-1,800
-2,000
-2,200
2,000
3,000
4,000
5,000
6,000
7,000
8,000
9,000
10,000
11,000
12,000
13,000
14,000
15,000
16,000

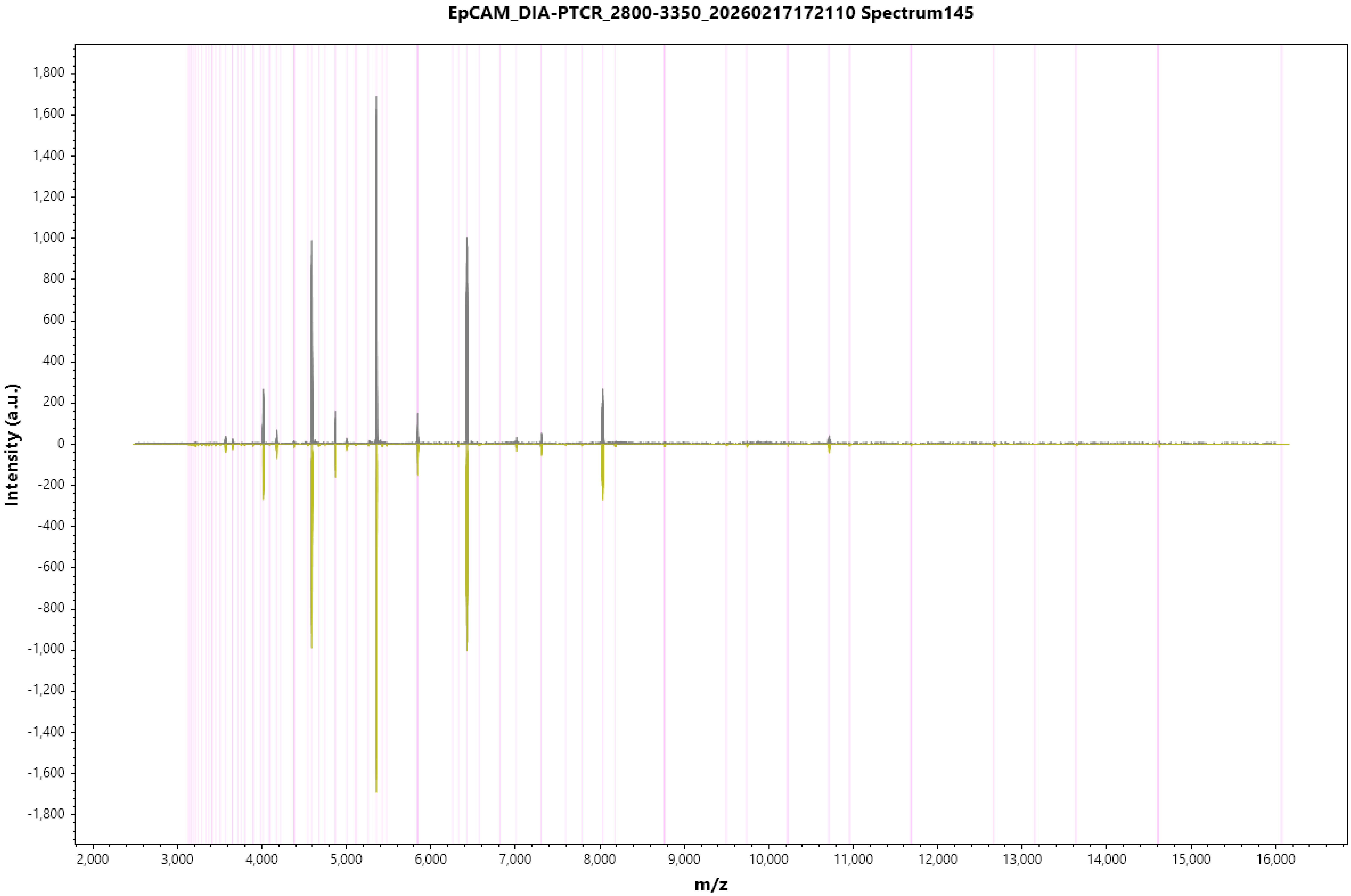
EpCAM_DIA-PTCR_2800-3350_20260217172110 Spectrum145
Intensity (a.u.)
1,800
1,600
1,400
1,200
1,000
800
600
400
200
0
-200
-400
-600
-800
-1,000
-1,200
-1,400
-1,600
-1,800
2,000
3,000
4,000
5,000
6,000
7,000
8,000
9,000
10,000
11,000
12,000
13,000
14,000
15,000
16,000
m/z

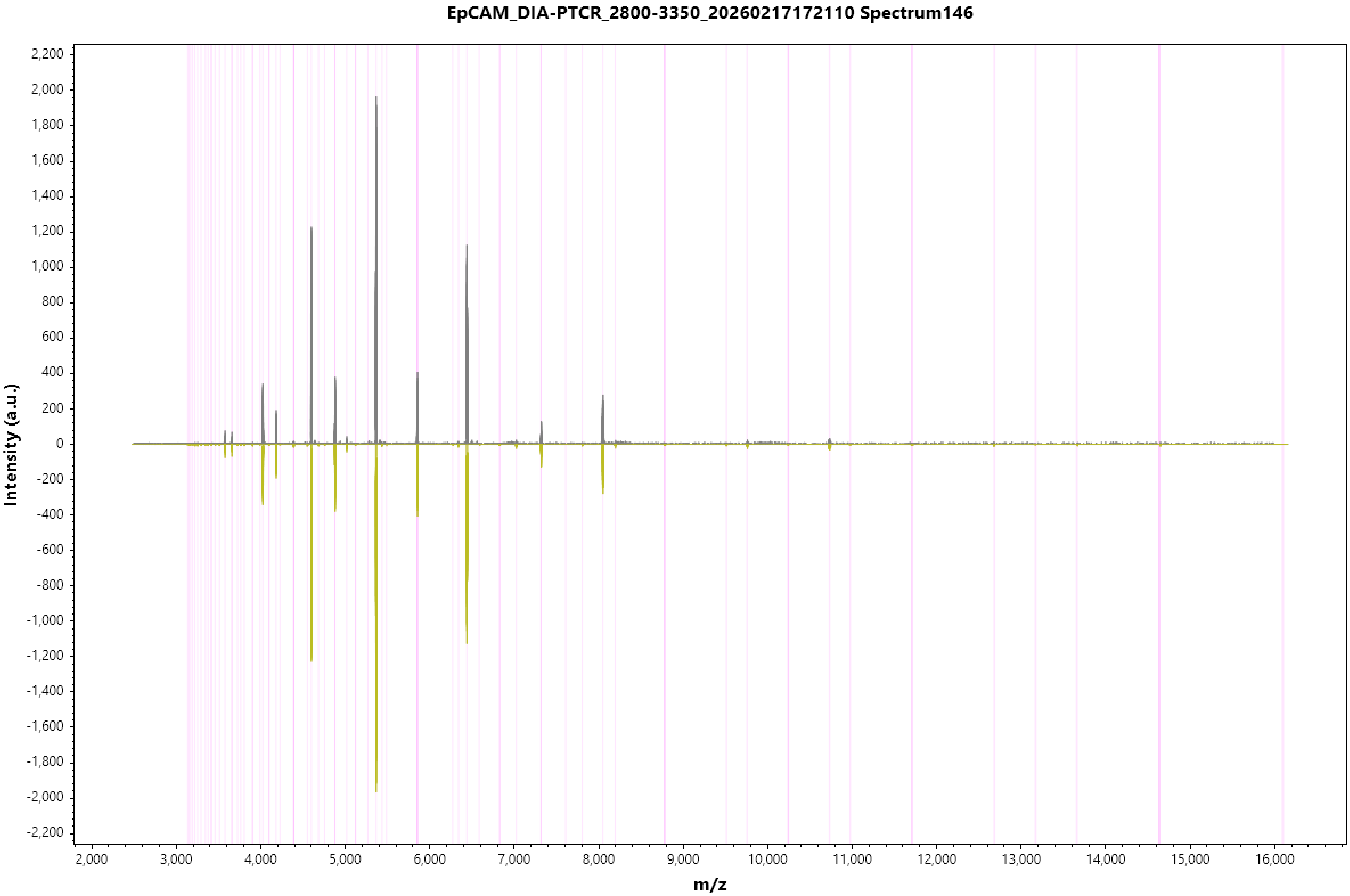
EpCAM_DIA-PTCR_2800-3350_20260217172110 Spectrum146
Intensity (a.u.)
m/z
2,200
2,000
1,800
1,600
1,400
1,200
1,000
800
600
400
200
0
-200
-400
-600
-800
-1,000
-1,200
-1,400
-1,600
-1,800
-2,000
-2,200
2,000
3,000
4,000
5,000
6,000
7,000
8,000
9,000
10,000
11,000
12,000
13,000
14,000
15,000
16,000

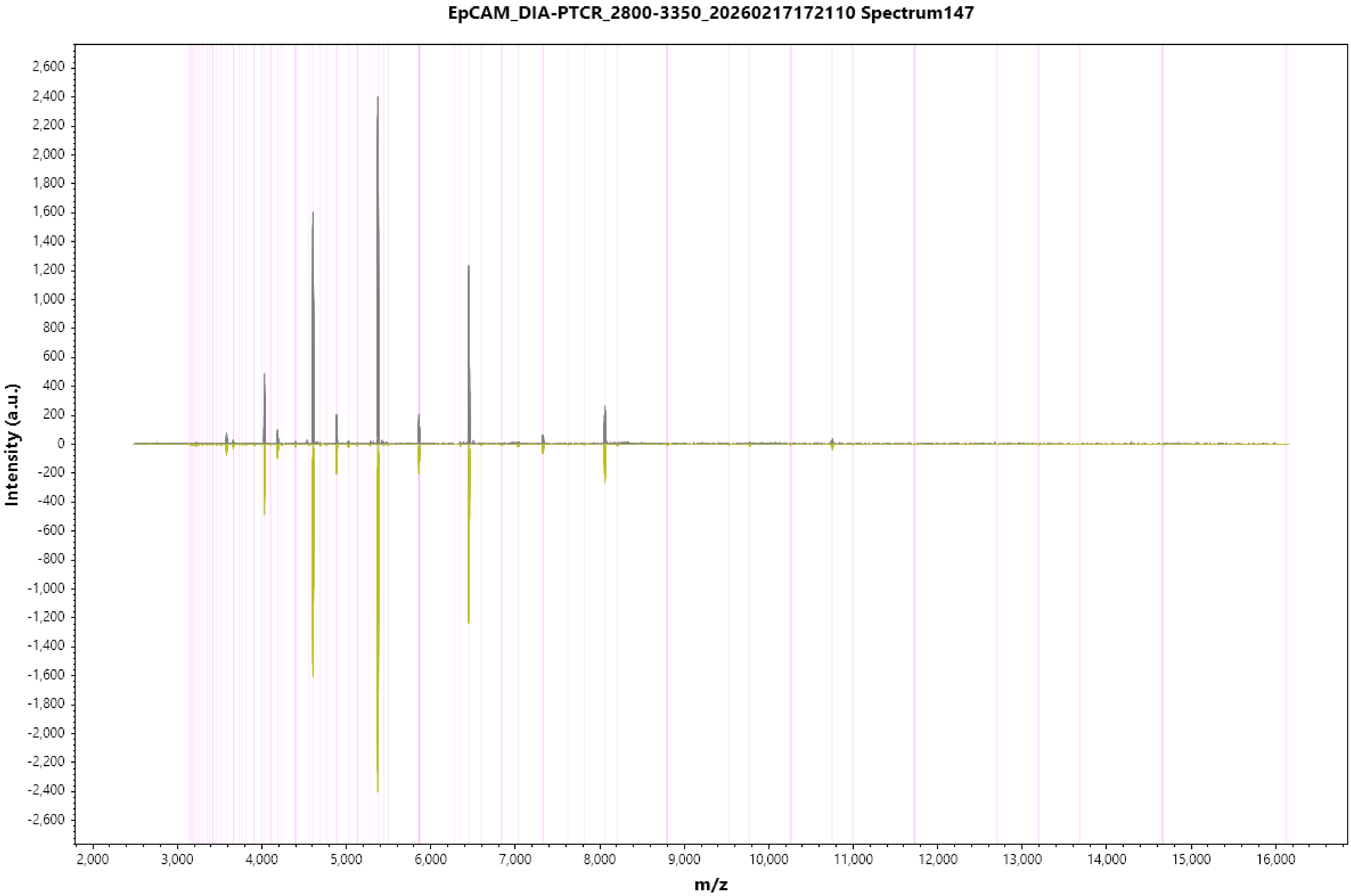
EpCAM_DIA-PTCR_2800-3350_20260217172110 Spectrum147
Intensity (a.u.)
m/z
2,600
2,400
2,200
2,000
1,800
1,600
1,400
1,200
1,000
800
600
400
200
0
-200
-400
-600
-800
-1,000
-1,200
-1,400
-1,600
-1,800
-2,000
-2,200
-2,400
-2,600
2,000
3,000
4,000
5,000
6,000
7,000
8,000
9,000
10,000
11,000
12,000
13,000
14,000
15,000
16,000

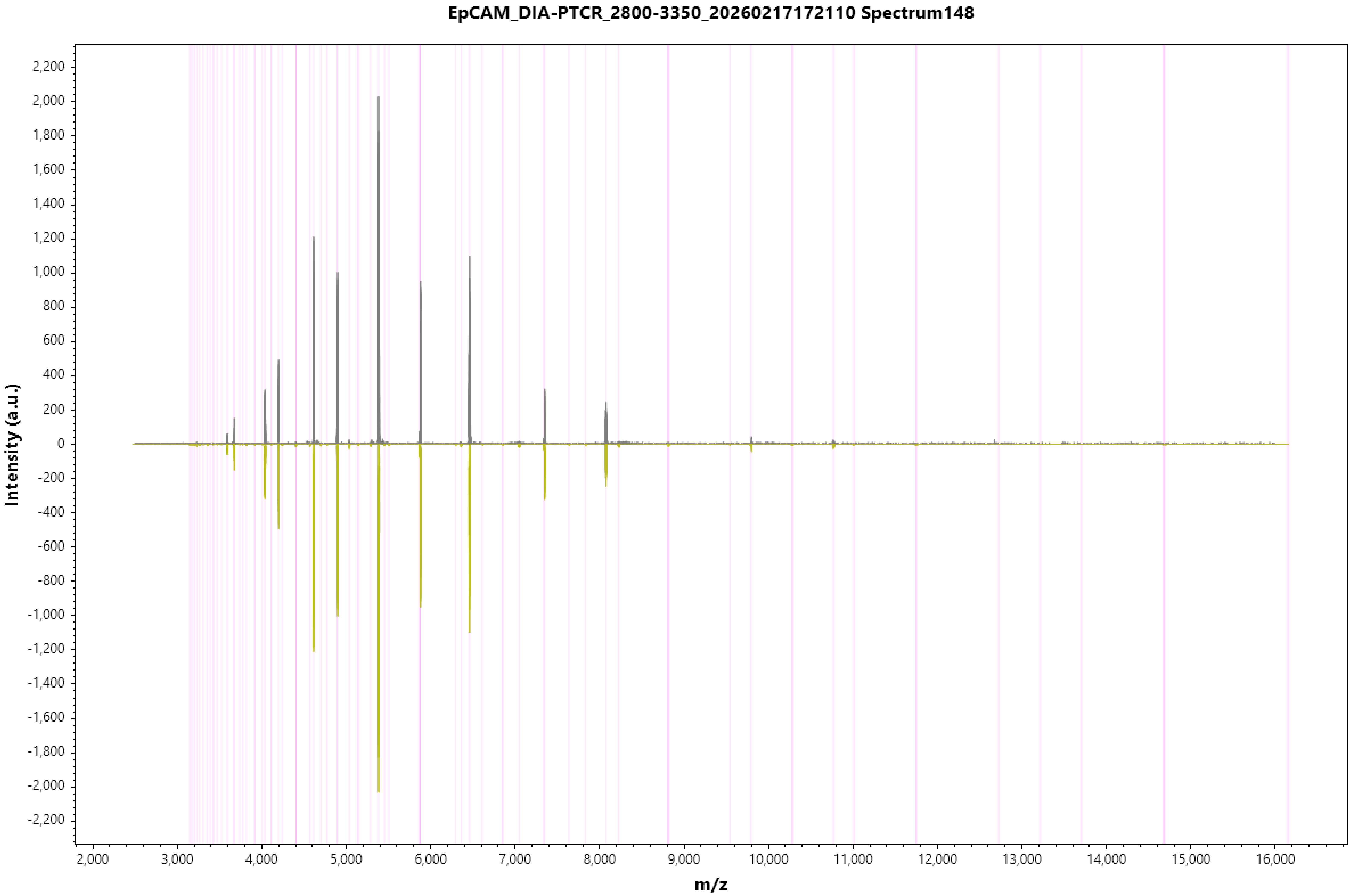
EpCAM_DIA-PTCR_2800-3350_20260217172110 Spectrum148
Intensity (a.u.)
m/z
2,200
2,000
1,800
1,600
1,400
1,200
1,000
800
600
400
200
0
-200
-400
-600
-800
-1,000
-1,200
-1,400
-1,600
-1,800
-2,000
-2,200
2,000
3,000
4,000
5,000
6,000
7,000
8,000
9,000
10,000
11,000
12,000
13,000
14,000
15,000
16,000

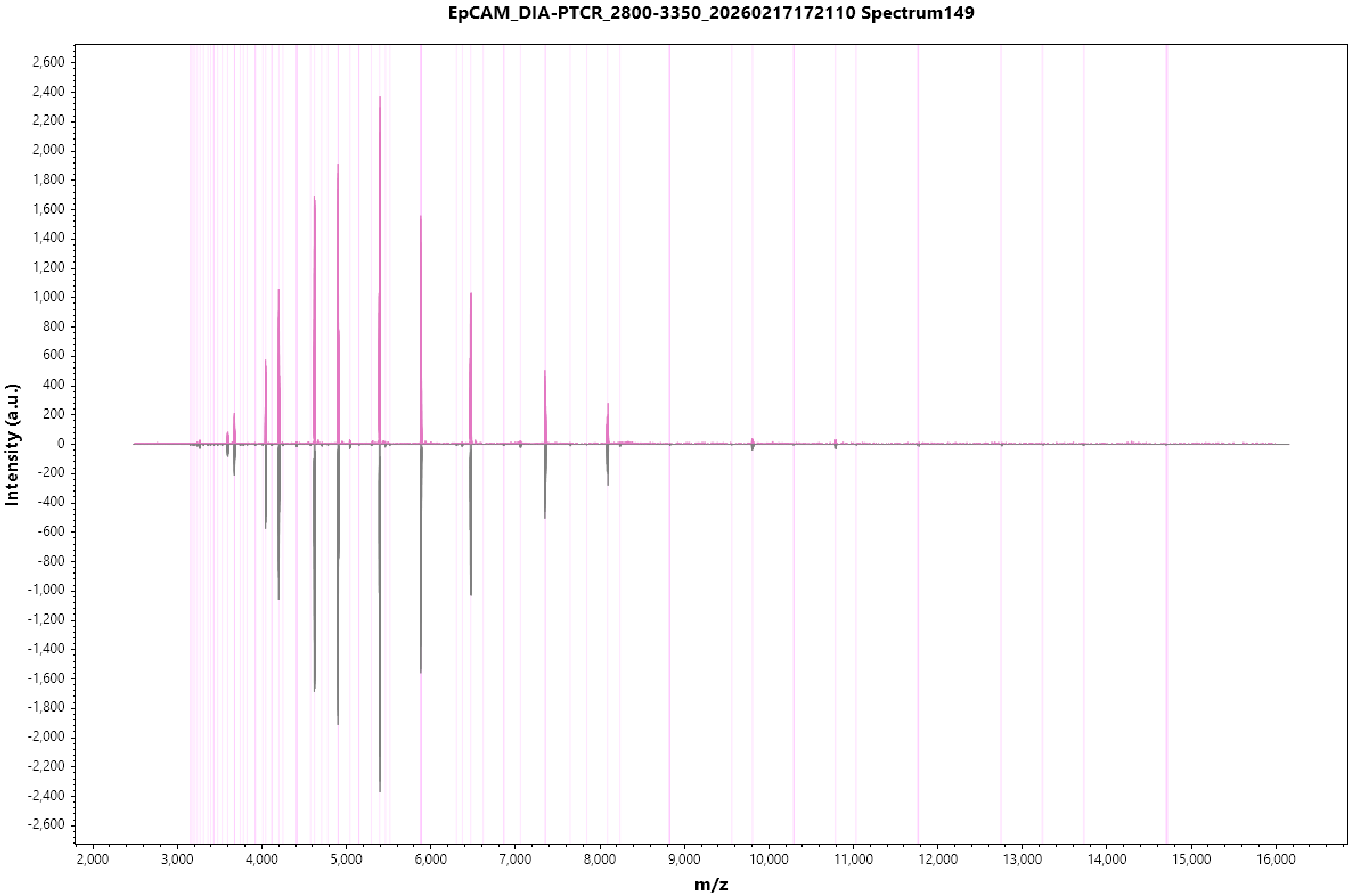
EpCAM_DIA-PTCR_2800-3350_20260217172110 Spectrum149
Intensity (a.u.)
m/z
2,600
2,400
2,200
2,000
1,800
1,600
1,400
1,200
1,000
800
600
400
200
0
-200
-400
-600
-800
-1,000
-1,200
-1,400
-1,600
-1,800
-2,000
-2,200
-2,400
-2,600
2,000
3,000
4,000
5,000
6,000
7,000
8,000
9,000
10,000
11,000
12,000
13,000
14,000
15,000
16,000

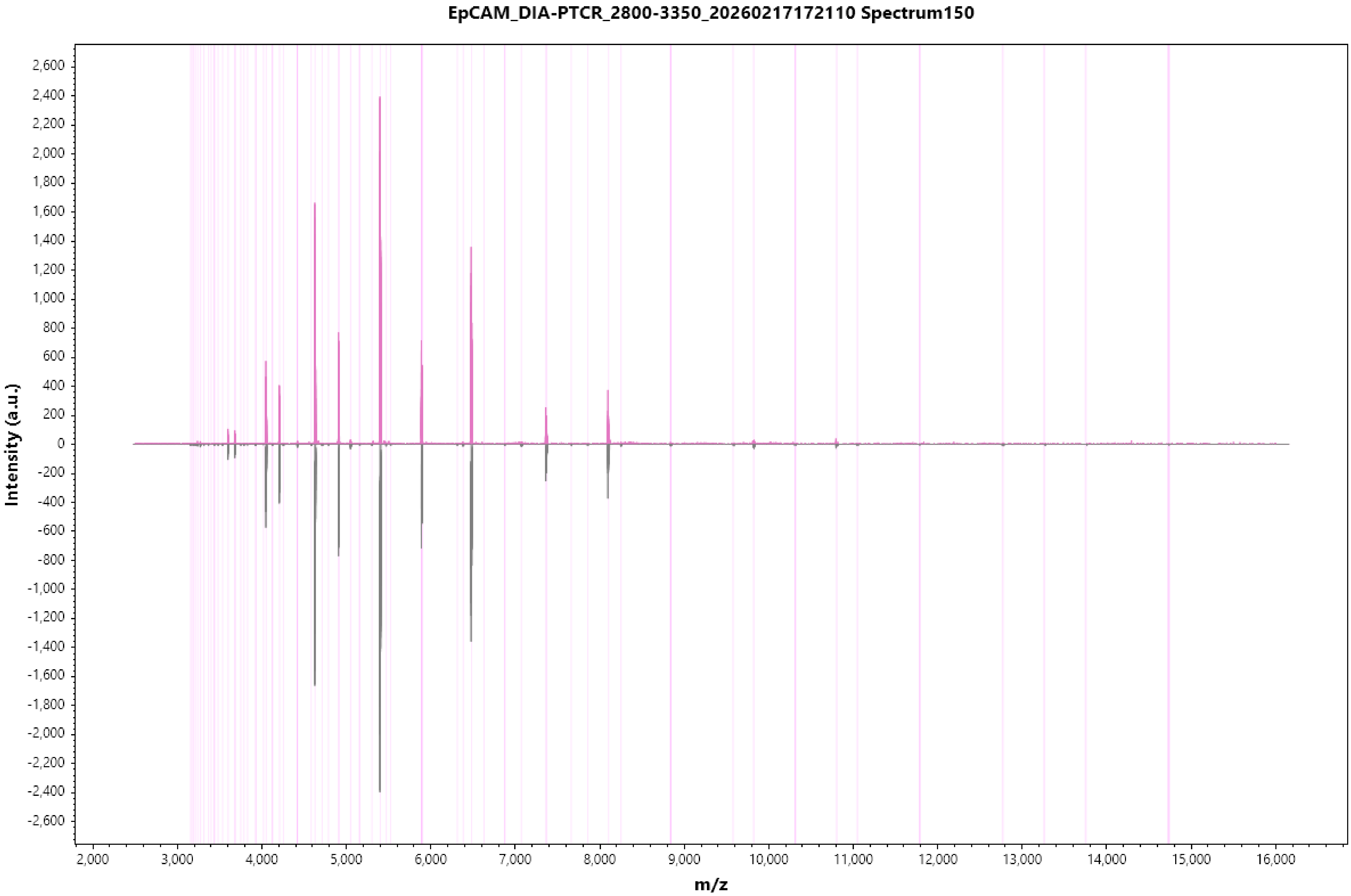
EpCAM_DIA-PTCR_2800-3350_20260217172110 Spectrum150
Intensity (a.u.)
2,600
2,400
2,200
2,000
1,800
1,600
1,400
1,200
1,000
800
600
400
200
0
-200
-400
-600
-800
-1,000
-1,200
-1,400
-1,600
-1,800
-2,000
-2,200
-2,400
-2,600
2,000
3,000
4,000
5,000
6,000
7,000
8,000
9,000
10,000
11,000
12,000
13,000
14,000
15,000
16,000
m/z

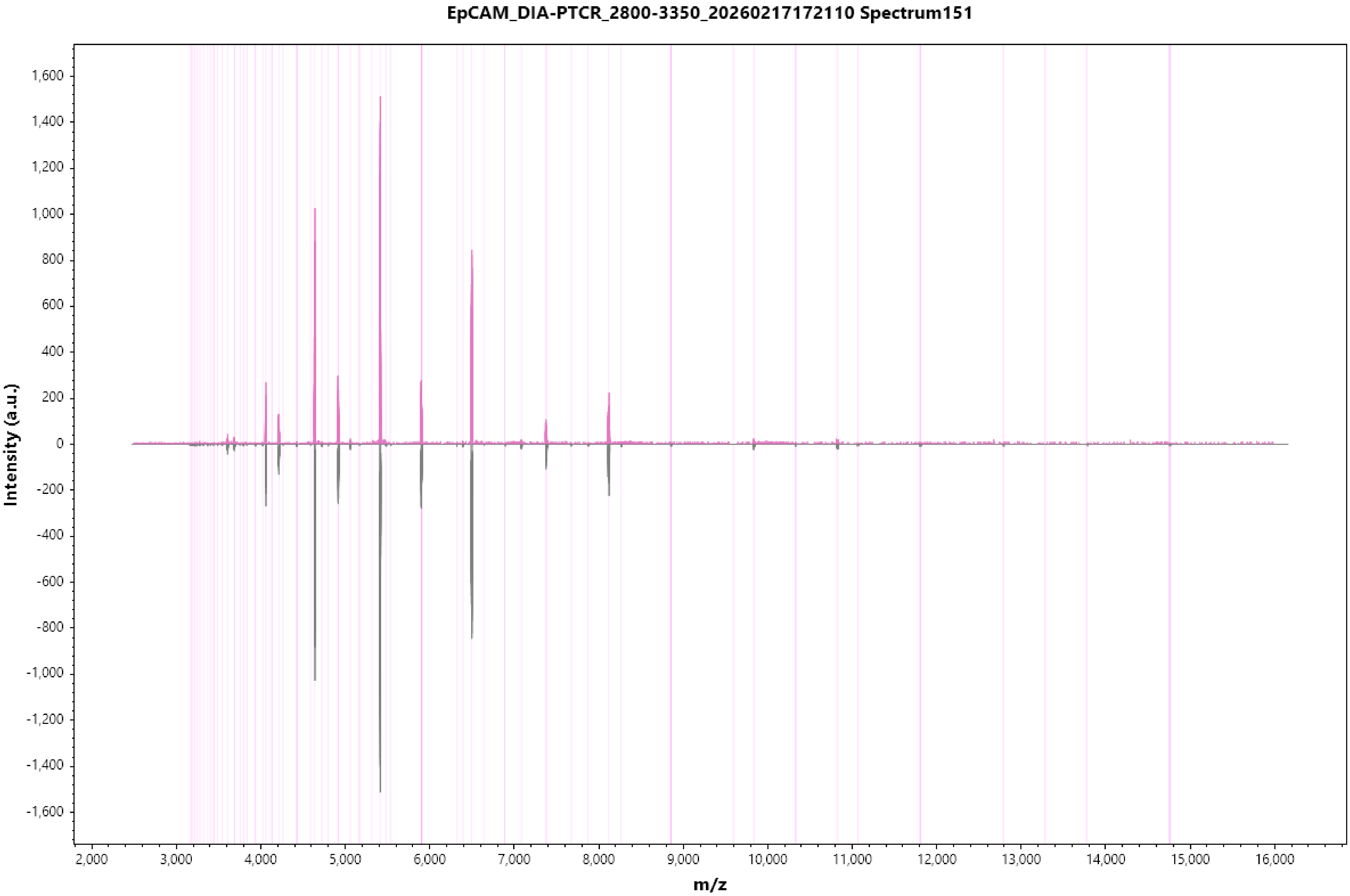
EpCAM_DIA-PTCR_2800-3350_20260217172110 Spectrum151
Intensity (a.u.)
1,600
1,400
1,200
1,000
800
600
400
200
0
-200
-400
-600
-800
-1,000
-1,200
-1,400
-1,600
2,000
3,000
4,000
5,000
6,000
7,000
8,000
9,000
10,000
11,000
12,000
13,000
14,000
15,000
16,000
m/z

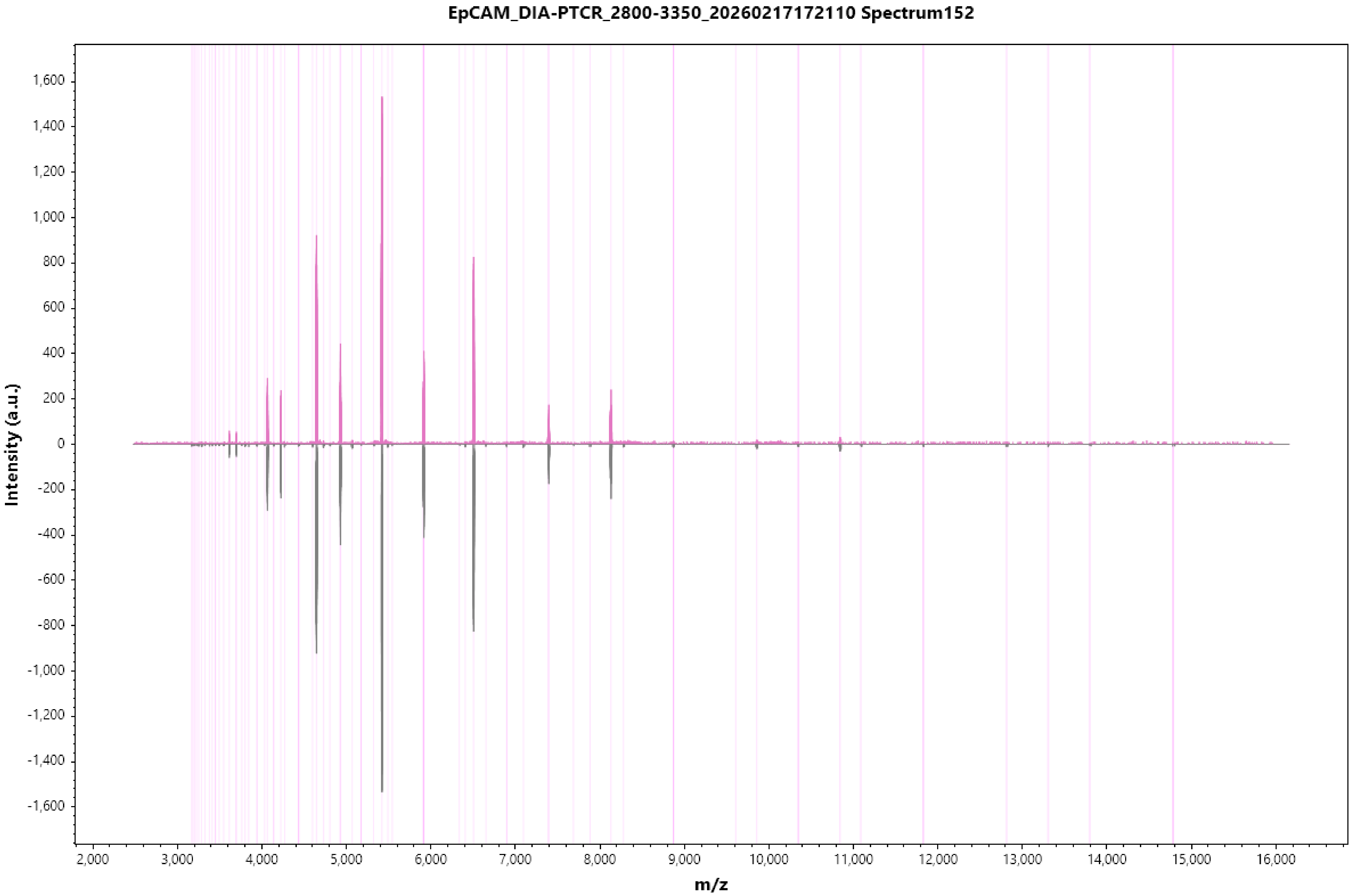
EpCAM_DIA-PTCR_2800-3350_20260217172110 Spectrum152
Intensity (a.u.)
1,600
1,400
1,200
1,000
800
600
400
200
0
-200
-400
-600
-800
-1,000
-1,200
-1,400
-1,600
2,000
3,000
4,000
5,000
6,000
7,000
8,000
9,000
10,000
11,000
12,000
13,000
14,000
15,000
16,000
m/z

#### EpCAM_DIA-PTCR_2800-3350_20260217172110 Spectrum153

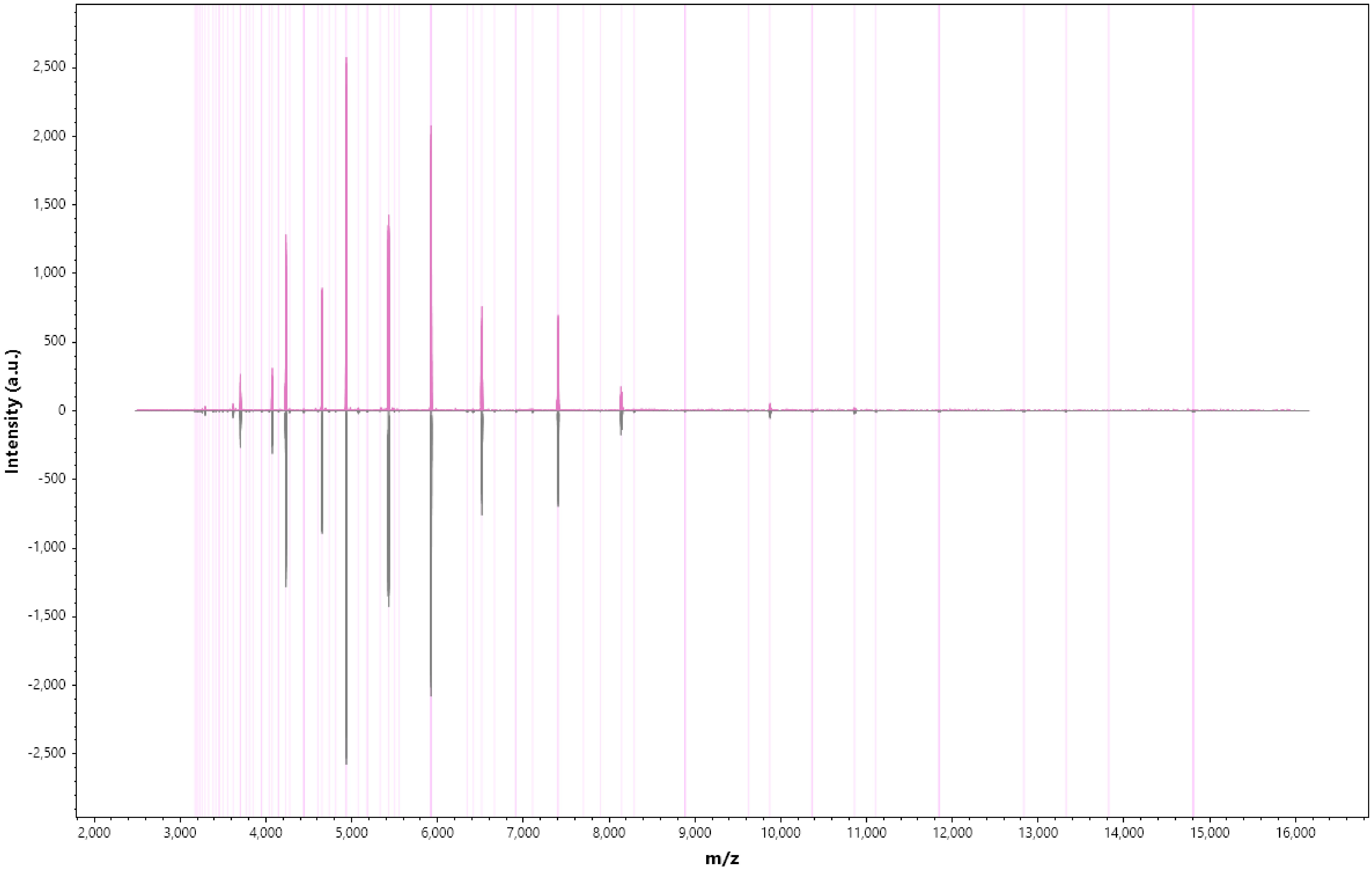

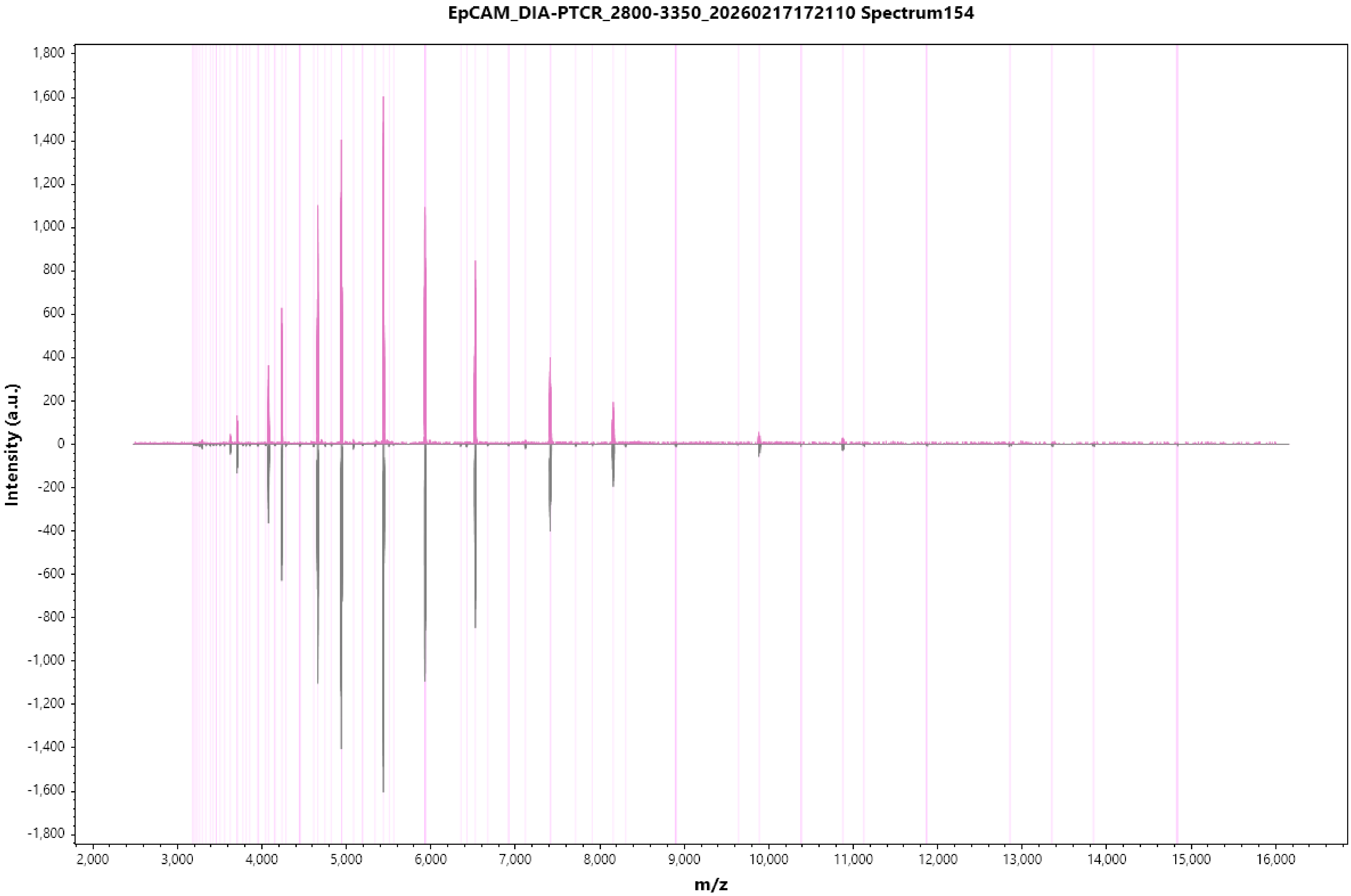

EpCAM_DIA-PTCR_2800-3350_20260217172110 Spectrum154
Intensity (a.u.)
1,800
1,600
1,400
1,200
1,000
800
600
400
200
0
-200
-400
-600
-800
-1,000
-1,200
-1,400
-1,600
-1,800
2,000
3,000
4,000
5,000
6,000
7,000
8,000
9,000
10,000
11,000
12,000
13,000
14,000
15,000
16,000
m/z

EpCAM_DIA-PTCR_2800-3350_20260217172110 Spectrum155

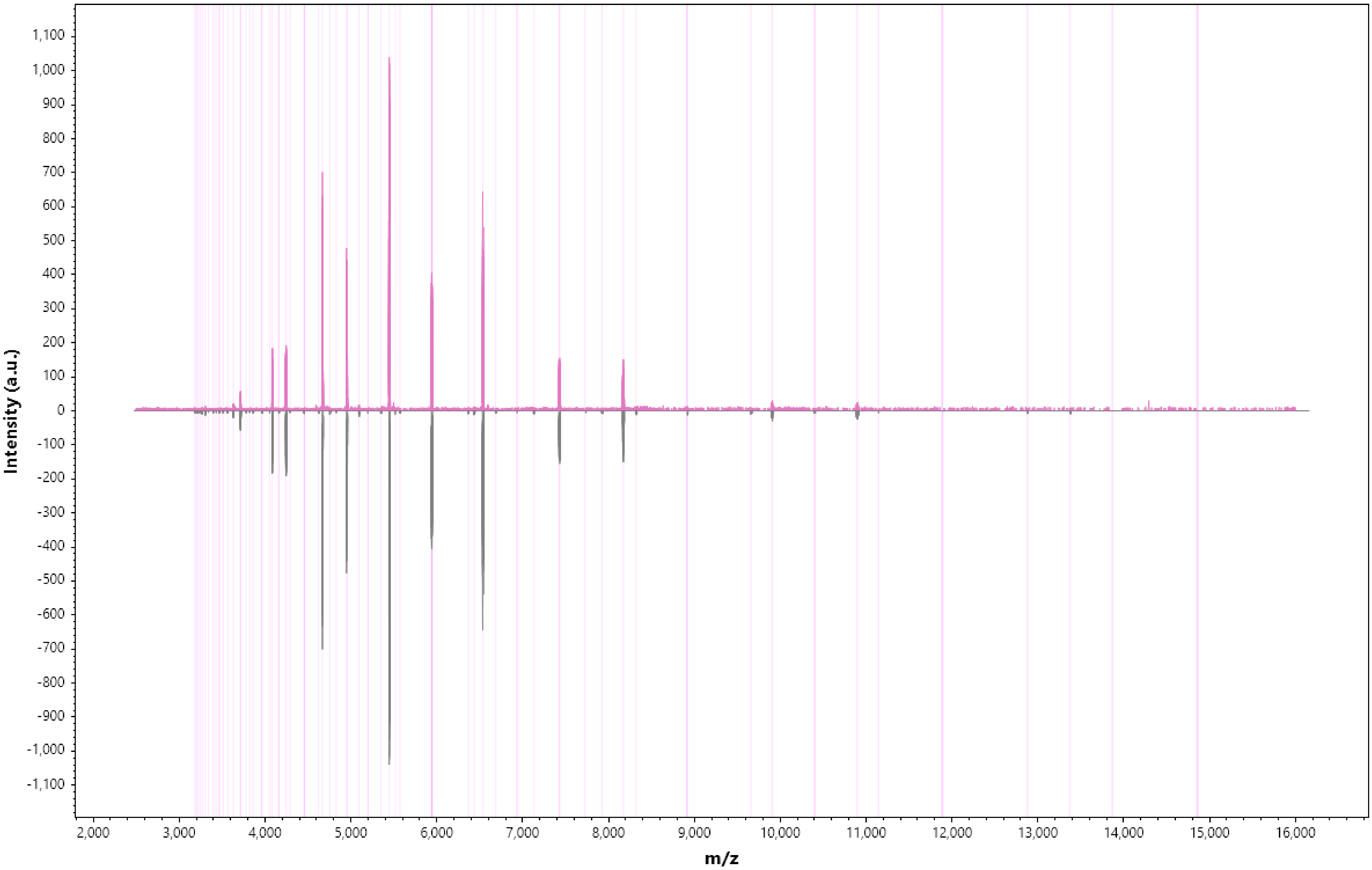

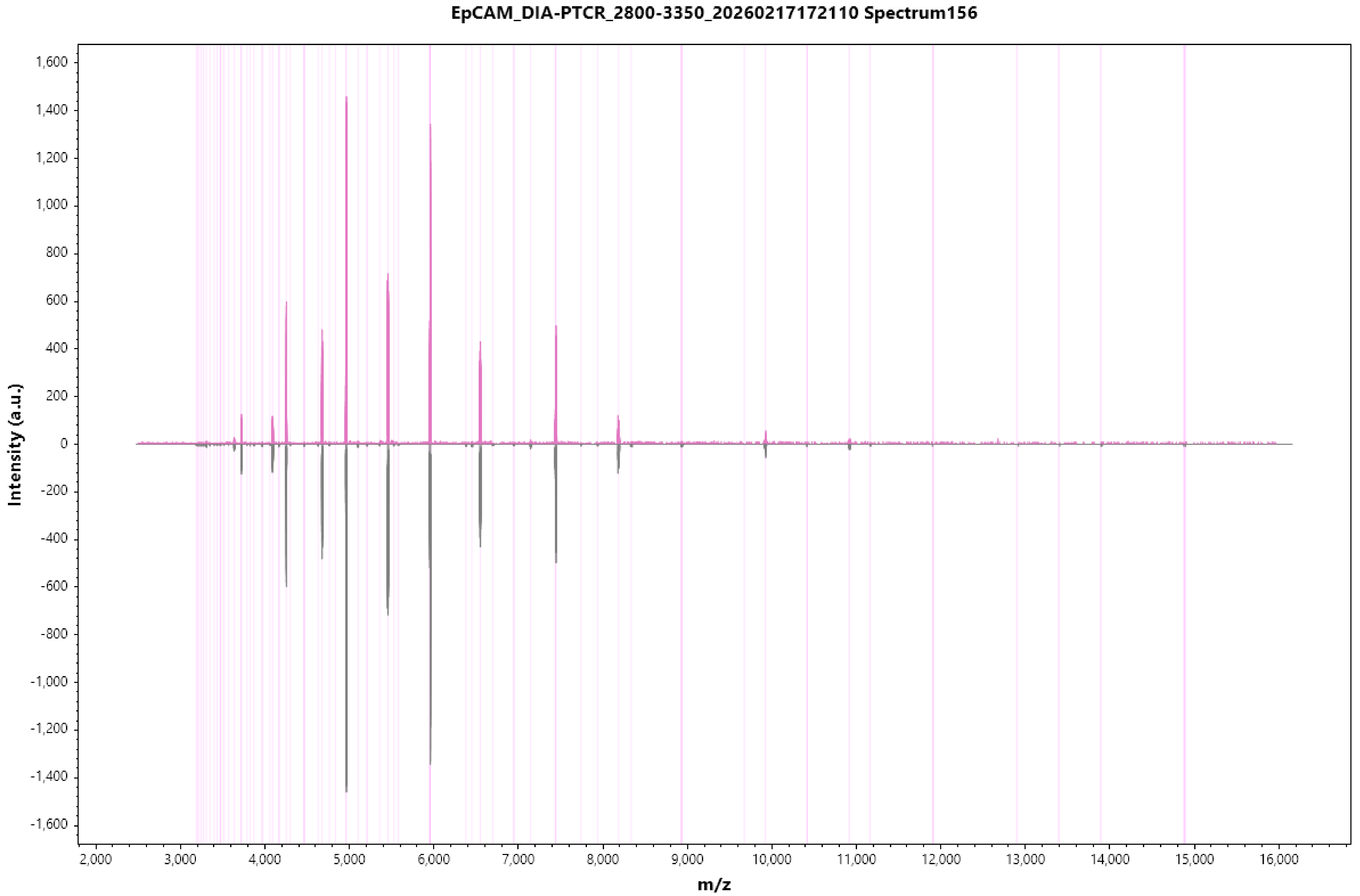
EpCAM_DIA-PTCR_2800-3350_20260217172110 Spectrum156
Intensity (a.u.)
m/z
1,600
1,400
1,200
1,000
800
600
400
200
0
-200
-400
-600
-800
-1,000
-1,200
-1,400
-1,600
2,000
3,000
4,000
5,000
6,000
7,000
8,000
9,000
10,000
11,000
12,000
13,000
14,000
15,000
16,000

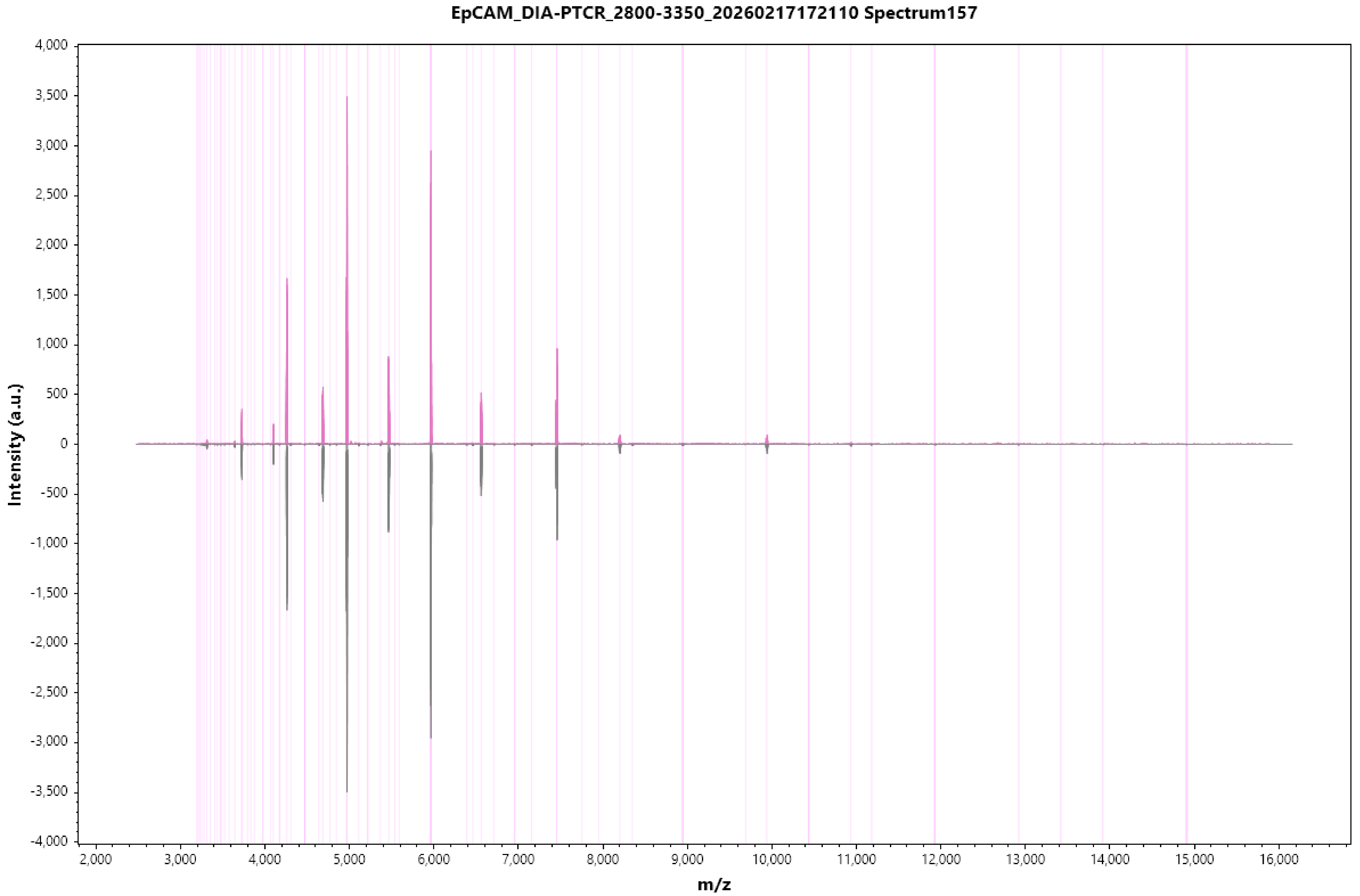
EpCAM_DIA-PTCR_2800-3350_20260217172110 Spectrum157
Intensity (a.u.)
m/z
4,000
3,500
3,000
2,500
2,000
1,500
1,000
500
0
-500
-1,000
-1,500
-2,000
-2,500
-3,000
-3,500
-4,000
2,000
3,000
4,000
5,000
6,000
7,000
8,000
9,000
10,000
11,000
12,000
13,000
14,000
15,000
16,000

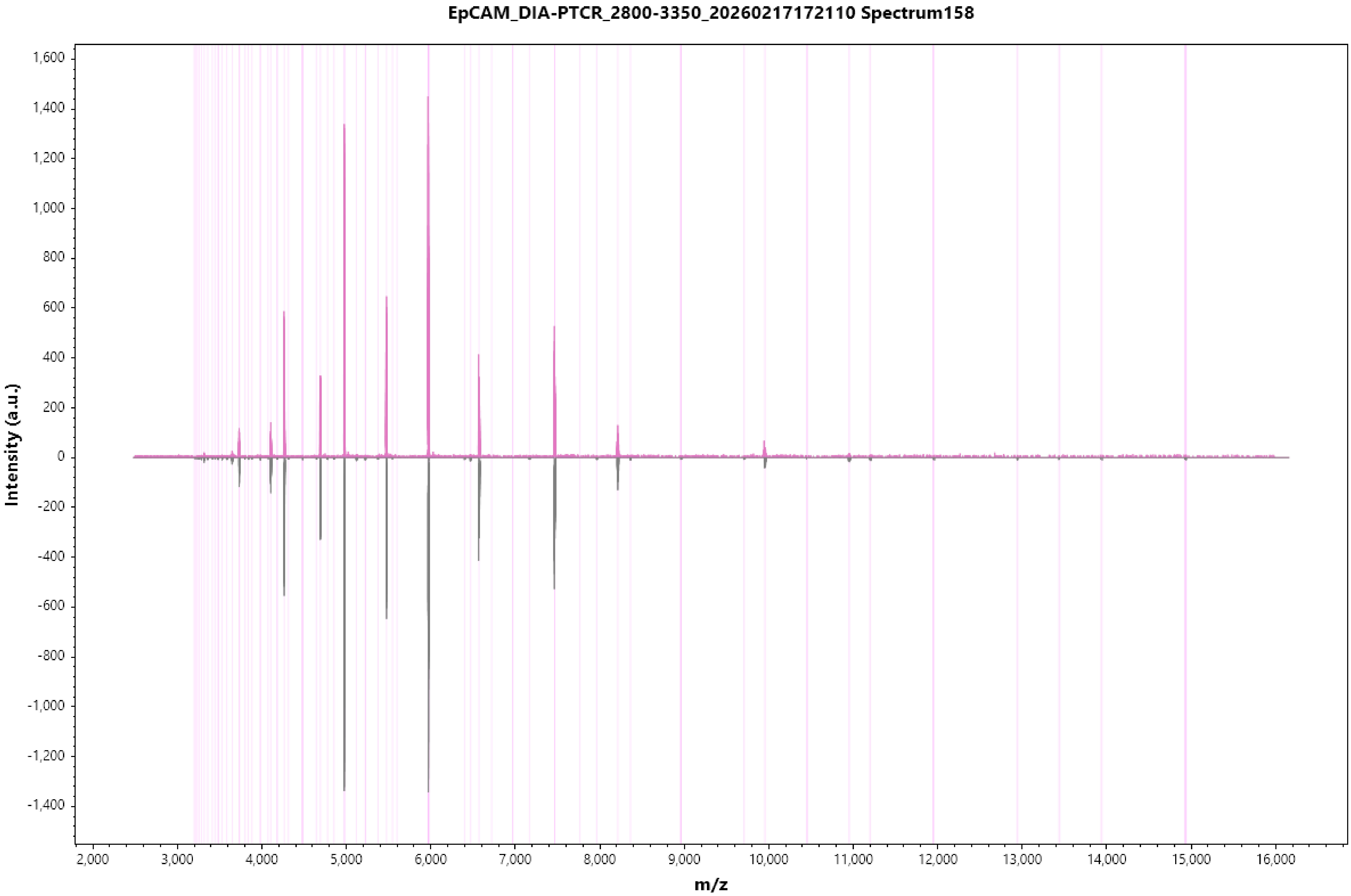
EpCAM_DIA-PTCR_2800-3350_20260217172110 Spectrum158
Intensity (a.u.)
m/z
1,600
1,400
1,200
1,000
800
600
400
200
0
-200
-400
-600
-800
-1,000
-1,200
-1,400
2,000
3,000
4,000
5,000
6,000
7,000
8,000
9,000
10,000
11,000
12,000
13,000
14,000
15,000
16,000

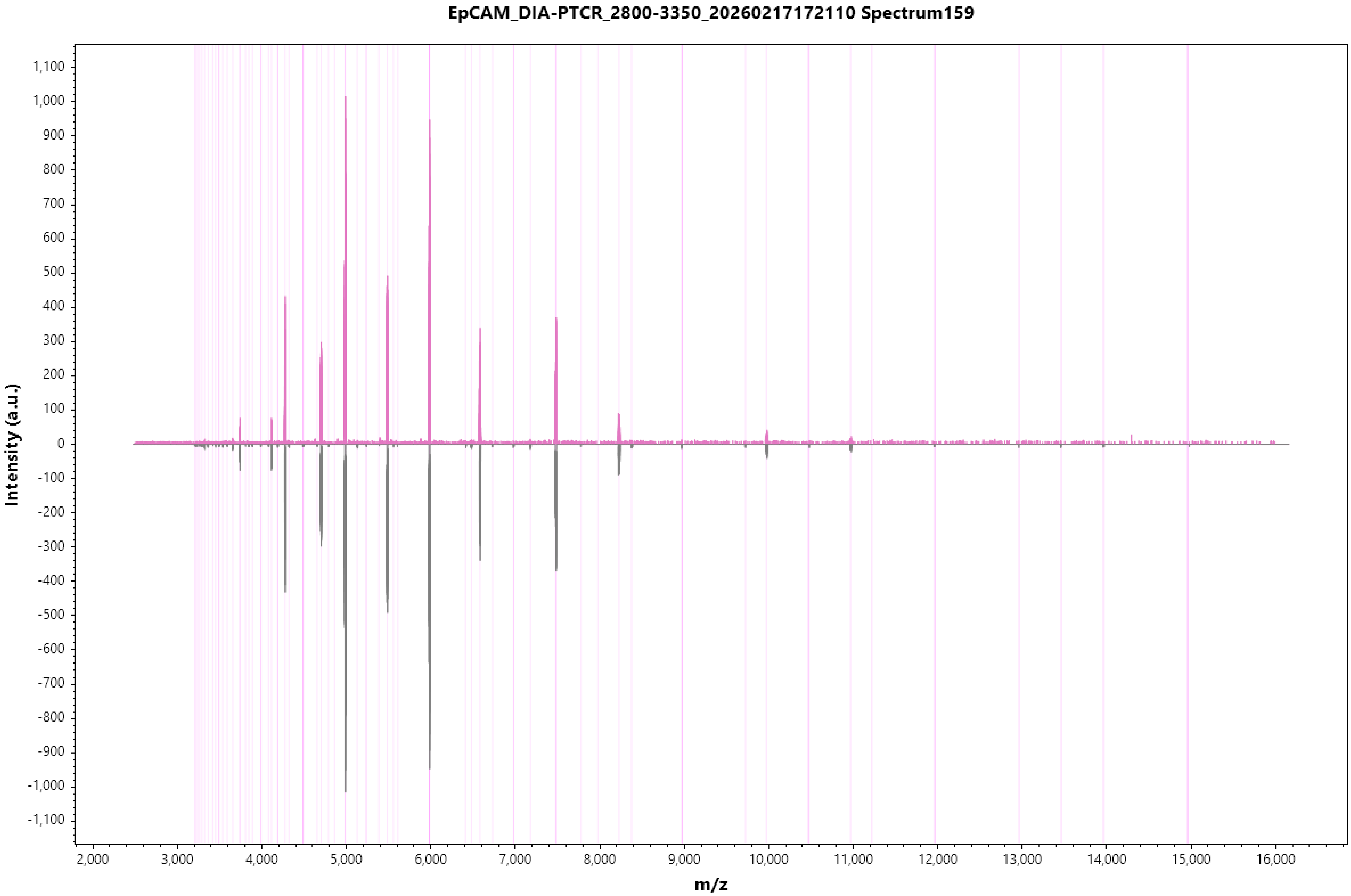
EpCAM_DIA-PTCR_2800-3350_20260217172110 Spectrum159
Intensity (a.u.)
m/z
1,100
1,000
900
800
700
600
500
400
300
200
100
0
-100
-200
-300
-400
-500
-600
-700
-800
-900
-1,000
-1,100
2,000
3,000
4,000
5,000
6,000
7,000
8,000
9,000
10,000
11,000
12,000
13,000
14,000
15,000
16,000

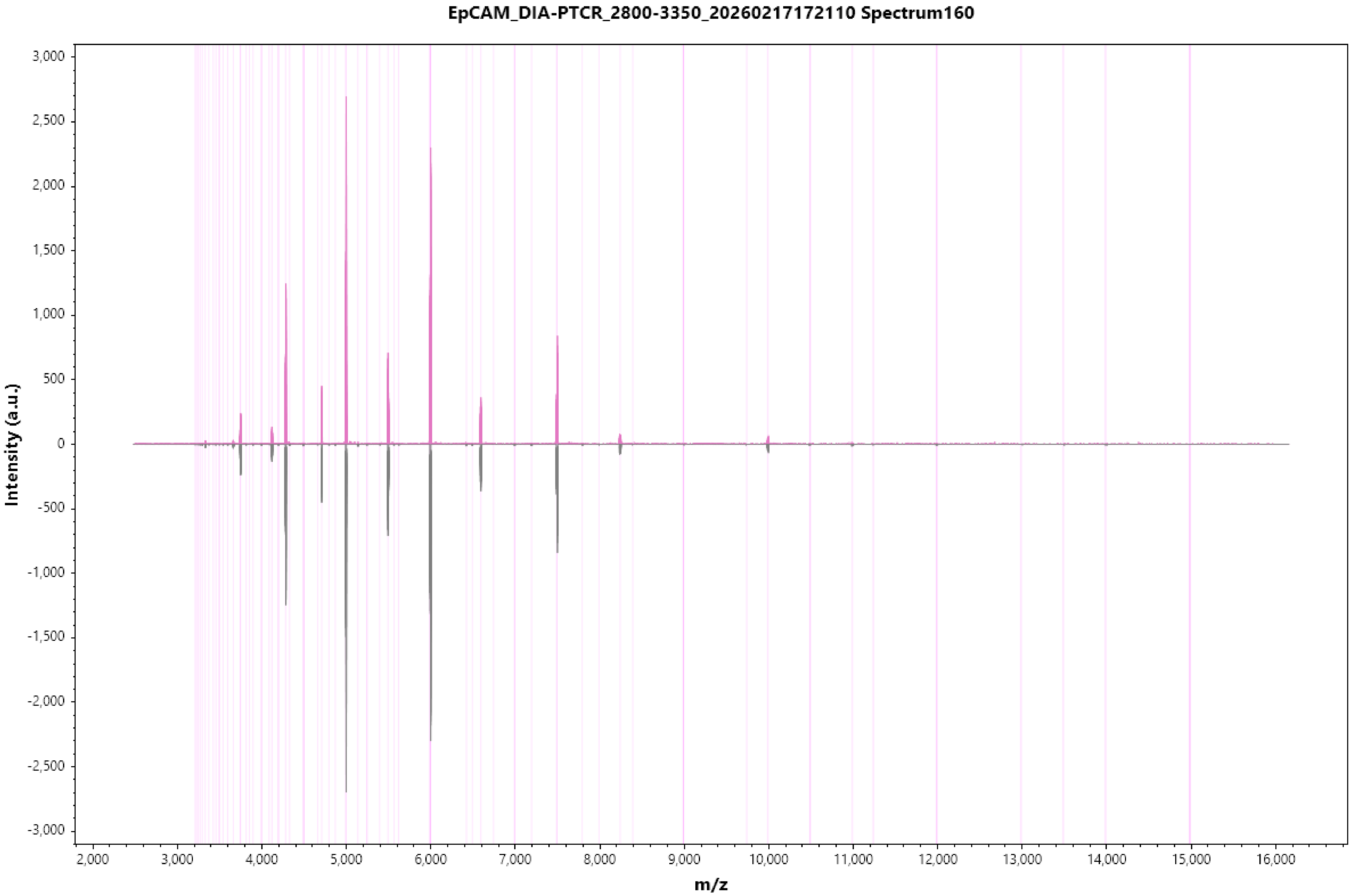
EpCAM_DIA-PTCR_2800-3350_20260217172110 Spectrum160
Intensity (a.u.)
3,000
2,500
2,000
1,500
1,000
500
0
-500
-1,000
-1,500
-2,000
-2,500
-3,000
2,000
3,000
4,000
5,000
6,000
7,000
8,000
9,000
10,000
11,000
12,000
13,000
14,000
15,000
16,000
m/z

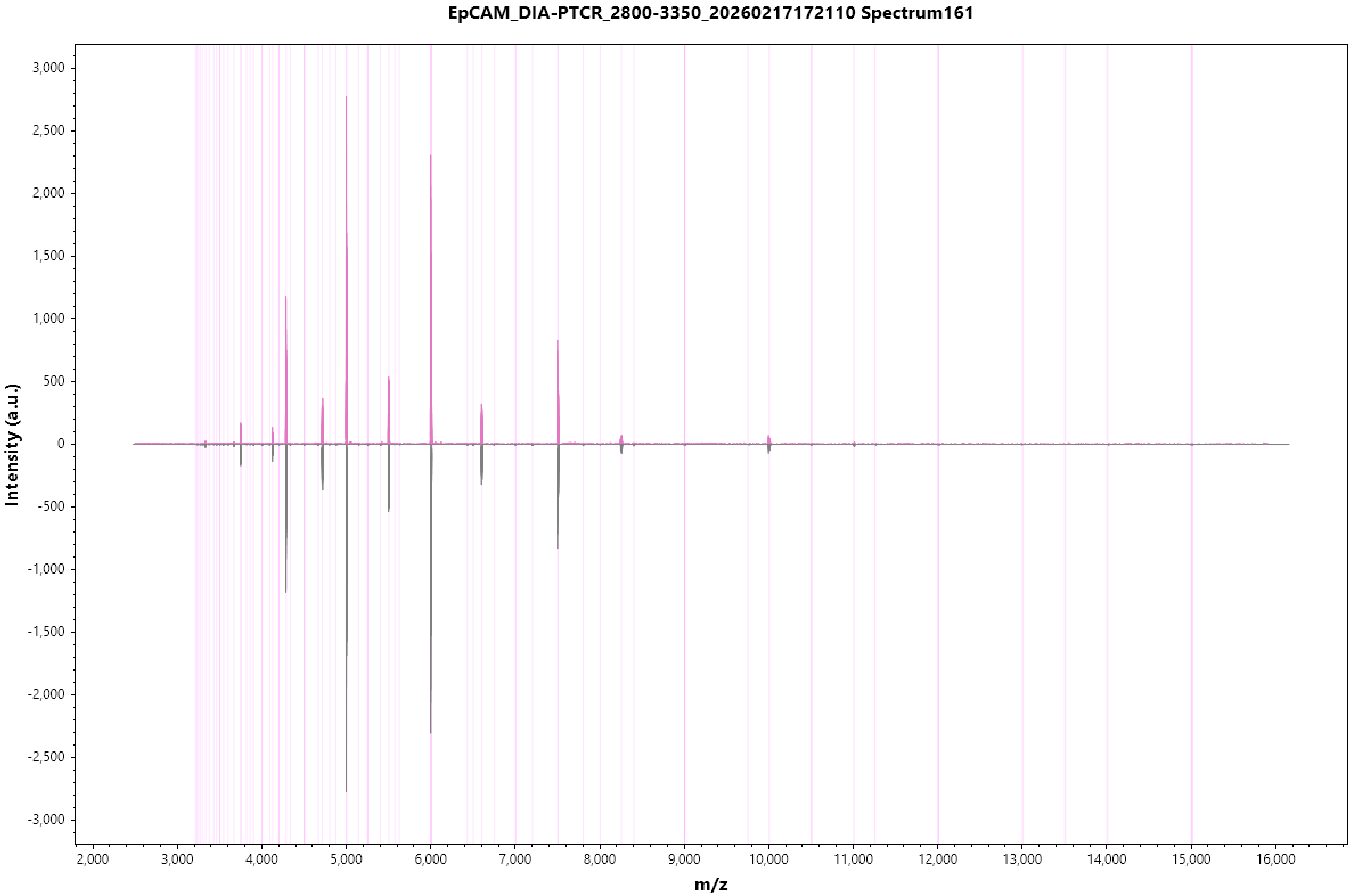
EpCAM_DIA-PTCR_2800-3350_20260217172110 Spectrum161
Intensity (a.u.)
3,000
2,500
2,000
1,500
1,000
500
0
-500
-1,000
-1,500
-2,000
-2,500
-3,000
2,000
3,000
4,000
5,000
6,000
7,000
8,000
9,000
10,000
11,000
12,000
13,000
14,000
15,000
16,000
m/z

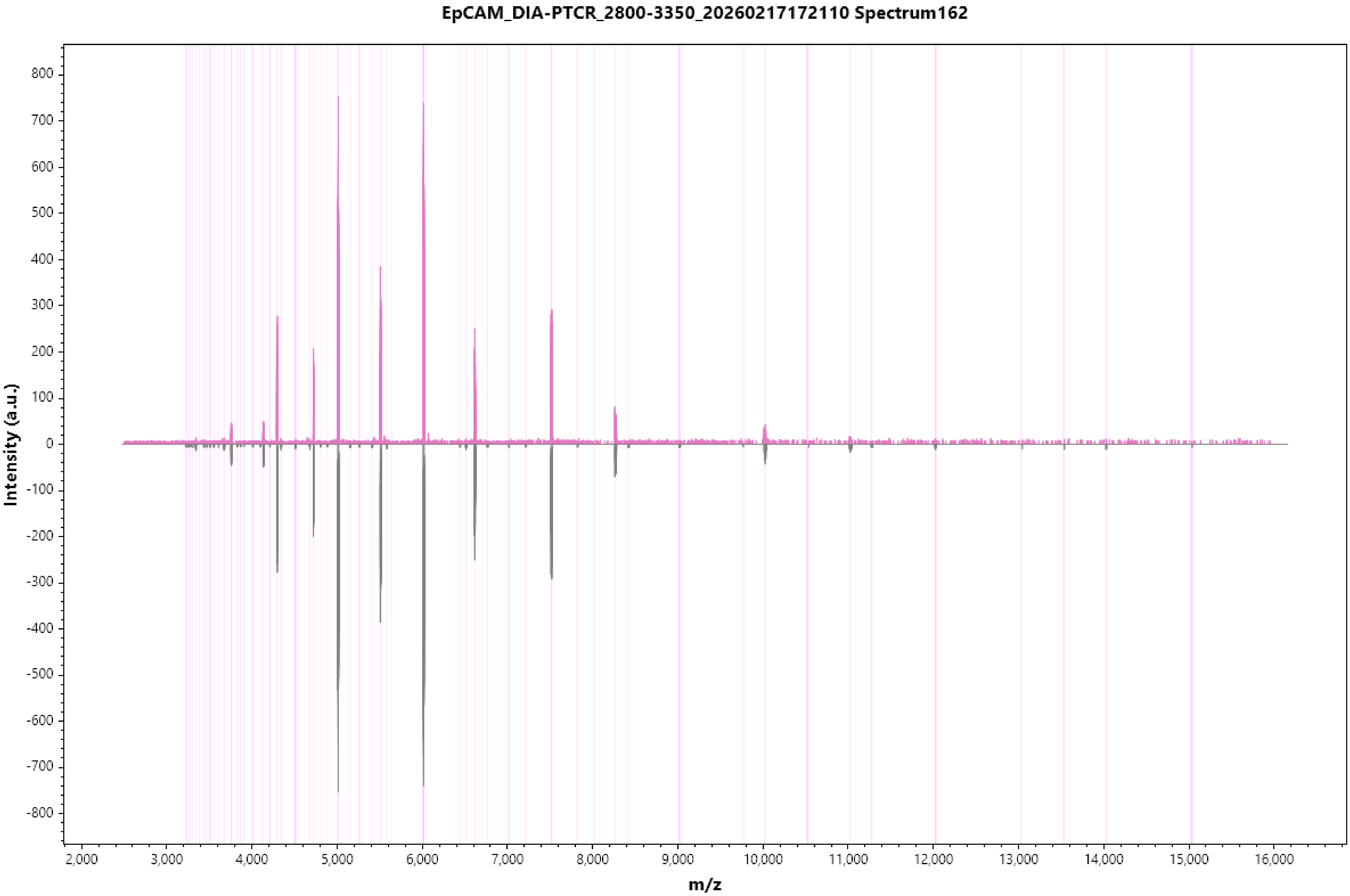

EpCAM_DIA-PTCR_2800-3350_20260217172110 Spectrum162
Intensity (a.u.)
m/z
800
700
600
500
400
300
200
100
0
-100
-200
-300
-400
-500
-600
-700
-800
2,000
3,000
4,000
5,000
6,000
7,000
8,000
9,000
10,000
11,000
12,000
13,000
14,000
15,000
16,000

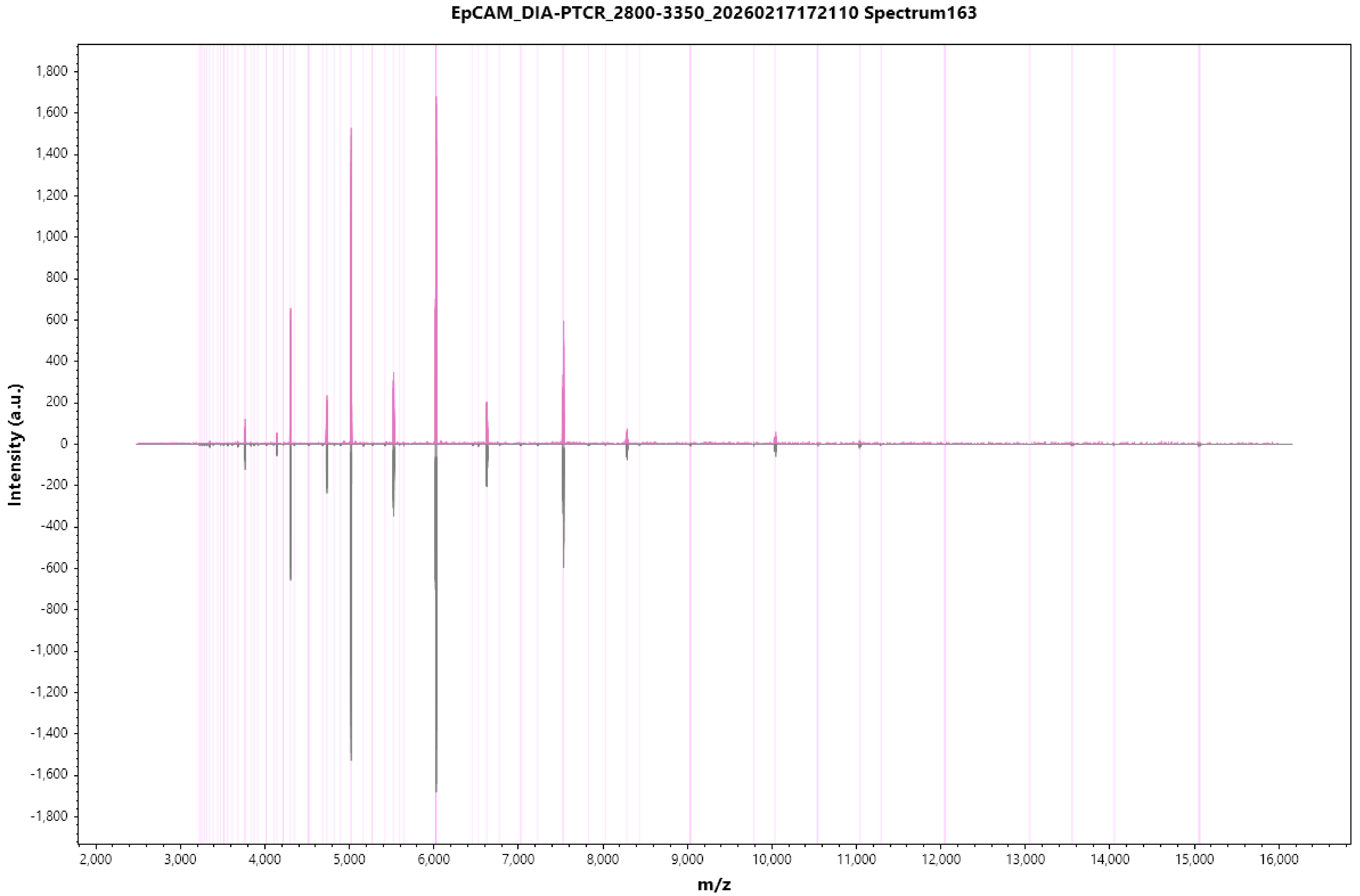
EpCAM_DIA-PTCR_2800-3350_20260217172110 Spectrum163
Intensity (a.u.)
m/z
1,800
1,600
1,400
1,200
1,000
800
600
400
200
0
-200
-400
-600
-800
-1,000
-1,200
-1,400
-1,600
-1,800
2,000
3,000
4,000
5,000
6,000
7,000
8,000
9,000
10,000
11,000
12,000
13,000
14,000
15,000
16,000

EpCAM_DIA-PTCR_2800-3350_20260217172110 Spectrum164

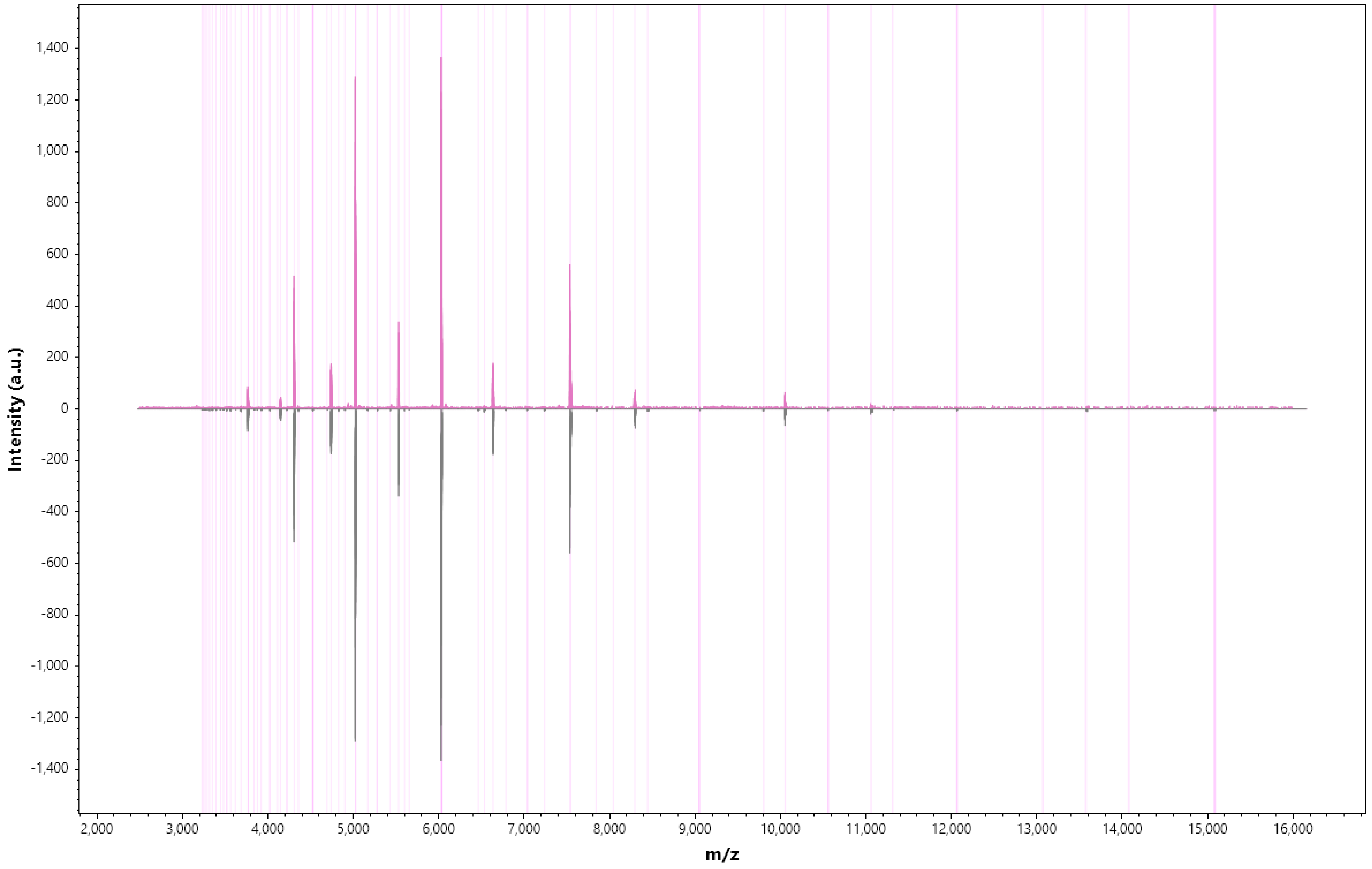

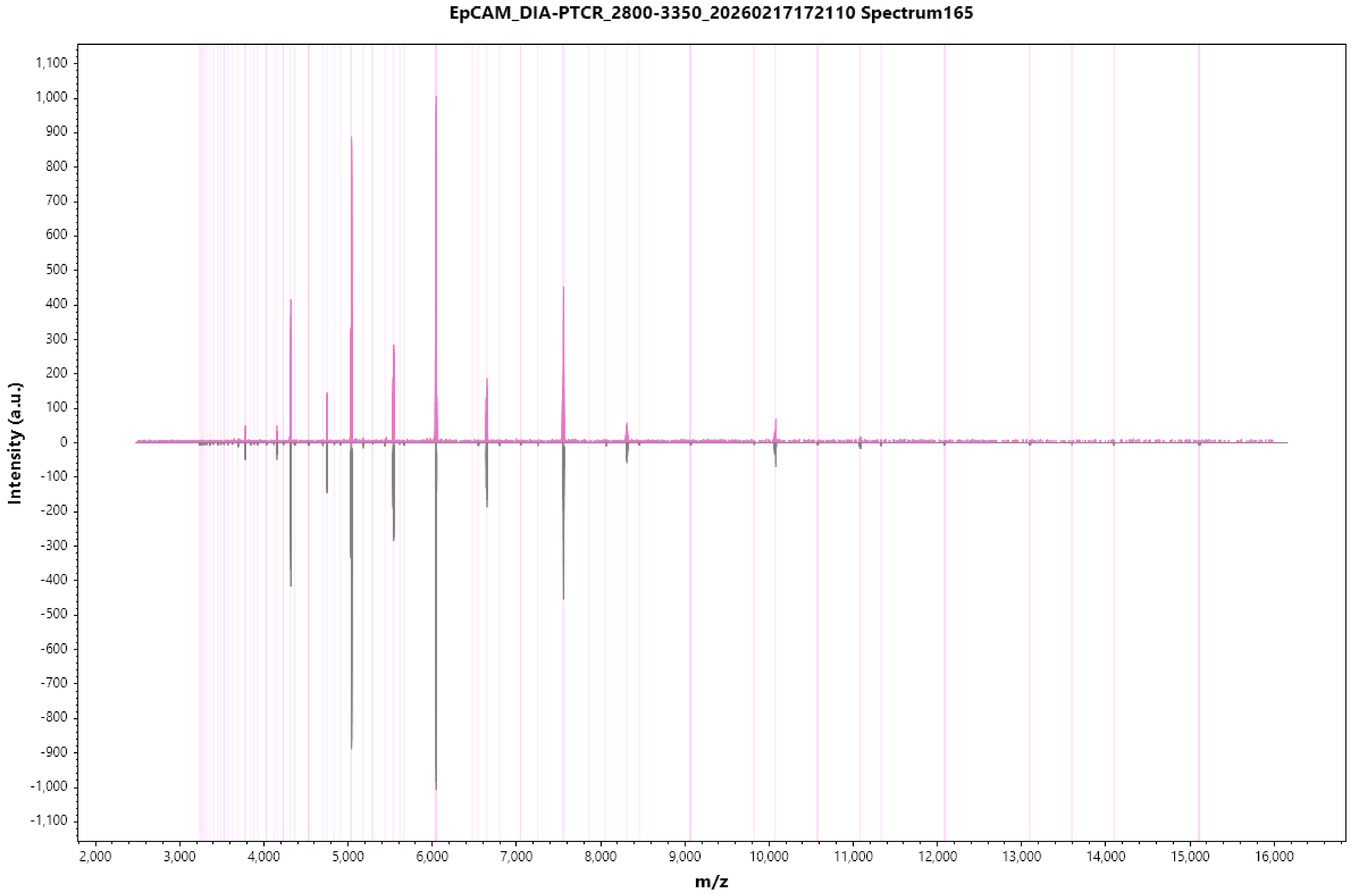
EpCAM_DIA-PTCR_2800-3350_20260217172110 Spectrum165
Intensity (a.u.)
m/z
1,100
1,000
900
800
700
600
500
400
300
200
100
0
-100
-200
-300
-400
-500
-600
-700
-800
-900
-1,000
-1,100
2,000
3,000
4,000
5,000
6,000
7,000
8,000
9,000
10,000
11,000
12,000
13,000
14,000
15,000
16,000

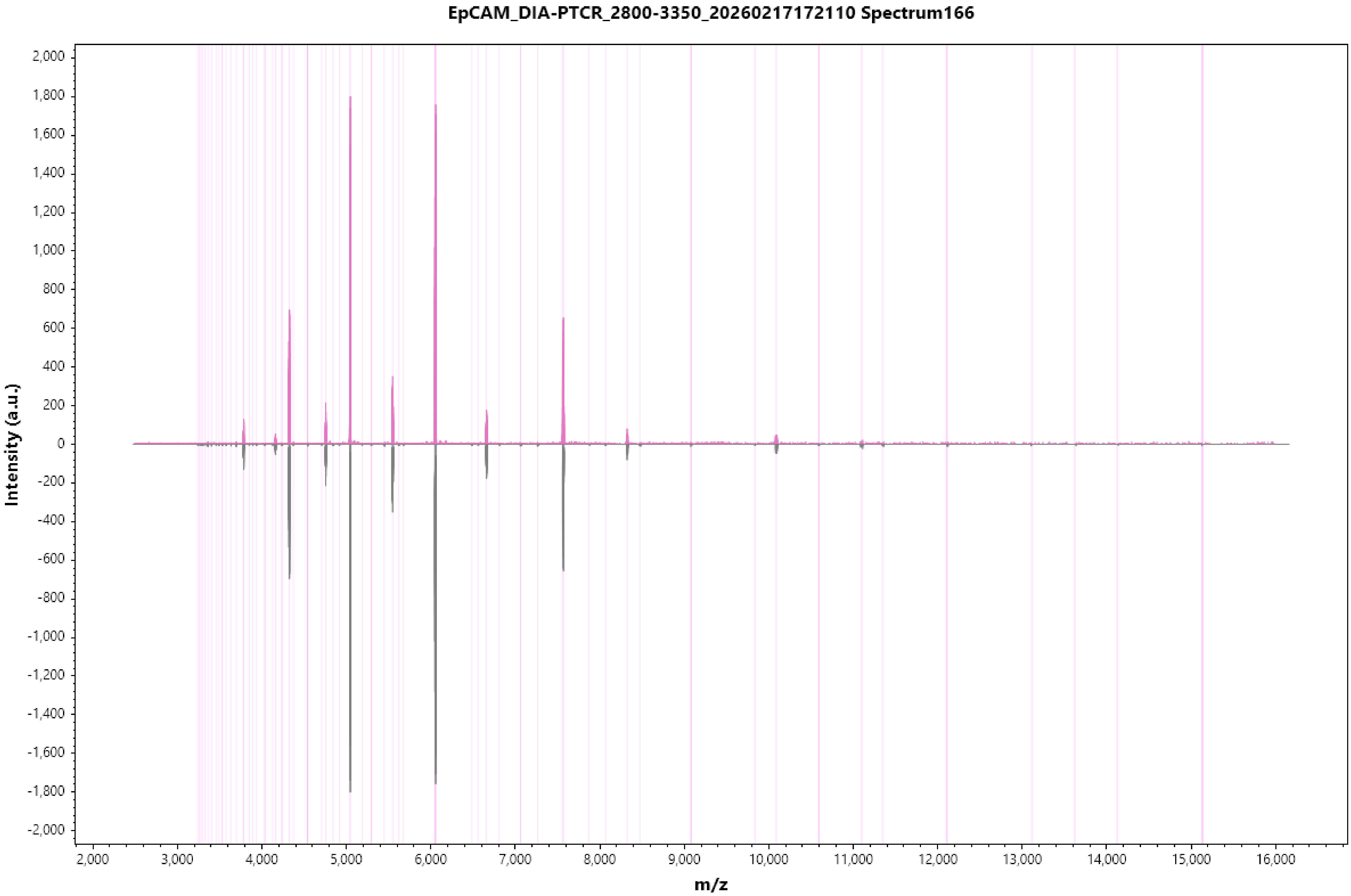
EpCAM_DIA-PTCR_2800-3350_20260217172110 Spectrum166
Intensity (a.u.)
m/z
2,000
1,800
1,600
1,400
1,200
1,000
800
600
400
200
0
-200
-400
-600
-800
-1,000
-1,200
-1,400
-1,600
-1,800
-2,000
2,000
3,000
4,000
5,000
6,000
7,000
8,000
9,000
10,000
11,000
12,000
13,000
14,000
15,000
16,000

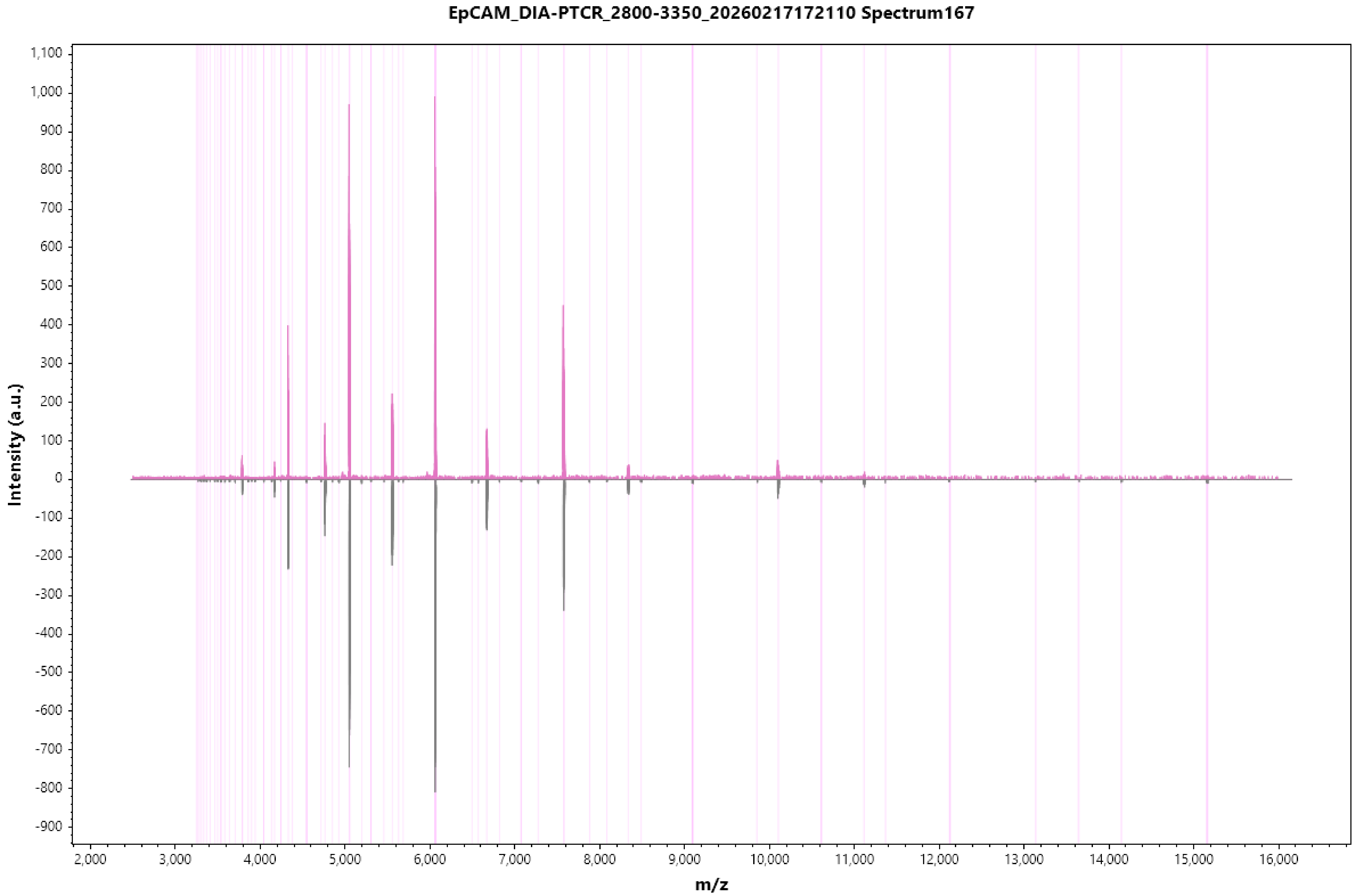
EpCAM_DIA-PTCR_2800-3350_20260217172110 Spectrum167
Intensity (a.u.)
m/z
1,100
1,000
900
800
700
600
500
400
300
200
100
0
-100
-200
-300
-400
-500
-600
-700
-800
-900
2,000
3,000
4,000
5,000
6,000
7,000
8,000
9,000
10,000
11,000
12,000
13,000
14,000
15,000
16,000

### EpCAM_DIA-PTCR_2800-3350_20260217172110 Spectrum168

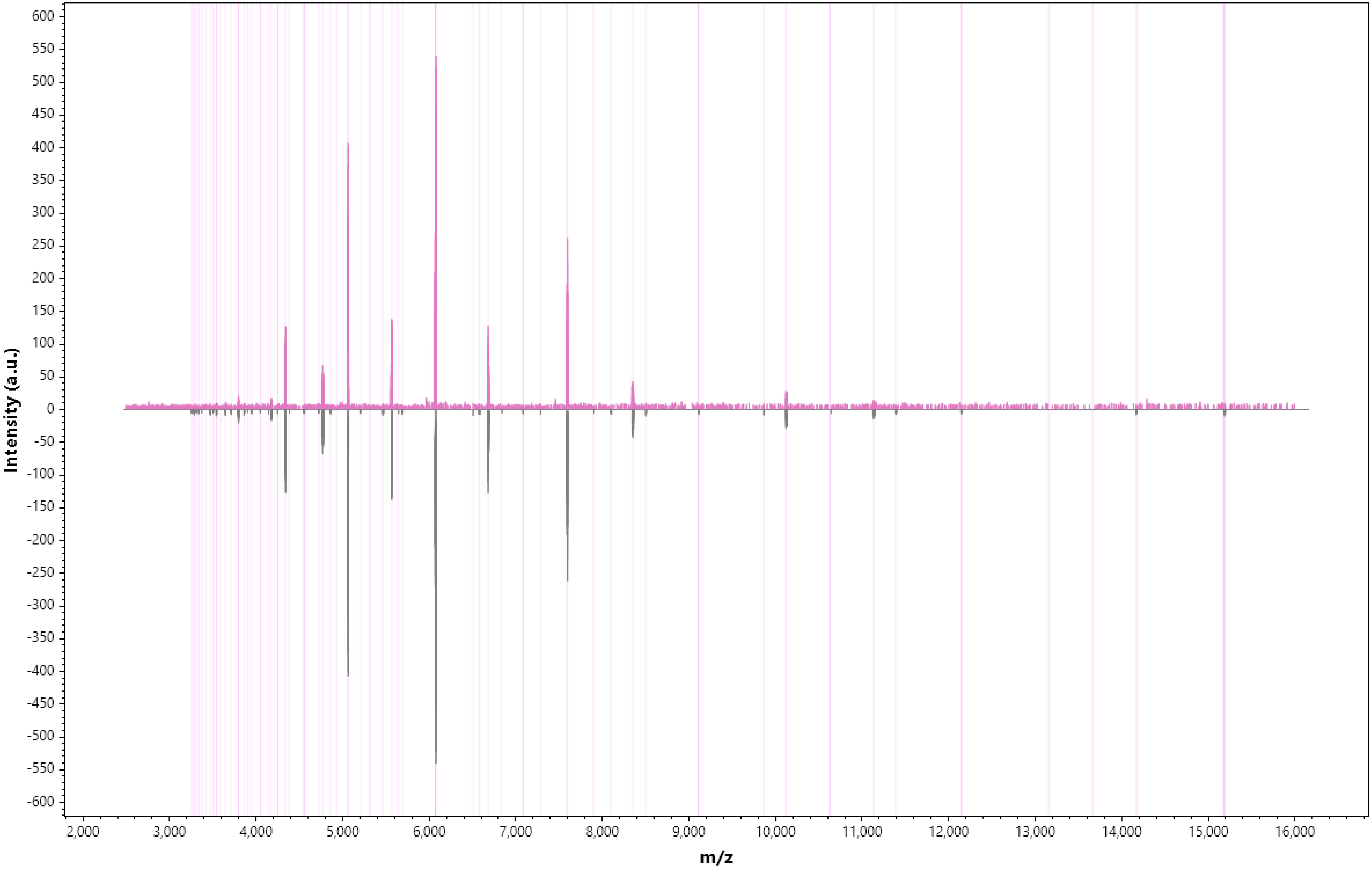

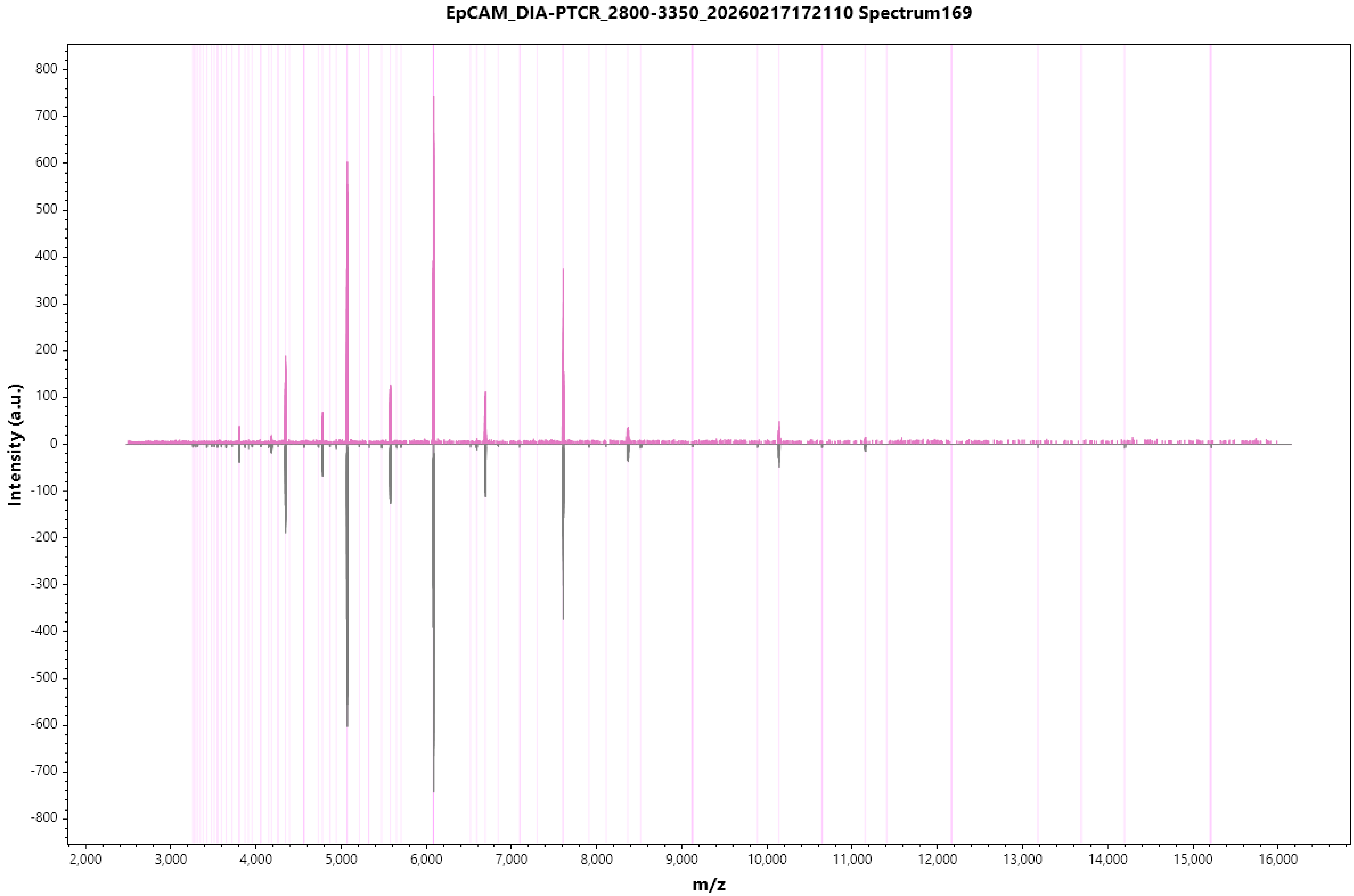

EpCAM_DIA-PTCR_2800-3350_20260217172110 Spectrum169
Intensity (a.u.)
m/z
800
700
600
500
400
300
200
100
0
-100
-200
-300
-400
-500
-600
-700
-800
2,000
3,000
4,000
5,000
6,000
7,000
8,000
9,000
10,000
11,000
12,000
13,000
14,000
15,000
16,000

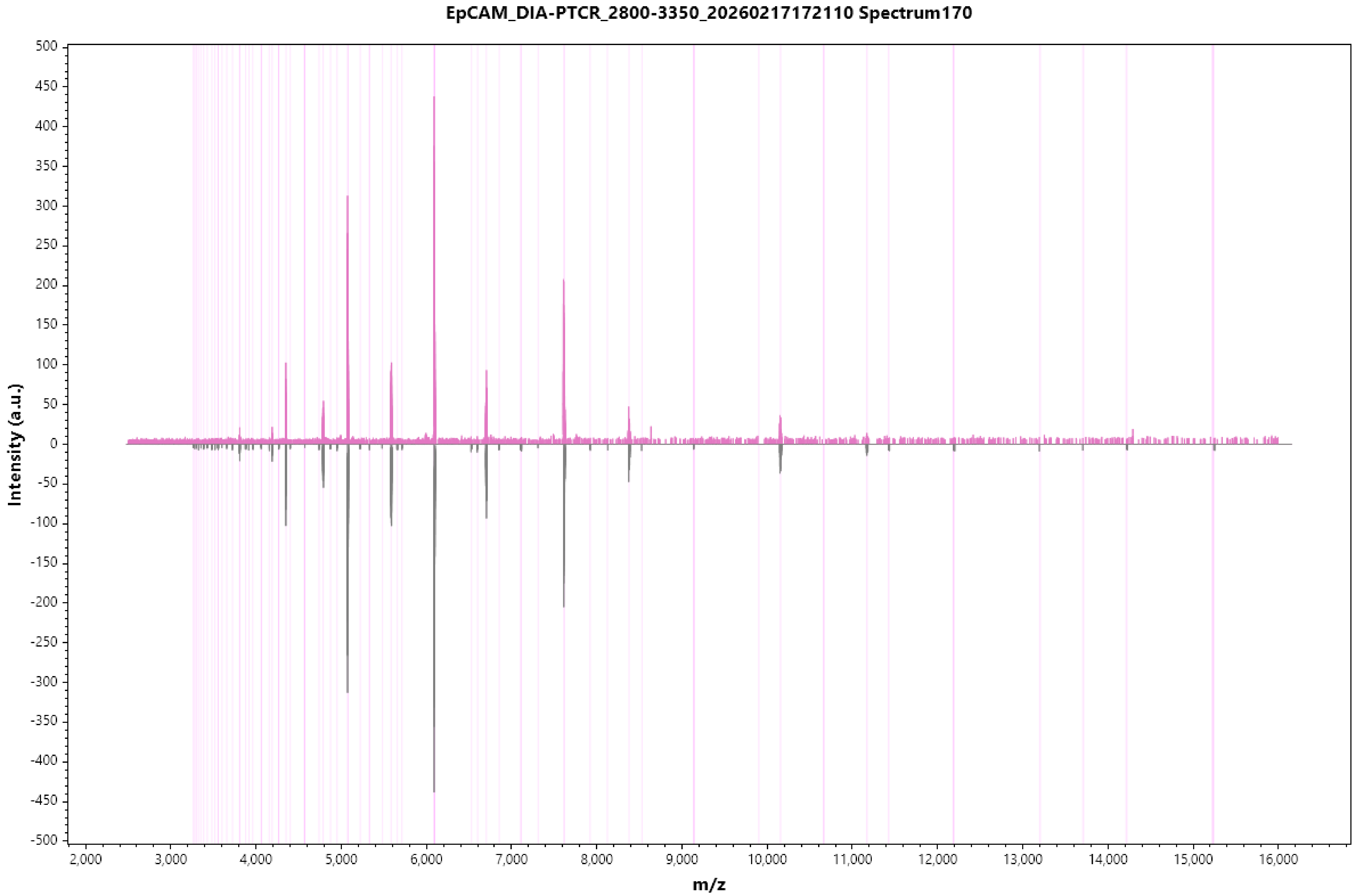

EpCAM_DIA-PTCR_2800-3350_20260217172110 Spectrum170
Intensity (a.u.)
m/z

EpCAM_DIA-PTCR_2800-3350_20260217172110 Spectrum171

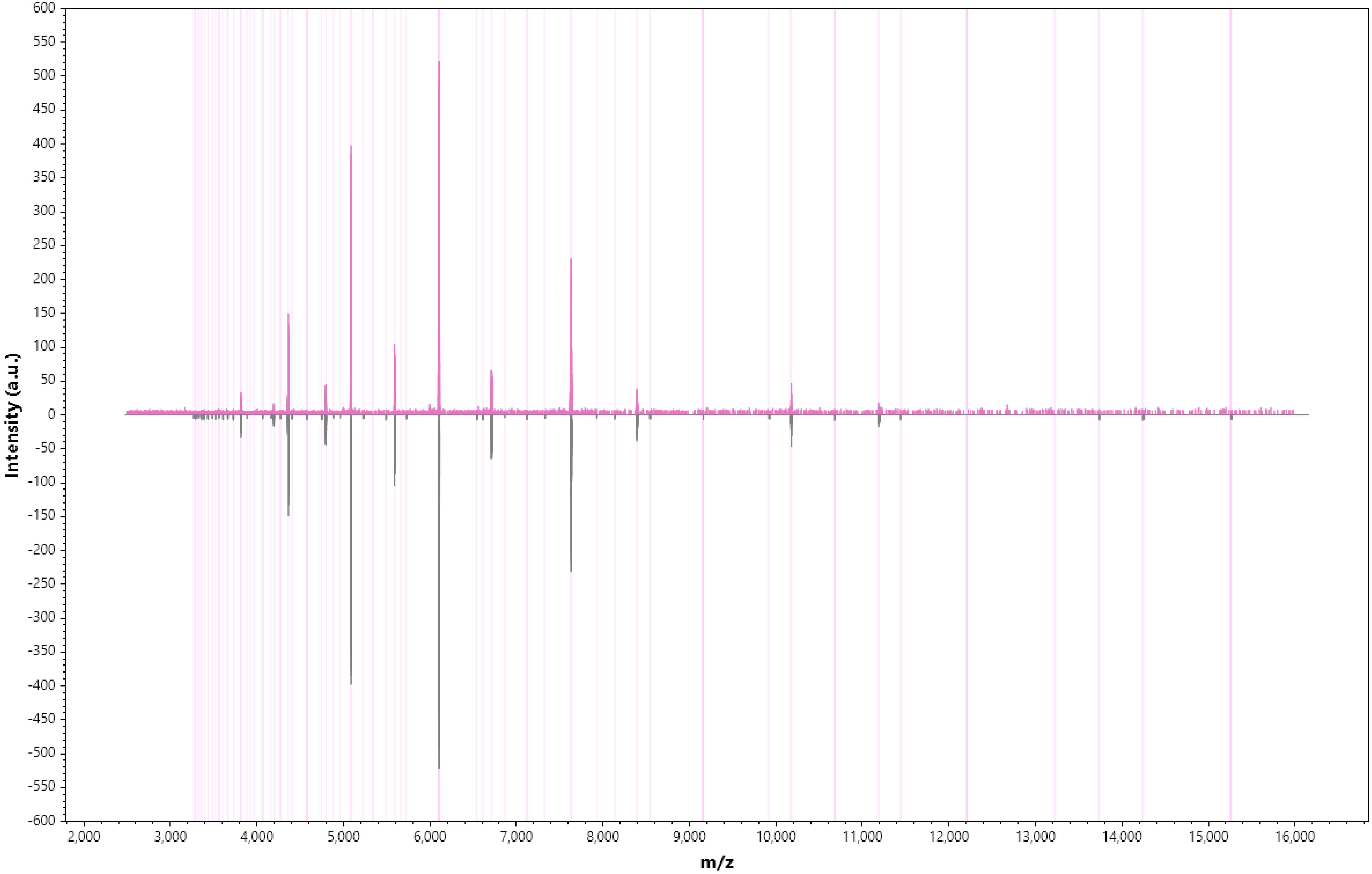

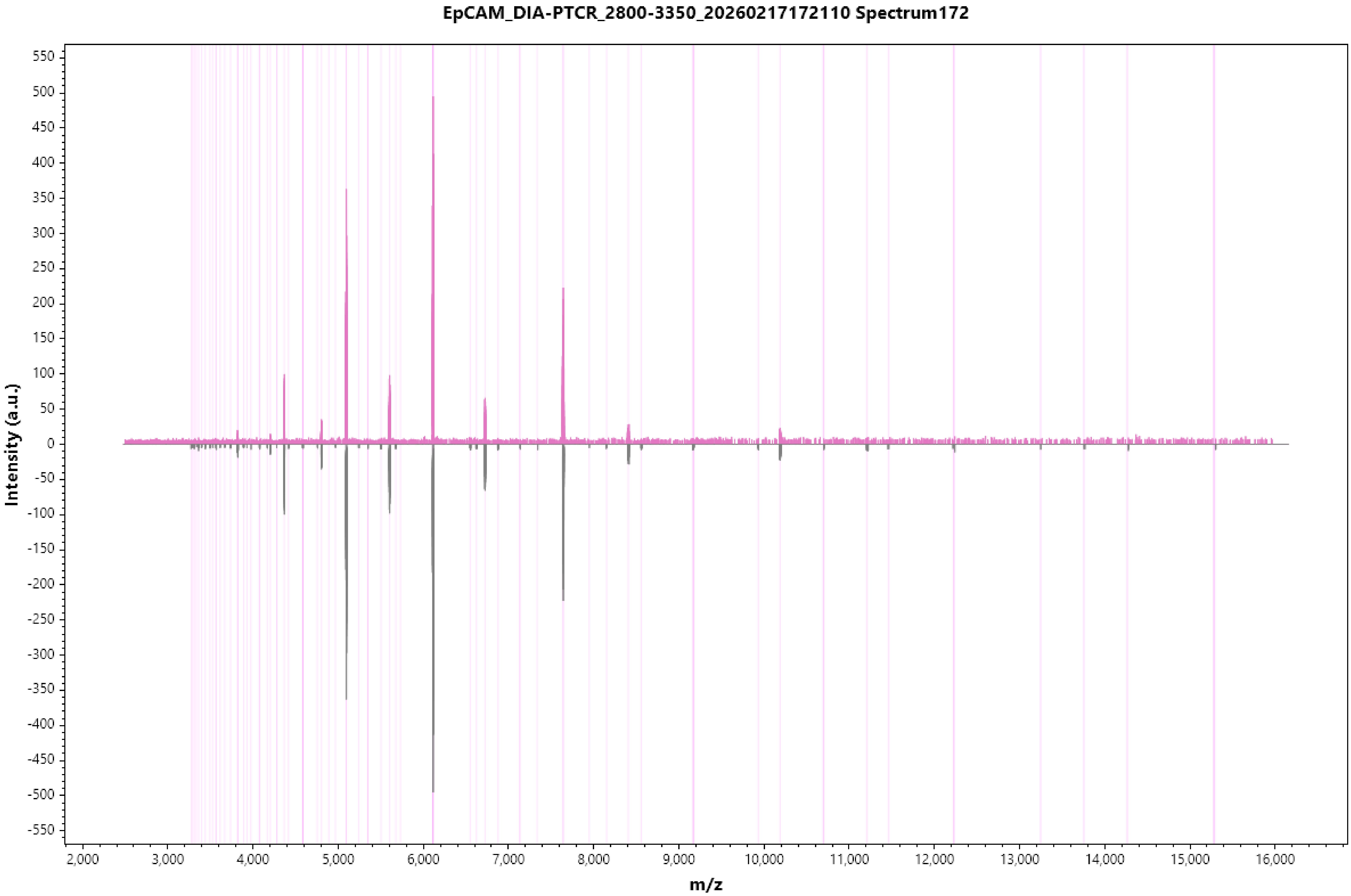

EpCAM_DIA-PTCR_2800-3350_20260217172110 Spectrum172
Intensity (a.u.)
m/z

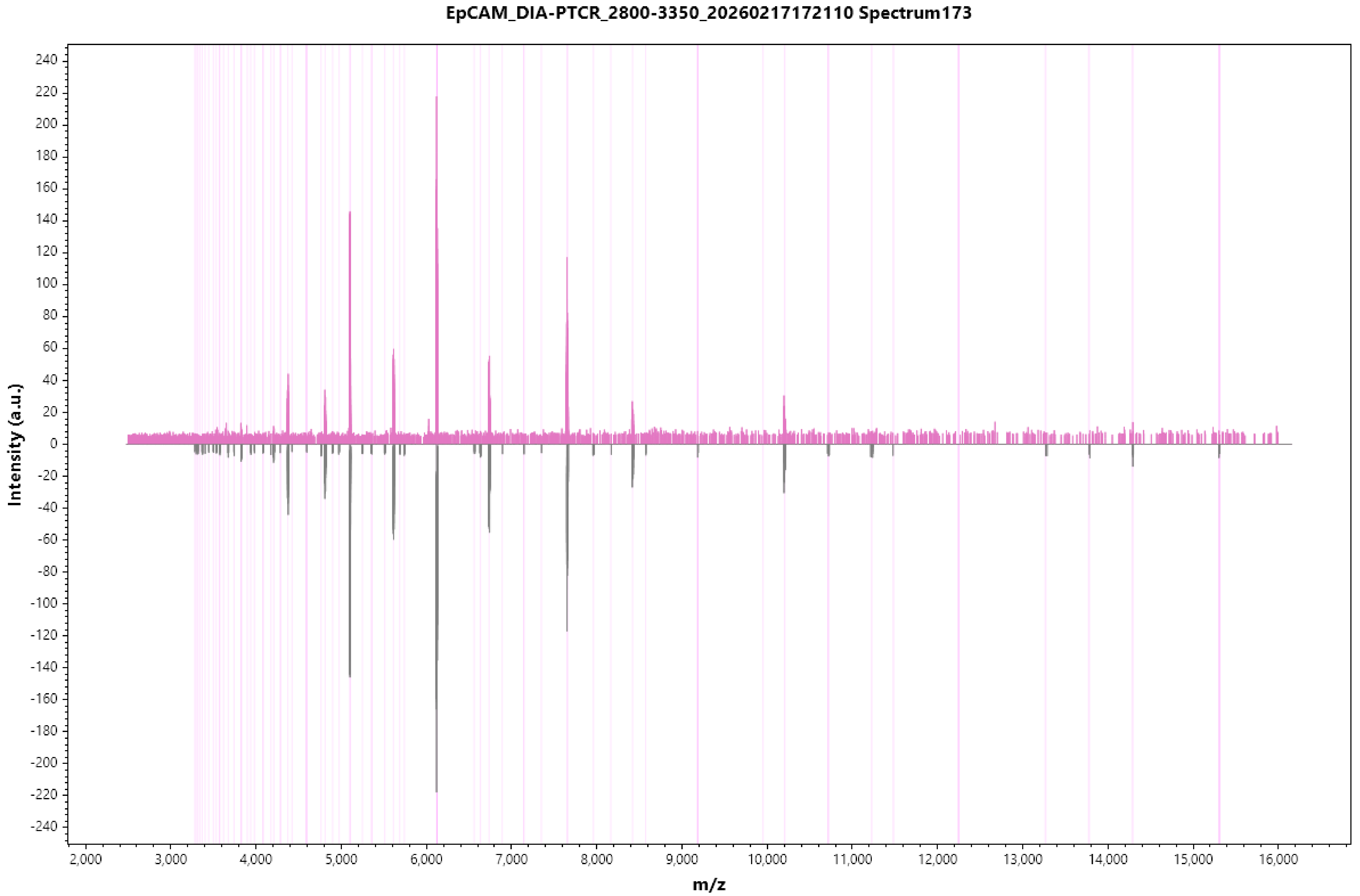

EpCAM_DIA-PTCR_2800-3350_20260217172110 Spectrum173
Intensity (a.u.)
m/z
240
220
200
180
160
140
120
100
80
60
40
20
0
-20
-40
-60
-80
-100
-120
-140
-160
-180
-200
-220
-240
2,000
3,000
4,000
5,000
6,000
7,000
8,000
9,000
10,000
11,000
12,000
13,000
14,000
15,000
16,000

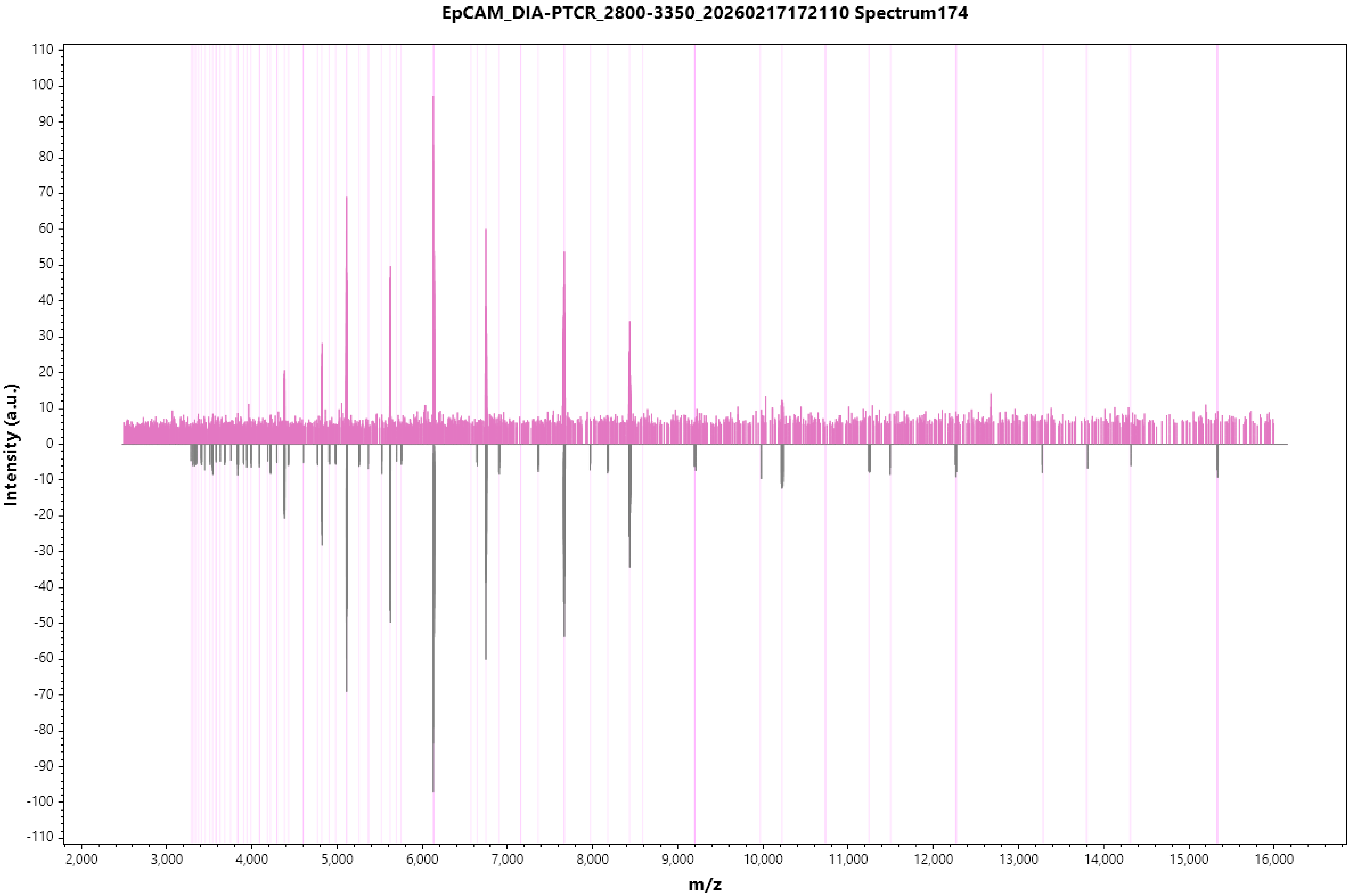

EpCAM_DIA-PTCR_2800-3350_20260217172110 Spectrum174
Intensity (a.u.)
m/z

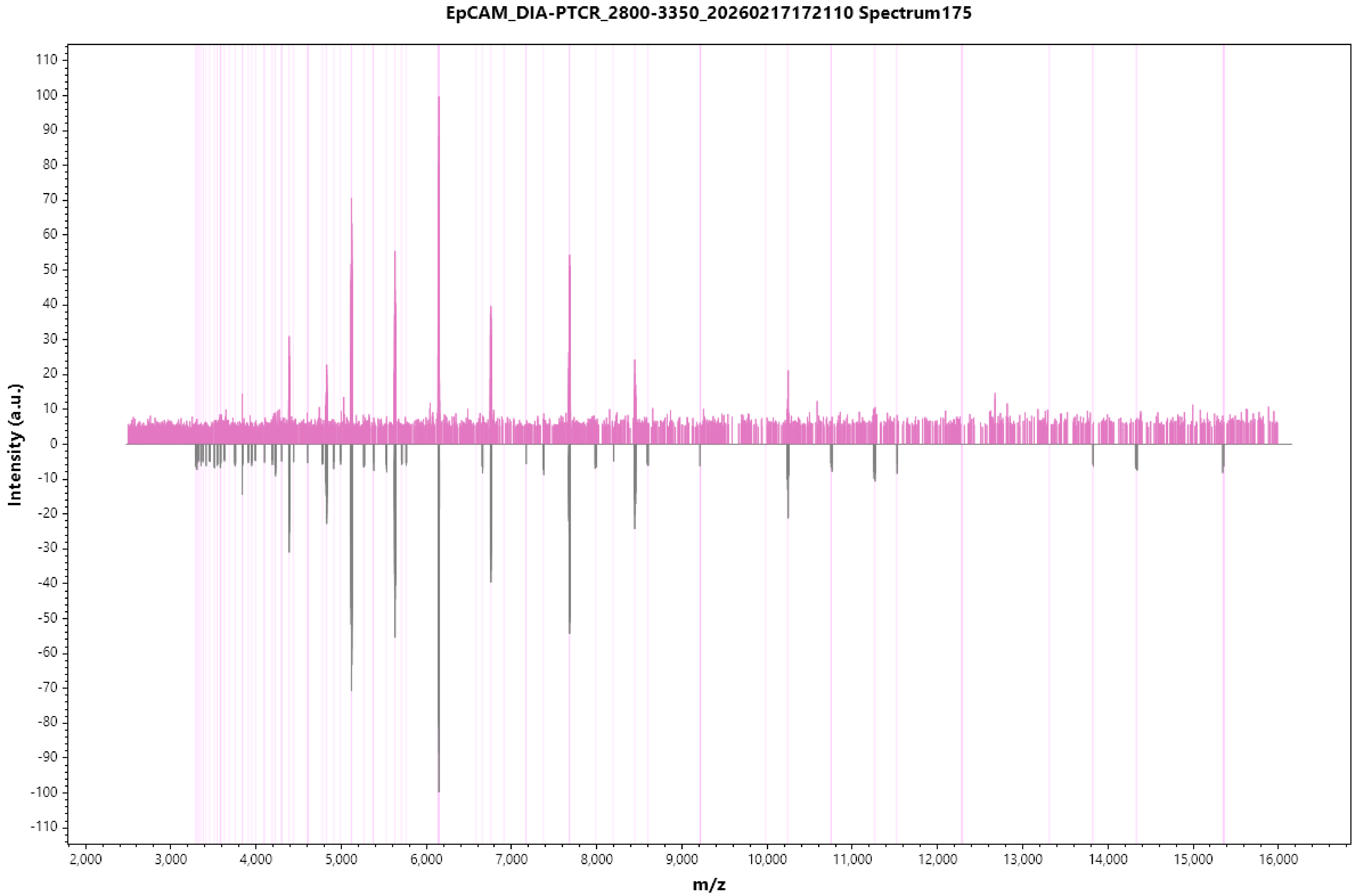
EpCAM_DIA-PTCR_2800-3350_20260217172110 Spectrum175
Intensity (a.u.)
m/z
110
100
90
80
70
60
50
40
30
20
10
0
-10
-20
-30
-40
-50
-60
-70
-80
-90
-100
-110
2,000
3,000
4,000
5,000
6,000
7,000
8,000
9,000
10,000
11,000
12,000
13,000
14,000
15,000
16,000

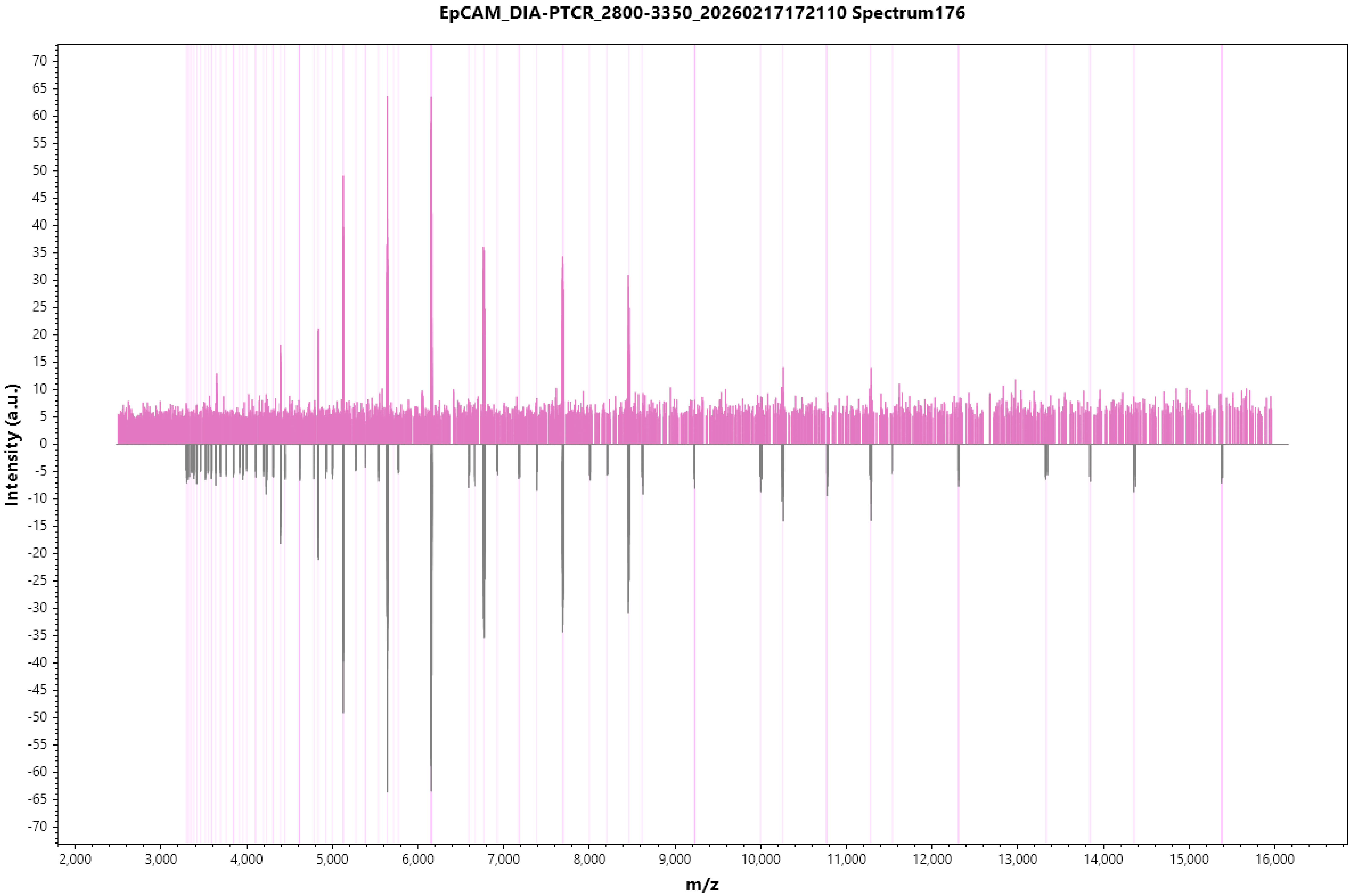
EpCAM_DIA-PTCR_2800-3350_20260217172110 Spectrum176
Intensity (a.u.)
70
65
60
55
50
45
40
35
30
25
20
15
10
5
0
-5
-10
-15
-20
-25
-30
-35
-40
-45
-50
-55
-60
-65
-70
2,000
3,000
4,000
5,000
6,000
7,000
8,000
9,000
10,000
11,000
12,000
13,000
14,000
15,000
16,000
m/z

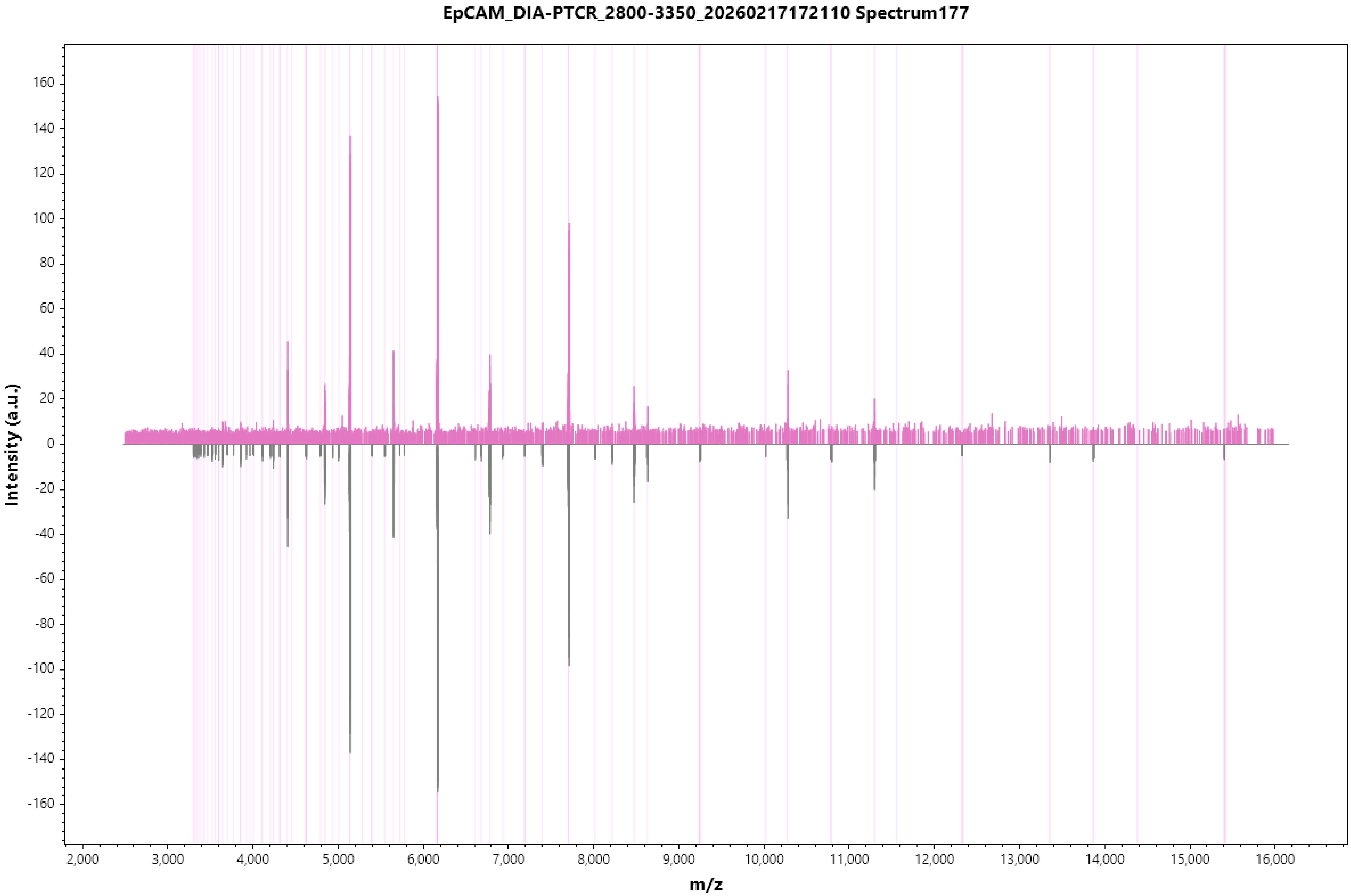
EpCAM_DIA-PTCR_2800-3350_20260217172110 Spectrum177
Intensity (a.u.)
160
140
120
100
80
60
40
20
0
-20
-40
-60
-80
-100
-120
-140
-160
2,000
3,000
4,000
5,000
6,000
7,000
8,000
9,000
10,000
11,000
12,000
13,000
14,000
15,000
16,000
m/z

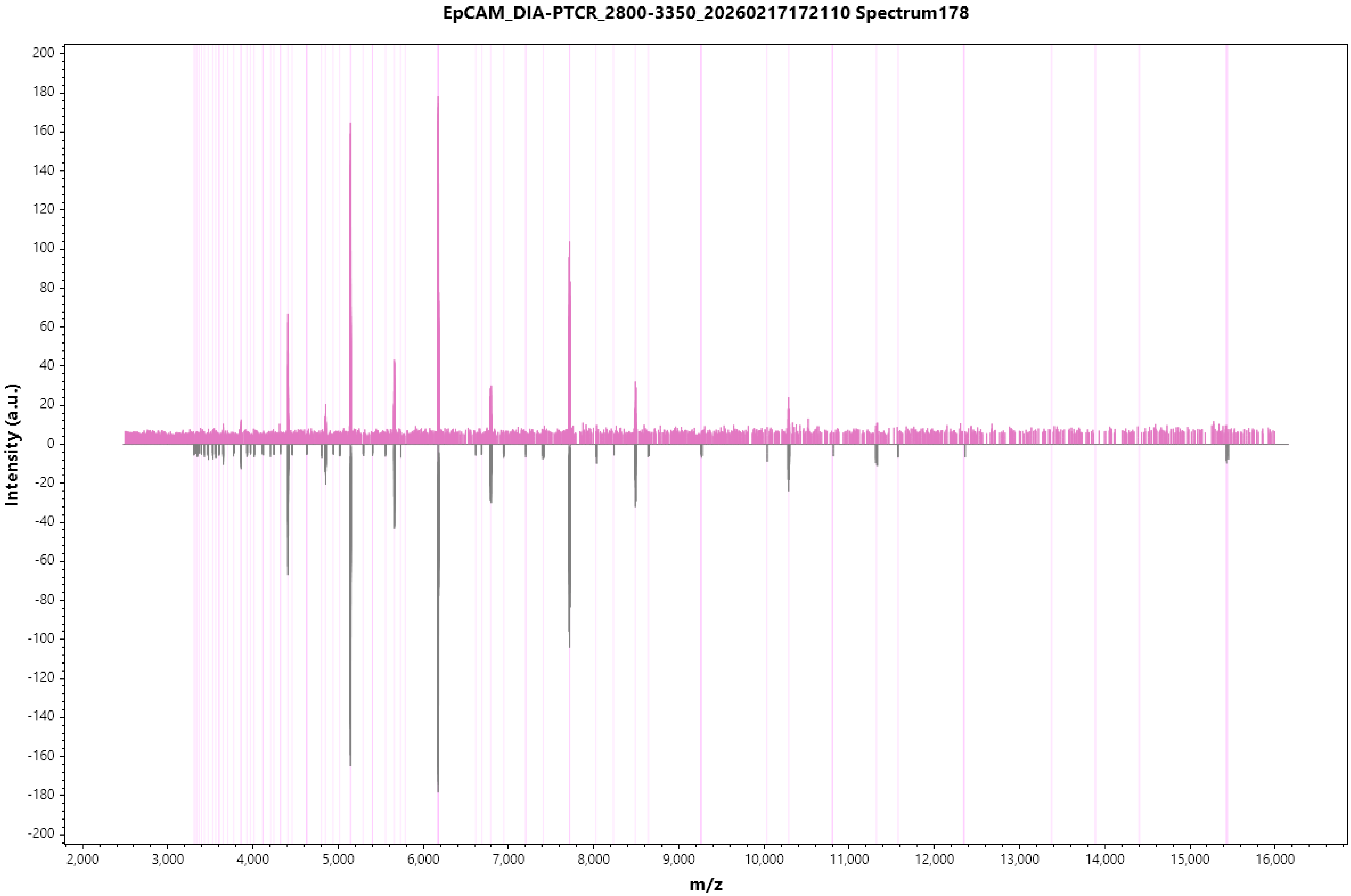
EpCAM_DIA-PTCR_2800-3350_20260217172110 Spectrum178
Intensity (a.u.)
m/z
200
180
160
140
120
100
80
60
40
20
0
-20
-40
-60
-80
-100
-120
-140
-160
-180
-200
2,000
3,000
4,000
5,000
6,000
7,000
8,000
9,000
10,000
11,000
12,000
13,000
14,000
15,000
16,000

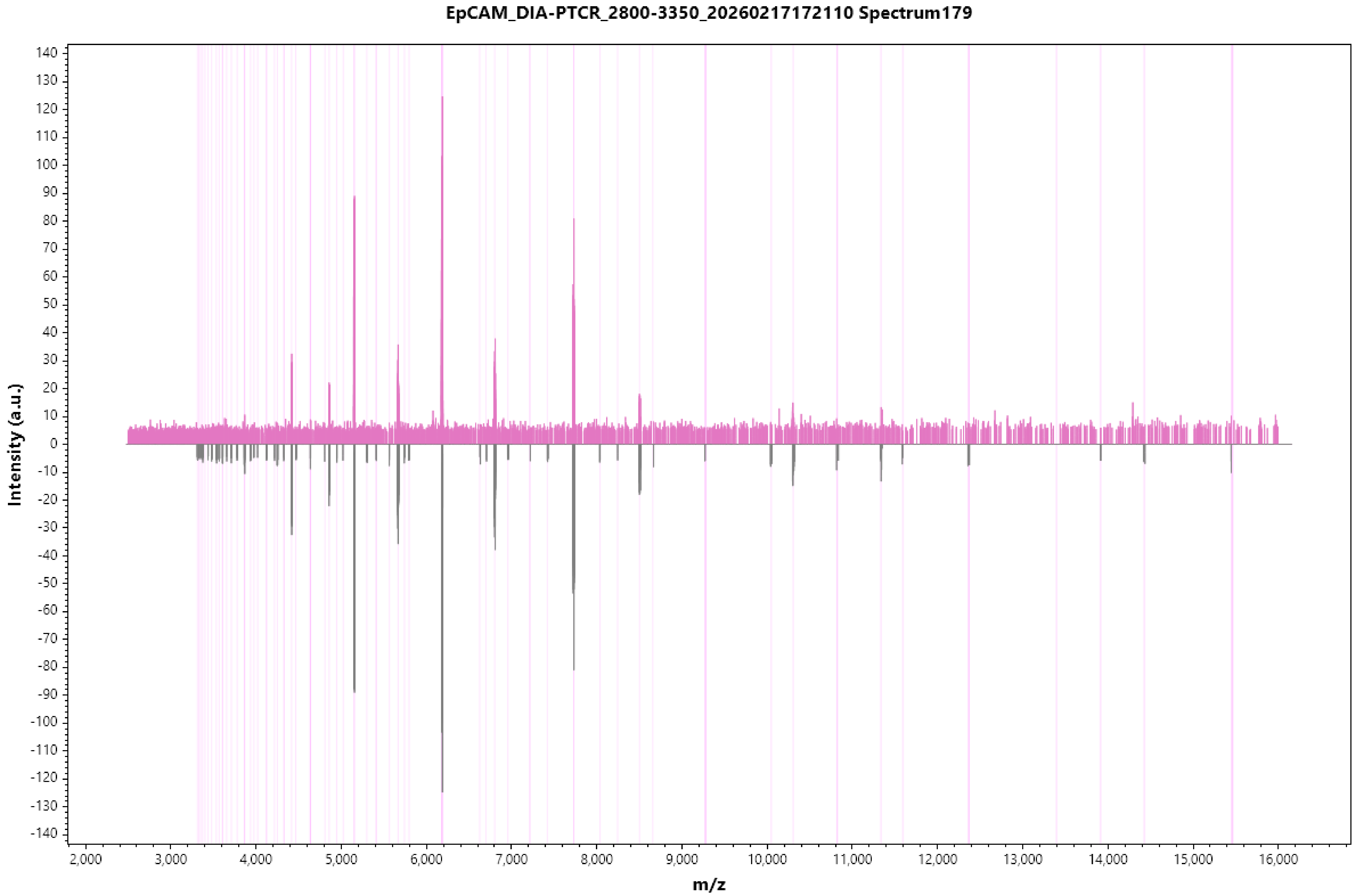

EpCAM_DIA-PTCR_2800-3350_20260217172110 Spectrum179
Intensity (a.u.)
m/z
140
130
120
110
100
90
80
70
60
50
40
30
20
10
0
-10
-20
-30
-40
-50
-60
-70
-80
-90
-100
-110
-120
-130
-140
2,000
3,000
4,000
5,000
6,000
7,000
8,000
9,000
10,000
11,000
12,000
13,000
14,000
15,000
16,000

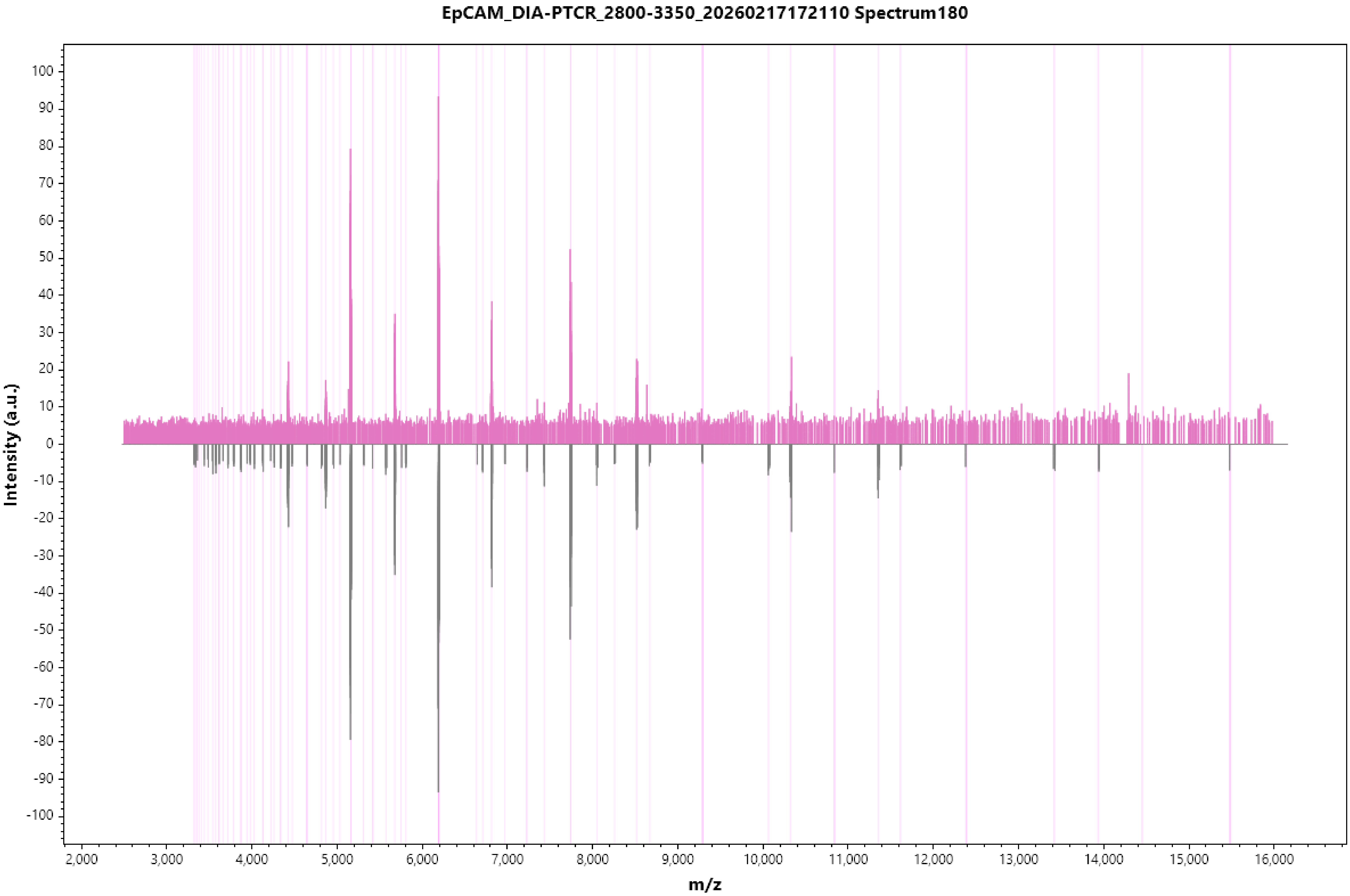

EpCAM_DIA-PTCR_2800-3350_20260217172110 Spectrum180
Intensity (a.u.)
m/z

EpCAM_DIA-PTCR_2800-3350_20260217172110 Spectrum181

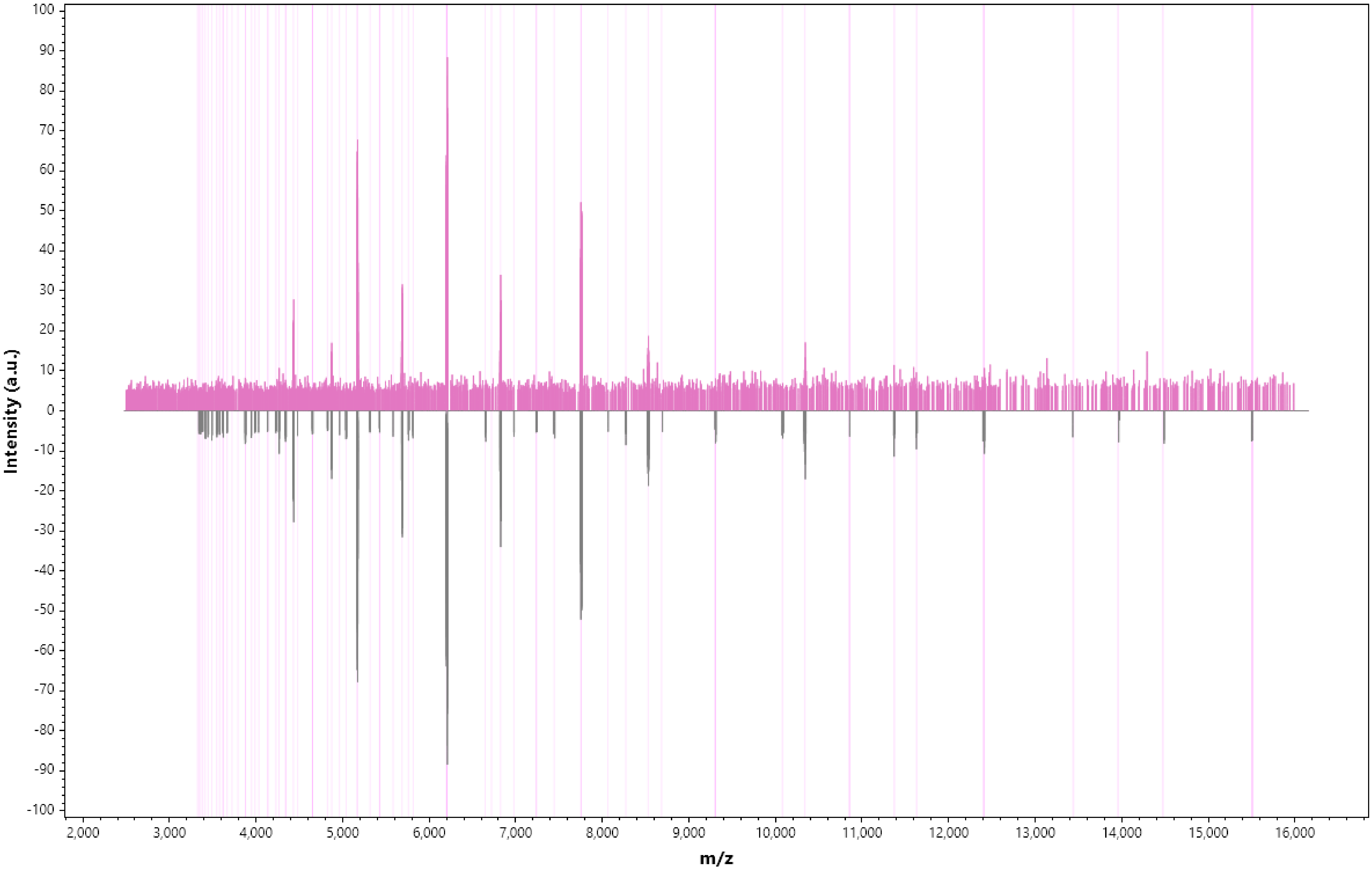

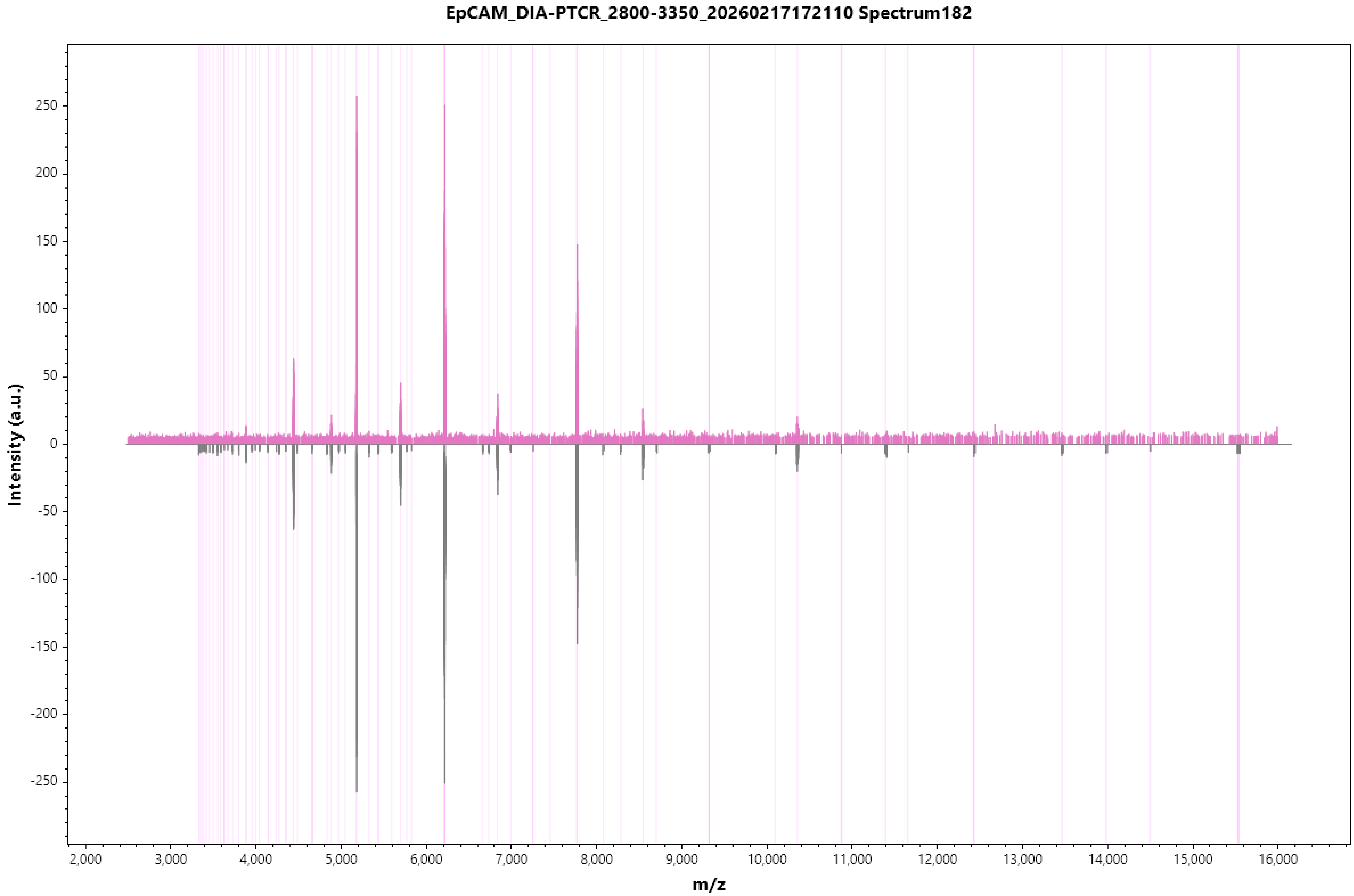

EpCAM_DIA-PTCR_2800-3350_20260217172110 Spectrum182
Intensity (a.u.)
m/z
250
200
150
100
50
0
-50
-100
-150
-200
-250
2,000
3,000
4,000
5,000
6,000
7,000
8,000
9,000
10,000
11,000
12,000
13,000
14,000
15,000
16,000

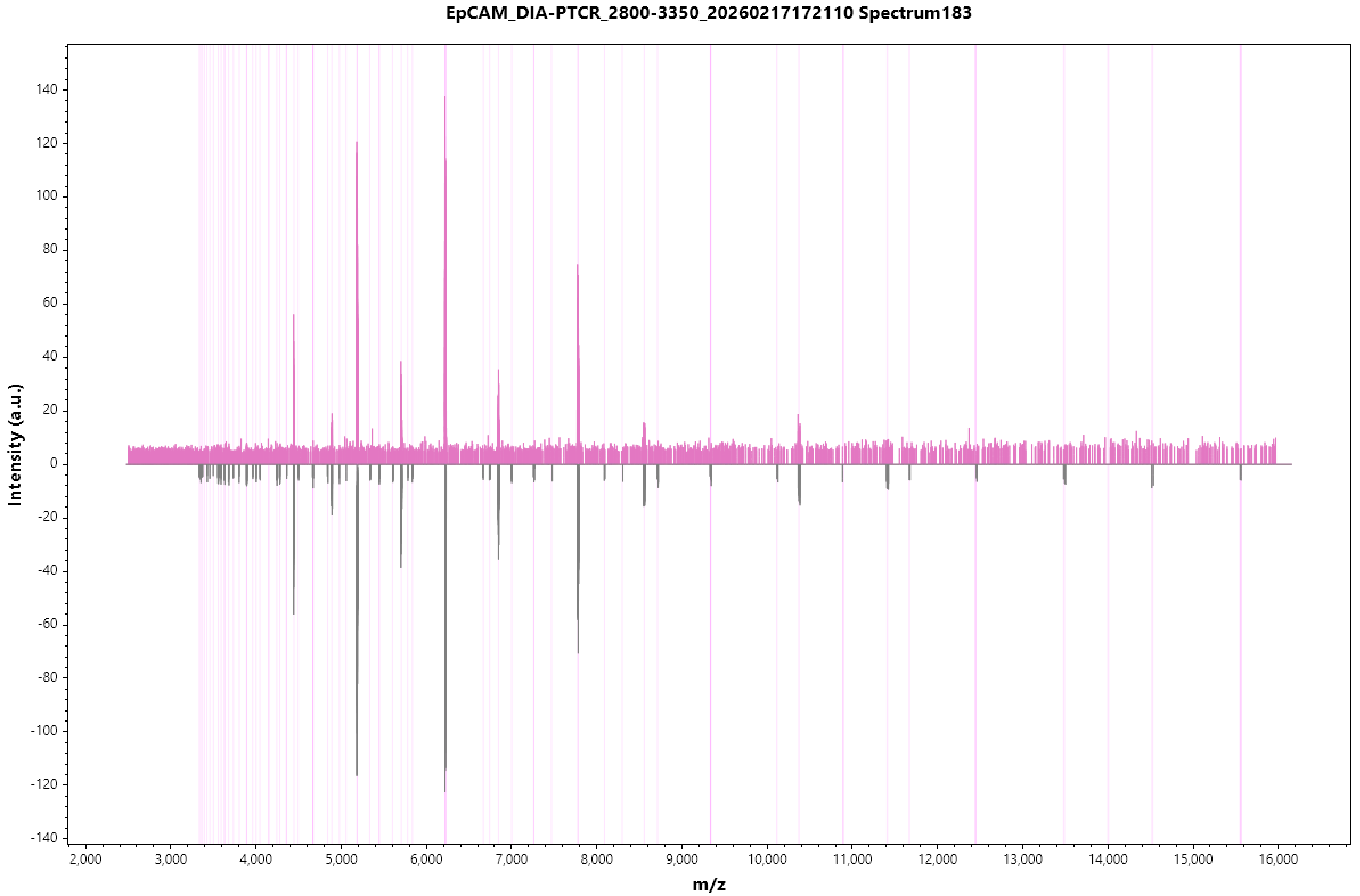

EpCAM_DIA-PTCR_2800-3350_20260217172110 Spectrum183
Intensity (a.u.)
m/z
140
120
100
80
60
40
20
0
-20
-40
-60
-80
-100
-120
-140
2,000
3,000
4,000
5,000
6,000
7,000
8,000
9,000
10,000
11,000
12,000
13,000
14,000
15,000
16,000

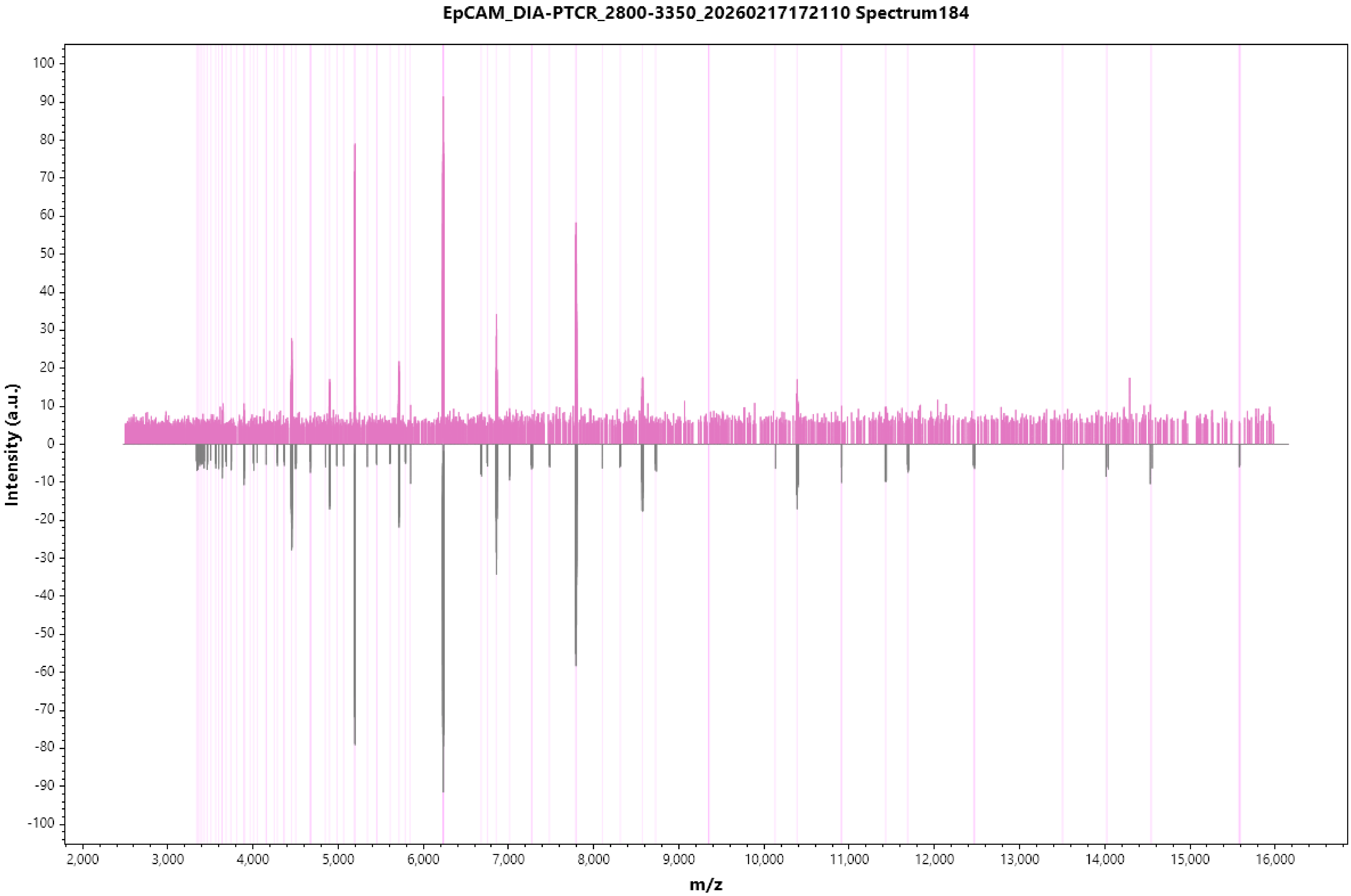
EpCAM_DIA-PTCR_2800-3350_20260217172110 Spectrum184
Intensity (a.u.)
m/z
100
90
80
70
60
50
40
30
20
10
0
-10
-20
-30
-40
-50
-60
-70
-80
-90
-100
2,000
3,000
4,000
5,000
6,000
7,000
8,000
9,000
10,000
11,000
12,000
13,000
14,000
15,000
16,000

#### EpCAM_DIA-PTCR_2800-3350_20260217172110 Spectrum185

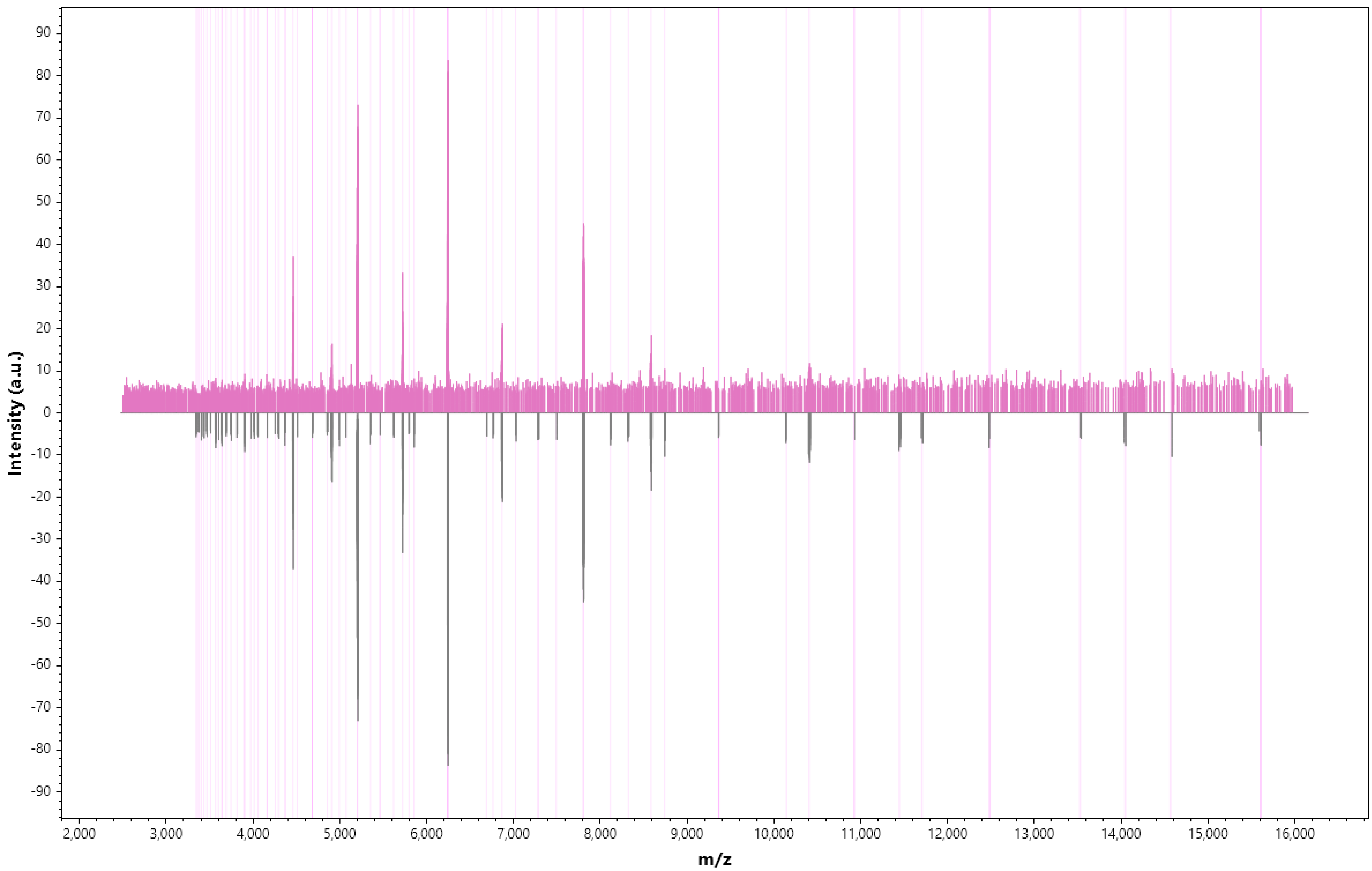

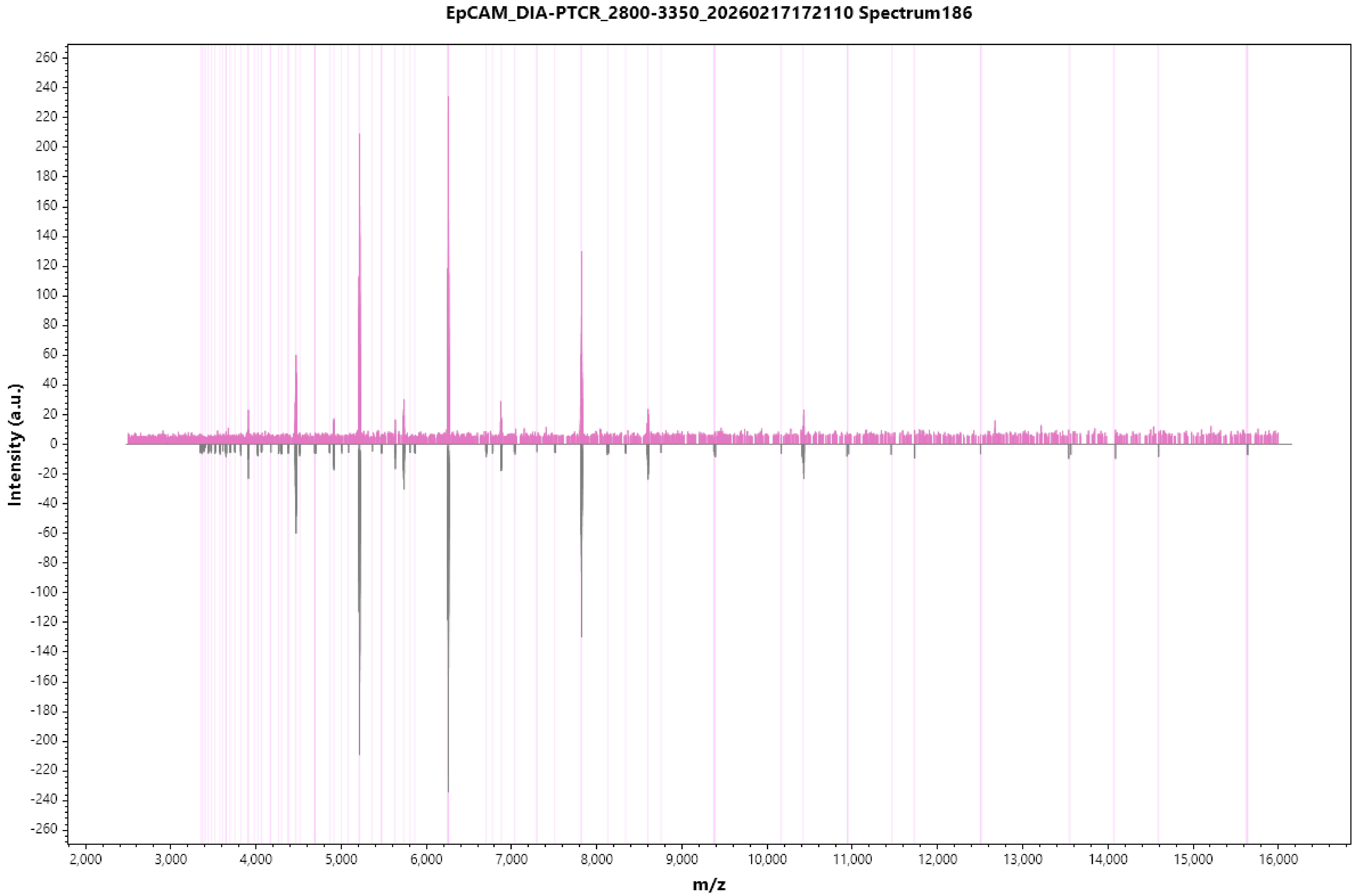

EpCAM_DIA-PTCR_2800-3350_20260217172110 Spectrum186
Intensity (a.u.)
m/z
260
240
220
200
180
160
140
120
100
80
60
40
20
0
-20
-40
-60
-80
-100
-120
-140
-160
-180
-200
-220
-240
-260
2,000
3,000
4,000
5,000
6,000
7,000
8,000
9,000
10,000
11,000
12,000
13,000
14,000
15,000
16,000

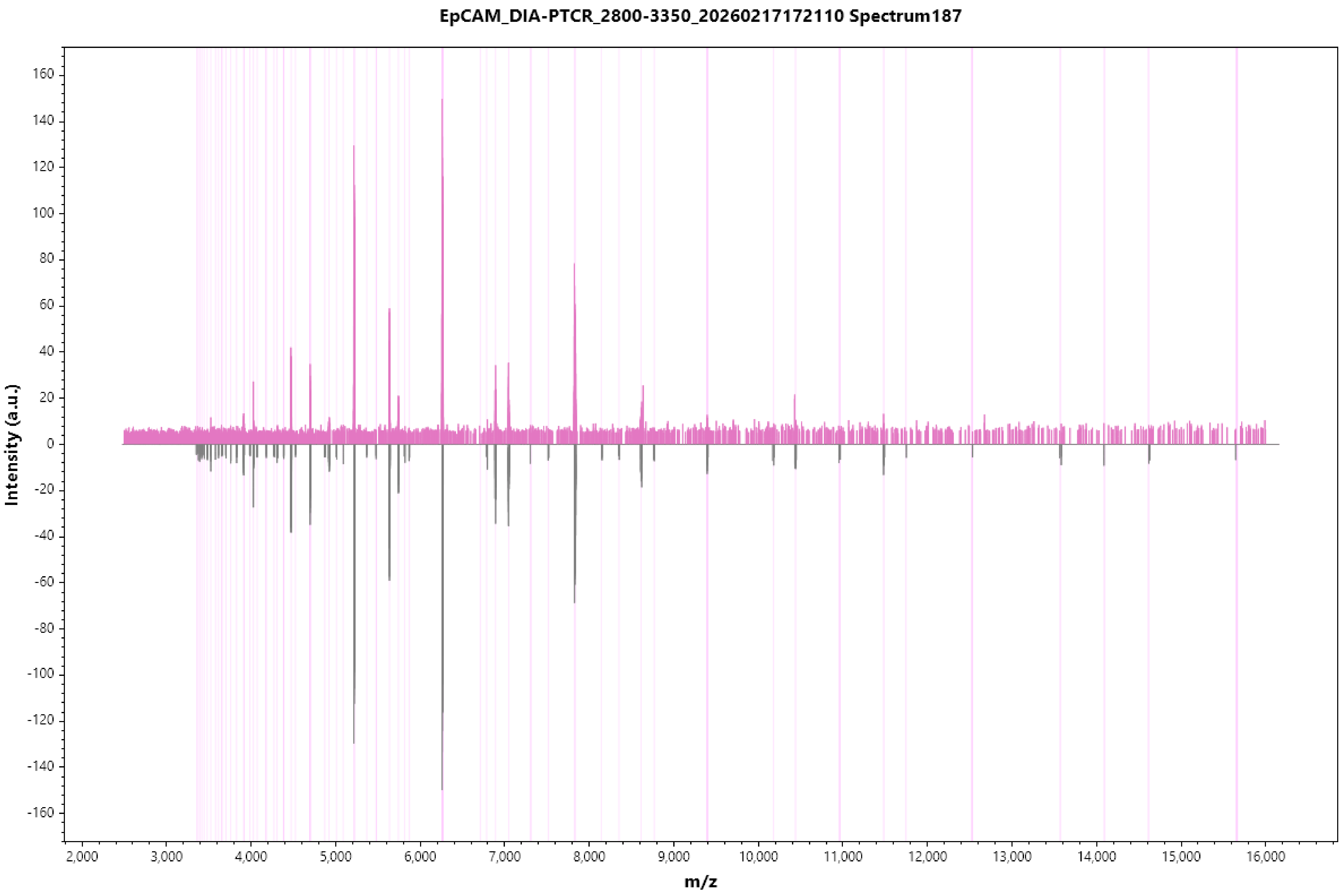
EpCAM_DIA-PTCR_2800-3350_20260217172110 Spectrum187
Intensity (a.u.)
m/z
160
140
120
100
80
60
40
20
0
-20
-40
-60
-80
-100
-120
-140
-160
2,000
3,000
4,000
5,000
6,000
7,000
8,000
9,000
10,000
11,000
12,000
13,000
14,000
15,000
16,000

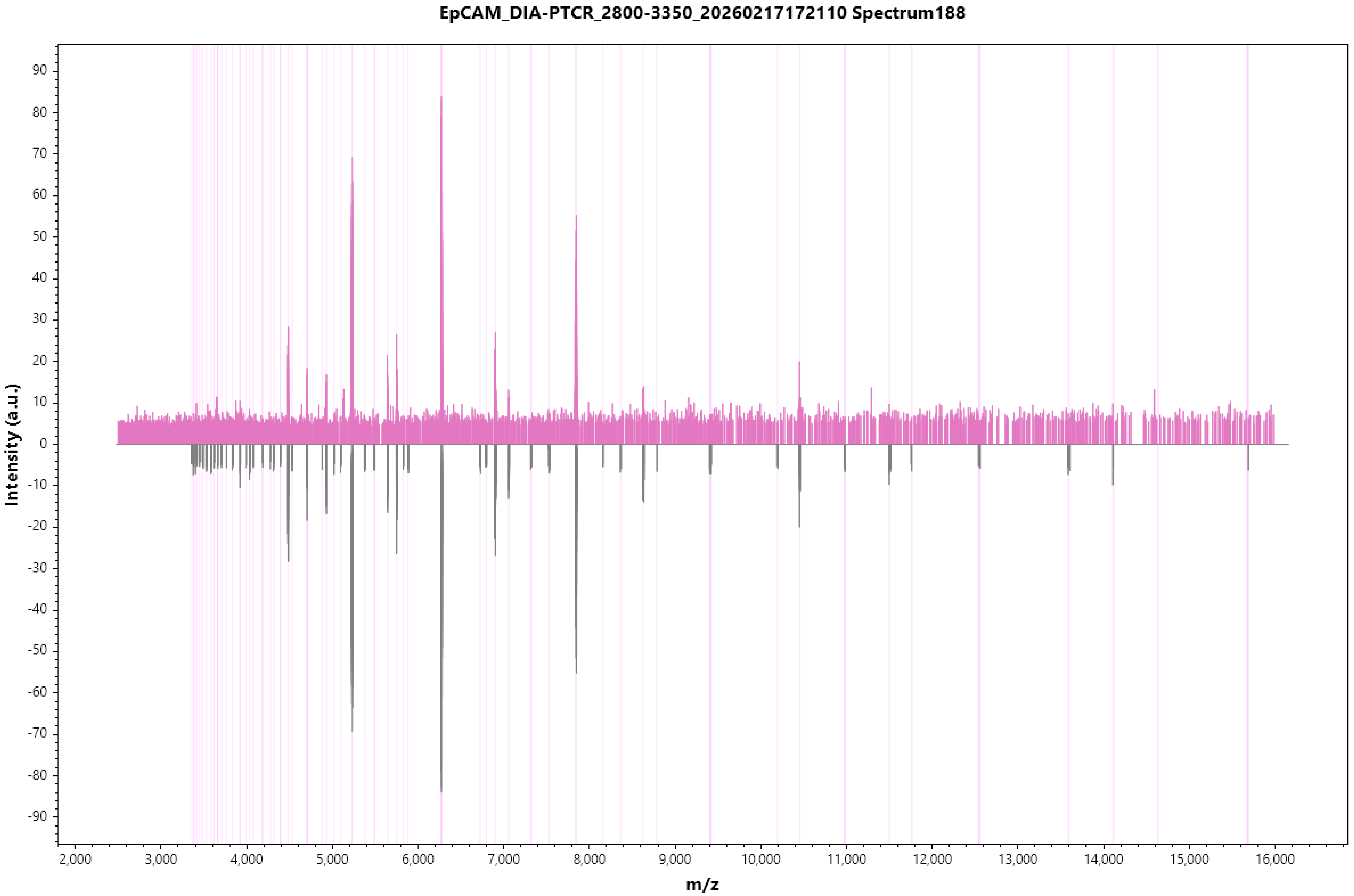
EpCAM_DIA-PTCR_2800-3350_20260217172110 Spectrum188
Intensity (a.u.)
m/z
90
80
70
60
50
40
30
20
10
0
-10
-20
-30
-40
-50
-60
-70
-80
-90
2,000
3,000
4,000
5,000
6,000
7,000
8,000
9,000
10,000
11,000
12,000
13,000
14,000
15,000
16,000

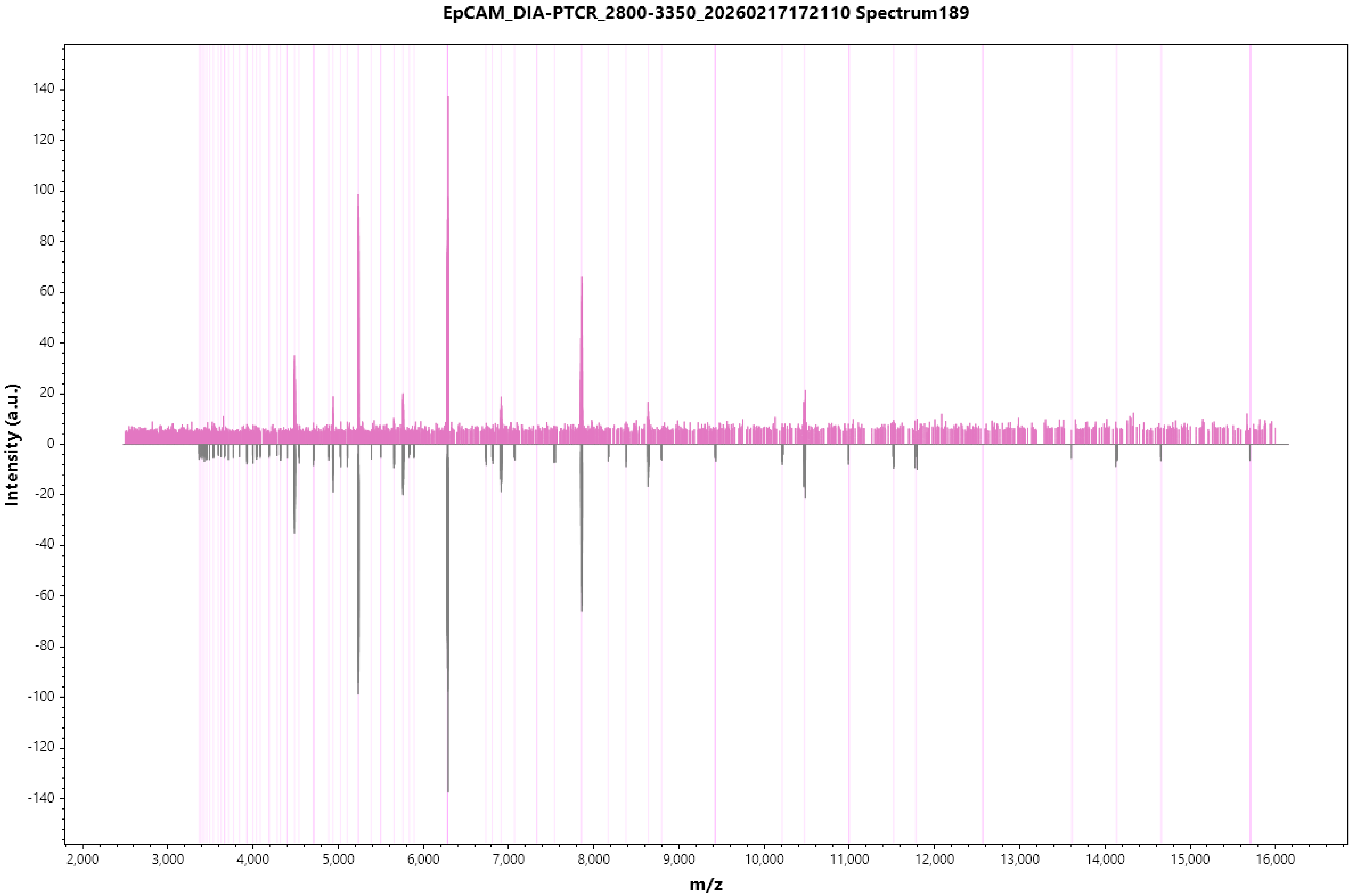
EpCAM_DIA-PTCR_2800-3350_20260217172110 Spectrum189
Intensity (a.u.)
m/z
140
120
100
80
60
40
20
0
-20
-40
-60
-80
-100
-120
-140
2,000
3,000
4,000
5,000
6,000
7,000
8,000
9,000
10,000
11,000
12,000
13,000
14,000
15,000
16,000

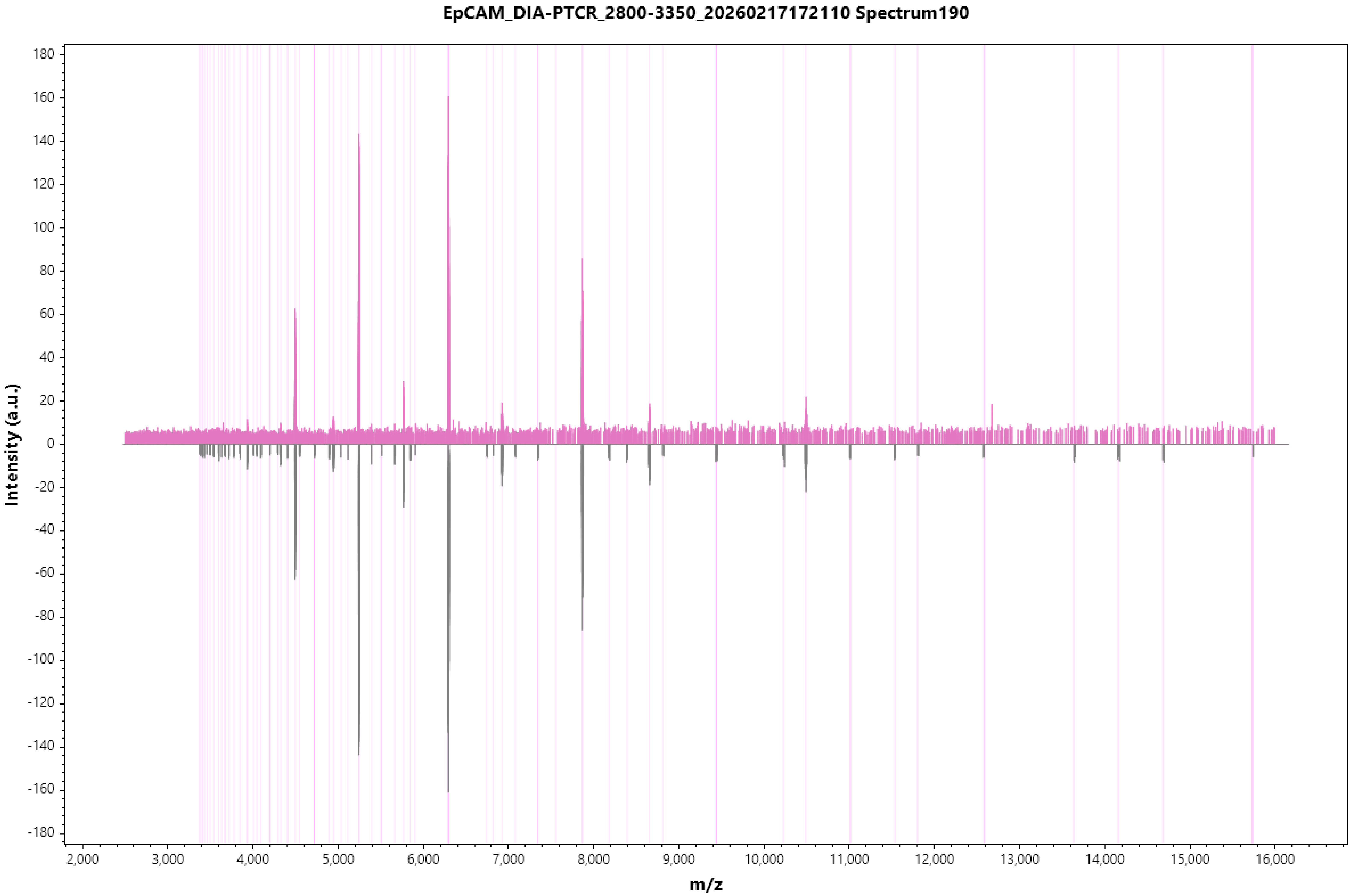
EpCAM_DIA-PTCR_2800-3350_20260217172110 Spectrum190
Intensity (a.u.)
m/z
180
160
140
120
100
80
60
40
20
0
-20
-40
-60
-80
-100
-120
-140
-160
-180
2,000
3,000
4,000
5,000
6,000
7,000
8,000
9,000
10,000
11,000
12,000
13,000
14,000
15,000
16,000

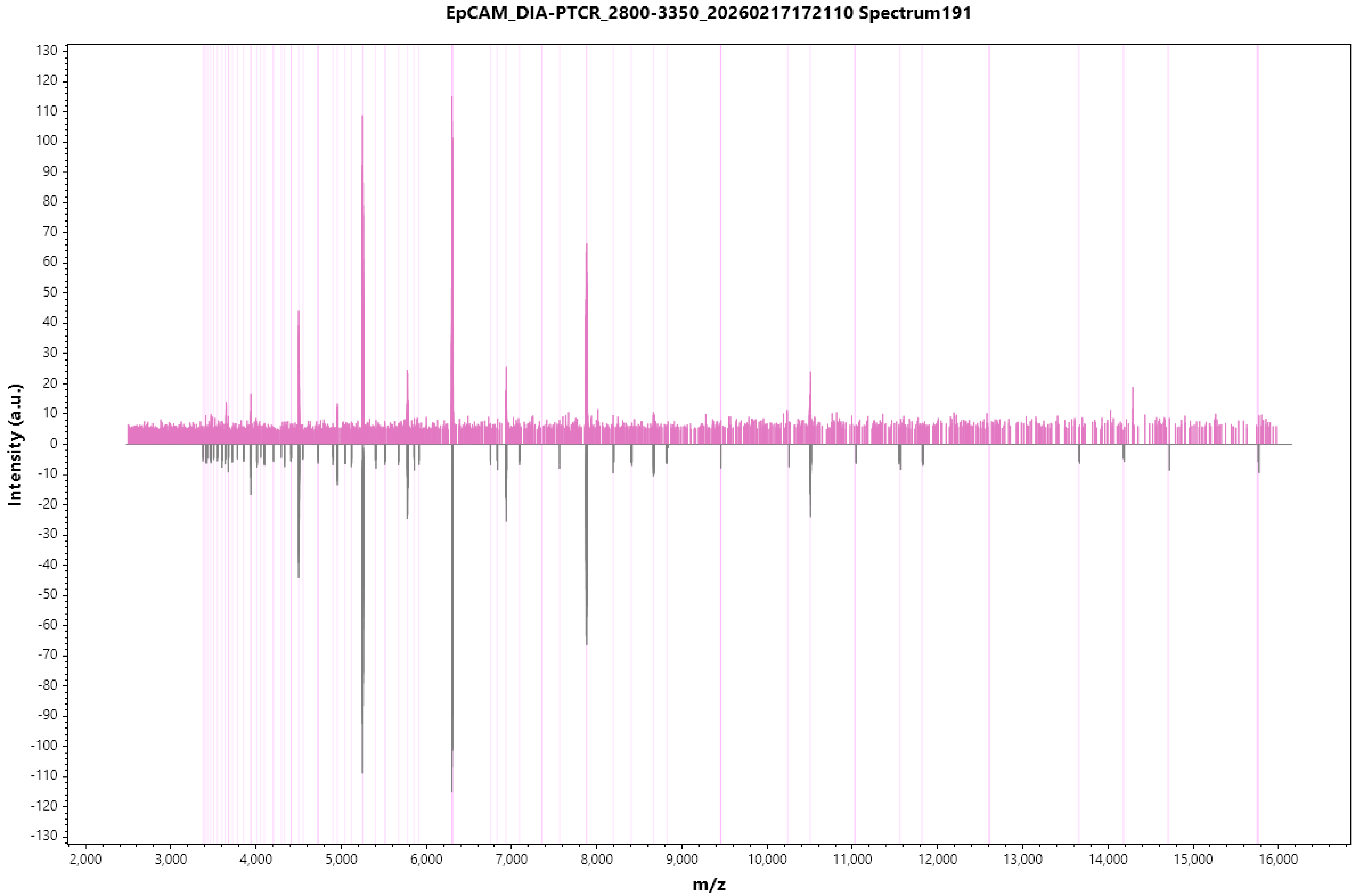
EpCAM_DIA-PTCR_2800-3350_20260217172110 Spectrum191
Intensity (a.u.)
m/z
130
120
110
100
90
80
70
60
50
40
30
20
10
0
-10
-20
-30
-40
-50
-60
-70
-80
-90
-100
-110
-120
-130
2,000
3,000
4,000
5,000
6,000
7,000
8,000
9,000
10,000
11,000
12,000
13,000
14,000
15,000
16,000

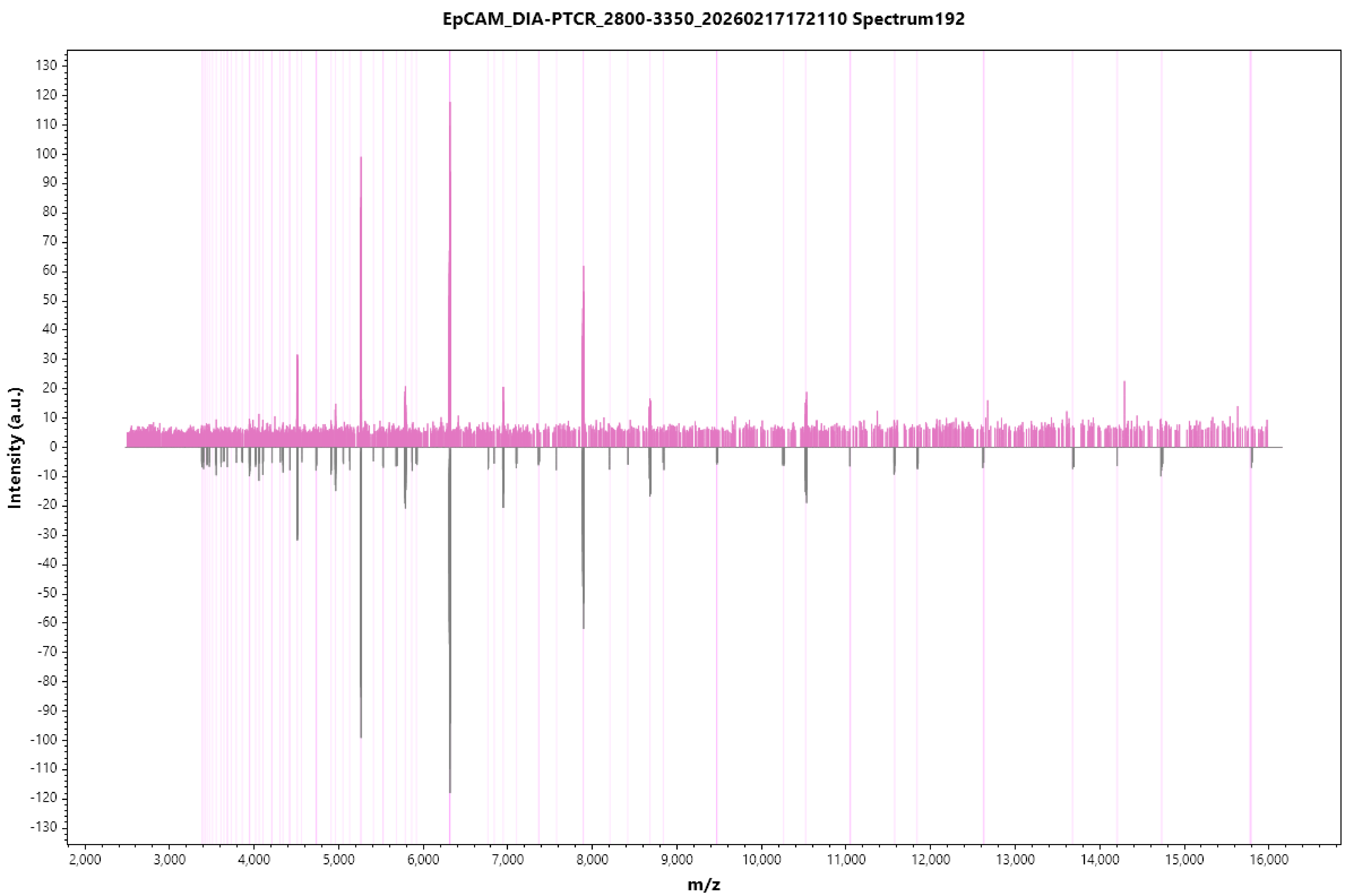
EpCAM_DIA-PTCR_2800-3350_20260217172110 Spectrum192
Intensity (a.u.)
m/z

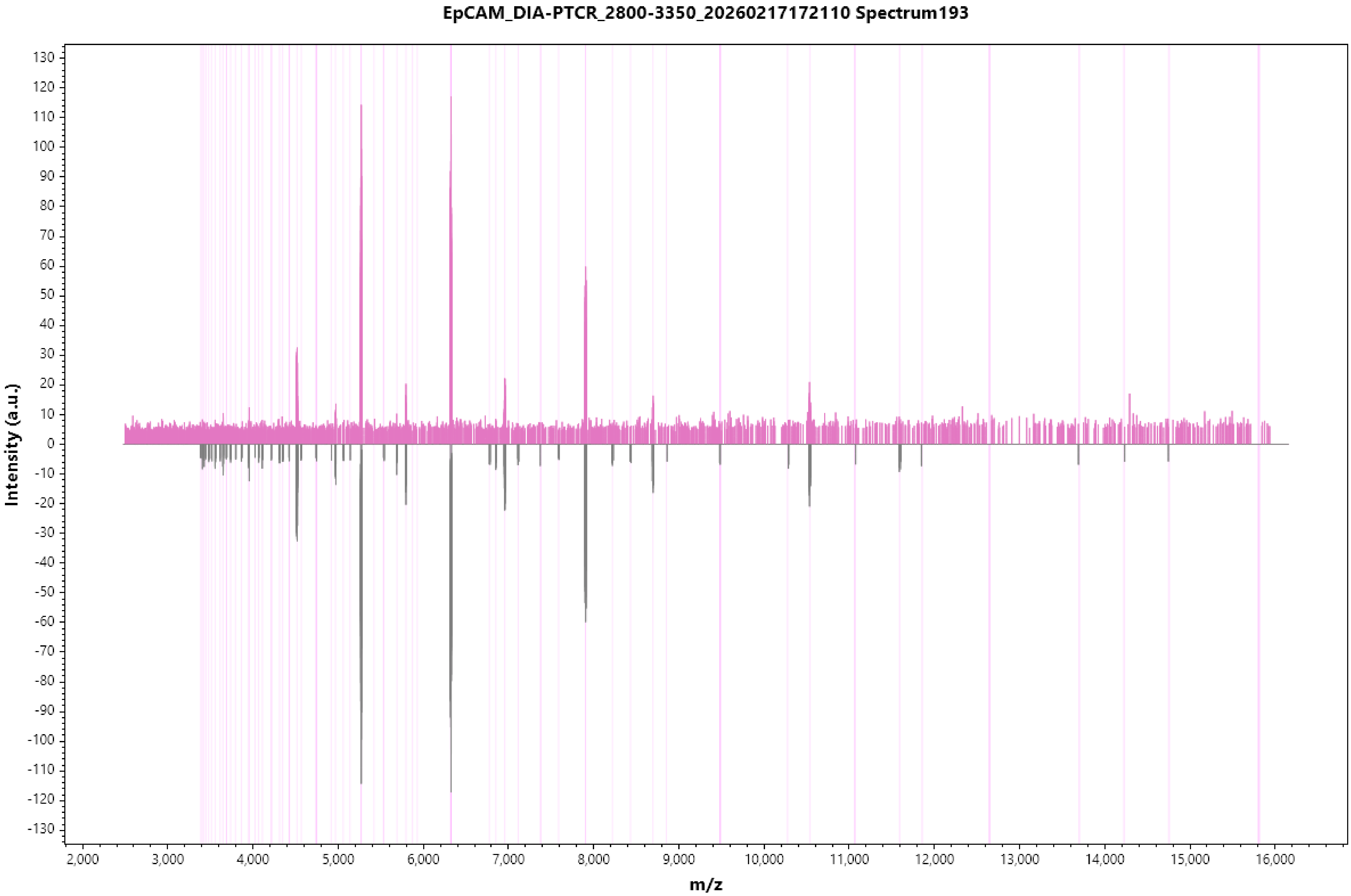
EpCAM_DIA-PTCR_2800-3350_20260217172110 Spectrum193
Intensity (a.u.)
m/z

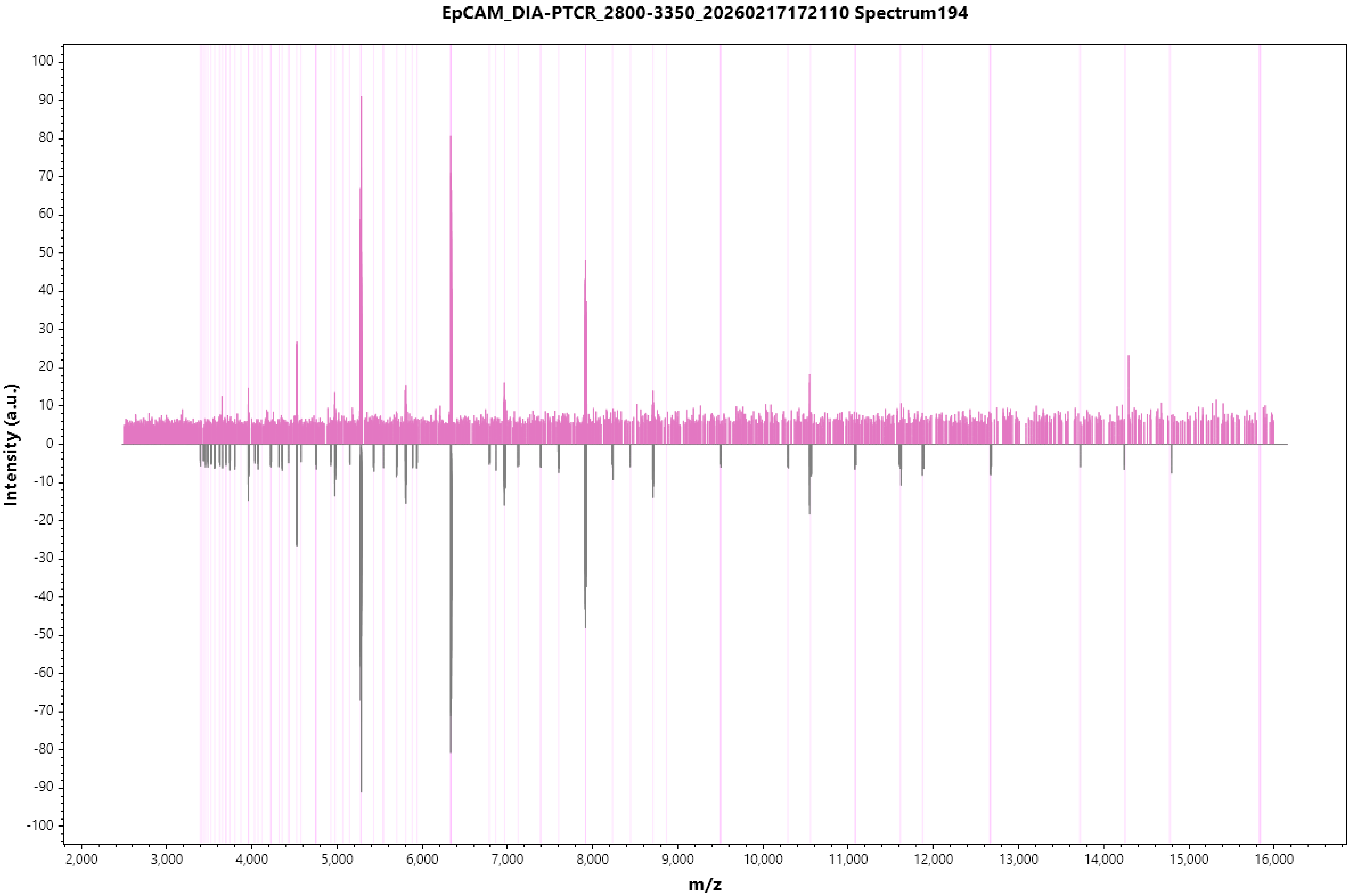
EpCAM_DIA-PTCR_2800-3350_20260217172110 Spectrum194
Intensity (a.u.)
m/z

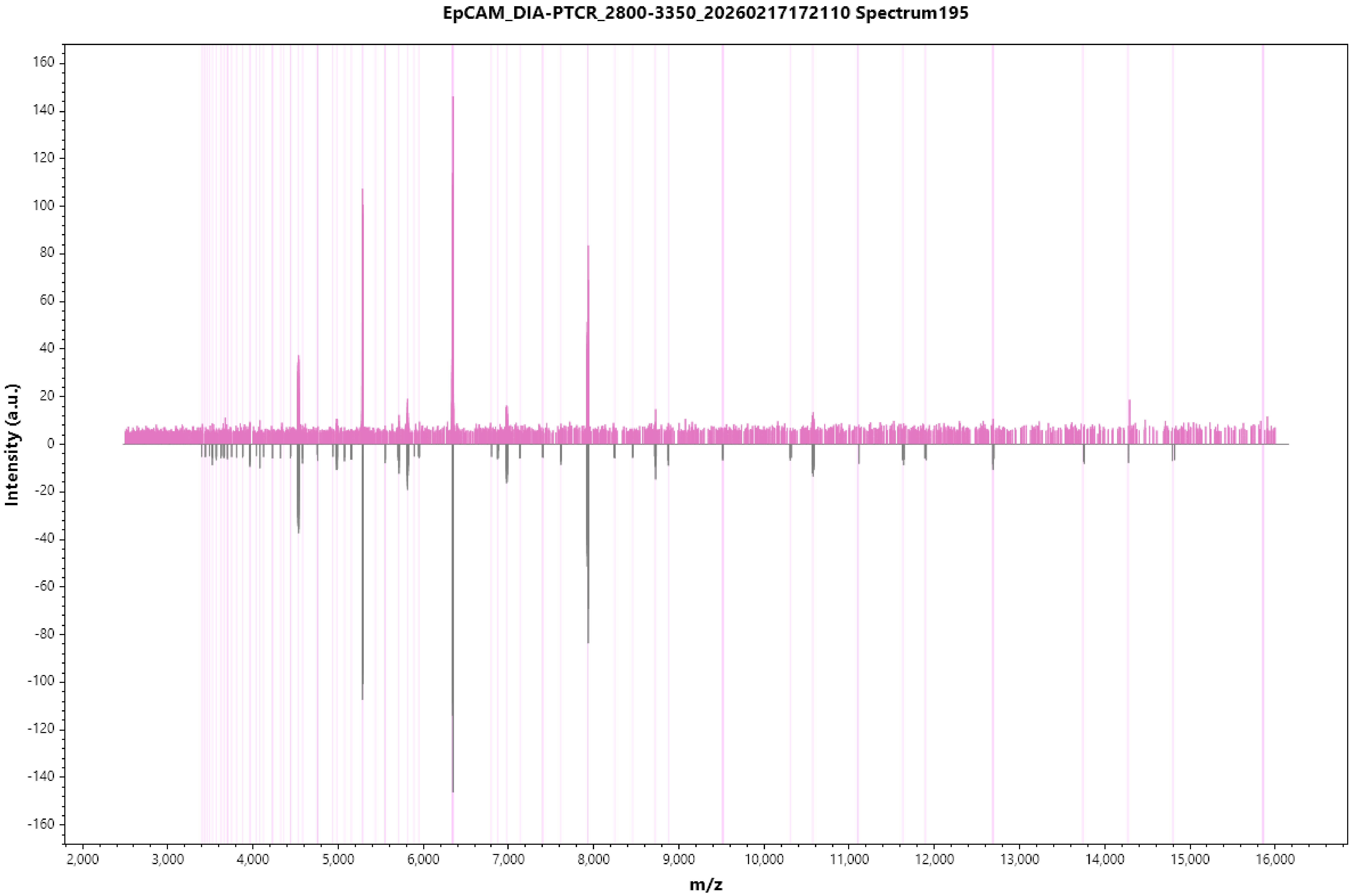
EpCAM_DIA-PTCR_2800-3350_20260217172110 Spectrum195
Intensity (a.u.)
160
140
120
100
80
60
40
20
0
-20
-40
-60
-80
-100
-120
-140
-160
2,000
3,000
4,000
5,000
6,000
7,000
8,000
9,000
10,000
11,000
12,000
13,000
14,000
15,000
16,000
m/z

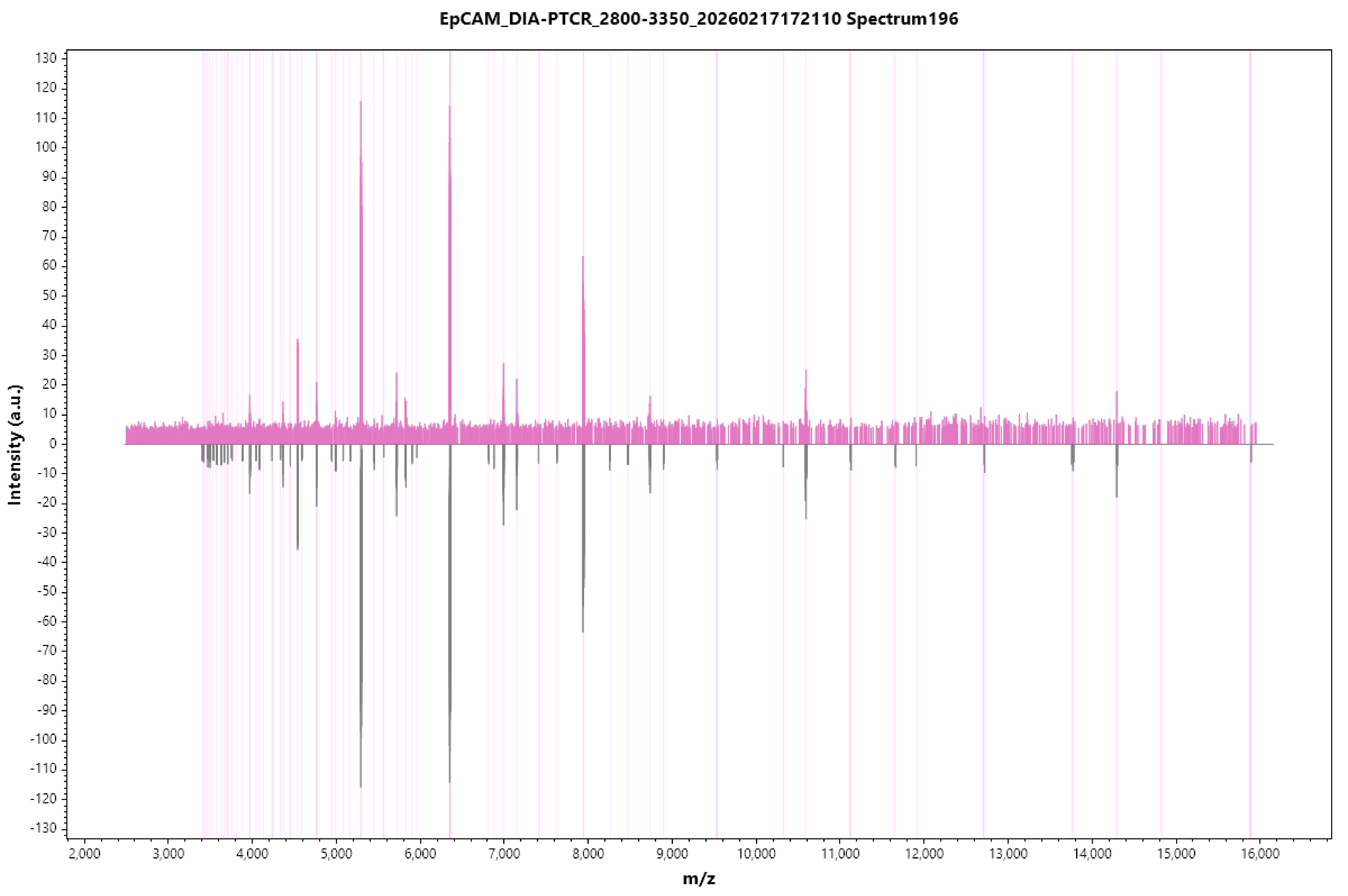
EpCAM_DIA-PTCR_2800-3350_20260217172110 Spectrum196
Intensity (a.u.)
m/z
130
120
110
100
90
80
70
60
50
40
30
20
10
0
-10
-20
-30
-40
-50
-60
-70
-80
-90
-100
-110
-120
-130
2,000
3,000
4,000
5,000
6,000
7,000
8,000
9,000
10,000
11,000
12,000
13,000
14,000
15,000
16,000

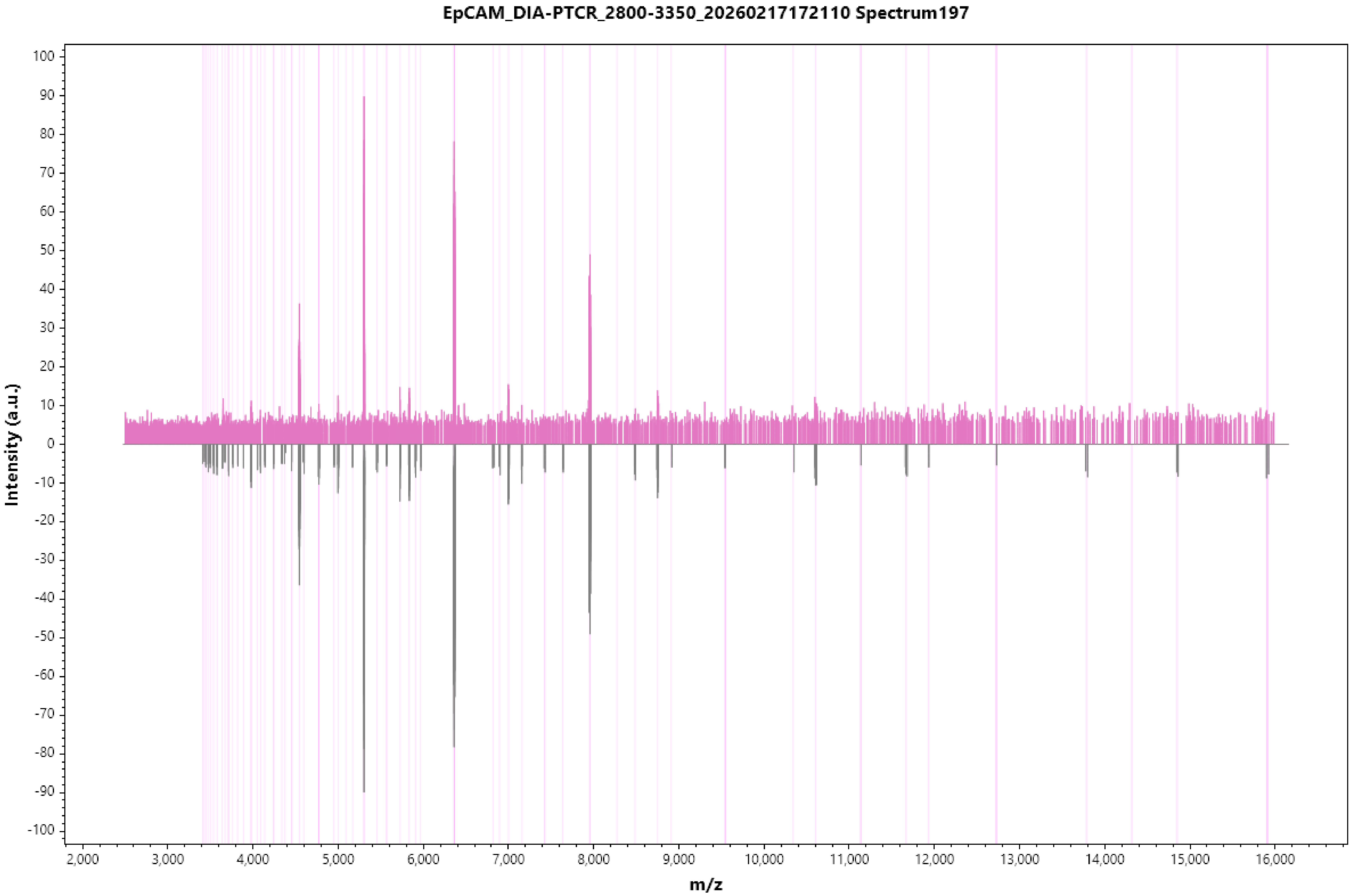
EpCAM_DIA-PTCR_2800-3350_20260217172110 Spectrum197
Intensity (a.u.)
m/z
100
90
80
70
60
50
40
30
20
10
0
-10
-20
-30
-40
-50
-60
-70
-80
-90
-100
2,000
3,000
4,000
5,000
6,000
7,000
8,000
9,000
10,000
11,000
12,000
13,000
14,000
15,000
16,000

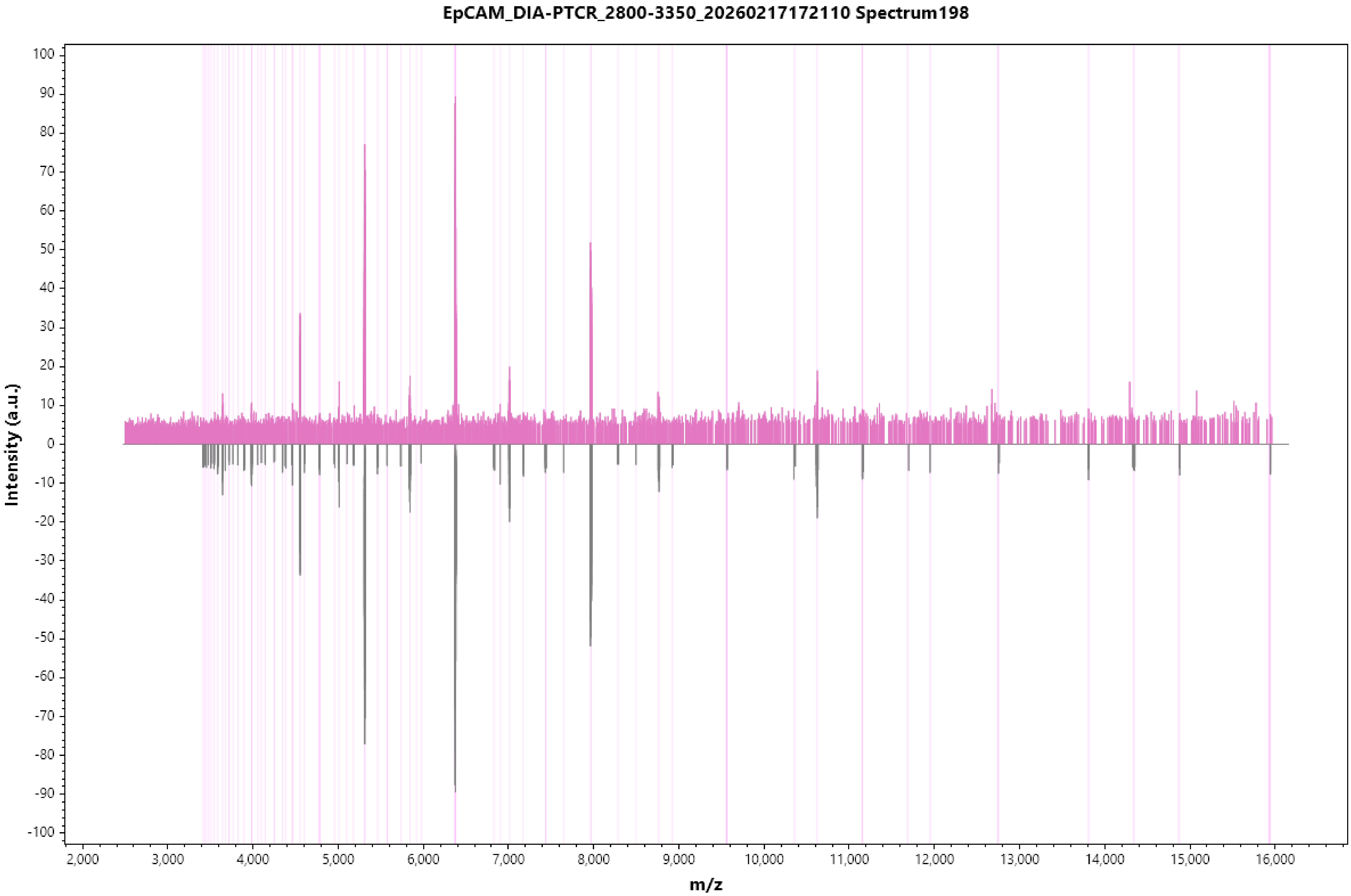
EpCAM_DIA-PTCR_2800-3350_20260217172110 Spectrum198
Intensity (a.u.)
100
90
80
70
60
50
40
30
20
10
0
-10
-20
-30
-40
-50
-60
-70
-80
-90
-100
2,000
3,000
4,000
5,000
6,000
7,000
8,000
9,000
10,000
11,000
12,000
13,000
14,000
15,000
16,000
m/z

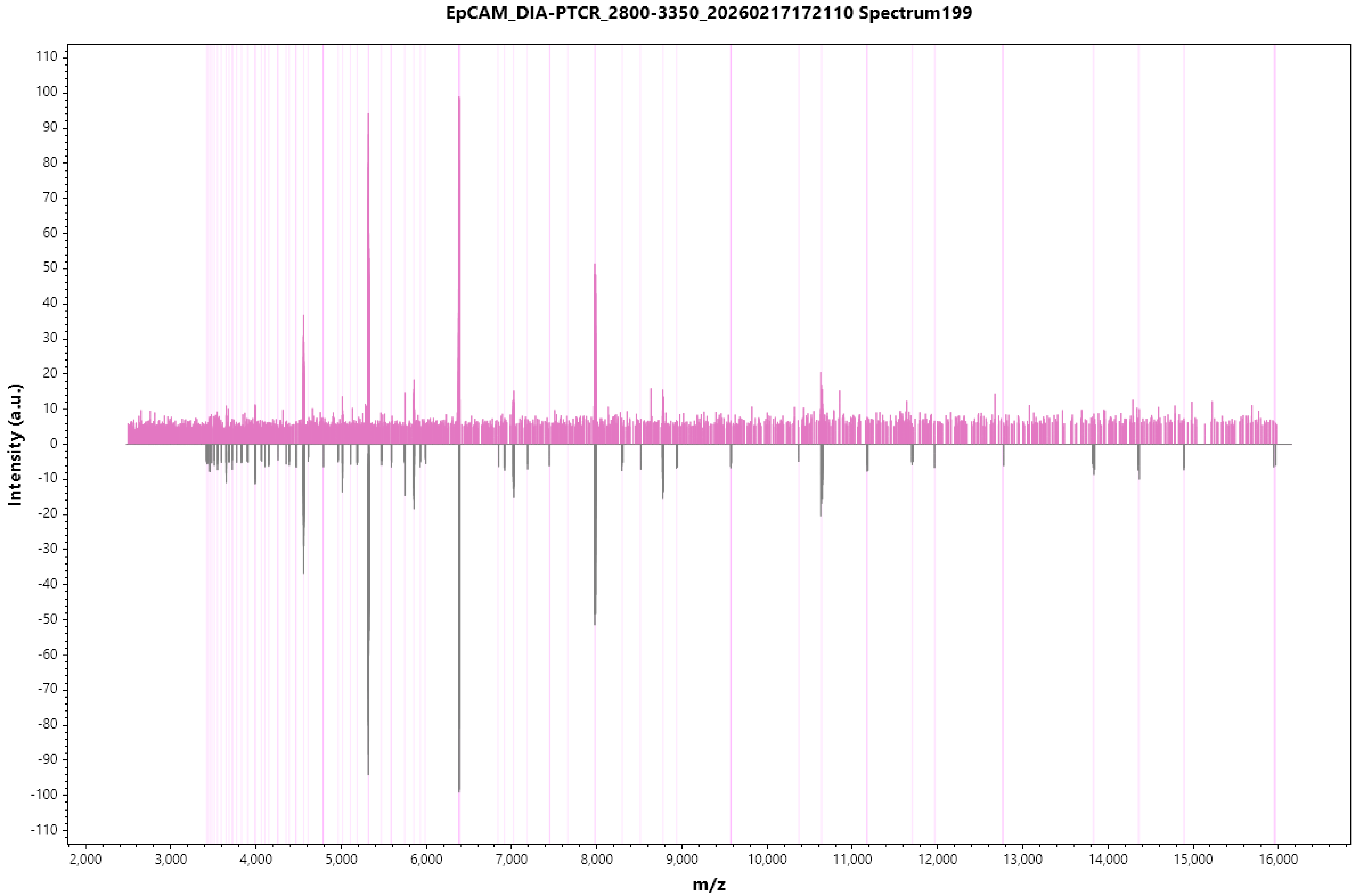

EpCAM_DIA-PTCR_2800-3350_20260217172110 Spectrum199
Intensity (a.u.)
m/z
110
100
90
80
70
60
50
40
30
20
10
0
-10
-20
-30
-40
-50
-60
-70
-80
-90
-100
-110
2,000
3,000
4,000
5,000
6,000
7,000
8,000
9,000
10,000
11,000
12,000
13,000
14,000
15,000
16,000

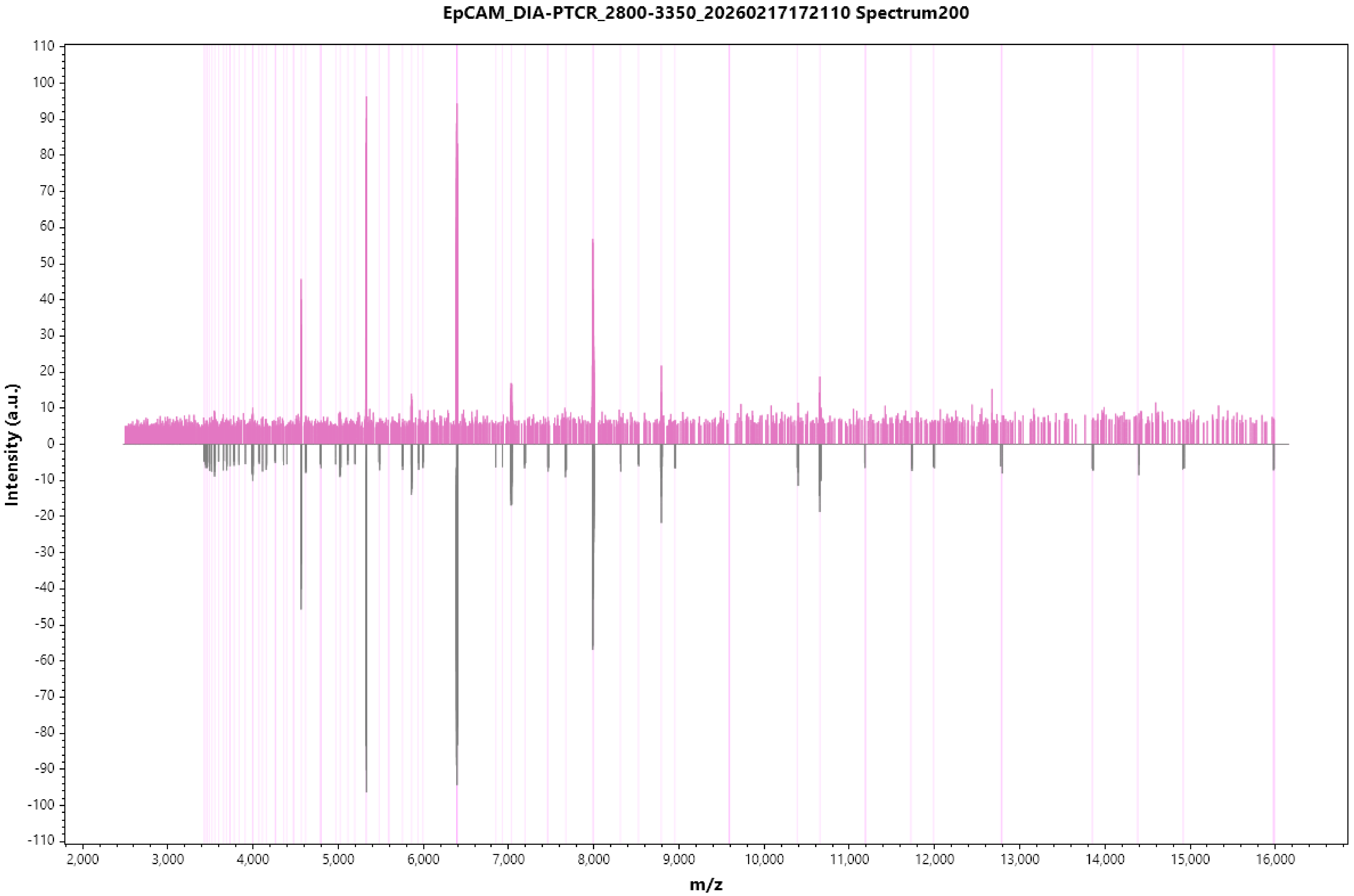
EpCAM_DIA-PTCR_2800-3350_20260217172110 Spectrum200
Intensity (a.u.)
m/z
110
100
90
80
70
60
50
40
30
20
10
0
-10
-20
-30
-40
-50
-60
-70
-80
-90
-100
-110
2,000
3,000
4,000
5,000
6,000
7,000
8,000
9,000
10,000
11,000
12,000
13,000
14,000
15,000
16,000

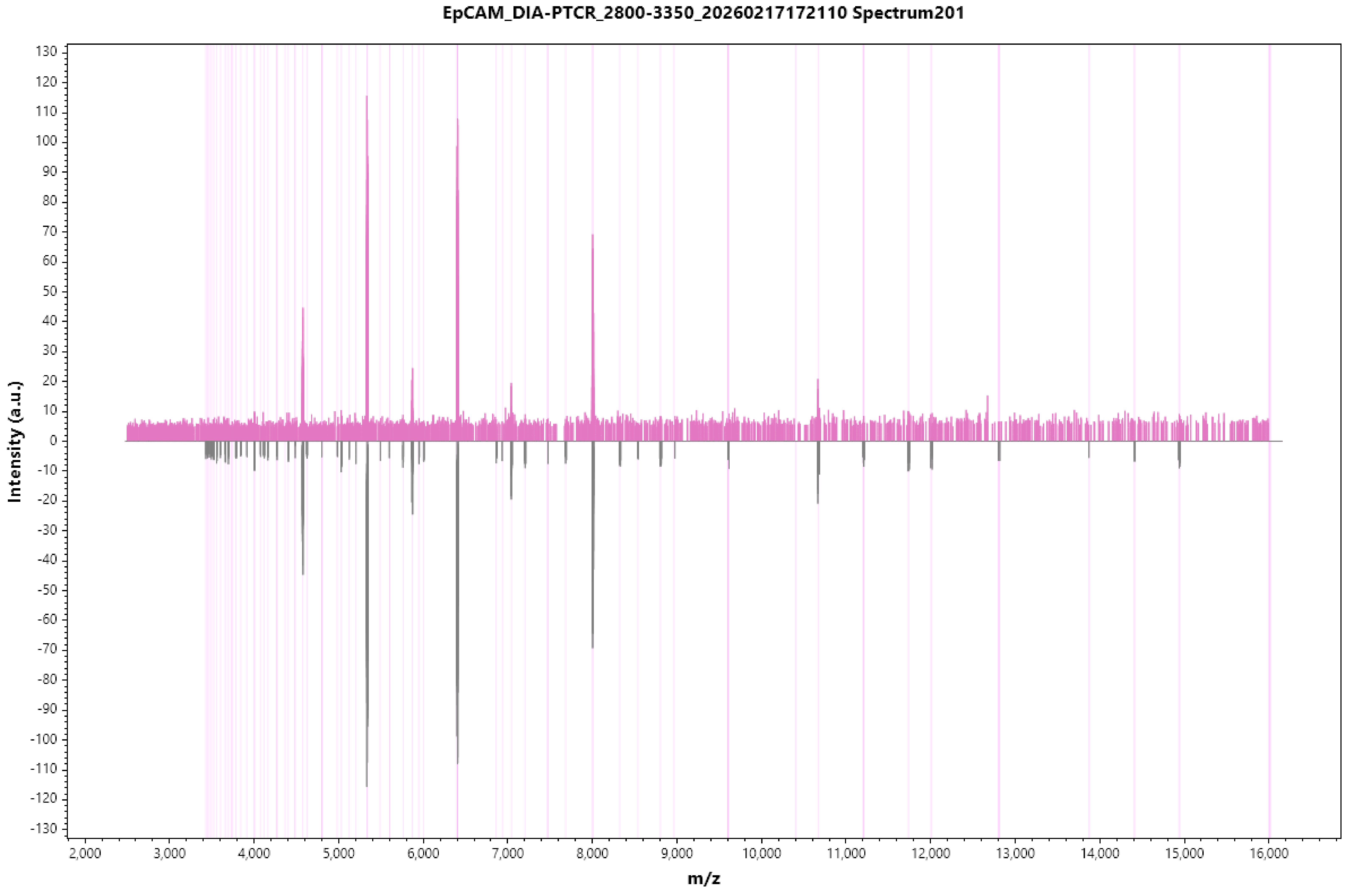
EpCAM_DIA-PTCR_2800-3350_20260217172110 Spectrum201
Intensity (a.u.)
m/z
130
120
110
100
90
80
70
60
50
40
30
20
10
0
-10
-20
-30
-40
-50
-60
-70
-80
-90
-100
-110
-120
-130
2,000
3,000
4,000
5,000
6,000
7,000
8,000
9,000
10,000
11,000
12,000
13,000
14,000
15,000
16,000

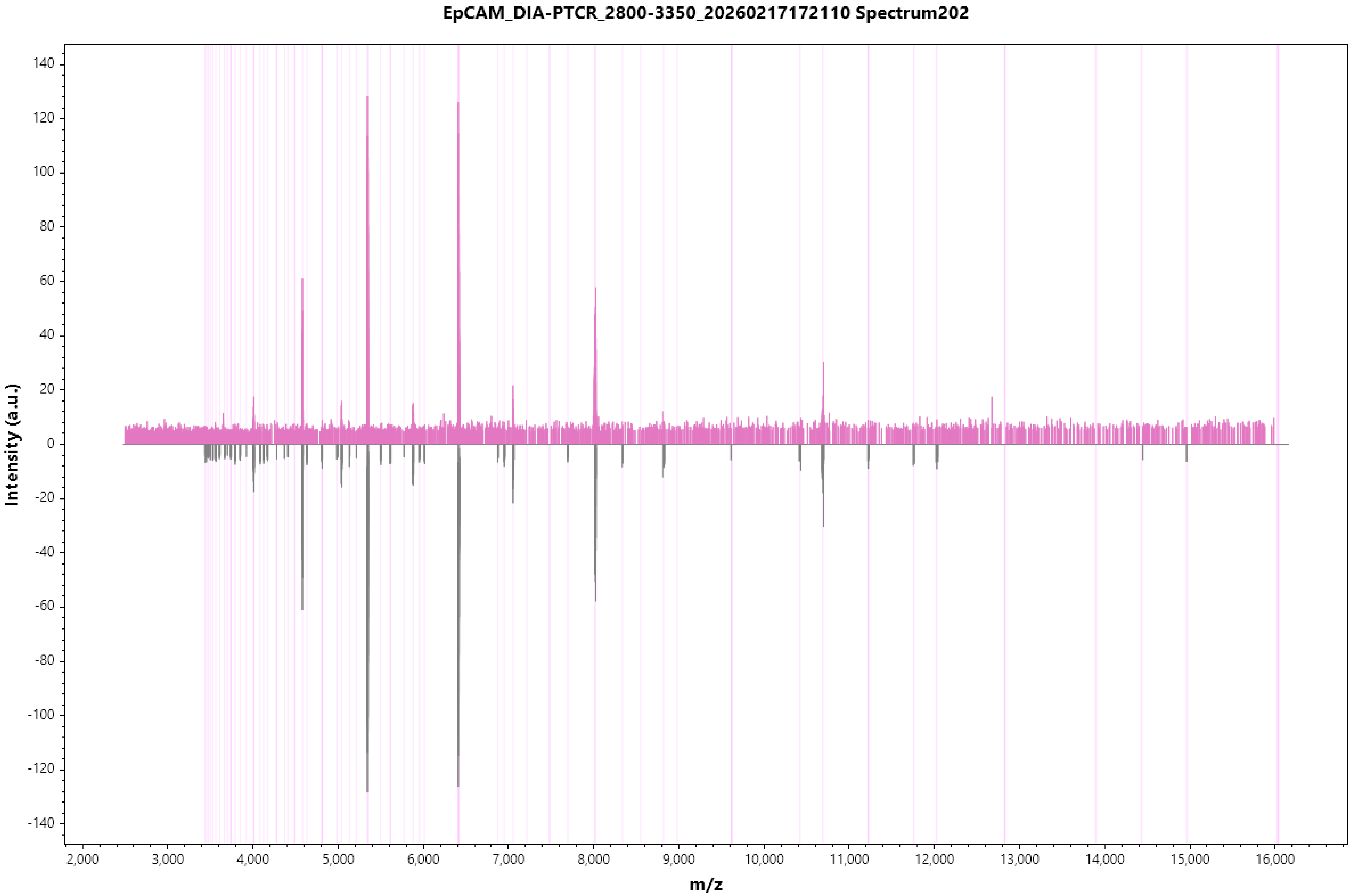
EpCAM_DIA-PTCR_2800-3350_20260217172110 Spectrum202
Intensity (a.u.)
m/z
140
120
100
80
60
40
20
0
-20
-40
-60
-80
-100
-120
-140
2,000
3,000
4,000
5,000
6,000
7,000
8,000
9,000
10,000
11,000
12,000
13,000
14,000
15,000
16,000

EpCAM_DIA-PTCR_2800-3350_20260217172110 Spectrum203

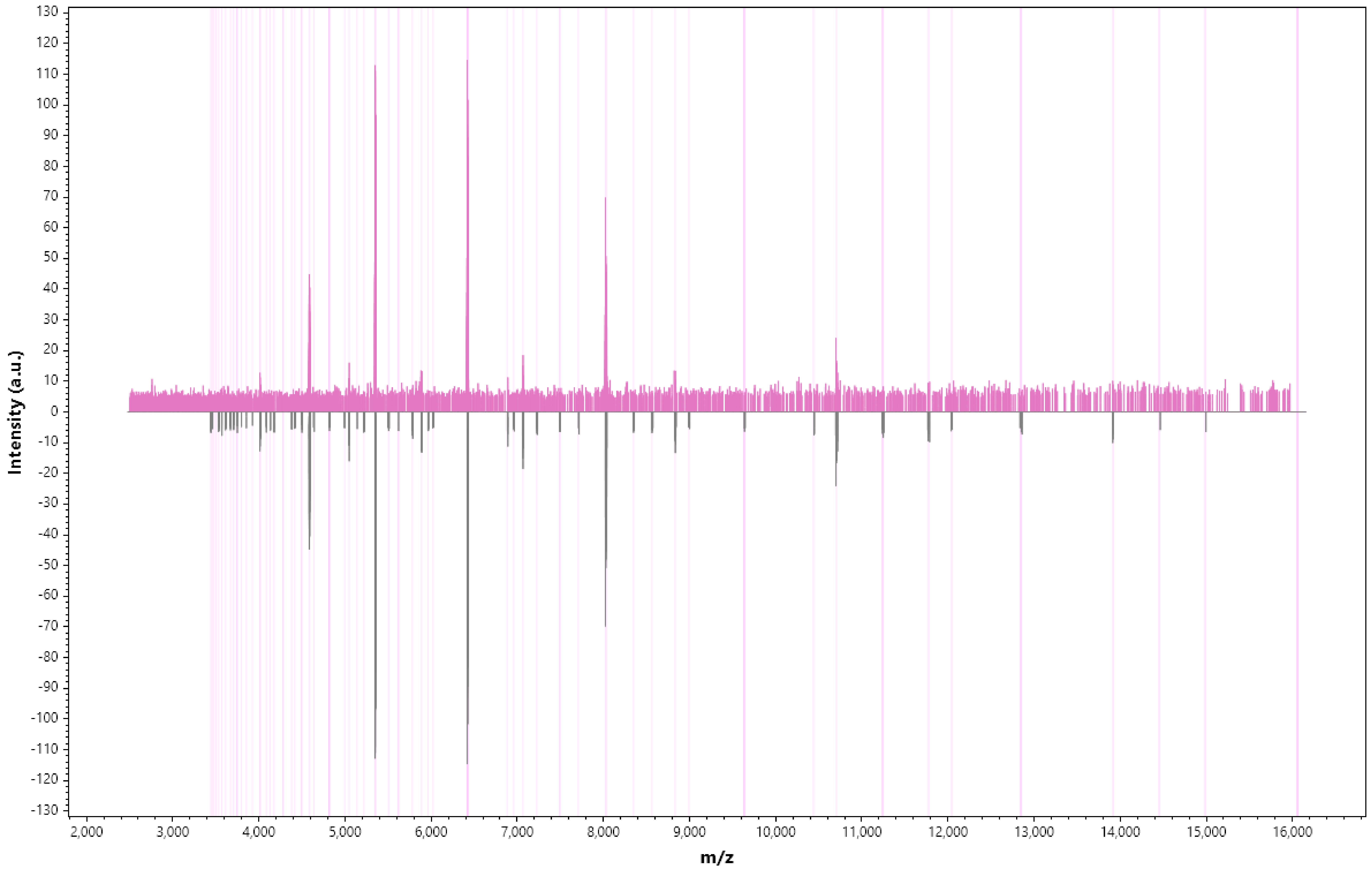

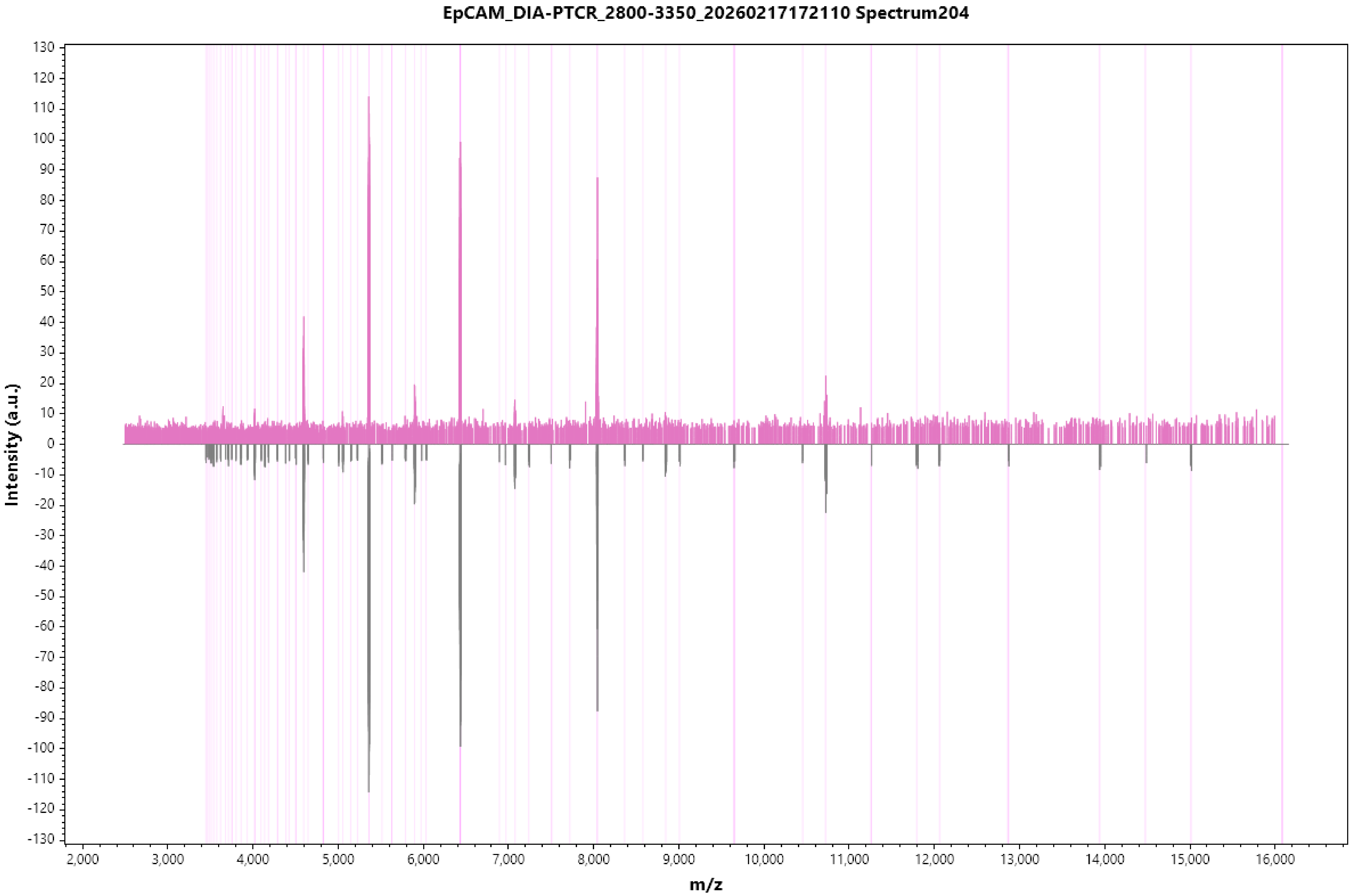
EpCAM_DIA-PTCR_2800-3350_20260217172110 Spectrum204
Intensity (a.u.)
m/z
130
120
110
100
90
80
70
60
50
40
30
20
10
0
-10
-20
-30
-40
-50
-60
-70
-80
-90
-100
-110
-120
-130
2,000
3,000
4,000
5,000
6,000
7,000
8,000
9,000
10,000
11,000
12,000
13,000
14,000
15,000
16,000

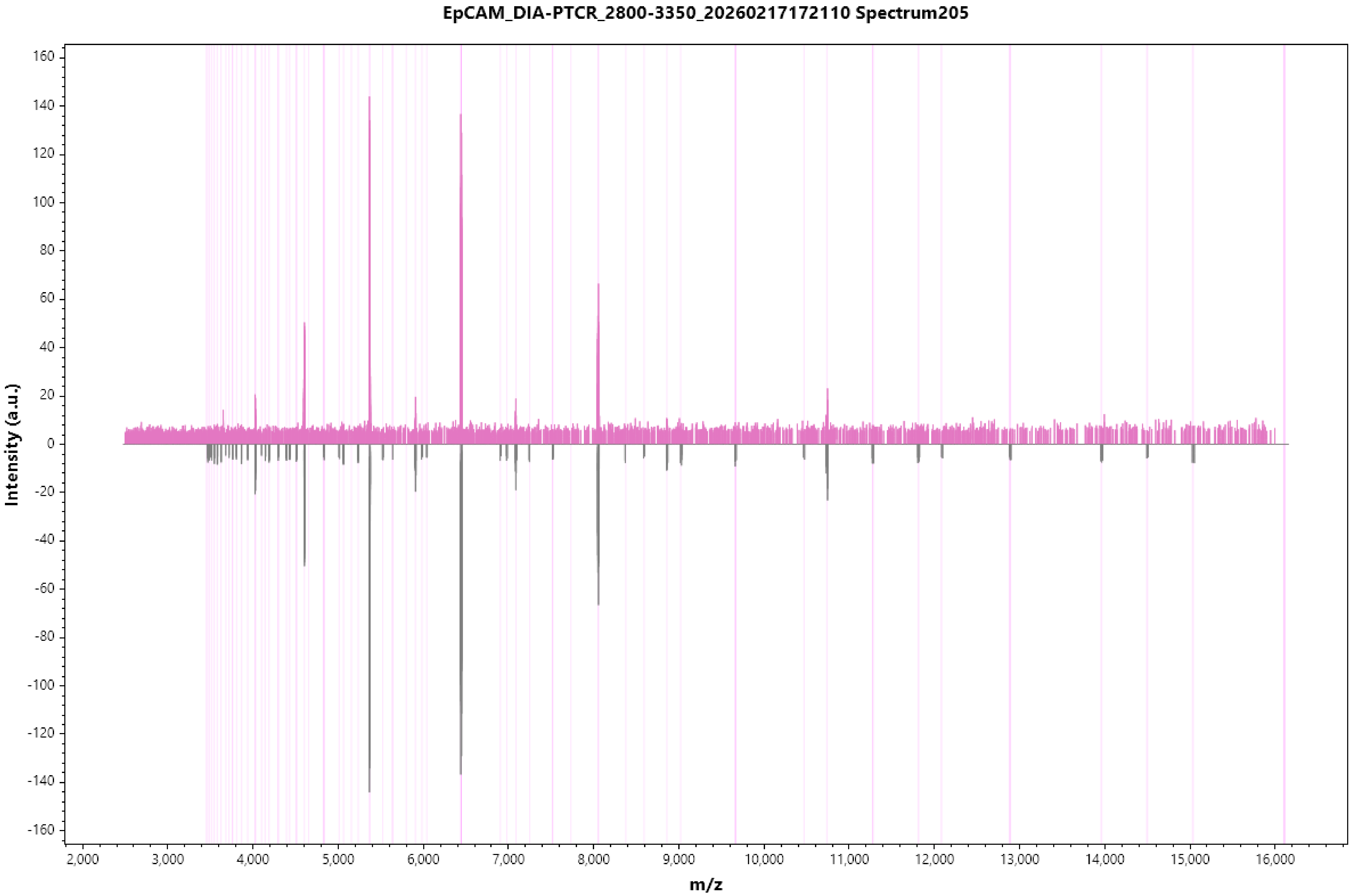
EpCAM_DIA-PTCR_2800-3350_20260217172110 Spectrum205
Intensity (a.u.)
m/z
160
140
120
100
80
60
40
20
0
-20
-40
-60
-80
-100
-120
-140
-160
2,000
3,000
4,000
5,000
6,000
7,000
8,000
9,000
10,000
11,000
12,000
13,000
14,000
15,000
16,000

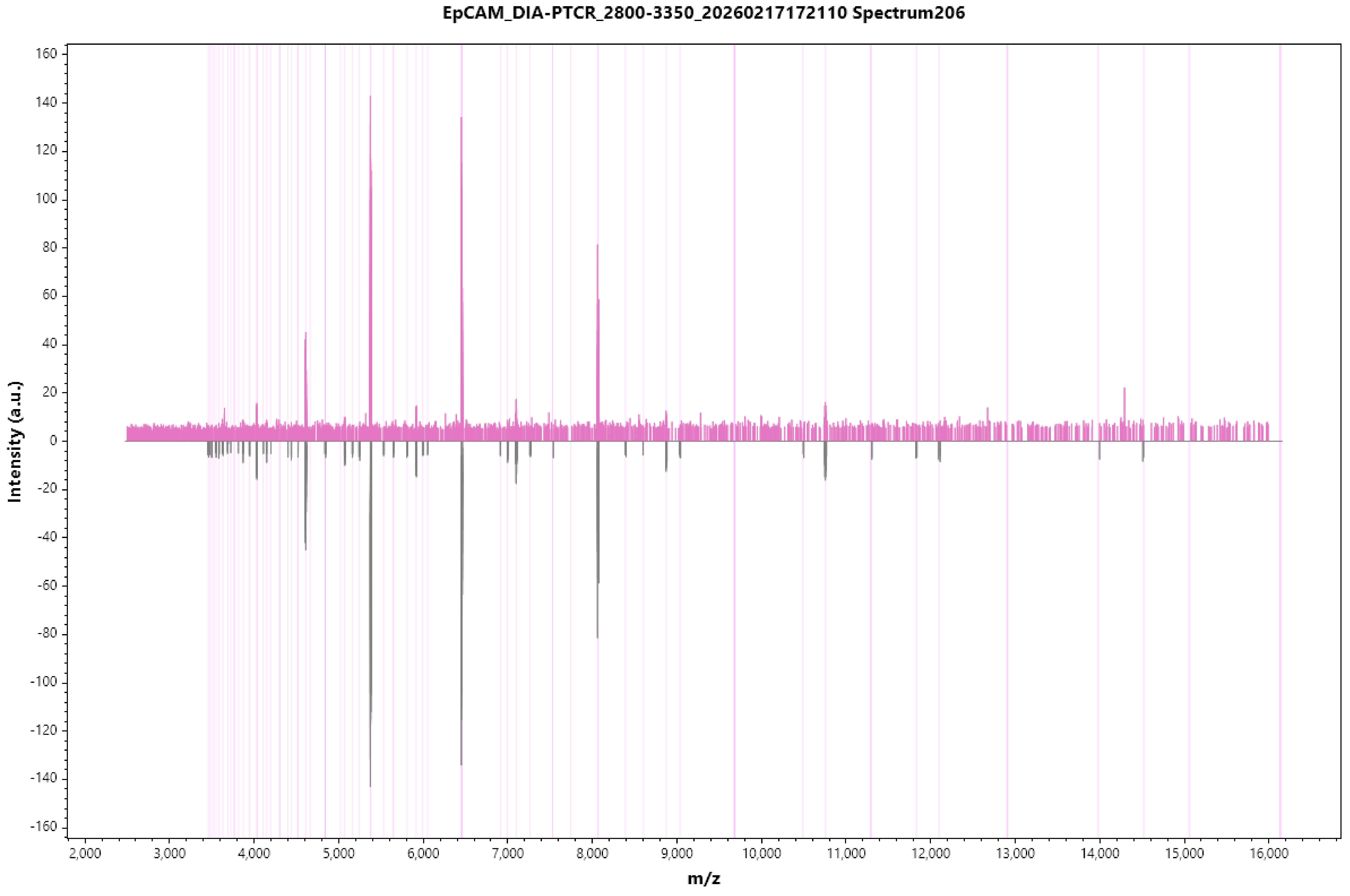

EpCAM_DIA-PTCR_2800-3350_20260217172110 Spectrum206
Intensity (a.u.)
m/z
160
140
120
100
80
60
40
20
0
-20
-40
-60
-80
-100
-120
-140
-160
2,000
3,000
4,000
5,000
6,000
7,000
8,000
9,000
10,000
11,000
12,000
13,000
14,000
15,000
16,000

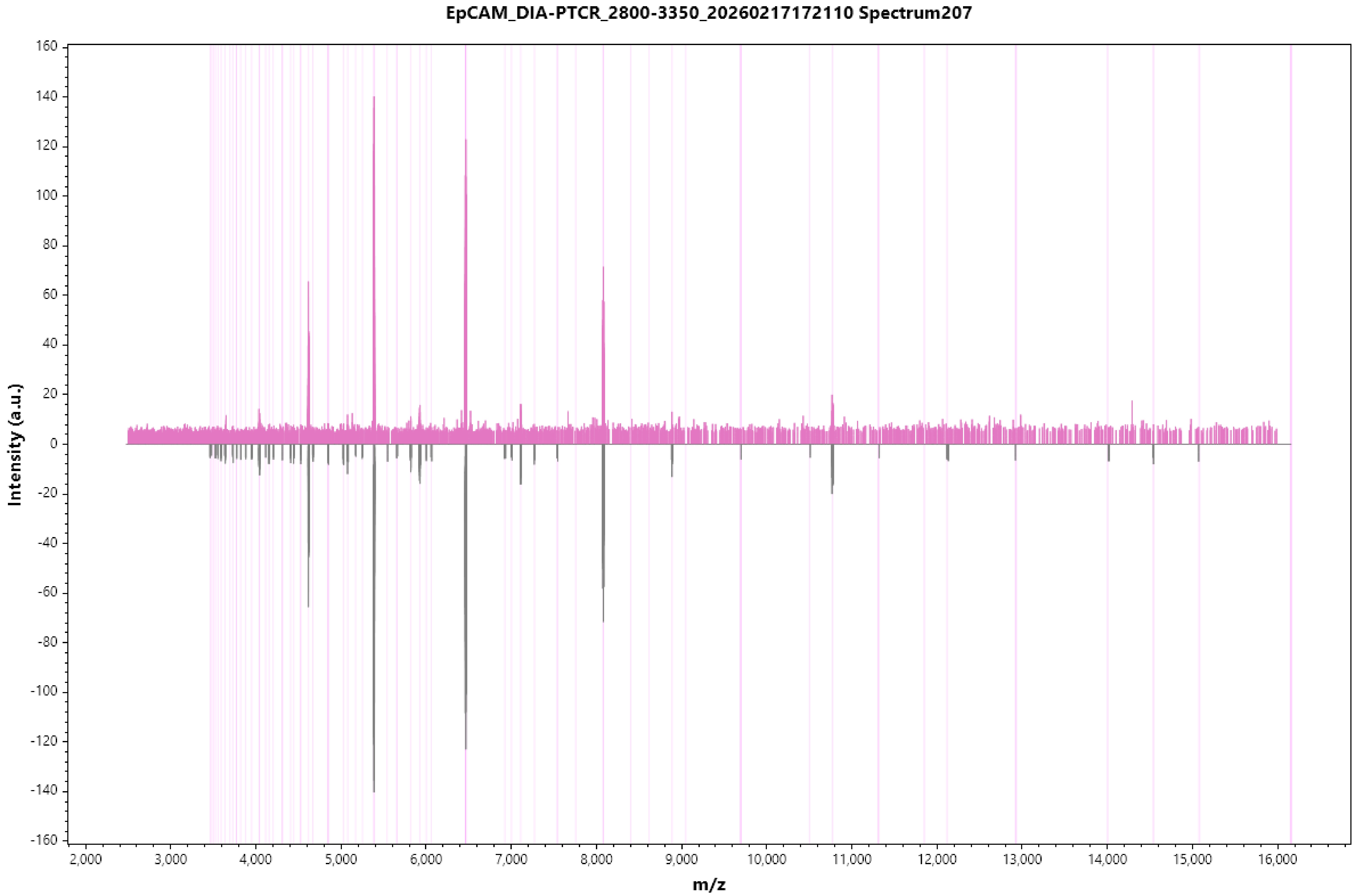
EpCAM_DIA-PTCR_2800-3350_20260217172110 Spectrum207
Intensity (a.u.)
m/z
160
140
120
100
80
60
40
20
0
-20
-40
-60
-80
-100
-120
-140
-160
2,000
3,000
4,000
5,000
6,000
7,000
8,000
9,000
10,000
11,000
12,000
13,000
14,000
15,000
16,000

EpCAM_DIA-PTCR_2800-3350_20260217172110 Spectrum208

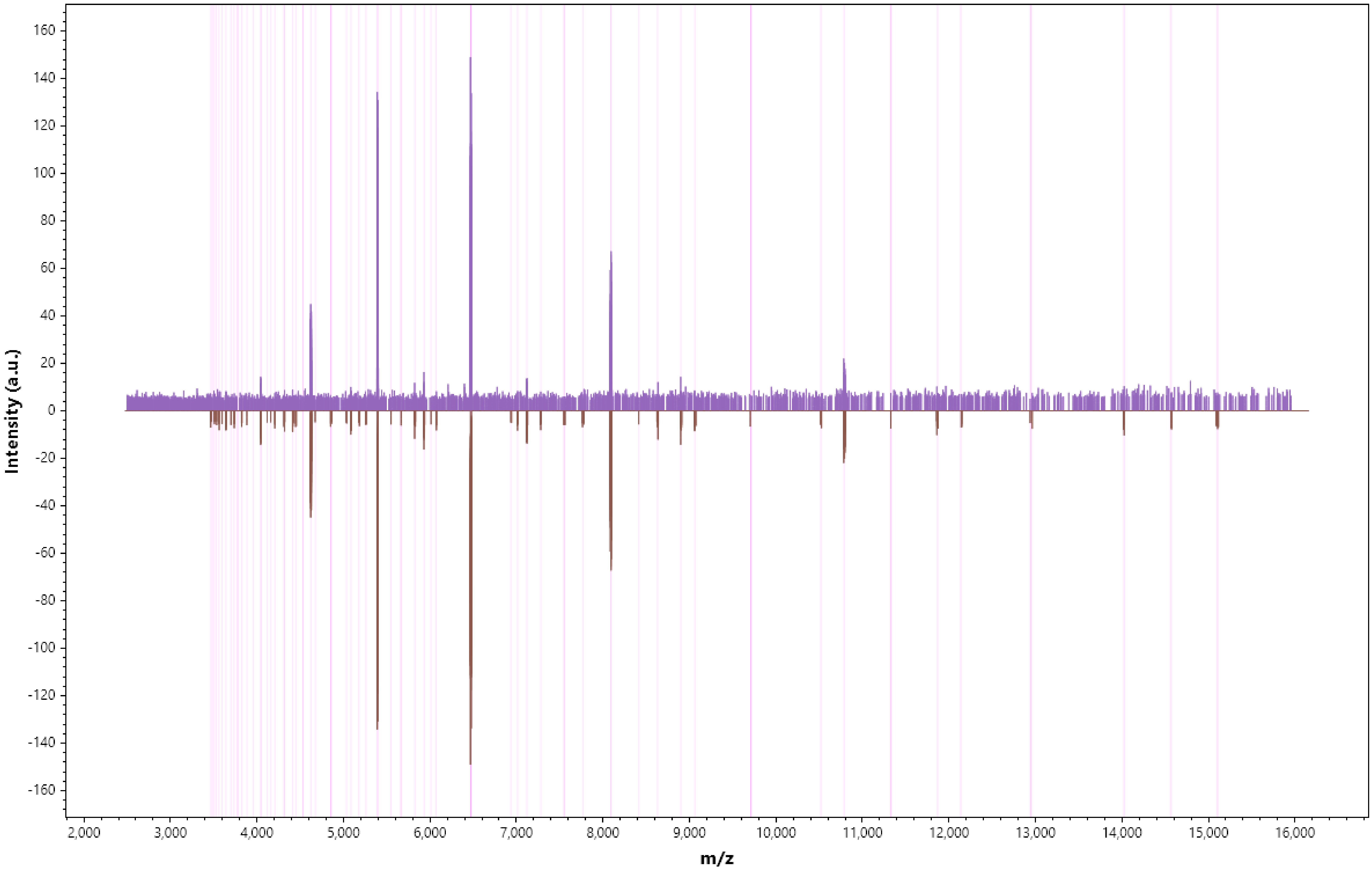

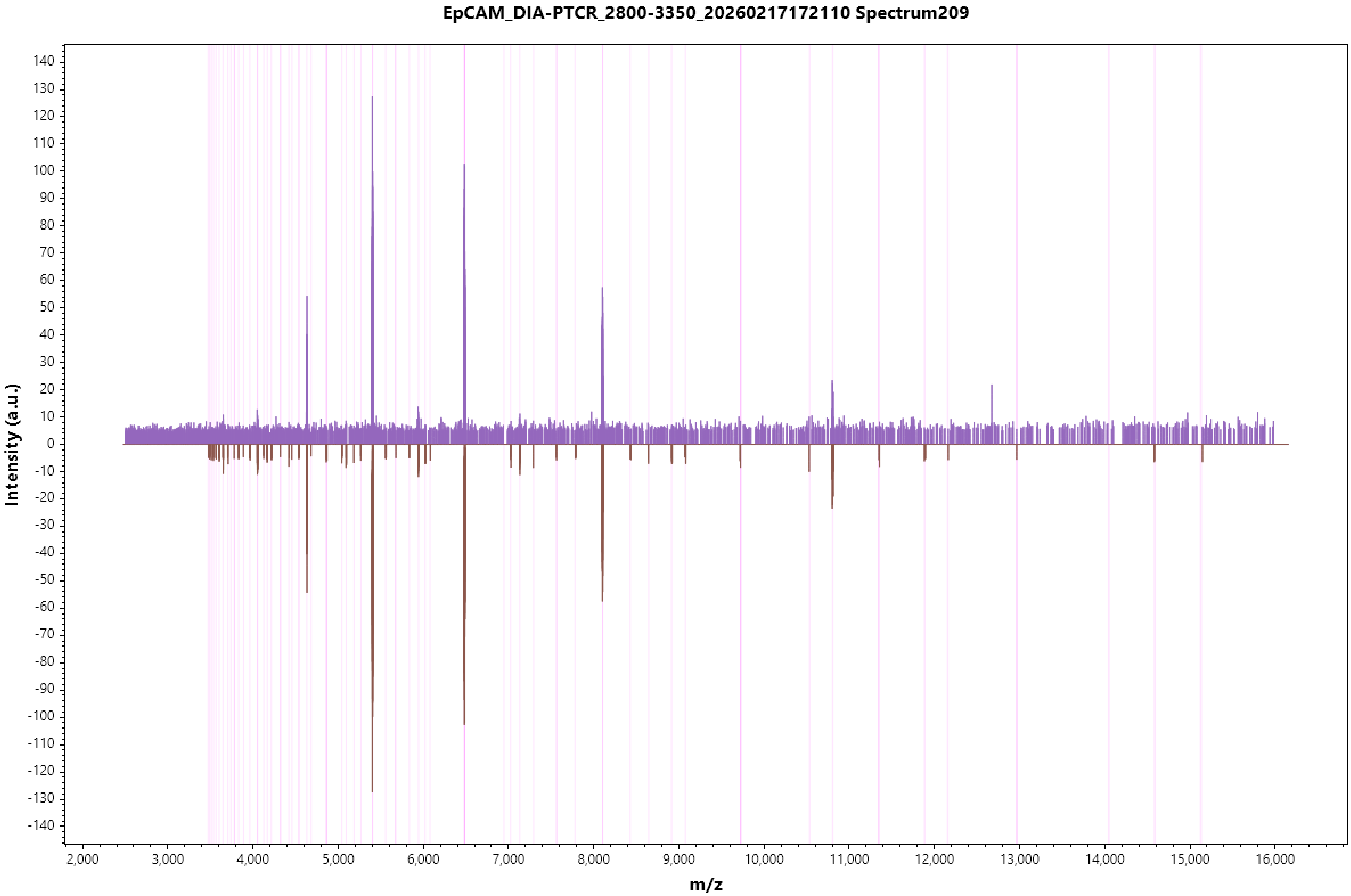
EpCAM_DIA-PTCR_2800-3350_20260217172110 Spectrum209
Intensity (a.u.)
m/z

EpCAM_DIA-PTCR_2800-3350_20260217172110 Spectrum210

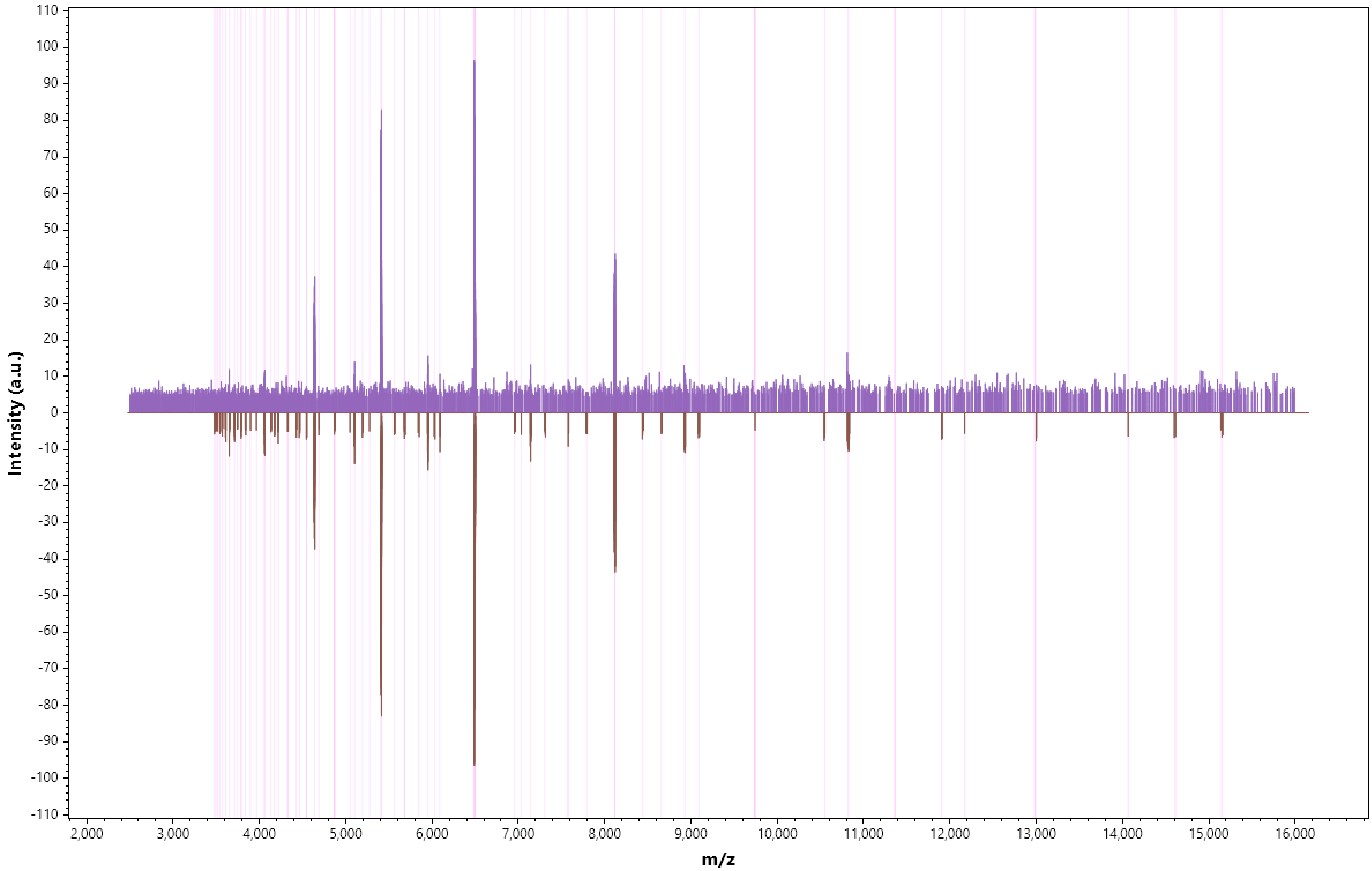

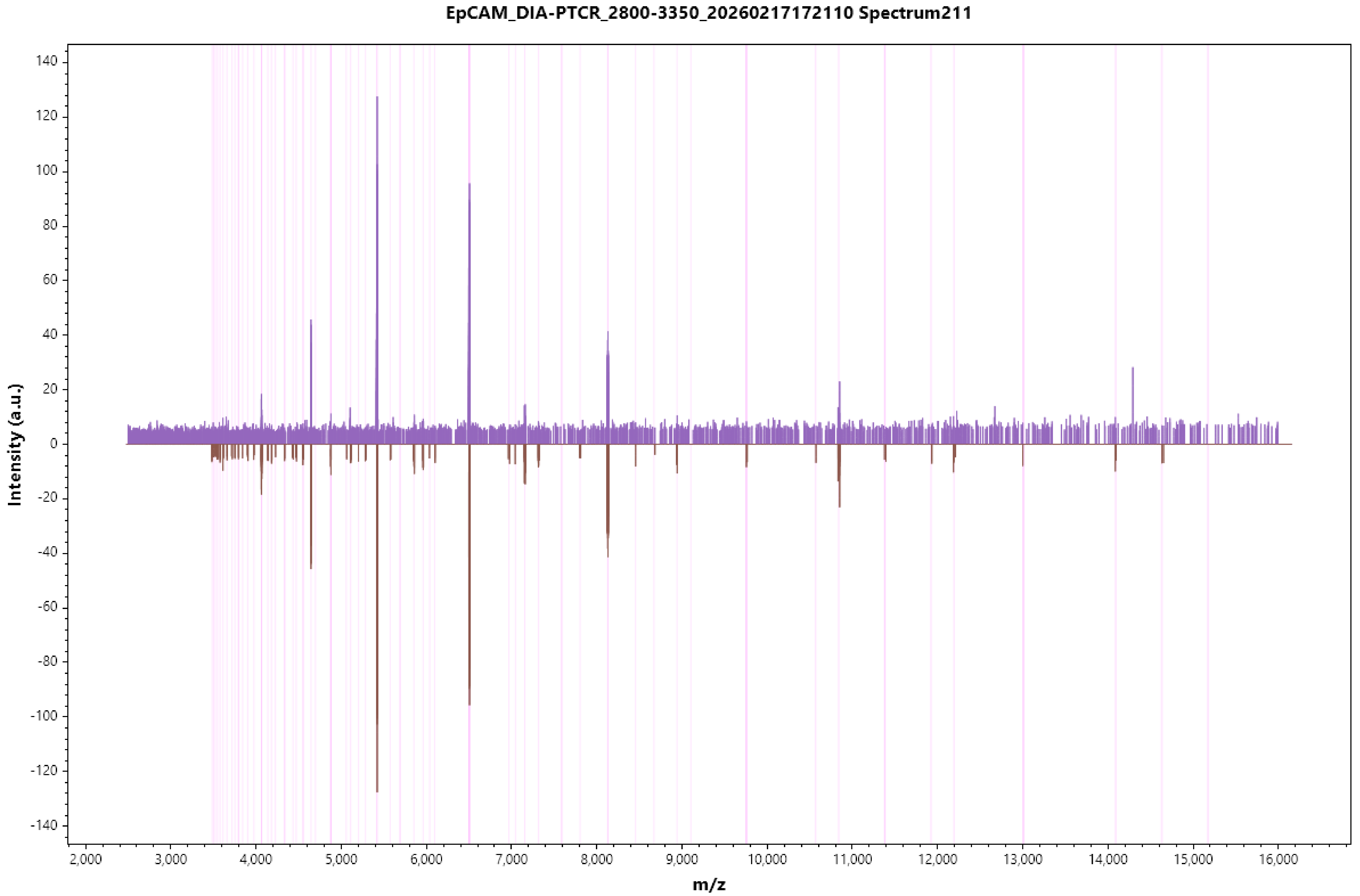
EpCAM_DIA-PTCR_2800-3350_20260217172110 Spectrum211
Intensity (a.u.)
m/z
140
120
100
80
60
40
20
0
-20
-40
-60
-80
-100
-120
-140
2,000
3,000
4,000
5,000
6,000
7,000
8,000
9,000
10,000
11,000
12,000
13,000
14,000
15,000
16,000

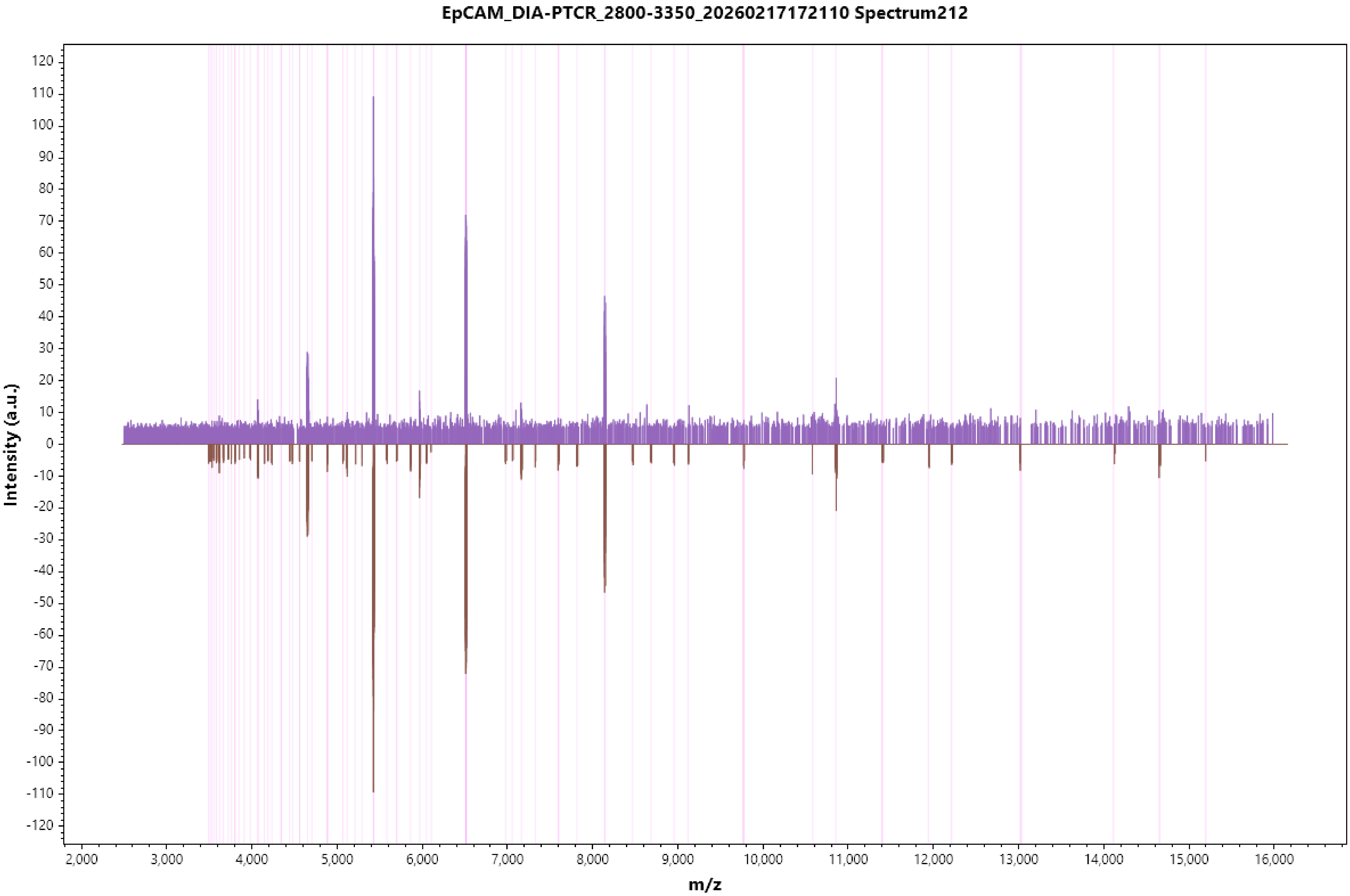

EpCAM_DIA-PTCR_2800-3350_20260217172110 Spectrum212
Intensity (a.u.)
m/z

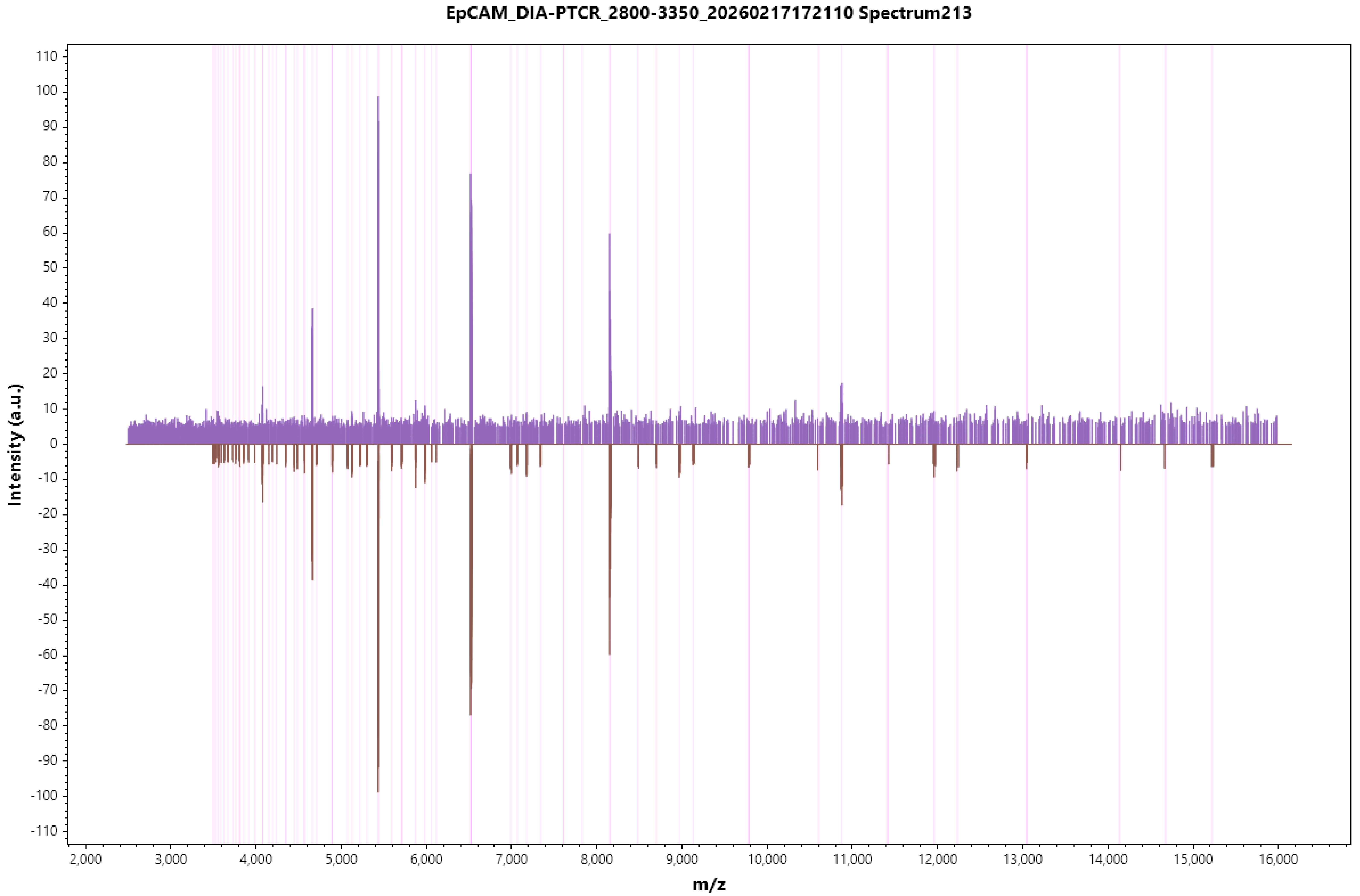
EpCAM_DIA-PTCR_2800-3350_20260217172110 Spectrum213
Intensity (a.u.)
110
100
90
80
70
60
50
40
30
20
10
0
-10
-20
-30
-40
-50
-60
-70
-80
-90
-100
-110
2,000
3,000
4,000
5,000
6,000
7,000
8,000
9,000
10,000
11,000
12,000
13,000
14,000
15,000
16,000
m/z

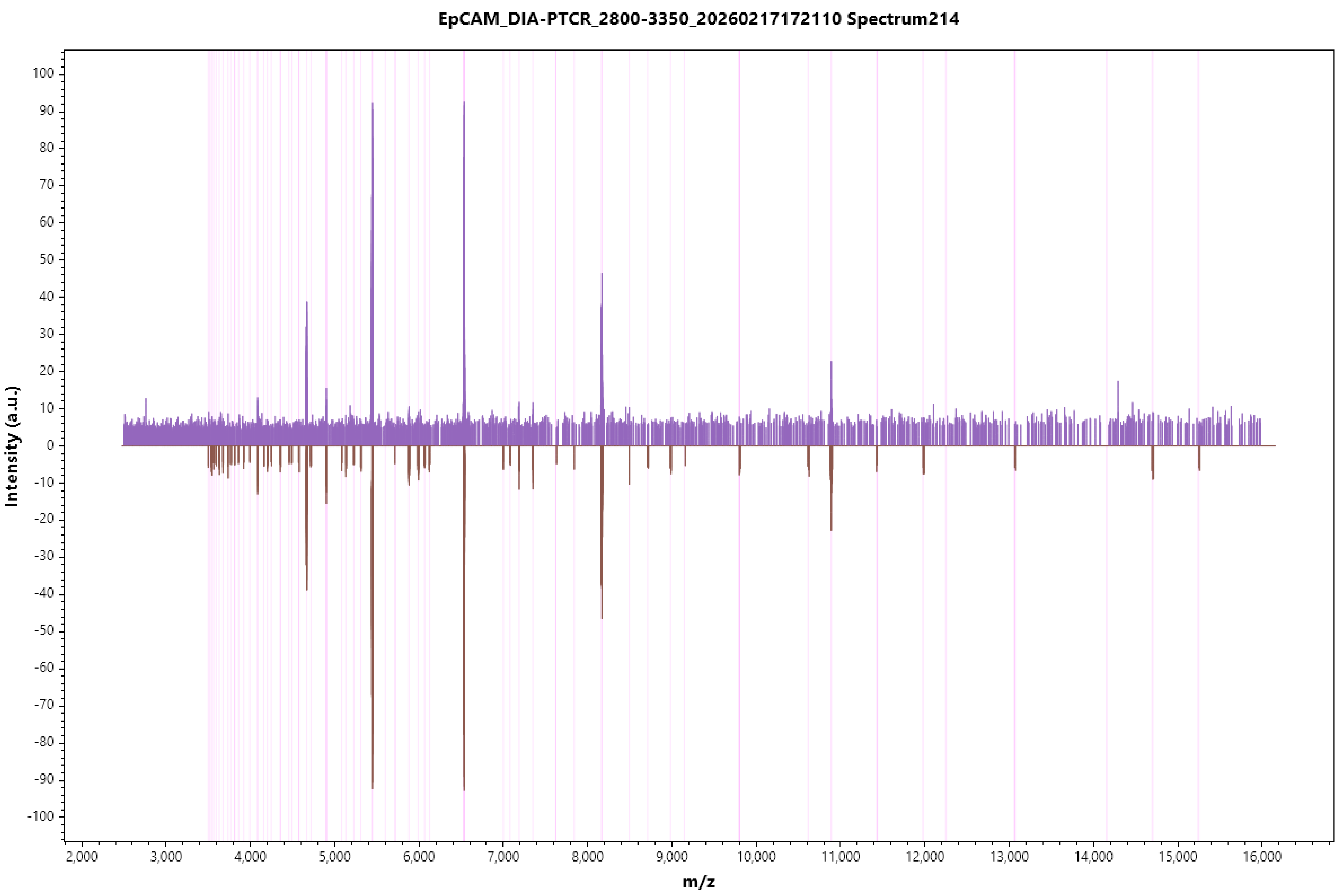
EpCAM_DIA-PTCR_2800-3350_20260217172110 Spectrum214
Intensity (a.u.)
m/z

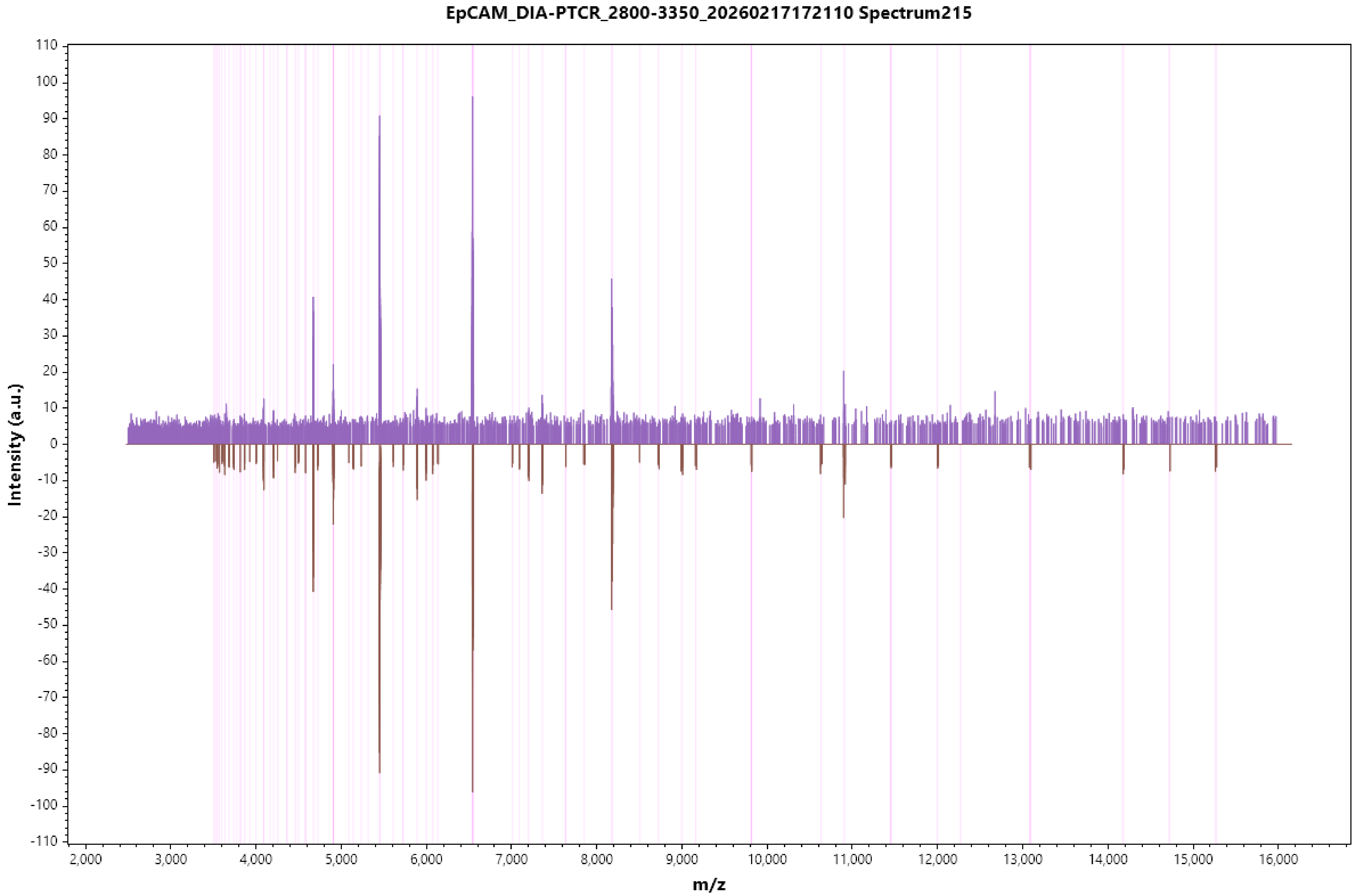
EpCAM_DIA-PTCR_2800-3350_20260217172110 Spectrum215
Intensity (a.u.)
110
100
90
80
70
60
50
40
30
20
10
0
-10
-20
-30
-40
-50
-60
-70
-80
-90
-100
-110
2,000
3,000
4,000
5,000
6,000
7,000
8,000
9,000
10,000
11,000
12,000
13,000
14,000
15,000
16,000
m/z

EpCAM_DIA-PTCR_2800-3350_20260217172110 Spectrum216

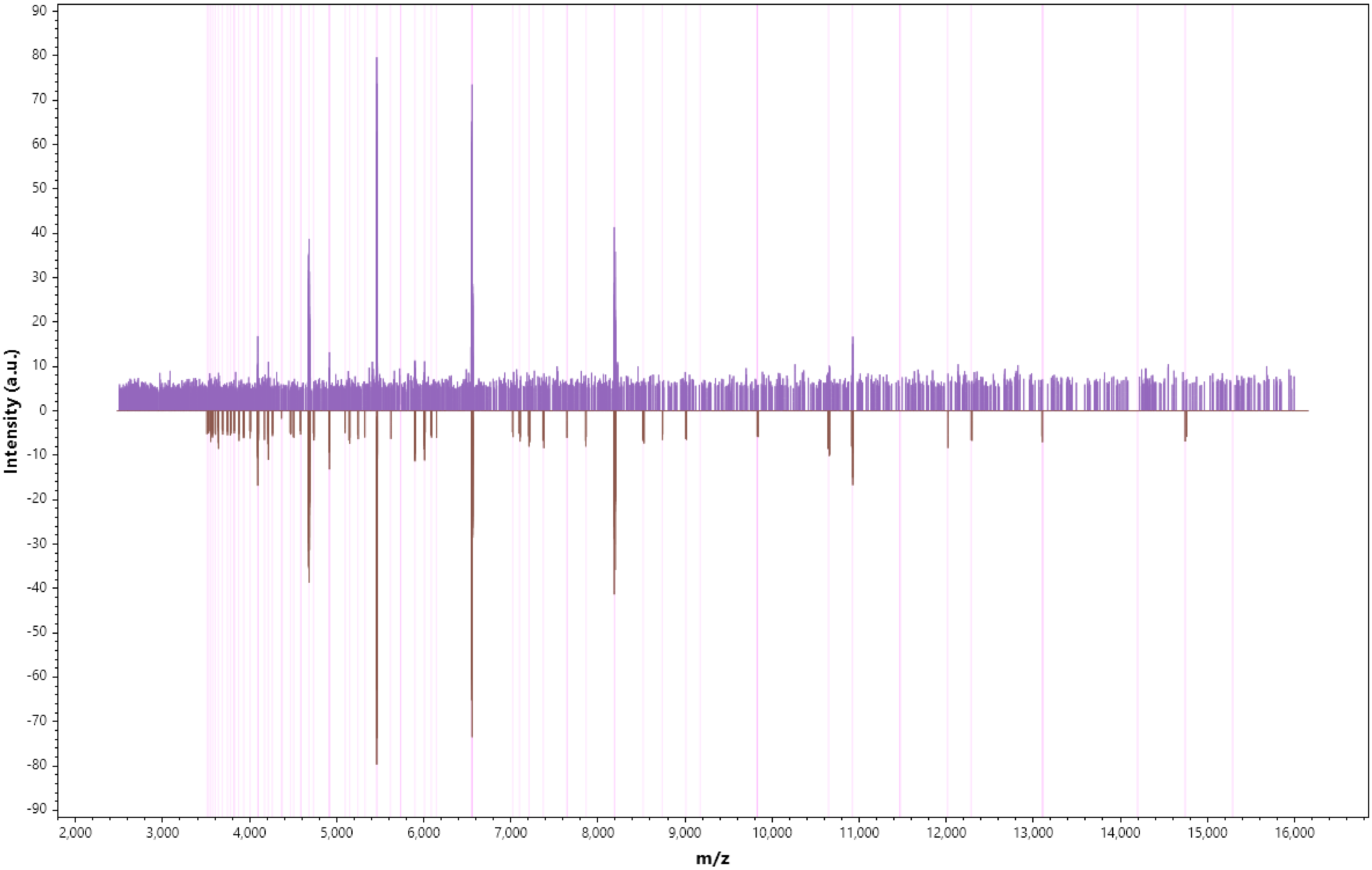

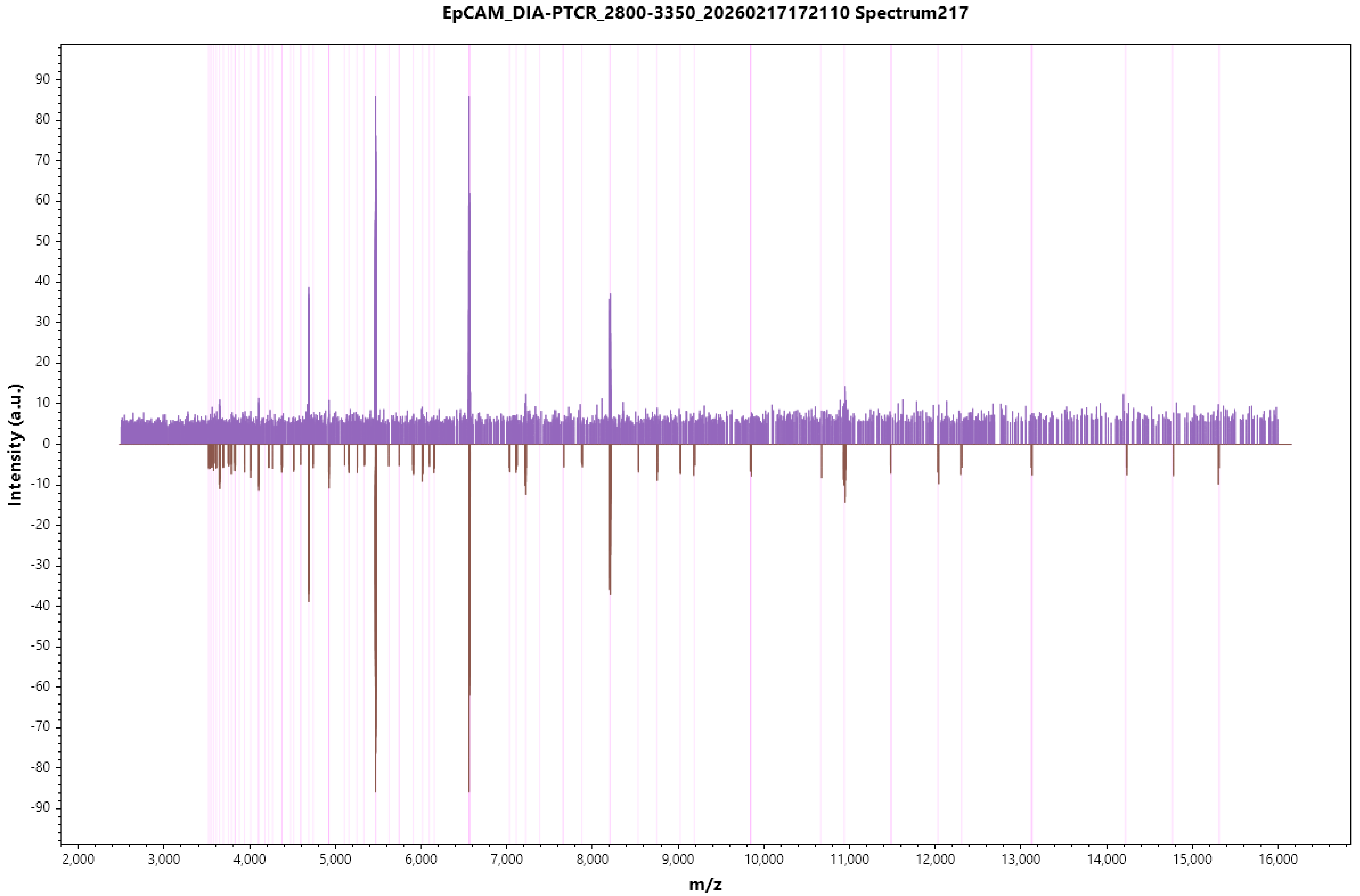

EpCAM_DIA-PTCR_2800-3350_20260217172110 Spectrum217
Intensity (a.u.)
m/z
90
80
70
60
50
40
30
20
10
0
-10
-20
-30
-40
-50
-60
-70
-80
-90
2,000
3,000
4,000
5,000
6,000
7,000
8,000
9,000
10,000
11,000
12,000
13,000
14,000
15,000
16,000

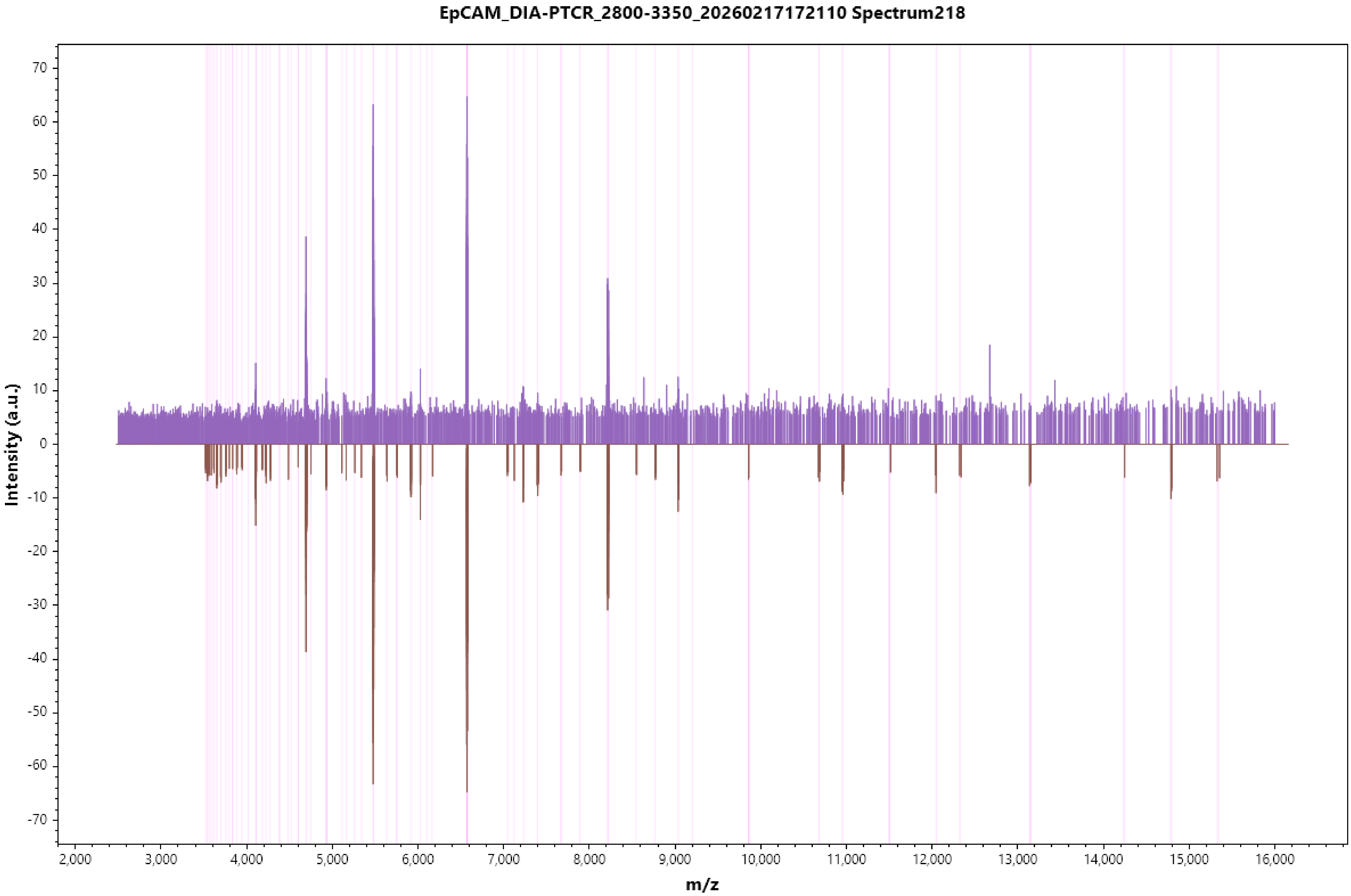
EpCAM_DIA-PTCR_2800-3350_20260217172110 Spectrum218
Intensity (a.u.)
70
60
50
40
30
20
10
0
-10
-20
-30
-40
-50
-60
-70
2,000
3,000
4,000
5,000
6,000
7,000
8,000
9,000
10,000
11,000
12,000
13,000
14,000
15,000
16,000
m/z

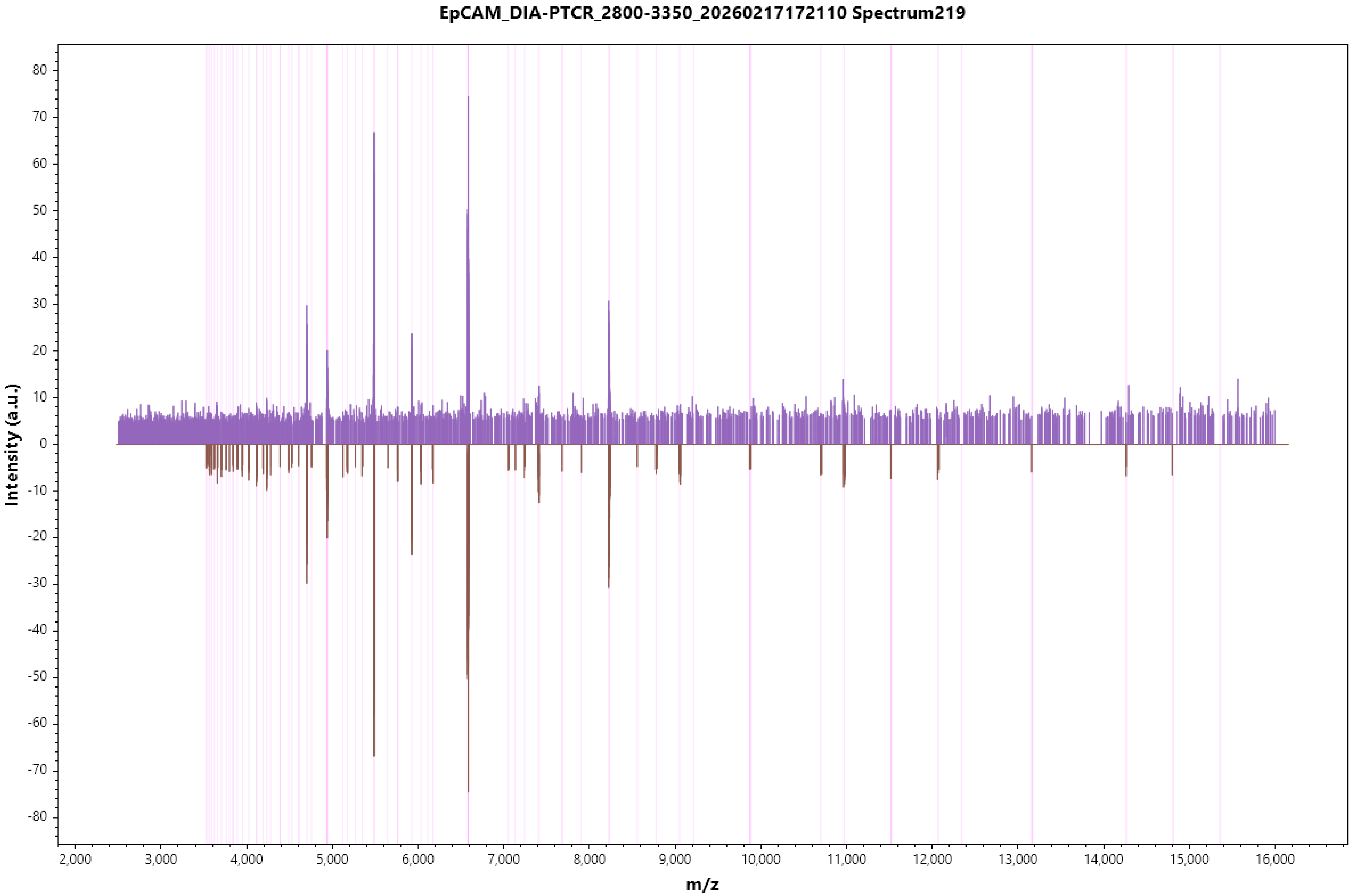

EpCAM_DIA-PTCR_2800-3350_20260217172110 Spectrum219
Intensity (a.u.)
m/z
80
70
60
50
40
30
20
10
0
-10
-20
-30
-40
-50
-60
-70
-80
2,000
3,000
4,000
5,000
6,000
7,000
8,000
9,000
10,000
11,000
12,000
13,000
14,000
15,000
16,000

EpCAM_DIA-PTCR_2800-3350_20260217172110 Spectrum220

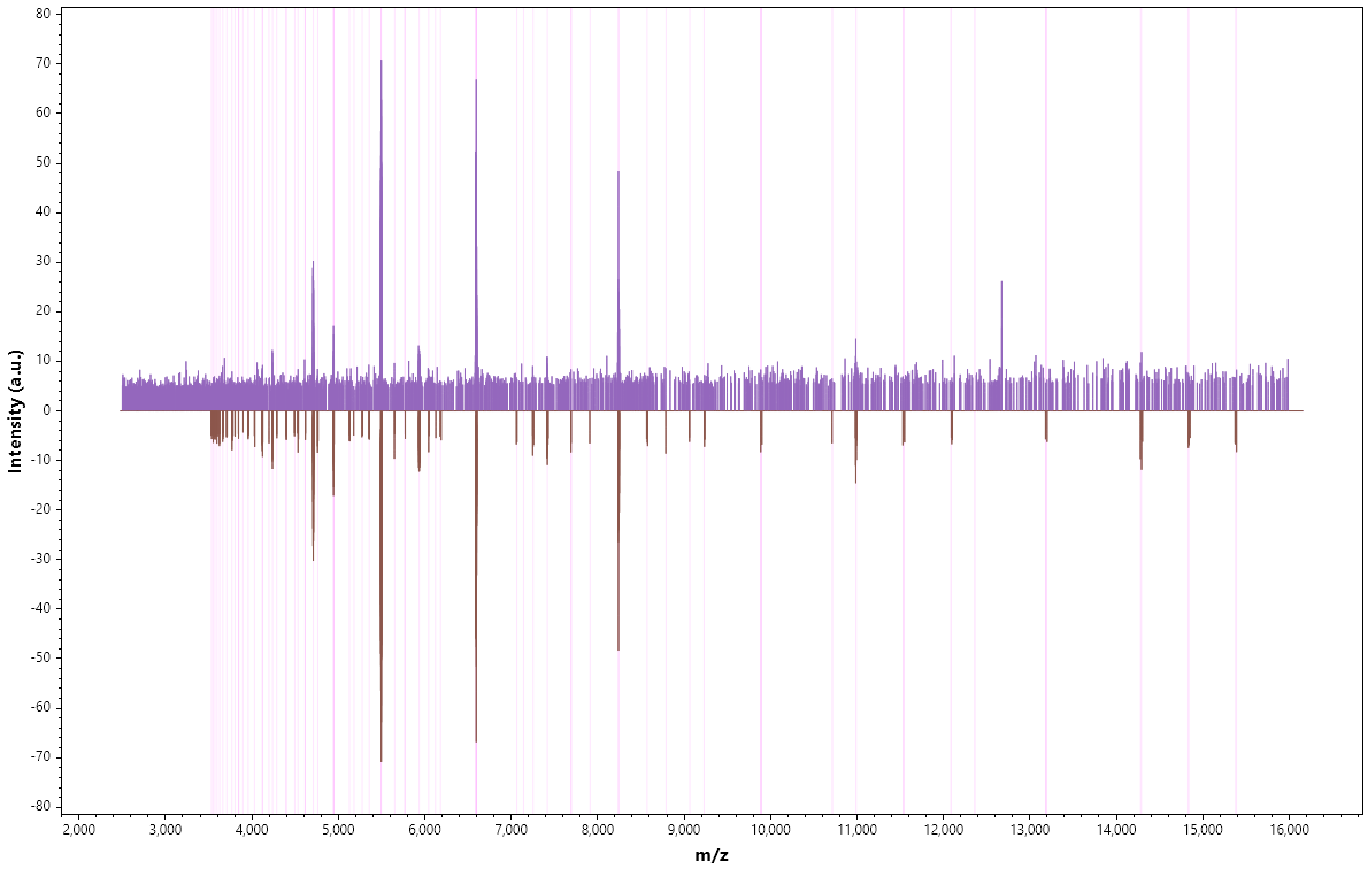

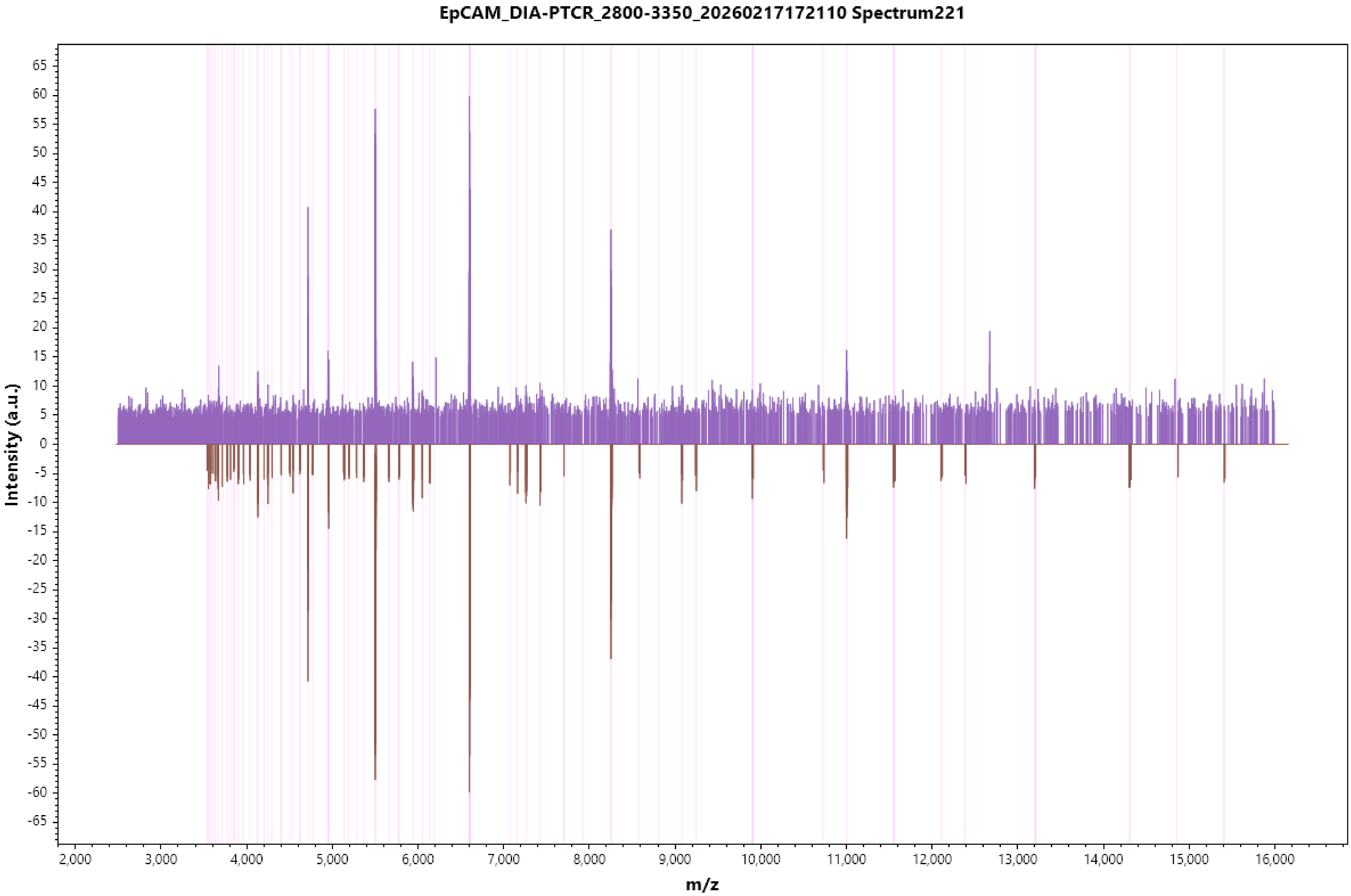

EpCAM_DIA-PTCR_2800-3350_20260217172110 Spectrum221
Intensity (a.u.)
m/z
65
60
55
50
45
40
35
30
25
20
15
10
5
0
-5
-10
-15
-20
-25
-30
-35
-40
-45
-50
-55
-60
-65
2,000
3,000
4,000
5,000
6,000
7,000
8,000
9,000
10,000
11,000
12,000
13,000
14,000
15,000
16,000

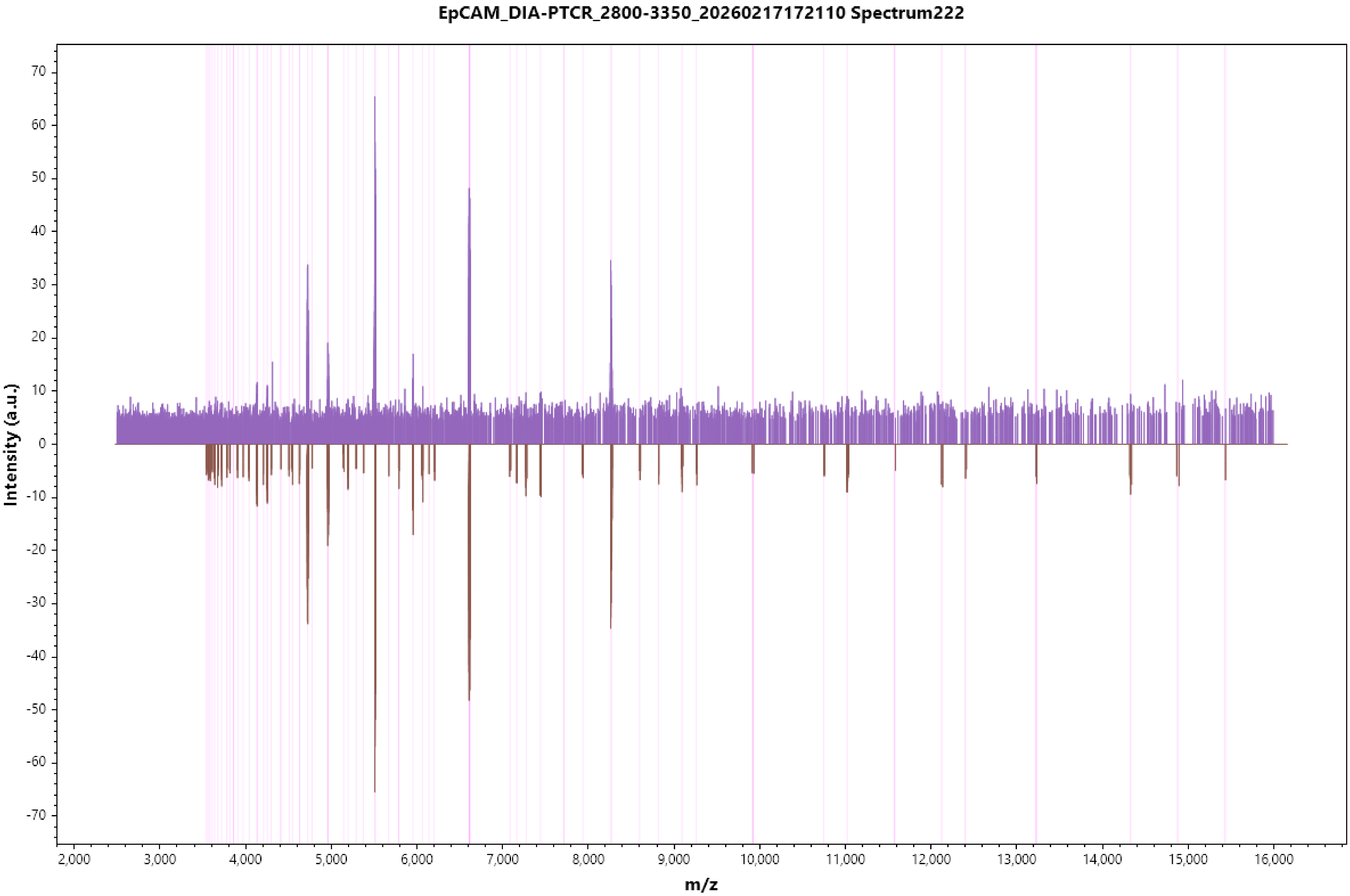
EpCAM_DIA-PTCR_2800-3350_20260217172110 Spectrum222
Intensity (a.u.)
m/z
70
60
50
40
30
20
10
0
-10
-20
-30
-40
-50
-60
-70
2,000
3,000
4,000
5,000
6,000
7,000
8,000
9,000
10,000
11,000
12,000
13,000
14,000
15,000
16,000

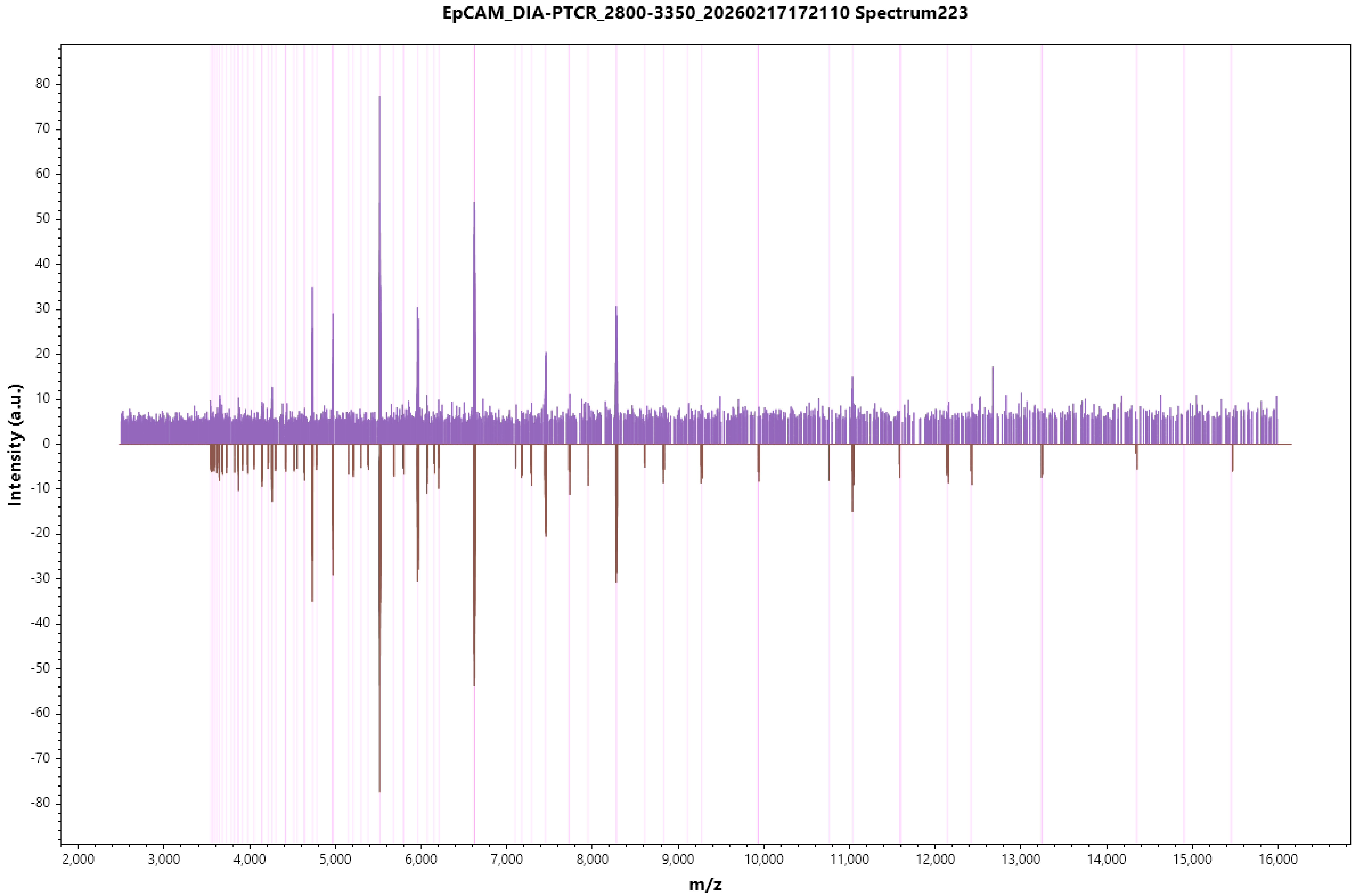
EpCAM_DIA-PTCR_2800-3350_20260217172110 Spectrum223
Intensity (a.u.)
m/z
80
70
60
50
40
30
20
10
0
-10
-20
-30
-40
-50
-60
-70
-80
2,000
3,000
4,000
5,000
6,000
7,000
8,000
9,000
10,000
11,000
12,000
13,000
14,000
15,000
16,000

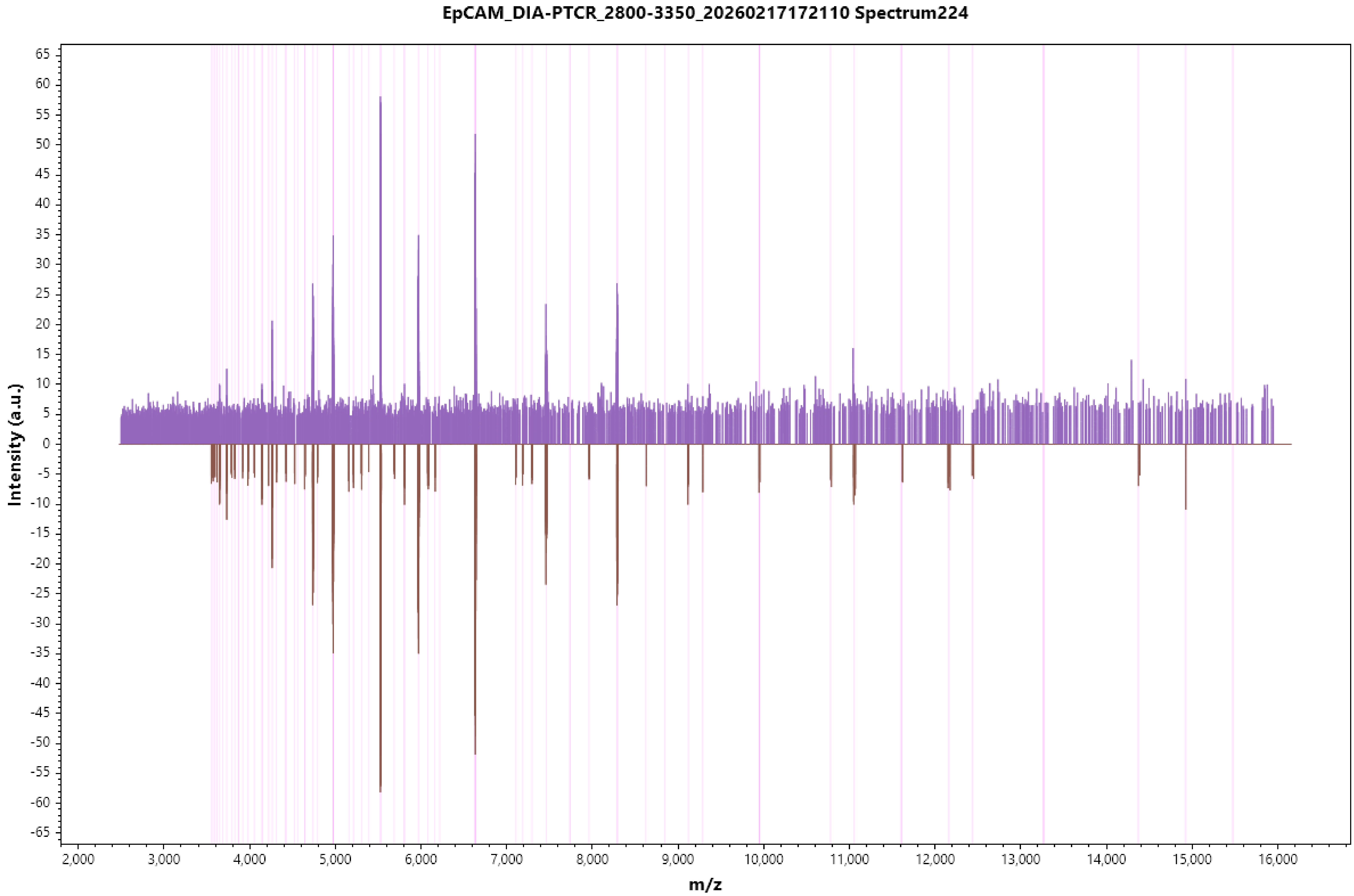

EpCAM_DIA-PTCR_2800-3350_20260217172110 Spectrum224
Intensity (a.u.)
m/z
65
60
55
50
45
40
35
30
25
20
15
10
5
0
-5
-10
-15
-20
-25
-30
-35
-40
-45
-50
-55
-60
-65
2,000
3,000
4,000
5,000
6,000
7,000
8,000
9,000
10,000
11,000
12,000
13,000
14,000
15,000
16,000

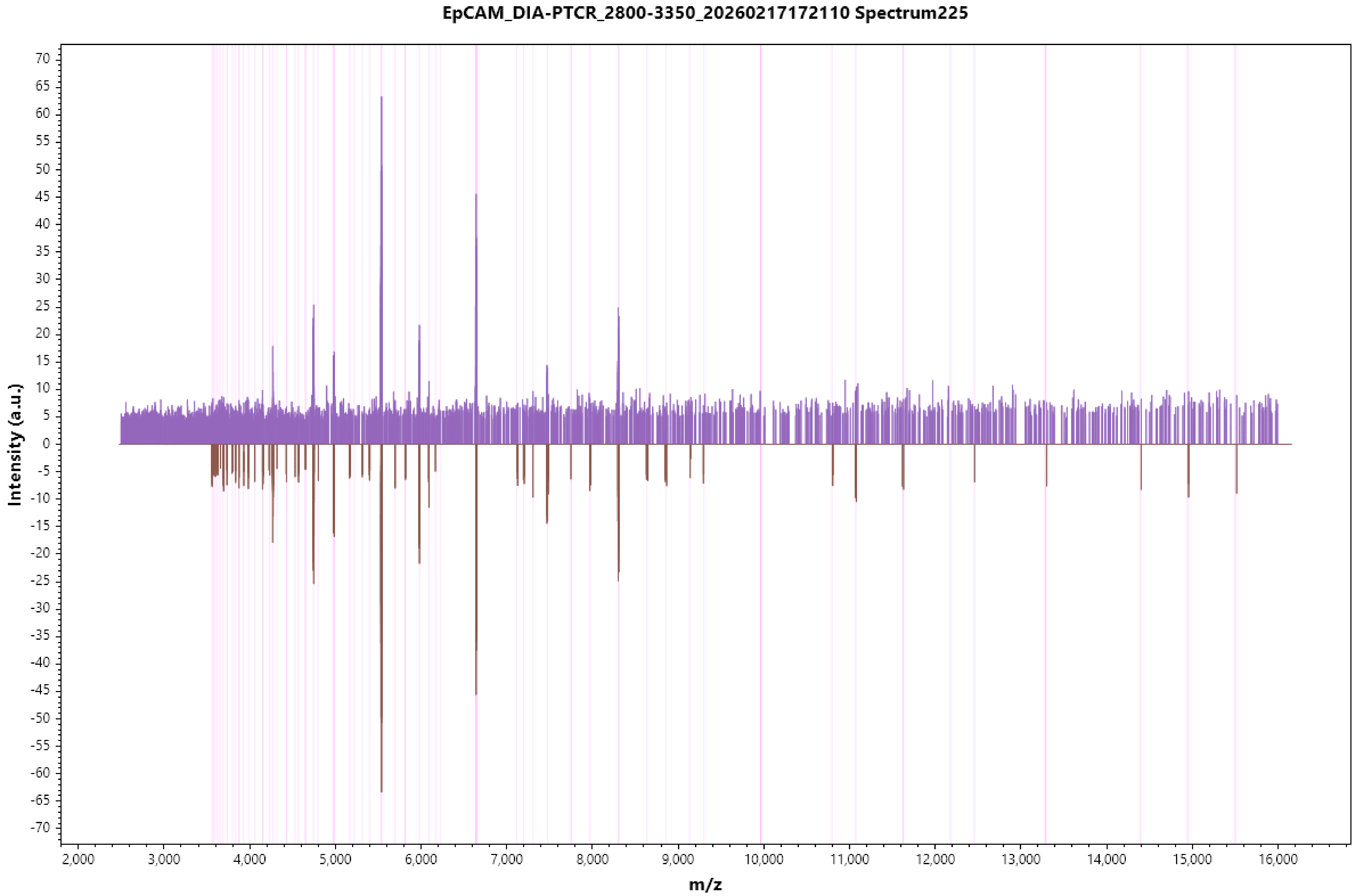
EpCAM_DIA-PTCR_2800-3350_20260217172110 Spectrum225
Intensity (a.u.)
m/z
70
65
60
55
50
45
40
35
30
25
20
15
10
5
0
-5
-10
-15
-20
-25
-30
-35
-40
-45
-50
-55
-60
-65
-70
2,000
3,000
4,000
5,000
6,000
7,000
8,000
9,000
10,000
11,000
12,000
13,000
14,000
15,000
16,000

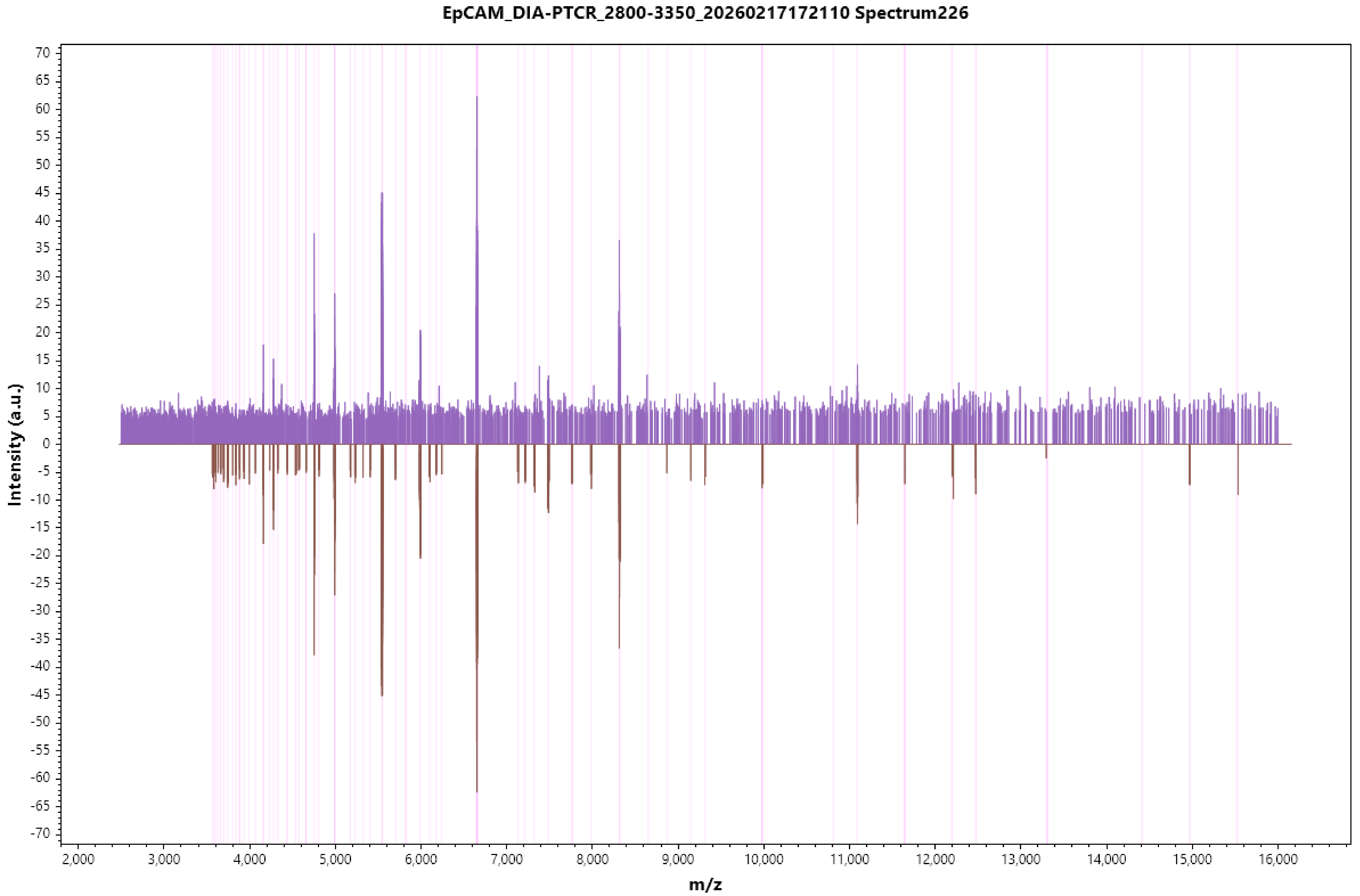

EpCAM_DIA-PTCR_2800-3350_20260217172110 Spectrum226
Intensity (a.u.)
m/z
70
65
60
55
50
45
40
35
30
25
20
15
10
5
0
-5
-10
-15
-20
-25
-30
-35
-40
-45
-50
-55
-60
-65
-70
2,000
3,000
4,000
5,000
6,000
7,000
8,000
9,000
10,000
11,000
12,000
13,000
14,000
15,000
16,000

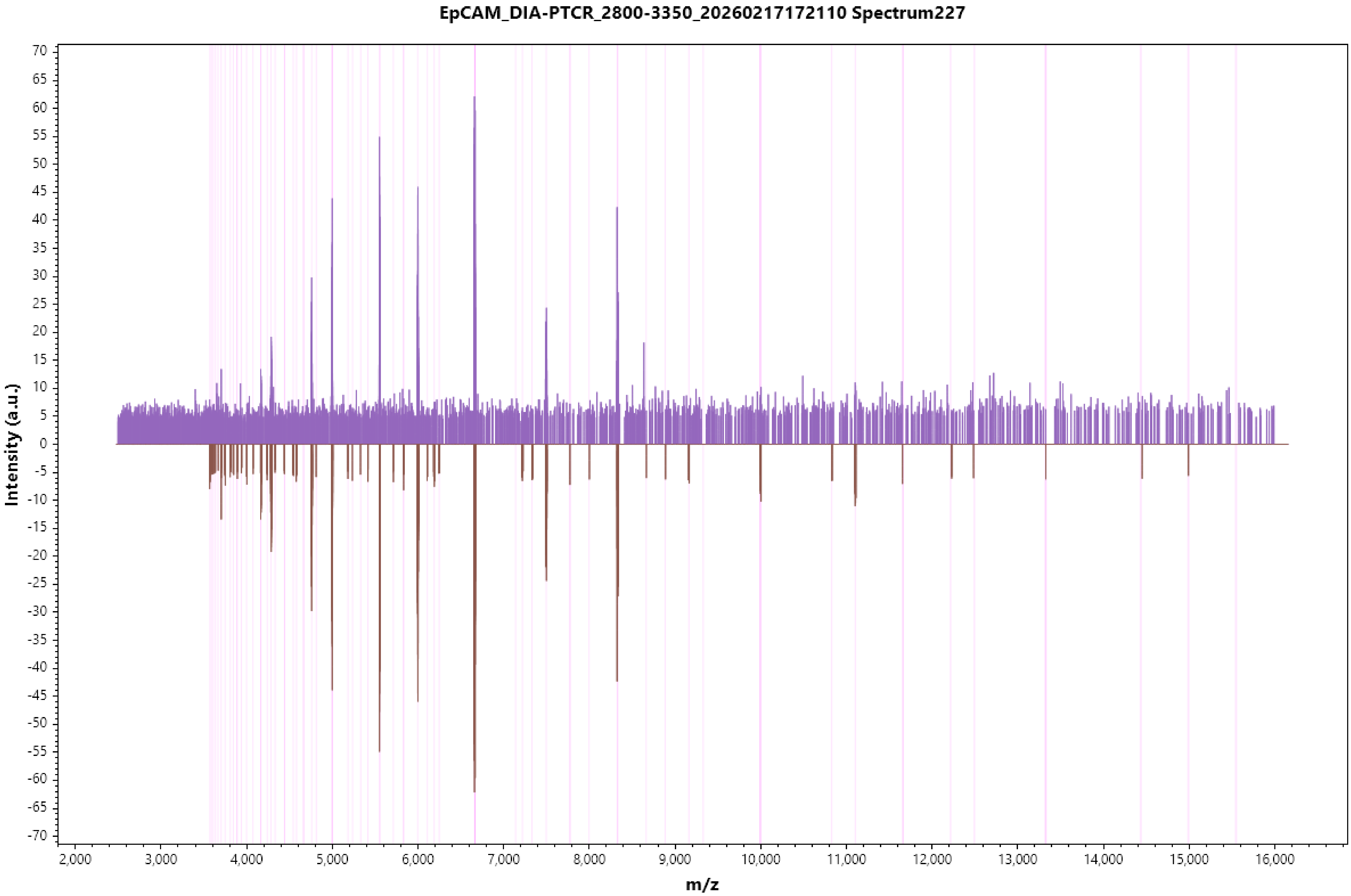
EpCAM_DIA-PTCR_2800-3350_20260217172110 Spectrum227
Intensity (a.u.)
m/z
70
65
60
55
50
45
40
35
30
25
20
15
10
5
0
-5
-10
-15
-20
-25
-30
-35
-40
-45
-50
-55
-60
-65
-70
2,000
3,000
4,000
5,000
6,000
7,000
8,000
9,000
10,000
11,000
12,000
13,000
14,000
15,000
16,000

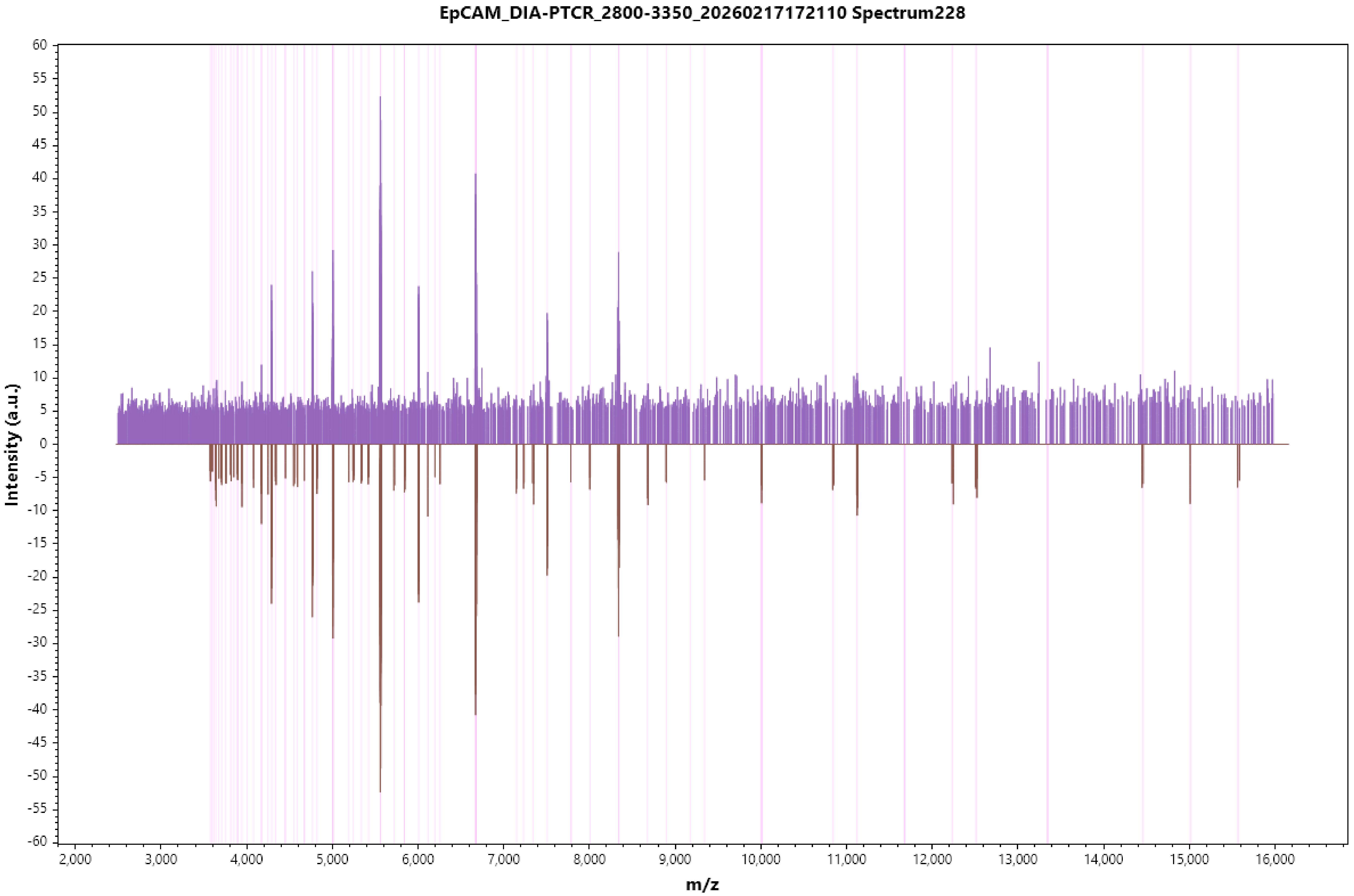
EpCAM_DIA-PTCR_2800-3350_20260217172110 Spectrum228
Intensity (a.u.)
m/z
60
55
50
45
40
35
30
25
20
15
10
5
0
-5
-10
-15
-20
-25
-30
-35
-40
-45
-50
-55
-60
2,000
3,000
4,000
5,000
6,000
7,000
8,000
9,000
10,000
11,000
12,000
13,000
14,000
15,000
16,000

EpCAM_DIA-PTCR_2800-3350_20260217172110 Spectrum229

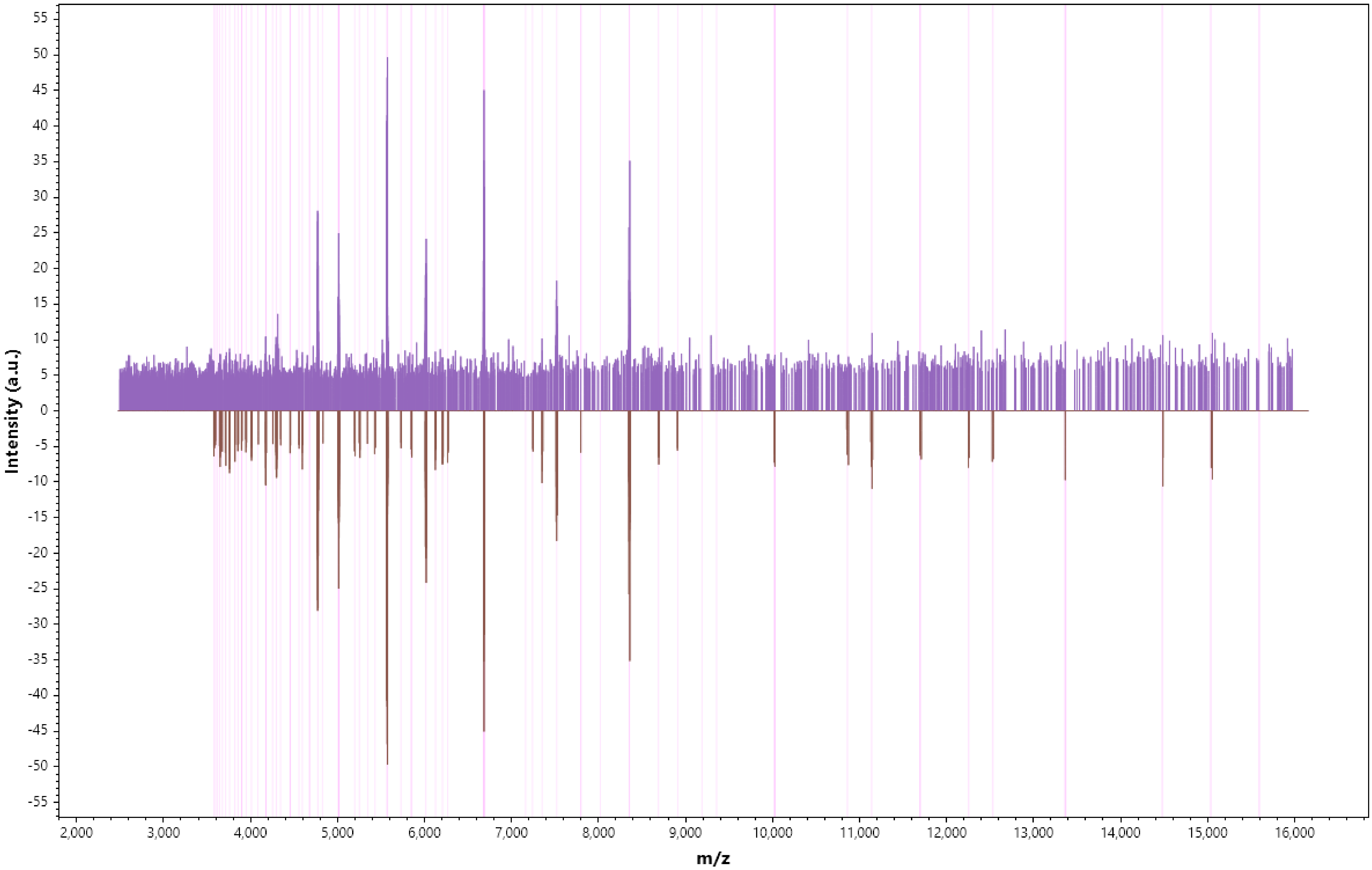

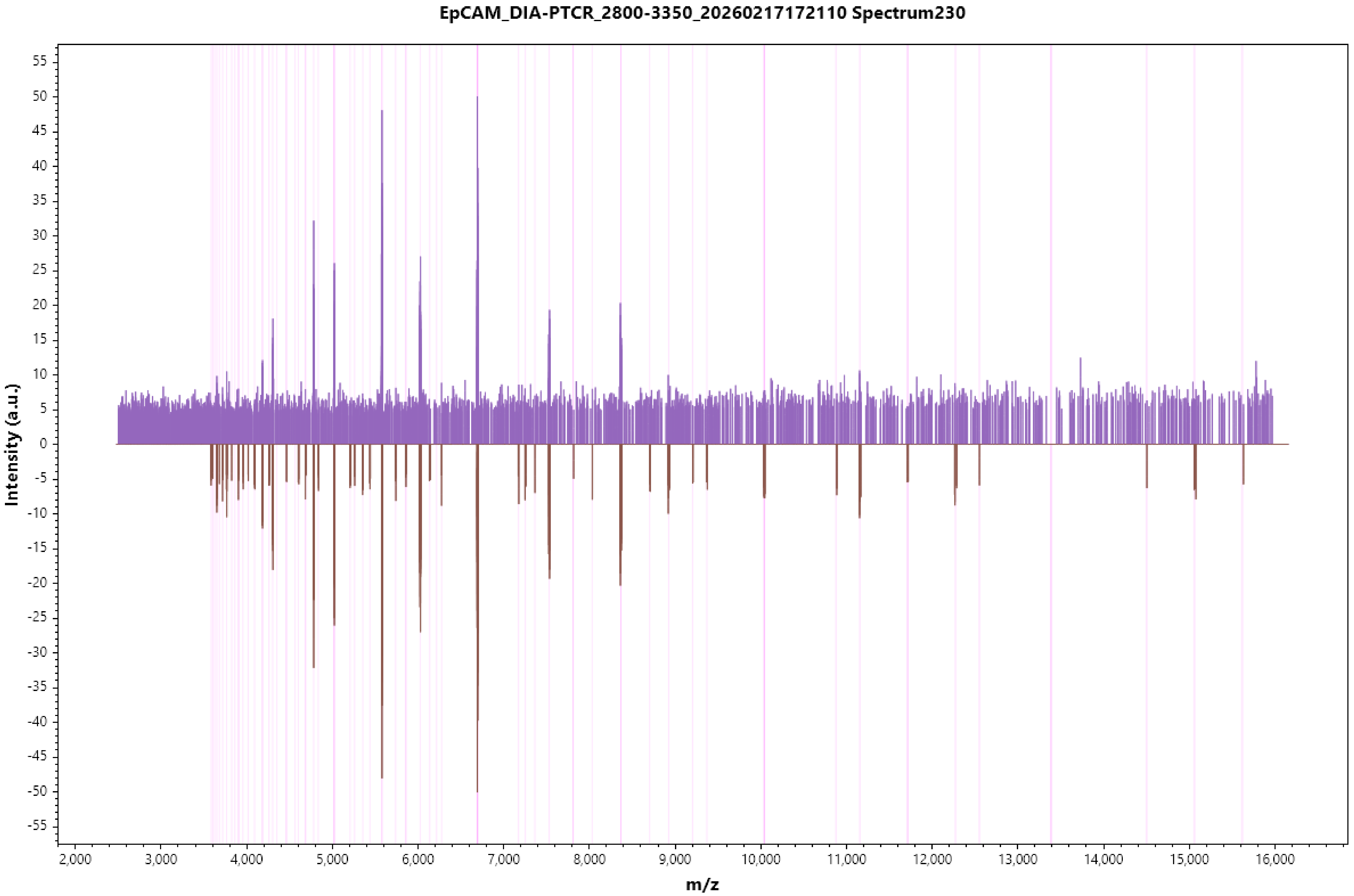
EpCAM_DIA-PTCR_2800-3350_20260217172110 Spectrum230
Intensity (a.u.)
m/z
55
50
45
40
35
30
25
20
15
10
5
0
-5
-10
-15
-20
-25
-30
-35
-40
-45
-50
-55
2,000
3,000
4,000
5,000
6,000
7,000
8,000
9,000
10,000
11,000
12,000
13,000
14,000
15,000
16,000

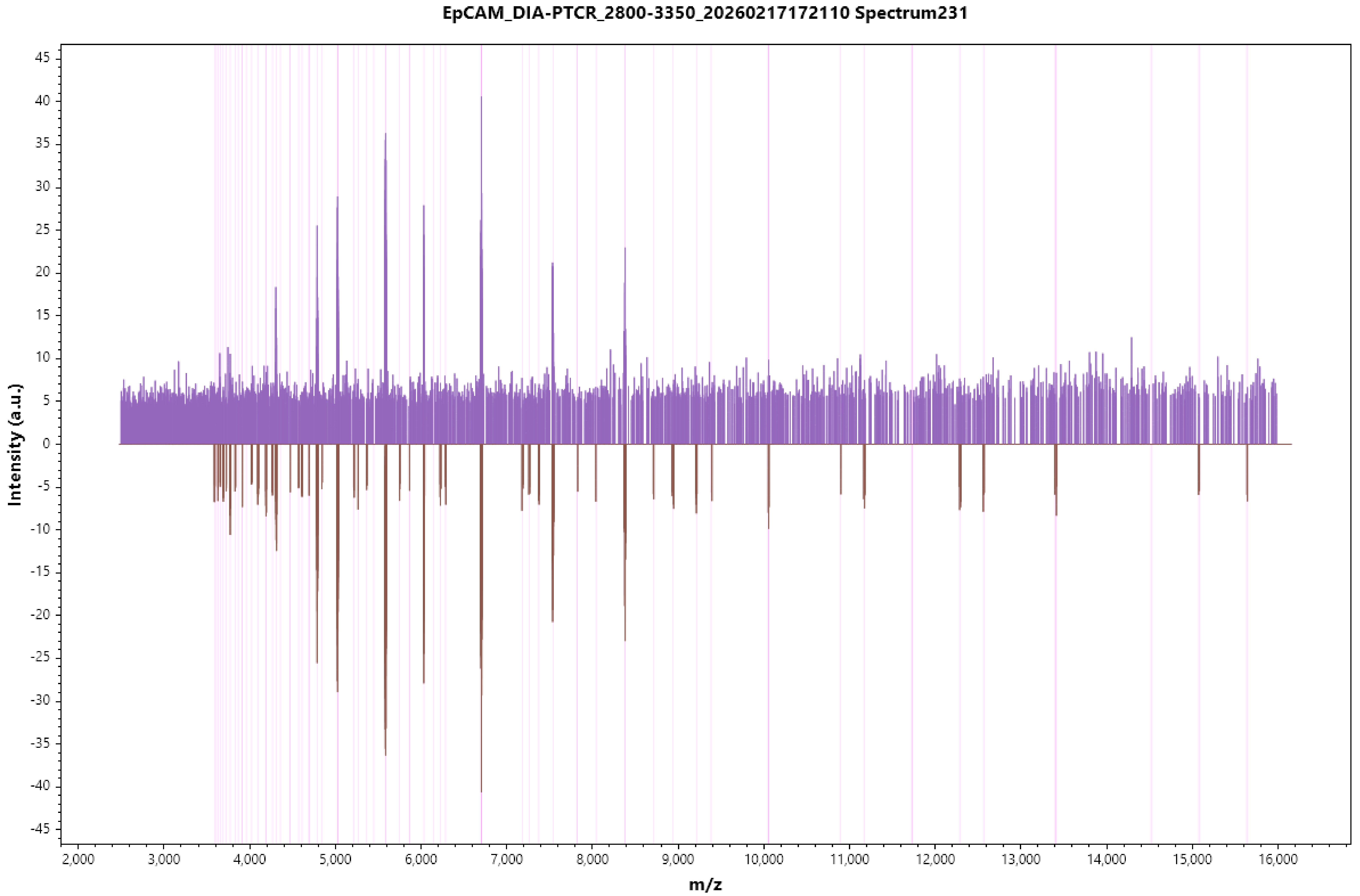

EpCAM_DIA-PTCR_2800-3350_20260217172110 Spectrum231
Intensity (a.u.)
m/z

EpCAM_DIA-PTCR_2800-3350_20260217172110 Spectrum232

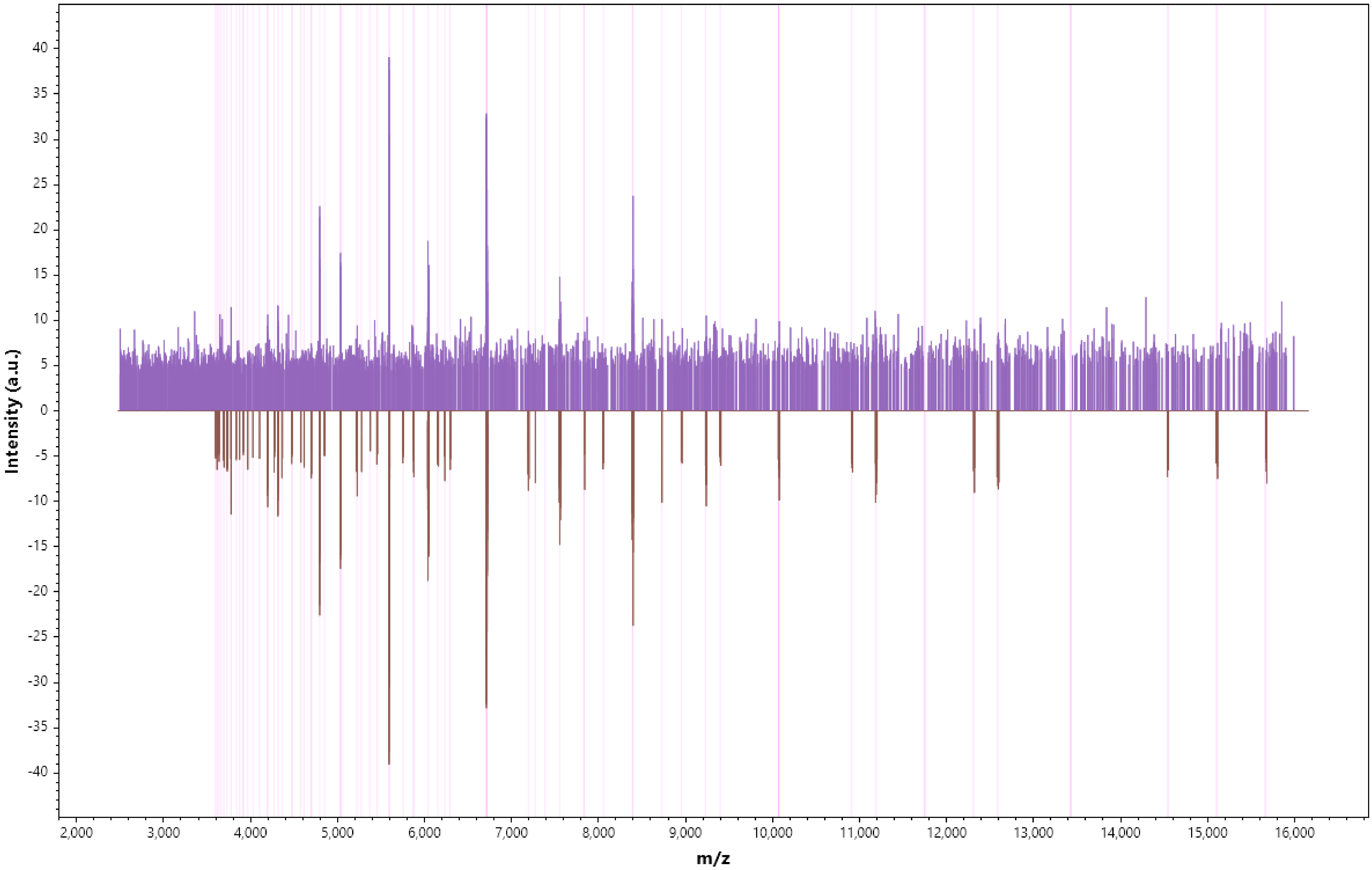

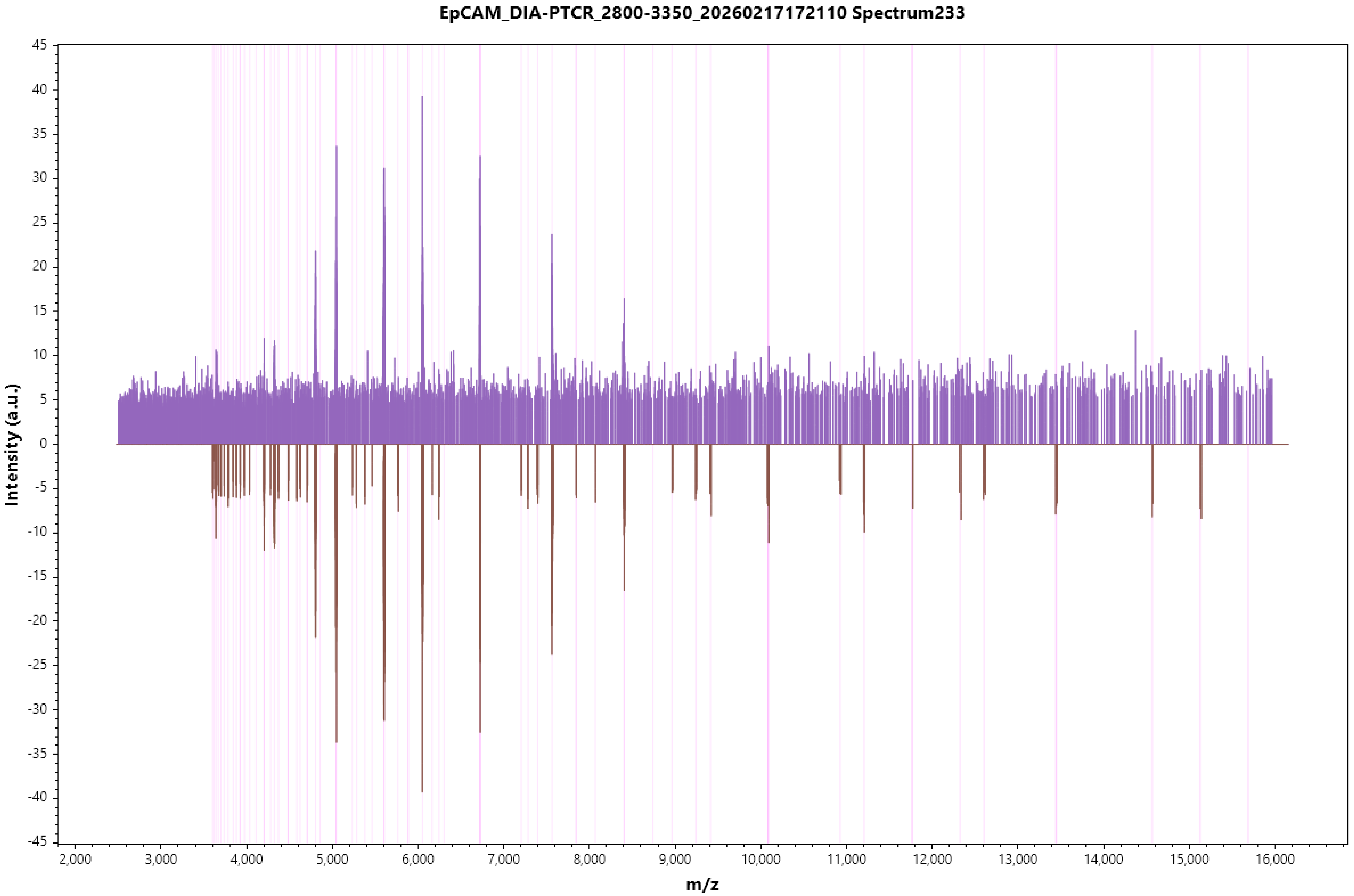

EpCAM_DIA-PTCR_2800-3350_20260217172110 Spectrum233
Intensity (a.u.)
m/z
45
40
35
30
25
20
15
10
5
0
-5
-10
-15
-20
-25
-30
-35
-40
-45
2,000
3,000
4,000
5,000
6,000
7,000
8,000
9,000
10,000
11,000
12,000
13,000
14,000
15,000
16,000

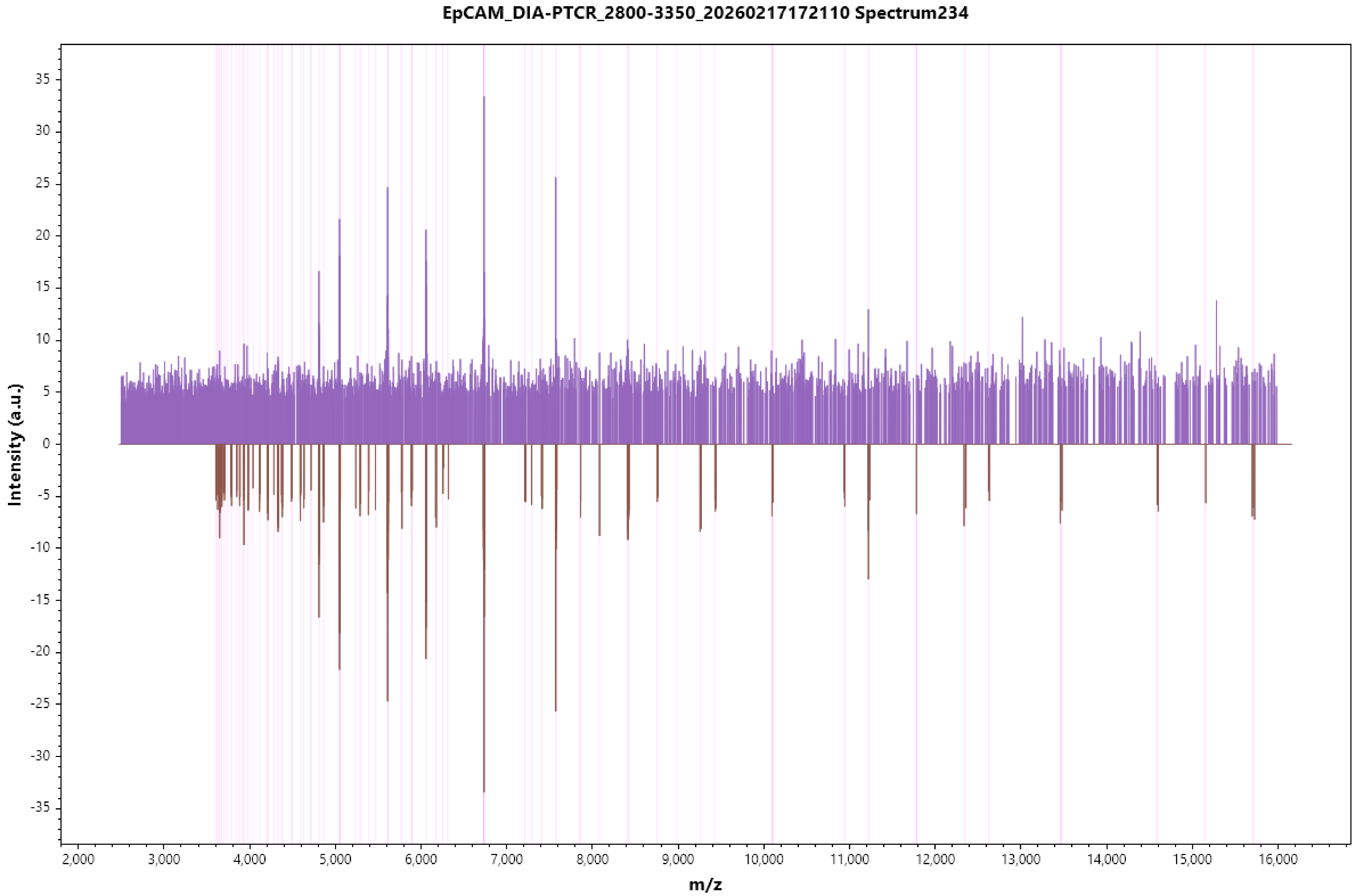

EpCAM_DIA-PTCR_2800-3350_20260217172110 Spectrum234
Intensity (a.u.)
m/z
35
30
25
20
15
10
5
0
-5
-10
-15
-20
-25
-30
-35
2,000
3,000
4,000
5,000
6,000
7,000
8,000
9,000
10,000
11,000
12,000
13,000
14,000
15,000
16,000

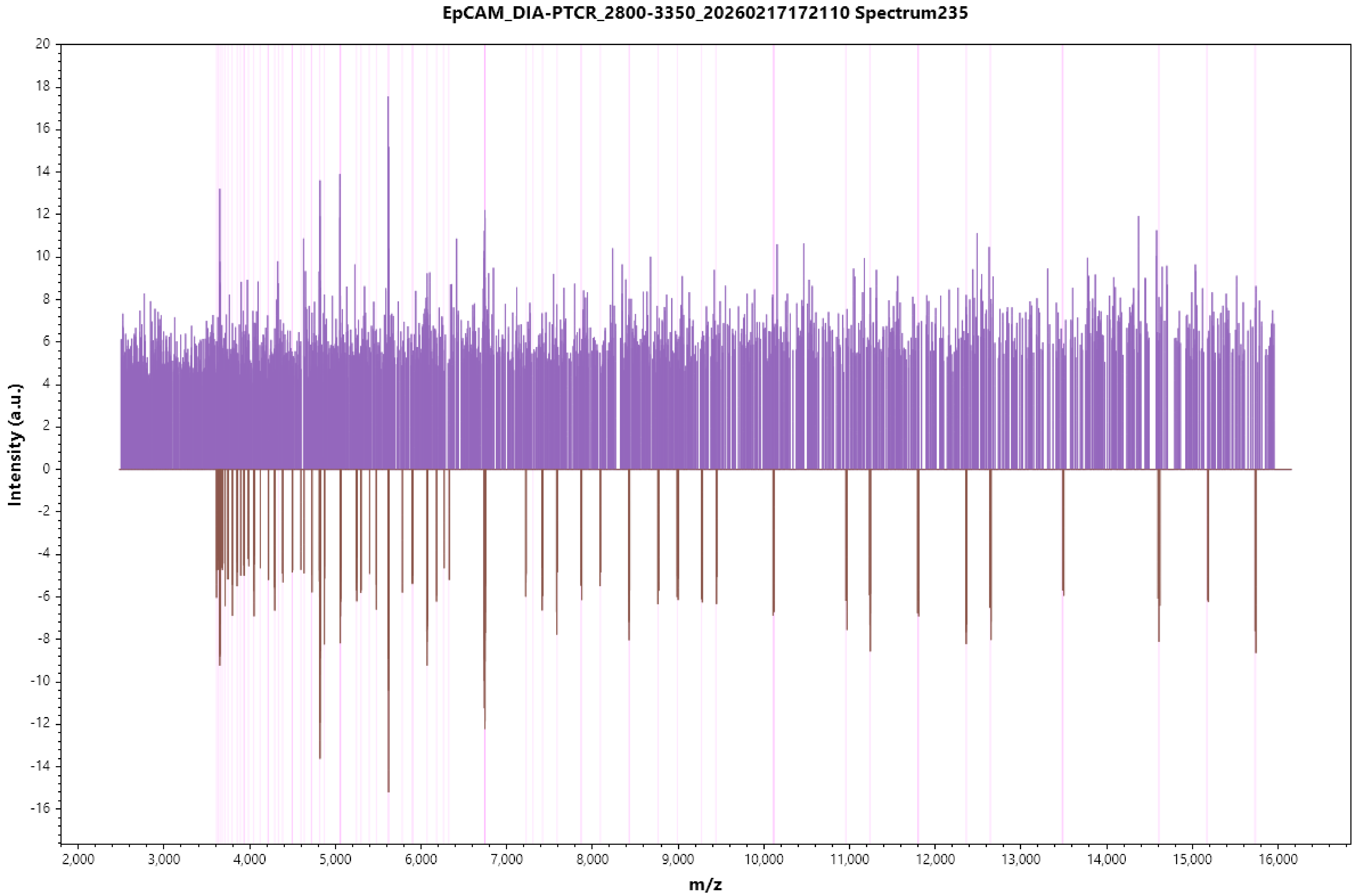
EpCAM_DIA-PTCR_2800-3350_20260217172110 Spectrum235
Intensity (a.u.)
m/z
20
18
16
14
12
10
8
6
4
2
0
-2
-4
-6
-8
-10
-12
-14
-16
2,000
3,000
4,000
5,000
6,000
7,000
8,000
9,000
10,000
11,000
12,000
13,000
14,000
15,000
16,000

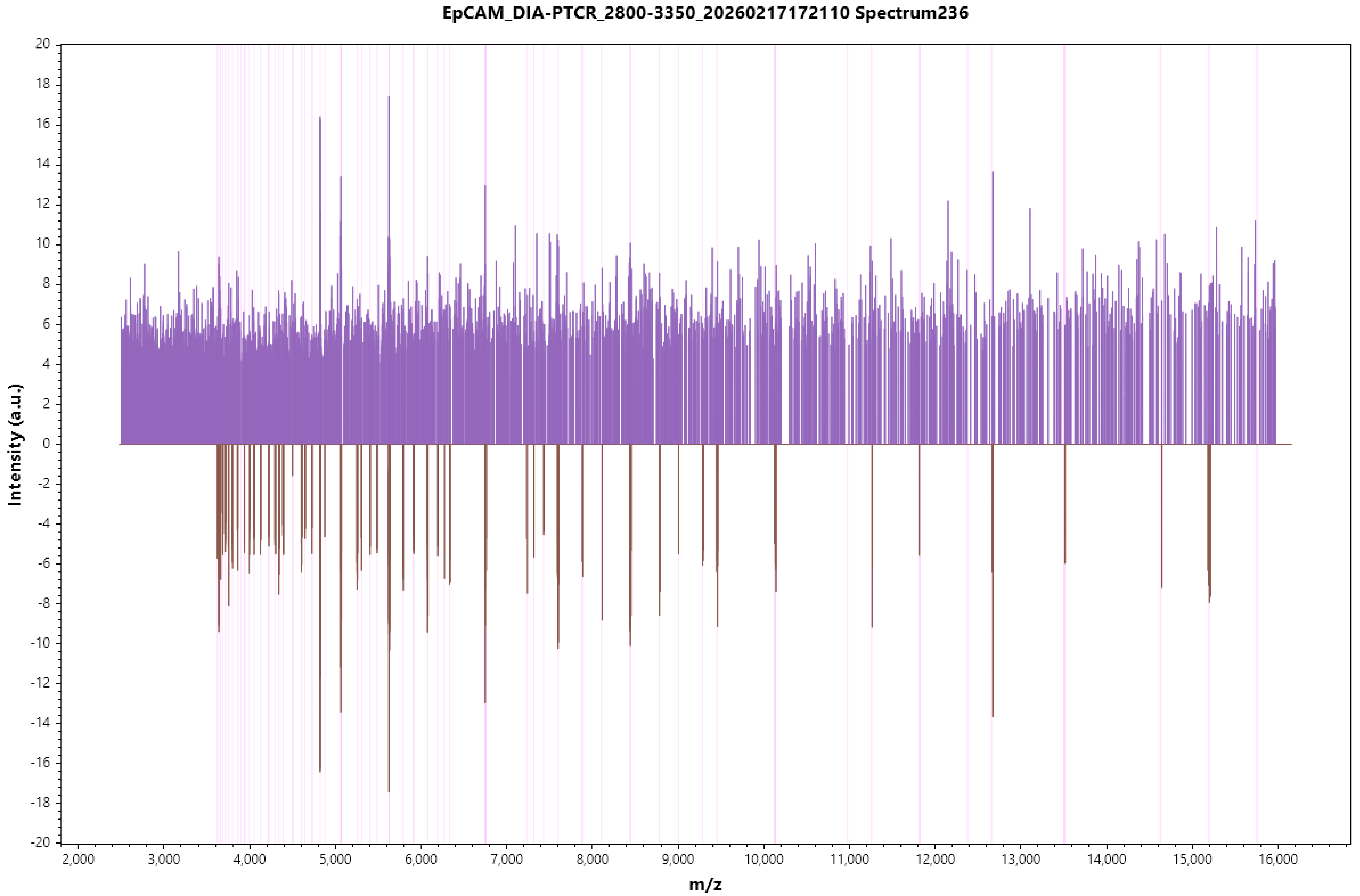
EpCAM_DIA-PTCR_2800-3350_20260217172110 Spectrum236
Intensity (a.u.)
m/z

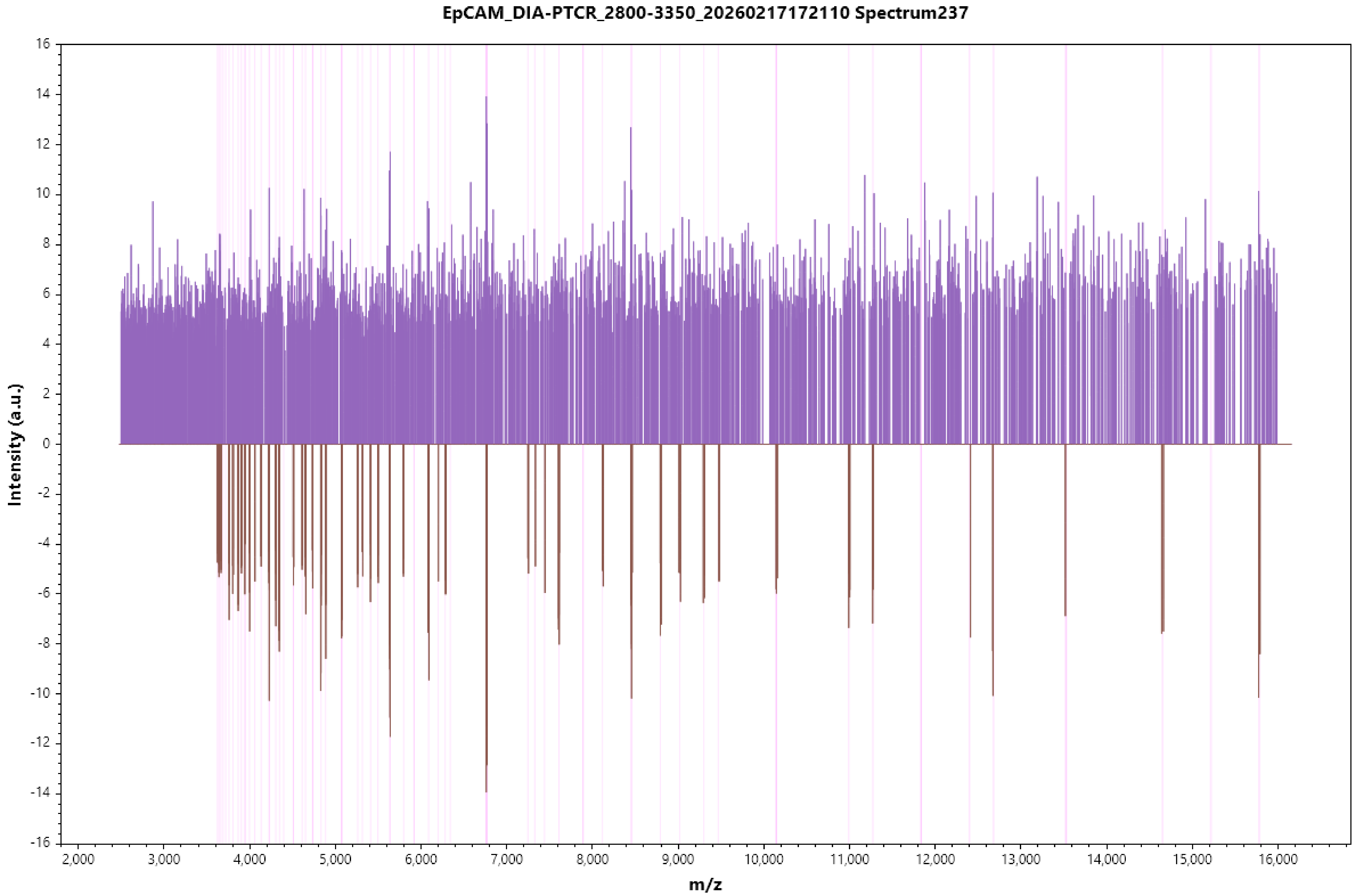
EpCAM_DIA-PTCR_2800-3350_20260217172110 Spectrum237
Intensity (a.u.)
m/z

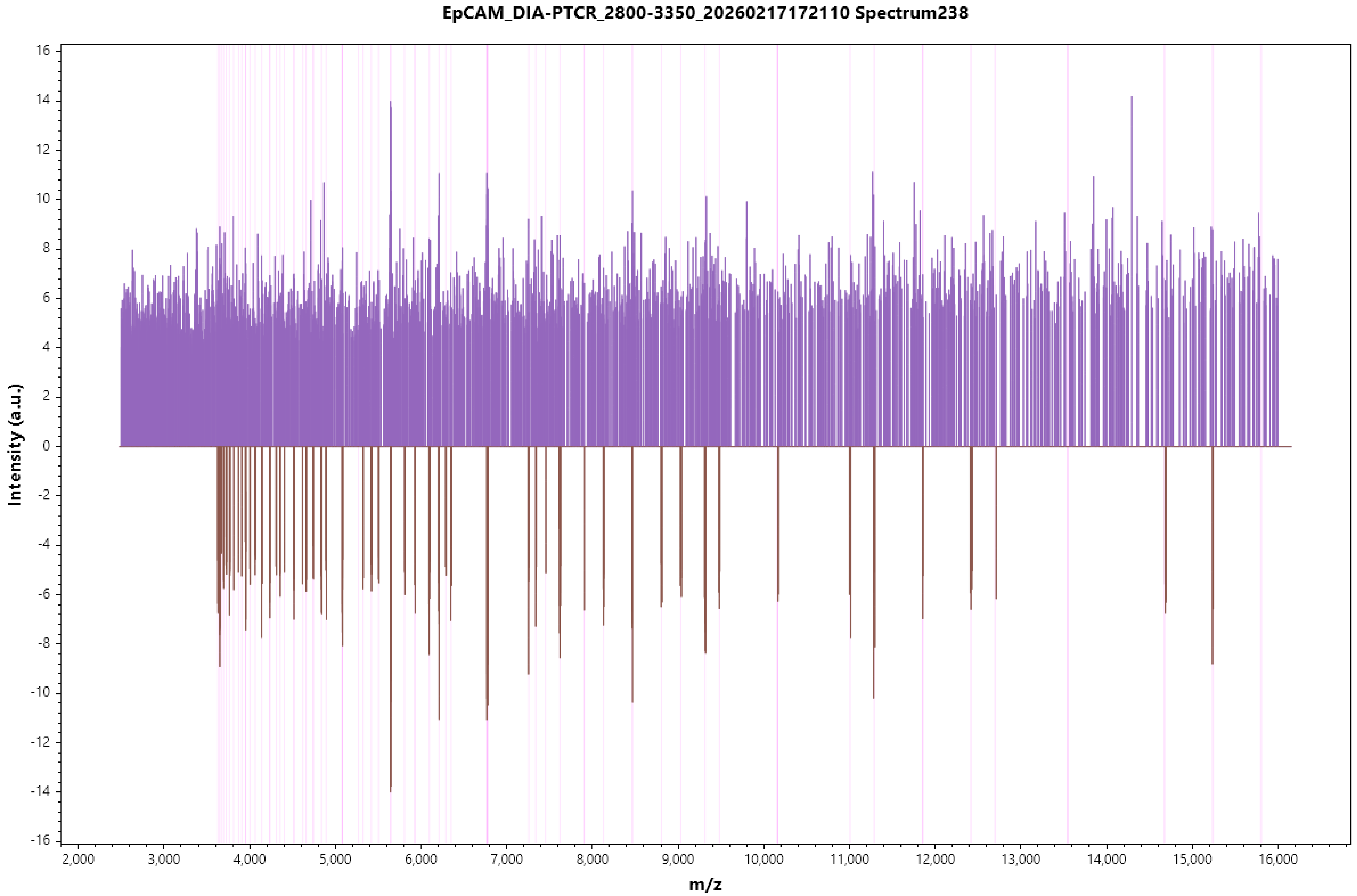
EpCAM_DIA-PTCR_2800-3350_20260217172110 Spectrum238
Intensity (a.u.)
m/z
16
14
12
10
8
6
4
2
0
-2
-4
-6
-8
-10
-12
-14
-16
2,000
3,000
4,000
5,000
6,000
7,000
8,000
9,000
10,000
11,000
12,000
13,000
14,000
15,000
16,000

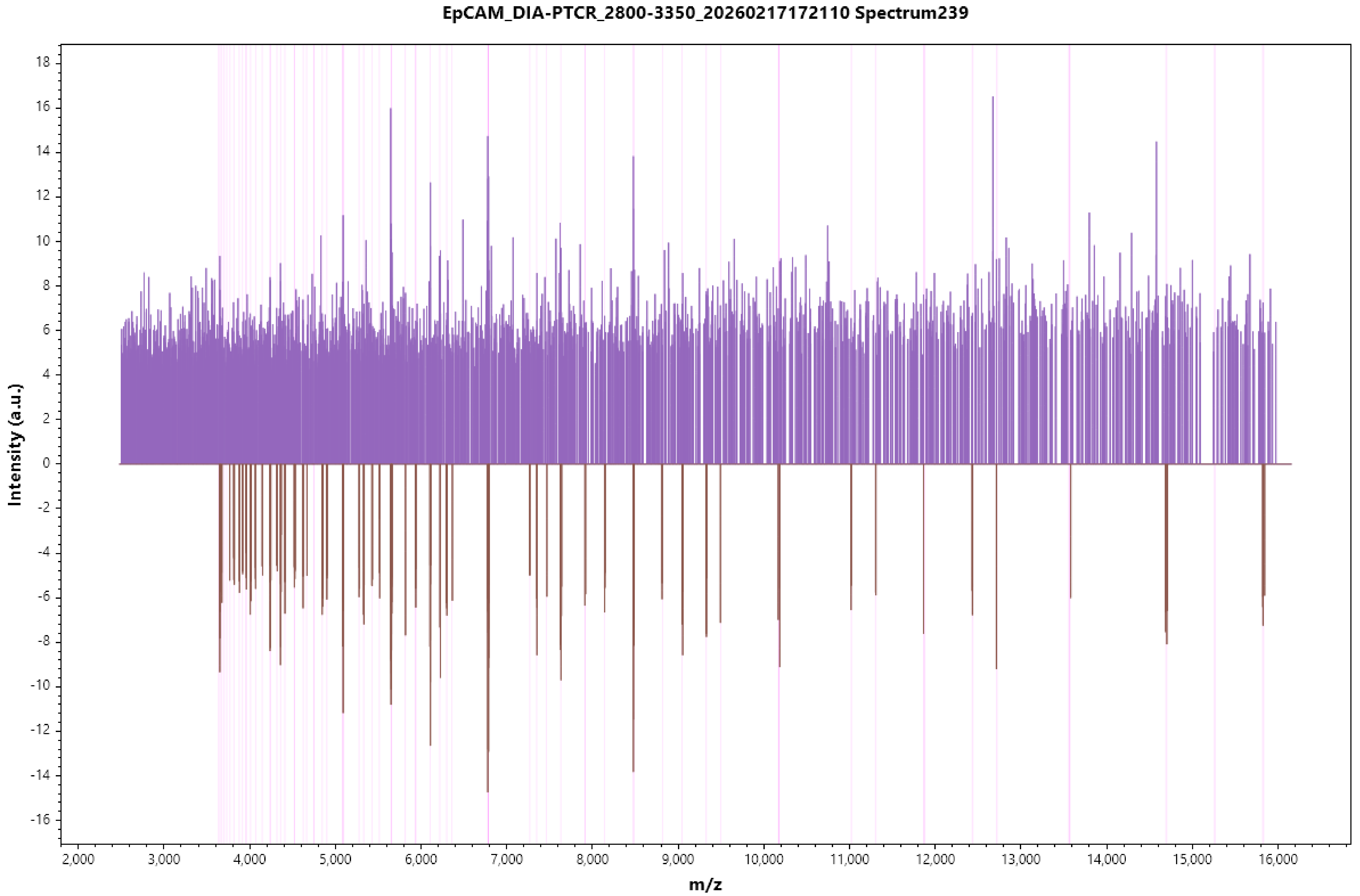

EpCAM_DIA-PTCR_2800-3350_20260217172110 Spectrum239
Intensity (a.u.)
m/z

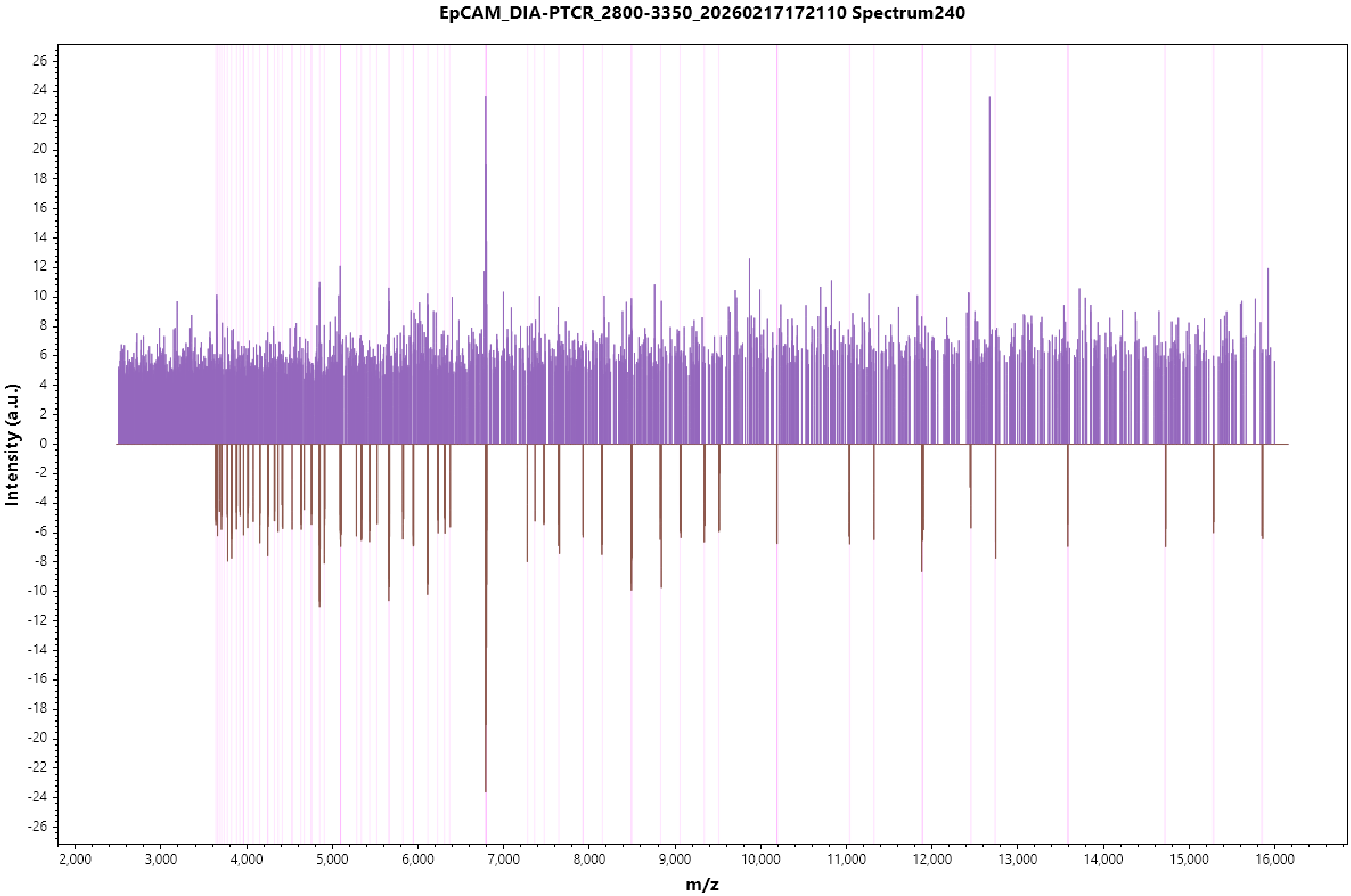

EpCAM_DIA-PTCR_2800-3350_20260217172110 Spectrum240
Intensity (a.u.)
m/z
26
24
22
20
18
16
14
12
10
8
6
4
2
0
-2
-4
-6
-8
-10
-12
-14
-16
-18
-20
-22
-24
-26
2,000
3,000
4,000
5,000
6,000
7,000
8,000
9,000
10,000
11,000
12,000
13,000
14,000
15,000
16,000
